# A nitrite-oxidizing bacterium constitutively consumes atmospheric hydrogen

**DOI:** 10.1101/2021.08.20.457082

**Authors:** Pok Man Leung, Anne Daebeler, Eleonora Chiri, Paul R. F. Cordero, Iresha Hanchapola, David L. Gillett, Ralf B. Schittenhelm, Holger Daims, Chris Greening

## Abstract

Chemolithoautotrophic nitrite-oxidizing bacteria (NOB) of the genus *Nitrospira* contribute to nitrification in diverse natural environments and engineered systems. *Nitrospira* are thought to be well-adapted to substrate limitation owing to their high affinity for nitrite and capacity to use alternative energy sources. Here, we demonstrate that the canonical nitrite oxidizer *Nitrospira moscoviensis* oxidizes hydrogen (H_2_) below atmospheric levels using a high-affinity group 2a nickel-iron hydrogenase [*K*_m(app)_ = 32 nM]. Atmospheric H_2_ oxidation occurred under both nitrite-replete and nitrite-deplete conditions, suggesting low-potential electrons derived from H_2_ oxidation promote nitrite-dependent growth and enable survival during nitrite limitation. Proteomic analyses confirmed the hydrogenase was abundant under both conditions and indicated extensive metabolic changes occur to reduce energy expenditure and growth under nitrite-deplete conditions. Respirometry analysis indicates the hydrogenase and nitrite oxidoreductase are *bona fide* components of the aerobic respiratory chain of *N. moscoviensis*, though they transfer electrons to distinct electron carriers in accord with the contrasting redox potentials of their substrates. Collectively, this study suggests atmospheric H_2_ oxidation enhances the growth and survival of NOB in amid variability of nitrite supply. These findings also extend the phenomenon of atmospheric H_2_ oxidation to a seventh phylum (Nitrospirota) and reveal unexpected new links between the global hydrogen and nitrogen cycles.

## Introduction

Bacteria of the genus *Nitrospira* are the most widespread group of nitrite-oxidizing bacteria. These bacteria are diverse and abundant in many natural environments, in wastewater treatment plants, and in drinking water treatment systems, where they play key roles in nitrification [1]. Reflecting the high standard redox potential of nitrite, chemolithoautotrophic growth on nitrite alone is a challenging lifestyle given little ATP is produced during catabolic processes and reverse electron flow is necessary for anabolic processes. However, it has recently been revealed that nitrite-oxidizing *Nitrospira* are more metabolically flexible than previously realized. In addition to mediating aerobic nitrite oxidation via the key enzyme nitrite oxidoreductase, cultured representatives can use ammonia (comammox bacteria), formate, and hydrogen (H_2_) as electron donors for aerobic respiration or (in case of formate and H_2_) nitrate respiration [2–6]. Such flexibility is thought to enhance the growth and survival of *Nitrospira* under different conditions, including amid variability or limitation in nitrite supply.

The model nitrite-oxidizing bacterium *Nitrospira moscoviensis* can aerobically grow on H_2_ as its sole energy and electron source in the absence of nitrite [3]. It possesses a group 2a [NiFe]-hydrogenase as its sole hydrogenase, and is predicted to use the high-energy electrons derived from H_2_ oxidation to support both aerobic respiration and carbon fixation through the reverse tricarboxylic acid (rTCA) cycle [3, 7]. Interestingly, *N. moscoviensis* was also found to transcribe hydrogenase genes even when grown on nitrite with air in the headspace, but without externally added H_2_ [3, 8]. Based on those results, we speculated that *N. moscoviensis* may be capable of both using H_2_ at elevated concentrations and potentially even scavenging atmospheric H_2_. However, the previous studies did not attempt to measure atmospheric H_2_ uptake or the kinetic parameters of this process.

Consistent with this hypothesis, recent studies have shown atmospheric H_2_ is a desirable energy source for bacteria. While the concentration of atmospheric H_2_ (530 ppbv) is thought to be too low to support growth, this gas nevertheless serves as a dependable lifeline for long-term survival and can be a useful supplement during mixotrophic growth. This reflects its ubiquitous availability, high diffusibility, low activation energy, and high energy yield [9, 10]. Culture-based studies have revealed bacteria from at least six phyla oxidize atmospheric H_2_, including various organotrophs, lithotrophs, and methanotrophs [11–15]. These bacteria each possess either high- or medium-affinity group 1h, 1f, or 2a [NiFe]-hydrogenases [16–18] that input electrons into aerobic respiratory chains. Bacterial atmospheric H_2_ oxidation is also biogeochemically significant, contributing to the net loss of three quarters of the H_2_ removed from the atmosphere each year [10]. In this study, we addressed the hypothesis that nitrite-oxidizing bacteria are also capable of oxidizing atmospheric H_2_. To do so, we performed kinetic, proteomic, and respirometric analyses of aerobic H_2_ oxidation by the *N. moscoviensis* group 2a [NiFe]-hydrogenase in the presence and absence of nitrite as the main substrate.

## Methods

### Bacterial strain and growth conditions

*Nitrospira moscoviensis* was maintained in mineral medium at 37ºC with shaking at 100 rpm in the dark. The medium base (per litre water) contained 0.15 g KH _2_PO_4_, 0.05 g MgSO_4_·7H_2_O, 0.5 g NaCl, 0.01 g CaCO_3_, 0.01 g NH_4_Cl, 34.4 μg MnSOg MnSO_4_ · H_2_O, 50 μg MnSOg H_3_BO_3_, 70 μg MnSOg ZnCl_2_, 72.6 μg MnSOg Na_2_MoO_4_ · 2H_2_O, 20 μg MnSOg CuCl_2_ · 2H_2_O, 24 μg MnSOg NiCl_2_ · 6H_2_O, 80 μg MnSOg CoCl_2_ · 6H_2_O, 1 mg FeSO_4_ · 7H_2_O, 3 μg MnSOg Na_2_SeO_3_ · 5H_2_O, and 4 μg MnSOg Na_2_WO_4_ · 2H_2_O [3]. The pH of the medium was adjusted to 8.4 – 8.6 before autoclaving. Filter-sterilized NaNO_2_ was added to the sterile medium at a final concentration of 1 mM. During incubation, nitrite consumption was monitored regularly with Quantofix nitrite indicator strips (Macherey-Nagel). Nitrite was replenished to a concentration of 1 mM when completely consumed until transfer of the culture to fresh medium. Heterotrophic contamination was tested regularly by inoculating 200 μl MnSOl of the culture onto nutrient agar, 5% R2A agar and nutrient broth at 37°C, and monitoring for two weeks. Biomass of *N. moscoviensis* was assessed by measuring total cell protein with the enhanced test tube protocol of the BCA protein assay (Sigma-Aldrich).

### Gas chromatography

To determine the ability of *N. moscoviensis* to oxidize H_2_ at sub-atmospheric levels in the presence and absence of nitrite, we prepared nitrite-replete and nitrite-deplete cultures as follows. Cells were harvested by centrifugation (4800 × *g*, 30 min) from a nitrite-oxidizing mother culture, re-suspended in fresh nitrite-free medium, and washed twice by centrifuging and re-suspending as described above. The absence of nitrite and nitrate in the final washed culture was confirmed by Quantofix nitrite and nitrate indicator strips (Macherey-Nagel). 30 ml of nitrite-free cultures was transferred into sterile 160 ml serum vials sealed with butyl rubber stoppers. One set of triplicate cultures was maintained with 1 mM nitrite as described in the above section (nitrite-replete condition) whereas another set was not supplied with any nitrite (nitrite-deplete condition). Cultures were allowed to adapt to the respective conditions for 48 hours. A medium only control and a heat-killed control (121°C, 15 p.s.i. for 20 min) were included in the experiment. For gas chromatography measurements, H_2_ in the air headspace of the vials was amended to give starting mixing ratios of approximately 10 parts per million (ppmv) via 1 % v/v H_2_ (in N_2_; Air Liquide). At each time interval, 2 ml of headspace gas was sampled using a gas-tight syringe and injected into a VICI gas chromatographic machine with a pulsed discharge helium ionization detector (model TGA-6791-W-4U-2, Valco Instruments Company Inc.) for H_2_ quantification [13]. The machine was calibrated against ultra-pure H_2_ standards down to the limit of quantification (20 ppbv). Calibration mixed gas (10.20 ppmv of H_2_, 10.10 ppmv of CH_4_, 9.95 ppmv of CO in N_2_, Air Liquide) and pressurised air (Air Liquide) with known trace gas concentrations were used as internal reference standards.

### Kinetics analysis

To ensure an accurate determination of H_2_ oxidation kinetics, *N. moscoviensis* kept under nitrite-deplete conditions (as described above) was used to minimise potential biomass deviations caused by nitrite-dependent growth. Triplicate cultures were incubated independently with approximately 10, 30, 90, 300, and 900 ppmv H_2_ in the vial headspace. Concentrations of H_2_ in the headspace were quantified by gas chromatography at three time points, 0 h, 8 h, and 24 h. Reaction rates of H_2_ consumption were calculated based on the concentration change between the second and third time points. Michaelis-Menten curves and parameters were estimated using the nonlinear fit (Michaelis-Menten, least squares regression) function in GraphPad Prism (version 9.0.0). Dissolved H_2_ was calculated by fitting Henry’s Law as previously described [3] with constants for H_2_ adopted from Sander [19].

### Shotgun proteomics

Triplicate 150 ml nitrite-replete and nitrite-deplete cultures were grown in Schott bottles and adapted for 48 hours as previously described. Cells were then harvested by centrifugation (4800 × g, 30 min, 4ºC), resuspended in phosphate-buffered saline (PBS; 137 mM NaCl, 2.7 mM KCl, 10 mM Na_2_HPO_4_ and 2 mM KH_2_PO_4_, pH 7.4), and centrifuged again. The cell pellets were immediately stored at -20ºC and sent to the Proteomics & Metabolomics Facility in Monash University for analysis. The samples were lysed in SDS lysis buffer (5% w/v sodium dodecyl sulphate, 100 mM HEPES, pH 8.1), heated at 95°C for 10 min, and then probe-sonicated before measuring the protein concentration using the BCA method. The lysed samples were denatured and alkylated by adding TCEP (Tris(2-carboxyethyl) phosphine hydrochloride) and CAA (2-chloroacetamide) to a final concentration of 10 mM and 40 mM, respectively, and the mixture was incubated at 55°C for 15 min. Sequencing grade trypsin was added at an enzyme to protein ratio of 1:50 and incubated overnight at 37°C after the proteins were trapped using S-Trap mini columns (Profiti). Tryptic peptides were sequentially eluted from the columns using (i) 50 mM TEAB, (ii) 0.2% formic acid and (iii) 50% acetonitrile, 0.2% formic acid. The fractions were pooled and concentrated in a vacuum concentrator prior to MS analysis. Using a Dionex UltiMate 3000 RSLCnano system equipped with a Dionex UltiMate 3000 RS autosampler, an Acclaim PepMap RSLC analytical column (75 µm × 50 cm, nanoViper, C18, 2 µm, 100Å; Thermo Scientific) and an Acclaim PepMap 100 trap column (100 µm × 2 cm, nanoViper, C18, 5 µm, 100Å; Thermo Scientific), the tryptic peptides were separated by increasing concentrations of 80% acetonitrile (ACN) / 0.1% formic acid at a flow of 250 nl/min for 158 min and analyzed with a QExactive HF mass spectrometer (ThermoFisher Scientific). The instrument was operated in data-dependent acquisition mode to automatically switch between full scan MS and MS/MS acquisition. Each survey full scan (375–1575 m/z) was acquired with a resolution of 120,000 (at 200 m/z), an AGC (automatic gain control) target of 3 × 106, and a maximum injection time of 54 ms. Dynamic exclusion was set to 15 seconds. The 12 most intense multiply charged ions (z ≥ 2) were sequentially isolated and fragmented in the collision cell by higher-energy collisional dissociation (HCD) with a fixed injection time of 54 ms, 30,000 resolution and an AGC target of 2 × 105. The raw data files were analyzed with the MaxQuant software suite v1.6.5.0 [20] and its implemented Andromeda search engine [21] to obtain protein identifications and their respective label-free quantification (LFQ) values using standard parameters. These data were further analyzed with LFQ-Analyst [22].

### Respirometry

Respirometry experiments were used to test for the fate of electrons from H_2_ oxidation and nitrite oxidation in the electron transport chain of *N. moscoviensis*. Rates of H_2_ oxidation or O_2_ consumption (for nitrite oxidation) were measured amperometrically according to previously established protocols [23, 24]. For each set of measurements, either a Unisense H_2_ microsensor or Unisense O_2_ microsensor electrode was polarised at +800 mV or -800 mV, respectively, with a Unisense multimeter. The microsensors were calibrated with either H_2_ or O_2_ standards of known concentration. Gas-saturated phosphate-buffered saline (PBS; 137 mM NaCl, 2.7 mM KCl, 10 mM Na_2_HPO_4_ and 2 mM KH_2_PO_4_, pH 7.4) was prepared by bubbling the solution with 100% (v/v) of either H_2_ or O_2_ for 10 min. To assess the electron flow from H_2_ to the respiratory chain, H_2_ oxidation was first measured in uncoupler/inhibitor-untreated cells in 1.1 ml microrespiration assay chambers sequentially amended with 0.9 ml of 3x concentrated *N. moscoviensis* culture [total cell protein = 4.67 (replicate 1), 4.86 (replicate 2), and 4.85 (replicate 3) µg/ ml], 0.1 ml O_2_-saturated PBS, and 0.1 ml H_2_-saturated PBS. Chambers were stirred (250 rpm) at room temperature. Following measurements in untreated cells, the assay mixtures were treated with either 10 µM nigericin, 10 µM valinomycin, 40 µM *N*-oxo-2-heptyl-4-hydroxyquinoline (HQNO), or 250 µM sodium azide, and the H_2_ oxidation rate was measured. To assess the integration of nitrite oxidation with the respiratory chain, O_2_ consumption was measured using nitrite as the electron source. Initial O_2_ consumption by uncoupler/inhibitor-untreated cells was measured in 1.1 ml microrespiration assay chambers sequentially amended with 1.0 ml of 5x concentrated *N. moscoviensis* culture [total cell protein = 8.87 (replicate 1), 10.38 (replicate 2), and 10.97 (replicate 3) µg/ml], 0.1 ml O_2_-saturated PBS, and 500 µM sodium nitrite with stirring (250 rpm) at room temperature. After initial measurements in untreated cells, the assay mixtures were treated with either 10 µM nigericin, 10 µM valinomycin, 40 µM HQNO, 250 µM sodium azide, or 500 µM sodium tungstate before further O_2_ consumption measurement. In both H_2_ and O_2_ measurements, changes in the gas concentrations over time were logged using Unisense Logger Software (Unisense, Denmark). Upon observing a linear change in either H_2_ or O_2_ concentration, rates of consumption were calculated over a period of 30 s and normalised against total cell protein concentration. Nitrite oxidation rates were calculated from oxygen uptake measurements using a nitrite oxidation to oxygen consumption ratio of 1:0.5 [25].

## Results and Discussion

### Nitrospira moscoviensis *oxidizes atmospheric H*_*2*_ *during growth and persistence using a high-affinity group 2a [NiFe]-hydrogenase*

Hydrogenase activity of *N. moscoviensis* cultures was measured during growth under nitrite-replete conditions and survival under nitrite-deplete cultures. The bacterium was incubated in closed vials with ambient air headspace supplemented with ∼10 ppmv of H_2_, and H_2_ mixing ratios were monitored using gas chromatography. Consistent with our hypothesis, the cultures oxidized H_2_ in a first-order kinetic process to final mixing ratios of 98 ppbv (nitrite-replete cultures) and 248 ppbv (nitrite-deplete cultures), which were five and two times below tropospheric H_2_ levels, respectively (Fig. 1a). No decrease of H_2_ was observed in heat-killed and medium-only controls, excluding H_2_ leakage as a source of bias in these experiments. This constitutes the first report of atmospheric H_2_ oxidation in the phylum Nitrospirota and by nitrifying microorganisms. The capacity of *N. moscoviensis* to oxidize atmospheric H_2_ is consistent with the physiology of other bacteria (*Mycobacterium smegmatis, Gemmatimonas aurantiaca, Acidithiobacillus ferrooxidans* and *Chloroflexus aggregans*) expressing group 2a [NiFe] hydrogenases [14, 26].

**Figure 1.**
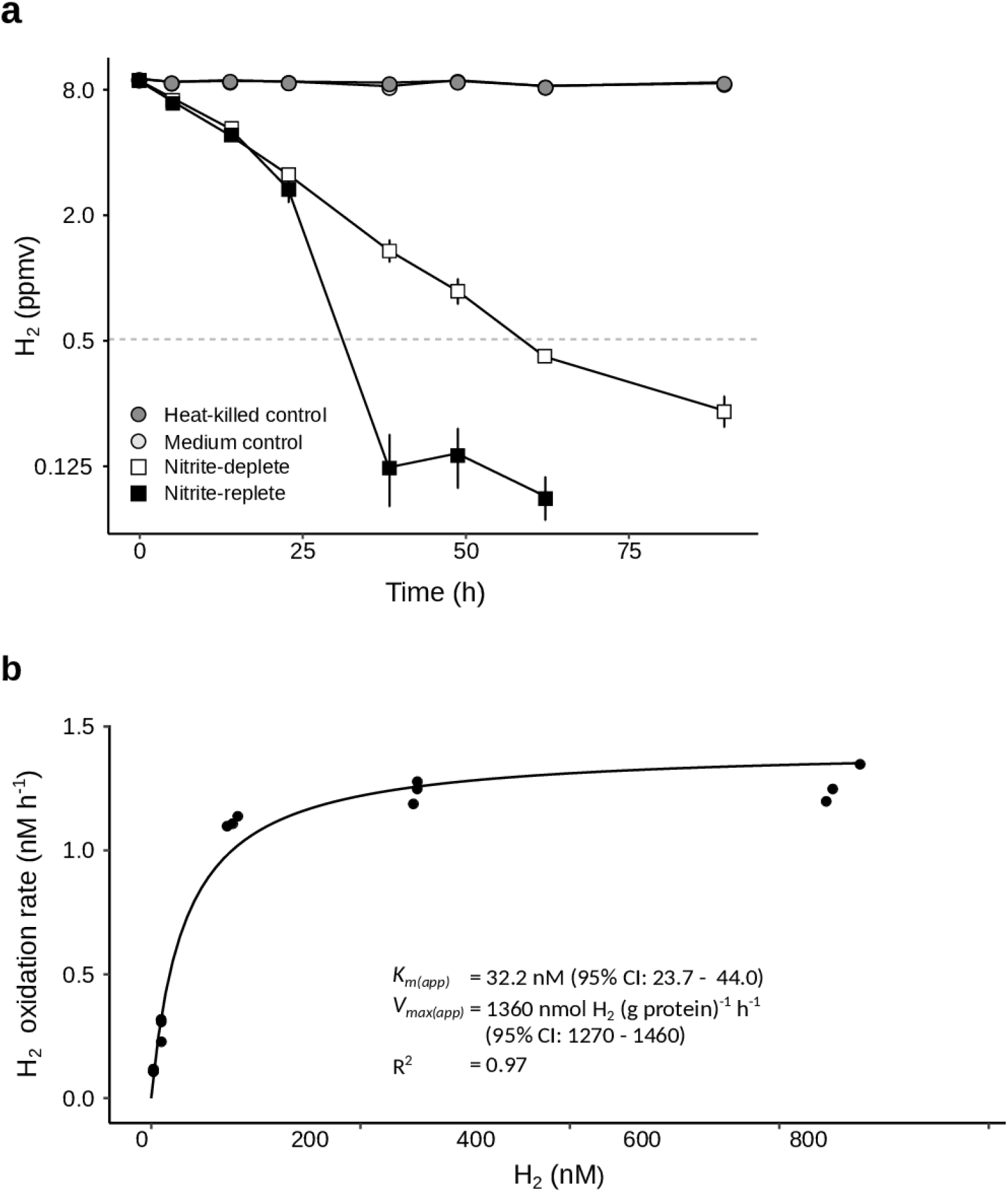
Hydrogen oxidizing activities of *Nitrospira moscoviensis* during nitrite-replete and nitrite-deplete conditions. **a** Oxidation of molecular hydrogen (H_2_) to sub-atmospheric levels by *N. moscoviensis* cultures. Error bars show standard errors of three biological replicates, with heat-killed cells and medium-only incubations as negative controls. The mixing ratio of H_2_ is on a logarithmic scale, and the grey dotted line indicates the average atmospheric mixing ratio (0.53 ppmv). **b** Kinetics of H_2_ oxidation by *N. moscoviensis* cells. The curve was fitted and kinetic parameters were calculated based on a Michaelis–Menten non-linear regression model.

Whole-cell kinetic measurements revealed that H_2_ oxidation by *N. moscoviensis* followed Michaelis-Menten-type kinetics (R^2^ = 0.97), with a mean apparent half-saturation constant of *K*_m(app)_ = 32.2 nM H_2_ (95% CI: 23.7 to 44.0) (Fig. 1b). The calculated mean maximum oxidation rate [*V*_max(app)_)] was 1360 nmol H_2_ per g of protein per h (95% CI: 1270 to 1460) (Fig. 1b). This suggests that *N. moscoviensis* possesses an unusually high-affinity group 2a [NiFe]-hydrogenase adapted for atmospheric H_2_ uptake. Its apparent H_2_ affinity is among the highest of any organism reported to date, comparable to the actinobacterium *Streptomyces avermitilis* and acidobacterium *Pyrinomonas methylaliphatogenes*, and in line with the high-affinity activities observed in whole soils [12, 27, 28].

### Proteome analysis reveals high hydrogenase expression and significant metabolic remodelling following nitrite depletion

To gain a holistic perspective of the *N. moscoviensis* metabolism, we compared shotgun proteomes from triplicate cultures grown under nitrite-deplete versus nitrite-replete conditions. Whole cell proteome analysis resulted in the detection of 2244 of the 4733 non-identical proteins (57.8%) encoded in the *N. moscoviensis* genome (Table S1). The proteome greatly changed under the nitrite-deplete condition, with 151 proteins significantly more abundant and 293 proteins less abundant by at least two-fold (p < 0.05; Table S1). Nitrite oxidoreductase and group 2a [NiFe] hydrogenase subunits were highly abundant in both conditions, though appear to be differentially regulated (Fig. 2). The *N. moscoviensis* genome encodes multiple copies of the nitrite oxidoreductase (Nxr) catalytic subunit, NxrA, and of the beta and gamma subunits NxrB and NxrC, respectively (see Supplementary Note for a discussion of the Nxr isoforms). One paralog each of NxrA, NxrB, and NxrC were among the ten most abundant proteins detected under nitrite-replete conditions, but their expression decreased by twofold (*p* < 0.001) following nitrite starvation (Fig. 2, Table S1, Supplementary Note). Conversely, the group 2a [NiFe]- hydrogenase catalytic subunit (HucL) increased 2.6-fold during starvation (*p* < 0.001) (Fig. 2). We also detected most other hydrogenase-related structural, maturation, and accessory proteins whose gene transcription had earlier been observed during growth on H_2_, with the exceptions of HypC, HypA, and UreH [3]. Altogether, this suggests that the hydrogenase appears to be constitutively expressed and slightly upregulated by *N. moscoviensis* under nitrite-deplete starvation conditions. These observations parallel those made regarding the group 2a [NiFe]-hydrogenase of *M. smegmatis*, but contrast with those of three other bacteria for which hydrogenase expression and activity peaked during growth [14, 18].

**Figure 2.**
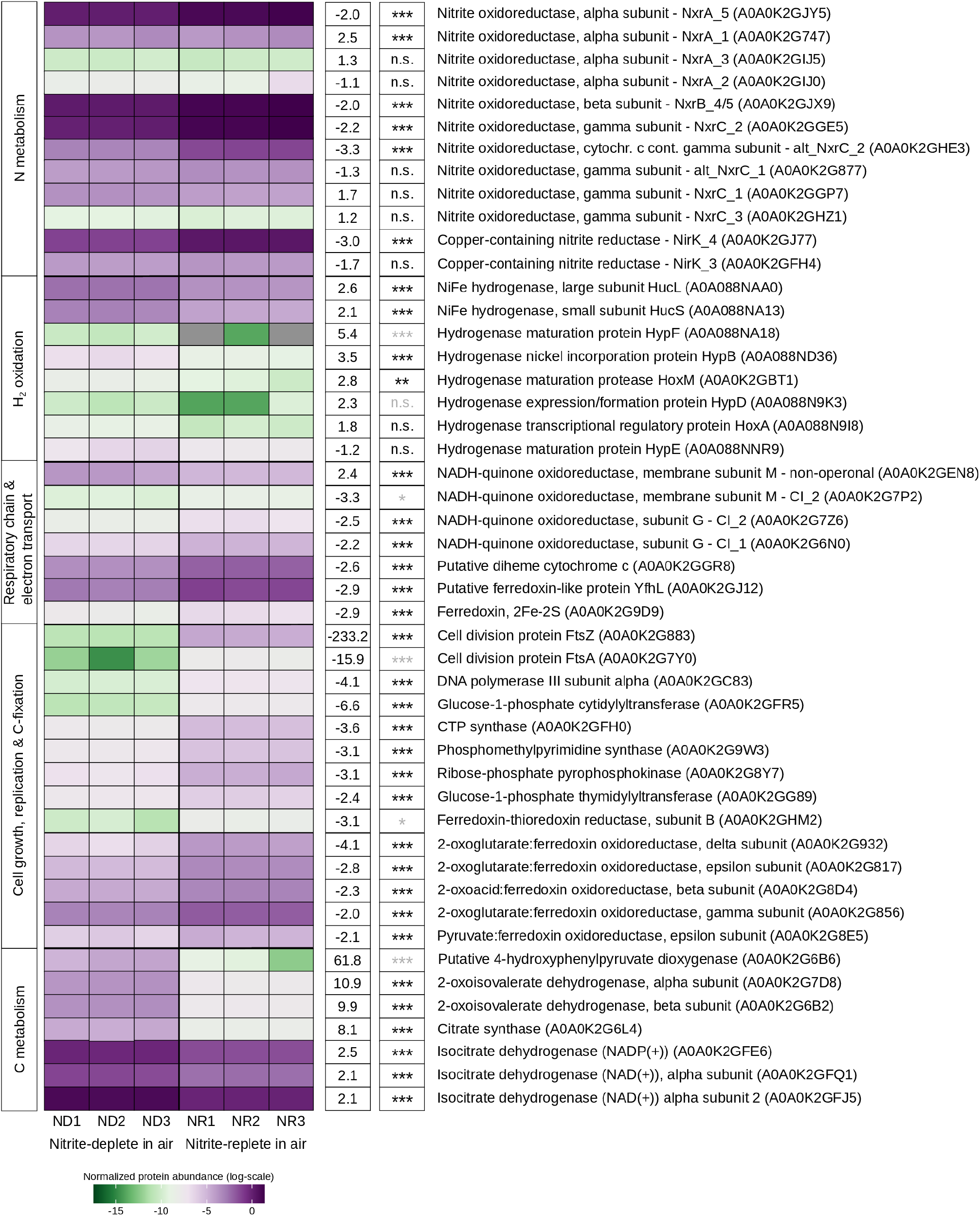
Heat map of selected *N. moscoviensis* proteins that were differentially abundant under nitrite-deplete versus nitrite-replete conditions. Notably, atmospheric H_2_ was oxidized under both conditions (see Fig. 1a). Relative protein abundance values were log_10_ transformed. The complete set of untransformed values is listed in Table S1. Fold-changes under nitrite-deplete conditions and the corresponding significance values (adj. p value ≤ 0.001, ***; ≤0.01, **; ≤0.05, *) are shown in white boxes next to each protein, with grey asterisks indicating that abundance values were imputed. Select low-abundance proteins of interest were included despite being not statistically significant. Depicted proteins are sorted based on their functional category as displayed on the left side. UniProtKB accession numbers are indicated in parenthesis. CI_1, 2M-type NADH-quinone oxidoreductase 1; CI_2, NADH-quinone oxidoreductase 2; nomenclature of nitrite oxidoreductase subunits according to Mundinger *et al*. [8].

Extensive other changes in the *N. moscoviensis* proteome were evident under nitrite-deplete conditions. Production of various proteins associated with biomass formation and growth decreased, including proteins responsible for cell division, DNA synthesis and replication, carbon fixation via the rTCA cycle, and amino acid synthesis (Fig. 2, Table S1). *N. moscoviensis* also appeared to generate additional reductant in nitrite-deplete conditions by catabolizing amino acid reserves. A putative 4-hydroxyphenylpyruvate dioxygenase, which catalyses the second step in the catabolism of tyrosine, and a putative branched-chain amino-acid dehydrogenase (2-oxoisovalerate dehydrogenase) were among the ten most differentially abundant proteins, upregulated by 62- and 10-fold respectively (p < 0.001) (Fig. 2, Table S1). Moreover, enzymes of the TCA cycle (citrate synthase and two forms of isocitrate dehydrogenase) were 8.1, 2.5, and 2.1 fold (p < 0.001) more abundant under nitrite-deplete conditions (Fig. 2, Table S1). Especially, the increased expression of citrate synthase, a key enzyme of the oxidative TCA cycle further indicates the increased mobilisation of energy harvesting from storage compounds. Rather unexpectedly, the relative abundance of proteins related to glycogen mobilization did not significantly vary (*e*.*g*., glycogen synthase GlgA, 1,4-alpha-glucan branching enzyme *GlgB*, glycogen debranching enzyme GlgX; Table S1). This observation potentially suggests that *N. moscoviensis* uses other compounds (*e*.*g*., amino acids) prior to glycogen under starvation, or that glycogen reserves play a role other than for persistence in this organism.

Finally, the proteome profiles indicate a broader remodelling of the respiratory chain and differential use of electron transport proteins. *N. moscoviensis* encodes three forms of complex I (NADH-quinone oxidoreductase) [3], designated as Cl_1 to Cl_3 [8]. Subunits M and G of CI_2 and the G subunit of CI_1 were less abundant (p < 0.05, 0.001 and 0.001, respectively) in the nitrite-deplete condition (Fig. 2, Table S1). Both forms of complex I are likely important for proton motive force-driven reverse electron transport under nitrite-oxidizing conditions. CI_2 may catalyse NADH production [8], whereas the 2M-type CI_1 might reduce ferredoxins for the rTCA cycle, interacting with ferredoxin *via* its G subunit [7, 29, 30]. Thus, we conclude that in the nitrite-deplete condition, reverse electron transport and ferredoxin reduction were downregulated, which is consistent with the downregulation of rTCA cycle enzymes (see above). Further analysis of the proteomic data is provided in the Supplementary Note.

### Hydrogenase and nitrite oxidoreductase differentially feed electrons into the aerobic respiratory chain

Having confirmed its production and activity, we subsequently investigated the integration of the hydrogenase in the respiratory chain of *N. moscoviensis*. We used H_2_ and oxygen microsensors to compare rates of H_2_- and nitrite-stimulated aerobic respiration in the presence and absence of selected uncouplers and inhibitors. Untreated cells rapidly oxidized H_2_ and nitrite, confirming oxidation of these electron donors can be coupled to O_2_ reduction as expected (Fig. 3a, b). The H_2_ and nitrite oxidation activities increased 2-3 (*p* < 0.001) and 1.5 to 2-fold (*p* < 0.01), respectively upon addition of valinomycin and nigericin (Fig. 3a, b). These two ionophores uncouple oxidative phosphorylation which induces the cellular response to increase the respiration rate to replenish the electrochemical gradient [31]. Hence, the increased oxidation rates show that both the hydrogenase and nitrite oxidoreductase are *bona fide* components of the respiratory chain of *N. moscoviensis*.

**Figure 3.**
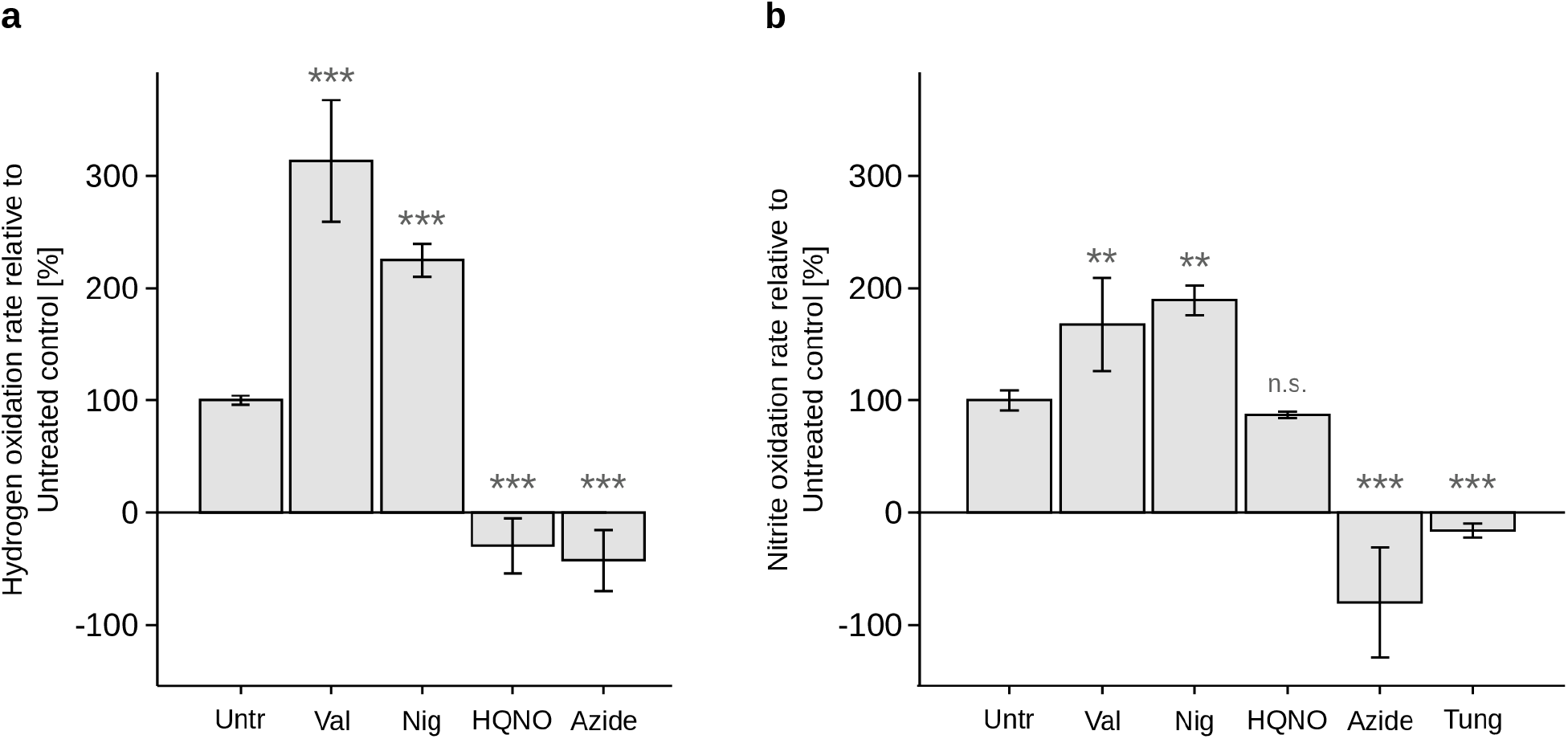
Amperometric measurements of hydrogen (a) and nitrite-oxidizing (b) activities of *N. moscoviensis* cells. The rates of H_2_ and nitrite oxidation were measured with a hydrogen or oxygen electrode, respectively, before and after treatment with the respiratory uncouplers and ionophores valinomycin (Val) and nigericin (Nig), the quinolone inhibitor 2-heptyl-4-hydroxyquinoline-N-oxide (HQNO), the terminal oxidase inhibitor azide, and the nitrite oxidoreductase inhibitor tungstate (Tung). Asterisks indicate significant differences to the untreated (Untr) control. ***, p < 0.001; **, p < 0.01; n.s., not significant. Negative rate values reflect electrode drift in the absence of any detectable hydrogen or oxygen consumption.

Upon addition of the cytochrome *bcc–aa*_*3*_ oxidase inhibitor azide [32] both H_2_ and nitrite oxidation ceased, suggesting the electrons derived were linked to cytochrome *c* reduction (Fig. 3a, b). This is remarkable, because *N. moscoviensis* and most other *Nitrospira* lack any characterized terminal oxidase and possess only novel, putative “cytochrome *bd*-like oxidases” [3, 7] whose sensitivity to azide inhibition is demonstrated hereby. Addition of the quinolone inhibitor 2-heptyl-4-hydroxyquinoline-N-oxide (HQNO) completely inhibited respiration with H_2_ but not with nitrite (Fig. 3a, b). Thus, electrons from H_2_ pass through the quinone pool, whereas electrons from nitrite are most likely directly transferred to the higher-potential cytochrome *c*. These observations are consistent with the different redox potentials of H_2_ (−420 mV) and nitrite (+420 mV) relative to quinones and cytochrome *c*. Thus, lower-potential electrons are transferred into the respiratory chain from H_2_, providing capacity for increased energy production and reducing costs of reverse electron transport.

## Conclusions

In conclusion, this study shows that *N. moscoviensis* can efficiently oxidize atmospheric H_2_ with a high-affinity hydrogenase, indicating that the environmentally widespread *Nitrospira* bacteria likely contribute to atmospheric H_2_ uptake in nature. Together with the previously demonstrated capabilities of *N. moscoviensis* to grow chemolithoautotrophically on elevated H_2_ concentrations and chemoorganoautotrophically on formate without nitrite [3, 4], the use of atmospheric H_2_ adds to the impressive metabolic versatility of this organism. Overall, our results suggest that atmospheric H_2_ serves as an alternative source of energy and electrons during nitrite starvation, likely improving the capacity of *N. moscoviensis* to persist during periods of low substrate availability. Under these conditions, *N. moscoviensis* remodels its metabolism extensively to reduce energy expenditure and growth, and to tap intracellular sources of energy and carbon (such as amino acids) in addition to scavenging H_2_ from air. Under nitrite-replete conditions, the use of atmospheric H_2_ likely serves as a convenient source of electrons for CO_2_ fixation, reducing the need for energetically expensive reverse electron transport from nitrite to the low-potential ferredoxins of the rTCA cycle. Hence, atmospheric H_2_ utilization may confer a selective advantage over other nitrite oxidizers and could contribute to the frequently observed predominance of *Nitrospira* in natural and engineered nitrifying microbial communities.

## Acknowledgements

We thank Ashleigh Kropp for technical assistance. This study was supported by the Austrian Science Fund (FWF) grants T938 (to AD) and P30570-B29 (to HD), the Czech Science Foundation (GACR) grant 21-17322M (to AD), the Comammox Research Platform of the University of Vienna, the NHMRC EL2 Fellowship APP1178715 (to CG), the ARC Discovery Project Grant DP200103074 (to CG), the Swiss National Foundation Early Mobility Postdoctoral Fellowship P2EZP3_178421 (to EC), Monash International Tuition Scholarships (to PML and DLG), and Australian Government Research Training Scholarships (to PML and DLG).

## Author contributions

HD, CG, PML, AD, EC, and PRFC designed the study. Different authors were responsible for bacterial culture preparation and maintenance (PML and AD), gas chromatography measurements (PML and EC), shotgun proteomics (PML, EC, IH, and RBS) and respirometry assays (PRFC, DLG, and PML). PML, AD, CG, EC, and PRFC analysed the data. The manuscript was written by AD, HD, CG, and PML with contributions from all co-authors.

## Supplementary Note

Nitrite oxidoreductase (Nxr), the key enzyme for nitrite oxidation in *Nitrospira*, consists of at least three subunits: alpha (NxrA), beta (NxrB), and gamma (NxrC) [1]. The *N. moscoviensis* genome encodes four paralogous alpha, four beta, and five gamma NXR subunits. A fifth set alpha and beta subunits were present in the originally sequenced genome [3] but have been lost from the laboratory strain after deletion of a genomic region [8]. In the proteomes, we detected four alpha, one beta, and five gamma subunits of NXR. Their protein expression was consistent with transcriptomic data from *N. moscoviensis* reported by Mundiger *et al*. [8]. Among these, the three subunits NxrA_5, NxrB_4/5, and NxrC_2 were among the 10 most abundant detected proteins in nitrite-replete conditions (nomenclature as in [8]). They were significantly downregulated (∼2×) in the nitrite-deplete cultures (p < 0.001; Fig. 2, Table S1). Two of the gamma subunits (alt_NxrC_1 and alt_NxrC_2) contain a predicted C-terminal transmembrane helix next to signal peptides for translocation into the periplasm *via* the Sec pathway, suggesting a membrane-anchored localization. The other three gamma subunits only contain the signal peptides and are likely soluble in the periplasm [8]. In line with previously reported transcriptomic and genomic analyses of *N. moscoviensis*, the NxrC_2 gamma subunit (lacking a transmembrane helix) showed the highest abundance in the proteomes, which was comparable to the most abundant alpha and beta subunits. Thus, the largest fraction of Nxr enzyme in *N. moscoviensis* is likely soluble in the periplasm. It may interact there with the membrane-anchored, also highly expressed (Table S1) alt_NxrC_1 and alt_NxrC_2 subunits, for example to transfer electrons from nitrite into the membrane-bound respiratory chain [1, 8].

Methylisocitrate lyase, which functions in the methylcitrate cycle (a modified version of the TCA cycle) allowing for the catabolism of propionic acid without a requirement for cobalamin (Vitamin B12), was upregulated ∼3× (p < 0.001, Fig. S1, Table S1) in the nitrite-deplete condition. Since many proteins involved in the energy-expensive biosynthesis of cobalamin were strongly downregulated under nitrite depletion (Table S1), we assume that *N. moscoviensis* uses the methylcitrate cycle for the catabolism of propionate, or of other fatty-acids with an odd number of carbon atoms via beta-oxidation. Since we did not detect any NADH-quinone oxidoreductase CI_3 subunits in the proteomes, we assume that this form of respiratory complex I in *N. moscoviensis* may not have a function under the tested conditions.

