## Supplemental Table S1 for "A nitrite-oxidizing bacterium constitutively consumes atmospheric hydrogen"

**Table S1. Differential expression analysis of N. moscoviensis grown under nitrite-deplete and nitrite-replete conditions in air. Also shown are global proteomic spectral counts and normalized spectral abundance (riBAQ) values.**

| | | | | | | | | | | | | MS/MS spectral counts | | | | | | Normalised spectral abundance (riBAQ) | | | | | |
|---|---|---|---|---|---|---|---|---|---|---|---|---|---|---|---|---|---|---|---|---|---|---|---|
| Gene Name | Protein UniProtKB IDs | Annotation | Mol. weight [kDa] | Length (aa) | Log2 fold change | Fold change | p value | adjusted p value | significant | imputed | number of NAs | Nitrite-deplete in air 1 | Nitrite-deplete in air 2 | Nitrite-deplete in air 3 | Nitrite-replete in air 1 | Nitrite-replete in air 2 | Nitrite-replete in air 3 | Nitrite-deplete in air 1 | Nitrite-deplete in air 2 | Nitrite-deplete in air 3 | Nitrite-replete in air 1 | Nitrite-replete in air 2 | Nitrite-replete in air 3 |
| NITMOv2_3655 | A0A0K2GHF5 | Uncharacterized protein | 28.679 | 265 | 0.213 | 1.16 | 2.55E-02 | 3.92E-02 | no | no | 0 | 541 | 645 | 592 | 527 | 527 | 525 | 3.977774854 | 3.984598421 | 3.789033185 | 3.12511338 | 3.29147352 | 3.346303807 |
| NITMOv2_4425 | A0A0K2GIM6 | Alginate_exp domain-containing protein | 62.347 | 568 | 0.278 | 1.21 | 1.16E-02 | 2.03E-02 | no | no | 0 | 537 | 584 | 571 | 419 | 420 | 477 | 2.896644696 | 2.929247895 | 3.094993812 | 2.342983606 | 2.413177565 | 2.682437409 |
| groEL | A0A0K2G9S5 | 60 kDa chaperonin | 58.328 | 545 | 0.135 | 1.10 | 5.35E-02 | 7.47E-02 | no | no | 0 | 1093 | 1145 | 1253 | 1592 | 1558 | 1438 | 2.489824417 | 2.401083772 | 2.393860667 | 2.30892646 | 2.425451646 | 2.38412908 |
| hupB.1 | A0A0K2GIF8 | DNA-binding protein HU-beta | 10.363 | 96 | 0.0835 | 1.06 | 3.24E-01 | 3.66E-01 | no | no | 0 | 113 | 170 | 155 | 143 | 126 | 118 | 2.255639273 | 2.199673536 | 2.262530012 | 2.230002725 | 1.601379163 | 1.592721936 |
| IDH | A0A0K2GFJ5 | Isocitrate dehydrogenase [NAD] catalytic subunit 5, mitochondrial | 37.011 | 337 | 1.06 | 2.08 | 1.22E-06 | 3.15E-05 | yes | no | 0 | 419 | 451 | 432 | 255 | 251 | 253 | 1.916207825 | 1.834319279 | 1.870321076 | 0.883901412 | 0.908592738 | 0.929359013 |
| NITMOv2_4557 | A0A0K2GJ23 | Orotate phosphoribosyltransferase (Modular protein) | 17.343 | 169 | 0.346 | 1.27 | 1.87E-01 | 2.24E-01 | no | no | 0 | 39 | 68 | 45 | 13 | 16 | 20 | 1.81494541 | 1.709092954 | 1.65353922 | 1.2993986 | 1.310716493 | 1.312035195 |
| rbr.1 | A0A0K2GJA4 | Rubrerythrin | 15.609 | 141 | 0.491 | 1.41 | 9.63E-04 | 2.73E-03 | no | no | 0 | 82 | 88 | 92 | 92 | 88 | 86 | 1.701014653 | 1.597261387 | 1.599377308 | 1.377670807 | 1.343079672 | 1.30469886 |
| bcp2 | A0A0K2GIB3 | Peroxiredoxin | 17.592 | 156 | 0.459 | 1.37 | 2.44E-04 | 1.01E-03 | no | no | 0 | 103 | 104 | 101 | 96 | 94 | 90 | 1.586072116 | 1.5169515 | 1.563965962 | 1.202039583 | 1.201694687 | 1.189978719 |
| groS | A0A0K2GAU8 | 10 kDa chaperonin | 10.935 | 99 | 0.297 | 1.23 | 1.17E-02 | 2.04E-02 | no | no | 0 | 81 | 88 | 81 | 62 | 65 | 59 | 1.536579161 | 1.459754904 | 1.408554825 | 1.23009971 | 1.220090272 | 1.269150001 |
| bcp3 | A0A0K2GCR5 | Peroxiredoxin | 23.587 | 211 | 0.577 | 1.49 | 2.52E-04 | 1.03E-03 | no | no | 0 | 175 | 175 | 150 | 141 | 123 | 116 | 1.371363755 | 1.369296287 | 1.266142547 | 0.932306982 | 0.937662112 | 0.907080289 |
| NITMOv2_4533 | A0A0K2GJX9 | Putative Nitrite oxidoreductase, beta subunit | 49.888 | 429 | -1.03 | -2.04 | 3.32E-05 | 2.66E-04 | yes | no | 0 | 325 | 351 | 344 | 545 | 520 | 606 | 1.158615721 | 1.153024714 | 1.176645978 | 2.132125447 | 2.125435816 | 2.513234192 |
| NITMOv2_4538 | A0A0K2GJY5 | Putative Nitrite oxidoreductase, alpha subunit | 131.83 | 1145 | -1.03 | -2.04 | 4.93E-05 | 3.48E-04 | yes | no | 0 | 782 | 833 | 823 | 1417 | 1359 | 1566 | 1.058512638 | 1.07339923 | 1.051776408 | 1.964653131 | 1.921126744 | 2.317418778 |
| NITMOv2_3685 | A0A0K2GHH6 | Putative Monoheme cytochrome c | 15.99 | 149 | 0.428 | 1.35 | 4.92E-03 | 9.81E-03 | no | no | 0 | 88 | 87 | 92 | 74 | 87 | 78 | 1.050797808 | 1.031415951 | 0.93367468 | 0.759888976 | 0.843011856 | 0.729012333 |
| NITMOv2_3003 | A0A0K2GEN6 | Nitrogen regulatory protein P-II | 11.086 | 98 | 0.654 | 1.57 | 1.73E-04 | 7.93E-04 | no | no | 0 | 90 | 96 | 75 | 47 | 41 | 43 | 1.043293765 | 1.019820197 | 1.009022049 | 0.665846829 | 0.65354044 | 0.586924785 |
| NITMOv2_3763 | A0A0K2GH25 | Uncharacterized protein | 22.251 | 210 | 0.574 | 1.49 | 1.88E-04 | 8.31E-04 | no | no | 0 | 70 | 76 | 72 | 56 | 62 | 62 | 1.036422086 | 1.046805257 | 0.992744717 | 0.709928623 | 0.74368502 | 0.717720131 |
| NITMOv2_3624 | A0A0K2GGE5 | Putative Nitrite oxidoreductase, membrane subunit | 34.243 | 316 | -1.11 | -2.16 | 5.09E-05 | 3.55E-04 | yes | no | 0 | 187 | 204 | 224 | 315 | 339 | 354 | 0.974071082 | 0.997665071 | 1.067114295 | 2.134938863 | 2.117978146 | 2.527079726 |
| NITMOv2_1768 | A0A0K2GB67 | Uncharacterized protein | 25.912 | 248 | -1.18 | -2.27 | 2.84E-05 | 2.37E-04 | yes | no | 0 | 116 | 133 | 138 | 259 | 289 | 290 | 0.862985959 | 0.903530228 | 0.919487135 | 2.010852391 | 2.367654707 | 2.655465589 |
| NITMOv2_3292 | A0A0K2GFE6 | Isocitrate dehydrogenase (NADP(+)) | 13.507 | 117 | 1.3 | 2.46 | 1.51E-06 | 3.61E-05 | yes | no | 0 | 64 | 67 | 85 | 68 | 73 | 77 | 0.8567888 | 0.802883774 | 0.826271129 | 0.356830051 | 0.342540084 | 0.345275259 |
| NITMOv2_4492 | A0A0K2GIV1 | Uncharacterized protein | 16.948 | 159 | -1.2 | -2.30 | 6.82E-05 | 4.34E-04 | yes | no | 0 | 62 | 67 | 74 | 150 | 151 | 157 | 0.852889228 | 0.825469096 | 0.853150775 | 2.128867807 | 2.229221717 | 2.529776908 |
| rbpA.1 | A0A0K2GGK5 | Putative RNA-binding protein RbpA | 10.147 | 99 | -0.393 | -1.31 | 3.30E-02 | 4.90E-02 | no | no | 0 | 65 | 94 | 104 | 85 | 85 | 85 | 0.786174918 | 0.925704909 | 1.058410044 | 1.168382238 | 1.134327072 | 1.177140133 |
| NITMOv2_4475 | A0A0K2GIZ3 | Uncharacterized protein | 23.354 | 218 | -1 | -2.00 | 8.68E-05 | 5.13E-04 | yes | no | 0 | 181 | 181 | 162 | 261 | 272 | 283 | 0.728081825 | 0.705737555 | 0.689610558 | 1.520133331 | 1.459807737 | 1.691636591 |
| NITMOv2_2966 | A0A0K2GEJ0 | Isocitrate dehydrogenase (NADP(+)) | 13.296 | 116 | 0.997 | 2.00 | 5.91E-06 | 8.24E-05 | no | no | 0 | 177 | 188 | 206 | 132 | 127 | 123 | 0.726985729 | 0.711154122 | 0.894448035 | 0.390709508 | 0.428241136 | 0.451346437 |
| rbpF | A0A0K2GHG7 | Putative RNA-binding protein RbpF | 11.195 | 110 | -0.933 | -1.91 | 1.06E-05 | 1.24E-04 | no | no | 0 | 34 | 41 | 44 | 71 | 65 | 68 | 0.725130797 | 0.753469826 | 0.785664074 | 1.386881044 | 1.294278547 | 1.365277568 |
| NITMOv2_4091 | A0A0K2GHP0 | Putative Response regulator, CheY-like | 16.041 | 144 | 4.15 | 17.75 | 1.51E-08 | 3.40E-06 | yes | no | 0 | 129 | 138 | 141 | 25 | 24 | 23 | 0.675363819 | 0.690563347 | 0.70425824 | 0.048149401 | 0.043994035 | 0.041033129 |
| NITMOv2_4476 | A0A0K2GIT2 | Uncharacterized protein | 17.752 | 169 | -1.27 | -2.41 | 6.56E-06 | 8.94E-05 | yes | no | 0 | 42 | 43 | 53 | 96 | 89 | 104 | 0.625259581 | 0.613323499 | 0.637231675 | 1.484891589 | 1.410726949 | 1.689119221 |
| tufB | A0A0K2G9S8 | Elongation factor Tu | 43.714 | 401 | -0.532 | -1.45 | 1.61E-03 | 4.08E-03 | no | no | 0 | 203 | 215 | 247 | 347 | 341 | 325 | 0.584956971 | 0.584813196 | 0.619765657 | 0.80798359 | 0.763743044 | 0.737193785 |
| modA | A0A0K2GAR7 | Molybdate ABC transporter, periplasmic binding protein | 27.524 | 257 | 0.985 | 1.98 | 3.76E-05 | 2.87E-04 | no | no | 0 | 93 | 89 | 85 | 60 | 60 | 58 | 0.581731919 | 0.564085541 | 0.5381681 | 0.327303986 | 0.307007396 | 0.273889842 |
| smbP | A0A0K2GGB0 | Metal-binding protein SmbP | 11.803 | 115 | 1.1 | 2.14 | 2.16E-05 | 1.99E-04 | yes | no | 0 | 59 | 56 | 58 | 51 | 48 | 40 | 0.526336909 | 0.529161397 | 0.463280868 | 0.310245797 | 0.342959577 | 0.323931559 |
| NITMOv2_0132 | A0A0K2G7I8 | FGE-sulfatase domain-containing protein | 35.065 | 321 | 0.486 | 1.40 | 5.06E-04 | 1.71E-03 | no | no | 0 | 159 | 164 | 172 | 124 | 137 | 133 | 0.494381492 | 0.483990752 | 0.47516773 | 0.318019711 | 0.346362139 | 0.356675348 |
| NITMOv2_0141 | A0A0K2G6I2 | PDZ domain-containing protein | 15.155 | 150 | 0.814 | 1.76 | 1.56E-04 | 7.54E-04 | no | no | 0 | 45 | 42 | 35 | 28 | 36 | 28 | 0.475368441 | 0.477146715 | 0.440369899 | 0.29783715 | 0.282210644 | 0.259396984 |
| NITMOv2_3174 | A0A0K2GF39 | Uncharacterized protein | 9.9944 | 89 | 0.653 | 1.57 | 4.58E-05 | 3.28E-04 | no | no | 0 | 29 | 28 | 27 | 24 | 23 | 24 | 0.474883629 | 0.485144461 | 0.479462338 | 0.351277255 | 0.348381925 | 0.319310387 |
| idh | A0A0K2G8E9 | Isocitrate dehydrogenase [NADP] | 81.549 | 743 | 0.221 | 1.17 | 1.45E-02 | 2.44E-02 | no | no | 0 | 392 | 426 | 417 | 414 | 421 | 392 | 0.450390098 | 0.45205843 | 0.451662419 | 0.433103252 | 0.433461505 | 0.413657813 |
| NITMOv2_0909 | A0A0K2G8Q7 | Uncharacterized protein | 16.123 | 149 | 1.04 | 2.06 | 6.30E-05 | 4.12E-04 | yes | no | 0 | 52 | 60 | 56 | 36 | 38 | 39 | 0.441789959 | 0.432582256 | 0.396043407 | 0.200981587 | 0.200145206 | 0.196211 |
| NITMOv2_0731 | A0A0K2G980 | Putative Cytochrome c | 19.55 | 182 | 0.454 | 1.37 | 5.04E-04 | 1.71E-03 | no | no | 0 | 47 | 50 | 50 | 33 | 38 | 41 | 0.435803588 | 0.439308967 | 0.396100924 | 0.262669444 | 0.289497409 | 0.300178376 |
| glnA | A0A0K2G9R8 | Glutamine synthetase | 52.98 | 469 | 0.171 | 1.13 | 3.71E-02 | 5.42E-02 | no | no | 0 | 150 | 150 | 196 | 228 | 213 | 196 | 0.434412389 | 0.433814183 | 0.43314192 | 0.406716367 | 0.385872251 | 0.36084699 |
| NITMOv2_4530 | A0A0K2GJ77 | Copper-containing nitrite reductase | 36.449 | 329 | -1.57 | -2.97 | 9.29E-07 | 2.93E-05 | yes | no | 0 | 91 | 95 | 101 | 223 | 213 | 221 | 0.432810403 | 0.435593632 | 0.433122748 | 1.262143043 | 1.29538166 | 1.297308582 |
| NITMOv2_2089 | A0A0K2GC20 | Heat shock protein, Hsp20 family | 16.999 | 148 | 2.53 | 5.78 | 3.30E-07 | 1.46E-05 | yes | no | 0 | 44 | 46 | 57 | 18 | 19 | 15 | 0.422755829 | 0.363359709 | 0.454384893 | 0.065881329 | 0.068675814 | 0.056667794 |
| hupB | A0A0K2G735 | HU, DNA-binding transcriptional regulator, beta subunit | 9.436 | 90 | -0.997 | -2.00 | 7.88E-06 | 1.01E-04 | no | no | 0 | 32 | 34 | 32 | 46 | 43 | 52 | 0.419530777 | 0.430548599 | 0.443034857 | 0.875313088 | 0.880797382 | 0.917797092 |
| NITMOv2_3764 | A0A0K2GGR9 | PepSY domain-containing protein | 12.388 | 117 | 0.859 | 1.81 | 7.98E-05 | 4.85E-04 | no | no | 0 | 18 | 21 | 20 | 18 | 13 | 12 | 0.41655867 | 0.425620892 | 0.408965576 | 0.214071377 | 0.203842967 | 0.185961709 |
| icd | A0A0K2GFQ1 | Isocitrate dehydrogenase (NAD(+)) | 37.189 | 338 | 1.08 | 2.11 | 1.75E-06 | 3.98E-05 | yes | no | 0 | 83 | 78 | 83 | 44 | 48 | 44 | 0.415926307 | 0.394118767 | 0.361686541 | 0.176282752 | 0.187389484 | 0.178395214 |
| fabI | A0A0K2G6S1 | Enoyl-[acyl-carrier-protein] reductase [NADH] | 27.612 | 259 | 0.112 | 1.08 | 1.19E-01 | 1.50E-01 | no | no | 0 | 40 | 45 | 45 | 49 | 49 | 45 | 0.403616305 | 0.406692241 | 0.367016456 | 0.263054438 | 0.275203542 | 0.294586218 |
| NITMOv2_4244 | A0A0K2GJ08 | Isochorismatase family protein | 29.246 | 268 | 0.772 | 1.71 | 9.51E-05 | 5.41E-04 | no | no | 0 | 104 | 108 | 137 | 119 | 104 | 103 | 0.390252364 | 0.414865977 | 0.396024235 | 0.28957459 | 0.278233221 | 0.258012431 |
| NITMOv2_2298 | A0A0K2GCW8 | LysM domain-containing protein | 24.26 | 218 | 0.765 | 1.70 | 5.82E-05 | 3.90E-04 | no | no | 0 | 87 | 87 | 78 | 58 | 70 | 68 | 0.35481895 | 0.347012237 | 0.332276143 | 0.166272912 | 0.20412263 | 0.222661365 |
| htrA | A0A0K2GGK4 | Putative periplasmic serine endoprotease DegP-like | 52.484 | 504 | 0.404 | 1.32 | 1.84E-03 | 4.50E-03 | no | no | 0 | 138 | 130 | 144 | 132 | 126 | 123 | 0.354039036 | 0.327223193 | 0.436458738 | 0.382595023 | 0.367026099 | 0.365701917 |
| NITMOv2_0430 | A0A0K2G7B1 | GYD family protein | 11.313 | 101 | 1.06 | 2.08 | 4.43E-06 | 7.00E-05 | yes | no | 0 | 21 | 20 | 24 | 19 | 17 | 17 | 0.342804051 | 0.351881281 | 0.321041141 | 0.174994503 | 0.171122442 | 0.172756305 |
| aclA | A0A0K2G940 | ATP citrate lyase, alpha subunit | 65.343 | 606 | 0.205 | 1.15 | 2.50E-02 | 3.87E-02 | no | no | 0 | 216 | 222 | 216 | 210 | 206 | 200 | 0.339326054 | 0.339972656 | 0.33338814 | 0.296075063 | 0.30177149 | 0.290288709 |
| NITMOv2_2239 | A0A0K2GCH1 | Putative Regulator protein, AsnC family | 8.6089 | 78 | 0.882 | 1.84 | 6.47E-05 | 4.19E-04 | no | no | 0 | 27 | 38 | 32 | 20 | 21 | 26 | 0.330957782 | 0.325521961 | 0.284824556 | 0.165858304 | 0.185027888 | 0.182832977 |
| NITMOv2_3932 | A0A0K2GI74 | Heat shock protein, Hsp20 family | 17.901 | 157 | 3.01 | 8.06 | 6.69E-08 | 7.15E-06 | yes | no | 0 | 54 | 44 | 60 | 14 | 11 | 13 | 0.327943517 | 0.390696749 | 0.523309522 | 0.057940091 | 0.050458903 | 0.057543479 |
| atpD | A0A0K2G7Z8 | ATP synthase subunit beta | 52.336 | 480 | -0.287 | -1.22 | 2.13E-03 | 5.08E-03 | no | no | 0 | 198 | 196 | 226 | 244 | 240 | 236 | 0.305157366 | 0.304109902 | 0.307083664 | 0.365196263 | 0.372137709 | 0.355974081 |
| NITMOv2_2049 | A0A0K2GBY7 | Putative Murein lipoprotein (Modular protein) | 14.666 | 135 | 0.493 | 1.41 | 1.84E-04 | 8.22E-04 | no | no | 0 | 19 | 20 | 18 | 16 | 19 | 18 | 0.295861628 | 0.298321802 | 0.326351884 | 0.211035849 | 0.235662357 | 0.23384568 |
| rbr | A0A0K2GF07 | Rubrerythrin | 15.701 | 141 | 0.665 | 1.59 | 1.22E-04 | 6.37E-04 | no | no | 0 | 44 | 48 | 47 | 46 | 46 | 41 | 0.295545447 | 0.300805209 | 0.291093917 | 0.197635102 | 0.214920714 | 0.200166867 |
| NITMOv2_1100 | A0A0K2G985 | Uncharacterized protein | 7.1031 | 58 | 0.917 | 1.89 | 6.07E-02 | 8.32E-02 | no | no | 0 | 18 | 20 | 16 | 14 | 11 | 14 | 0.29050762 | 0.294234934 | 0.293586324 | 0.150651047 | 0.244394045 | 0.252168536 |
| atpA | A0A0K2G725 | ATP synthase subunit alpha | 54.446 | 505 | -0.0557 | -1.04 | 4.64E-01 | 5.08E-01 | no | no | 0 | 90 | 88 | 99 | 111 | 115 | 113 | 0.285448715 | 0.281896112 | 0.270291908 | 0.31180058 | 0.313750372 | 0.293992838 |
| aclB | A0A0K2G827 | ATP citrate synthase | 43.167 | 398 | 0.187 | 1.14 | 1.16E-02 | 2.02E-02 | no | no | 0 | 182 | 198 | 199 | 191 | 190 | 182 | 0.274267967 | 0.283616899 | 0.30363264 | 0.276499607 | 0.280579279 | 0.275184489 |
| ihfA | A0A0K2GJN3 | Integration host factor subunit alpha | 10.102 | 91 | 0.663 | 1.58 | 2.73E-04 | 1.09E-03 | no | no | 0 | 10 | 11 | 9 | 11 | 10 | 12 | 0.271389175 | 0.267034775 | 0.250410173 | 0.174802006 | 0.190838656 | 0.193891424 |
| NITMOv2_0369 | A0A0K2G791 | Uncharacterized protein | 19.376 | 175 | 0.667 | 1.59 | 9.92E-05 | 5.55E-04 | no | no | 0 | 68 | 69 | 61 | 58 | 58 | 56 | 0.269618558 | 0.269244421 | 0.257331394 | 0.167664813 | 0.178688869 | 0.178458148 |
| bfrB | A0A0K2G8K8 | Bacterioferritin | 18.231 | 158 | -0.0861 | -1.06 | 4.12E-01 | 4.56E-01 | no | no | 0 | 71 | 78 | 76 | 83 | 82 | 78 | 0.263821896 | 0.285943872 | 0.305108911 | 0.328962421 | 0.313191047 | 0.299297296 |
| NITMOv2_0186 | A0A0K2G6M7 | Uncharacterized protein | 14.127 | 139 | -0.18 | -1.13 | 2.78E-01 | 3.21E-01 | no | no | 0 | 24 | 32 | 27 | 24 | 23 | 22 | 0.26375866 | 0.289013911 | 0.298954584 | 0.338483614 | 0.296877394 | 0.276011625 |
| NITMOv2_0198 | A0A0K2G6R5 | Putative Outer membrane protein, OmpA/MotB family | 44.52 | 415 | 0.371 | 1.29 | 2.47E-03 | 5.66E-03 | no | no | 0 | 95 | 99 | 108 | 94 | 96 | 86 | 0.259859088 | 0.247715036 | 0.266227368 | 0.221282608 | 0.220016788 | 0.207647052 |
| rplS | A0A0K2GG95 | 50S ribosomal protein L19 | 15.135 | 134 | -0.0604 | -1.04 | 2.83E-01 | 3.26E-01 | no | no | 0 | 31 | 27 | 29 | 27 | 28 | 29 | 0.257877683 | 0.26687834 | 0.272458385 | 0.298858864 | 0.306183945 | 0.276389231 |
| NITMOv2_1176 | A0A0K2GAH5 | Uncharacterized protein | 21.875 | 207 | 0.859 | 1.81 | 1.05E-04 | 5.76E-04 | no | no | 0 | 35 | 35 | 36 | 33 | 33 | 33 | 0.254399686 | 0.260484053 | 0.245521221 | 0.181598628 | 0.176234053 | 0.179141434 |
| dnaK | A0A0K2GDF1 | Chaperone protein DnaK | 69.374 | 639 | -0.0128 | -1.01 | 8.63E-01 | 8.80E-01 | no | no | 0 | 204 | 200 | 199 | 215 | 224 | 210 | 0.252228572 | 0.243158862 | 0.246671563 | 0.274367334 | 0.2724846 | 0.248104782 |
| NITMOv2_1689 | A0A0K2GB62 | Putative Periplasmic chaperone Skp | 20.542 | 182 | 0.406 | 1.33 | 1.41E-03 | 3.68E-03 | no | no | 0 | 53 | 58 | 57 | 51 | 47 | 48 | 0.248329 | 0.260171183 | 0.258692632 | 0.210236246 | 0.216769594 | 0.218309911 |
| NITMOv2_3005 | A0A0K2GEN9 | UPF0753 protein NITMOv2_3005 | 121.19 | 1083 | 0.984 | 1.98 | 3.69E-06 | 6.29E-05 | no | no | 0 | 301 | 308 | 302 | 203 | 205 | 191 | 0.244408348 | 0.246698207 | 0.246901631 | 0.131269569 | 0.127300865 | 0.123122762 |
| NITMOv2_4118 | A0A0K2GHS3 | Uncharacterized protein | 16.295 | 144 | 0.809 | 1.75 | 1.19E-05 | 1.34E-04 | no | no | 0 | 30 | 37 | 36 | 28 | 29 | 30 | 0.239939649 | 0.23664725 | 0.256468638 | 0.138300149 | 0.148768078 | 0.157545997 |
| acnA | A0A0K2G845 | Aconitate hydratase A | 80.839 | 748 | -0.175 | -1.13 | 3.34E-02 | 4.95E-02 | no | no | 0 | 218 | 230 | 236 | 282 | 287 | 273 | 0.238780317 | 0.23424206 | 0.225486107 | 0.303153026 | 0.297840677 | 0.275795851 |
| rpsG | A0A0K2G9L3 | 30S ribosomal protein S7 | 17.754 | 156 | -0.088 | -1.06 | 3.44E-01 | 3.87E-01 | no | no | 0 | 29 | 31 | 30 | 30 | 32 | 31 | 0.237410197 | 0.24016704 | 0.219542676 | 0.245759331 | 0.263488787 | 0.247475439 |
| NITMOv2_4282 | A0A0K2GIG8 | Uncharacterized protein | 25.421 | 230 | 0.835 | 1.78 | 1.34E-05 | 1.42E-04 | no | no | 0 | 42 | 44 | 43 | 30 | 32 | 34 | 0.231234117 | 0.2268505 | 0.221785842 | 0.132523761 | 0.140674953 | 0.133449373 |
| rplJL | A0A0K2G9H7 | Multifunctional fusion protein | 32.782 | 312 | -0.191 | -1.14 | 1.09E-01 | 1.39E-01 | no | no | 0 | 74 | 77 | 69 | 83 | 90 | 86 | 0.23066499 | 0.210131494 | 0.163879568 | 0.249786959 | 0.256932252 | 0.242206944 |
| rplP | A0A0K2G9J2 | 50S ribosomal protein L16 | 16.262 | 144 | -0.205 | -1.15 | 2.83E-02 | 4.30E-02 | no | no | 0 | 15 | 16 | 18 | 16 | 14 | 16 | 0.228177695 | 0.236353934 | 0.225141004 | 0.290314963 | 0.283142853 | 0.261195105 |
| NITMOv2_1735 | A0A0K2GB32 | Uncharacterized protein | 15.559 | 144 | 0.574 | 1.49 | 1.42E-04 | 7.06E-04 | no | no | 0 | 39 | 40 | 39 | 30 | 31 | 34 | 0.221896221 | 0.228649504 | 0.224546661 | 0.13260224 | 0.165886537 | 0.144444885 |
| rpoA | A0A0K2G9L2 | DNA-directed RNA polymerase subunit alpha | 36.552 | 332 | -0.505 | -1.42 | 8.78E-04 | 2.57E-03 | no | no | 0 | 81 | 88 | 92 | 135 | 130 | 131 | 0.22088444 | 0.217894588 | 0.220731362 | 0.310852903 | 0.301445217 | 0.284444814 |
| iscS | A0A0K2GIC1 | Cysteine desulfurase IscS | 44.16 | 405 | 1.22 | 2.33 | 2.28E-06 | 4.57E-05 | yes | no | 0 | 129 | 154 | 138 | 93 | 85 | 80 | 0.220336392 | 0.214765885 | 0.217433716 | 0.100524851 | 0.102112586 | 0.100790095 |
| rpmI | A0A0K2G751 | 50S ribosomal protein L35 | 7.2185 | 64 | -0.507 | -1.42 | 3.06E-04 | 1.18E-03 | no | no | 0 | 10 | 11 | 10 | 10 | 11 | 11 | 0.212326459 | 0.235610867 | 0.230700988 | 0.272116601 | 0.297638698 | 0.286332842 |
| NITMOv2_3786 | A0A0K2GHR9 | Putative dCTP deaminase | 34.646 | 304 | 0.695 | 1.62 | 4.71E-05 | 3.34E-04 | no | no | 0 | 78 | 80 | 85 | 75 | 74 | 71 | 0.208555467 | 0.210952779 | 0.215842411 | 0.136974882 | 0.129601867 | 0.135754565 |

| | | | | | | | | | | | | | | | | | | | | | | | |
|---|---|---|---|---|---|---|---|---|---|---|---|---|---|---|---|---|---|---|---|---|---|---|---|
| NITMOv2_0279 | A0A0K2G6Y9 | Uncharacterized protein | 21.648 | 206 | 0.482 | 1.40 | 1.01E-03 | 2.83E-03 | no | no | 0 | 23 | 27 | 30 | 24 | 27 | 23 | 0.20435236 | 0.234046516 | 0.255241607 | 0.180754603 | 0.181547643 | 0.176530562 |
| NITMOv2_3538 | A0A0K2GG53 | Putative Outer membrane chaperone Skp | 20.102 | 179 | 0.389 | 1.31 | 1.05E-03 | 2.92E-03 | no | no | 0 | 34 | 37 | 34 | 31 | 33 | 33 | 0.202185462 | 0.201253801 | 0.195615572 | 0.141120969 | 0.155896367 | 0.151432385 |
| rpsJ | A0A0K2G9K4 | 30S ribosomal protein S10 | 11.918 | 103 | -0.255 | -1.19 | 1.61E-02 | 2.65E-02 | no | no | 0 | 32 | 29 | 32 | 31 | 34 | 33 | 0.201257997 | 0.210248821 | 0.191131158 | 0.246558933 | 0.241814934 | 0.245677318 |
| rpsH | A0A0K2G9M0 | 30S ribosomal protein S8 | 14.428 | 131 | -0.233 | -1.18 | 6.15E-02 | 8.40E-02 | no | no | 0 | 50 | 42 | 46 | 51 | 54 | 54 | 0.193903613 | 0.187870775 | 0.168971746 | 0.236001218 | 0.234326191 | 0.238934363 |
| cspE | A0A0K2GBE9 | DNA-binding transcriptional repressor | 7.8308 | 72 | -1.72 | -3.29 | 4.79E-07 | 1.85E-05 | yes | no | 0 | 12 | 11 | 12 | 17 | 18 | 18 | 0.193229093 | 0.204186959 | 0.20242176 | 0.611592314 | 0.645849718 | 0.619866368 |
| NITMOv2_0954 | A0A0K2G8V4 | Uncharacterized protein | 25.449 | 229 | 0.219 | 1.16 | 2.06E-02 | 3.26E-02 | no | no | 0 | 48 | 50 | 47 | 45 | 46 | 43 | 0.188842601 | 0.192133632 | 0.193180683 | 0.160379545 | 0.159330002 | 0.153843665 |
| NITMOv2_3178 | A0A0K2GG27 | Uncharacterized protein | 21.088 | 188 | 0.504 | 1.42 | 2.69E-04 | 1.08E-03 | no | no | 0 | 48 | 46 | 46 | 33 | 39 | 39 | 0.187310174 | 0.172876467 | 0.167877004 | 0.126276496 | 0.123140107 | 0.132739115 |
| NITMOv2_0375 | A0A0K2G754 | PHB domain-containing protein | 28.474 | 254 | 0.68 | 1.60 | 1.02E-04 | 5.64E-04 | no | no | 0 | 86 | 88 | 85 | 55 | 61 | 62 | 0.186393247 | 0.201468899 | 0.192701374 | 0.12185647 | 0.111983123 | 0.112592964 |
| prcA | A0A0K2GH61 | Proteasome endopeptidase complex | 25.38 | 231 | 0.68 | 1.60 | 2.83E-04 | 1.11E-03 | no | no | 0 | 38 | 37 | 38 | 32 | 34 | 32 | 0.182809856 | 0.188735079 | 0.173385223 | 0.127236019 | 0.103852709 | 0.116502079 |
| NITMOv2_0584 | A0A0K2G7W4 | Uncharacterized protein | 23.363 | 212 | 0.591 | 1.51 | 2.58E-04 | 1.05E-03 | no | no | 0 | 33 | 39 | 34 | 38 | 30 | 34 | 0.182795101 | 0.190641632 | 0.191411074 | 0.099578654 | 0.090306163 | 0.114326353 |
| NITMOv2_0460 | A0A0K2G7E2 | OsmC family protein | 20.35 | 186 | 2.11 | 4.32 | 3.41E-07 | 1.46E-05 | yes | no | 0 | 31 | 30 | 35 | 16 | 15 | 16 | 0.180976003 | 0.156836 | 0.194158473 | 0.045829073 | 0.043785842 | 0.040497289 |
| purE | A0A0K2GI72 | N5-carboxyaminoimidazole ribonucleotide mutase | 20.376 | 196 | 0.434 | 1.35 | 4.00E-03 | 8.32E-03 | no | no | 0 | 62 | 53 | 69 | 67 | 68 | 63 | 0.180166578 | 0.172387607 | 0.195097919 | 0.169959969 | 0.160805999 | 0.139406549 |
| clpP | A0A0K2G6Z7 | ATP-dependent Clp protease proteolytic subunit | 23.175 | 210 | 0.433 | 1.35 | 4.89E-04 | 1.66E-03 | no | no | 0 | 60 | 60 | 70 | 54 | 57 | 62 | 0.177723549 | 0.184728384 | 0.200753765 | 0.157121905 | 0.159500907 | 0.151421596 |
| hucL | A0A088NAA0 | Putative NiFe hydrogenase large subunit | 59.503 | 531 | 1.17 | 2.25 | 1.64E-05 | 1.64E-04 | yes | no | 0 | 126 | 132 | 126 | 93 | 88 | 85 | 0.176503088 | 0.169452493 | 0.160338433 | 0.079818105 | 0.076677272 | 0.072707036 |
| NITMOv2_1470 | A0A0K2GBD3 | Uncharacterized protein | 9.711 | 80 | 1.97 | 3.92 | 3.24E-07 | 1.46E-05 | yes | no | 0 | 13 | 14 | 19 | 10 | 6 | 8 | 0.175019143 | 0.178529642 | 0.200427834 | 0.042316745 | 0.040258986 | 0.040900068 |
| porC | A0A0K2G9G6 | Pyruvate:ferredoxin oxidoreductase gamma subunit | 25.449 | 235 | -0.272 | -1.21 | 2.39E-03 | 5.51E-03 | no | no | 0 | 59 | 54 | 59 | 67 | 60 | 64 | 0.174650264 | 0.19023099 | 0.17752837 | 0.226805788 | 0.220529503 | 0.217213057 |
| fbaB | A0A0K2GFX6 | Fructose-bisphosphate aldolase class I | 32.864 | 307 | -0.0954 | -1.07 | 3.11E-01 | 3.53E-01 | no | no | 0 | 69 | 70 | 88 | 81 | 82 | 82 | 0.173387646 | 0.174726313 | 0.168373568 | 0.214071377 | 0.196447445 | 0.195851376 |
| NITMOv2_3663 | A0A0K2GGH1 | Uncharacterized protein | 39.565 | 371 | 0.788 | 1.73 | 3.49E-05 | 2.74E-04 | no | no | 0 | 122 | 127 | 121 | 90 | 86 | 82 | 0.17272788 | 0.177776798 | 0.169385869 | 0.10562898 | 0.109550058 | 0.105835624 |
| fecA | A0A0K2G9R9 | Iron(III) dicitrate transport protein FecA | 89.323 | 810 | 0.998 | 2.00 | 7.33E-06 | 9.68E-05 | no | no | 0 | 142 | 135 | 148 | 94 | 96 | 101 | 0.170655837 | 0.1510479 | 0.16665956 | 0.085247999 | 0.084681838 | 0.084049586 |
| ahcY | A0A0K2G8F8 | Adenosylhomocysteinase | 45.876 | 418 | 0.186 | 1.14 | 2.84E-02 | 4.30E-02 | no | no | 0 | 91 | 101 | 108 | 102 | 99 | 84 | 0.170147839 | 0.182422921 | 0.190009575 | 0.16773885 | 0.162421828 | 0.149387921 |
| NITMOv2_4015 | A0A0K2GHT1 | Uncharacterized protein | 21.494 | 203 | 0.276 | 1.21 | 4.88E-02 | 6.88E-02 | no | no | 0 | 23 | 26 | 25 | 18 | 20 | 18 | 0.169591359 | 0.181851933 | 0.152203601 | 0.143895886 | 0.132816433 | 0.134975978 |
| NITMOv2_0998 | A0A0K2G927 | Uncharacterized protein | 18.198 | 182 | 0.733 | 1.66 | 3.19E-04 | 1.21E-03 | no | no | 0 | 23 | 22 | 18 | 17 | 18 | 16 | 0.168027314 | 0.160371434 | 0.135782476 | 0.097665531 | 0.090299949 | 0.08064754 |
| NITMOv2_4529 | A0A0K2GIY6 | Plastocyanin-like domain-containing protein | 179.2 | 1610 | -1.93 | -3.81 | 7.66E-08 | 7.47E-06 | yes | no | 0 | 303 | 321 | 337 | 932 | 890 | 884 | 0.167510884 | 0.164479811 | 0.171205326 | 0.673768818 | 0.662707159 | 0.663237055 |
| NITMOv2_2960 | A0A0K2GEH7 | Uncharacterized protein | 63.357 | 590 | 1.32 | 2.50 | 7.50E-05 | 4.61E-04 | yes | no | 0 | 136 | 144 | 156 | 76 | 73 | 95 | 0.166838472 | 0.167176362 | 0.16828921 | 0.063088643 | 0.062235806 | 0.080048766 |
| fkpA | A0A0K2GB55 | Peptidyl-prolyl cis-trans isomerase | 23.989 | 226 | 0.292 | 1.22 | 1.66E-02 | 2.72E-02 | no | no | 0 | 35 | 36 | 38 | 42 | 36 | 37 | 0.166448514 | 0.172446271 | 0.160016337 | 0.152546402 | 0.13683736 | 0.127619864 |
| rplF | A0A0K2G9V7 | 50S ribosomal protein L6 | 19.166 | 178 | -0.352 | -1.28 | 1.07E-03 | 2.95E-03 | no | no | 0 | 38 | 42 | 40 | 47 | 44 | 47 | 0.16600586 | 0.166525201 | 0.158854493 | 0.216973638 | 0.223481497 | 0.205920855 |
| pal.2 | A0A0K2GHI3 | Peptidoglycan-associated protein | 25.849 | 246 | 0.199 | 1.15 | 3.44E-02 | 5.08E-02 | no | no | 0 | 56 | 63 | 52 | 57 | 61 | 57 | 0.164555641 | 0.172158821 | 0.162094621 | 0.156174228 | 0.158801751 | 0.165283313 |
| sucC | A0A0K2G945 | Succinate--CoA ligase [ADP-forming] subunit beta | 42.123 | 392 | 0.206 | 1.15 | 2.55E-02 | 3.92E-02 | no | no | 0 | 110 | 111 | 112 | 97 | 105 | 89 | 0.164500836 | 0.160349924 | 0.162200069 | 0.15127296 | 0.146625551 | 0.145843823 |
| rplW | A0A0K2G9I7 | 50S ribosomal protein L23 | 10.975 | 97 | -0.48 | -1.39 | 2.03E-04 | 8.87E-04 | no | no | 0 | 13 | 13 | 16 | 22 | 20 | 18 | 0.164138281 | 0.16689869 | 0.150317041 | 0.249860996 | 0.234357265 | 0.221438643 |
| rpsS | A0A0K2G9L0 | 30S ribosomal protein S19 | 10.817 | 95 | -0.243 | -1.18 | 1.20E-02 | 2.08E-02 | no | no | 0 | 15 | 17 | 19 | 21 | 23 | 23 | 0.162165308 | 0.165281541 | 0.162447392 | 0.207037836 | 0.215557723 | 0.20628048 |
| NITMOv2_0465 | A0A0K2G7H1 | CrtC domain-containing protein | 43.493 | 394 | 1.52 | 2.87 | 2.40E-03 | 5.53E-03 | yes | no | 0 | 5 | 5 | 6 | 0 | 4 | 2 | 0.158503926 | 0.14557267 | 0.129229364 | 0.018136171 | 0.094673561 | 0.11084519 |
| NITMOv2_4499 | A0A0K2GIT7 | Uncharacterized protein | 43.884 | 412 | 0.602 | 1.52 | 4.64E-05 | 3.31E-04 | no | no | 0 | 89 | 88 | 91 | 74 | 77 | 72 | 0.158377453 | 0.1549666 | 0.168526947 | 0.106887614 | 0.114653901 | 0.096961895 |
| NITMOv2_4065 | A0A0K2GHX6 | Urate_ox_N domain-containing protein | 20.857 | 195 | 0.0191 | 1.01 | 8.37E-01 | 8.57E-01 | no | no | 0 | 30 | 32 | 29 | 29 | 26 | 21 | 0.158213039 | 0.170134941 | 0.169234407 | 0.136446256 | 0.131483375 | 0.128639399 |
| ppiA | A0A0K2G8B9 | Peptidyl-prolyl cis-trans isomerase | 22.909 | 210 | 0.595 | 1.51 | 2.52E-04 | 1.03E-03 | no | no | 0 | 49 | 45 | 52 | 36 | 41 | 40 | 0.157175963 | 0.155400707 | 0.142772718 | 0.101525835 | 0.103364853 | 0.088309335 |
| NITMOv2_4457 | A0A0K2GIR6 | Transcriptional regulator, LuxR family | 10.419 | 92 | 0.538 | 1.45 | 7.09E-04 | 2.20E-03 | no | no | 0 | 22 | 24 | 27 | 20 | 18 | 23 | 0.153529336 | 0.164802459 | 0.142851325 | 0.097714396 | 0.086246394 | 0.08443798 |
| mrp | A0A0K2GD16 | Iron-sulfur cluster carrier protein | 30.563 | 290 | 0.62 | 1.54 | 6.03E-04 | 1.96E-03 | no | no | 0 | 78 | 85 | 86 | 70 | 66 | 60 | 0.153407079 | 0.168423932 | 0.161408251 | 0.113476931 | 0.110281842 | 0.096758707 |
| rplE | A0A0K2G9J7 | 50S ribosomal protein L5 | 24.03 | 215 | -0.155 | -1.11 | 6.61E-02 | 8.94E-02 | no | no | 0 | 48 | 50 | 49 | 56 | 61 | 63 | 0.153175212 | 0.151718615 | 0.1573092 | 0.175083348 | 0.183101324 | 0.185494197 |
| NITMOv2_1414 | A0A0K2GA41 | Dienelactone hydrolase-like enzyme | 25.785 | 235 | 0.53 | 1.44 | 9.22E-05 | 5.37E-04 | no | no | 0 | 53 | 57 | 60 | 43 | 45 | 46 | 0.152662998 | 0.154481651 | 0.153438301 | 0.109280499 | 0.110434103 | 0.108883439 |
| yfhL | A0A0K2GJ12 | Putative ferredoxin-like protein YfhL | 12.374 | 115 | -1.53 | -2.89 | 5.89E-06 | 8.24E-05 | yes | no | 0 | 5 | 7 | 7 | 14 | 13 | 14 | 0.147064477 | 0.118808574 | 0.127206681 | 0.451967949 | 0.372355225 | 0.404882981 |
| rpsD | A0A0K2G9P3 | 30S ribosomal protein S4 | 23.543 | 208 | -0.337 | -1.26 | 3.34E-03 | 7.19E-03 | no | no | 0 | 38 | 40 | 46 | 44 | 43 | 46 | 0.146876876 | 0.152643538 | 0.146653204 | 0.194081313 | 0.190652214 | 0.191427997 |
| NITMOv2_2386 | A0A0K2GCW2 | Uncharacterized protein | 11.215 | 104 | 0.789 | 1.73 | 1.05E-02 | 1.85E-02 | no | no | 0 | 11 | 11 | 13 | 6 | 5 | 8 | 0.146427898 | 0.147958306 | 0.139603527 | 0.109308633 | 0.124174859 | 0.123557908 |
| NITMOv2_1775 | A0A0K2GB69 | Uncharacterized protein | 14.418 | 130 | 0.192 | 1.14 | 3.13E-02 | 4.68E-02 | no | no | 0 | 48 | 43 | 52 | 45 | 49 | 42 | 0.146145442 | 0.136651956 | 0.146403963 | 0.141397869 | 0.117069875 | 0.102607996 |
| NITMOv2_2507 | A0A0K2GD97 | OmpA-like domain-containing protein | 56.908 | 513 | 0.578 | 1.49 | 1.40E-04 | 7.01E-04 | no | no | 0 | 83 | 82 | 85 | 57 | 67 | 65 | 0.146067451 | 0.143959433 | 0.137981546 | 0.105158103 | 0.107684087 | 0.106362473 |
| ndk | A0A0K2GCA5 | Nucleoside diphosphate kinase | 15.096 | 139 | 0.429 | 1.35 | 6.34E-04 | 2.04E-03 | no | no | 0 | 16 | 19 | 19 | 13 | 15 | 14 | 0.145378175 | 0.1501132 | 0.138545213 | 0.105522367 | 0.114688082 | 0.113925372 |
| NITMOv2_2963 | A0A0K2GEJ1 | Uncharacterized protein | 8.7486 | 75 | 1.16 | 2.23 | 1.26E-05 | 1.39E-04 | yes | no | 0 | 27 | 26 | 31 | 22 | 19 | 18 | 0.144686791 | 0.141573797 | 0.130295348 | 0.031932277 | 0.039249093 | 0.060053656 |
| NITMOv2_1097 | A0A0K2G998 | IPT/TIG domain-containing protein | 16.944 | 157 | 1.05 | 2.07 | 1.93E-02 | 3.09E-02 | yes | no | 0 | 14 | 16 | 22 | 27 | 28 | 27 | 0.143809914 | 0.143668072 | 0.129407667 | 0.050614843 | 0.076417807 | 0.124836372 |
| rplD | A0A0K2G9L7 | 50S ribosomal protein L4 | 22.486 | 207 | -0.305 | -1.24 | 1.81E-02 | 2.92E-02 | no | no | 0 | 34 | 41 | 33 | 40 | 38 | 36 | 0.14330824 | 0.183586408 | 0.145525869 | 0.226746559 | 0.208239885 | 0.197487666 |
| rpsE | A0A0K2G9K2 | 30S ribosomal protein S5 | 17.945 | 169 | -0.409 | -1.33 | 5.25E-03 | 1.04E-02 | no | no | 0 | 24 | 26 | 23 | 30 | 31 | 26 | 0.142918282 | 0.1450623 | 0.137329686 | 0.206889761 | 0.196929086 | 0.173464765 |
| rpsO | A0A0K2GIR7 | 30S ribosomal protein S15 | 10.49 | 89 | -0.534 | -1.45 | 2.15E-03 | 5.10E-03 | no | no | 0 | 10 | 8 | 9 | 12 | 11 | 11 | 0.142559943 | 0.146368534 | 0.139871941 | 0.25495476 | 0.249101699 | 0.17558475 |
| ihfB | A0A0K2G866 | Integration host factor (IHF), DNA-binding protein, beta subunit | 10.406 | 92 | 0.472 | 1.39 | 2.63E-03 | 5.97E-03 | no | no | 0 | 5 | 5 | 6 | 4 | 5 | 6 | 0.141940227 | 0.160031187 | 0.154350906 | 0.088892114 | 0.120489527 | 0.120560439 |
| rpsB | A0A0K2GB56 | 30S ribosomal protein S2 | 28.641 | 257 | -0.363 | -1.29 | 1.11E-03 | 3.03E-03 | no | no | 0 | 63 | 63 | 59 | 64 | 59 | 64 | 0.14069658 | 0.143734557 | 0.142498554 | 0.193977661 | 0.183443134 | 0.18168218 |
| NITMOv2_2241 | A0A0K2GDG4 | Putative Carboxymethylenebutenolidase | 29.432 | 266 | 0.387 | 1.31 | 3.67E-03 | 7.78E-03 | no | no | 0 | 69 | 66 | 69 | 72 | 67 | 61 | 0.140232847 | 0.132989419 | 0.138280635 | 0.118278989 | 0.116087948 | 0.099919805 |
| NITMOv2_0733 | A0A0K2G8I5 | Putative Monoheme cytochrome c | 29.076 | 268 | 0.0699 | 1.05 | 3.53E-01 | 3.96E-01 | no | no | 0 | 53 | 55 | 57 | 56 | 61 | 54 | 0.139608915 | 0.140903081 | 0.133744455 | 0.12717827 | 0.13426291 | 0.130784557 |
| NITMOv2_0196 | A0A0K2G6N6 | Uncharacterized protein | 39.263 | 359 | 0.883 | 1.84 | 6.03E-05 | 4.00E-04 | no | no | 0 | 105 | 115 | 124 | 78 | 80 | 79 | 0.139589944 | 0.147375585 | 0.132026611 | 0.070847749 | 0.081293259 | 0.077757959 |
| glnB | A0A0K2G9Q0 | Nitrogen regulatory protein PII | 12.436 | 112 | 0.345 | 1.27 | 5.34E-03 | 1.05E-02 | no | no | 0 | 19 | 20 | 18 | 18 | 15 | 17 | 0.137452557 | 0.126994042 | 0.117647343 | 0.073347248 | 0.080598763 | 0.072442712 |
| glnB.1 | A0A0K2GFB2 | Nitrogen regulatory protein P-II | 11.94 | 109 | 0.921 | 1.89 | 2.04E-05 | 1.91E-04 | no | no | 0 | 35 | 31 | 29 | 20 | 20 | 20 | 0.134925212 | 0.127850525 | 0.12189402 | 0.076447929 | 0.068100952 | 0.063858481 |
| NITMOv2_4164 | A0A0K2GI61 | Putative Thiosulfate sulfurtransferase GlpE | 11.674 | 106 | 0.144 | 1.10 | 1.82E-01 | 2.20E-01 | no | no | 0 | 29 | 37 | 32 | 32 | 29 | 28 | 0.132402084 | 0.150922752 | 0.16104781 | 0.149673755 | 0.150139978 | 0.136560123 |
| NITMOv2_0376 | A0A0K2G882 | Membrane-bound serine protease | 46.597 | 446 | 1.01 | 2.01 | 6.96E-06 | 9.34E-05 | yes | no | 0 | 36 | 36 | 37 | 28 | 28 | 26 | 0.130838039 | 0.068649603 | 0.066811834 | 0.033272351 | 0.03347406 | 0.033320986 |
| rpsM | A0A0K2G9N0 | 30S ribosomal protein S13 | 14.373 | 130 | -0.352 | -1.28 | 2.29E-03 | 5.35E-03 | no | no | 0 | 13 | 15 | 11 | 17 | 15 | 16 | 0.130753724 | 0.128038247 | 0.121631359 | 0.180843448 | 0.16265488 | 0.185422272 |
| NITMOv2_3860 | A0A0K2GHZ6 | Uncharacterized protein | 18.335 | 176 | 0.866 | 1.82 | 5.82E-05 | 3.90E-04 | no | no | 0 | 14 | 13 | 15 | 11 | 10 | 10 | 0.130540828 | 0.124917366 | 0.120452259 | 0.072184862 | 0.079093246 | 0.077693227 |
| NMRAL | A0A0K2GHQ7 | NmrA-like family domain-containing protein 1 | 33.895 | 316 | 0.408 | 1.33 | 2.29E-03 | 5.34E-03 | no | no | 0 | 87 | 78 | 82 | 74 | 69 | 67 | 0.130041261 | 0.121520768 | 0.124430523 | 0.108038153 | 0.096651397 | 0.093829568 |
| NITMOv2_3173 | A0A0K2GG26 | Uncharacterized protein | 16.831 | 147 | 0.149 | 1.11 | 2.47E-01 | 2.88E-01 | no | no | 0 | 29 | 28 | 41 | 42 | 37 | 35 | 0.129453164 | 0.103346916 | 0.14029565 | 0.101752389 | 0.090223818 | 0.121544012 |
| NITMOv2_4243 | A0A0K2GI60 | Putative peroxidase-related enzyme | 19.809 | 189 | 1.68 | 3.20 | 9.15E-07 | 2.93E-05 | yes | no | 0 | 15 | 14 | 15 | 8 | 8 | 7 | 0.129398359 | 0.110593774 | 0.117444116 | 0.040824153 | 0.041372975 | 0.041146411 |
| rplM | A0A0K2GC42 | 50S ribosomal protein L13 | 15.827 | 142 | -0.287 | -1.22 | 6.84E-03 | 1.29E-02 | no | no | 0 | 28 | 24 | 26 | 28 | 36 | 30 | 0.128723838 | 0.131399647 | 0.122520956 | 0.147864284 | 0.182899345 | 0.188047529 |
| cld1 | A0A0K2GA28 | Chlorite dismutase | 33.743 | 297 | 0.528 | 1.44 | 1.12E-03 | 3.06E-03 | no | no | 0 | 93 | 95 | 91 | 79 | 82 | 81 | 0.127071262 | 0.125742561 | 0.122099164 | 0.094878768 | 0.085865743 | 0.084576435 |
| NITMOv2_3819 | A0A0K2GH77 | Putative Universal stress protein UspA | 30.386 | 285 | 0.539 | 1.45 | 2.85E-04 | 1.11E-03 | no | no | 0 | 58 | 62 | 62 | 53 | 45 | 45 | 0.126407281 | 0.131610834 | 0.132791589 | 0.090901486 | 0.088485249 | 0.088931485 |
| rpsC | A0A0K2G9M2 | 30S ribosomal protein S3 | 25.224 | 223 | -0.167 | -1.12 | 1.31E-01 | 1.64E-01 | no | no | 0 | 41 | 43 | 39 | 46 | 50 | 45 | 0.126261838 | 0.125429691 | 0.110166288 | 0.124672848 | 0.131376171 | 0.131226895 |
| NITMOv2_2413 | A0A0K2GD02 | Uncharacterized protein | 11.691 | 100 | 0.221 | 1.17 | 4.15E-02 | 5.96E-02 | no | no | 0 | 11 | 13 | 14 | 13 | 11 | 11 | 0.125094074 | 0.125766026 | 0.126930599 | 0.107275569 | 0.09343217 | 0.095911792 |
| NITMOv2_1993 | A0A0K2GBU2 | Group 1 truncated hemoglobin | 15.364 | 147 | 0.318 | 1.25 | 4.35E-03 | 8.89E-03 | no | no | 0 | 34 | 33 | 35 | 39 | 34 | 30 | 0.125064563 | 0.131820066 | 0.136967328 | 0.127674319 | 0.113769856 | 0.10319778 |
| bcp | A0A0K2GAH9 | Peroxiredoxin | 17.109 | 155 | 0.323 | 1.25 | 9.44E-03 | 1.70E-02 | no | no | 0 | 37 | 41 | 37 | 41 | 30 | 35 | 0.124860099 | 0.119917308 | 0.123052031 | 0.102652682 | 0.101331085 | 0.090193766 |
| NITMOv2_0136 | A0A0K2G6H7 | FGE-sulfatase domain-containing protein | 30.976 | 282 | 0.134 | 1.10 | 1.56E-01 | 1.92E-01 | no | no | 0 | 65 | 65 | 62 | 57 | 60 | 59 | 0.123825132 | 0.141292214 | 0.126959357 | 0.131888521 | 0.123210023 | 0.115732484 |
| NITMOv2_0734 | A0A0K2G8A8 | Putative Quinol-cytochrome c reductase, FeS subunit, modulated with PRC-barrel (Modular p | 35.247 | 312 | -0.0817 | -1.06 | 2.28E-01 | 2.70E-01 | no | no | 0 | 78 | 84 | 82 | 80 | 81 | 78 | 0.122360157 | 0.135611663 | 0.129093241 | 0.14098326 | 0.139884129 | 0.140084441 |
| rpsA | A0A0K2G782 | 30S ribosomal protein S1 | 63.548 | 569 | -0.566 | -1.48 | 1.45E-04 | 7.17E-04 | no | no | 0 | 85 | 82 | 84 | 118 | 114 | 115 | 0.122345402 | 0.120672107 | 0.119048842 | 0.182176119 | 0.188414913 | 0.190798655 |
| recA | A0A0K2GE94 | Protein RecA | 38.533 | 357 | -0.0943 | -1.07 | 1.85E-01 | 2.22E-01 | no | no | 0 | 75 | 89 | 93 | 90 | 90 | 87 | 0.122322215 | 0.119837135 | 0.126407193 | 0.118671387 | 0.115472691 | 0.116766403 |
| NITMOv2_1317 | A0A0K2G9X7 | Putative Thiol:disulfide interchange protein DsbE | 18.97 | 177 | 0.755 | 1.69 | 1.95E-04 | 8.58E-04 | no | no | 0 | 30 | 34 | 29 | 23 | 26 | 24 | 0.122214714 | 0.128703096 | 0.114758068 | 0.080903492 | 0.078036743 | 0.070892732 |
| NITMOv2_3829 | A0A0K2GGX5 | Putative Blue copper protein (Modular protein) | 18.401 | 167 | 0.889 | 1.85 | 1.57E-05 | 1.58E-04 | no | no | 0 | 17 | 19 | 21 | 11 | 12 | 10 | 0.121807893 | 0.120533271 | 0.118069135 | 0.046271816 | 0.045748142 | 0.047393084 |
| ccA1 | A0A0K2G8U7 | Cytochrome c peroxidase | 35.994 | 331 | 0.632 | 1.55 | 1.31E-04 | 6.66E-04 | no | no | 0 | 48 | 44 | 46 | 40 | 34 | 36 | 0.121457986 | 0.118908301 | 0.120247114 | 0.089109784 | 0.090189637 | 0.084952243 |
| rpoB | A0A0K2GAJ6 | DNA-directed RNA polymerase subunit beta | 147.11 | 1318 | -0.374 | -1.30 | 1.09E-03 | 3.01E-03 | no | no | 0 | 332 | 336 | 330 | 428 | 407 | 393 | 0.120193259 | 0.125148108 | 0.127185591 | 0.179333088 | 0.176886599 | 0.169422589 |
| sgaA | A0A0K2GGS4 | Serine-glyoxylate transaminase | 40.913 | 380 | -0.104 | -1.07 | 2.34E-01 | 2.76E-01 | no | no | 0 | 113 | 116 | 142 | 149 | 146 | 151 | 0.119773792 | 0.118781198 | 0.119223311 | 0.137262146 | 0.134099774 | 0.123716142 |
| trxA | A0A0K2GBA9 | Thioredoxin | 14.634 | 132 | -0.153 | -1.11 | 2.52E-01 | 2.94E-01 | no | no | 0 | 26 | 28 | 27 | 28 | 27 | 27 | 0.118070627 | 0.112398644 | 0.10808417 | 0.12684066 | 0.120340374 | 0.100728959 |

| | | | | | | | | | | | | | | | | | | | | | |
|---|---|---|---|---|---|---|---|---|---|---|---|---|---|---|---|---|---|---|---|---|---|
| NITMOv2_0449 | A0A0K2G7H0 | Uncharacterized protein | 22.745 | 203 | 0.34 | 1.27 | 1.62E-03 | 4.09E-03 | no | no | 0 | 53 | 53 | 58 | 56 | 56 | 51 | 0.117933615 | 0.120830498 | 0.119863667 | 0.112894998 | 0.108005699 | 0.107457529 |
| rpsL | A0A0K2G9T8 | 30S ribosomal protein S12 | 13.594 | 123 | -0.607 | -1.52 | 9.53E-05 | 5.41E-04 | no | no | 0 | 7 | 7 | 8 | 10 | 10 | 10 | 0.117794495 | 0.12262168 | 0.126338173 | 0.224451403 | 0.216614226 | 0.189144383 |
| NITMOv2_0135 | A0A0K2G6L8 | Uncharacterized protein | 13.196 | 122 | 0.332 | 1.26 | 1.15E-01 | 1.46E-01 | no | no | 0 | 12 | 11 | 9 | 10 | 12 | 7 | 0.116951345 | 0.11576591 | 0.103511563 | 0.106930556 | 0.09894463 | 0.075085951 |
| atpF | A0A0K2GI75 | ATP synthase subunit b | 19.479 | 168 | 0.0267 | 1.02 | 6.67E-01 | 7.05E-01 | no | no | 0 | 28 | 27 | 26 | 24 | 25 | 29 | 0.115935348 | 0.11587737 | 0.126042918 | 0.112087992 | 0.125883908 | 0.122065467 |
| hucS | A0A088NA13 | Putative NiFe hydrogenase small subunit | 34.696 | 320 | 1.09 | 2.13 | 1.27E-04 | 6.52E-04 | yes | no | 0 | 51 | 55 | 54 | 39 | 34 | 36 | 0.115878435 | 0.1193033 | 0.101028742 | 0.051534386 | 0.045313111 | 0.043976654 |
| ppiB | A0A0K2GAE0 | Peptidyl-prolyl cis-trans isomerase | 18.439 | 169 | 0.564 | 1.48 | 6.96E-04 | 2.17E-03 | no | no | 0 | 23 | 27 | 22 | 22 | 22 | 20 | 0.115043716 | 0.113573863 | 0.100486165 | 0.089287473 | 0.089998534 | 0.082409699 |
| serA | A0A0K2GHQ9 | D-3-phosphoglycerate dehydrogenase | 56.496 | 531 | 0.0991 | 1.07 | 2.14E-01 | 2.55E-01 | no | no | 0 | 95 | 100 | 103 | 114 | 114 | 111 | 0.115024745 | 0.118466372 | 0.11842574 | 0.119875233 | 0.117633861 | 0.10620244 |
| NITMOv2_3640 | A0A0K2GHE3 | Putative Nitrite oxidoreductase, cytochrome c containing membrane subunit | 66.788 | 591 | -1.74 | -3.34 | 2.47E-07 | 1.33E-05 | yes | no | 0 | 110 | 114 | 122 | 289 | 290 | 290 | 0.114607385 | 0.10763715 | 0.107000932 | 0.387940514 | 0.419121028 | 0.412524997 |
| forC | A0A0K2G856 | 2-oxoglutarate:ferredoxin oxidoreductase, gamma subunit | 25.967 | 239 | -1.03 | -2.04 | 1.65E-06 | 3.80E-05 | yes | no | 0 | 28 | 28 | 26 | 40 | 39 | 41 | 0.113711537 | 0.108829968 | 0.115398425 | 0.268636849 | 0.255953433 | 0.25445215 |
| pepA | A0A0K2G6Y2 | Probable cytosol aminopeptidase | 53.962 | 502 | 0.369 | 1.29 | 1.93E-03 | 4.67E-03 | no | no | 0 | 111 | 113 | 122 | 123 | 122 | 118 | 0.113374277 | 0.112124883 | 0.110400191 | 0.091311652 | 0.088921833 | 0.083903938 |
| rplC | A0A0K2G9U3 | 50S ribosomal protein L3 | 22.433 | 206 | -0.185 | -1.14 | 2.24E-02 | 3.51E-02 | no | no | 0 | 37 | 34 | 36 | 45 | 42 | 40 | 0.112575392 | 0.108790859 | 0.106067238 | 0.112786904 | 0.129996502 | 0.124622395 |
| NITMOv2_1740 | A0A0K2GB37 | Uncharacterized protein | 17.072 | 147 | 0.275 | 1.21 | 6.78E-03 | 1.28E-02 | no | no | 0 | 47 | 49 | 50 | 43 | 51 | 47 | 0.112377251 | 0.117959913 | 0.122764445 | 0.108762238 | 0.109775342 | 0.10056533 |
| NITMOv2_0024 | A0A0K2G6A9 | RND efflux system, membrane fusion protein | 44.711 | 415 | 0.31 | 1.24 | 3.70E-03 | 7.82E-03 | no | no | 0 | 88 | 95 | 97 | 91 | 93 | 82 | 0.112280289 | 0.121796485 | 0.124031738 | 0.104570247 | 0.105644104 | 0.096079017 |
| NITMOv2_3180 | A0A0K2GFF4 | TPR_REGION domain-containing protein | 50.048 | 457 | 0.711 | 1.64 | 5.43E-05 | 3.72E-04 | no | no | 0 | 92 | 97 | 98 | 75 | 71 | 64 | 0.112143277 | 0.119125355 | 0.113101577 | 0.072911908 | 0.075241671 | 0.073963923 |
| NITMOv2_2302 | A0A0K2GCP3 | Uncharacterized protein | 14.31 | 136 | 0.0725 | 1.05 | 6.40E-01 | 6.80E-01 | no | no | 0 | 24 | 22 | 27 | 29 | 24 | 25 | 0.111038749 | 0.104303126 | 0.114449393 | 0.105541616 | 0.089832291 | 0.065991053 |
| NITMOv2_0741 | A0A0K2G990 | Putative dihem cytochrome c | 36.78 | 333 | 0.191 | 1.14 | 3.90E-02 | 5.65E-02 | no | no | 0 | 93 | 88 | 90 | 85 | 88 | 84 | 0.109424115 | 0.11086558 | 0.106515871 | 0.098758321 | 0.09774985 | 0.104285643 |
| gcvH | A0A0K2GBJ0 | Glycine cleavage system H protein | 14.029 | 129 | -0.272 | -1.21 | 1.34E-02 | 2.27E-02 | no | no | 0 | 17 | 16 | 21 | 47 | 37 | 32 | 0.108502973 | 0.102300756 | 0.12271268 | 0.203898656 | 0.170796169 | 0.150687962 |
| NITMOv2_1283 | A0A0K2G9R5 | Uncharacterized protein | 31.345 | 283 | 0.573 | 1.49 | 7.11E-05 | 4.48E-04 | no | no | 0 | 28 | 30 | 35 | 25 | 30 | 27 | 0.108361745 | 0.110515556 | 0.11096961 | 0.069852688 | 0.071422723 | 0.069044263 |
| prcB | A0A0K2GG69 | Proteasome subunit beta | 30.391 | 275 | 0.577 | 1.49 | 4.83E-04 | 1.65E-03 | no | no | 0 | 33 | 40 | 39 | 47 | 50 | 41 | 0.10752281 | 0.112361491 | 0.100024111 | 0.092493287 | 0.098817229 | 0.093353065 |
| NITMOv2_0557 | A0A0K2G7R3 | Putative Disulfide oxidoreductase dsbA | 23.932 | 217 | 0.103 | 1.07 | 3.75E-01 | 4.18E-01 | no | no | 0 | 33 | 37 | 31 | 37 | 32 | 33 | 0.107408985 | 0.097603791 | 0.098624529 | 0.102292861 | 0.099544351 | 0.08520398 |
| rplA | A0A0K2G9K8 | 50S ribosomal protein L1 | 24.73 | 230 | -0.153 | -1.11 | 5.49E-02 | 7.61E-02 | no | no | 0 | 42 | 44 | 45 | 53 | 52 | 56 | 0.107219276 | 0.111747483 | 0.1072866 | 0.150029134 | 0.153032932 | 0.137349498 |
| NITMOv2_3177 | A0A0K2GF46 | Uncharacterized protein | 23.038 | 203 | 0.312 | 1.24 | 5.79E-03 | 1.12E-02 | no | no | 0 | 25 | 29 | 26 | 16 | 22 | 21 | 0.106664904 | 0.105296489 | 0.107725647 | 0.081642384 | 0.08007828 | 0.075826777 |
| dnaN | A0A0K2G663 | Beta sliding clamp | 40.866 | 375 | 0.195 | 1.14 | 3.13E-02 | 4.68E-02 | no | no | 0 | 73 | 79 | 73 | 63 | 66 | 69 | 0.106167445 | 0.103333228 | 0.099408678 | 0.106902421 | 0.101809619 | 0.099941382 |
| gspG | A0A0K2GCE2 | Type II secretion system core protein G | 19.615 | 178 | 0.571 | 1.49 | 2.84E-04 | 1.11E-03 | no | no | 0 | 16 | 17 | 18 | 12 | 14 | 14 | 0.105996707 | 0.110366943 | 0.099456609 | 0.075510617 | 0.080991845 | 0.067102292 |
| rpsP | A0A0K2GGK0 | 30S ribosomal protein S16 | 10.973 | 96 | -1.45 | -2.73 | 5.36E-03 | 1.05E-02 | yes | no | 0 | 11 | 11 | 22 | 38 | 32 | 34 | 0.105448659 | 0.101569422 | 0.112773729 | 0.187506803 | 0.180848486 | 0.154685186 |
| NITMOv2_0033 | A0A0K2G7B8 | Appr-1-p processing enzyme family protein | 17.698 | 161 | -0.134 | -1.10 | 1.87E-01 | 2.24E-01 | no | no | 0 | 45 | 39 | 40 | 45 | 39 | 39 | 0.105115615 | 0.095849762 | 0.098770239 | 0.114548991 | 0.105860065 | 0.097485148 |
| erpA | A0A0K2GHP6 | Iron-sulfur cluster insertion protein ErpA 2 | 11.332 | 106 | -0.866 | -1.82 | 1.38E-04 | 6.94E-04 | no | no | 0 | 32 | 35 | 35 | 41 | 45 | 43 | 0.104748844 | 0.125748427 | 0.125034452 | 0.271953719 | 0.262230305 | 0.220018127 |
| forA | A0A0K2G835 | Pyruvate:ferredoxin oxidoreductase alpha subunit | 48.41 | 444 | -0.972 | -1.96 | 4.63E-06 | 7.17E-05 | no | no | 0 | 85 | 87 | 98 | 142 | 136 | 150 | 0.102166695 | 0.098896337 | 0.108699603 | 0.21079893 | 0.214609978 | 0.212627848 |
| NITMOv2_4116 | A0A0K2GHR1 | Uncharacterized protein | 14.934 | 138 | 1.4 | 2.64 | 2.18E-05 | 1.99E-04 | yes | no | 0 | 26 | 25 | 26 | 17 | 15 | 18 | 0.101163345 | 0.102838502 | 0.09883159 | 0.056320155 | 0.059706413 | 0.05874822 |
| ppdK | A0A0K2GF53 | Pyruvate, phosphate dikinase | 102.92 | 935 | 0.0594 | 1.04 | 2.92E-01 | 3.34E-01 | no | no | 0 | 213 | 220 | 249 | 232 | 239 | 244 | 0.100965205 | 0.102572562 | 0.105877432 | 0.099285467 | 0.098338695 | 0.095027116 |
| rpsQ | A0A0K2G9L5 | 30S ribosomal protein S17 | 10.184 | 87 | -0.709 | -1.63 | 1.25E-04 | 6.48E-04 | no | no | 0 | 11 | 14 | 13 | 14 | 12 | 14 | 0.100876674 | 0.102298801 | 0.104903476 | 0.139301133 | 0.146398714 | 0.143927026 |
| NITMOv2_4640 | A0A0K2GJ93 | Uncharacterized protein | 22.157 | 199 | 0.824 | 1.77 | 4.81E-05 | 3.41E-04 | no | no | 0 | 35 | 33 | 36 | 23 | 23 | 24 | 0.100549953 | 0.091624058 | 0.090537628 | 0.058052627 | 0.05790725 | 0.059402736 |
| NITMOv2_0172 | A0A0K2G7M4 | Signal transduction response regulator, receiver domain | 13.697 | 123 | -0.0335 | -1.02 | 5.95E-01 | 6.36E-01 | no | no | 0 | 17 | 19 | 23 | 31 | 32 | 31 | 0.10042348 | 0.10645802 | 0.107560765 | 0.111241006 | 0.104980682 | 0.092311953 |
| rpsU | A0A0K2GJ41 | 30S ribosomal protein S21 | 7.8402 | 64 | 0.875 | 1.83 | 6.54E-02 | 8.86E-02 | no | no | 0 | 4 | 4 | 3 | 4 | 3 | 5 | 0.100170535 | 0.097511886 | 0.09476705 | 0.090020442 | 0.104927857 | 0.101074198 |
| NITMOv2_3940 | A0A0K2GH89 | Uncharacterized protein | 34.082 | 317 | 7.62 | 196.72 | 8.15E-06 | 1.02E-04 | yes | yes | 3 | 46 | 49 | 55 | 0 | 0 | 0 | 0.099316845 | 0.104958198 | 0.10342337 | 0 | 0 | 0 |
| ttgC | A0A0K2G6B8 | RND efflux system, outer membrane factor lipoprotein | 52.965 | 478 | 0.49 | 1.40 | 4.23E-04 | 1.50E-03 | no | no | 0 | 86 | 80 | 85 | 78 | 79 | 74 | 0.09919248 | 0.088565751 | 0.086952397 | 0.074422269 | 0.074432203 | 0.07035869 |
| rplU | A0A0K2G982 | 50S ribosomal protein L21 | 11.445 | 104 | -0.3 | -1.23 | 7.69E-03 | 1.42E-02 | no | no | 0 | 10 | 14 | 13 | 20 | 18 | 17 | 0.099129244 | 0.113262948 | 0.103001578 | 0.137379125 | 0.147320047 | 0.15225952 |
| fabZ | A0A0K2GGH0 | 3-hydroxyacyl-[acyl-carrier-protein] dehydratase FabZ | 16.19 | 144 | 0.447 | 1.36 | 5.92E-04 | 1.94E-03 | no | no | 0 | 41 | 39 | 40 | 37 | 33 | 36 | 0.09912292 | 0.103002759 | 0.102922971 | 0.087674941 | 0.081288598 | 0.078649827 |
| NITMOv2_1342 | A0A0K2G9Z7 | Putative NHL repeat protein | 107.26 | 989 | -0.272 | -1.21 | 3.85E-03 | 8.05E-03 | no | no | 0 | 109 | 115 | 118 | 143 | 142 | 151 | 0.09867605 | 0.096678869 | 0.102092808 | 0.124350046 | 0.122304227 | 0.113216912 |
| porA | A0A0K2G8Q9 | Pyruvate:ferredoxin oxidoreductase alpha subunit | 44.803 | 403 | -0.356 | -1.28 | 1.65E-03 | 4.14E-03 | no | no | 0 | 60 | 63 | 68 | 77 | 74 | 73 | 0.098058442 | 0.096227162 | 0.098563177 | 0.138615548 | 0.138359968 | 0.130122849 |
| sucD | A0A0K2G850 | Succinate--CoA ligase [ADP-forming] subunit alpha | 29.984 | 290 | 0.0911 | 1.07 | 2.36E-01 | 2.77E-01 | no | no | 0 | 39 | 37 | 34 | 37 | 35 | 35 | 0.097636867 | 0.097263545 | 0.096837665 | 0.096375802 | 0.101373034 | 0.093854741 |
| NITMOv2_0201 | A0A0K2G6P0 | Uncharacterized protein | 57.985 | 549 | 0.532 | 1.45 | 4.66E-04 | 1.62E-03 | no | no | 0 | 100 | 95 | 95 | 80 | 86 | 77 | 0.097579954 | 0.097756316 | 0.097077319 | 0.073970642 | 0.070336699 | 0.066129508 |
| dksA | A0A0K2G9N8 | DnaK suppressor protein (Modular protein) | 20.789 | 183 | -0.104 | -1.07 | 1.33E-01 | 1.66E-01 | no | no | 0 | 24 | 22 | 26 | 29 | 28 | 25 | 0.097383922 | 0.061318662 | 0.095068056 | 0.101938963 | 0.1019075 | 0.10282377 |
| accB | A0A0K2GC14 | Biotin carboxyl carrier protein of acetyl-CoA carboxylase | 20.487 | 189 | 0.48 | 1.39 | 5.61E-04 | 1.85E-03 | no | no | 0 | 25 | 25 | 23 | 19 | 17 | 21 | 0.096861168 | 0.098374234 | 0.094914677 | 0.069038278 | 0.071817358 | 0.074564496 |
| rplI | A0A0K2G6G8 | 50S ribosomal protein L9 | 17.07 | 158 | -0.346 | -1.27 | 3.85E-03 | 8.05E-03 | no | no | 0 | 21 | 26 | 30 | 34 | 33 | 32 | 0.096614546 | 0.090695225 | 0.092713691 | 0.128198503 | 0.132226034 | 0.116363624 |
| rpoC | A0A0K2G9J9 | DNA-directed RNA polymerase subunit beta | 154.09 | 1395 | -0.39 | -1.31 | 4.86E-04 | 1.66E-03 | no | no | 0 | 253 | 263 | 269 | 352 | 350 | 333 | 0.096163461 | 0.098867005 | 0.100913708 | 0.137903309 | 0.13347364 | 0.12999698 |
| ctpA | A0A0K2GG72 | Carboxy-terminal-processing protease | 47.922 | 442 | -0.149 | -1.11 | 8.81E-02 | 1.16E-01 | no | no | 0 | 63 | 57 | 58 | 68 | 71 | 65 | 0.09604542 | 0.102421993 | 0.088825536 | 0.093757844 | 0.097659736 | 0.095908196 |
| hflC | A0A0K2GHW4 | Protein HflC | 33.019 | 286 | 0.583 | 1.50 | 1.30E-04 | 6.63E-04 | no | no | 0 | 28 | 39 | 32 | 30 | 30 | 26 | 0.095326633 | 0.100578014 | 0.096632521 | 0.059123206 | 0.064965624 | 0.063779364 |
| NITMOv2_1049 | A0A0K2G967 | Uncharacterized protein | 9.0634 | 77 | 1.03 | 2.04 | 1.56E-04 | 7.54E-04 | yes | no | 0 | 4 | 3 | 3 | 2 | 5 | 4 | 0.095273937 | 0.092214601 | 0.087452795 | 0.026646015 | 0.043178353 | 0.033677015 |
| NITMOv2_2220 | A0A0K2GCF0 | Uncharacterized protein | 10.3 | 92 | 0.655 | 1.57 | 2.20E-03 | 5.17E-03 | no | no | 0 | 7 | 7 | 8 | 5 | 6 | 7 | 0.095058933 | 0.098389878 | 0.08195608 | 0.057161219 | 0.05518054 | 0.05135974 |
| frr | A0A0K2GB51 | Ribosome-recycling factor | 21.179 | 187 | 0.437 | 1.35 | 1.18E-03 | 3.18E-03 | no | no | 0 | 29 | 34 | 30 | 25 | 23 | 21 | 0.094917705 | 0.091213416 | 0.091739735 | 0.074691765 | 0.079346496 | 0.074188688 |
| NITMOv2_2388 | A0A0K2GCX5 | Uncharacterized protein | 25.36 | 224 | -0.281 | -1.22 | 7.52E-03 | 1.40E-02 | no | no | 0 | 33 | 32 | 36 | 28 | 35 | 32 | 0.094685839 | 0.095597511 | 0.095434248 | 0.123294274 | 0.125644641 | 0.115655164 |
| rplX | A0A0K2G9M7 | 50S ribosomal protein L24 | 12.335 | 113 | -0.321 | -1.25 | 2.99E-02 | 4.51E-02 | no | no | 0 | 15 | 15 | 15 | 13 | 13 | 12 | 0.09452564 | 0.095145804 | 0.098528667 | 0.136021282 | 0.111232695 | 0.107871097 |
| rplY | A0A0K2G7Z5 | 50S ribosomal protein L25 | 27.06 | 260 | -0.607 | -1.52 | 2.56E-04 | 1.04E-03 | no | no | 0 | 45 | 48 | 61 | 63 | 70 | 71 | 0.09404926 | 0.099995293 | 0.101437114 | 0.155419048 | 0.158599772 | 0.135056894 |
| NITMOv2_3472 | A0A0K2GG75 | Putative periplasmic serine endoprotease DegP-like | 54.779 | 513 | 0.397 | 1.32 | 1.34E-03 | 3.54E-03 | no | no | 0 | 71 | 71 | 69 | 58 | 59 | 64 | 0.093836364 | 0.096938942 | 0.091578687 | 0.07313254 | 0.069364095 | 0.07103838 |
| NITMOv2_1129 | A0A0K2G9E6 | Uncharacterized protein | 23.143 | 211 | 0.913 | 1.88 | 9.27E-06 | 1.12E-04 | no | no | 0 | 28 | 30 | 26 | 20 | 22 | 18 | 0.093608714 | 0.102476746 | 0.090023809 | 0.064865537 | 0.063244145 | 0.057065179 |
| NITMOv2_2068 | A0A0K2GC11 | Uncharacterized protein | 33.138 | 297 | 0.568 | 1.48 | 3.76E-04 | 1.36E-03 | no | no | 0 | 51 | 51 | 45 | 39 | 40 | 32 | 0.093290424 | 0.087726868 | 0.095602965 | 0.071919809 | 0.062408265 | 0.061497548 |
| NITMOv2_1528 | A0A0K2GAI7 | Putative Anion-selective porin | 64.722 | 581 | 1.02 | 2.03 | 2.87E-06 | 5.32E-05 | yes | no | 0 | 83 | 80 | 87 | 57 | 56 | 53 | 0.091840205 | 0.093131702 | 0.096766727 | 0.049017118 | 0.04769335 | 0.045098681 |
| moeA | A0A0K2GGH8 | Molybdopterin molybdenumtransferase | 46.612 | 432 | 0.351 | 1.28 | 1.17E-03 | 3.17E-03 | no | no | 0 | 98 | 103 | 110 | 100 | 108 | 104 | 0.091728487 | 0.094246302 | 0.091983224 | 0.076603407 | 0.075505797 | 0.076150439 |
| rpsF | A0A0K2G8Z5 | 30S ribosomal protein S6 | 13.287 | 115 | -0.26 | -1.20 | 1.10E-02 | 1.93E-02 | no | no | 0 | 16 | 16 | 20 | 21 | 21 | 19 | 0.091680006 | 0.089175848 | 0.119482137 | 0.287279435 | 0.118987117 | 0.356639386 |
| devR.2 | A0A0K2G7Q0 | Transcriptional regulatory protein DevR (DosR) | 25.111 | 224 | -0.0199 | -1.01 | 7.69E-01 | 7.97E-01 | no | no | 0 | 41 | 37 | 39 | 37 | 36 | 40 | 0.090701951 | 0.082900844 | 0.089655699 | 0.100473024 | 0.093209993 | 0.092236432 |
| NITMOv2_1899 | A0A0K2GBS1 | Uncharacterized protein | 21.522 | 199 | 0.445 | 1.36 | 9.76E-04 | 2.76E-03 | no | no | 0 | 28 | 34 | 38 | 28 | 28 | 23 | 0.090354151 | 0.095681595 | 0.101943264 | 0.073827009 | 0.062787363 | 0.063319045 |
| infC | A0A0K2G714 | Translation initiation factor IF-3 | 18.818 | 163 | 0.515 | 1.43 | 2.40E-04 | 1.00E-03 | no | no | 0 | 29 | 26 | 24 | 22 | 22 | 22 | 0.090202384 | 0.093972541 | 0.094410444 | 0.071277165 | 0.075154665 | 0.073802092 |
| NITMOv2_3693 | A0A0K2GGJ8 | Putative Tol-Pal system protein YbgF | 28.407 | 259 | 0.363 | 1.29 | 3.34E-03 | 7.19E-03 | no | no | 0 | 37 | 38 | 37 | 24 | 24 | 24 | 0.089957871 | 0.09069718 | 0.087866918 | 0.073930661 | 0.081026026 | 0.074566294 |
| NITMOv2_4591 | A0A0K2GJ39 | Uncharacterized protein | 62.189 | 566 | -0.0868 | -1.06 | 1.87E-01 | 2.24E-01 | no | no | 0 | 133 | 135 | 150 | 178 | 169 | 151 | 0.089900958 | 0.091571261 | 0.091258509 | 0.102623067 | 0.100860319 | 0.094921027 |
| NITMOv2_1745 | A0A0K2GB39 | Precorrin-2 dehydrogenase | 23.961 | 217 | 0.382 | 1.30 | 6.35E-04 | 2.04E-03 | no | no | 0 | 39 | 39 | 43 | 38 | 35 | 33 | 0.089365557 | 0.086985757 | 0.0919583 | 0.079741107 | 0.07565495 | 0.076686279 |
| rplN | A0A0K2G9V3 | 50S ribosomal protein L14 | 13.607 | 122 | 0.0072 | 1.01 | 9.29E-01 | 9.41E-01 | no | no | 0 | 24 | 22 | 23 | 24 | 22 | 26 | 0.088941874 | 0.089850475 | 0.091475156 | 0.078275169 | 0.088701211 | 0.09578952 |
| NITMOv2_3374 | A0A0K2GFN6 | Fibronectin type-III domain-containing protein | 83.523 | 779 | 0.534 | 1.45 | 4.40E-04 | 1.54E-03 | no | no | 0 | 72 | 80 | 82 | 64 | 66 | 62 | 0.088372747 | 0.088634192 | 0.08047214 | 0.062145408 | 0.060183393 | 0.05744638 |
| rpoD.1 | A0A0K2G9K6 | RNA polymerase sigma factor SigA | 68.868 | 605 | 0.00844 | 1.01 | 8.98E-01 | 9.12E-01 | no | no | 0 | 78 | 78 | 78 | 87 | 84 | 80 | 0.088227303 | 0.092187225 | 0.090926827 | 0.092790917 | 0.102400017 | 0.089995973 |
| NITMOv2_1336 | A0A0K2GA63 | Uncharacterized protein | 16.686 | 153 | 0.996 | 1.99 | 3.20E-03 | 6.97E-03 | no | no | 0 | 9 | 11 | 10 | 11 | 9 | 8 | 0.088020731 | 0.076885913 | 0.079904638 | 0.059783619 | 0.065469016 | 0.064750349 |
| NITMOv2_2962 | A0A0K2GFI9 | FolC bifunctional protein | 50.627 | 467 | 1.88 | 3.68 | 1.01E-07 | 8.38E-06 | yes | no | 0 | 68 | 64 | 68 | 26 | 25 | 29 | 0.088016516 | 0.090040153 | 0.091578687 | 0.025550264 | 0.02640015 | 0.025696952 |
| NITMOv2_2968 | A0A0K2GEJ5 | Uncharacterized protein | 87.485 | 795 | 1.5 | 2.83 | 2.94E-06 | 5.41E-05 | yes | no | 0 | 141 | 135 | 151 | 73 | 75 | 85 | 0.087995437 | 0.084230543 | 0.095012456 | 0.030734354 | 0.030334071 | 0.036694262 |
| NITMOv2_3642 | A0A0K2GGR8 | Putative dihem cytochrome c | 47.183 | 437 | -1.38 | -2.60 | 1.11E-06 | 3.12E-05 | yes | no | 0 | 50 | 48 | 49 | 95 | 95 | 89 | 0.086897233 | 0.083297798 | 0.081576468 | 0.234091056 | 0.247516944 | 0.236327087 |
| NITMOv2_0572 | A0A0K2G7S6 | Uncharacterized protein | 11.259 | 110 | 0.826 | 1.77 | 6.02E-03 | 1.16E-02 | no | no | 0 | 7 | 9 | 9 | 4 | 6 | 8 | 0.086756005 | 0.102236227 | 0.07892493 | 0.070059993 | 0.068522 | 0.062657336 |
| NITMOv2_2977 | A0A0K2GFK5 | AIRC domain-containing protein | 26.752 | 252 | 1.19 | 2.28 | 4.67E-06 | 7.17E-05 | yes | no | 0 | 30 | 28 | 32 | 16 | 16 | 18 | 0.086728603 | 0.090591586 | 0.095363311 | 0.049778222 | 0.047816091 | 0.044918869 |
| galU | A0A0K2GG59 | UTP--glucose-1-phosphate uridylyltransferase | 32.288 | 297 | 0.406 | 1.33 | 3.11E-04 | 1.19E-03 | no | no | 0 | 41 | 44 | 47 | 47 | 44 | 44 | 0.08650306 | 0.084981431 | 0.086141406 | 0.065277185 | 0.065683424 | 0.067015982 |
| rplO | A0A0K2G9M5 | 50S ribosomal protein L15 | 15.944 | 149 | -0.476 | -1.39 | 1.03E-03 | 2.87E-03 | no | no | 0 | 33 | 35 | 33 | 33 | 33 | 35 | 0.085482847 | 0.090126192 | 0.095455338 | 0.122826359 | 0.135609952 | 0.118444051 |
| NITMOv2_0915 | A0A0K2G903 | ABC transporter, periplasmic binding component, putative Fe3+ transport system | 37.681 | 349 | 0.712 | 1.64 | 1.28E-05 | 1.39E-04 | no | no | 0 | 71 | 75 | 71 | 54 | 61 | 49 | 0.085398532 | 0.090067529 | 0.089661451 | 0.050866569 | 0.053858357 | 0.051134975 |
| NITMOv2_4495 | A0A0K2GJ17 | Uncharacterized protein | 25.767 | 235 | -0.259 | -1.20 | 3.76E-02 | 5.47E-02 | no | no | 0 | 84 | 84 | 75 | 82 | 82 | 75 | 0.085109753 | 0.081876194 | 0.085618001 | 0.110373289 | 0.104935625 | 0.091884 |
| NITMOv2_1739 | A0A0K2GBB0 | 2Fe-2S Ferredoxin | 11.47 | 108 | 0.195 | 1.14 | 2.43E-01 | 2.84E-01 | no | no | 0 | 8 | 10 | 6 | 9 | 10 | 6 | 0.085071811 | 0.105711042 | 0.082937705 | 0.091558937 | 0.085861081 | 0.070009854 |

| | | | | | | | | | | | | | | | | | | | | | | | |
|---|---|---|---|---|---|---|---|---|---|---|---|---|---|---|---|---|---|---|---|---|---|---|---|
| pdxJ | A0A0K2GIY9 | Pyridoxine 5-phosphate synthase | 25.809 | 237 | 0.244 | 1.18 | 6.87E-02 | 9.28E-02 | no | no | 0 | 23 | 31 | 30 | 29 | 33 | 26 | 0.085065488 | 0.072190904 | 0.079254695 | 0.08187338 | 0.068060557 | 0.062842543 |
| NITMOv2_3649 | A0A0K2GGH3 | Uncharacterized protein | 23.957 | 217 | 0.393 | 1.31 | 1.61E-03 | 4.08E-03 | no | no | 0 | 29 | 26 | 29 | 28 | 26 | 24 | 0.084783032 | 0.083170695 | 0.084609535 | 0.073874393 | 0.066489785 | 0.071930248 |
| pnp | A0A0K2GJQ2 | Polyribonucleotide nucleotidyltransferase | 75.523 | 705 | -0.207 | -1.15 | 1.22E-02 | 2.11E-02 | no | no | 0 | 85 | 86 | 95 | 137 | 146 | 137 | 0.084698717 | 0.084443686 | 0.092418436 | 0.112804673 | 0.113424939 | 0.105995657 |
| NITMOv2_1179 | A0A0K2G9J4 | Uncharacterized protein | 30.555 | 274 | -0.416 | -1.33 | 2.75E-03 | 6.17E-03 | no | no | 0 | 51 | 49 | 59 | 69 | 67 | 65 | 0.084285573 | 0.079158134 | 0.085581573 | 0.117599327 | 0.11678089 | 0.106391243 |
| porD | A0A0K2G8I3 | Pyruvate:ferredoxin oxidoreductase delta subunit (PorD) | 23.347 | 210 | -0.527 | -1.44 | 1.18E-03 | 3.18E-03 | no | no | 0 | 28 | 27 | 29 | 35 | 35 | 33 | 0.083948313 | 0.083796435 | 0.079149247 | 0.12106131 | 0.111813771 | 0.102314902 |
| NITMOv2_4177 | A0A0K2GIW3 | Putative Zn-dependent peptidase, M16 family | 50.41 | 461 | 0.317 | 1.25 | 1.01E-02 | 1.80E-02 | no | no | 0 | 69 | 67 | 69 | 58 | 56 | 59 | 0.08376282 | 0.087415953 | 0.084883699 | 0.065952405 | 0.071520604 | 0.066679734 |
| NITMOv2_1527 | A0A0K2GBJ4 | Uncharacterized protein | 24.8 | 220 | 0.93 | 1.91 | 1.80E-04 | 8.10E-04 | no | no | 0 | 31 | 32 | 30 | 20 | 19 | 19 | 0.083739633 | 0.088688944 | 0.090627738 | 0.057589154 | 0.054945934 | 0.051663622 |
| NITMOv2_2087 | A0A0K2GCC5 | Putative Ferredoxin-NADP(+) reductase | 49.133 | 450 | -0.00388 | -1.00 | 9.53E-01 | 9.59E-01 | no | no | 0 | 67 | 63 | 69 | 77 | 73 | 73 | 0.083301195 | 0.079070139 | 0.081026221 | 0.081027875 | 0.086681425 | 0.083988449 |
| yrbD | A0A0K2G8H8 | Putative ABC transporter, periplasmic binding component, ATP-dependent toluene efflux trans | 15.562 | 147 | 0.118 | 1.09 | 1.56E-01 | 1.92E-01 | no | no | 0 | 20 | 19 | 21 | 22 | 20 | 22 | 0.083041926 | 0.086823455 | 0.086149075 | 0.069769767 | 0.075395485 | 0.071730656 |
| NITMOv2_1476 | A0A0K2GAD8 | Putative Membrane-fusion protein of multidrug efflux transporter | 44.624 | 412 | 0.64 | 1.56 | 5.27E-05 | 3.64E-04 | no | no | 0 | 53 | 62 | 59 | 58 | 51 | 46 | 0.082824814 | 0.077359129 | 0.090947916 | 0.060128633 | 0.059317993 | 0.051320181 |
| tyrS | A0A0K2GCB0 | Tyrosine--tRNA ligase | 45.564 | 403 | 0.203 | 1.15 | 1.16E-02 | 2.03E-02 | no | no | 0 | 60 | 67 | 72 | 59 | 64 | 60 | 0.082639321 | 0.082187109 | 0.085794386 | 0.064197721 | 0.063427479 | 0.064527382 |
| NITMOv2_2289 | A0A0K2GCM1 | Uncharacterized protein | 22.729 | 205 | 0.324 | 1.25 | 7.69E-03 | 1.42E-02 | no | no | 0 | 39 | 44 | 48 | 37 | 38 | 35 | 0.082472799 | 0.083207848 | 0.085685104 | 0.070052589 | 0.066076506 | 0.060522966 |
| liaR | A0A0K2G7I0 | Transcriptional regulatory protein LiaR | 25.011 | 223 | 0.236 | 1.18 | 1.36E-02 | 2.30E-02 | no | no | 0 | 36 | 37 | 43 | 38 | 38 | 35 | 0.082124999 | 0.081647408 | 0.076817888 | 0.068309751 | 0.067010268 | 0.06368766 |
| NITMOv2_4014 | A0A0K2GHF9 | Uncharacterized protein | 22.066 | 212 | 0.443 | 1.36 | 3.64E-03 | 7.73E-03 | no | no | 0 | 24 | 23 | 18 | 16 | 19 | 18 | 0.081543225 | 0.086543827 | 0.071117946 | 0.066808276 | 0.060938482 | 0.056813442 |
| NITMOv2_3171 | A0A0K2GF33 | Uncharacterized protein | 7.3956 | 64 | 0.189 | 1.14 | 7.94E-02 | 1.06E-01 | no | no | 0 | 13 | 15 | 11 | 11 | 7 | 11 | 0.081039443 | 0.087435508 | 0.078857827 | 0.071197205 | 0.059858674 | 0.06122783 |
| rpoZ | A0A0K2GC18 | DNA-directed RNA polymerase subunit omega | 13.374 | 121 | -0.159 | -1.12 | 4.42E-02 | 6.31E-02 | no | no | 0 | 23 | 26 | 24 | 24 | 30 | 31 | 0.08094248 | 0.083575471 | 0.086175916 | 0.096408378 | 0.101551707 | 0.104122014 |
| eno | A0A0K2GFS9 | Enolase | 45.958 | 428 | 0.135 | 1.10 | 1.01E-01 | 1.30E-01 | no | no | 0 | 59 | 64 | 70 | 68 | 64 | 60 | 0.08026796 | 0.07584562 | 0.080493229 | 0.084540203 | 0.078137732 | 0.074893552 |
| NITMOv2_0061 | A0A0K2G6D8 | Putative Orotate phosphoribosyltransferase (Modular protein) | 27.422 | 250 | 2.61 | 6.11 | 1.28E-08 | 3.40E-06 | yes | no | 0 | 30 | 32 | 31 | 6 | 3 | 4 | 0.079892757 | 0.081074464 | 0.078372767 | 0.006726583 | 0.007338658 | 0.008497562 |
| bkdAB | A0A0K2G6B2 | 2-oxoisovalerate dehydrogenase, beta subunit | 36.056 | 331 | 3.31 | 9.92 | 2.34E-08 | 4.25E-06 | yes | no | 0 | 57 | 58 | 68 | 5 | 5 | 5 | 0.079538634 | 0.082249683 | 0.087029086 | 0.005018987 | 0.005463209 | 0.004869133 |
| NITMOv2_2404 | A0A0K2GD74 | Uncharacterized protein | 12.12 | 109 | -0.207 | -1.15 | 1.64E-01 | 2.00E-01 | no | no | 0 | 16 | 13 | 21 | 29 | 26 | 26 | 0.079378435 | 0.0701944 | 0.081309972 | 0.082638926 | 0.070339807 | 0.07017708 |
| NITMOv2_3617 | A0A0K2GGP7 | Putative Nitrite oxidoreductase, membrane subunit | 30.847 | 277 | 0.681 | 1.60 | 1.97E-04 | 8.67E-04 | no | no | 0 | 48 | 52 | 44 | 36 | 34 | 37 | 0.079017988 | 0.081248498 | 0.073978462 | 0.05801857 | 0.052512869 | 0.051976495 |
| NITMOv2_1415 | A0A0K2GB78 | Uncharacterized protein | 17.368 | 152 | 0.485 | 1.40 | 4.55E-04 | 1.59E-03 | no | no | 0 | 24 | 29 | 26 | 23 | 23 | 23 | 0.078836711 | 0.089369437 | 0.077630796 | 0.066960792 | 0.064042737 | 0.063480876 |
| fumC | A0A0K2GC26 | Fumarate hydratase class II | 51.575 | 485 | 0.446 | 1.36 | 5.06E-04 | 1.71E-03 | no | no | 0 | 44 | 48 | 50 | 43 | 47 | 52 | 0.078697591 | 0.076872225 | 0.077688313 | 0.060038307 | 0.071651114 | 0.068251292 |
| NITMOv2_1098 | A0A0K2G9J6 | Putative Phosphate-binding protein PstS | 29.014 | 273 | 0.507 | 1.42 | 2.71E-03 | 6.11E-03 | no | no | 0 | 29 | 32 | 30 | 26 | 30 | 24 | 0.078577442 | 0.084672472 | 0.080481726 | 0.073795914 | 0.069430903 | 0.056108578 |
| nusA | A0A0K2GIS2 | Transcription termination/antitermination protein NusA | 43.11 | 389 | -0.2 | -1.15 | 1.97E-02 | 3.14E-02 | no | no | 0 | 53 | 56 | 64 | 64 | 63 | 62 | 0.078457293 | 0.076322747 | 0.078543401 | 0.092534748 | 0.092291768 | 0.091407498 |
| pal.1 | A0A0K2GHI1 | Peptidoglycan-associated protein | 24.42 | 233 | 0.246 | 1.19 | 1.31E-02 | 2.23E-02 | no | no | 0 | 27 | 32 | 29 | 30 | 33 | 26 | 0.078368762 | 0.08185664 | 0.066503159 | 0.073668569 | 0.074457062 | 0.053598401 |
| leuB | A0A0K2G8M7 | 3-isopropylmalate dehydrogenase | 38.553 | 357 | -0.0727 | -1.05 | 3.59E-01 | 4.02E-01 | no | no | 0 | 41 | 44 | 43 | 46 | 45 | 46 | 0.076819473 | 0.077789326 | 0.072254867 | 0.074885742 | 0.073649148 | 0.066875729 |
| NITMOv2_0063 | A0A0K2G6F0 | Putative dioxygenase | 41.331 | 369 | 2.1 | 4.29 | 3.52E-07 | 1.46E-05 | yes | no | 0 | 67 | 70 | 71 | 22 | 20 | 19 | 0.076545449 | 0.080251224 | 0.077433321 | 0.017563122 | 0.01527455 | 0.013640728 |
| iscX | A0A0K2GIC7 | Iron-sulfur cluster assembly protein IscX | 8.2662 | 70 | 0.497 | 1.41 | 1.99E-02 | 3.17E-02 | no | no | 0 | 9 | 11 | 10 | 6 | 6 | 4 | 0.075837202 | 0.078221478 | 0.060043992 | 0.044865108 | 0.044189799 | 0.034633615 |
| NITMOv2_3723 | A0A0K2GGM7 | DUF5069 domain-containing protein | 16.468 | 146 | 0.0288 | 1.02 | 8.22E-01 | 8.43E-01 | no | no | 0 | 36 | 39 | 47 | 35 | 35 | 35 | 0.07570019 | 0.074150254 | 0.075075121 | 0.072284072 | 0.068341773 | 0.060415079 |
| NITMOv2_1069 | A0A0K2G986 | TPR_REGION domain-containing protein | 46.634 | 420 | 0.664 | 1.58 | 1.88E-04 | 8.31E-04 | no | no | 0 | 87 | 82 | 89 | 65 | 64 | 57 | 0.075603228 | 0.071666846 | 0.075186321 | 0.049699742 | 0.048670616 | 0.046591122 |
| arsC | A0A0K2GC24 | Arsenate reductase | 13.032 | 116 | 0.223 | 1.17 | 7.85E-03 | 1.45E-02 | no | no | 0 | 22 | 25 | 27 | 21 | 25 | 21 | 0.075485186 | 0.077353263 | 0.07751001 | 0.061961795 | 0.063713357 | 0.062123294 |
| NITMOv2_0255 | A0A0K2G747 | Putative Nitrite oxidoreductase, alpha subunit | 131.78 | 1145 | 1.3 | 2.46 | 1.23E-05 | 1.38E-04 | yes | no | 0 | 91 | 85 | 91 | 53 | 52 | 60 | 0.075312341 | 0.070896403 | 0.095039298 | 0.065588141 | 0.078791832 | 0.091745545 |
| NITMOv2_0348 | A0A0K2G732 | Probable transcriptional regulatory protein NITMOv2_0348 | 27.467 | 252 | 0.126 | 1.09 | 1.19E-01 | 1.50E-01 | no | no | 0 | 43 | 42 | 44 | 43 | 43 | 41 | 0.075230133 | 0.07664735 | 0.075360789 | 0.070673021 | 0.07370508 | 0.068170376 |
| atpH | A0A0K2G789 | ATP synthase subunit delta | 21.689 | 198 | -0.136 | -1.10 | 1.07E-01 | 1.37E-01 | no | no | 0 | 22 | 21 | 20 | 23 | 25 | 25 | 0.073904279 | 0.078467863 | 0.079822197 | 0.085615224 | 0.08284694 | 0.075736871 |
| NITMOv2_2235 | A0A0K2GCG4 | Uncharacterized protein | 15.381 | 141 | 0.273 | 1.21 | 5.05E-03 | 1.00E-02 | no | no | 0 | 19 | 17 | 17 | 17 | 14 | 17 | 0.073421575 | 0.07771893 | 0.076651089 | 0.06246821 | 0.064726357 | 0.061209848 |
| NITMOv2_3573 | A0A0K2GG87 | Uncharacterized protein | 25.977 | 243 | 0.508 | 1.42 | 3.54E-04 | 1.31E-03 | no | no | 0 | 23 | 24 | 23 | 18 | 17 | 20 | 0.073215003 | 0.059062085 | 0.06913169 | 0.050497864 | 0.049355789 | 0.047297784 |
| NITMOv2_3620 | A0A0K2GHC3 | TPR_REGION domain-containing protein | 30.798 | 270 | -0.435 | -1.35 | 4.59E-04 | 1.60E-03 | no | no | 0 | 42 | 43 | 41 | 56 | 49 | 49 | 0.072822938 | 0.074865945 | 0.078815648 | 0.109681781 | 0.104278418 | 0.093950042 |
| NITMOv2_2572 | A0A0K2GDF8 | PepSY domain-containing protein | 9.4488 | 92 | 0.521 | 1.43 | 3.49E-04 | 1.30E-03 | no | no | 0 | 9 | 9 | 9 | 8 | 7 | 9 | 0.072797643 | 0.067875249 | 0.07112945 | 0.047024035 | 0.041654192 | 0.043091978 |
| metK | A0A0K2G880 | S-adenosylmethionine synthase | 41.966 | 382 | -1.19 | -2.28 | 1.81E-06 | 4.01E-05 | yes | no | 0 | 46 | 50 | 51 | 81 | 91 | 82 | 0.072489893 | 0.068858835 | 0.075985808 | 0.160261086 | 0.167067334 | 0.161361611 |
| NITMOv2_2254 | A0A0K2GCI7 | Aklaviketone reductase | 20.861 | 200 | 0.389 | 1.31 | 1.10E-03 | 3.01E-03 | no | no | 0 | 55 | 59 | 61 | 51 | 47 | 42 | 0.072420333 | 0.076882002 | 0.074739605 | 0.055168135 | 0.055283083 | 0.052895335 |
| lpd | A0A0K2GBB3 | Dihydrolipoyl dehydrogenase | 49.634 | 477 | 0.0848 | 1.06 | 2.63E-01 | 3.04E-01 | no | no | 0 | 67 | 64 | 65 | 72 | 67 | 62 | 0.071874393 | 0.067116539 | 0.067569143 | 0.071752485 | 0.072631486 | 0.069741934 |
| NITMOv2_1481 | A0A0K2GAE2 | Uncharacterized protein | 16.865 | 150 | 5.24 | 37.79 | 2.47E-09 | 1.79E-06 | yes | no | 0 | 29 | 39 | 34 | 1 | 0 | 0 | 0.071804833 | 0.083722129 | 0.076637668 | 0.006099339 | 0.004848884 | 0.004343182 |
| fusA | A0A0K2G9I2 | Elongation factor G | 77.034 | 693 | -0.808 | -1.75 | 1.27E-05 | 1.39E-04 | no | no | 0 | 114 | 109 | 118 | 175 | 178 | 169 | 0.071781646 | 0.071565163 | 0.072764852 | 0.140265098 | 0.135237069 | 0.130825914 |
| NITMOv2_4176 | A0A0K2GHZ8 | Putative Zn-dependent peptidase, M16 family | 51.581 | 459 | 0.414 | 1.33 | 1.64E-03 | 4.13E-03 | no | no | 0 | 56 | 67 | 69 | 55 | 54 | 49 | 0.071722626 | 0.074691768 | 0.068608284 | 0.049599051 | 0.050516389 | 0.044650949 |
| tal | A0A0K2G788 | Probable transaldolase | 23.467 | 215 | 0.14 | 1.10 | 1.34E-01 | 1.68E-01 | no | no | 0 | 32 | 32 | 29 | 27 | 26 | 31 | 0.071254677 | 0.079234396 | 0.075817091 | 0.078041211 | 0.079363587 | 0.070673361 |
| NITMOv2_0491 | A0A0K2G8L8 | Putative Outer membrane lipoprotein carrier protein LolA | 26.304 | 235 | 0.286 | 1.22 | 1.09E-02 | 1.92E-02 | no | no | 0 | 19 | 22 | 19 | 15 | 11 | 16 | 0.070830994 | 0.054709277 | 0.039157625 | 0.024559645 | 0.026440546 | 0.061359092 |
| NITMOv2_4518 | A0A0K2GJU5 | Uncharacterized protein | 19.037 | 176 | 0.539 | 1.45 | 1.53E-03 | 3.93E-03 | no | no | 0 | 52 | 59 | 60 | 41 | 39 | 42 | 0.070799376 | 0.073675082 | 0.068809594 | 0.044496402 | 0.03497647 | 0.031896874 |
| bkdAA | A0A0K2G7D8 | 2-oxoisovalerate dehydrogenase, alpha subunit (TPP-binding module) | 36.597 | 332 | 3.44 | 10.85 | 1.62E-09 | 1.79E-06 | yes | no | 0 | 57 | 56 | 69 | 8 | 5 | 7 | 0.06993093 | 0.075182726 | 0.079385067 | 0.005352895 | 0.00437066 | 0.004770056 |
| NITMOv2_3006 | A0A0K2GEN8 | Putative NADH-quinone oxidoreductase, membrane subunit M | 39.22 | 379 | 1.24 | 2.36 | 1.65E-05 | 1.64E-04 | yes | no | 0 | 10 | 11 | 9 | 7 | 7 | 7 | 0.069363911 | 0.066283522 | 0.046115274 | 0.030953504 | 0.02934593 | 0.029036063 |
| pup | A0A0K2GHB5 | Bacterial ubiquitin-like modifier | 7.5274 | 65 | -2.08 | -4.23 | 9.64E-03 | 1.73E-02 | yes | no | 0 | 8 | 8 | 10 | 6 | 8 | 5 | 0.069319646 | 0.069335963 | 0.067160771 | 0.051759459 | 0.081809081 | 0.149641456 |
| NITMOv2_4502 | A0A0K2GIW1 | Uncharacterized protein | 36.926 | 342 | 0.386 | 1.31 | 1.03E-03 | 2.88E-03 | no | no | 0 | 52 | 54 | 54 | 49 | 53 | 51 | 0.069132045 | 0.072300408 | 0.069079924 | 0.057664672 | 0.057475327 | 0.05812607 |
| NITMOv2_2883 | A0A0K2GEK4 | Uncharacterized protein | 42.507 | 388 | 0.523 | 1.44 | 8.03E-04 | 2.41E-03 | no | no | 0 | 37 | 43 | 52 | 36 | 34 | 34 | 0.068746303 | 0.066422358 | 0.068339871 | 0.048494415 | 0.047124703 | 0.05172296 |
| NITMOv2_0928 | A0A0K2G9U2 | Uncharacterized protein | 17.794 | 160 | -0.0884 | -1.06 | 2.21E-01 | 2.62E-01 | no | no | 0 | 20 | 20 | 20 | 25 | 26 | 24 | 0.068716793 | 0.074586317 | 0.074753025 | 0.083650275 | 0.097566515 | 0.079410433 |
| NITMOv2_3924 | A0A0K2GHH2 | Uncharacterized protein | 12.528 | 120 | 2.13 | 4.38 | 4.14E-02 | 5.95E-02 | no | no | 0 | 11 | 11 | 7 | 2 | 2 | 2 | 0.068613507 | 0.065777063 | 0.028309904 | 0.016690963 | 0.023098578 | 0.015722593 |
| NITMOv2_0058 | A0A0K2G6E4 | Dihydrolipoamide acetyltransferase component of pyruvate dehydrogenase complex | 43.584 | 404 | 1.85 | 3.61 | 3.16E-07 | 1.46E-05 | yes | no | 0 | 59 | 53 | 52 | 28 | 24 | 21 | 0.06735932 | 0.06324868 | 0.051606237 | 0.013437321 | 0.013822635 | 0.015042544 |
| rpsR | A0A0K2G8B0 | 30S ribosomal protein S18 | 9.2367 | 80 | -0.567 | -1.48 | 6.48E-04 | 2.06E-03 | no | no | 0 | 12 | 11 | 10 | 15 | 14 | 14 | 0.06675436 | 0.069861976 | 0.074438599 | 0.128045986 | 0.123258187 | 0.118379318 |
| folD | A0A0K2G8C4 | Bifunctional protein FolD | 30.512 | 286 | -0.12 | -1.09 | 1.83E-01 | 2.21E-01 | no | no | 0 | 46 | 47 | 50 | 49 | 50 | 54 | 0.066138859 | 0.066827134 | 0.081124 | 0.081035278 | 0.078939432 | 0.066303926 |
| rplK | A0A0K2G9T3 | 50S ribosomal protein L11 | 14.997 | 141 | 0.174 | 1.13 | 8.21E-02 | 1.09E-01 | no | no | 0 | 21 | 21 | 21 | 22 | 23 | 21 | 0.06608827 | 0.064830631 | 0.059852268 | 0.067840355 | 0.070984584 | 0.068155991 |
| gpmA | A0A0K2G771 | 2,3-bisphosphoglycerate-dependent phosphoglycerate mutase | 22.935 | 201 | 0.0727 | 1.05 | 3.00E-01 | 3.42E-01 | no | no | 0 | 24 | 26 | 25 | 24 | 26 | 25 | 0.065902777 | 0.0722613 | 0.069240972 | 0.072945966 | 0.079522062 | 0.073397515 |
| NITMOv2_2401 | A0A0K2GCX8 | NIL domain-containing protein | 9.1694 | 79 | 0.39 | 1.31 | 2.30E-02 | 3.58E-02 | no | no | 0 | 9 | 9 | 9 | 9 | 8 | 7 | 0.065649832 | 0.069349651 | 0.054451415 | 0.05491937 | 0.051997047 | 0.050377965 |
| rplR | A0A0K2G9N2 | 50S ribosomal protein L18 | 13.438 | 121 | -0.672 | -1.59 | 8.36E-05 | 5.03E-04 | no | no | 0 | 9 | 10 | 11 | 9 | 12 | 13 | 0.065616106 | 0.067843962 | 0.071948109 | 0.101820503 | 0.116853913 | 0.114110578 |
| gapA | A0A0K2G8Y1 | Glyceraldehyde-3-phosphate dehydrogenase | 36.41 | 337 | -0.344 | -1.27 | 4.21E-03 | 8.66E-03 | no | no | 0 | 53 | 51 | 56 | 66 | 71 | 67 | 0.065597135 | 0.066670699 | 0.069948432 | 0.097860989 | 0.09774985 | 0.095341788 |
| NITMOv2_3419 | A0A0K2GFT1 | HTH luxR-type domain-containing protein | 22.305 | 202 | 0.607 | 1.52 | 5.06E-04 | 1.71E-03 | no | no | 0 | 31 | 30 | 33 | 25 | 30 | 26 | 0.065548654 | 0.063577164 | 0.064633855 | 0.050272791 | 0.046705209 | 0.049034769 |
| rpsI | A0A0K2GB35 | 30S ribosomal protein S9 | 14.171 | 130 | -0.561 | -1.48 | 1.52E-04 | 7.40E-04 | no | no | 0 | 13 | 11 | 14 | 18 | 18 | 18 | 0.065299924 | 0.066262012 | 0.078637345 | 0.102146267 | 0.111995552 | 0.113682625 |
| NITMOv2_2172 | A0A0K2GCK4 | Heat shock protein, Hsp20 family | 17.085 | 153 | 1.65 | 3.14 | 3.48E-05 | 2.74E-04 | yes | no | 0 | 15 | 12 | 14 | 7 | 6 | 5 | 0.065285169 | 0.051326368 | 0.064578255 | 0.017911098 | 0.017680892 | 0.018225758 |
| NITMOv2_4561 | A0A0K2GJ25 | Uncharacterized protein | 24.294 | 227 | 0.181 | 1.13 | 5.49E-02 | 7.60E-02 | no | no | 0 | 30 | 31 | 44 | 42 | 43 | 39 | 0.064680208 | 0.063455957 | 0.063521858 | 0.060707604 | 0.060546954 | 0.055324597 |
| NITMOv2_3738 | A0A0K2GH05 | Uncharacterized protein | 48.343 | 440 | -0.307 | -1.24 | 5.05E-03 | 1.00E-02 | no | no | 0 | 71 | 71 | 74 | 90 | 88 | 88 | 0.064382998 | 0.064955779 | 0.067454108 | 0.087329927 | 0.085213196 | 0.074839608 |
| rplB | A0A0K2GAK5 | 50S ribosomal protein L2 | 30.666 | 280 | -0.269 | -1.20 | 4.46E-03 | 9.05E-03 | no | no | 0 | 13 | 16 | 17 | 15 | 19 | 14 | 0.064007796 | 0.062896701 | 0.066267339 | 0.079480496 | 0.082348208 | 0.098810364 |
| NITMOv2_1746 | A0A0K2GB19 | Thiamine_BP domain-containing protein | 11.639 | 102 | 0.1 | 1.07 | 6.53E-01 | 6.91E-01 | no | no | 0 | 9 | 12 | 20 | 16 | 15 | 15 | 0.063546171 | 0.062077372 | 0.059238753 | 0.050585228 | 0.045619186 | 0.042957119 |
| NITMOv2_3200 | A0A0K2GFH4 | Copper-containing nitrite reductase | 34.085 | 316 | -0.0756 | -1.05 | 2.85E-01 | 3.28E-01 | no | no | 0 | 35 | 40 | 40 | 49 | 47 | 45 | 0.063225773 | 0.059202877 | 0.058634824 | 0.072581702 | 0.065150512 | 0.063766777 |
| hppA | A0A0K2GGC2 | K(+)-insensitive pyrophosphate-energized proton pump | 69.329 | 685 | 0.485 | 1.40 | 1.05E-03 | 2.93E-03 | no | no | 0 | 31 | 33 | 32 | 28 | 25 | 26 | 0.063126703 | 0.063966327 | 0.064440214 | 0.051540309 | 0.050350145 | 0.050426515 |
| bfrA | A0A0K2G9N6 | Bacterioferritin | 18.034 | 156 | -0.135 | -1.10 | 1.21E-01 | 1.53E-01 | no | no | 0 | 23 | 29 | 32 | 33 | 31 | 33 | 0.062972828 | 0.067654285 | 0.079770431 | 0.107216339 | 0.092184564 | 0.083947093 |
| pdaD | A0A0K2GIA2 | Biodegradative arginine decarboxylase | 21.051 | 189 | 0.0717 | 1.05 | 3.49E-01 | 3.92E-01 | no | no | 0 | 12 | 12 | 14 | 13 | 13 | 16 | 0.062833708 | 0.059146169 | 0.063939815 | 0.055591628 | 0.051482779 | 0.053697297 |
| NITMOv2_0027 | A0A0K2G684 | Nitroreductase | 22.036 | 197 | 0.0307 | 1.02 | 6.80E-01 | 7.17E-01 | no | no | 0 | 16 | 21 | 25 | 28 | 26 | 26 | 0.062768364 | 0.069439601 | 0.060841562 | 0.062369001 | 0.066067184 | 0.065214265 |
| dapL | A0A0K2G7A0 | Aminotransferase | 42.383 | 391 | 0.12 | 1.09 | 1.10E-01 | 1.40E-01 | no | no | 0 | 59 | 64 | 74 | 81 | 85 | 73 | 0.062648215 | 0.062933855 | 0.062549819 | 0.05865381 | 0.057378999 | 0.054439922 |
| cld2 | A0A0K2GDB5 | Chlorite dismutase | 27.847 | 240 | -0.344 | -1.27 | 3.27E-03 | 7.09E-03 | no | no | 0 | 40 | 48 | 49 | 62 | 62 | 67 | 0.062213992 | 0.062595564 | 0.060210791 | 0.085829932 | 0.093306322 | 0.086383547 |
| NITMOv2_0197 | A0A0K2G7P8 | Putative Leu/Ile/Val-binding protein, periplasmic-binding component of ABC transport system | 44.107 | 413 | 0.387 | 1.31 | 1.18E-02 | 2.05E-02 | no | no | 0 | 40 | 39 | 44 | 44 | 36 | 36 | 0.062144432 | 0.065931543 | 0.063117321 | 0.059395664 | 0.055808227 | 0.050403139 |
| NITMOv2_3879 | A0A0K2GH50 | CBM20 domain-containing protein | 11.064 | 98 | 0.477 | 1.39 | 1.45E-03 | 3.74E-03 | no | no | 0 | 8 | 8 | 13 | 18 | 17 | 11 | 0.061973694 | 0.058455899 | 0.057517075 | 0.04173037 | 0.041289077 | 0.041752377 |
| NITMOv2_4221 | A0A0K2GI29 | Uncharacterized protein | 19.824 | 181 | 0.264 | 1.20 | 2.00E-02 | 3.18E-02 | no | no | 0 | 15 | 17 | 15 | 13 | 10 | 12 | 0.061939968 | 0.065661692 | 0.059307773 | 0.050903588 | 0.04858361 | 0.045672282 |

Table S1

| | | | | | | | | | | | | | | | | | | | | | | | |
|---|---|---|---|---|---|---|---|---|---|---|---|---|---|---|---|---|---|---|---|---|---|---|---|
| secD | A0A0K2GGE0 | Protein translocase subunit SecD | 59.945 | 547 | 0.473 | 1.39 | 1.95E-04 | 8.60E-04 | no | no | 0 | 52 | 55 | 50 | 48 | 47 | 46 | 0.06164065 | 0.062096927 | 0.064302173 | 0.047642986 | 0.050486869 | 0.048682337 |
| NITMOv2_2889 | A0A0K2GEA5 | Uncharacterized protein | 27.643 | 254 | 0.325 | 1.25 | 9.88E-03 | 1.77E-02 | no | no | 0 | 25 | 26 | 25 | 24 | 23 | 22 | 0.061541579 | 0.06257992 | 0.060195454 | 0.047333511 | 0.046782893 | 0.04531086 |
| iscU | A0A0K2GID4 | Iron-sulfur cluster assembly scaffold protein IscU | 14.732 | 136 | 1.41 | 2.66 | 2.00E-04 | 8.79E-04 | yes | no | 0 | 39 | 45 | 29 | 18 | 21 | 15 | 0.061282311 | 0.067654285 | 0.052315614 | 0.020985125 | 0.028531801 | 0.029084613 |
| rho | A0A0K2G8R5 | Transcription termination factor Rho | 47.728 | 423 | -1.45 | -2.73 | 2.84E-07 | 1.39E-05 | yes | no | 0 | 81 | 88 | 82 | 149 | 155 | 155 | 0.061010394 | 0.064902982 | 0.067074496 | 0.178237336 | 0.185571677 | 0.179139636 |
| NITMOv2_3002 | A0A0K2GFP7 | Putative Sensor histidine kinase | 73.284 | 667 | 1.04 | 2.06 | 2.22E-06 | 4.54E-05 | yes | no | 0 | 89 | 97 | 93 | 59 | 53 | 51 | 0.060843872 | 0.05910706 | 0.065381577 | 0.032438692 | 0.032551174 | 0.028234101 |
| atpG | A0A0K2G6Y5 | ATP synthase gamma chain | 32.792 | 292 | -0.286 | -1.22 | 4.25E-03 | 8.73E-03 | no | no | 0 | 29 | 32 | 33 | 51 | 53 | 45 | 0.060664703 | 0.062671826 | 0.064576337 | 0.086238618 | 0.079142964 | 0.069655624 |
| lpxA | A0A0K2GG54 | Acyl-[acyl-carrier-protein]--UDP-N-acetylglucosamine O-acyltransferase | 28.896 | 270 | 0.00603 | 1.00 | 9.33E-01 | 9.44E-01 | no | no | 0 | 32 | 32 | 30 | 36 | 39 | 35 | 0.060418081 | 0.060538442 | 0.062626509 | 0.065136514 | 0.062425355 | 0.059827093 |
| fbp | A0A0K2G7T4 | Peptidyl-prolyl cis-trans isomerase | 15.168 | 146 | -0.122 | -1.09 | 3.17E-01 | 3.59E-01 | no | no | 0 | 13 | 11 | 13 | 20 | 20 | 20 | 0.060352737 | 0.062339401 | 0.061338126 | 0.082125107 | 0.072101681 | 0.062642951 |
| cysK | A0A0K2GDY0 | Cysteine synthase A, O-acetylserine sulfhydrolase A subunit | 32.889 | 309 | -0.0456 | -1.03 | 5.88E-01 | 6.28E-01 | no | no | 0 | 36 | 39 | 39 | 41 | 38 | 41 | 0.059631843 | 0.06204413 | 0.0607457 | 0.066248554 | 0.062579169 | 0.058744624 |
| NITMOv2_4038 | A0A0K2GIJ7 | Response regulator, LuxR family | 23.81 | 214 | 0.228 | 1.17 | 8.42E-02 | 1.11E-01 | no | no | 0 | 46 | 44 | 49 | 49 | 42 | 44 | 0.059496939 | 0.057266992 | 0.07179473 | 0.056693303 | 0.054142681 | 0.052902528 |
| NITMOv2_0740 | A0A0K2G877 | Putative nitrite oxidoreductase, membrane subunit | 30.566 | 279 | -0.382 | -1.30 | 4.16E-02 | 5.97E-02 | no | no | 0 | 19 | 18 | 19 | 33 | 26 | 30 | 0.059389437 | 0.062931899 | 0.06863896 | 0.088040685 | 0.077892251 | 0.078755916 |
| nusG | A0A0K2G9J3 | Transcription termination/antitermination protein NusG | 19.889 | 178 | -0.435 | -1.35 | 1.16E-03 | 3.16E-03 | no | no | 0 | 28 | 26 | 22 | 34 | 34 | 32 | 0.059216591 | 0.058105875 | 0.057988715 | 0.091615205 | 0.097162558 | 0.082120201 |
| NITMOv2_1239 | A0A0K2GAN6 | Putative Ribosomal protein S30Ae-family protein, sigma-54 modulation protein | 20.935 | 186 | 0.274 | 1.21 | 3.07E-03 | 6.74E-03 | no | no | 0 | 16 | 17 | 18 | 11 | 14 | 14 | 0.058580012 | 0.063510709 | 0.067015061 | 0.050724418 | 0.056392411 | 0.054792353 |
| NITMOv2_2436 | A0A0K2GDA1 | Response regulator, CheY like | 13.325 | 120 | 0.597 | 1.51 | 3.11E-04 | 1.19E-03 | no | no | 0 | 10 | 13 | 10 | 9 | 8 | 11 | 0.058548394 | 0.067227999 | 0.056019714 | 0.04574319 | 0.051232636 | 0.048319117 |
| NITMOv2_3687 | A0A0K2GGV8 | DUF4115 domain-containing protein | 32.495 | 305 | 0.199 | 1.15 | 3.76E-02 | 5.47E-02 | no | no | 0 | 24 | 23 | 26 | 20 | 21 | 17 | 0.058179516 | 0.059136392 | 0.055864418 | 0.053348299 | 0.052745921 | 0.045497864 |
| NITMOv2_1147 | A0A0K2G9F0 | PHB domain-containing protein | 30.976 | 278 | 0.77 | 1.71 | 1.35E-05 | 1.43E-04 | no | no | 0 | 29 | 24 | 23 | 19 | 16 | 16 | 0.058143682 | 0.058313152 | 0.056207603 | 0.028391814 | 0.036365461 | 0.032080283 |
| dnaK.1 | A0A0K2GIK4 | Chaperone protein DnaK | 66.016 | 606 | 0.789 | 1.73 | 1.74E-05 | 1.69E-04 | no | no | 0 | 70 | 73 | 79 | 54 | 51 | 51 | 0.058086769 | 0.061432077 | 0.064854337 | 0.037528014 | 0.033296941 | 0.034901535 |
| yvqK | A0A0K2GBC6 | Corrinoid adenosyltransferase | 22.317 | 198 | 0.251 | 1.19 | 4.84E-03 | 9.68E-03 | no | no | 0 | 23 | 33 | 32 | 30 | 30 | 29 | 0.058034072 | 0.058792234 | 0.054165747 | 0.049693819 | 0.050409185 | 0.051194313 |
| osmC | A0A0K2GCW1 | Osmotically inducible, stress-inducible membrane protein | 14.86 | 141 | -0.0205 | -1.01 | 8.06E-01 | 8.29E-01 | no | no | 0 | 10 | 11 | 9 | 11 | 10 | 11 | 0.058031964 | 0.055096454 | 0.056000542 | 0.061484995 | 0.057566994 | 0.054806738 |
| ssb | A0A0K2G806 | Single-stranded DNA-binding protein | 14.967 | 136 | -0.146 | -1.11 | 1.94E-01 | 2.32E-01 | no | no | 0 | 15 | 16 | 13 | 18 | 16 | 14 | 0.057218324 | 0.056576722 | 0.058377914 | 0.063661691 | 0.055893679 | 0.056644418 |
| NITMOv2_1178 | A0A0K2G9S1 | Uncharacterized protein | 7.4966 | 69 | 0.557 | 1.47 | 2.76E-01 | 3.19E-01 | no | no | 0 | 7 | 10 | 10 | 6 | 10 | 10 | 0.057087635 | 0.064568602 | 0.064315593 | 0.022569523 | 0.063972821 | 0.070819009 |
| NITMOv2_4410 | A0A0K2GIL1 | VOC domain-containing protein | 22.042 | 198 | 1.19 | 2.28 | 3.30E-05 | 2.66E-04 | yes | no | 0 | 40 | 36 | 39 | 25 | 20 | 22 | 0.056512185 | 0.059590054 | 0.055536571 | 0.0283237 | 0.026053679 | 0.023600343 |
| NITMOv2_3643 | A0A0K2GGF2 | Uncharacterized protein | 71.241 | 634 | -1.24 | -2.36 | 4.05E-06 | 6.54E-05 | yes | no | 0 | 26 | 28 | 27 | 50 | 53 | 47 | 0.056438409 | 0.054734698 | 0.0501338 | 0.140873685 | 0.116777783 | 0.135639485 |
| NITMOv2_3779 | A0A0K2GGT6 | TPR_REGION domain-containing protein | 28.243 | 256 | 0.0303 | 1.02 | 7.25E-01 | 7.59E-01 | no | no | 0 | 23 | 26 | 32 | 40 | 39 | 31 | 0.055833448 | 0.058614289 | 0.058715348 | 0.061344325 | 0.058769543 | 0.056153531 |
| porB | A0A0K2G8G1 | Pyruvate:ferredoxin oxidoreductase, beta subunit | 33.318 | 302 | -0.532 | -1.45 | 2.71E-04 | 1.08E-03 | no | no | 0 | 35 | 41 | 41 | 54 | 51 | 45 | 0.055211625 | 0.054697545 | 0.06309048 | 0.081861534 | 0.086930014 | 0.087059641 |
| NITMOv2_3236 | A0A0K2GFA1 | Cupin_2 domain-containing protein | 11.827 | 105 | 0.617 | 1.53 | 2.22E-04 | 9.51E-04 | no | no | 0 | 14 | 15 | 14 | 11 | 10 | 10 | 0.054819559 | 0.054034651 | 0.0523252 | 0.04157193 | 0.041318597 | 0.038096797 |
| NITMOv2_3641 | A0A0K2GGE8 | Putative monohem cytochrome c | 39.157 | 345 | -1.62 | -3.07 | 7.27E-07 | 2.60E-05 | yes | no | 0 | 25 | 26 | 23 | 62 | 59 | 59 | 0.054815344 | 0.054738609 | 0.053648093 | 0.205334979 | 0.201745498 | 0.191787622 |
| rpmA | A0A0K2G8Y6 | 50S ribosomal protein L27 | 9.3014 | 87 | -0.319 | -1.25 | 7.69E-03 | 1.42E-02 | no | no | 0 | 14 | 11 | 12 | 13 | 12 | 10 | 0.054747892 | 0.061475097 | 0.065207108 | 0.069375888 | 0.085626476 | 0.081611333 |
| NITMOv2_4415 | A0A0K2GIL6 | EVE domain-containing protein | 18.866 | 164 | 0.676 | 1.60 | 1.17E-04 | 6.20E-04 | no | no | 0 | 26 | 27 | 26 | 17 | 16 | 14 | 0.054735244 | 0.057380407 | 0.056704167 | 0.034208183 | 0.035327602 | 0.033754334 |
| NITMOv2_0694 | A0A0K2G870 | Ferredoxin | 26.399 | 237 | 0.0821 | 1.06 | 2.36E-01 | 2.78E-01 | no | no | 0 | 24 | 27 | 27 | 30 | 28 | 22 | 0.054351611 | 0.053166436 | 0.054878959 | 0.055280672 | 0.054916414 | 0.053490514 |
| NITMOv2_2266 | A0A0K2GDI8 | Putative Thioredoxin | 9.6881 | 87 | 0.0455 | 1.03 | 4.69E-01 | 5.12E-01 | no | no | 0 | 12 | 9 | 9 | 7 | 8 | 7 | 0.054052292 | 0.051349833 | 0.058671251 | 0.053733293 | 0.055472632 | 0.051764317 |
| dapB | A0A0K2G7A9 | 4-hydroxy-tetrahydrodipicolinate reductase | 28.619 | 267 | 0.426 | 1.34 | 8.13E-04 | 2.43E-03 | no | no | 0 | 40 | 40 | 44 | 36 | 31 | 31 | 0.053942683 | 0.054417917 | 0.057292759 | 0.038256541 | 0.044425959 | 0.040603378 |
| NITMOv2_3614 | A0A0K2GGD7 | MULTIHEME_CYTC domain-containing protein | 47.543 | 432 | 0.783 | 1.72 | 1.60E-04 | 7.61E-04 | no | no | 0 | 31 | 33 | 37 | 23 | 31 | 30 | 0.053696061 | 0.053244653 | 0.047951986 | 0.024498934 | 0.034614462 | 0.034534718 |
| NITMOv2_3818 | A0A0K2GGW6 | Transcriptional regulator, MarR family | 24.276 | 220 | -0.121 | -1.09 | 1.24E-01 | 1.56E-01 | no | no | 0 | 12 | 12 | 12 | 11 | 14 | 11 | 0.053386203 | 0.052593492 | 0.054372808 | 0.05505856 | 0.063498949 | 0.058091906 |
| folE | A0A0K2GHX1 | GTP cyclohydrolase 1 | 24.718 | 216 | 0.616 | 1.53 | 9.08E-04 | 2.64E-03 | no | no | 0 | 25 | 27 | 26 | 19 | 19 | 17 | 0.053346153 | 0.054421828 | 0.050177896 | 0.034950036 | 0.035113194 | 0.034290174 |
| NITMOv2_3785 | A0A0K2GGW2 | Uncharacterized protein | 10.317 | 90 | 1.19 | 2.28 | 9.45E-05 | 5.41E-04 | yes | no | 0 | 8 | 8 | 10 | 6 | 7 | 6 | 0.053124826 | 0.04857702 | 0.050572847 | 0.026201792 | 0.030090143 | 0.028381547 |
| NITMOv2_0306 | A0A0K2G798 | OMP_b-brl domain-containing protein | 23.646 | 215 | 0.835 | 1.78 | 1.76E-04 | 7.99E-04 | no | no | 0 | 20 | 21 | 21 | 11 | 12 | 10 | 0.052677956 | 0.056416376 | 0.05864441 | 0.039035413 | 0.031246082 | 0.027185796 |
| NITMOv2_0455 | A0A0K2G7D7 | OsmC family protein | 16.44 | 148 | 0.526 | 1.44 | 9.67E-05 | 5.45E-04 | no | no | 0 | 7 | 7 | 6 | 5 | 6 | 5 | 0.052538836 | 0.052055746 | 0.049573967 | 0.024949081 | 0.026524445 | 0.030708316 |
| rpmC | A0A0K2GAK8 | 50S ribosomal protein L29 | 8.2406 | 72 | -0.243 | -1.18 | 1.01E-01 | 1.30E-01 | no | no | 0 | 8 | 8 | 7 | 11 | 11 | 11 | 0.052178389 | 0.061635443 | 0.051126928 | 0.062749552 | 0.073995619 | 0.065624236 |
| NITMOv2_1149 | A0A0K2G9G4 | Uncharacterized protein | 12.213 | 115 | -0.344 | -1.27 | 6.95E-03 | 1.31E-02 | no | no | 0 | 8 | 8 | 8 | 9 | 8 | 8 | 0.051938091 | 0.05098221 | 0.053598245 | 0.082148799 | 0.072707617 | 0.066782226 |
| nrdR | A0A0K2G6T1 | Transcriptional repressor NrdR | 18.253 | 154 | 0.287 | 1.22 | 1.91E-02 | 3.05E-02 | no | no | 0 | 14 | 12 | 16 | 18 | 13 | 12 | 0.051919121 | 0.049709219 | 0.047395987 | 0.050970222 | 0.04422398 | 0.040632148 |
| hisD | A0A0K2G8J1 | Histidinol dehydrogenase | 46.148 | 427 | 0.24 | 1.18 | 5.27E-03 | 1.04E-02 | no | no | 0 | 38 | 39 | 47 | 43 | 43 | 38 | 0.051910689 | 0.050620454 | 0.048676701 | 0.038531959 | 0.040836955 | 0.039272768 |
| atpC | A0A0K2G708 | ATP synthase epsilon chain | 15.509 | 141 | -0.288 | -1.22 | 4.95E-03 | 9.86E-03 | no | no | 0 | 15 | 13 | 16 | 16 | 18 | 18 | 0.051870639 | 0.049439369 | 0.049317057 | 0.050363116 | 0.081782668 | 0.059239107 |
| moaB | A0A0K2GB33 | Molybdenum cofactor biosynthesis protein B | 18.598 | 170 | 0.339 | 1.26 | 4.78E-03 | 9.59E-03 | no | no | 0 | 27 | 24 | 27 | 26 | 22 | 24 | 0.051670391 | 0.049341597 | 0.04869779 | 0.046867076 | 0.043677085 | 0.041353194 |
| yqjG | A0A0K2GH94 | S-transferase | 35.997 | 314 | 0.875 | 1.83 | 1.34E-05 | 1.42E-04 | no | no | 0 | 41 | 49 | 39 | 30 | 30 | 25 | 0.051229845 | 0.052902452 | 0.052363545 | 0.032702264 | 0.03071783 | 0.028536186 |
| NITMOv2_4628 | A0A0K2GJH1 | Putative Peptidylprolyl isomerase, PpiC-type | 33.441 | 292 | 0.262 | 1.20 | 8.68E-03 | 1.58E-02 | no | no | 0 | 49 | 51 | 46 | 46 | 41 | 42 | 0.050901016 | 0.048197665 | 0.052578276 | 0.046523543 | 0.047362416 | 0.042599293 |
| fabG.4 | A0A0K2GHN6 | 3-oxoacyl-(Acyl-carrier-protein) reductase | 26.021 | 244 | 0.479 | 1.39 | 8.23E-04 | 2.45E-03 | no | no | 0 | 24 | 24 | 25 | 15 | 13 | 13 | 0.050848319 | 0.04973464 | 0.050072448 | 0.033760997 | 0.032173629 | 0.032495649 |
| tcmO | A0A0K2GC41 | Tetracenomycin polyketide synthesis 8-O-methyl transferase TcmO | 37.29 | 337 | 0.0894 | 1.06 | 3.06E-01 | 3.47E-01 | no | no | 0 | 47 | 49 | 67 | 63 | 68 | 62 | 0.050591158 | 0.046312622 | 0.05452427 | 0.053207628 | 0.049797035 | 0.048108736 |
| NITMOv2_4526 | A0A0K2GIZ1 | FGE-sulfatase domain-containing protein | 30.929 | 275 | 0.628 | 1.55 | 6.73E-04 | 2.12E-03 | no | no | 0 | 29 | 31 | 28 | 23 | 26 | 20 | 0.049821783 | 0.052055746 | 0.048624935 | 0.037692377 | 0.035019973 | 0.032020945 |
| coaD | A0A0K2GC93 | Phosphopantetheine adenylyltransferase | 18.195 | 162 | 0.198 | 1.15 | 4.57E-02 | 6.50E-02 | no | no | 0 | 33 | 38 | 31 | 28 | 27 | 30 | 0.049672124 | 0.054058116 | 0.056763601 | 0.04300233 | 0.047778803 | 0.041311838 |
| adk | A0A0K2G9N7 | Adenylate kinase | 24.168 | 218 | -0.112 | -1.08 | 2.28E-01 | 2.70E-01 | no | no | 0 | 25 | 29 | 26 | 27 | 36 | 33 | 0.049617319 | 0.049773749 | 0.047614552 | 0.055875931 | 0.059171947 | 0.055712991 |
| NITMOv2_4500 | A0A0K2GJ22 | Uncharacterized protein | 48.263 | 454 | -0.225 | -1.17 | 1.96E-02 | 3.14E-02 | no | no | 0 | 55 | 55 | 65 | 78 | 82 | 70 | 0.049448689 | 0.048700213 | 0.046642513 | 0.046530947 | 0.052629395 | 0.042788095 |
| NITMOv2_0037 | A0A0K2G692 | Putative pterin-4-alpha-carbinolamine dehydratase | 12.849 | 116 | 0.852 | 1.81 | 2.50E-05 | 2.20E-04 | no | no | 0 | 14 | 14 | 15 | 10 | 10 | 7 | 0.049349618 | 0.048358011 | 0.049111913 | 0.033870573 | 0.032510778 | 0.029350735 |
| efp | A0A0K2GC09 | Elongation factor P | 21.06 | 186 | 0.158 | 1.12 | 8.97E-02 | 1.17E-01 | no | no | 0 | 17 | 13 | 15 | 20 | 20 | 17 | 0.049343295 | 0.047002892 | 0.044280479 | 0.041143994 | 0.042945301 | 0.044623977 |
| NITMOv2_1249 | A0A0K2GAP7 | ANTAR domain-containing protein | 47.529 | 420 | 0.567 | 1.48 | 6.45E-05 | 4.18E-04 | no | no | 0 | 47 | 49 | 52 | 38 | 37 | 38 | 0.049292706 | 0.052110499 | 0.051901491 | 0.032415 | 0.033589033 | 0.034667779 |
| NITMOv2_0739 | A0A0K2G8B3 | Putative Cytochrome c | 25.936 | 239 | -0.0738 | -1.05 | 5.78E-01 | 6.18E-01 | no | no | 0 | 20 | 20 | 26 | 27 | 30 | 32 | 0.049143046 | 0.04784373 | 0.042625904 | 0.062714014 | 0.065029325 | 0.061128933 |
| NITMOv2_0735 | A0A0K2G872 | Putative Quinol-cytochrome c reductase, cytochrome b subunit | 42.026 | 373 | 0.447 | 1.36 | 1.20E-03 | 3.19E-03 | no | no | 0 | 14 | 16 | 17 | 15 | 14 | 15 | 0.049031329 | 0.046991159 | 0.049084174 | 0.039094643 | 0.042805469 | 0.039030022 |
| Tktl | A0A0K2GG43 | Transketolase | 66.006 | 621 | -0.211 | -1.16 | 1.33E-01 | 1.67E-01 | no | no | 0 | 50 | 50 | 54 | 78 | 70 | 66 | 0.049008142 | 0.050950923 | 0.049142589 | 0.057294486 | 0.056544672 | 0.052260599 |
| htpX | A0A0K2G9D3 | Protease HtpX | 31.982 | 292 | 0.4 | 1.32 | 8.64E-03 | 1.58E-02 | no | no | 0 | 19 | 23 | 21 | 16 | 16 | 16 | 0.048930151 | 0.05592165 | 0.045829605 | 0.03730146 | 0.041783147 | 0.037636478 |
| rplQ | A0A0K2GAN0 | 50S ribosomal protein L17 | 15.724 | 142 | -0.102 | -1.07 | 2.38E-01 | 2.78E-01 | no | no | 0 | 15 | 19 | 16 | 18 | 21 | 21 | 0.048913288 | 0.048291526 | 0.069994446 | 0.080598458 | 0.072496316 | 0.08022678 |
| NITMOv2_0599 | A0A0K2G7X9 | Uncharacterized protein | 20.289 | 185 | 0.97 | 1.96 | 5.29E-05 | 3.64E-04 | no | no | 0 | 14 | 15 | 13 | 9 | 11 | 9 | 0.048887993 | 0.050196124 | 0.04361003 | 0.02387554 | 0.026592807 | 0.023997727 |
| NITMOv2_0065 | A0A0K2G7E3 | Uncharacterized protein | 27.975 | 243 | 7.17 | 144.01 | 3.41E-05 | 2.70E-04 | yes | yes | 1 | 20 | 24 | 28 | 0 | 0 | 0 | 0.048272493 | 0.051568842 | 0.048214647 | 0.000380063 | 0.000305236 | 0 |
| purH | A0A0K2GBZ2 | Bifunctional purine biosynthesis protein PurH | 55.66 | 517 | -0.388 | -1.31 | 4.87E-04 | 1.66E-03 | no | no | 0 | 49 | 49 | 54 | 66 | 68 | 64 | 0.047855134 | 0.045594975 | 0.047039381 | 0.069473617 | 0.07130775 | 0.067771193 |
| NITMOv2_3597 | A0A0K2GGM9 | Sulfurtransferase | 33.081 | 292 | 0.224 | 1.17 | 1.58E-02 | 2.62E-02 | no | no | 0 | 33 | 36 | 33 | 34 | 33 | 37 | 0.047694935 | 0.046778016 | 0.044299651 | 0.041204705 | 0.041208285 | 0.039839176 |
| NITMOv2_0252 | A0A0K2G6U1 | Uncharacterized protein | 13.768 | 122 | 0.025 | 1.02 | 8.42E-01 | 8.61E-01 | no | no | 0 | 13 | 15 | 17 | 25 | 26 | 21 | 0.04768018 | 0.044149906 | 0.041778486 | 0.047718505 | 0.05263561 | 0.046866235 |
| atoC.4 | A0A0K2GEN3 | Sigma-54 dependent transcriptional regulator | 52.172 | 470 | 1.06 | 2.08 | 7.68E-06 | 9.90E-05 | yes | no | 0 | 70 | 75 | 74 | 37 | 47 | 48 | 0.047654885 | 0.046688066 | 0.04658162 | 0.02238443 | 0.022514394 | 0.021624207 |
| grxD | A0A0K2GHQ0 | Glutaredoxin | 12.238 | 107 | 0.278 | 1.21 | 6.78E-03 | 1.28E-02 | no | no | 0 | 18 | 18 | 18 | 24 | 18 | 19 | 0.047513657 | 0.035522508 | 0.03640064 | 0.041637083 | 0.044918476 | 0.030282161 |
| NITMOv2_2277 | A0A0K2GCL8 | Uncharacterized protein | 14.959 | 134 | 0.725 | 1.65 | 8.43E-04 | 2.49E-03 | no | no | 0 | 12 | 16 | 15 | 13 | 11 | 11 | 0.047138455 | 0.050464019 | 0.036780252 | 0.03203889 | 0.028034623 | 0.026387431 |
| NITMOv2_0851 | A0A0K2G8I6 | Putative Peptidase M48, Ste24p | 30.835 | 289 | 0.274 | 1.21 | 2.15E-02 | 3.39E-02 | no | no | 0 | 23 | 28 | 26 | 17 | 20 | 15 | 0.047127916 | 0.05131268 | 0.04786571 | 0.043533918 | 0.039769576 | 0.036681675 |
| hpt | A0A0K2GBF0 | Hypoxanthine phosphoribosyltransferase | 20.194 | 180 | 0.492 | 1.41 | 2.30E-04 | 9.72E-04 | no | no | 0 | 21 | 24 | 26 | 19 | 20 | 20 | 0.0471237 | 0.047096753 | 0.048784066 | 0.035999885 | 0.036433823 | 0.03429377 |
| modC | A0A0K2GAG6 | Molybdate ABC transporter, ATP-binding protein | 38.89 | 362 | 0.491 | 1.41 | 1.56E-03 | 3.99E-03 | no | no | 0 | 35 | 31 | 40 | 26 | 22 | 24 | 0.047041493 | 0.044224213 | 0.045591868 | 0.030196843 | 0.030354269 | 0.025941497 |
| NITMOv2_0732 | A0A0K2G884 | Uncharacterized protein | 63.087 | 567 | -0.169 | -1.12 | 2.60E-01 | 3.02E-01 | no | no | 0 | 48 | 49 | 60 | 61 | 63 | 60 | 0.046938207 | 0.053377623 | 0.051590899 | 0.051556597 | 0.05136936 | 0.048081765 |
| NITMOv2_2231 | A0A0K2GDF4 | Uncharacterized protein | 28.505 | 254 | 0.454 | 1.37 | 7.65E-04 | 2.32E-03 | no | no | 0 | 26 | 24 | 26 | 21 | 20 | 21 | 0.046837029 | 0.046774105 | 0.048544411 | 0.040554658 | 0.040240342 | 0.035615389 |
| NITMOv2_2895 | A0A0K2GEB8 | Sigma-54 dependent transcriptional regulator | 52.643 | 478 | 0.881 | 1.84 | 7.54E-05 | 4.62E-04 | no | no | 0 | 10 | 10 | 10 | 7 | 7 | 6 | 0.046809627 | 0.037016464 | 0.022075053 | 0.003588735 | 0.003575331 | 0.003141678 |
| NITMOv2_0866 | A0A0K2G8J9 | Putative Peptidyl-prolyl cis-trans isomerase D | 26.268 | 232 | -0.0656 | -1.05 | 4.07E-01 | 4.51E-01 | no | no | 0 | 24 | 24 | 23 | 28 | 31 | 30 | 0.046267902 | 0.044763914 | 0.05247091 | 0.052083742 | 0.052856233 | 0.053432974 |
| yajC | A0A0K2GGF7 | Preprotein translocase, subunit YajC | 12.076 | 111 | 0.571 | 1.49 | 1.90E-03 | 4.62E-03 | no | no | 0 | 7 | 7 | 6 | 4 | 4 | 4 | 0.046044467 | 0.046785838 | 0.048469639 | 0.034812327 | 0.034629999 | 0.031887884 |
| NITMOv2_3615 | A0A0K2GHB7 | MULTIHEME_CYTC domain-containing protein | 48.239 | 441 | 0.689 | 1.61 | 7.02E-04 | 2.19E-03 | no | no | 0 | 31 | 30 | 28 | 29 | 23 | 21 | 0.045907455 | 0.045196066 | 0.038906467 | 0.031341459 | 0.032495241 | 0.027462707 |
| NITMOv2_0409 | A0A0K2G7D5 | Uncharacterized protein | 11.496 | 103 | 0.698 | 1.62 | 3.87E-03 | 8.08E-03 | no | no | 0 | 11 | 13 | 11 | 9 | 9 | 8 | 0.045831572 | 0.04094103 | 0.034979968 | 0.02647721 | 0.025662152 | 0.028624294 |
| tsf | A0A0K2GC36 | Elongation factor Ts | 21.906 | 198 | -0.0803 | -1.06 | 3.00E-01 | 3.42E-01 | no | no | 0 | 21 | 24 | 26 | 25 | 28 | 26 | 0.045726178 | 0.043357953 | 0.044905498 | 0.053459355 | 0.052433632 | 0.051424472 |
| NITMOv2_4057 | A0A0K2GHP2 | PilZ domain-containing protein | 14.184 | 128 | 0.394 | 1.31 | 9.54E-03 | 1.72E-02 | no | no | 0 | 7 | 8 | 9 | 10 | 9 | 9 | 0.04569456 | 0.045467872 | 0.044433858 | 0.037625743 | 0.034642429 | 0.033110606 |

| | | | | | | | | | | | | | | | | | | | | | | | |
|---|---|---|---|---|---|---|---|---|---|---|---|---|---|---|---|---|---|---|---|---|---|---|---|
| NITMOv2_3787 | A0A0K2GGT4 | Putative helicase | 66.862 | 593 | 0.492 | 1.41 | 3.83E-04 | 1.38E-03 | no | no | 0 | 55 | 54 | 63 | 57 | 56 | 53 | 0.045673481 | 0.046177696 | 0.042516622 | 0.03517659 | 0.035605711 | 0.033551146 |
| NITMOv2_0633 | A0A0K2G888 | Putative Cell division protein DivIVA | 19.848 | 174 | -0.286 | -1.22 | 6.54E-03 | 1.24E-02 | no | no | 0 | 31 | 31 | 28 | 34 | 32 | 29 | 0.045498527 | 0.04988912 | 0.046985699 | 0.057073855 | 0.05766643 | 0.05743739 |
| NITMOv2_3431 | A0A0K2GFT8 | Uncharacterized protein | 14.081 | 135 | 0.223 | 1.17 | 1.69E-01 | 2.05E-01 | no | no | 0 | 17 | 15 | 19 | 19 | 14 | 12 | 0.04547534 | 0.044232034 | 0.038605461 | 0.039377465 | 0.040025934 | 0.033455846 |
| NITMOv2_3831 | A0A0K2GGY3 | Putative Universal stress protein | 31.305 | 287 | 1.22 | 2.33 | 1.66E-06 | 3.80E-05 | yes | no | 0 | 24 | 21 | 26 | 16 | 16 | 14 | 0.045458477 | 0.04318783 | 0.048182054 | 0.020178119 | 0.018714089 | 0.018306673 |
| NITMOv2_1591 | A0A0K2GAP3 | Uncharacterized protein | 13.765 | 124 | -0.0195 | -1.01 | 8.95E-01 | 9.10E-01 | no | no | 0 | 17 | 18 | 18 | 22 | 23 | 20 | 0.045294063 | 0.04147291 | 0.040283042 | 0.057415907 | 0.053553836 | 0.048941267 |
| NITMOv2_3628 | A0A0K2GGD6 | WD_REPEATS_REGION domain-containing protein | 40.565 | 366 | -0.191 | -1.14 | 4.56E-02 | 6.49E-02 | no | no | 0 | 39 | 35 | 43 | 62 | 60 | 55 | 0.045036902 | 0.04746242 | 0.049991924 | 0.056225387 | 0.056403287 | 0.053747645 |
| NITMOv2_4245 | A0A0K2GI26 | Uncharacterized protein | 12.944 | 124 | 0.78 | 1.72 | 1.17E-04 | 6.20E-04 | no | no | 0 | 11 | 13 | 13 | 11 | 9 | 9 | 0.04489989 | 0.043582828 | 0.044445361 | 0.031334056 | 0.029886611 | 0.029025275 |
| pyrF | A0A0K2GJ42 | Orotidine 5-phosphate decarboxylase | 25.383 | 238 | 0.151 | 1.11 | 1.58E-01 | 1.94E-01 | no | no | 0 | 31 | 32 | 32 | 28 | 29 | 26 | 0.044813467 | 0.042024343 | 0.047018292 | 0.046083761 | 0.043010555 | 0.037559158 |
| NITMOv2_2288 | A0A0K2GCV8 | Uncharacterized protein | 38.349 | 368 | 0.503 | 1.42 | 3.01E-04 | 1.16E-03 | no | no | 0 | 28 | 25 | 35 | 34 | 36 | 34 | 0.044779741 | 0.047378336 | 0.048038261 | 0.038028506 | 0.039761808 | 0.037524994 |
| iscA | A0A0K2GJ83 | FeS cluster assembly protein | 12.641 | 118 | 0.542 | 1.46 | 7.71E-04 | 2.34E-03 | no | no | 0 | 16 | 17 | 19 | 15 | 14 | 13 | 0.044566845 | 0.038858487 | 0.050638033 | 0.035089226 | 0.033460077 | 0.030898917 |
| NITMOv2_4788 | A0A0K2GJP0 | Uncharacterized protein | 9.0254 | 80 | 0.202 | 1.15 | 2.54E-02 | 3.91E-02 | no | no | 0 | 4 | 4 | 4 | 4 | 5 | 4 | 0.044526796 | 0.04646319 | 0.047580042 | 0.045880899 | 0.043886831 | 0.039163083 |
| NITMOv2_4544 | A0A0K2GIZ7 | YceI domain-containing protein | 22.368 | 203 | 0.411 | 1.33 | 2.20E-03 | 5.17E-03 | no | no | 0 | 28 | 27 | 31 | 23 | 22 | 18 | 0.044509933 | 0.04183271 | 0.041023095 | 0.030530011 | 0.029547908 | 0.028989312 |
| NITMOv2_0020 | A0A0K2G6J2 | Uncharacterized protein | 62.039 | 558 | 0.383 | 1.30 | 4.08E-03 | 8.47E-03 | no | no | 0 | 74 | 70 | 76 | 62 | 57 | 61 | 0.044438265 | 0.046119033 | 0.043496329 | 0.031242249 | 0.031160629 | 0.029611462 |
| NITMOv2_0569 | A0A0K2G7U8 | Histidine kinase domain-containing protein | 68.108 | 627 | 0.0669 | 1.05 | 2.37E-01 | 2.78E-01 | no | no | 0 | 79 | 83 | 104 | 93 | 90 | 89 | 0.044233801 | 0.044286787 | 0.04528511 | 0.044249118 | 0.040941052 | 0.037881022 |
| ilvD | A0A0K2GEK7 | Dihydroxy-acid dehydratase | 58.621 | 557 | 1.04 | 2.06 | 5.57E-06 | 7.92E-05 | yes | no | 0 | 49 | 48 | 50 | 28 | 30 | 26 | 0.044189535 | 0.046961827 | 0.047997999 | 0.02008039 | 0.020298844 | 0.018768791 |
| NITMOv2_0200 | A0A0K2G6T4 | Uncharacterized protein | 39.964 | 356 | 0.364 | 1.29 | 8.56E-03 | 1.56E-02 | no | no | 0 | 42 | 42 | 45 | 36 | 38 | 38 | 0.044119975 | 0.044419756 | 0.04316273 | 0.025773856 | 0.026196618 | 0.025416445 |
| NITMOv2_1319 | A0A0K2GAX6 | Uncharacterized protein | 21.869 | 198 | 0.52 | 1.43 | 3.76E-04 | 1.36E-03 | no | no | 0 | 31 | 29 | 30 | 22 | 24 | 22 | 0.044031445 | 0.051979484 | 0.050101207 | 0.038469768 | 0.035936645 | 0.033412691 |
| tig | A0A0K2G6S9 | Trigger factor | 49.329 | 440 | -0.254 | -1.19 | 2.67E-02 | 4.07E-02 | no | no | 0 | 44 | 50 | 53 | 57 | 63 | 58 | 0.043923943 | 0.048584842 | 0.043086041 | 0.069343312 | 0.065046415 | 0.058411972 |
| NITMOv2_3612 | A0A0K2GGP3 | Uncharacterized protein | 9.5548 | 84 | 0.535 | 1.45 | 1.77E-03 | 4.38E-03 | no | no | 0 | 7 | 7 | 8 | 8 | 6 | 7 | 0.043740557 | 0.042673549 | 0.040785358 | 0.029955481 | 0.028365557 | 0.024520981 |
| NITMOv2_3370 | A0A0K2GGL3 | UDP-glucose/GDP-mannose dehydrogenase family protein | 47.222 | 431 | -0.0251 | -1.02 | 7.68E-01 | 7.96E-01 | no | no | 0 | 47 | 47 | 47 | 54 | 58 | 50 | 0.043702616 | 0.041746671 | 0.035917496 | 0.041398683 | 0.045303789 | 0.038997656 |
| NITMOv2_3836 | A0A0K2GGY8 | Uncharacterized protein | 49.296 | 468 | 0.765 | 1.70 | 6.43E-05 | 4.18E-04 | no | no | 0 | 37 | 42 | 49 | 39 | 35 | 36 | 0.043428592 | 0.044206614 | 0.043417723 | 0.028362199 | 0.026929955 | 0.026694909 |
| NITMOv2_4189 | A0A0K2GI85 | Uncharacterized protein | 8.3196 | 73 | -0.366 | -1.29 | 3.26E-02 | 4.86E-02 | no | no | 0 | 15 | 15 | 13 | 15 | 16 | 13 | 0.043116626 | 0.045845272 | 0.045557358 | 0.026315809 | 0.03606094 | 0.036548614 |
| infB | A0A0K2GJQ7 | Translation initiation factor IF-2 | 89.767 | 838 | -0.346 | -1.27 | 1.79E-03 | 4.40E-03 | no | no | 0 | 69 | 73 | 74 | 90 | 89 | 83 | 0.043089223 | 0.042542535 | 0.042606732 | 0.059238705 | 0.057250043 | 0.054835508 |
| NITMOv2_2530 | A0A0K2GDC5 | Uncharacterized protein | 44.016 | 394 | -0.533 | -1.45 | 1.70E-04 | 7.81E-04 | no | no | 0 | 26 | 21 | 31 | 50 | 44 | 39 | 0.042998585 | 0.041212836 | 0.043231751 | 0.063161199 | 0.063961946 | 0.058300488 |
| forB | A0A0K2G8D4 | 2-oxoacid:ferredoxin oxidoreductase, beta subunit | 32.939 | 299 | -1.17 | -2.25 | 1.12E-06 | 3.12E-05 | yes | no | 0 | 27 | 30 | 29 | 67 | 66 | 67 | 0.042996477 | 0.043293423 | 0.042880897 | 0.103077656 | 0.112309396 | 0.11135226 |
| NITMOv2_1173 | A0A0K2G9R6 | ABC_transp_aux domain-containing protein | 56.49 | 507 | 0.258 | 1.20 | 1.87E-02 | 3.00E-02 | no | no | 0 | 56 | 59 | 62 | 59 | 53 | 52 | 0.042867896 | 0.044343494 | 0.043045779 | 0.038936203 | 0.038164624 | 0.039229613 |
| acrB.1 | A0A0K2GAM3 | RND-type permease AcrB | 117.63 | 1089 | 0.576 | 1.49 | 2.35E-04 | 9.83E-04 | no | no | 0 | 109 | 117 | 122 | 116 | 103 | 108 | 0.042726669 | 0.043625848 | 0.041178391 | 0.030581837 | 0.02890313 | 0.028872434 |
| atoB | A0A0K2G6M4 | Acetyl-CoA acetyltransferase | 41.363 | 398 | 2.19 | 4.56 | 1.13E-06 | 3.12E-05 | yes | no | 0 | 34 | 31 | 29 | 10 | 10 | 8 | 0.042614951 | 0.04361216 | 0.039443293 | 0.0083748 | 0.008617648 | 0.007963879 |
| proS | A0A0K2GAC6 | Proline--tRNA ligase | 62.763 | 567 | 0.492 | 1.41 | 5.11E-04 | 1.72E-03 | no | no | 0 | 66 | 65 | 70 | 80 | 71 | 74 | 0.042608627 | 0.042118205 | 0.03830062 | 0.029983616 | 0.033143126 | 0.03035049 |
| NITMOv2_3647 | A0A0K2GGS3 | FGE-sulfatase domain-containing protein | 34.009 | 304 | 0.404 | 1.32 | 3.55E-03 | 7.59E-03 | no | no | 0 | 34 | 40 | 36 | 37 | 32 | 32 | 0.042562254 | 0.045090472 | 0.044167362 | 0.036623278 | 0.034048922 | 0.033668024 |
| NITMOv2_4554 | A0A0K2GJ06 | OMP_b-brl domain-containing protein | 23.901 | 233 | 0.829 | 1.78 | 7.42E-04 | 2.28E-03 | no | no | 0 | 14 | 15 | 14 | 13 | 10 | 11 | 0.042267151 | 0.044570325 | 0.048527156 | 0.032893281 | 0.023092363 | 0.029057641 |
| prpC | A0A0K2G6L4 | Citrate synthase | 43.334 | 392 | 3.01 | 8.06 | 1.30E-08 | 3.40E-06 | yes | no | 0 | 36 | 39 | 44 | 11 | 9 | 9 | 0.04221867 | 0.039099006 | 0.044748284 | 0.003276001 | 0.003153041 | 0.003371478 |
| infA | A0A0K2GAM5 | Translation initiation factor IF-1 | 8.2917 | 72 | -0.498 | -1.41 | 3.81E-02 | 5.54E-02 | no | no | 0 | 4 | 6 | 6 | 6 | 7 | 7 | 0.04192146 | 0.046449502 | 0.052156484 | 0.058692309 | 0.066489785 | 0.061456191 |
| arc | A0A0K2GG68 | Proteasome-associated ATPase | 65.56 | 589 | -0.0685 | -1.05 | 4.44E-01 | 4.87E-01 | no | no | 0 | 45 | 60 | 56 | 71 | 70 | 64 | 0.04170224 | 0.046602027 | 0.042919241 | 0.048911985 | 0.051025996 | 0.044703095 |
| NITMOv2_3791 | A0A0K2GGW7 | Uncharacterized protein | 15.073 | 140 | 0.181 | 1.13 | 2.99E-02 | 4.51E-02 | no | no | 0 | 14 | 11 | 12 | 10 | 9 | 10 | 0.04167273 | 0.041883552 | 0.039489307 | 0.032191407 | 0.034031832 | 0.032218738 |
| ltaA | A0A0K2GAZ3 | L-allo-threonine aldolase | 36.928 | 343 | 0.334 | 1.26 | 3.58E-03 | 7.63E-03 | no | no | 0 | 30 | 29 | 31 | 29 | 30 | 27 | 0.041407138 | 0.040860857 | 0.039017667 | 0.033204237 | 0.028121629 | 0.026944848 |
| NITMOv2_3725 | A0A0K2GHL0 | Uncharacterized protein | 7.761 | 69 | 1.59 | 3.01 | 1.66E-04 | 7.81E-04 | yes | no | 0 | 8 | 7 | 6 | 4 | 2 | 1 | 0.040939189 | 0.042759589 | 0.030292326 | 0.023915521 | 0.01534633 | 0.009583267 |
| NITMOv2_1616 | A0A0K2GAS0 | Uncharacterized protein | 40.153 | 361 | 0.272 | 1.21 | 1.02E-02 | 1.81E-02 | no | no | 0 | 35 | 37 | 45 | 33 | 38 | 36 | 0.040795853 | 0.040731798 | 0.038473172 | 0.03175903 | 0.032558942 | 0.032323029 |
| smtA | A0A0K2GG42 | SAM-dependent methyltransferase | 25.26 | 223 | 0.696 | 1.62 | 3.12E-04 | 1.19E-03 | no | no | 0 | 23 | 23 | 21 | 12 | 14 | 11 | 0.040781098 | 0.039789276 | 0.037717781 | 0.024626278 | 0.026755943 | 0.02279838 |
| NITMOv2_3280 | A0A0K2GFE0 | Uncharacterized protein | 85.804 | 786 | 1.27 | 2.41 | 3.93E-05 | 2.98E-04 | yes | no | 0 | 89 | 87 | 96 | 44 | 44 | 52 | 0.040410112 | 0.038187772 | 0.039151873 | 0.014530852 | 0.014040772 | 0.018466706 |
| NITMOv2_0736 | A0A0K2G984 | Uncharacterized protein | 34.853 | 323 | 0.32 | 1.25 | 2.34E-02 | 3.64E-02 | no | no | 0 | 25 | 26 | 22 | 21 | 24 | 20 | 0.04012344 | 0.042065408 | 0.037092762 | 0.035477181 | 0.04210942 | 0.035453558 |
| rpmD | A0A0K2GAL7 | 50S ribosomal protein L30 | 7.8191 | 72 | -0.428 | -1.35 | 1.19E-03 | 3.18E-03 | no | no | 0 | 11 | 11 | 12 | 11 | 9 | 11 | 0.03989579 | 0.041480731 | 0.039113528 | 0.04516718 | 0.052130664 | 0.05141728 |
| ilvC | A0A0K2G8M2 | Ketol-acid reductoisomerase (NADP(+)) | 36.594 | 337 | -1.17 | -2.25 | 1.82E-06 | 4.01E-05 | yes | no | 0 | 24 | 25 | 31 | 52 | 54 | 60 | 0.039741915 | 0.040956674 | 0.039640768 | 0.099202545 | 0.099741669 | 0.099727406 |
| NITMOv2_1456 | A0A0K2GAA6 | TPR_REGION domain-containing protein | 15.447 | 147 | -0.0388 | -1.03 | 7.18E-01 | 7.52E-01 | no | no | 0 | 15 | 14 | 17 | 12 | 17 | 14 | 0.038848175 | 0.040948852 | 0.040633896 | 0.055510187 | 0.051832357 | 0.04283125 |
| NITMOv2_2058 | A0A0K2GC01 | Putative periplasmic serine endoprotease DegP-like | 48.979 | 463 | 0.319 | 1.25 | 4.76E-03 | 9.55E-03 | no | no | 0 | 48 | 48 | 44 | 54 | 50 | 37 | 0.038839743 | 0.040655536 | 0.038488509 | 0.035266916 | 0.032962899 | 0.03105895 |
| purS | A0A0K2GAV8 | Phosphoribosylformylglycinamidine synthase subunit PurS | 8.9043 | 79 | 0.243 | 1.18 | 3.09E-02 | 4.64E-02 | no | no | 0 | 7 | 8 | 6 | 9 | 9 | 5 | 0.038738565 | 0.037910099 | 0.039648437 | 0.03904874 | 0.037059957 | 0.031672109 |
| NITMOv2_0208 | A0A0K2G7Q6 | Putative RNA polymerase-binding protein DksA | 17.005 | 150 | 0.295 | 1.23 | 4.50E-03 | 9.10E-03 | no | no | 0 | 9 | 10 | 11 | 6 | 9 | 8 | 0.038725918 | 0.039857717 | 0.036920211 | 0.029095168 | 0.032931826 | 0.030179669 |
| hflK | A0A0K2GI56 | Protein HflK | 38.232 | 343 | 0.588 | 1.50 | 6.61E-05 | 4.24E-04 | no | no | 0 | 23 | 20 | 24 | 26 | 25 | 23 | 0.038711163 | 0.036224511 | 0.037188624 | 0.026209195 | 0.026807215 | 0.025254614 |
| glyA | A0A0K2G6K8 | Serine hydroxymethyltransferase | 45.402 | 420 | -0.384 | -1.30 | 5.12E-03 | 1.01E-02 | no | no | 0 | 27 | 30 | 36 | 52 | 47 | 53 | 0.038702731 | 0.037200275 | 0.035911744 | 0.057833477 | 0.052115127 | 0.054031748 |
| NITMOv2_0246 | A0A0K2G6X8 | SmpA_OmlA domain-containing protein | 13.933 | 125 | 0.54 | 1.45 | 1.18E-04 | 6.25E-04 | no | no | 0 | 14 | 13 | 10 | 10 | 11 | 12 | 0.038388658 | 0.036359436 | 0.038248855 | 0.026123312 | 0.025943368 | 0.024697197 |
| glgC1 | A0A0K2GD78 | Glucose-1-phosphate adenylyltransferase | 46.271 | 412 | 0.524 | 1.44 | 1.58E-04 | 7.58E-04 | no | no | 0 | 38 | 44 | 43 | 28 | 26 | 29 | 0.03803875 | 0.039466629 | 0.0394107 | 0.028451044 | 0.026808768 | 0.026813585 |
| clpC | A0A0K2GD33 | Negative regulator of genetic competence ClpC/MecB | 91.53 | 808 | -0.86 | -1.82 | 3.35E-05 | 2.67E-04 | no | no | 0 | 72 | 75 | 74 | 125 | 129 | 115 | 0.037950219 | 0.038537795 | 0.039447127 | 0.073798875 | 0.073503102 | 0.062597998 |
| NITMOv2_4541 | A0A0K2GJ04 | FGE-sulfatase domain-containing protein | 33.845 | 304 | -0.185 | -1.14 | 1.42E-01 | 1.75E-01 | no | no | 0 | 22 | 20 | 20 | 23 | 22 | 22 | 0.037935464 | 0.039896826 | 0.033762523 | 0.032151427 | 0.032495241 | 0.034683962 |
| NITMOv2_2969 | A0A0K2GET9 | Uncharacterized protein | 17.193 | 156 | 2.49 | 5.62 | 2.46E-04 | 1.01E-03 | yes | no | 0 | 11 | 10 | 10 | 1 | 3 | 3 | 0.03777105 | 0.037079038 | 0.033565048 | 0.004998849 | 0.004825578 | 0.01192586 |
| hemL | A0A0K2GJ57 | Glutamate-1-semialdehyde 2,1-aminomutase | 45.305 | 427 | 0.355 | 1.28 | 3.09E-03 | 6.76E-03 | no | no | 0 | 40 | 40 | 46 | 38 | 33 | 37 | 0.037766834 | 0.038357895 | 0.038486592 | 0.029688947 | 0.030001583 | 0.027930219 |
| NITMOv2_3301 | A0A0K2GFR2 | Uncharacterized protein | 39.53 | 375 | 0.336 | 1.26 | 7.65E-03 | 1.42E-02 | no | no | 0 | 32 | 31 | 29 | 27 | 28 | 32 | 0.037728892 | 0.037096637 | 0.03635846 | 0.037492476 | 0.034224488 | 0.030852166 |
| NITMOv2_2409 | A0A0K2GD76 | Uncharacterized protein | 29.64 | 264 | 0.00675 | 1.00 | 9.34E-01 | 9.44E-01 | no | no | 0 | 44 | 41 | 34 | 39 | 37 | 35 | 0.037627714 | 0.036390723 | 0.031122489 | 0.039067989 | 0.037260381 | 0.034392667 |
| NITMOv2_4597 | A0A0K2GJE1 | Uncharacterized protein | 17.584 | 159 | 0.615 | 1.53 | 5.88E-04 | 1.93E-03 | no | no | 0 | 11 | 12 | 16 | 9 | 10 | 9 | 0.037612959 | 0.035884264 | 0.043041945 | 0.02943574 | 0.028778836 | 0.022048564 |
| rfaEa | A0A0K2GFG7 | D-beta-D-heptose 7-phosphate kinase, domain I of bifunctional protein HldE | 36.094 | 338 | -0.325 | -1.25 | 1.52E-02 | 2.54E-02 | no | no | 0 | 22 | 22 | 29 | 39 | 29 | 33 | 0.037433789 | 0.035978126 | 0.037573988 | 0.044275771 | 0.040641192 | 0.040042364 |
| mtnP | A0A0K2GHH8 | S-methyl-5-thioadenosine phosphorylase | 32.01 | 299 | -0.0487 | -1.03 | 4.16E-01 | 4.59E-01 | no | no | 0 | 19 | 18 | 22 | 25 | 25 | 24 | 0.037060695 | 0.036543247 | 0.039075184 | 0.042378936 | 0.042260127 | 0.039693528 |
| NITMOv2_2480 | A0A0K2GD69 | IPT/TIG domain-containing protein | 60.375 | 568 | -0.242 | -1.18 | 1.14E-02 | 1.99E-02 | no | no | 0 | 60 | 60 | 59 | 70 | 64 | 69 | 0.036978488 | 0.040933208 | 0.039822906 | 0.056685899 | 0.05555187 | 0.056887165 |
| NITMOv2_0360 | A0A0K2G741 | Putative Peptidase S49, signal peptide peptidase SppA | 32.604 | 300 | 0.162 | 1.12 | 9.77E-02 | 1.26E-01 | no | no | 0 | 21 | 25 | 23 | 22 | 21 | 22 | 0.03662858 | 0.038455667 | 0.037081258 | 0.036661778 | 0.03577817 | 0.032857071 |
| ntrX | A0A0K2G9K9 | Nitrogen assimilation regulatory protein NtrX | 52.353 | 464 | 0.174 | 1.13 | 2.57E-02 | 3.94E-02 | no | no | 0 | 46 | 45 | 47 | 40 | 35 | 34 | 0.03645995 | 0.035037559 | 0.034815086 | 0.031513226 | 0.030411755 | 0.034788253 |
| NITMOv2_4858 | A0A0K2GJV9 | Uncharacterized protein | 10.178 | 89 | 0.029 | 1.02 | 7.03E-01 | 7.39E-01 | no | no | 0 | 8 | 9 | 8 | 15 | 13 | 10 | 0.036440979 | 0.035452112 | 0.035486118 | 0.060821621 | 0.05958678 | 0.055038696 |
| NITMOv2_3784 | A0A0K2GGU1 | FGE-sulfatase domain-containing protein | 30.707 | 281 | 0.548 | 1.46 | 4.89E-04 | 1.66E-03 | no | no | 0 | 11 | 11 | 12 | 9 | 11 | 11 | 0.035888715 | 0.035199861 | 0.032178886 | 0.026031506 | 0.025366952 | 0.024992088 |
| NITMOv2_0185 | A0A0K2G6R9 | Putative Gamma-glutamyl-gamma-aminobutyrate hydrolase | 26.985 | 246 | 0.0865 | 1.06 | 3.26E-01 | 3.68E-01 | no | no | 0 | 50 | 47 | 41 | 36 | 34 | 36 | 0.035825479 | 0.038275767 | 0.034533252 | 0.040442121 | 0.039713644 | 0.039346491 |
| aspC | A0A0K2GBZ8 | Aminotransferase | 43.002 | 398 | 0.344 | 1.27 | 1.55E-03 | 3.97E-03 | no | no | 0 | 29 | 29 | 32 | 29 | 31 | 23 | 0.035779106 | 0.035808002 | 0.035541718 | 0.02889971 | 0.028272336 | 0.027849303 |
| yrdA | A0A0K2G6T3 | Protein YrdA | 20.665 | 191 | 2.32 | 4.99 | 1.42E-01 | 1.76E-01 | no | yes | 1 | 4 | 4 | 4 | 4 | 3 | 2 | 0.035599936 | 0.034073528 | 0.028516966 | 0.023749677 | 0.023847452 | 0.00900661 |
| NITMOv2_1475 | A0A0K2GBF3 | Putative Outer membrane protein TolC | 49.119 | 453 | 0.458 | 1.37 | 9.21E-04 | 2.67E-03 | no | no | 0 | 38 | 40 | 41 | 47 | 47 | 39 | 0.035568318 | 0.041330163 | 0.0442939 | 0.037787144 | 0.035270116 | 0.031909461 |
| NITMOv2_2080 | A0A0K2GD28 | Uncharacterized protein | 31.976 | 289 | 0.343 | 1.27 | 1.18E-03 | 3.17E-03 | no | no | 0 | 22 | 21 | 19 | 18 | 17 | 21 | 0.035399688 | 0.032200217 | 0.036128392 | 0.026543844 | 0.026597468 | 0.028527195 |
| dapA | A0A0K2G8B5 | 4-hydroxy-tetrahydrodipicolinate synthase | 30.99 | 290 | -0.298 | -1.23 | 5.38E-03 | 1.05E-02 | no | no | 0 | 22 | 25 | 28 | 40 | 36 | 34 | 0.035205763 | 0.034984762 | 0.036511839 | 0.046249605 | 0.044429066 | 0.041394551 |
| NITMOv2_1429 | A0A0K2GA59 | Uncharacterized protein | 24.97 | 246 | 0.463 | 1.38 | 1.81E-03 | 4.45E-03 | no | no | 0 | 9 | 8 | 8 | 6 | 7 | 4 | 0.035197332 | 0.0343512 | 0.034466149 | 0.020884435 | 0.02809677 | 0.024794295 |
| NITMOv2_3625 | A0A0K2GHC7 | Uncharacterized protein | 37.924 | 337 | -0.686 | -1.61 | 3.70E-04 | 1.35E-03 | no | no | 0 | 24 | 28 | 25 | 37 | 30 | 28 | 0.03515939 | 0.032055515 | 0.033206525 | 0.053183936 | 0.052561033 | 0.046821281 |
| NITMOv2_4438 | A0A0K2GJK9 | Uncharacterized protein | 41.194 | 376 | 0.44 | 1.36 | 2.69E-03 | 6.06E-03 | no | no | 0 | 39 | 44 | 40 | 39 | 40 | 33 | 0.035066643 | 0.035053203 | 0.032986043 | 0.027584808 | 0.026670491 | 0.024835652 |
| ftsH | A0A0K2GB75 | ATP-dependent zinc metalloprotease FtsH | 67.383 | 603 | -0.294 | -1.23 | 9.55E-03 | 1.72E-02 | no | no | 0 | 43 | 45 | 56 | 64 | 55 | 64 | 0.034809482 | 0.032540463 | 0.033866054 | 0.039608461 | 0.039858136 | 0.036785966 |
| NITMOv2_0963 | A0A0K2G8X6 | Putative Regulatory protein, FmdB family | 12.894 | 117 | 0.786 | 1.72 | 2.45E-03 | 5.62E-03 | no | no | 0 | 3 | 3 | 3 | 4 | 3 | 3 | 0.03478208 | 0.03430427 | 0.034746065 | 0.025116405 | 0.024464263 | 0.018793964 |
| ppa | A0A0K2GBL8 | Inorganic diphosphatase | 25.257 | 228 | 0.126 | 1.09 | 1.84E-01 | 2.22E-01 | no | no | 0 | 24 | 24 | 27 | 26 | 23 | 22 | 0.034777864 | 0.03724525 | 0.034519831 | 0.028113434 | 0.02855666 | 0.02874297 |
| NITMOv2_0447 | A0A0K2G7E8 | Uncharacterized protein | 43.461 | 393 | 0.146 | 1.11 | 1.46E-01 | 1.81E-01 | no | no | 0 | 33 | 38 | 42 | 49 | 47 | 42 | 0.034746246 | 0.034048107 | 0.038296786 | 0.038546767 | 0.037793294 | 0.036636722 |
| NITMOv2_3859 | A0A0K2GH30 | Uncharacterized protein | 15.949 | 147 | 0.289 | 1.22 | 9.64E-02 | 1.25E-01 | no | no | 0 | 16 | 16 | 18 | 16 | 13 | 14 | 0.03467247 | 0.03274774 | 0.041310681 | 0.028094184 | 0.030080821 | 0.025380483 |
| hscB | A0A0K2GIA3 | Co-chaperone protein HscB (Hsc20) | 27.401 | 236 | 0.916 | 1.89 | 8.38E-05 | 5.03E-04 | no | no | 0 | 20 | 24 | 28 | 30 | 28 | 26 | 0.034400554 | 0.034317958 | 0.035591566 | 0.01952511 | 0.018266629 | 0.016618957 |

| | | | | | | | | | | | | | | | | | | | | | | | |
|---|---|---|---|---|---|---|---|---|---|---|---|---|---|---|---|---|---|---|---|---|---|---|---|
| fadB | A0A0K2G7D1 | Putative enoyl-CoA hydratase | 27.154 | 257 | 2.46 | 5.50 | 7.29E-07 | 2.60E-05 | yes | no | 0 | 46 | 51 | 39 | 9 | 11 | 12 | 0.034387907 | 0.034691447 | 0.031872129 | 0.005103389 | 0.005744736 | 0.005318663 |
| sdhA or nadB | A0A0K2G833 | Succinate dehydrogenase/fumarate reductase, flavoprotein subunit | 57.101 | 527 | 0.0517 | 1.04 | 5.68E-01 | 6.09E-01 | no | no | 0 | 34 | 35 | 41 | 41 | 42 | 41 | 0.0342256 | 0.034709045 | 0.036114971 | 0.043591667 | 0.041062239 | 0.036527037 |
| NITMOv2_0944 | A0A0K2G8U3 | Putative Aminomethyltransferase, glycine cleavage system T protein | 41.016 | 373 | 0.175 | 1.13 | 7.81E-02 | 1.04E-01 | no | no | 0 | 29 | 25 | 33 | 29 | 33 | 27 | 0.034143393 | 0.033244422 | 0.034684714 | 0.027598135 | 0.030407094 | 0.028548772 |
| rng | A0A0K2G8M1 | Ribonuclease G | 57.343 | 502 | -0.433 | -1.35 | 1.63E-03 | 4.10E-03 | no | no | 0 | 47 | 50 | 63 | 77 | 74 | 69 | 0.034101236 | 0.034759887 | 0.039537237 | 0.049674569 | 0.049599717 | 0.046414906 |
| gdhA | A0A0K2G876 | Glutamate dehydrogenase | 45.45 | 418 | 0.518 | 1.43 | 1.72E-04 | 7.89E-04 | no | no | 0 | 28 | 30 | 33 | 29 | 29 | 26 | 0.034059078 | 0.034003132 | 0.03737843 | 0.027045817 | 0.026342664 | 0.025189882 |
| thiG | A0A0K2GH65 | Thiazole synthase | 27.471 | 257 | 0.0387 | 1.03 | 6.44E-01 | 6.83E-01 | no | no | 0 | 21 | 23 | 20 | 25 | 23 | 20 | 0.03399795 | 0.035946838 | 0.035108423 | 0.037214096 | 0.039182285 | 0.03689745 |
| def | A0A0K2GCZ5 | Peptide deformylase | 19.947 | 179 | 0.659 | 1.58 | 2.31E-04 | 9.74E-04 | no | no | 0 | 14 | 18 | 18 | 11 | 12 | 12 | 0.03362064 | 0.034591719 | 0.038262276 | 0.024495973 | 0.023595756 | 0.019559964 |
| NITMOv2_4165 | A0A0K2GHX0 | Putative UDP-glucose 4-epimerase | 33.768 | 307 | -0.513 | -1.43 | 3.72E-04 | 1.35E-03 | no | no | 0 | 19 | 17 | 17 | 36 | 41 | 32 | 0.033546864 | 0.031026954 | 0.031584543 | 0.050791051 | 0.055722775 | 0.050052506 |
| atoC.1 | A0A0K2G9H9 | Acetoacetate metabolism regulatory protein AtoC | 51.633 | 467 | -0.333 | -1.26 | 3.33E-03 | 7.18E-03 | no | no | 0 | 16 | 15 | 14 | 25 | 31 | 28 | 0.033424607 | 0.015256336 | 0.032389782 | 0.036418936 | 0.037766881 | 0.034477178 |
| NITMOv2_0036 | A0A0K2G6C6 | Sulfiredoxin | 36.637 | 330 | 0.48 | 1.39 | 6.56E-04 | 2.08E-03 | no | no | 0 | 22 | 26 | 28 | 35 | 34 | 25 | 0.033409852 | 0.03613065 | 0.034792079 | 0.025896758 | 0.027170776 | 0.028613505 |
| NITMOv2_0064 | A0A0K2G6B6 | Putative 4-hydroxyphenylpyruvate dioxygenase | 21.694 | 194 | 5.95 | 61.82 | 7.41E-08 | 7.47E-06 | yes | yes | 3 | 18 | 19 | 19 | 0 | 0 | 0 | 0.0333424 | 0.047065466 | 0.049432092 | 0.002157002 | 0.001600913 | 0.000198279 |
| glnB.2 | A0A0K2GJU3 | Nitrogen regulatory protein P-II | 12.451 | 113 | -0.0057 | -1.00 | 9.65E-01 | 9.69E-01 | no | no | 0 | 11 | 8 | 10 | 18 | 14 | 13 | 0.03324333 | 0.024873186 | 0.028938758 | 0.040602042 | 0.037965753 | 0.033867615 |
| NITMOv2_0712 | A0A0K2G864 | Uncharacterized protein | 23.22 | 205 | 0.184 | 1.14 | 1.98E-02 | 3.15E-02 | no | no | 0 | 24 | 29 | 29 | 23 | 25 | 26 | 0.033230682 | 0.035565528 | 0.039132701 | 0.027170199 | 0.029125307 | 0.030066387 |
| NITMOv2_4428 | A0A0K2GIL4 | Uncharacterized protein | 54.643 | 486 | -0.162 | -1.12 | 5.43E-02 | 7.54E-02 | no | no | 0 | 64 | 50 | 64 | 67 | 70 | 65 | 0.033051513 | 0.033244422 | 0.033946578 | 0.041839945 | 0.038400783 | 0.035726873 |
| apgM | A0A0K2GIS0 | Putative 2,3-bisphosphoglycerate-independent phosphoglycerate mutase | 42.988 | 403 | -0.0423 | -1.03 | 5.31E-01 | 5.72E-01 | no | no | 0 | 38 | 37 | 42 | 34 | 35 | 46 | 0.032990384 | 0.032417271 | 0.034872603 | 0.029431298 | 0.03808694 | 0.036692464 |
| comE | A0A0K2G7V5 | Sulfopyruvate decarboxylase, beta subunit | 21.036 | 192 | 0.25 | 1.19 | 2.22E-02 | 3.49E-02 | no | no | 0 | 12 | 15 | 13 | 18 | 11 | 12 | 0.032874451 | 0.034538922 | 0.036366129 | 0.034745693 | 0.031314444 | 0.032225931 |
| NITMOv2_4574 | A0A0K2GK18 | Uncharacterized protein | 26.637 | 245 | -1.61 | -3.05 | 3.47E-07 | 1.46E-05 | yes | no | 0 | 28 | 26 | 28 | 58 | 61 | 53 | 0.032720576 | 0.033007813 | 0.032372527 | 0.116078601 | 0.106727019 | 0.108387158 |
| NITMOv2_3929 | A0A0K2GHH4 | Putative Toluene tolerance-type auxiliary component of ABC-type transport system | 22.938 | 197 | 0.948 | 1.93 | 1.82E-05 | 1.74E-04 | no | no | 0 | 12 | 10 | 13 | 12 | 11 | 11 | 0.032669987 | 0.032419226 | 0.029740162 | 0.016314854 | 0.016340065 | 0.016101098 |
| anmK | A0A0K2G7A3 | Anhydro-N-acetylmuramic acid kinase | 41.83 | 398 | 0.429 | 1.35 | 2.05E-03 | 4.89E-03 | no | no | 0 | 18 | 19 | 18 | 17 | 15 | 16 | 0.032621506 | 0.032587394 | 0.033287049 | 0.02586122 | 0.024652259 | 0.024139779 |
| NITMOv2_1450 | A0A0K2GBB4 | SnoaL-like domain-containing protein | 33.089 | 292 | 0.374 | 1.30 | 1.24E-02 | 2.13E-02 | no | no | 0 | 22 | 18 | 17 | 9 | 12 | 9 | 0.032606751 | 0.029564285 | 0.032029342 | 0.023591237 | 0.024616524 | 0.020637039 |
| NITMOv2_3007 | A0A0K2GFQ2 | Putative NADH-quinone oxidoreductase, subunit L | 63.175 | 571 | 0.481 | 1.40 | 1.45E-02 | 2.44E-02 | no | no | 0 | 35 | 29 | 28 | 24 | 24 | 21 | 0.032594103 | 0.031576432 | 0.03434728 | 0.024729931 | 0.023452817 | 0.020820447 |
| NITMOv2_4299 | A0A0K2GIB6 | NAD-dependent aldehyde dehydrogenase | 51.016 | 480 | 0.0953 | 1.07 | 2.55E-01 | 2.97E-01 | no | no | 0 | 38 | 37 | 44 | 52 | 42 | 39 | 0.032562485 | 0.032640191 | 0.033611061 | 0.0326534 | 0.031323766 | 0.028262871 |
| msrB | A0A0K2GDQ0 | Peptide-methionine (R)-S-oxide reductase | 15.672 | 139 | 0.404 | 1.32 | 4.27E-03 | 8.76E-03 | no | no | 0 | 10 | 9 | 7 | 7 | 6 | 7 | 0.032539299 | 0.031901035 | 0.031910473 | 0.022990055 | 0.026165544 | 0.021386855 |
| czcA.1 | A0A0K2GDU9 | Cation efflux system protein CzcA | 114.9 | 1055 | 0.453 | 1.37 | 5.26E-04 | 1.77E-03 | no | no | 0 | 78 | 75 | 80 | 65 | 68 | 62 | 0.032514004 | 0.032006629 | 0.035419015 | 0.025767933 | 0.026810322 | 0.025367896 |
| NITMOv2_0066 | A0A0K2G6E3 | Uncharacterized protein | 36.527 | 320 | 1.81 | 3.51 | 1.99E-07 | 1.31E-05 | yes | no | 0 | 20 | 21 | 24 | 5 | 7 | 5 | 0.032480278 | 0.032499399 | 0.030915428 | 0.008202738 | 0.00917138 | 0.008209683 |
| metF | A0A0K2G846 | Methylenetetrahydrofolate reductase | 32.047 | 296 | -0.235 | -1.18 | 9.93E-03 | 1.77E-02 | no | no | 0 | 33 | 39 | 42 | 47 | 39 | 40 | 0.032450768 | 0.036113051 | 0.039278411 | 0.042353764 | 0.041759842 | 0.039947064 |
| NITMOv2_3861 | A0A0K2GH07 | Putative Universal stress protein | 32.437 | 299 | 0.988 | 1.98 | 5.25E-06 | 7.69E-05 | no | no | 0 | 29 | 29 | 29 | 22 | 18 | 23 | 0.032277922 | 0.033541648 | 0.035474615 | 0.01807398 | 0.017753915 | 0.017826035 |
| NITMOv2_0481 | A0A0K2G8K9 | AIRC domain-containing protein | 27.261 | 253 | 0.201 | 1.15 | 2.35E-02 | 3.65E-02 | no | no | 0 | 20 | 20 | 23 | 14 | 15 | 18 | 0.032218901 | 0.034278849 | 0.033049311 | 0.031230403 | 0.029317963 | 0.026696707 |
| fusA.1 | A0A0K2GC98 | Elongation factor G (EF-G) | 75.533 | 700 | 1.97 | 3.92 | 2.94E-07 | 1.40E-05 | yes | no | 0 | 36 | 44 | 53 | 18 | 14 | 14 | 0.032117723 | 0.032829868 | 0.033064649 | 0.007354123 | 0.008155272 | 0.00711193 |
| NITMOv2_0519 | A0A0K2G7Q4 | Uncharacterized protein | 48.71 | 438 | -0.0646 | -1.05 | 4.13E-01 | 4.57E-01 | no | no | 0 | 32 | 33 | 33 | 44 | 48 | 41 | 0.032046055 | 0.033381302 | 0.033181601 | 0.040095627 | 0.039809972 | 0.035992995 |
| pdxA | A0A0K2G7I5 | 4-hydroxythreonine-4-phosphate dehydrogenase | 36.594 | 350 | 0.305 | 1.24 | 1.20E-02 | 2.07E-02 | no | no | 0 | 35 | 31 | 32 | 24 | 24 | 22 | 0.031894288 | 0.032440736 | 0.030694946 | 0.030013231 | 0.029030532 | 0.024916567 |
| NITMOv2_1765 | A0A0K2GB59 | Putative Fe(2+)-trafficking protein YggX | 10.395 | 93 | 0.379 | 1.30 | 1.59E-03 | 4.04E-03 | no | no | 0 | 8 | 9 | 9 | 7 | 5 | 6 | 0.031879533 | 0.031091483 | 0.031356392 | 0.038785167 | 0.023862989 | 0.024308802 |
| lpxD | A0A0K2GBE5 | UDP-3-O-acylglucosamine N-acyltransferase | 38.275 | 363 | 0.45 | 1.37 | 1.67E-04 | 7.81E-04 | no | no | 0 | 12 | 14 | 15 | 9 | 14 | 13 | 0.031879533 | 0.031664427 | 0.037573988 | 0.020246233 | 0.027107075 | 0.022783995 |
| degP | A0A0K2G813 | Serine protease Do | 49.087 | 469 | 0.629 | 1.55 | 7.15E-05 | 4.48E-04 | no | no | 0 | 28 | 31 | 26 | 19 | 21 | 19 | 0.031816297 | 0.032779027 | 0.030167706 | 0.018207247 | 0.0191196 | 0.018364213 |
| NITMOv2_3627 | A0A0K2GGQ6 | Uncharacterized protein | 39.015 | 332 | -0.986 | -1.98 | 1.31E-05 | 1.42E-04 | no | no | 0 | 25 | 26 | 34 | 57 | 63 | 52 | 0.031795218 | 0.02809575 | 0.031502102 | 0.065974616 | 0.066374812 | 0.060932938 |
| greA | A0A0K2GFI0 | Transcription elongation factor GreA | 17.604 | 158 | 0.366 | 1.29 | 2.09E-03 | 5.00E-03 | no | no | 0 | 9 | 9 | 8 | 8 | 7 | 7 | 0.031786786 | 0.031068018 | 0.029880121 | 0.028135645 | 0.028137166 | 0.027063524 |
| NITMOv2_1172 | A0A0K2G9H5 | Uncharacterized protein | 50.253 | 457 | 0.346 | 1.27 | 2.62E-03 | 5.95E-03 | no | no | 0 | 30 | 35 | 40 | 33 | 33 | 36 | 0.031649774 | 0.0317446 | 0.031749425 | 0.029857752 | 0.029366127 | 0.025887553 |
| NITMOv2_1384 | A0A0K2GA10 | Ferredoxin--NAD(+) reductase | 56.245 | 519 | 0.274 | 1.21 | 3.28E-03 | 7.11E-03 | no | no | 0 | 44 | 46 | 48 | 30 | 38 | 38 | 0.031546489 | 0.030199803 | 0.030746711 | 0.024657374 | 0.026727977 | 0.02480868 |
| NITMOv2_4119 | A0A0K2GHU6 | Putative Ubiquinone/menaquinone biosynthesis methyltransferase UbiE | 31.054 | 286 | 0.531 | 1.44 | 1.39E-03 | 3.64E-03 | no | no | 0 | 47 | 40 | 42 | 25 | 25 | 26 | 0.031527518 | 0.029055871 | 0.032865257 | 0.024743257 | 0.023802395 | 0.020559719 |
| NITMOv2_0803 | A0A0K2G8F0 | Putative Proteasome, beta subunit | 27.937 | 257 | 0.117 | 1.08 | 1.37E-01 | 1.70E-01 | no | no | 0 | 20 | 18 | 17 | 18 | 19 | 22 | 0.031405261 | 0.034572165 | 0.035323153 | 0.034054185 | 0.031563033 | 0.033358747 |
| NITMOv2_1830 | A0A0K2GBC1 | Putative Vitamin B12 transporter BtuB | 74.08 | 674 | -0.131 | -1.10 | 8.93E-02 | 1.17E-01 | no | no | 0 | 56 | 58 | 58 | 68 | 62 | 63 | 0.031367319 | 0.031973386 | 0.033793199 | 0.037550225 | 0.039873673 | 0.036694262 |
| NITMOv2_3000 | A0A0K2GEN5 | Uncharacterized protein | 14.868 | 136 | 1.01 | 2.01 | 5.69E-05 | 3.86E-04 | yes | no | 0 | 10 | 9 | 8 | 7 | 7 | 7 | 0.031219768 | 0.030545915 | 0.027228583 | 0.011607416 | 0.013828228 | 0.014860214 |
| NITMOv2_3595 | A0A0K2GHA1 | Putative N-carbamoyl-D-amino acid hydrolase | 29.363 | 263 | 0.328 | 1.26 | 3.70E-02 | 5.42E-02 | no | no | 0 | 17 | 16 | 20 | 16 | 19 | 18 | 0.031141776 | 0.031685937 | 0.030802311 | 0.025724992 | 0.023849006 | 0.022334465 |
| clpB | A0A0K2GCG7 | Chaperone protein ClpB | 96.166 | 865 | 0.106 | 1.08 | 2.93E-01 | 3.35E-01 | no | no | 0 | 62 | 63 | 73 | 67 | 69 | 60 | 0.030975254 | 0.029255326 | 0.033457683 | 0.033428814 | 0.032993973 | 0.027816937 |
| NITMOv2_0804 | A0A0K2G8P9 | Putative Proteasome, alpha subunit | 29.941 | 272 | 0.236 | 1.18 | 1.59E-02 | 2.62E-02 | no | no | 0 | 20 | 14 | 20 | 19 | 18 | 15 | 0.030893047 | 0.029748096 | 0.028737448 | 0.028770885 | 0.026496478 | 0.027701857 |
| NITMOv2_4307 | A0A0K2GI88 | Uncharacterized protein | 17.342 | 167 | 0.992 | 1.99 | 1.08E-05 | 1.26E-04 | no | no | 0 | 12 | 11 | 11 | 8 | 8 | 12 | 0.030888831 | 0.031253784 | 0.031709164 | 0.019334094 | 0.020886136 | 0.020029274 |
| aroQ | A0A0K2GD24 | 3-dehydroquinate dehydratase | 17.934 | 165 | -0.252 | -1.19 | 1.54E-01 | 1.90E-01 | no | no | 0 | 19 | 18 | 19 | 20 | 17 | 16 | 0.030852997 | 0.03345952 | 0.040154587 | 0.049911489 | 0.051380236 | 0.042669419 |
| NITMOv2_3948 | A0A0K2GH95 | Uncharacterized protein | 11.159 | 103 | 1.42 | 2.68 | 1.35E-02 | 2.28E-02 | yes | no | 0 | 4 | 4 | 5 | 2 | 3 | 4 | 0.030825595 | 0.021783592 | 0.107999812 | 0.021553732 | 0.020556755 | 0.019345987 |
| prpD | A0A0K2G6A2 | 2-methylcitrate dehydratase | 51.748 | 475 | 1.49 | 2.81 | 1.20E-06 | 3.14E-05 | yes | no | 0 | 41 | 44 | 38 | 19 | 19 | 20 | 0.030815055 | 0.032204128 | 0.030535815 | 0.012128786 | 0.011459331 | 0.011919926 |
| NITMOv2_3689 | A0A0K2GGL4 | Putative Carbohydrate/purine kinase, PfkB family | 33.762 | 304 | 0.0599 | 1.04 | 4.25E-01 | 4.69E-01 | no | no | 0 | 22 | 22 | 24 | 26 | 29 | 25 | 0.030718093 | 0.029585795 | 0.035093085 | 0.034114896 | 0.03350358 | 0.031206396 |
| NITMOv2_2479 | A0A0K2GE69 | Uncharacterized protein | 34.806 | 330 | 0.479 | 1.39 | 1.58E-04 | 7.58E-04 | no | no | 0 | 28 | 29 | 24 | 22 | 23 | 19 | 0.030446177 | 0.032409449 | 0.030771635 | 0.025237826 | 0.025081075 | 0.024555145 |
| phnW | A0A0K2G858 | 2-aminoethylphosphonate aminotransferase | 39.633 | 360 | 0.0836 | 1.06 | 4.87E-01 | 5.30E-01 | no | no | 0 | 18 | 21 | 27 | 35 | 37 | 34 | 0.030441961 | 0.029687478 | 0.030307664 | 0.029970289 | 0.028081233 | 0.026306515 |
| rpsT | A0A0K2GK23 | 30S ribosomal protein S20 | 10.214 | 92 | -0.899 | -1.86 | 1.58E-03 | 4.03E-03 | no | no | 0 | 3 | 3 | 3 | 5 | 5 | 3 | 0.030420882 | 0.032540463 | 0.034351114 | 0.070917344 | 0.070454779 | 0.052125739 |
| wrbA | A0A0K2GHB0 | NAD(P)H:quinone oxidoreductase WrbA | 22.093 | 209 | 0.645 | 1.56 | 1.79E-03 | 4.40E-03 | no | no | 0 | 9 | 10 | 9 | 7 | 10 | 7 | 0.030336567 | 0.025240809 | 0.022669397 | 0.020289175 | 0.020701248 | 0.018630335 |
| NITMOv2_4517 | A0A0K2GIX5 | Putative Sulfurtransferase TusA | 9.1077 | 81 | -0.39 | -1.31 | 7.53E-03 | 1.40E-02 | no | no | 0 | 13 | 12 | 10 | 9 | 6 | 10 | 0.030201663 | 0.031488437 | 0.029166909 | 0.038443115 | 0.036884391 | 0.034315348 |
| NITMOv2_3300 | A0A0K2GFG0 | Uncharacterized protein | 44.336 | 413 | 0.377 | 1.30 | 2.52E-03 | 5.75E-03 | no | no | 0 | 19 | 16 | 19 | 13 | 11 | 12 | 0.030197447 | 0.030794256 | 0.026442517 | 0.02329805 | 0.023502535 | 0.02339176 |
| NITMOv2_0275 | A0A0K2G769 | Glucose-6-phosphate isomerase | 61.421 | 564 | 0.592 | 1.51 | 8.41E-05 | 5.03E-04 | no | no | 0 | 47 | 44 | 54 | 57 | 55 | 43 | 0.030182692 | 0.032495488 | 0.032865257 | 0.025054214 | 0.024111578 | 0.022388409 |
| dmpA | A0A0K2G6X3 | Peptidase S58 | 45.629 | 436 | 0.493 | 1.41 | 1.78E-03 | 4.38E-03 | no | no | 0 | 9 | 12 | 6 | 10 | 10 | 9 | 0.030167937 | 0.033140783 | 0.02705028 | 0.030007308 | 0.027992673 | 0.02540206 |
| acsA | A0A0K2GF43 | Acetyl-CoA synthetase | 70.26 | 629 | 0.198 | 1.15 | 4.33E-02 | 6.19E-02 | no | no | 0 | 38 | 35 | 45 | 42 | 40 | 41 | 0.030161613 | 0.029941685 | 0.032637106 | 0.030430801 | 0.031303568 | 0.029762504 |
| NITMOv2_3827 | A0A0K2GGZ9 | Uncharacterized protein | 7.9872 | 70 | 2.02 | 4.06 | 1.05E-05 | 1.24E-04 | yes | no | 0 | 13 | 9 | 11 | 2 | 2 | 3 | 0.030083622 | 0.024315886 | 0.03431277 | 0.012205489 | 0.009420435 | 0.007477667 |
| fur | A0A0K2G8T7 | Ferric uptake regulation protein | 16.721 | 143 | 0.352 | 1.28 | 3.81E-02 | 5.54E-02 | no | no | 0 | 11 | 16 | 14 | 14 | 13 | 11 | 0.030011954 | 0.030207624 | 0.029882038 | 0.030457454 | 0.025175849 | 0.022050362 |
| NITMOv2_1066 | A0A0K2GA87 | Uncharacterized protein | 41.783 | 383 | 0.535 | 1.45 | 1.44E-04 | 7.13E-04 | no | no | 0 | 34 | 34 | 36 | 31 | 27 | 31 | 0.029957149 | 0.029040227 | 0.028490125 | 0.018688489 | 0.022307754 | 0.021399442 |
| gmhA.1 | A0A0K2GJ66 | Phosphoheptose isomerase | 20.402 | 192 | -0.129 | -1.09 | 2.31E-01 | 2.72E-01 | no | no | 0 | 13 | 14 | 15 | 13 | 15 | 14 | 0.029933963 | 0.030170471 | 0.031816529 | 0.036633644 | 0.036216308 | 0.034646202 |
| NITMOv2_3282 | A0A0K2GFD6 | Antitoxin | 10.441 | 94 | 0.887 | 1.85 | 8.79E-05 | 5.18E-04 | no | no | 0 | 5 | 5 | 4 | 2 | 4 | 1 | 0.029790627 | 0.03096829 | 0.027855519 | 0.017453547 | 0.018322562 | 0.015900967 |
| NITMOv2_0373 | A0A0K2G7G1 | Trypsin-like serine peptidase, S1C | 34.314 | 331 | 0.183 | 1.14 | 3.36E-02 | 4.97E-02 | no | no | 0 | 19 | 20 | 21 | 18 | 15 | 16 | 0.02976744 | 0.035696542 | 0.036070875 | 0.024607029 | 0.021749982 | 0.021232217 |
| NITMOv2_1111 | A0A0K2GAC3 | Uncharacterized protein | 13.77 | 131 | 0.824 | 1.77 | 2.71E-03 | 6.11E-03 | no | no | 0 | 12 | 11 | 7 | 8 | 6 | 6 | 0.029729498 | 0.030594801 | 0.023610759 | 0.018168747 | 0.01712312 | 0.009524469 |
| NITMOv2_1154 | A0A0K2G9G9 | Uncharacterized protein | 40.908 | 364 | 0.308 | 1.24 | 3.93E-03 | 8.19E-03 | no | no | 0 | 41 | 41 | 47 | 32 | 35 | 36 | 0.029723175 | 0.028920946 | 0.03039394 | 0.02278275 | 0.022298432 | 0.021958658 |
| NITMOv2_0139 | A0A0K2G6U5 | Uncharacterized protein | 33.042 | 309 | 0.2 | 1.15 | 1.01E-01 | 1.30E-01 | no | no | 0 | 25 | 28 | 26 | 25 | 22 | 22 | 0.029497632 | 0.030207624 | 0.031308461 | 0.025485111 | 0.023354935 | 0.021672756 |
| NITMOv2_0724 | A0A0K2G898 | Putative ABC transporter, auxiliary component, ATP-dependent toluene efflux transporter | 25.152 | 220 | -1.24 | -2.36 | 8.56E-07 | 2.82E-05 | yes | no | 0 | 18 | 20 | 21 | 38 | 36 | 33 | 0.029267873 | 0.030195892 | 0.030397774 | 0.077607352 | 0.079501865 | 0.07315297 |
| NITMOv2_2974 | A0A0K2GEU4 | Uncharacterized protein | 27.837 | 254 | 0.858 | 1.81 | 7.66E-05 | 4.67E-04 | no | no | 0 | 19 | 23 | 33 | 19 | 18 | 15 | 0.029215176 | 0.032734052 | 0.033311973 | 0.020070025 | 0.019464517 | 0.017221507 |
| ilvE | A0A0K2GA12 | Branched-chain-amino-acid aminotransferase | 33.793 | 304 | 0.129 | 1.09 | 5.56E-02 | 7.69E-02 | no | no | 0 | 38 | 45 | 48 | 33 | 35 | 35 | 0.029187774 | 0.02981067 | 0.032199976 | 0.024439704 | 0.025606219 | 0.026229196 |
| devR.4 | A0A0K2GH63 | Transcriptional regulatory protein DevR (DosR) | 23.595 | 219 | 0.307 | 1.24 | 7.14E-03 | 1.34E-02 | no | no | 0 | 17 | 19 | 19 | 12 | 16 | 16 | 0.029116106 | 0.029867378 | 0.026406089 | 0.020607535 | 0.023821039 | 0.022115094 |
| NITMOv2_3694 | A0A0K2GGL8 | Putative Tol-Pal system protein YbgF (Modular protein) | 58.124 | 529 | 0.207 | 1.15 | 3.03E-02 | 4.56E-02 | no | no | 0 | 27 | 27 | 24 | 27 | 25 | 24 | 0.029088704 | 0.02794978 | 0.026611233 | 0.027475233 | 0.029036747 | 0.026970022 |
| NITMOv2_0083 | A0A0K2G6G9 | Dyp-type peroxidase | 32.334 | 294 | -0.0366 | -1.03 | 7.38E-01 | 7.70E-01 | no | no | 0 | 24 | 26 | 25 | 32 | 28 | 28 | 0.029057086 | 0.030606534 | 0.028162277 | 0.034433256 | 0.032422218 | 0.03114526 |
| NITMOv2_2970 | A0A0K2GEI7 | Uncharacterized protein | 36.776 | 344 | 2.03 | 4.08 | 1.36E-04 | 6.85E-04 | yes | no | 0 | 36 | 29 | 32 | 12 | 9 | 21 | 0.029046546 | 0.027996023 | 0.02933946 | 0.005436261 | 0.004939774 | 0.010300358 |
| mreB | A0A0K2G8L5 | Cell shape-determining protein MreB | 37.672 | 349 | -0.86 | -1.82 | 4.95E-05 | 3.48E-04 | no | no | 0 | 24 | 28 | 31 | 50 | 54 | 47 | 0.029025468 | 0.028750822 | 0.029651969 | 0.059185398 | 0.054414575 | 0.053447359 |
| NITMOv2_4603 | A0A0K2GJ59 | Putative Peptidase M16 | 59.822 | 543 | 0.311 | 1.24 | 2.76E-02 | 4.19E-02 | no | no | 0 | 41 | 41 | 40 | 36 | 38 | 41 | 0.028911642 | 0.029478246 | 0.029153488 | 0.026839993 | 0.02572896 | 0.023413338 |
| NITMOv2_0790 | A0A0K2G8G0 | mRNA interferase | 12.437 | 112 | 0.244 | 1.18 | 3.13E-02 | 4.68E-02 | no | no | 0 | 9 | 8 | 12 | 7 | 9 | 10 | 0.028852622 | 0.027065234 | 0.031456088 | 0.014622214 | 0.014909901 | 0.014658645 |
| NITMOv2_0457 | A0A0K2G7F8 | YCII domain-containing protein | 12.913 | 119 | 0.598 | 1.51 | 2.26E-03 | 5.29E-03 | no | no | 0 | 8 | 8 | 8 | 6 | 7 | 5 | 0.028787277 | 0.028342136 | 0.026379248 | 0.021023625 | 0.019922853 | 0.017383877 |
| forE | A0A0K2G817 | 2-oxoglutarate:ferredoxin oxidoreductase, epsilon subunit | 11.573 | 107 | -1.46 | -2.75 | 2.35E-06 | 4.65E-05 | yes | no | 0 | 8 | 7 | 4 | 17 | 18 | 15 | 0.028669236 | 0.027153228 | 0.026956336 | 0.097807683 | 0.091984139 | 0.090801531 |

| | | | | | | | | | | | | | | | | | | | | | | |
|---|---|---|---|---|---|---|---|---|---|---|---|---|---|---|---|---|---|---|---|---|---|---|
| secY | A0A0K2G9W1 | Protein translocase subunit SecY | 47.636 | 439 | -0.0282 | -1.02 | 6.28E-01 | 6.68E-01 | no | no | 0 | 13 | 17 | 18 | 21 | 18 | 20 | 0.028589137 | 0.029315944 | 0.029030785 | 0.030312341 | 0.030506529 | 0.029246444 |
| NITMOv2_3830 | A0A0K2GH90 | Putative Universal stress protein | 18.31 | 169 | 1.07 | 2.10 | 2.60E-05 | 2.24E-04 | yes | no | 0 | 14 | 13 | 14 | 3 | 6 | 5 | 0.028555411 | 0.029493889 | 0.032435796 | 0.016821269 | 0.014567314 | 0.013472064 |
| grpE | A0A0K2GD65 | Protein GrpE | 21.977 | 204 | -0.0919 | -1.07 | 3.10E-01 | 3.52E-01 | no | no | 0 | 11 | 12 | 11 | 10 | 8 | 11 | 0.028525901 | 0.028412531 | 0.03420157 | 0.035647467 | 0.037142302 | 0.0321594 |
| NITMOv2_1696 | A0A0K2GAX1 | Uncharacterized protein | 29.051 | 260 | 0.233 | 1.18 | 5.34E-03 | 1.05E-02 | no | no | 0 | 24 | 25 | 27 | 18 | 26 | 29 | 0.028416291 | 0.029202529 | 0.030000906 | 0.023098149 | 0.024335308 | 0.024177539 |
| tolB | A0A0K2GGW8 | Protein TolB | 48.034 | 442 | -0.118 | -1.09 | 1.49E-01 | 1.84E-01 | no | no | 0 | 26 | 31 | 30 | 31 | 34 | 32 | 0.028405752 | 0.027018303 | 0.0263294 | 0.025491034 | 0.029378557 | 0.030141908 |
| rpoN | A0A0K2G9L8 | RNA polymerase, sigma-54 (Sigma N) factor | 56.605 | 502 | -0.0313 | -1.02 | 7.22E-01 | 7.56E-01 | no | no | 0 | 35 | 36 | 44 | 58 | 47 | 52 | 0.028355163 | 0.024231802 | 0.03024823 | 0.029560122 | 0.027740977 | 0.027523843 |
| ilvI | A0A0K2G8J7 | Acetolactate synthase | 64.849 | 591 | -1.83 | -3.56 | 5.97E-08 | 7.05E-06 | yes | no | 0 | 17 | 20 | 24 | 62 | 58 | 61 | 0.028264524 | 0.019192831 | 0.027270763 | 0.049227384 | 0.052014138 | 0.050368975 |
| NITMOv2_0848 | A0A0K2G8J3 | zf-RING_7 domain-containing protein | 29.092 | 257 | 0.105 | 1.08 | 2.26E-01 | 2.67E-01 | no | no | 0 | 28 | 26 | 30 | 27 | 30 | 27 | 0.028138051 | 0.028232631 | 0.028557228 | 0.034048262 | 0.031911057 | 0.032878649 |
| NITMOv2_4593 | A0A0K2GJ51 | Putative Peptidase M20 | 49.678 | 455 | 0.342 | 1.27 | 2.66E-03 | 6.01E-03 | no | no | 0 | 31 | 30 | 35 | 29 | 33 | 30 | 0.028095894 | 0.029400028 | 0.028885075 | 0.023838522 | 0.023569343 | 0.022571817 |
| mmgB | A0A0K2G6D3 | Putative 3-hydroxybutyryl-CoA dehydrogenase | 33.317 | 311 | 3.08 | 8.46 | 1.41E-06 | 3.47E-05 | yes | no | 0 | 19 | 24 | 23 | 4 | 5 | 4 | 0.02804952 | 0.029591661 | 0.02841727 | 0.002093774 | 0.002506243 | 0.001591697 |
| pycB | A0A0K2G978 | Pyruvate carboxylase subunit B | 71.477 | 651 | 0.078 | 1.06 | 2.14E-01 | 2.55E-01 | no | no | 0 | 66 | 68 | 67 | 59 | 62 | 63 | 0.028043197 | 0.028269784 | 0.028208291 | 0.026472768 | 0.025248872 | 0.026590618 |
| folK | A0A0K2G7E5 | 2-amino-4-hydroxy-6-hydroxymethyldihydropteridine pyrophosphokinase | 20.982 | 184 | 0.631 | 1.55 | 2.68E-04 | 1.08E-03 | no | no | 0 | 5 | 6 | 5 | 3 | 5 | 3 | 0.028007363 | 0.03056547 | 0.034795913 | 0.009400957 | 0.00864375 | 0.025382281 |
| rpsK | A0A0K2G9W5 | 30S ribosomal protein S11 | 13.7 | 127 | 0.374 | 1.30 | 2.29E-01 | 2.70E-01 | no | no | 0 | 4 | 3 | 4 | 2 | 2 | 3 | 0.027948342 | 0.026738675 | 0.024745763 | 0.014816339 | 0.026101843 | 0.028289843 |
| NITMOv2_0062 | A0A0K2G6M8 | Putative Fumarylacetoacetase | 36.286 | 332 | 1.86 | 3.63 | 3.35E-07 | 1.46E-05 | yes | no | 0 | 47 | 58 | 60 | 17 | 18 | 14 | 0.027927264 | 0.025805931 | 0.028793048 | 0.006512023 | 0.007063967 | 0.006726772 |
| pmbA | A0A0K2GA68 | Peptidase U62 | 49.452 | 461 | 0.271 | 1.21 | 1.25E-02 | 2.16E-02 | no | no | 0 | 34 | 32 | 38 | 34 | 39 | 31 | 0.027815546 | 0.027571692 | 0.025336272 | 0.022436256 | 0.026089414 | 0.025029849 |
| NITMOv2_1023 | A0A0K2G948 | Uncharacterized protein | 32.789 | 298 | 0.516 | 1.43 | 6.59E-04 | 2.08E-03 | no | no | 0 | 15 | 16 | 14 | 13 | 10 | 13 | 0.027724907 | 0.028148547 | 0.027098211 | 0.022311873 | 0.021358455 | 0.021509127 |
| rfaEb | A0A0K2GK64 | D-beta-D-heptose 1-phosphate adenylyltransferase | 17.09 | 160 | 1.04 | 2.06 | 3.59E-05 | 2.79E-04 | yes | no | 0 | 7 | 7 | 7 | 3 | 4 | 5 | 0.027714368 | 0.02889748 | 0.026436765 | 0.014859281 | 0.015244409 | 0.013630659 |
| apt | A0A0K2G9E4 | Adenine phosphoribosyltransferase | 19.151 | 174 | -0.283 | -1.22 | 3.44E-02 | 5.08E-02 | no | no | 0 | 12 | 9 | 14 | 26 | 25 | 27 | 0.027646916 | 0.028359734 | 0.029974065 | 0.031867124 | 0.029397201 | 0.031984982 |
| NITMOv2_3746 | A0A0K2GHM8 | Uncharacterized protein | 41.687 | 416 | 0.2 | 1.15 | 4.92E-02 | 6.92E-02 | no | no | 0 | 2 | 3 | 2 | 3 | 2 | 1 | 0.027632161 | 0.029298345 | 0.026166435 | 0.026321732 | 0.028739994 | 0.024213502 |
| NITMOv2_4549 | A0A0K2GJ00 | Putative VacJ-like lipoprotein | 32.996 | 299 | 0.448 | 1.36 | 9.67E-04 | 2.74E-03 | no | no | 0 | 14 | 18 | 17 | 16 | 19 | 22 | 0.02761319 | 0.028375378 | 0.028737448 | 0.028639099 | 0.027060465 | 0.025675375 |
| fabD | A0A0K2G8B4 | Malonyl CoA-acyl carrier protein transacylase | 32.785 | 316 | -0.66 | -1.58 | 7.50E-04 | 2.30E-03 | no | no | 0 | 21 | 18 | 19 | 30 | 33 | 26 | 0.027522551 | 0.026363231 | 0.027335949 | 0.050219484 | 0.050535033 | 0.043674569 |
| NITMOv2_4586 | A0A0K2GJD1 | Uncharacterized protein | 35.559 | 327 | 0.0744 | 1.05 | 5.02E-01 | 5.45E-01 | no | no | 0 | 21 | 23 | 22 | 19 | 17 | 23 | 0.02751412 | 0.027037858 | 0.023555159 | 0.025529533 | 0.02388474 | 0.025186286 |
| NITMOv2_4842 | A0A0K2GJS9 | Uncharacterized protein | 55.31 | 490 | 0.572 | 1.49 | 1.28E-04 | 6.57E-04 | no | no | 0 | 43 | 44 | 51 | 43 | 41 | 41 | 0.02751412 | 0.026222439 | 0.027057949 | 0.017742293 | 0.017213233 | 0.019178762 |
| ppnK | A0A0K2GC27 | NAD kinase | 30.916 | 286 | -0.00881 | -1.01 | 9.48E-01 | 9.54E-01 | no | no | 0 | 8 | 13 | 18 | 14 | 19 | 12 | 0.027499365 | 0.030205669 | 0.029262771 | 0.018578914 | 0.018386263 | 0.016415409 |
| NITMOv2_1454 | A0A0K2GA85 | Uncharacterized protein | 10.655 | 93 | 0.354 | 1.28 | 5.54E-03 | 1.08E-02 | no | no | 0 | 7 | 7 | 6 | 7 | 6 | 6 | 0.027463531 | 0.030235001 | 0.02834058 | 0.022097165 | 0.021468766 | 0.020494987 |
| gyrA | A0A0K2G694 | DNA gyrase subunit A | 91.943 | 824 | -0.319 | -1.25 | 1.01E-03 | 2.83E-03 | no | no | 0 | 65 | 58 | 69 | 72 | 74 | 79 | 0.027366568 | 0.026537265 | 0.029743997 | 0.037005311 | 0.038156855 | 0.035660342 |
| proA | A0A0K2GAX5 | Gamma-glutamyl phosphate reductase | 49.114 | 454 | 0.16 | 1.12 | 5.37E-02 | 7.47E-02 | no | no | 0 | 33 | 34 | 34 | 39 | 38 | 37 | 0.02730544 | 0.027552138 | 0.027424141 | 0.028873057 | 0.027323037 | 0.025725722 |
| NITMOv2_4501 | A0A0K2GIV7 | Uncharacterized protein | 16.287 | 145 | 0.248 | 1.19 | 1.28E-01 | 1.61E-01 | no | no | 0 | 10 | 13 | 13 | 14 | 11 | 12 | 0.027259067 | 0.026977239 | 0.025689043 | 0.01553154 | 0.022808039 | 0.025506351 |
| secF | A0A0K2GGR1 | Protein-export membrane protein SecF | 33.179 | 305 | 0.393 | 1.31 | 2.32E-03 | 5.40E-03 | no | no | 0 | 19 | 20 | 18 | 15 | 14 | 15 | 0.027244311 | 0.027393748 | 0.025598933 | 0.017885925 | 0.016843457 | 0.019398133 |
| NITMOv2_3054 | A0A0K2GES8 | RHH_3 domain-containing protein | 11.182 | 92 | 0.742 | 1.67 | 3.87E-04 | 1.39E-03 | no | no | 0 | 4 | 4 | 4 | 3 | 2 | 4 | 0.027219017 | 0.025545857 | 0.024337392 | 0.020917011 | 0.017859565 | 0.019052894 |
| ribE | A0A0K2G9T0 | 6,7-dimethyl-8-ribityllumazine synthase | 19.339 | 178 | 0.4 | 1.32 | 2.96E-03 | 6.55E-03 | no | no | 0 | 16 | 17 | 18 | 16 | 16 | 18 | 0.027183183 | 0.02544613 | 0.027140391 | 0.024457473 | 0.022969622 | 0.020390696 |
| leuA | A0A0K2G8K3 | 2-isopropylmalate synthase | 56.276 | 514 | 0.166 | 1.12 | 4.01E-02 | 5.78E-02 | no | no | 0 | 27 | 33 | 34 | 25 | 33 | 31 | 0.027155781 | 0.028703892 | 0.029155405 | 0.024355302 | 0.023687423 | 0.021383259 |
| NITMOv2_4588 | A0A0K2GJ48 | Putative RNA polymerase sigma-E factor (Sigma-24) | 23.537 | 207 | 0.399 | 1.32 | 4.64E-02 | 6.59E-02 | no | no | 0 | 20 | 17 | 20 | 13 | 14 | 13 | 0.027084113 | 0.019096819 | 0.030405443 | 0.016757597 | 0.017273827 | 0.017609542 |
| NITMOv2_0918 | A0A0K2G9S9 | Metal-dependent carboxypeptidase | 58.995 | 510 | 0.193 | 1.14 | 2.01E-02 | 3.19E-02 | no | no | 0 | 42 | 44 | 50 | 46 | 39 | 38 | 0.027082005 | 0.02655682 | 0.028225546 | 0.027820246 | 0.024844915 | 0.022837939 |
| NITMOv2_0987 | A0A0K2G992 | Sigma-54 dependent DNA-binding response regulator | 48.739 | 446 | 2.36 | 5.13 | 2.10E-07 | 1.33E-05 | yes | no | 0 | 37 | 51 | 48 | 8 | 8 | 6 | 0.026993474 | 0.030860741 | 0.02727268 | 0.004544556 | 0.003793623 | 0.00367536 |
| NITMOv2_4832 | A0A0K2GJR9 | Putative Lipoprotein, TraT related | 23.926 | 225 | 0.507 | 1.42 | 5.52E-04 | 1.84E-03 | no | no | 0 | 16 | 14 | 15 | 13 | 14 | 13 | 0.026902835 | 0.02804882 | 0.023135285 | 0.017734889 | 0.020645315 | 0.020110189 |
| NITMOv2_2973 | A0A0K2GEK1 | Uncharacterized protein | 35.079 | 334 | 0.958 | 1.94 | 9.03E-06 | 1.10E-04 | no | no | 0 | 12 | 12 | 12 | 8 | 11 | 8 | 0.026812197 | 0.026210707 | 0.025006507 | 0.007690992 | 0.010473209 | 0.012589187 |
| rpmE | A0A0K2G8I7 | 50S ribosomal protein L31 | 7.6498 | 68 | 0.661 | 1.58 | 2.98E-01 | 3.41E-01 | no | no | 0 | 6 | 6 | 7 | 3 | 7 | 6 | 0.026784794 | 0.031858015 | 0.029684562 | 0.015931341 | 0.030337178 | 0.038274811 |
| katA | A0A0K2G6D1 | Catalase | 54.981 | 486 | -3.96 | -15.56 | 4.90E-02 | 6.91E-02 | no | yes | 4 | 1 | 4 | 3 | 1 | 5 | 4 | 0.026782686 | 0.025676872 | 0.024889556 | 7.612661E-05 | 0.02388474 | 0.021054203 |
| NITMOv2_0577 | A0A0K2G7T1 | HEPN domain-containing protein | 14.738 | 130 | 0.587 | 1.50 | 3.64E-04 | 1.33E-03 | no | no | 0 | 8 | 9 | 9 | 4 | 6 | 6 | 0.026736313 | 0.028799708 | 0.026973591 | 0.014042354 | 0.016851226 | 0.020588489 |
| ppiC | A0A0K2GG93 | Peptidyl-prolyl cis-trans isomerase | 20.865 | 188 | 0.416 | 1.33 | 2.72E-03 | 6.11E-03 | no | no | 0 | 10 | 14 | 10 | 6 | 8 | 10 | 0.026700479 | 0.026824715 | 0.023271409 | 0.014875569 | 0.014702174 | 0.016230562 |
| NITMOv2_1394 | A0A0K2GA19 | HTH cro/C1-type domain-containing protein | 17.364 | 156 | 0.701 | 1.63 | 2.87E-04 | 1.12E-03 | no | no | 0 | 14 | 16 | 16 | 11 | 12 | 12 | 0.026651998 | 0.027683152 | 0.02772323 | 0.01759866 | 0.015381754 | 0.017674274 |
| guaB | A0A0K2G9X3 | Inosine-5-monophosphate dehydrogenase | 52.4 | 488 | -0.196 | -1.15 | 3.78E-02 | 5.51E-02 | no | no | 0 | 45 | 47 | 42 | 55 | 58 | 54 | 0.026350571 | 0.030485297 | 0.028804551 | 0.036636605 | 0.037944001 | 0.033412691 |
| vapC.3 | A0A0K2GGQ0 | Ribonuclease VapC | 15.061 | 131 | 0.195 | 1.14 | 9.21E-02 | 1.20E-01 | no | no | 0 | 19 | 15 | 15 | 6 | 8 | 8 | 0.026346356 | 0.029611216 | 0.030426533 | 0.019622839 | 0.020295737 | 0.016499561 |
| NITMOv2_1043 | A0A0K2G9E5 | Putative Universal stress protein | 30.701 | 281 | 0.453 | 1.37 | 7.33E-04 | 2.27E-03 | no | no | 0 | 11 | 12 | 12 | 19 | 19 | 15 | 0.026335816 | 0.029267058 | 0.027997395 | 0.024923908 | 0.024812288 | 0.024880605 |
| ilvH | A0A0K2G8V2 | Acetolactate synthase | 19.012 | 172 | -0.61 | -1.53 | 7.27E-05 | 4.52E-04 | no | no | 0 | 11 | 9 | 8 | 17 | 14 | 14 | 0.026289443 | 0.026906843 | 0.024523364 | 0.042028 | 0.043245161 | 0.042155157 |
| lysC | A0A0K2GIY5 | Aspartokinase | 43.383 | 406 | -0.173 | -1.13 | 3.48E-02 | 5.13E-02 | no | no | 0 | 30 | 31 | 33 | 40 | 40 | 30 | 0.026264149 | 0.02827174 | 0.028311822 | 0.041348337 | 0.037041312 | 0.033340766 |
| trpF | A0A0K2GCN3 | N-(5-phosphoribosyl)anthranilate isomerase | 21.959 | 211 | 0.159 | 1.12 | 5.36E-02 | 7.47E-02 | no | no | 0 | 9 | 10 | 13 | 13 | 12 | 12 | 0.025971154 | 0.024345218 | 0.027213245 | 0.029046304 | 0.026735745 | 0.026126703 |
| NITMOv2_1737 | A0A0K2GC16 | Uncharacterized protein | 8.7509 | 74 | 0.105 | 1.08 | 2.46E-01 | 2.88E-01 | no | no | 0 | 5 | 4 | 6 | 8 | 8 | 9 | 0.025969046 | 0.023512201 | 0.02431055 | 0.024771391 | 0.027378969 | 0.02406246 |
| NITMOv2_0917 | A0A0K2G8Q2 | Uncharacterized protein | 22.2 | 198 | 0.444 | 1.36 | 6.04E-04 | 1.96E-03 | no | no | 0 | 27 | 28 | 33 | 26 | 26 | 25 | 0.025962722 | 0.024615068 | 0.025418713 | 0.017629756 | 0.018485698 | 0.018700462 |
| lon.1 | A0A0K2GGH6 | Lon protease | 92.659 | 835 | -0.139 | -1.10 | 4.53E-02 | 6.45E-02 | no | no | 0 | 78 | 82 | 96 | 108 | 101 | 99 | 0.025893162 | 0.026816893 | 0.028248553 | 0.034628714 | 0.033354427 | 0.03124056 |
| NITMOv2_3471 | A0A0K2GFY0 | TRAM domain-containing protein | 36.775 | 341 | 0.212 | 1.16 | 3.68E-02 | 5.39E-02 | no | no | 0 | 15 | 14 | 16 | 11 | 11 | 12 | 0.025863652 | 0.027078922 | 0.02618369 | 0.024737334 | 0.021886706 | 0.022686897 |
| NITMOv2_4017 | A0A0K2GHK1 | Response regulator, LuxR family | 23.211 | 214 | 0.46 | 1.38 | 9.36E-04 | 2.69E-03 | no | no | 0 | 19 | 19 | 16 | 11 | 16 | 13 | 0.025791984 | 0.025371823 | 0.025315182 | 0.01912679 | 0.018131459 | 0.019045701 |
| NITMOv2_1932 | A0A0K2GBK6 | Uncharacterized protein | 8.0632 | 68 | 0.605 | 1.52 | 1.17E-04 | 6.20E-04 | no | no | 0 | 4 | 4 | 5 | 4 | 4 | 3 | 0.025743503 | 0.024826256 | 0.026661082 | 0.022744251 | 0.021596168 | 0.022586202 |
| NITMOv2_1541 | A0A0K2GAH6 | Antitoxin | 9.8414 | 89 | 0.382 | 1.30 | 6.96E-04 | 2.17E-03 | no | no | 0 | 6 | 6 | 5 | 8 | 6 | 4 | 0.025728748 | 0.026376919 | 0.024329723 | 0.022856788 | 0.022626259 | 0.020536344 |
| NITMOv2_3603 | A0A0K2GGB6 | Putative Oligoendopeptidase F | 68.375 | 603 | 0.409 | 1.33 | 2.43E-03 | 5.59E-03 | no | no | 0 | 36 | 37 | 43 | 26 | 35 | 32 | 0.02572664 | 0.026523577 | 0.029301115 | 0.023172186 | 0.022250268 | 0.020370917 |
| NITMOv2_1526 | A0A0K2GAG1 | Putative Molybdate ABC transporter, periplasmic binding protein | 31.615 | 294 | 0.923 | 1.90 | 1.28E-05 | 1.39E-04 | no | no | 0 | 19 | 20 | 26 | 19 | 21 | 20 | 0.025701345 | 0.02753845 | 0.02565645 | 0.014983664 | 0.016414641 | 0.015374117 |
| NITMOv2_3711 | A0A0K2GGL1 | Uncharacterized protein | 27.486 | 252 | 0.449 | 1.37 | 1.70E-03 | 4.26E-03 | no | no | 0 | 21 | 22 | 19 | 13 | 16 | 14 | 0.025568549 | 0.026611572 | 0.026147262 | 0.017148514 | 0.017242753 | 0.017280845 |
| proA.1 | A0A0K2GFW8 | Gamma-glutamyl phosphate reductase | 46.744 | 437 | -0.0739 | -1.05 | 3.70E-01 | 4.13E-01 | no | no | 0 | 35 | 32 | 37 | 45 | 39 | 39 | 0.025541147 | 0.025428531 | 0.024975831 | 0.032249156 | 0.030088589 | 0.027581383 |
| ribE.1 | A0A0K2GFF9 | Riboflavin synthase | 23.194 | 217 | 0.348 | 1.27 | 1.72E-02 | 2.80E-02 | no | no | 0 | 11 | 12 | 13 | 13 | 14 | 12 | 0.025410458 | 0.025698382 | 0.024772604 | 0.025831605 | 0.024882204 | 0.01506538 |
| argJ | A0A0K2GBD1 | Arginine biosynthesis bifunctional protein ArgJ | 42.042 | 401 | 0.101 | 1.07 | 2.20E-01 | 2.62E-01 | no | no | 0 | 26 | 24 | 29 | 24 | 23 | 22 | 0.025393595 | 0.023539577 | 0.023154457 | 0.024306437 | 0.023606631 | 0.021627803 |
| NITMOv2_0845 | A0A0K2G8L1 | Putative HIT-like protein | 12.699 | 114 | 0.15 | 1.11 | 1.09E-01 | 1.40E-01 | no | no | 0 | 6 | 9 | 9 | 5 | 6 | 6 | 0.02538938 | 0.027839588 | 0.030156202 | 0.031727934 | 0.030828141 | 0.028503819 |
| NITMOv2_2093 | A0A0K2GC31 | Uncharacterized protein | 13.411 | 121 | 0.162 | 1.12 | 2.41E-01 | 2.82E-01 | no | no | 0 | 7 | 6 | 7 | 8 | 6 | 5 | 0.025372517 | 0.026564641 | 0.027383879 | 0.028559138 | 0.027972475 | 0.021374268 |
| NITMOv2_0146 | A0A0K2G6I7 | CbiX domain-containing protein | 18.608 | 167 | 0.0209 | 1.01 | 8.16E-01 | 8.38E-01 | no | no | 0 | 7 | 6 | 6 | 8 | 6 | 5 | 0.025353546 | 0.027622534 | 0.027811423 | 0.029398721 | 0.029024318 | 0.02462707 |
| NITMOv2_0573 | A0A0K2G829 | Putative Cytochrome c | 22.657 | 206 | 0.437 | 1.35 | 6.84E-03 | 1.29E-02 | no | no | 0 | 9 | 11 | 8 | 9 | 8 | 6 | 0.025343006 | 0.02633781 | 0.018922159 | 0.020305463 | 0.018098832 | 0.015929377 |
| NITMOv2_3878 | A0A0K2GH27 | HTH tetR-type domain-containing protein | 23.188 | 209 | 0.255 | 1.19 | 8.08E-02 | 1.07E-01 | no | no | 0 | 17 | 18 | 19 | 22 | 14 | 15 | 0.025340898 | 0.024304154 | 0.025696712 | 0.025554706 | 0.01940237 | 0.019845865 |
| NITMOv2_3330 | A0A0K2GFJ0 | Uncharacterized protein | 20.865 | 181 | 0.487 | 1.40 | 1.19E-03 | 3.18E-03 | no | no | 0 | 7 | 7 | 6 | 4 | 4 | 4 | 0.025233397 | 0.014356443 | 0.00803226 | 0.00816646 | 0.007212654 | 0.006877814 |
| NITMOv2_3545 | A0A0K2GH56 | HD domain-containing protein | 54.582 | 472 | 0.492 | 1.41 | 1.82E-03 | 4.46E-03 | no | no | 0 | 40 | 46 | 44 | 34 | 31 | 30 | 0.025224965 | 0.025121527 | 0.02538612 | 0.01952511 | 0.018397139 | 0.017028209 |
| devR.3 | A0A0K2GBH8 | CRISPR-associated protein Cas7/Cst2/DevR | 34.878 | 314 | -0.369 | -1.29 | 1.33E-02 | 2.26E-02 | no | no | 0 | 54 | 56 | 59 | 48 | 50 | 49 | 0.025161729 | 0.023836804 | 0.031858708 | 0.022896768 | 0.029192115 | 0.024436469 |
| resA | A0A0K2G859 | Thiol-disulfide oxidoreductase ResA | 23.524 | 211 | 0.624 | 1.54 | 8.98E-05 | 5.26E-04 | no | no | 0 | 31 | 27 | 29 | 21 | 21 | 17 | 0.025119571 | 0.02660375 | 0.02567754 | 0.017937751 | 0.01940237 | 0.018790368 |
| NITMOv2_3606 | A0A0K2GGB8 | Uncharacterized protein | 7.6968 | 61 | -0.109 | -1.08 | 5.53E-01 | 5.95E-01 | no | no | 0 | 1 | 3 | 3 | 2 | 4 | 2 | 0.025085845 | 0.060730075 | 0.065895396 | 0.026360231 | 0.076344784 | 0.050359984 |
| NITMOv2_4462 | A0A0K2GIS1 | Transcriptional regulator, MerR family | 13.416 | 115 | 0.414 | 1.33 | 1.37E-03 | 3.59E-03 | no | no | 0 | 6 | 9 | 10 | 7 | 9 | 6 | 0.025035256 | 0.024738261 | 0.025223155 | 0.022804961 | 0.022598292 | 0.016725765 |
| hemE | A0A0K2G989 | Uroporphyrinogen decarboxylase | 37.971 | 341 | -0.299 | -1.23 | 1.97E-02 | 3.14E-02 | no | no | 0 | 21 | 18 | 21 | 23 | 27 | 21 | 0.025018393 | 0.024779325 | 0.027176818 | 0.034738289 | 0.031061194 | 0.028899406 |
| NITMOv2_3539 | A0A0K2GG71 | Putative Outer membrane chaperone Skp (Modular protein) | 26.978 | 237 | 0.487 | 1.40 | 3.30E-03 | 7.15E-03 | no | no | 0 | 7 | 9 | 9 | 7 | 8 | 8 | 0.024953049 | 0.024927939 | 0.02289563 | 0.011696409 | 0.009901765 | 0.014621783 |
| accC | A0A0K2GCB2 | Biotin carboxylase | 48.945 | 446 | -0.263 | -1.20 | 4.46E-03 | 9.05E-03 | no | no | 0 | 32 | 26 | 29 | 37 | 33 | 34 | 0.024908784 | 0.020451938 | 0.026500034 | 0.028750155 | 0.02946867 | 0.02782413 |
| NITMOv2_0600 | A0A0K2G7T7 | Uncharacterized protein | 19.82 | 174 | 0.771 | 1.71 | 5.97E-05 | 3.98E-04 | no | no | 0 | 14 | 14 | 12 | 5 | 8 | 7 | 0.024830792 | 0.024939671 | 0.023351932 | 0.010588367 | 0.012754945 | 0.013068386 |
| fabG.1 | A0A0K2G836 | 3-oxoacyl-[acyl-carrier-protein] reductase | 25.864 | 247 | -0.626 | -1.54 | 2.14E-04 | 9.25E-04 | no | no | 0 | 16 | 14 | 14 | 24 | 22 | 24 | 0.024700104 | 0.023850492 | 0.026691757 | 0.048481088 | 0.046012267 | 0.04184588 |
| NITMOv2_2412 | A0A0K2GDZ7 | Putative Outer membrane protein TolC | 48.719 | 442 | 0.458 | 1.37 | 3.55E-04 | 1.31E-03 | no | no | 0 | 47 | 50 | 56 | 44 | 41 | 39 | 0.024685349 | 0.024409747 | 0.024906811 | 0.016622849 | 0.016385121 | 0.016633521 |
| mdmC | A0A0K2G8R7 | O-methyltransferase MdmC | 23.399 | 213 | 0.232 | 1.17 | 2.87E-02 | 4.33E-02 | no | no | 0 | 17 | 18 | 18 | 19 | 17 | 19 | 0.024643191 | 0.021423792 | 0.02304134 | 0.021068047 | 0.01954997 | 0.018076514 |
| NITMOv2_1175 | A0A0K2G9F7 | ABC-type transport system, ATPase component | 36.149 | 331 | -0.0591 | -1.04 | 3.68E-01 | 4.11E-01 | no | no | 0 | 18 | 22 | 24 | 30 | 30 | 27 | 0.024464021 | 0.02471284 | 0.02730719 | 0.03189822 | 0.031502439 | 0.029496382 |

| | | | | | | | | | | | | | | | | | | | | | | | |
|---|---|---|---|---|---|---|---|---|---|---|---|---|---|---|---|---|---|---|---|---|---|---|---|
| NITMOv2_0340 | A0A0K2G743 | Uncharacterized protein | 26.888 | 250 | -1.68 | -3.20 | 1.88E-07 | 1.28E-05 | yes | no | 0 | 20 | 16 | 16 | 52 | 49 | 51 | 0.024312254 | 0.022276363 | 0.024471598 | 0.092577689 | 0.094084716 | 0.090359193 |
| NITMOv2_0893 | A0A0K2G8P2 | Uncharacterized protein | 22.664 | 212 | 0.368 | 1.29 | 1.90E-02 | 3.05E-02 | no | no | 0 | 13 | 16 | 14 | 13 | 12 | 12 | 0.024263773 | 0.026142266 | 0.026413758 | 0.025680569 | 0.02273657 | 0.020315175 |
| NITMOv2_4602 | A0A0K2GJE6 | Putative Peptidase M16 | 53.632 | 492 | 0.183 | 1.14 | 1.07E-01 | 1.37E-01 | no | no | 0 | 34 | 31 | 38 | 35 | 31 | 30 | 0.024255342 | 0.026081648 | 0.022443163 | 0.019741299 | 0.021568202 | 0.019290246 |
| NITMOv2_2978 | A0A0K2GEK6 | Uncharacterized protein | 30.892 | 287 | 0.547 | 1.46 | 3.28E-04 | 1.23E-03 | no | no | 0 | 22 | 22 | 26 | 18 | 19 | 20 | 0.024202645 | 0.021414014 | 0.024954742 | 0.01674427 | 0.016955322 | 0.017064531 |
| gcvT | A0A0K2G6S5 | Aminomethyltransferase | 39.828 | 367 | 0.287 | 1.22 | 6.19E-03 | 1.19E-02 | no | no | 0 | 25 | 24 | 23 | 25 | 21 | 22 | 0.024105682 | 0.023701878 | 0.023018333 | 0.021863207 | 0.022295325 | 0.020673001 |
| NITMOv2_1109 | A0A0K2G9C6 | Uncharacterized protein | 21.909 | 205 | 0.18 | 1.13 | 9.64E-02 | 1.25E-01 | no | no | 0 | 20 | 20 | 20 | 13 | 13 | 12 | 0.023991857 | 0.0261364 | 0.023601173 | 0.02039875 | 0.018815079 | 0.018865889 |
| NITMOv2_4089 | A0A0K2GHR8 | Uncharacterized protein | 8.9933 | 79 | 0.388 | 1.31 | 1.91E-03 | 4.64E-03 | no | no | 0 | 3 | 5 | 6 | 7 | 8 | 5 | 0.023966563 | 0.023633438 | 0.01716252 | 0.020951068 | 0.020732321 | 0.019858452 |
| NITMOv2_3935 | A0A0K2GH84 | Putative Universal stress protein | 30.816 | 285 | 1.18 | 2.27 | 3.86E-06 | 6.39E-05 | yes | no | 0 | 16 | 16 | 16 | 10 | 10 | 10 | 0.023941268 | 0.022790644 | 0.025334354 | 0.01028896 | 0.00960004 | 0.009550901 |
| dapF | A0A0K2G7Y1 | Diaminopimelate epimerase | 30.764 | 289 | -0.241 | -1.18 | 3.96E-02 | 5.72E-02 | no | no | 0 | 22 | 22 | 29 | 34 | 25 | 28 | 0.023934944 | 0.025137171 | 0.026687923 | 0.028881941 | 0.027896345 | 0.025923515 |
| lysU | A0A0K2GCL3 | Lysine--tRNA ligase | 56.483 | 494 | -0.0575 | -1.04 | 4.15E-01 | 4.59E-01 | no | no | 0 | 33 | 28 | 28 | 31 | 28 | 36 | 0.023907542 | 0.023070271 | 0.023884924 | 0.029179571 | 0.028449456 | 0.028002144 |
| gmhA | A0A088ND31 | Phosphoheptose isomerase | 24.388 | 229 | 3.04 | 8.22 | 3.51E-06 | 6.20E-05 | yes | no | 0 | 13 | 10 | 14 | 1 | 3 | 1 | 0.023804256 | 0.021597826 | 0.022665562 | 0.002733752 | 0.003193126 | 0.002215106 |
| NITMOv2_3680 | A0A0K2GHH3 | Uncharacterized protein | 28.341 | 256 | 0.314 | 1.24 | 4.17E-02 | 5.99E-02 | no | no | 0 | 16 | 16 | 21 | 16 | 13 | 14 | 0.023772638 | 0.025976054 | 0.029586783 | 0.0191875 | 0.014377299 | 0.015102421 |
| fbp.1 | A0A0K2GFZ5 | Fructose-1,6-bisphosphatase class 1 | 35.482 | 325 | 0.219 | 1.16 | 2.71E-02 | 4.12E-02 | no | no | 0 | 29 | 28 | 37 | 33 | 37 | 30 | 0.023648273 | 0.022086686 | 0.024193599 | 0.022704271 | 0.02407895 | 0.021710517 |
| NITMOv2_1258 | A0A0K2G9P0 | Uncharacterized protein | 34.696 | 310 | -0.016 | -1.01 | 8.44E-01 | 8.63E-01 | no | no | 0 | 20 | 22 | 22 | 25 | 23 | 27 | 0.023604008 | 0.022798465 | 0.021678186 | 0.024941677 | 0.023372026 | 0.021620611 |
| NITMOv2_1377 | A0A0K2GAA3 | HTH cro/C1-type domain-containing protein | 11.934 | 105 | 0.553 | 1.47 | 1.19E-03 | 3.18E-03 | no | no | 0 | 4 | 6 | 5 | 8 | 7 | 6 | 0.023507045 | 0.023875912 | 0.021137525 | 0.019054233 | 0.017042329 | 0.017040436 |
| cysE | A0A0K2GG70 | Serine acetyltransferase | 23.523 | 216 | 0.16 | 1.12 | 5.46E-02 | 7.57E-02 | no | no | 0 | 15 | 13 | 17 | 11 | 12 | 13 | 0.023479643 | 0.022131661 | 0.024605805 | 0.020721553 | 0.021067916 | 0.018993556 |
| NITMOv2_0483 | A0A0K2G7U4 | Putative nickel insertion protein | 43.06 | 401 | 0.308 | 1.24 | 3.81E-03 | 8.00E-03 | no | no | 0 | 22 | 27 | 24 | 16 | 23 | 19 | 0.023370033 | 0.025645585 | 0.026595896 | 0.021269428 | 0.019299827 | 0.019703814 |
| yliI | A0A0K2G8A6 | Soluble aldose sugar dehydrogenase YliI | 42.181 | 384 | 0.724 | 1.65 | 3.35E-05 | 2.67E-04 | no | no | 0 | 32 | 31 | 29 | 21 | 20 | 21 | 0.023340523 | 0.022340893 | 0.022560114 | 0.012097543 | 0.016291901 | 0.015555188 |
| NITMOv2_4523 | A0A0K2GJV1 | Putative 37.6 kDa protein in cld 5region | 38.116 | 340 | -0.257 | -1.19 | 7.40E-03 | 1.38E-02 | no | no | 0 | 29 | 32 | 28 | 32 | 29 | 30 | 0.023222482 | 0.023304924 | 0.023211974 | 0.035902155 | 0.035761079 | 0.033236475 |
| NITMOv2_0611 | A0A0K2G8W6 | UPF0234 protein NITMOv2_0611 | 18.401 | 165 | -0.176 | -1.13 | 8.18E-02 | 1.08E-01 | no | no | 0 | 7 | 5 | 8 | 11 | 11 | 9 | 0.023203511 | 0.025074597 | 0.027439479 | 0.041361664 | 0.038274935 | 0.0347433 |
| pfp | A0A0K2GGX4 | Pyrophosphate--fructose 6-phosphate 1-phosphotransferase | 44.982 | 416 | 0.429 | 1.35 | 7.46E-04 | 2.29E-03 | no | no | 0 | 38 | 38 | 33 | 29 | 32 | 25 | 0.023180324 | 0.024744127 | 0.024187847 | 0.017355818 | 0.016639925 | 0.015737518 |
| NITMOv2_3938 | A0A0K2GH82 | Uncharacterized protein | 10.7 | 93 | 0.767 | 1.70 | 1.59E-04 | 7.59E-04 | no | no | 0 | 9 | 9 | 8 | 3 | 6 | 4 | 0.023112872 | 0.027530628 | 0.026022642 | 0.014660269 | 0.016296562 | 0.016118 |
| exbD | A0A0K2G8T1 | Biopolymer transport protein exbD | 13.576 | 124 | 0.632 | 1.55 | 1.42E-03 | 3.68E-03 | no | no | 0 | 5 | 6 | 6 | 2 | 2 | 4 | 0.02308547 | 0.023275593 | 0.022262943 | 0.008881067 | 0.013616772 | 0.015279356 |
| gmk | A0A0K2GC13 | Guanylate kinase | 27.576 | 238 | 0.0692 | 1.05 | 3.40E-01 | 3.83E-01 | no | no | 0 | 16 | 15 | 15 | 17 | 19 | 14 | 0.023041204 | 0.022876683 | 0.023382608 | 0.021644057 | 0.023368918 | 0.022823554 |
| purC | A0A0K2G9W0 | Phosphoribosylaminoimidazole-succinocarboxamide synthase | 26.806 | 235 | 0.435 | 1.35 | 3.57E-04 | 1.31E-03 | no | no | 0 | 20 | 19 | 26 | 17 | 17 | 17 | 0.022996939 | 0.022309606 | 0.035708517 | 0.022055704 | 0.020810005 | 0.01925788 |
| NITMOv2_4239 | A0A0K2GI21 | Putative 3-oxoacyl-(Acyl-carrier-protein) reductase | 27.339 | 255 | 0.291 | 1.22 | 4.50E-03 | 9.10E-03 | no | no | 0 | 19 | 22 | 28 | 17 | 19 | 18 | 0.022967429 | 0.022587278 | 0.022090391 | 0.018912082 | 0.018538524 | 0.017699088 |
| NITMOv2_3816 | A0A0K2GGZ1 | Uncharacterized protein | 18.057 | 160 | 0.288 | 1.22 | 3.52E-02 | 5.18E-02 | no | no | 0 | 10 | 13 | 10 | 5 | 9 | 8 | 0.022786151 | 0.024736306 | 0.02683555 | 0.018093229 | 0.021140939 | 0.020584893 |
| NITMOv2_0025 | A0A0K2G6J6 | Multidrug efflux system protein | 117.31 | 1084 | 0.305 | 1.24 | 3.32E-03 | 7.17E-03 | no | no | 0 | 47 | 51 | 57 | 52 | 51 | 45 | 0.022729239 | 0.023512201 | 0.024711253 | 0.019135674 | 0.018263522 | 0.019367565 |
| hisB | A0A0K2G8G2 | Imidazoleglycerol-phosphate dehydratase | 22.384 | 201 | -0.00245 | -1.00 | 9.65E-01 | 9.69E-01 | no | no | 0 | 17 | 17 | 16 | 13 | 11 | 10 | 0.022703944 | 0.023236484 | 0.024881887 | 0.023395779 | 0.024439405 | 0.018714847 |
| NITMOv2_0841 | A0A0K2G8H7 | Uncharacterized protein | 26 | 244 | 0.678 | 1.60 | 2.39E-03 | 5.52E-03 | no | no | 0 | 16 | 15 | 14 | 15 | 15 | 13 | 0.022659679 | 0.024967047 | 0.019991018 | 0.018838044 | 0.015553902 | 0.013871787 |
| gatA | A0A0K2GG32 | Glutamyl-tRNA(Gln) amidotransferase subunit A | 52.684 | 488 | 0.133 | 1.10 | 9.65E-02 | 1.25E-01 | no | no | 0 | 46 | 49 | 56 | 43 | 42 | 48 | 0.022649139 | 0.021914607 | 0.022452749 | 0.025402189 | 0.025182064 | 0.021581052 |
| NITMOv2_4464 | A0A0K2GIQ1 | (R)-citramalate synthase | 57.248 | 520 | -0.0853 | -1.06 | 2.22E-01 | 2.63E-01 | no | no | 0 | 36 | 34 | 35 | 31 | 34 | 29 | 0.022518451 | 0.021797281 | 0.025085114 | 0.022876037 | 0.023976408 | 0.022156451 |
| tldD | A0A0K2GAC9 | Putative peptidase | 53.282 | 494 | 0.344 | 1.27 | 1.23E-03 | 3.27E-03 | no | no | 0 | 37 | 34 | 35 | 34 | 34 | 37 | 0.022488941 | 0.022333071 | 0.022103812 | 0.018823237 | 0.018474823 | 0.01985126 |
| NITMOv2_1108 | A0A0K2G9K5 | Uncharacterized protein | 15.494 | 138 | 0.339 | 1.26 | 2.56E-02 | 3.93E-02 | no | no | 0 | 15 | 18 | 17 | 14 | 14 | 10 | 0.022341389 | 0.018122619 | 0.020015942 | 0.018076941 | 0.019298274 | 0.017692435 |
| NITMOv2_4187 | A0A0K2GIX1 | Uncharacterized protein | 19.673 | 172 | 0.741 | 1.67 | 2.79E-05 | 2.34E-04 | no | no | 0 | 22 | 19 | 26 | 18 | 17 | 22 | 0.022305555 | 0.022800421 | 0.017577793 | 0.012618173 | 0.012761937 | 0.01684516 |
| glgE | A0A0K2GI06 | Alpha-1,4-glucan:maltose-1-phosphate maltosyltransferase | 76.535 | 664 | 0.5 | 1.41 | 2.63E-04 | 1.06E-03 | no | no | 0 | 31 | 27 | 33 | 29 | 28 | 30 | 0.022115846 | 0.021079634 | 0.021641758 | 0.016291162 | 0.015724807 | 0.017283722 |
| mdh | A0A0K2GHM5 | Malate dehydrogenase | 32.643 | 312 | -0.545 | -1.46 | 2.01E-04 | 8.80E-04 | no | no | 0 | 21 | 22 | 30 | 34 | 38 | 38 | 0.022035747 | 0.021873543 | 0.021260228 | 0.033550732 | 0.034482399 | 0.029075622 |
| ubact | A0A0K2G9F6 | Prokaryotic ubiquitin-like protein UBact | 7.6115 | 66 | -0.223 | -1.17 | 4.18E-01 | 4.62E-01 | no | no | 0 | 5 | 4 | 3 | 3 | 3 | 2 | 0.021841822 | 0.017535987 | 0.012930414 | 0.019116424 | 0.021185996 | 0.013531582 |
| deaD | A0A0K2GCU7 | Cold-shock DEAD box protein A | 61.932 | 576 | 0.413 | 1.33 | 7.40E-04 | 2.28E-03 | no | no | 0 | 56 | 52 | 50 | 46 | 44 | 42 | 0.021820744 | 0.022735891 | 0.023378774 | 0.018636663 | 0.018822847 | 0.019241696 |
| Hmgcll | A0A0K2G6L9 | 3-hydroxymethyl-3-methylglutaryl-CoA lyase, cytoplasmic | 36.523 | 339 | 2.72 | 6.59 | 2.69E-07 | 1.39E-05 | yes | no | 0 | 19 | 17 | 19 | 3 | 5 | 4 | 0.021749076 | 0.021945894 | 0.01959415 | 0.003528913 | 0.003443579 | 0.002390782 |
| rplV | A0A0K2G9U8 | 50S ribosomal protein L22 | 13.023 | 118 | -0.252 | -1.19 | 1.67E-02 | 2.73E-02 | no | no | 0 | 3 | 8 | 7 | 12 | 12 | 11 | 0.021736429 | 0.022043666 | 0.024797528 | 0.054603971 | 0.051863431 | 0.050645886 |
| NITMOv2_1166 | A0A0K2GAG5 | Putative Transcriptional regulator, AsnC family (Modular protein) | 14.853 | 138 | 0.421 | 1.34 | 4.10E-02 | 5.90E-02 | no | no | 0 | 4 | 3 | 5 | 4 | 2 | 2 | 0.021725889 | 0.023293192 | 0.023300167 | 0.018586318 | 0.014176874 | 0.014874779 |
| gatB | A0A0K2GFU3 | Aspartyl/glutamyl-tRNA(Asn/Gln) amidotransferase subunit B | 52.808 | 476 | 0.336 | 1.26 | 2.93E-03 | 6.49E-03 | no | no | 0 | 31 | 37 | 36 | 31 | 30 | 27 | 0.021706918 | 0.022010423 | 0.021127939 | 0.015753651 | 0.016723824 | 0.016303926 |
| NITMOv2_3237 | A0A0K2GG79 | Uncharacterized protein | 16.48 | 151 | 0.496 | 1.41 | 4.28E-04 | 1.51E-03 | no | no | 0 | 6 | 6 | 7 | 4 | 6 | 4 | 0.02163525 | 0.020358077 | 0.01921262 | 0.014308148 | 0.015133942 | 0.013448509 |
| NITMOv2_0256 | A0A0K2G6Y8 | Uncharacterized protein | 10.624 | 101 | 0.634 | 1.55 | 5.19E-03 | 1.03E-02 | no | no | 0 | 5 | 5 | 4 | 3 | 3 | 2 | 0.021580446 | 0.020602507 | 0.019481033 | 0.017634198 | 0.01105817 | 0.010048262 |
| NITMOv2_4009 | A0A0K2GHF6 | Uncharacterized protein | 8.3034 | 70 | 1.2 | 2.30 | 1.70E-05 | 1.66E-04 | yes | no | 0 | 7 | 5 | 2 | 2 | 2 | 2 | 0.021561475 | 0.021292777 | 0.018931362 | 0.008325195 | 0.008400754 | 0.007718975 |
| NITMOv2_3302 | A0A0K2GFF6 | Uncharacterized protein | 66.018 | 615 | 0.0719 | 1.05 | 3.95E-01 | 4.38E-01 | no | no | 0 | 46 | 44 | 50 | 42 | 43 | 40 | 0.021458189 | 0.023363587 | 0.022970403 | 0.02361641 | 0.022177245 | 0.019446682 |
| NITMOv2_1184 | A0A0K2G9I3 | Uncharacterized protein | 30.397 | 277 | 0.355 | 1.28 | 5.57E-03 | 1.08E-02 | no | no | 0 | 14 | 13 | 13 | 11 | 10 | 14 | 0.021447649 | 0.021067902 | 0.019607571 | 0.016202317 | 0.016197126 | 0.016171404 |
| purL | A0A0K2G9U4 | Phosphoribosylformylglycinamidine synthase subunit PurL | 80.028 | 743 | 0.317 | 1.25 | 2.75E-03 | 6.18E-03 | no | no | 0 | 44 | 47 | 51 | 42 | 43 | 40 | 0.021401276 | 0.022765223 | 0.023351932 | 0.018509319 | 0.019581044 | 0.018272509 |
| thrC | A0A0K2GD70 | Threonine synthase | 45.076 | 416 | -1.41 | -2.66 | 1.02E-06 | 3.05E-05 | yes | no | 0 | 16 | 18 | 18 | 53 | 54 | 45 | 0.021321177 | 0.019162131 | 0.023396029 | 0.067123674 | 0.070763962 | 0.062421782 |
| NITMOv2_4077 | A0A0K2GHR0 | Uncharacterized protein | 54.084 | 491 | 0.138 | 1.10 | 1.47E-01 | 1.82E-01 | no | no | 0 | 44 | 45 | 40 | 50 | 47 | 42 | 0.021310637 | 0.02257359 | 0.022772927 | 0.021910591 | 0.020910995 | 0.019948358 |
| rplT | A0A0K2G7B9 | 50S ribosomal protein L20 | 13.597 | 122 | -0.385 | -1.31 | 2.37E-03 | 5.49E-03 | no | no | 0 | 5 | 5 | 4 | 7 | 9 | 7 | 0.021310637 | 0.023234528 | 0.02388109 | 0.021122835 | 0.03088252 | 0.023204756 |
| NITMOv2_4099 | A0A0K2GHS6 | Uncharacterized protein | 13.1 | 121 | 0.227 | 1.17 | 6.20E-02 | 8.47E-02 | no | no | 0 | 8 | 5 | 4 | 5 | 5 | 5 | 0.02122843 | 0.020223152 | 0.019116759 | 0.018546337 | 0.016281025 | 0.014719601 |
| NITMOv2_3010 | A0A0K2GEP4 | Transcriptional regulator, AbrB family | 10.516 | 93 | 0.858 | 1.81 | 2.75E-04 | 1.09E-03 | no | no | 0 | 8 | 7 | 8 | 3 | 4 | 3 | 0.021194704 | 0.023860269 | 0.026189442 | 0.015038451 | 0.014599631 | 0.012823662 |
| cusR | A0A0K2G6T7 | DNA-binding response regulator in two-component regulatory system with CusS | 25.341 | 223 | 0.359 | 1.28 | 1.71E-03 | 4.27E-03 | no | no | 0 | 22 | 20 | 20 | 15 | 15 | 14 | 0.021160978 | 0.024210293 | 0.023892593 | 0.015885438 | 0.016184697 | 0.017506869 |
| NITMOv2_4037 | A0A0K2GHM4 | Nitrogen regulation protein B | 55.654 | 504 | 0.532 | 1.45 | 1.74E-04 | 7.94E-04 | no | no | 0 | 22 | 26 | 28 | 22 | 20 | 18 | 0.021150439 | 0.020582953 | 0.020234507 | 0.015430849 | 0.015922124 | 0.015049736 |
| NITMOv2_4338 | A0A0K2GIL9 | 6-carboxy-5,6,7,8-tetrahydropterin synthase | 28.103 | 250 | 0.329 | 1.26 | 1.76E-02 | 2.86E-02 | no | no | 0 | 25 | 24 | 24 | 22 | 22 | 19 | 0.021016167 | 0.021730796 | 0.019486785 | 0.019203788 | 0.016265488 | 0.01470108 |
| trpS | A0A0K2GHJ2 | Tryptophan--tRNA ligase | 37.585 | 330 | 0.123 | 1.09 | 1.26E-01 | 1.58E-01 | no | no | 0 | 16 | 23 | 21 | 23 | 21 | 21 | 0.020959465 | 0.020270083 | 0.01996226 | 0.019837547 | 0.019257878 | 0.017125667 |
| NITMOv2_1125 | A0A0K2G9A9 | Uncharacterized protein | 16.722 | 149 | 0.414 | 1.33 | 1.11E-03 | 3.02E-03 | no | no | 0 | 22 | 24 | 24 | 21 | 19 | 17 | 0.020925739 | 0.021693642 | 0.024866549 | 0.018152459 | 0.020109295 | 0.020228865 |
| NITMOv2_3250 | A0A0K2GFA7 | Uncharacterized protein | 119.97 | 1119 | 0.567 | 1.48 | 3.14E-04 | 1.19E-03 | no | no | 0 | 64 | 68 | 77 | 56 | 56 | 58 | 0.020890327 | 0.021773815 | 0.021032077 | 0.014919992 | 0.015364353 | 0.014644979 |
| proC | A0A0K2G8X8 | Pyrroline-5-carboxylate reductase | 28.225 | 272 | 0.0477 | 1.03 | 6.03E-01 | 6.43E-01 | no | no | 0 | 15 | 17 | 15 | 14 | 13 | 15 | 0.020803482 | 0.019912237 | 0.020673554 | 0.02333803 | 0.02187117 | 0.020471611 |
| Mccc | A0A0K2G6A7 | Methylcrotonoyl-CoA carboxylase beta chain, mitochondrial | 58.1 | 534 | 2.43 | 5.39 | 1.41E-07 | 1.06E-05 | yes | no | 0 | 27 | 30 | 26 | 7 | 5 | 5 | 0.020750574 | 0.021308421 | 0.019442689 | 0.003989573 | 0.002837799 | 0.003028935 |
| rsfS | A0A0K2G977 | Ribosomal silencing factor RsfS | 14.42 | 131 | 0.418 | 1.34 | 3.37E-03 | 7.23E-03 | no | no | 0 | 7 | 7 | 9 | 8 | 9 | 9 | 0.020673848 | 0.025862639 | 0.021760627 | 0.022938229 | 0.021347579 | 0.017503273 |
| fabG | A0A0K2G7G5 | 3-oxoacyl-[acyl-carrier-protein] reductase FabG | 26.657 | 251 | 0.477 | 1.39 | 1.84E-04 | 8.22E-04 | no | no | 0 | 14 | 13 | 13 | 10 | 9 | 10 | 0.020535782 | 0.020749165 | 0.020685057 | 0.013449019 | 0.014063766 | 0.014256585 |
| rpoH | A0A0K2G9U6 | RNA polymerase sigma factor | 38.245 | 332 | 0.69 | 1.61 | 4.23E-05 | 3.08E-04 | no | no | 0 | 11 | 9 | 6 | 9 | 8 | 8 | 0.020474443 | 0.020539933 | 0.020203831 | 0.013367874 | 0.01326766 | 0.013350691 |
| mrcA | A0A0K2GID9 | DD-transpeptidase | 87.691 | 800 | 0.331 | 1.26 | 2.86E-03 | 6.37E-03 | no | no | 0 | 64 | 67 | 65 | 50 | 56 | 52 | 0.020469805 | 0.020131246 | 0.018233296 | 0.014573497 | 0.015267403 | 0.014533496 |
| NITMOv2_0738 | A0A0K2G8I9 | Putative Cytochrome c | 22.214 | 207 | -0.07 | -1.05 | 6.88E-01 | 7.25E-01 | no | no | 0 | 5 | 6 | 10 | 26 | 24 | 22 | 0.020412682 | 0.019867262 | 0.015642152 | 0.022422929 | 0.025528535 | 0.021710517 |
| NITMOv2_0010 | A0A0K2G6I5 | Uncharacterized protein | 16.29 | 148 | 0.729 | 1.66 | 8.04E-04 | 2.41E-03 | no | no | 0 | 3 | 5 | 6 | 4 | 5 | 3 | 0.020354926 | 0.018139436 | 0.021919757 | 0.0122585 | 0.01075194 | 0.010650812 |
| NITMOv2_4613 | A0A0K2GJ68 | Uncharacterized protein | 8.664 | 76 | 0.56 | 1.47 | 5.80E-02 | 7.98E-02 | no | no | 0 | 9 | 8 | 7 | 8 | 5 | 3 | 0.020301175 | 0.020403052 | 0.015880081 | 0.017228474 | 0.015413915 | 0.013383777 |
| NITMOv2_0054 | A0A0K2G7D3 | Lactamase_B domain-containing protein | 32.219 | 282 | 4.5 | 22.63 | 5.15E-08 | 6.80E-06 | yes | no | 0 | 11 | 12 | 11 | 0 | 0 | 0 | 0.020272508 | 0.020872358 | 0.022226515 | 0.00090589 | 0.001048844 | 0.001504182 |
| NITMOv2_0686 | A0A0K2G937 | Uncharacterized protein | 16.114 | 138 | 0.502 | 1.42 | 4.05E-04 | 1.44E-03 | no | no | 0 | 9 | 10 | 10 | 6 | 5 | 6 | 0.020184609 | 0.019040893 | 0.019296979 | 0.012080662 | 0.012512571 | 0.01263396 |
| serS | A0A0K2GAW0 | Serine--tRNA ligase | 48.51 | 428 | -0.232 | -1.17 | 1.10E-02 | 1.93E-02 | no | no | 0 | 37 | 28 | 31 | 46 | 45 | 41 | 0.020181026 | 0.016469882 | 0.01862748 | 0.022550273 | 0.02392669 | 0.022249953 |
| NITMOv2_1736 | A0A0K2GB09 | Plasmid stabilization system protein | 10.007 | 84 | 0.637 | 1.56 | 3.82E-03 | 8.01E-03 | no | no | 0 | 3 | 4 | 2 | 1 | 2 | 3 | 0.020103456 | 0.020135157 | 0.016709094 | 0.012305735 | 0.012932687 | 0.012733576 |
| NITMOv2_3540 | A0A0K2GH52 | Putative Outer membrane protein assembly factor YaeT | 87.549 | 780 | 0.483 | 1.40 | 1.82E-04 | 8.17E-04 | no | no | 0 | 50 | 50 | 47 | 44 | 45 | 41 | 0.020087015 | 0.020197731 | 0.020439651 | 0.014339243 | 0.015167502 | 0.015410439 |
| NITMOv2_3944 | A0A0K2GHI6 | Hemerythrin domain-containing protein | 19.095 | 161 | 1.9 | 3.73 | 2.49E-05 | 2.20E-04 | yes | no | 0 | 9 | 8 | 10 | 2 | 1 | 1 | 0.020048862 | 0.015526969 | 0.020006356 | 0.00385364 | 0.003759442 | 0.003430816 |
| NITMOv2_2293 | A0A0K2GCW3 | Uncharacterized protein | 11.668 | 104 | 0.0965 | 1.07 | 2.94E-01 | 3.36E-01 | no | no | 0 | 2 | 3 | 3 | 2 | 2 | 2 | 0.020020406 | 0.019878995 | 0.019496371 | 0.015883957 | 0.018025809 | 0.0080777 |
| NITMOv2_3600 | A0A0K2GHA6 | TPR_REGION domain-containing protein | 11.68 | 105 | 0.421 | 1.34 | 2.24E-02 | 3.51E-02 | no | no | 0 | 9 | 7 | 9 | 10 | 8 | 7 | 0.020015136 | 0.019545592 | 0.019083782 | 0.016384449 | 0.014788248 | 0.01285405 |
| NITMOv2_4059 | A0A0K2GHL3 | Putative 3-oxoadipate enol-lactonase | 28.73 | 263 | 0.697 | 1.62 | 4.19E-04 | 1.49E-03 | no | no | 0 | 10 | 11 | 11 | 7 | 7 | 6 | 0.019952321 | 0.020166444 | 0.019117142 | 0.013165456 | 0.013144453 | 0.01212653 |
| queF | A0A0K2G922 | NADPH-dependent 7-cyano-7-deazaguanine reductase | 15.753 | 138 | 0.189 | 1.14 | 3.92E-02 | 5.68E-02 | no | no | 0 | 8 | 9 | 9 | 6 | 8 | 8 | 0.019899203 | 0.020107781 | 0.020315031 | 0.018832121 | 0.019369743 | 0.020248644 |
| NITMOv2_0118 | A0A0K2G6J0 | Putative Tn7-like transposition protein C | 59.573 | 530 | 0.819 | 1.76 | 5.71E-05 | 3.86E-04 | no | no | 0 | 20 | 25 | 23 | 15 | 15 | 17 | 0.019894566 | 0.02001392 | 0.020169321 | 0.010334419 | 0.011809064 | 0.013267079 |

| | | | | | | | | | | | | | | | | | | | | | | | |
|---|---|---|---|---|---|---|---|---|---|---|---|---|---|---|---|---|---|---|---|---|---|---|---|
| NITMOv2_1133 | A0A0K2G9M8 | Cupredoxin_1 domain-containing protein | 15.461 | 145 | 1.07 | 2.10 | 1.55E-05 | 1.57E-04 | yes | no | 0 | 4 | 4 | 4 | 2 | 4 | 3 | 0.019869903 | 0.020764809 | 0.020708064 | 0.012363633 | 0.013735007 | 0.013088166 |
| pgk | A0A0K2G913 | Phosphoglycerate kinase | 42.478 | 399 | -0.718 | -1.64 | 5.08E-05 | 3.55E-04 | no | no | 0 | 28 | 32 | 38 | 49 | 53 | 51 | 0.019814255 | 0.020084316 | 0.022698155 | 0.036336014 | 0.038643158 | 0.036009178 |
| NITMOv2_0635 | A0A0K2G7X3 | Uncharacterized protein | 11.741 | 105 | -0.685 | -1.61 | 1.89E-05 | 1.78E-04 | no | no | 0 | 6 | 7 | 8 | 9 | 10 | 9 | 0.019752705 | 0.0240871 | 0.024670991 | 0.041308357 | 0.044127652 | 0.043850785 |
| nudF | A0A0K2G6W8 | ADP-ribose pyrophosphatase | 18.631 | 171 | 0.387 | 1.31 | 2.22E-03 | 5.20E-03 | no | no | 0 | 11 | 8 | 11 | 11 | 8 | 10 | 0.019712867 | 0.01972647 | 0.018846428 | 0.015374581 | 0.015187699 | 0.012707503 |
| NITMOv2_4416 | A0A0K2GIM4 | Uncharacterized protein | 14.4 | 127 | 0.875 | 1.83 | 3.13E-01 | 3.54E-01 | no | yes | 1 | 3 | 3 | 1 | 2 | 2 | 2 | 0.019511775 | 0.01826478 | 0.019354496 | 0.021732902 | 0.024649151 | 0.024280032 |
| NITMOv2_3497 | A0A0K2GGA7 | Putative SAM-dependent methyltransferase | 35.659 | 314 | -0.432 | -1.35 | 1.01E-03 | 2.84E-03 | no | no | 0 | 14 | 16 | 20 | 22 | 21 | 21 | 0.019475309 | 0.020027608 | 0.021735703 | 0.033205718 | 0.031006815 | 0.028126214 |
| NITMOv2_3221 | A0A0K2GF84 | Uncharacterized protein | 11.962 | 105 | 0.858 | 1.81 | 5.77E-05 | 3.89E-04 | no | no | 0 | 6 | 5 | 4 | 1 | 1 | 0 | 0.01942746 | 0.021949805 | 0.02152289 | 0.009091925 | 0.010309762 | 0.009831948 |
| pyrC | A0A0K2GFT9 | Dihydroorotase | 44.971 | 428 | -0.501 | -1.42 | 3.44E-04 | 1.28E-03 | no | no | 0 | 11 | 10 | 13 | 23 | 19 | 21 | 0.019342934 | 0.01880585 | 0.019423516 | 0.034491005 | 0.029608502 | 0.034459197 |
| NITMOv2_0018 | A0A0K2G7A5 | Putative oxidoreductase | 25.534 | 243 | -0.952 | -1.93 | 2.79E-05 | 2.34E-04 | no | no | 0 | 21 | 20 | 25 | 27 | 27 | 26 | 0.019311738 | 0.020811739 | 0.020318865 | 0.040701252 | 0.046136562 | 0.041653481 |
| NITMOv2_2064 | A0A0K2GC00 | Dioxygenase, ferredoxin subunit | 10.588 | 101 | -0.819 | -1.76 | 1.54E-04 | 7.50E-04 | no | no | 0 | 4 | 4 | 4 | 4 | 5 | 5 | 0.019177044 | 0.020164489 | 0.017115356 | 0.033621807 | 0.038427196 | 0.033072846 |
| NITMOv2_3269 | A0A0K2GFC9 | Uma2 domain-containing protein | 20.973 | 184 | 0.217 | 1.16 | 8.46E-02 | 1.12E-01 | no | no | 0 | 20 | 20 | 20 | 22 | 17 | 18 | 0.019157862 | 0.017539898 | 0.01981655 | 0.019221557 | 0.017227217 | 0.015442086 |
| NITMOv2_2615 | A0A0K2GDK0 | Putative Heavy metal efflux system, membrane fusion protein | 49.148 | 442 | -0.349 | -1.27 | 6.39E-03 | 1.22E-02 | no | no | 0 | 25 | 31 | 27 | 47 | 49 | 48 | 0.019142897 | 0.019677584 | 0.017961816 | 0.024045826 | 0.026052126 | 0.026229196 |
| NITMOv2_0068 | A0A0K2G6F5 | Putative Acyl-CoA dehydrogenase | 74.215 | 659 | 2.29 | 4.89 | 6.46E-08 | 7.15E-06 | yes | no | 0 | 34 | 37 | 37 | 6 | 5 | 7 | 0.01906891 | 0.019704961 | 0.01878546 | 0.002315738 | 0.002092653 | 0.002261317 |
| NITMOv2_2526 | A0A0K2GDL8 | Uncharacterized protein | 21.338 | 199 | 0.618 | 1.53 | 5.59E-05 | 3.80E-04 | no | no | 0 | 17 | 18 | 24 | 17 | 19 | 18 | 0.019064484 | 0.018384844 | 0.018311328 | 0.012651341 | 0.011937088 | 0.010806709 |
| zwf1 | A0A0K2GB65 | Glucose-6-phosphate 1-dehydrogenase | 57.795 | 514 | 0.291 | 1.22 | 6.99E-03 | 1.31E-02 | no | no | 0 | 18 | 21 | 18 | 14 | 17 | 18 | 0.019043616 | 0.018645895 | 0.016870333 | 0.013427844 | 0.013010526 | 0.014454918 |
| NITMOv2_0556 | A0A0K2G8S3 | Putative Outer membrane lipoprotein Slp | 19.674 | 176 | 0.494 | 1.41 | 4.06E-03 | 8.42E-03 | no | no | 0 | 11 | 10 | 10 | 6 | 6 | 9 | 0.019043405 | 0.020962308 | 0.01944844 | 0.01385504 | 0.013365853 | 0.012252938 |
| NITMOv2_1741 | A0A0K2GB14 | Uncharacterized protein | 20.449 | 186 | 0.426 | 1.34 | 5.59E-03 | 1.08E-02 | no | no | 0 | 7 | 6 | 9 | 4 | 5 | 4 | 0.018904706 | 0.018738583 | 0.017780253 | 0.014573793 | 0.013511277 | 0.012919142 |
| argC | A0A0K2GB54 | N-acetyl-gamma-glutamyl-phosphate reductase | 37.726 | 349 | -0.36 | -1.28 | 8.78E-03 | 1.60E-02 | no | no | 0 | 12 | 15 | 23 | 38 | 33 | 26 | 0.018896275 | 0.019413991 | 0.021313911 | 0.040526524 | 0.03543636 | 0.030210237 |
| NITMOv2_3283 | A0A0K2GFE7 | PINc domain-containing protein | 17.345 | 150 | 0.977 | 1.97 | 4.53E-06 | 7.06E-05 | no | no | 0 | 4 | 6 | 4 | 3 | 3 | 3 | 0.01886023 | 0.018429819 | 0.018747691 | 0.010075289 | 0.008755615 | 0.007458248 |
| pheS | A0A0K2G826 | Phenylalanine--tRNA ligase alpha subunit | 57.487 | 521 | -0.0316 | -1.02 | 7.44E-01 | 7.75E-01 | no | no | 0 | 28 | 30 | 29 | 44 | 35 | 34 | 0.018763479 | 0.018129072 | 0.018418118 | 0.023275839 | 0.020213392 | 0.021796827 |
| nuoD | A0A0K2G6W7 | NADH-quinone oxidoreductase subunit D | 47.474 | 414 | -0.483 | -1.40 | 1.51E-04 | 7.40E-04 | no | no | 0 | 19 | 23 | 19 | 30 | 31 | 30 | 0.018756944 | 0.019294123 | 0.01973986 | 0.027102085 | 0.032642841 | 0.031522865 |
| cbbE | A0A0K2G709 | Ribulose-phosphate 3-epimerase | 24.459 | 233 | 0.0465 | 1.03 | 6.43E-01 | 6.82E-01 | no | no | 0 | 11 | 9 | 11 | 9 | 10 | 11 | 0.018714998 | 0.019609144 | 0.019762867 | 0.01906756 | 0.016046419 | 0.017803919 |
| NITMOv2_1483 | A0A0K2GAF0 | Uncharacterized protein | 46.208 | 419 | 1.31 | 2.48 | 7.46E-05 | 4.60E-04 | yes | no | 0 | 18 | 21 | 19 | 11 | 12 | 10 | 0.0187131 | 0.02045976 | 0.01580991 | 0.007691584 | 0.007339124 | 0.006572853 |
| NITMOv2_4414 | A0A0K2GIU4 | Peptidase_M78 domain-containing protein | 11.038 | 92 | 2.14 | 4.41 | 3.54E-03 | 7.57E-03 | yes | yes | 1 | 5 | 5 | 4 | 1 | 1 | 1 | 0.018664408 | 0.01895329 | 0.016022723 | 0.004981968 | 0.004796369 | 0.003810399 |
| NITMOv2_2976 | A0A0K2GEK2 | Uncharacterized protein | 54.843 | 488 | 0.93 | 1.91 | 9.79E-06 | 1.17E-04 | no | no | 0 | 28 | 32 | 33 | 18 | 18 | 20 | 0.018581569 | 0.017162108 | 0.01842502 | 0.009688962 | 0.00921566 | 0.009243962 |
| rph | A0A0K2GCU8 | Ribonuclease PH | 27.06 | 247 | -0.295 | -1.23 | 2.61E-02 | 3.99E-02 | no | no | 0 | 18 | 26 | 20 | 19 | 19 | 18 | 0.018520019 | 0.019035223 | 0.019628661 | 0.021993513 | 0.017634281 | 0.017597494 |
| NITMOv2_0086 | A0A0K2G7F6 | Cu-NIR | 26.869 | 252 | -0.692 | -1.62 | 4.36E-03 | 8.89E-03 | no | no | 0 | 6 | 6 | 8 | 10 | 9 | 5 | 0.018450248 | 0.01827573 | 0.019203034 | 0.026422423 | 0.025303251 | 0.02093193 |
| NITMOv2_4821 | A0A0K2GK06 | 3-hydroxyacid dehydrogenase | 31.031 | 289 | 0.592 | 1.51 | 1.85E-04 | 8.22E-04 | no | no | 0 | 14 | 14 | 14 | 7 | 9 | 8 | 0.018324829 | 0.017321085 | 0.017569741 | 0.011035996 | 0.012529817 | 0.012142893 |
| NITMOv2_0051 | A0A0K2G6E1 | Putative Enoyl-CoA hydratase | 30.428 | 283 | 1.5 | 2.83 | 2.47E-07 | 1.33E-05 | yes | no | 0 | 23 | 29 | 25 | 11 | 16 | 12 | 0.018298692 | 0.017353349 | 0.02300683 | 0.00539021 | 0.006040712 | 0.006386747 |
| NITMOv2_1315 | A0A0K2G9W6 | PilZ domain-containing protein | 11.698 | 106 | 0.61 | 1.53 | 8.34E-05 | 5.03E-04 | no | no | 0 | 5 | 6 | 5 | 3 | 3 | 3 | 0.018262436 | 0.018071778 | 0.019436937 | 0.012004108 | 0.012468602 | 0.012197736 |
| yjjK | A0A0K2GK24 | Energy-dependent translational throttle protein EttA | 63.069 | 560 | -0.485 | -1.40 | 9.38E-04 | 2.70E-03 | no | no | 0 | 26 | 24 | 23 | 42 | 37 | 35 | 0.018169901 | 0.014914721 | 0.013088969 | 0.029509777 | 0.030024888 | 0.02774681 |
| NITMOv2_0994 | A0A0K2G8Y9 | Uncharacterized protein | 36.411 | 345 | 0.0307 | 1.02 | 7.93E-01 | 8.19E-01 | no | no | 0 | 13 | 15 | 13 | 12 | 13 | 14 | 0.0181484 | 0.017811313 | 0.013168918 | 0.018128767 | 0.01540537 | 0.013743761 |
| NITMOv2_1350 | A0A0K2G9Z8 | Putative Serine protease | 36.058 | 336 | 0.398 | 1.32 | 3.73E-03 | 7.88E-03 | no | no | 0 | 11 | 12 | 13 | 11 | 12 | 13 | 0.01813765 | 0.018694194 | 0.018327624 | 0.014311109 | 0.016893175 | 0.015005502 |
| pabA | A0A0K2GDM0 | Aminodeoxychorismate synthase, subunit II | 20.715 | 187 | -0.0275 | -1.02 | 7.07E-01 | 7.42E-01 | no | no | 0 | 4 | 8 | 7 | 8 | 6 | 6 | 0.01793972 | 0.019235655 | 0.019154528 | 0.020626785 | 0.025648168 | 0.016984335 |
| NITMOv2_2519 | A0A0K2GEC1 | Putative Small-conductance mechanosensitive channel MscS | 60.161 | 554 | 0.392 | 1.31 | 6.49E-04 | 2.06E-03 | no | no | 0 | 21 | 19 | 23 | 18 | 18 | 13 | 0.017924122 | 0.020563398 | 0.020299693 | 0.017500931 | 0.017475806 | 0.016923379 |
| NITMOv2_4067 | A0A0K2GHQ1 | Uncharacterized protein | 15.943 | 137 | 0.0112 | 1.01 | 8.82E-01 | 8.98E-01 | no | no | 0 | 5 | 5 | 4 | 4 | 8 | 8 | 0.017836224 | 0.018173852 | 0.017357119 | 0.016876056 | 0.018993752 | 0.016840126 |
| alaS | A0A0K2GD80 | Alanine--tRNA ligase | 95.86 | 877 | -0.14 | -1.10 | 6.38E-02 | 8.68E-02 | no | no | 0 | 70 | 73 | 72 | 85 | 73 | 82 | 0.0178299 | 0.016396162 | 0.015485514 | 0.020264002 | 0.020002091 | 0.019585138 |
| NITMOv2_0555 | A0A0K2G7P9 | TPR_REGION domain-containing protein | 39.446 | 353 | -0.0237 | -1.02 | 7.04E-01 | 7.39E-01 | no | no | 0 | 22 | 25 | 29 | 31 | 28 | 27 | 0.01779512 | 0.01870006 | 0.017395081 | 0.021892822 | 0.023549145 | 0.020768301 |
| NITMOv2_0453 | A0A0K2G7P0 | Putative Lactoylglutathione lyase | 13.248 | 122 | 0.289 | 1.22 | 1.92E-02 | 3.08E-02 | no | no | 0 | 4 | 5 | 7 | 2 | 1 | 2 | 0.017793434 | 0.019260294 | 0.02166093 | 0.00929464 | 0.009191422 | 0.007864263 |
| NITMOv2_4136 | A0A0K2GHS5 | Transcriptional regulator | 26.662 | 234 | 0.38 | 1.30 | 7.39E-03 | 1.38E-02 | no | no | 0 | 10 | 12 | 13 | 12 | 11 | 10 | 0.017692466 | 0.018747578 | 0.017392397 | 0.015414561 | 0.013163719 | 0.011787764 |
| NITMOv2_1306 | A0A0K2GA33 | DUF262 domain-containing protein | 41.101 | 352 | 0.535 | 1.45 | 6.92E-04 | 2.17E-03 | no | no | 0 | 19 | 23 | 21 | 17 | 17 | 15 | 0.017692045 | 0.018600137 | 0.01883109 | 0.015925418 | 0.0131429 | 0.013008868 |
| NITMOv2_0746 | A0A0K2G995 | Uncharacterized protein | 93.487 | 836 | 0.0702 | 1.05 | 3.42E-01 | 3.84E-01 | no | no | 0 | 38 | 39 | 46 | 48 | 48 | 46 | 0.017689305 | 0.01779919 | 0.018620961 | 0.018198362 | 0.017609422 | 0.017184825 |
| glgB | A0A0K2GH17 | 1,4-alpha-glucan branching enzyme GlgB | 74.023 | 645 | 0.293 | 1.23 | 2.60E-02 | 3.97E-02 | no | no | 0 | 23 | 24 | 34 | 32 | 24 | 27 | 0.0175837 | 0.01479759 | 0.017884743 | 0.013501141 | 0.014399983 | 0.014521268 |
| NITMOv2_4347 | A0A0K2GID2 | Uncharacterized protein | 21.322 | 186 | -0.241 | -1.18 | 1.43E-02 | 2.41E-02 | no | no | 0 | 8 | 11 | 11 | 14 | 12 | 13 | 0.01756768 | 0.017609903 | 0.017658317 | 0.025994487 | 0.025388704 | 0.023287469 |
| NITMOv2_0470 | A0A0K2G7H4 | Putative Ribonuclease Z | 27.932 | 252 | 0.562 | 1.48 | 1.60E-03 | 4.07E-03 | no | no | 0 | 14 | 10 | 13 | 8 | 10 | 7 | 0.01750065 | 0.01848066 | 0.019011502 | 0.01306136 | 0.012940766 | 0.012571565 |
| NITMOv2_3298 | A0A0K2GFG2 | Uncharacterized protein | 30.189 | 279 | 0.347 | 1.27 | 4.65E-02 | 6.59E-02 | no | no | 0 | 9 | 6 | 8 | 6 | 9 | 8 | 0.017454487 | 0.020070628 | 0.018937689 | 0.017286223 | 0.01600447 | 0.013046809 |
| fmt | A0A0K2G7B3 | Methionyl-tRNA formyltransferase | 32.97 | 308 | 0.676 | 1.60 | 7.51E-04 | 2.30E-03 | no | no | 0 | 10 | 10 | 12 | 10 | 9 | 7 | 0.017449217 | 0.017796843 | 0.016104398 | 0.010190787 | 0.009042424 | 0.008261648 |
| metG | A0A0K2GGT8 | Methionine--tRNA ligase | 73.775 | 662 | 0.225 | 1.17 | 3.32E-02 | 4.93E-02 | no | no | 0 | 32 | 30 | 31 | 28 | 32 | 24 | 0.017399682 | 0.016687327 | 0.015638318 | 0.018641105 | 0.015934554 | 0.01666355 |
| NITMOv2_0799 | A0A0K2G8P6 | Peptidase S1C, HtrA family | 39.446 | 372 | 0.189 | 1.14 | 1.88E-02 | 3.03E-02 | no | no | 0 | 16 | 20 | 22 | 15 | 16 | 20 | 0.017355628 | 0.01731561 | 0.017850999 | 0.015581885 | 0.01482274 | 0.014352065 |
| NITMOv2_0502 | A0A0K2G7L3 | Uncharacterized protein | 10.076 | 89 | 0.114 | 1.08 | 1.82E-01 | 2.19E-01 | no | no | 0 | 11 | 12 | 11 | 10 | 12 | 11 | 0.017287332 | 0.02223921 | 0.02318705 | 0.020610497 | 0.02152936 | 0.019426903 |
| ureC | A0A0K2G9Q9 | Urease subunit alpha | 61.62 | 575 | 0.254 | 1.19 | 2.54E-02 | 3.91E-02 | no | no | 0 | 27 | 27 | 25 | 19 | 20 | 18 | 0.01725466 | 0.017358629 | 0.015455413 | 0.011764079 | 0.010461712 | 0.011588532 |
| argG | A0A0K2G8A9 | Argininosuccinate synthase | 45.209 | 404 | -0.276 | -1.21 | 1.39E-02 | 2.35E-02 | no | no | 0 | 22 | 23 | 22 | 25 | 24 | 28 | 0.017198802 | 0.01769868 | 0.016436463 | 0.020650477 | 0.021456337 | 0.020286405 |
| purB | A0A0K2G9V1 | Adenylosuccinate lyase | 51.025 | 450 | 0.127 | 1.09 | 1.73E-01 | 2.09E-01 | no | no | 0 | 36 | 39 | 36 | 32 | 32 | 29 | 0.017166762 | 0.019230376 | 0.017438602 | 0.018438243 | 0.020443337 | 0.017414086 |
| NITMOv2_1370 | A0A0K2GB31 | Uncharacterized protein | 26.258 | 232 | 0.428 | 1.35 | 8.40E-04 | 2.49E-03 | no | no | 0 | 16 | 15 | 16 | 10 | 12 | 10 | 0.017148212 | 0.01665428 | 0.016124337 | 0.011644731 | 0.013835065 | 0.013894084 |
| NITMOv2_2282 | A0A0K2GCM3 | PilZ domain-containing protein | 14.374 | 126 | -3.49 | -11.24 | 8.63E-04 | 2.54E-03 | yes | yes | 2 | 3 | 3 | 5 | 9 | 11 | 8 | 0.017130928 | 0.01614919 | 0.016159422 | 0.017773388 | 0.017537953 | 0.015760534 |
| dop.1 | A0A0K2GGI6 | Pup deamidase/depupylase | 56.341 | 500 | -0.487 | -1.40 | 2.76E-03 | 6.19E-03 | no | no | 0 | 25 | 32 | 25 | 45 | 43 | 42 | 0.017118491 | 0.019644342 | 0.016215022 | 0.032044813 | 0.032821514 | 0.030323518 |
| NITMOv2_3898 | A0A0K2GH47 | Uncharacterized protein | 15.421 | 143 | 0.508 | 1.42 | 5.54E-04 | 1.84E-03 | no | no | 0 | 8 | 7 | 9 | 5 | 5 | 7 | 0.01708877 | 0.016573716 | 0.015347856 | 0.011152087 | 0.011462127 | 0.01126541 |
| NITMOv2_4505 | A0A0K2GJ28 | Uncharacterized protein | 45.585 | 421 | 0.453 | 1.37 | 3.58E-03 | 7.63E-03 | no | no | 0 | 17 | 16 | 19 | 18 | 18 | 16 | 0.017035019 | 0.01719359 | 0.015447361 | 0.013734063 | 0.012085775 | 0.012545672 |
| NITMOv2_2515 | A0A0K2GDB2 | fn3_3 domain-containing protein | 34.548 | 312 | -0.357 | -1.28 | 2.60E-02 | 3.98E-02 | no | no | 0 | 19 | 17 | 14 | 25 | 23 | 23 | 0.016915081 | 0.016715485 | 0.013826913 | 0.024285707 | 0.022860864 | 0.02119086 |
| NITMOv2_3931 | A0A0K2GHA4 | Uncharacterized protein | 20.524 | 181 | 0.0639 | 1.05 | 6.00E-01 | 6.40E-01 | no | no | 0 | 9 | 11 | 11 | 9 | 9 | 7 | 0.01687735 | 0.016772193 | 0.019576895 | 0.017543873 | 0.016737807 | 0.013623107 |
| NITMOv2_4041 | A0A0K2GHK2 | Response regulatory domain-containing protein | 20.083 | 183 | -0.211 | -1.16 | 4.15E-02 | 5.96E-02 | no | no | 0 | 13 | 13 | 11 | 15 | 16 | 11 | 0.016870394 | 0.017261639 | 0.018097364 | 0.022260047 | 0.023022447 | 0.020131766 |
| NITMOv2_1764 | A0A0K2GBD7 | BFN domain-containing protein | 17.121 | 151 | -0.693 | -1.62 | 1.24E-04 | 6.45E-04 | no | no | 0 | 10 | 8 | 10 | 18 | 18 | 17 | 0.016788609 | 0.016544971 | 0.018769547 | 0.034094165 | 0.033104284 | 0.030548283 |
| NITMOv2_3208 | A0A0K2GG52 | NTP_transf_2 domain-containing protein | 11.748 | 105 | 0.735 | 1.66 | 8.24E-04 | 2.45E-03 | no | no | 0 | 7 | 8 | 11 | 9 | 7 | 4 | 0.016787344 | 0.017052016 | 0.020644795 | 0.014871127 | 0.012537119 | 0.011494311 |
| NITMOv2_3502 | A0A0K2GGB2 | C-methyltransferase | 48.175 | 425 | -0.426 | -1.34 | 1.53E-03 | 3.92E-03 | no | no | 0 | 29 | 30 | 29 | 38 | 41 | 42 | 0.016781442 | 0.016779428 | 0.017979646 | 0.025844932 | 0.024687994 | 0.022197808 |
| NITMOv2_4327 | A0A0K2GIA8 | Uncharacterized protein | 27.779 | 249 | 0.16 | 1.12 | 8.32E-02 | 1.10E-01 | no | no | 0 | 13 | 14 | 12 | 14 | 11 | 14 | 0.016767951 | 0.018551643 | 0.018859849 | 0.018152459 | 0.018518326 | 0.018506265 |
| gyrB | A0A0K2G792 | DNA gyrase subunit B | 91.19 | 822 | -0.622 | -1.54 | 1.21E-04 | 6.35E-04 | no | no | 0 | 39 | 39 | 47 | 70 | 64 | 60 | 0.016736755 | 0.016584275 | 0.016540185 | 0.032035929 | 0.030466134 | 0.029003697 |
| zraR.2 | A0A0K2GHG5 | Transcriptional regulatory protein ZraR | 51.553 | 459 | -0.00667 | -1.00 | 9.43E-01 | 9.50E-01 | no | no | 0 | 45 | 49 | 53 | 52 | 55 | 51 | 0.016691436 | 0.016928433 | 0.018277009 | 0.018521165 | 0.017214787 | 0.016675957 |
| NITMOv2_1951 | A0A0K2GBP1 | Uncharacterized protein | 23.151 | 211 | 0.693 | 1.62 | 3.00E-04 | 1.16E-03 | no | no | 0 | 10 | 12 | 10 | 6 | 5 | 4 | 0.016685744 | 0.011953013 | 0.014814098 | 0.009109694 | 0.008954797 | 0.008423839 |
| NITMOv2_3708 | A0A0K2GGL2 | BFN domain-containing protein | 18.187 | 169 | 1.04 | 2.06 | 2.83E-04 | 1.11E-03 | yes | no | 0 | 4 | 7 | 5 | 3 | 3 | 6 | 0.016673308 | 0.017622613 | 0.014169715 | 0.008065917 | 0.007157498 | 0.007661615 |
| ywlC | A0A0K2GES6 | Threonylcarbamoyl-AMP synthase | 37.765 | 358 | 3.05 | 8.28 | 9.06E-09 | 3.39E-06 | yes | no | 0 | 25 | 21 | 30 | 1 | 0 | 1 | 0.0166419 | 0.014011895 | 0.014553737 | 0.000829987 | 0.001396744 | 0.005181107 |
| NITMOv2_1453 | A0A0K2GAA9 | Cyclophil_like domain-containing protein | 14.511 | 132 | 0.767 | 1.70 | 6.43E-03 | 1.23E-02 | no | no | 0 | 5 | 6 | 5 | 9 | 11 | 10 | 0.016551051 | 0.016430578 | 0.017243619 | 0.012604254 | 0.012932687 | 0.011946179 |
| NITMOv2_2411 | A0A0K2GCZ0 | Putative Acriflavine resistance protein A | 42.995 | 392 | 0.17 | 1.13 | 1.17E-01 | 1.48E-01 | no | no | 0 | 14 | 14 | 16 | 17 | 17 | 13 | 0.016472427 | 0.018610892 | 0.01624589 | 0.016169741 | 0.017264505 | 0.014841334 |
| cfp | A0A0K2GF34 | Pyruvate decarboxylase | 60.134 | 551 | 0.446 | 1.36 | 5.27E-03 | 1.04E-02 | no | no | 0 | 24 | 31 | 25 | 24 | 24 | 21 | 0.016432167 | 0.018163879 | 0.018177888 | 0.011986339 | 0.012697925 | 0.012560956 |
| NITMOv2_2972 | A0A0K2GFK0 | Nitrogen regulatory protein P-II | 10.751 | 99 | 0.0301 | 1.02 | 9.33E-01 | 9.44E-01 | no | no | 0 | 6 | 6 | 7 | 8 | 9 | 3 | 0.016391906 | 0.014283701 | 0.012644746 | 0.010015911 | 0.009656905 | 0.008018542 |
| NITMOv2_3025 | A0A0K2GEQ2 | Uncharacterized protein | 34.556 | 317 | 0.443 | 1.36 | 8.35E-04 | 2.48E-03 | no | no | 0 | 11 | 9 | 10 | 8 | 7 | 8 | 0.016389588 | 0.016190841 | 0.016223074 | 0.009830373 | 0.012404124 | 0.012172382 |
| lepB | A0A0K2GFH6 | Signal peptidase I | 28.438 | 253 | -0.282 | -1.22 | 1.78E-02 | 2.88E-02 | no | no | 0 | 10 | 10 | 11 | 12 | 16 | 18 | 0.01638748 | 0.017932159 | 0.016239946 | 0.020110005 | 0.01975661 | 0.020241452 |
| NITMOv2_2331 | A0A0K2GCR0 | Putative 3-hydroxyisobutyrate dehydrogenase | 31.111 | 296 | 0.335 | 1.26 | 1.33E-03 | 3.51E-03 | no | no | 0 | 6 | 6 | 6 | 4 | 4 | 7 | 0.01637968 | 0.01864433 | 0.01750513 | 0.007070116 | 0.010865203 | 0.013929147 |
| oxaA | A0A0K2G780 | Membrane protein insertase YidC | 62.412 | 563 | 0.357 | 1.28 | 1.83E-03 | 4.48E-03 | no | no | 0 | 24 | 18 | 20 | 18 | 18 | 19 | 0.016279556 | 0.017309352 | 0.016147919 | 0.016551773 | 0.016733146 | 0.016486615 |
| NITMOv2_3703 | A0A0K2GGK7 | Putative Extracellular ligand-binding receptor | 71.871 | 651 | 0.169 | 1.12 | 8.30E-02 | 1.10E-01 | no | no | 0 | 36 | 40 | 46 | 43 | 43 | 39 | 0.016278713 | 0.014770214 | 0.015519257 | 0.015827689 | 0.015199507 | 0.014188616 |
| NITMOv2_4302 | A0A0K2GII3 | Putative Deoxyhypusine synthase | 40.569 | 368 | -1.22 | -2.33 | 7.98E-06 | 1.01E-04 | yes | no | 0 | 19 | 17 | 21 | 38 | 34 | 34 | 0.016269017 | 0.015588175 | 0.01654958 | 0.045256025 | 0.043215641 | 0.037551966 |

| | | | | | | | | | | | | | | | | | | | | | |
|---|---|---|---|---|---|---|---|---|---|---|---|---|---|---|---|---|---|---|---|---|---|
| NITMOv2_3175 | A0A0K2GFE9 | Uncharacterized protein | 41.268 | 376 | 0.513 | 1.43 | 2.76E-04 | 1.10E-03 | no | no | 0 | 17 | 17 | 18 | 11 | 10 | 12 | 0.01616826 | 0.016910834 | 0.016226526 | 0.011538857 | 0.010346274 | 0.011760792 |
| NITMOv2_0482 | A0A0K2G7J4 | Uncharacterized protein | 18.235 | 166 | 0.289 | 1.22 | 1.13E-02 | 1.99E-02 | no | no | 0 | 11 | 10 | 11 | 11 | 12 | 11 | 0.016137064 | 0.017910063 | 0.016432053 | 0.012393988 | 0.014587978 | 0.015044881 |
| ispF | A0A0K2GFY9 | 2-C-methyl-D-erythritol 2,4-cyclodiphosphate synthase | 16.329 | 157 | 0.0614 | 1.04 | 5.21E-01 | 5.63E-01 | no | no | 0 | 11 | 13 | 11 | 11 | 10 | 8 | 0.016109872 | 0.016932539 | 0.01705132 | 0.016816827 | 0.015970289 | 0.014267733 |
| thiD | A0A0K2G988 | Hydroxymethylpyrimidine/phosphomethylpyrimidine kinase | 28.128 | 266 | 0.0484 | 1.03 | 6.43E-01 | 6.82E-01 | no | no | 0 | 12 | 13 | 12 | 16 | 14 | 15 | 0.016096803 | 0.01718264 | 0.014558147 | 0.015204295 | 0.014372327 | 0.013168182 |
| msrA | A0A0K2GK10 | Peptide methionine sulfoxide reductase MsrA | 23.035 | 206 | 0.635 | 1.55 | 1.13E-04 | 6.07E-04 | no | no | 0 | 10 | 12 | 10 | 8 | 6 | 4 | 0.016083735 | 0.016706295 | 0.015848255 | 0.011093153 | 0.011648569 | 0.010392062 |
| glyQ | A0A0K2GF60 | Glycine--tRNA ligase alpha subunit | 35.402 | 307 | -0.0149 | -1.01 | 8.95E-01 | 9.10E-01 | no | no | 0 | 10 | 11 | 9 | 13 | 12 | 11 | 0.016081205 | 0.016002728 | 0.015526351 | 0.019542879 | 0.017244307 | 0.014060769 |
| pyrR | A0A0K2GFJ4 | Bifunctional protein PyrR | 20.906 | 186 | 0.253 | 1.19 | 1.65E-01 | 2.01E-01 | no | no | 0 | 9 | 11 | 13 | 11 | 13 | 8 | 0.016069401 | 0.014661883 | 0.013794512 | 0.014003558 | 0.01428268 | 0.011739934 |
| NITMOv2_4411 | A0A0K2GIL8 | Aldehyde dehydrogenase | 55.364 | 508 | 0.961 | 1.95 | 2.69E-05 | 2.29E-04 | no | no | 0 | 40 | 40 | 38 | 19 | 23 | 22 | 0.016043263 | 0.019716693 | 0.018178847 | 0.009985111 | 0.008438198 | 0.008073205 |
| pheT | A0A0K2G731 | Phenylalanine--tRNA ligase beta subunit | 63.163 | 571 | -0.222 | -1.17 | 4.64E-02 | 6.59E-02 | no | no | 0 | 33 | 28 | 38 | 39 | 41 | 38 | 0.015981924 | 0.015761818 | 0.016474616 | 0.020684534 | 0.0192672 | 0.015733562 |
| NITMOv2_1763 | A0A0K2GB60 | BFN domain-containing protein | 17.927 | 162 | -0.515 | -1.43 | 2.66E-04 | 1.07E-03 | no | no | 0 | 11 | 12 | 12 | 11 | 11 | 14 | 0.015973914 | 0.01689871 | 0.019077839 | 0.020249195 | 0.022102668 | 0.025963074 |
| NITMOv2_4287 | A0A0K2GIH1 | Uncharacterized protein | 39.732 | 351 | -0.275 | -1.21 | 9.70E-02 | 1.26E-01 | no | no | 0 | 17 | 17 | 22 | 23 | 18 | 21 | 0.015905408 | 0.011603772 | 0.014923572 | 0.018284245 | 0.019273415 | 0.012751377 |
| NITMOv2_4618 | A0A0K2GJ71 | Uncharacterized protein | 46.013 | 418 | 0.598 | 1.51 | 3.74E-04 | 1.36E-03 | no | no | 0 | 18 | 21 | 26 | 16 | 15 | 17 | 0.015898031 | 0.015936047 | 0.015449278 | 0.010796115 | 0.01140402 | 0.009922034 |
| NITMOv2_3925 | A0A0K2GH75 | Uncharacterized protein | 8.3405 | 73 | 1.18 | 2.27 | 2.09E-02 | 3.30E-02 | yes | yes | 1 | 2 | 2 | 1 | 2 | 2 | 1 | 0.015818774 | 0.015240497 | 0.015590387 | 0.008940149 | 0.004858361 | 0.009602867 |
| NITMOv2_3727 | A0A0K2GF74 | Uncharacterized protein | 13.634 | 114 | 0.733 | 1.66 | 5.27E-04 | 1.77E-03 | no | no | 0 | 5 | 5 | 4 | 2 | 3 | 2 | 0.015752798 | 0.016943685 | 0.015197737 | 0.009786691 | 0.011264965 | 0.009180848 |
| gpo | A0A0K2GAD2 | Glutathione peroxidase | 21.731 | 196 | -0.0551 | -1.04 | 5.23E-01 | 5.65E-01 | no | no | 0 | 12 | 14 | 11 | 14 | 12 | 13 | 0.015725817 | 0.017375446 | 0.015854965 | 0.020397269 | 0.018951803 | 0.018259922 |
| NITMOv2_3616 | A0A0K2GGC6 | Uncharacterized protein | 13.906 | 128 | 1.59 | 3.01 | 5.26E-02 | 7.36E-02 | no | yes | 1 | 2 | 2 | 2 | 1 | 0 | 0 | 0.015670801 | 0.016775517 | 0.015061996 | 0.01294986 | 0.009718431 | 0.008504574 |
| NITMOv2_1827 | A0A0K2GCB6 | Putative Vitamin B12 import system, periplasmic binding protein BtuF | 32.296 | 290 | -0.152 | -1.11 | 5.96E-02 | 8.17E-02 | no | no | 0 | 16 | 15 | 16 | 11 | 17 | 12 | 0.01567038 | 0.014030863 | 0.014708842 | 0.021309409 | 0.019804774 | 0.016000403 |
| der | A0A0K2GJ36 | GTPase Der | 49.194 | 450 | 0.649 | 1.57 | 4.59E-05 | 3.28E-04 | no | no | 0 | 25 | 27 | 29 | 21 | 19 | 19 | 0.015670169 | 0.014972798 | 0.017376484 | 0.01013363 | 0.010844694 | 0.010558209 |
| NITMOv2_0316 | A0A0K2G7A8 | Uncharacterized protein | 14.149 | 125 | -2.89 | -7.41 | 2.84E-08 | 4.25E-06 | yes | no | 0 | 4 | 2 | 3 | 14 | 13 | 14 | 0.015616418 | 0.013747911 | 0.017056113 | 0.107820484 | 0.139436669 | 0.132474791 |
| NITMOv2_3684 | A0A0K2GGK9 | Uncharacterized protein | 67.771 | 630 | -0.315 | -1.24 | 2.38E-03 | 5.51E-03 | no | no | 0 | 35 | 30 | 32 | 40 | 42 | 40 | 0.015594707 | 0.016052982 | 0.014404768 | 0.017382471 | 0.020867492 | 0.020111987 |
| NITMOv2_1380 | A0A0K2GB41 | Activator of Hsp90 ATPase 1 family protein | 16.54 | 148 | 0.306 | 1.24 | 7.68E-03 | 1.42E-02 | no | no | 0 | 6 | 7 | 3 | 6 | 5 | 4 | 0.015590281 | 0.015252621 | 0.017103277 | 0.011356874 | 0.011639402 | 0.011606514 |
| mtnA | A0A0K2GCA0 | Methylthioribose-1-phosphate isomerase | 36.372 | 342 | 0.222 | 1.17 | 5.68E-02 | 7.82E-02 | no | no | 0 | 19 | 19 | 21 | 23 | 19 | 15 | 0.015550231 | 0.012626075 | 0.016481326 | 0.01910754 | 0.016360263 | 0.011946898 |
| NITMOv2_1458 | A0A0K2GAB3 | Uncharacterized protein | 30.434 | 287 | 0.407 | 1.33 | 4.85E-03 | 9.70E-03 | no | no | 0 | 12 | 10 | 9 | 6 | 11 | 7 | 0.015548334 | 0.013915883 | 0.015160734 | 0.012899959 | 0.013722267 | 0.01116975 |
| NITMOv2_0310 | A0A0K2G717 | Putative Manganese transport system, periplasmic binding component | 34.545 | 314 | 0.392 | 1.31 | 6.80E-04 | 2.14E-03 | no | no | 0 | 16 | 18 | 15 | 14 | 15 | 14 | 0.015471185 | 0.016777864 | 0.01617361 | 0.011904009 | 0.0130966 | 0.012868795 |
| sqr | A0A0K2GHC5 | Sulfide:quinone oxidoreductase | 45.297 | 415 | 0.812 | 1.76 | 4.27E-05 | 3.10E-04 | no | no | 0 | 16 | 18 | 17 | 9 | 8 | 8 | 0.015435352 | 0.015016404 | 0.015693917 | 0.006105558 | 0.007969918 | 0.007268006 |
| acpP | A0A0K2GBD2 | Acyl carrier protein | 11.534 | 104 | 0.117 | 1.08 | 1.67E-01 | 2.03E-01 | no | no | 0 | 3 | 3 | 4 | 4 | 3 | 4 | 0.015395091 | 0.013736374 | 0.013308301 | 0.012169063 | 0.010537066 | 0.010894637 |
| rny | A0A0K2GHX4 | Ribonuclease Y | 59.613 | 533 | -0.856 | -1.81 | 6.59E-05 | 4.24E-04 | no | no | 0 | 24 | 21 | 19 | 38 | 34 | 37 | 0.015380547 | 0.015692791 | 0.015874904 | 0.034483601 | 0.033012617 | 0.033721968 |
| amtB | A0A0K2G9S7 | Ammonium transporter | 45.642 | 441 | 0.371 | 1.29 | 5.43E-02 | 7.54E-02 | no | no | 0 | 3 | 3 | 1 | 1 | 2 | 1 | 0.015371272 | 0.016530696 | 0.013616209 | 0.009212754 | 0.01259445 | 0.012670822 |
| MUR | A0A0K2GG12 | GDP-mannose 4,6-dehydratase | 38.975 | 342 | -0.534 | -1.45 | 9.52E-04 | 2.72E-03 | no | no | 0 | 11 | 13 | 17 | 25 | 22 | 26 | 0.015343237 | 0.016985727 | 0.014473213 | 0.022839019 | 0.025850147 | 0.026813585 |
| NITMOv2_0042 | A0A0K2G697 | HARE-HTH domain-containing protein | 19.328 | 170 | -0.697 | -1.62 | 9.08E-05 | 5.30E-04 | no | no | 0 | 14 | 15 | 15 | 22 | 20 | 22 | 0.015331855 | 0.015031461 | 0.015230521 | 0.029768908 | 0.027935187 | 0.028831078 |
| NITMOv2_4575 | A0A0K2GJ26 | Putative general secretion pathway protein D | 61.241 | 567 | 0.000887 | 1.00 | 9.92E-01 | 9.93E-01 | no | no | 0 | 42 | 43 | 51 | 54 | 50 | 53 | 0.015309511 | 0.015983955 | 0.016538268 | 0.018878024 | 0.019173979 | 0.015360452 |
| NITMOv2_3503 | A0A0K2GG09 | Methyltransf_21 domain-containing protein | 26.27 | 232 | -0.203 | -1.15 | 4.18E-02 | 6.00E-02 | no | no | 0 | 12 | 10 | 11 | 13 | 12 | 10 | 0.015302977 | 0.016800938 | 0.01552405 | 0.017455028 | 0.018667479 | 0.019182358 |
| NITMOv2_0737 | A0A0K2G889 | Uncharacterized protein | 67.438 | 611 | -0.0188 | -1.01 | 8.48E-01 | 8.66E-01 | no | no | 0 | 9 | 9 | 10 | 14 | 13 | 14 | 0.015294967 | 0.012227166 | 0.015689124 | 0.02133162 | 0.021697157 | 0.020478804 |
| ftsH.1 | A0A0K2GHB2 | ATP-dependent zinc metalloprotease FtsH | 66.018 | 601 | 0.646 | 1.56 | 1.33E-04 | 6.72E-04 | no | no | 0 | 24 | 24 | 27 | 23 | 19 | 20 | 0.015278736 | 0.015965379 | 0.015084236 | 0.010856974 | 0.011559077 | 0.010653509 |
| pyrH | A0A0K2GB29 | Uridylate kinase | 25.91 | 241 | -0.118 | -1.09 | 1.05E-01 | 1.35E-01 | no | no | 0 | 16 | 17 | 19 | 20 | 18 | 16 | 0.015268408 | 0.014315966 | 0.015442951 | 0.018556703 | 0.018442195 | 0.01860696 |
| NITMOv2_1496 | A0A0K2GBG9 | Putative_PNPOx domain-containing protein | 22.807 | 209 | -0.0392 | -1.03 | 7.53E-01 | 7.83E-01 | no | no | 0 | 18 | 16 | 16 | 9 | 11 | 11 | 0.015248383 | 0.015317933 | 0.014197707 | 0.010247795 | 0.010961842 | 0.010584282 |
| nuoCD | A0A0K2G8R0 | NADH-quinone oxidoreductase, subunits C and D | 66.312 | 583 | 0.0559 | 1.04 | 4.04E-01 | 4.48E-01 | no | no | 0 | 24 | 23 | 26 | 25 | 31 | 25 | 0.015227936 | 0.014034578 | 0.011963935 | 0.014088553 | 0.013776802 | 0.012141994 |
| dxr | A0A0K2GA35 | 1-deoxy-D-xylulose 5-phosphate reductoisomerase | 42.543 | 386 | -0.433 | -1.35 | 7.18E-04 | 2.23E-03 | no | no | 0 | 19 | 21 | 20 | 37 | 35 | 27 | 0.015226461 | 0.015355477 | 0.014894622 | 0.023456489 | 0.021526252 | 0.020160536 |
| NITMOv2_3934 | A0A0K2GHH7 | Putative Phosphoribosyltransferase | 24.041 | 221 | 1.69 | 3.23 | 1.14E-05 | 1.30E-04 | yes | no | 0 | 12 | 9 | 13 | 3 | 5 | 5 | 0.015225407 | 0.013256313 | 0.015574857 | 0.003360108 | 0.004572328 | 0.004474445 |
| NITMOv2_0347 | A0A0K2G773 | Uncharacterized protein | 44.381 | 397 | 0.507 | 1.42 | 3.82E-03 | 8.01E-03 | no | no | 0 | 11 | 10 | 12 | 14 | 10 | 9 | 0.015223721 | 0.016112232 | 0.013684654 | 0.012697689 | 0.011711804 | 0.010664658 |
| NITMOv2_3033 | A0A0K2GEQ9 | Putative Metallophosphoesterase | 37.72 | 336 | 0.307 | 1.24 | 1.97E-03 | 4.75E-03 | no | no | 0 | 15 | 15 | 18 | 11 | 12 | 11 | 0.015191681 | 0.015415509 | 0.015354758 | 0.013249711 | 0.013312872 | 0.011308385 |
| NITMOv2_4333 | A0A0K2GIL5 | Putative Thioredoxin | 11.95 | 109 | -0.693 | -1.62 | 4.24E-04 | 1.50E-03 | no | no | 0 | 6 | 6 | 6 | 6 | 7 | 8 | 0.015177137 | 0.01474538 | 0.016154437 | 0.036925351 | 0.033714881 | 0.031172231 |
| lysA | A0A0K2G781 | Diaminopimelate decarboxylase | 46.056 | 420 | -1.47 | -2.77 | 1.87E-06 | 4.08E-05 | yes | no | 0 | 14 | 13 | 15 | 45 | 39 | 40 | 0.015156479 | 0.013615136 | 0.013510953 | 0.034662771 | 0.035790599 | 0.035187436 |
| secA | A0A0K2G7Q5 | Protein translocase subunit SecA | 102.51 | 907 | -0.557 | -1.47 | 2.42E-04 | 1.01E-03 | no | no | 0 | 52 | 52 | 49 | 73 | 75 | 70 | 0.015144254 | 0.015269633 | 0.01593453 | 0.026151446 | 0.02631159 | 0.023612929 |
| manB | A0A0K2GFY5 | Bifunctional Phosphomannomutase/phosphoglucomutase | 51.894 | 477 | -0.339 | -1.26 | 7.36E-03 | 1.38E-02 | no | no | 0 | 14 | 16 | 18 | 21 | 25 | 20 | 0.015122543 | 0.01494288 | 0.015133893 | 0.024713642 | 0.024548162 | 0.021848972 |
| NITMOv2_3534 | A0A0K2GG66 | Putative Oxidoreductase, Gfo/Idh/MocA family | 34.25 | 317 | -0.182 | -1.13 | 1.11E-01 | 1.41E-01 | no | no | 0 | 13 | 15 | 16 | 19 | 20 | 19 | 0.015016516 | 0.016199249 | 0.01598227 | 0.02216676 | 0.020068899 | 0.018191594 |
| paeR7IR | A0A0K2GCR1 | Type-2 restriction enzyme PaeR7I | 28.215 | 252 | 0.489 | 1.40 | 8.14E-03 | 1.50E-02 | no | no | 0 | 9 | 10 | 13 | 11 | 9 | 10 | 0.015006609 | 0.012836481 | 0.012350833 | 0.006543711 | 0.006825321 | 0.007598142 |
| glyS | A0A0K2GG37 | Glycine--tRNA ligase beta subunit | 80.556 | 736 | -0.0997 | -1.07 | 1.39E-01 | 1.73E-01 | no | no | 0 | 44 | 37 | 48 | 65 | 61 | 60 | 0.014995016 | 0.01440748 | 0.015418219 | 0.014379223 | 0.014441156 | 0.013586065 |
| NITMOv2_2389 | A0A0K2GD62 | DUF5069 domain-containing protein | 16.536 | 146 | 0.342 | 1.27 | 7.04E-03 | 1.32E-02 | no | no | 0 | 8 | 9 | 12 | 9 | 11 | 8 | 0.014995016 | 0.016517986 | 0.014964026 | 0.010664181 | 0.010932011 | 0.010849505 |
| NITMOv2_0188 | A0A0K2G6Q5 | Peptidase M24 | 45.349 | 403 | 0.577 | 1.49 | 3.61E-04 | 1.32E-03 | no | no | 0 | 30 | 30 | 27 | 15 | 22 | 15 | 0.014986374 | 0.015940936 | 0.014264426 | 0.008476231 | 0.00997339 | 0.009320023 |
| NITMOv2_0262 | A0A0K2G6V1 | Cofac_haem_bdg domain-containing protein | 36.46 | 329 | 0.341 | 1.27 | 4.09E-03 | 8.47E-03 | no | no | 0 | 13 | 14 | 14 | 12 | 9 | 9 | 0.014981315 | 0.016232883 | 0.017718135 | 0.008070655 | 0.010850288 | 0.008235396 |
| NITMOv2_3656 | A0A0K2GGG0 | Uncharacterized protein | 48.449 | 455 | 0.239 | 1.18 | 9.54E-03 | 1.72E-02 | no | no | 0 | 36 | 31 | 30 | 28 | 30 | 26 | 0.014950961 | 0.013607706 | 0.014119675 | 0.013440431 | 0.013880587 | 0.01222219 |
| NITMOv2_2613 | A0A0K2GDJ4 | Uncharacterized protein | 24.16 | 232 | 2.08 | 4.23 | 2.43E-07 | 1.33E-05 | yes | no | 0 | 8 | 10 | 10 | 4 | 3 | 2 | 0.014944005 | 0.01536193 | 0.015746833 | 0.002338393 | 0.003517689 | 0.002506761 |
| NITMOv2_3875 | A0A0K2GI12 | Uncharacterized protein | 25.701 | 224 | 0.716 | 1.64 | 1.04E-04 | 5.73E-04 | no | no | 0 | 11 | 14 | 12 | 8 | 10 | 8 | 0.014926932 | 0.014876004 | 0.014572143 | 0.009594046 | 0.009552342 | 0.010286693 |
| NITMOv2_2410 | A0A0K2GCY9 | Putative Heavy metal efflux pump, CzcA family, Acriflavine resistance protein B | 112.88 | 1025 | 0.628 | 1.55 | 3.98E-05 | 2.99E-04 | no | no | 0 | 24 | 28 | 26 | 23 | 22 | 21 | 0.014925456 | 0.015090124 | 0.01536492 | 0.010486047 | 0.010029789 | 0.010152553 |
| argB | A0A0K2G6U8 | Acetylglutamate kinase | 32.111 | 298 | -0.701 | -1.63 | 3.24E-05 | 2.61E-04 | no | no | 0 | 25 | 27 | 22 | 30 | 32 | 35 | 0.014914074 | 0.013993123 | 0.014149584 | 0.02329953 | 0.021732892 | 0.021361681 |
| NITMOv2_2063 | A0A0K2GC07 | Uncharacterized protein | 13.289 | 121 | 0.934 | 1.91 | 3.61E-04 | 1.32E-03 | no | no | 0 | 6 | 6 | 7 | 4 | 2 | 2 | 0.014907539 | 0.019194591 | 0.014907084 | 0.013133916 | 0.006520644 | 0.007864983 |
| NITMOv2_0582 | A0A0K2G7T6 | Uncharacterized protein | 13.455 | 123 | 0.39 | 1.31 | 1.80E-02 | 2.91E-02 | no | no | 0 | 3 | 5 | 5 | 1 | 4 | 4 | 0.014868543 | 0.016621624 | 0.014924531 | 0.006961873 | 0.011292931 | 0.012818447 |
| rnhC | A0A0K2G8L7 | Ribonuclease | 23.738 | 217 | 0.534 | 1.45 | 1.04E-04 | 5.73E-04 | no | no | 0 | 9 | 12 | 11 | 6 | 6 | 7 | 0.014855264 | 0.018268886 | 0.017302095 | 0.010076917 | 0.011062831 | 0.013448329 |
| purM | A0A0K2G9D1 | Phosphoribosylformylglycinamidine cyclo-ligase | 36.935 | 345 | -0.147 | -1.11 | 1.01E-01 | 1.30E-01 | no | no | 0 | 18 | 16 | 16 | 18 | 17 | 16 | 0.014852734 | 0.013581503 | 0.015060463 | 0.016884941 | 0.018154764 | 0.01400269 |
| glgC3 | A0A0K2GHD2 | Glucose-1-phosphate adenylyltransferase | 46.897 | 418 | 0.538 | 1.45 | 9.57E-04 | 2.72E-03 | no | no | 0 | 29 | 25 | 30 | 25 | 20 | 16 | 0.014822592 | 0.015850399 | 0.016263145 | 0.01180243 | 0.011076038 | 0.01052872 |
| NITMOv2_4555 | A0A0K2GJA1 | Uncharacterized protein | 17.997 | 165 | -1.2 | -2.30 | 1.90E-06 | 4.10E-05 | yes | no | 0 | 7 | 6 | 6 | 10 | 9 | 10 | 0.014813528 | 0.014756331 | 0.014368724 | 0.03636711 | 0.037114335 | 0.035539868 |
| NITMOv2_2458 | A0A0K2GD38 | Uncharacterized protein | 36.154 | 333 | 0.273 | 1.21 | 1.23E-02 | 2.13E-02 | no | no | 0 | 14 | 13 | 12 | 14 | 11 | 9 | 0.014718884 | 0.013392803 | 0.013926993 | 0.012932683 | 0.011701705 | 0.010014637 |
| yacO | A0A0K2GJM8 | Putative TrmH family tRNA/rRNA methyltransferase YacO | 27.229 | 256 | 0.739 | 1.67 | 1.04E-04 | 5.73E-04 | no | no | 0 | 14 | 15 | 13 | 9 | 10 | 10 | 0.014710031 | 0.016069213 | 0.012087597 | 0.010095279 | 0.009825635 | 0.007626012 |
| comD | A0A0K2G8W0 | Sulfopyruvate decarboxylase, alpha subunit | 18.102 | 165 | -0.269 | -1.20 | 3.30E-02 | 4.90E-02 | no | no | 0 | 9 | 8 | 12 | 10 | 10 | 9 | 0.014686634 | 0.015159933 | 0.014732232 | 0.01720182 | 0.017447839 | 0.015571731 |
| aroF | A0A0K2G860 | Phospho-2-dehydro-3-deoxyheptonate aldolase | 36.888 | 337 | -1.62 | -3.07 | 1.28E-06 | 3.23E-05 | yes | no | 0 | 15 | 20 | 19 | 54 | 54 | 51 | 0.014670825 | 0.016025802 | 0.014947921 | 0.05857385 | 0.061146675 | 0.052451199 |
| pheA | A0A0K2G777 | Bifunctional chorismate mutase/prephenate dehydratase | 40.101 | 358 | -0.304 | -1.23 | 8.32E-03 | 1.52E-02 | no | no | 0 | 14 | 16 | 16 | 22 | 24 | 23 | 0.014636466 | 0.013414313 | 0.018439016 | 0.023770408 | 0.024768785 | 0.022748033 |
| NITMOv2_4420 | A0A0K2GIM0 | Uncharacterized protein | 28.139 | 250 | 0.173 | 1.13 | 4.60E-02 | 6.54E-02 | no | no | 0 | 15 | 16 | 15 | 11 | 16 | 12 | 0.014627613 | 0.016531087 | 0.015996265 | 0.014346499 | 0.014681976 | 0.012742027 |
| ileS | A0A0K2G8W7 | Isoleucine--tRNA ligase | 105.51 | 937 | 0.178 | 1.13 | 2.74E-02 | 4.17E-02 | no | no | 0 | 28 | 30 | 37 | 39 | 42 | 39 | 0.01461349 | 0.01510127 | 0.014561598 | 0.013989195 | 0.014780169 | 0.014321857 |
| clpX | A0A0K2G6R0 | ATP-dependent Clp protease ATP-binding subunit ClpX | 46.613 | 418 | 0.112 | 1.08 | 2.64E-01 | 3.06E-01 | no | no | 0 | 15 | 16 | 17 | 19 | 15 | 18 | 0.01452517 | 0.015817939 | 0.015002562 | 0.014285196 | 0.013026218 | 0.012636118 |
| NITMOv2_0317 | A0A0K2G742 | Uncharacterized protein | 13.817 | 129 | 0.749 | 1.68 | 1.58E-02 | 2.62E-02 | no | yes | 3 | 1 | 1 | 1 | 0 | 1 | 0 | 0.014502827 | 0.012637417 | 0.01082874 | 0.011113291 | 0.010943353 | 0.009489406 |
| NITMOv2_4439 | A0A0K2GIM5 | Putative Peptidase M | 81.277 | 733 | 0.355 | 1.28 | 9.40E-03 | 1.69E-02 | no | no | 0 | 45 | 44 | 41 | 34 | 32 | 31 | 0.014472052 | 0.018056525 | 0.01394444 | 0.011725579 | 0.010836615 | 0.010936893 |
| aroC | A0A0K2GA01 | Chorismate synthase | 41.093 | 386 | 0.492 | 1.41 | 4.15E-04 | 1.48E-03 | no | no | 0 | 15 | 14 | 13 | 14 | 14 | 17 | 0.014442541 | 0.014532824 | 0.012781828 | 0.008836349 | 0.010365695 | 0.00955234 |
| yrbF | A0A0K2G874 | Toluene transporter subunit: ATP-binding component of ABC superfamily | 27.381 | 252 | -0.0123 | -1.01 | 9.22E-01 | 9.35E-01 | no | no | 0 | 25 | 26 | 24 | 20 | 20 | 20 | 0.014426943 | 0.018880156 | 0.016564342 | 0.011340734 | 0.013208154 | 0.01255754 |
| porE | A0A0K2G8E5 | Pyruvate:ferredoxin oxidoreductase epsilon subunit | 9.8655 | 92 | -1.07 | -2.10 | 2.53E-05 | 2.22E-04 | yes | no | 0 | 5 | 6 | 4 | 13 | 11 | 10 | 0.01442062 | 0.017684405 | 0.012430782 | 0.040850807 | 0.03253253 | 0.033074644 |
| NITMOv2_1088 | A0A0K2G9I6 | FlgO domain-containing protein | 34.696 | 322 | 0.348 | 1.27 | 3.26E-02 | 4.86E-02 | no | no | 0 | 14 | 13 | 12 | 16 | 13 | 10 | 0.014368977 | 0.014520309 | 0.013896892 | 0.012460177 | 0.012242541 | 0.01049006 |
| NITMOv2_3487 | A0A0K2GG94 | UDP-N-acetylglucosamine 2-epimerase | 43.209 | 376 | -0.38 | -1.30 | 8.05E-04 | 2.41E-03 | no | no | 0 | 16 | 18 | 22 | 33 | 32 | 32 | 0.014331667 | 0.012749463 | 0.013077274 | 0.023816311 | 0.022279788 | 0.022366831 |
| treT | A0A0K2GH21 | Trehalose synthase | 44.858 | 401 | 0.439 | 1.36 | 2.21E-02 | 3.47E-02 | no | no | 0 | 25 | 19 | 26 | 17 | 19 | 13 | 0.014288456 | 0.01135856 | 0.015079827 | 0.010558308 | 0.010669128 | 0.009108384 |
| NITMOv2_2369 | A0A0K2GD43 | Uncharacterized protein | 16.855 | 145 | 0.48 | 1.39 | 7.64E-03 | 1.42E-02 | no | no | 0 | 7 | 7 | 7 | 13 | 10 | 11 | 0.01428424 | 0.013995469 | 0.012221995 | 0.01208466 | 0.012978831 | 0.013215293 |
| NITMOv2_4092 | A0A0K2GHZ9 | Putative Histidine kinase | 40.811 | 382 | 1.52 | 2.87 | 5.01E-07 | 1.87E-05 | yes | no | 0 | 19 | 21 | 19 | 8 | 6 | 6 | 0.0142815 | 0.015812072 | 0.015761021 | 0.004926144 | 0.00406241 | 0.005177331 |

Table S1

| | | | | | | | | | | | | | | | | | | | | | | | |
|---|---|---|---|---|---|---|---|---|---|---|---|---|---|---|---|---|---|---|---|---|---|---|---|
| NITMOv2_2434 | A0A0K2GE16 | Peptidyl-prolyl cis-trans isomerase | 19.275 | 179 | -0.442 | -1.36 | 1.68E-02 | 2.75E-02 | no | no | 0 | 6 | 7 | 7 | 7 | 12 | 12 | 0.014281289 | 0.014843152 | 0.013605856 | 0.021913553 | 0.023081487 | 0.020322367 |
| NITMOv2_0604 | A0A0K2G7Y4 | Putative Sugar nucleotidyltransferase | 27.04 | 239 | 0.00942 | 1.01 | 8.69E-01 | 8.86E-01 | no | no | 0 | 13 | 15 | 14 | 15 | 18 | 16 | 0.014247774 | 0.014541623 | 0.015411125 | 0.017264012 | 0.016789079 | 0.016612304 |
| ald | A0A0K2G9F4 | Alanine dehydrogenase | 38.805 | 367 | 0.354 | 1.28 | 1.64E-02 | 2.70E-02 | no | no | 0 | 13 | 14 | 17 | 16 | 14 | 15 | 0.014234705 | 0.013859371 | 0.015029595 | 0.012609584 | 0.01078581 | 0.010611613 |
| glk | A0A0K2G7X1 | Glucokinase | 38.398 | 355 | 0.0472 | 1.03 | 5.41E-01 | 5.82E-01 | no | no | 0 | 29 | 31 | 30 | 28 | 27 | 33 | 0.014189807 | 0.01469141 | 0.015900787 | 0.018580395 | 0.015786954 | 0.015395335 |
| nifA | A0A0K2GIZ6 | Nif-specific regulatory protein NifA | 56.731 | 508 | -0.394 | -1.31 | 2.23E-03 | 5.22E-03 | no | no | 0 | 16 | 17 | 17 | 40 | 34 | 26 | 0.014165566 | 0.013620612 | 0.014704624 | 0.020135177 | 0.019352652 | 0.015622078 |
| gnd | A0A0K2GAZ1 | 6-phosphogluconate dehydrogenase (Decarboxylating) | 32.411 | 297 | -0.0713 | -1.05 | 5.98E-01 | 6.38E-01 | no | no | 0 | 18 | 18 | 28 | 32 | 29 | 26 | 0.014133105 | 0.014806194 | 0.01485551 | 0.021407138 | 0.018087956 | 0.017679309 |
| NITMOv2_2022 | A0A0K2GC65 | Putative 2,5-diketo-D-gluconic acid reductase | 31.405 | 279 | 0.463 | 1.38 | 2.50E-03 | 5.72E-03 | no | no | 0 | 8 | 5 | 8 | 9 | 10 | 9 | 0.014132051 | 0.012967299 | 0.01460742 | 0.012069409 | 0.011696267 | 0.010179704 |
| NITMOv2_4833 | A0A0K2GJT5 | Putative Lipoprotein, TraT related | 24.349 | 236 | 0.438 | 1.35 | 1.68E-03 | 4.22E-03 | no | no | 0 | 6 | 7 | 7 | 4 | 5 | 5 | 0.014083148 | 0.013502308 | 0.012458582 | 0.009181659 | 0.009657371 | 0.009305638 |
| NITMOv2_4510 | A0A0K2GJ34 | Uncharacterized protein | 15.681 | 143 | 0.561 | 1.48 | 1.75E-02 | 2.84E-02 | no | no | 0 | 5 | 6 | 3 | 5 | 5 | 4 | 0.014019491 | 0.01963652 | 0.008256768 | 0.011497545 | 0.010749609 | 0.009664183 |
| purU | A0A0K2G6V3 | Formyltetrahydrofolate deformylase | 32.89 | 289 | -0.15 | -1.11 | 1.07E-01 | 1.37E-01 | no | no | 0 | 15 | 11 | 16 | 16 | 15 | 13 | 0.013962789 | 0.01539967 | 0.015278836 | 0.016991555 | 0.016736253 | 0.016451552 |
| prpB | A0A0K2G6D4 | Methylisocitrate lyase | 32.837 | 305 | 1.65 | 3.14 | 2.63E-06 | 5.00E-05 | yes | no | 0 | 30 | 30 | 35 | 11 | 14 | 14 | 0.013902503 | 0.01514253 | 0.024335474 | 0.003955812 | 0.004276507 | 0.004135859 |
| cbbQ | A0A0K2GIM7 | Protein CbbQ | 30.133 | 272 | 0.154 | 1.11 | 1.30E-01 | 1.62E-01 | no | no | 0 | 16 | 15 | 16 | 16 | 15 | 14 | 0.013874469 | 0.016499409 | 0.015914975 | 0.012692802 | 0.014913785 | 0.014850684 |
| NITMOv2_4288 | A0A0K2GI92 | Uncharacterized protein | 32.359 | 291 | 0.602 | 1.52 | 1.23E-04 | 6.38E-04 | no | no | 0 | 11 | 11 | 9 | 11 | 9 | 7 | 0.013850228 | 0.01393094 | 0.014598792 | 0.009201056 | 0.011455757 | 0.011844765 |
| hisA | A0A0K2G9K1 | 1-(5-phosphoribosyl)-5-[(5-phosphoribosylamino)methylideneamino] imidazole-4-carboxamid | 25.631 | 240 | 0.348 | 1.27 | 1.07E-03 | 2.96E-03 | no | no | 0 | 12 | 14 | 11 | 11 | 9 | 12 | 0.013831889 | 0.014080335 | 0.013278392 | 0.010814032 | 0.013194326 | 0.01239445 |
| nth | A0A0K2GC23 | Endonuclease III | 24.645 | 221 | 0.141 | 1.10 | 2.84E-01 | 3.27E-01 | no | no | 0 | 8 | 8 | 10 | 6 | 9 | 9 | 0.013807438 | 0.014927041 | 0.013376938 | 0.014069451 | 0.027343235 | 0.012521038 |
| modB | A0A0K2GAJ2 | Molybdenum transport system permease | 24.574 | 224 | 1.14 | 2.20 | 1.57E-04 | 7.57E-04 | yes | no | 0 | 3 | 3 | 3 | 2 | 2 | 1 | 0.013782354 | 0.012511487 | 0.012673504 | 0.008036746 | 0.0071286 | 0.00632705 |
| NITMOv2_3138 | A0A0K2GG02 | Uncharacterized protein | 25.257 | 220 | 0.0783 | 1.06 | 3.70E-01 | 4.13E-01 | no | no | 0 | 29 | 29 | 31 | 24 | 27 | 26 | 0.013769075 | 0.014008571 | 0.015307211 | 0.013647883 | 0.013906223 | 0.012475366 |
| NITMOv2_0607 | A0A0K2G7W0 | Putative Heat shock protein Hsp20 | 16.704 | 146 | 2.72 | 6.59 | 2.63E-04 | 1.06E-03 | yes | yes | 3 | 3 | 2 | 4 | 0 | 0 | 0 | 0.0137383 | 0.005470732 | 0.016917497 | 0 | 0.000173127 | 0.000225808 |
| yfdZ | A0A0K2GIQ6 | Putative aminotransferase, PLP-dependent | 44.624 | 398 | 0.446 | 1.36 | 2.12E-03 | 5.06E-03 | no | no | 0 | 15 | 15 | 14 | 16 | 15 | 13 | 0.013733241 | 0.014037902 | 0.012886317 | 0.011542115 | 0.013260358 | 0.010813722 |
| NITMOv2_0922 | A0A0K2G8Q8 | SnoaL-like domain-containing protein | 15.132 | 130 | 0.417 | 1.34 | 1.94E-03 | 4.69E-03 | no | no | 0 | 3 | 2 | 2 | 2 | 1 | 1 | 0.013718064 | 0.012649345 | 0.012951887 | 0.010751101 | 0.010231146 | 0.010320677 |
| NITMOv2_0920 | A0A0K2G908 | Uncharacterized protein | 14.188 | 129 | 0.95 | 1.93 | 4.05E-02 | 5.84E-02 | no | no | 0 | 2 | 1 | 1 | 3 | 1 | 3 | 0.013710054 | 0.01196455 | 0.00982986 | 0.009203574 | 0.008824598 | 0.001568052 |
| exbB | A0A0K2G8P5 | Biopolymer transport protein exbB | 15.487 | 141 | 0.419 | 1.34 | 1.21E-02 | 2.10E-02 | no | no | 0 | 4 | 5 | 3 | 1 | 1 | 1 | 0.013694456 | 0.014072123 | 0.017860969 | 0.011545225 | 0.011128863 | 0.010634629 |
| NITMOv2_4601 | A0A0K2GJ50 | Putative Hydrolase with N-terminal DNA-binding domain | 30.298 | 280 | 0.63 | 1.55 | 4.69E-04 | 1.62E-03 | no | no | 0 | 11 | 12 | 13 | 10 | 11 | 9 | 0.013668318 | 0.01352851 | 0.014989333 | 0.009808754 | 0.009563684 | 0.008018003 |
| NITMOv2_2958 | A0A0K2GEI4 | Response regulator receiver domain protein | 16.651 | 148 | 3.28 | 9.71 | 5.48E-05 | 3.74E-04 | yes | yes | 1 | 7 | 10 | 11 | 0 | 0 | 0 | 0.013663259 | 0.012683761 | 0.013936004 | 0.000113129 | 0.000450894 | 0.000265565 |
| NITMOv2_0796 | A0A0K2G8D0 | Uncharacterized protein | 23.032 | 212 | 0.67 | 1.59 | 9.50E-05 | 5.41E-04 | no | no | 0 | 11 | 11 | 12 | 7 | 5 | 6 | 0.01360993 | 0.013081888 | 0.01348948 | 0.00895392 | 0.007447415 | 0.007861566 |
| capD | A0A0K2GG11 | UDP-galactose 4-epimerase | 38.484 | 341 | -1.3 | -2.46 | 4.20E-06 | 6.74E-05 | yes | no | 0 | 12 | 9 | 14 | 33 | 28 | 34 | 0.013603817 | 0.012723065 | 0.012755178 | 0.035182513 | 0.036901481 | 0.038012285 |
| NITMOv2_3773 | A0A0K2GH35 | TPR_REGION domain-containing protein | 66.333 | 592 | -0.447 | -1.36 | 2.94E-03 | 6.51E-03 | no | no | 0 | 32 | 24 | 28 | 32 | 32 | 34 | 0.013522242 | 0.013896524 | 0.01283992 | 0.019473284 | 0.021692496 | 0.020493189 |
| aspS | A0A0K2GA72 | Aspartate--tRNA(Asp/Asn) ligase | 65.733 | 587 | -0.528 | -1.44 | 4.42E-04 | 1.55E-03 | no | no | 0 | 41 | 40 | 43 | 59 | 61 | 58 | 0.013519713 | 0.013347632 | 0.013522456 | 0.02339726 | 0.021296308 | 0.018761598 |
| NITMOv2_4336 | A0A0K2GJ99 | Uncharacterized protein | 24.499 | 225 | 0.458 | 1.37 | 2.48E-03 | 5.69E-03 | no | no | 0 | 21 | 20 | 20 | 13 | 11 | 10 | 0.013482193 | 0.01453126 | 0.013571537 | 0.008795924 | 0.008060964 | 0.007527835 |
| kdsB | A0A0K2GFS3 | 3-deoxy-manno-octulosonate cytidylyltransferase | 29.537 | 266 | -0.248 | -1.19 | 2.15E-02 | 3.38E-02 | no | no | 0 | 10 | 10 | 10 | 10 | 13 | 11 | 0.01345205 | 0.015827129 | 0.015020968 | 0.020523133 | 0.022021877 | 0.018819138 |
| fabF.1 | A0A0K2G906 | 3-oxoacyl-[acyl-carrier-protein] synthase 2 | 43.661 | 416 | -0.532 | -1.45 | 9.44E-05 | 5.41E-04 | no | no | 0 | 15 | 18 | 21 | 25 | 27 | 25 | 0.013431393 | 0.014572128 | 0.015834259 | 0.031411054 | 0.030492546 | 0.024276436 |
| NITMOv2_3986 | A0A0K2GHD6 | fn3_3 domain-containing protein | 34.441 | 309 | -0.0591 | -1.04 | 5.54E-01 | 5.95E-01 | no | no | 0 | 15 | 15 | 13 | 11 | 11 | 16 | 0.013412422 | 0.014988246 | 0.013400903 | 0.01408959 | 0.015029379 | 0.014780557 |
| zwf2 | A0A0K2GC32 | Glucose-6-phosphate 1-dehydrogenase | 55.77 | 488 | 0.846 | 1.80 | 2.13E-04 | 9.20E-04 | no | no | 0 | 27 | 34 | 30 | 21 | 20 | 15 | 0.013396191 | 0.013621003 | 0.015632566 | 0.005782459 | 0.005660527 | 0.005195851 |
| dnaJ.1 | A0A0K2GHP4 | Chaperone protein DnaJ | 37.542 | 345 | 0.587 | 1.50 | 3.01E-04 | 1.16E-03 | no | no | 0 | 17 | 17 | 22 | 12 | 18 | 14 | 0.013393873 | 0.013902781 | 0.013614675 | 0.008489706 | 0.010068631 | 0.007653883 |
| nadC | A0A0K2GD52 | Quinolinate phosphoribosyltransferase [decarboxylating] | 31.015 | 293 | 0.364 | 1.29 | 6.81E-04 | 2.14E-03 | no | no | 0 | 10 | 10 | 9 | 7 | 7 | 6 | 0.013329793 | 0.010829027 | 0.011357897 | 0.008670505 | 0.009832471 | 0.009245221 |
| NITMOv2_0458 | A0A0K2G7P6 | DNA-binding protein Fis / transcriptional regulator, Fis family | 71.261 | 647 | 0.559 | 1.47 | 2.78E-04 | 1.10E-03 | no | no | 0 | 28 | 31 | 31 | 21 | 19 | 19 | 0.013311876 | 0.011499742 | 0.011487119 | 0.008175788 | 0.006706309 | 0.007806184 |
| NITMOv2_4830 | A0A0K2GJR6 | Putative Peptidase U62, TldE-like | 48.716 | 448 | 0.496 | 1.41 | 6.50E-04 | 2.06E-03 | no | no | 0 | 22 | 27 | 22 | 15 | 16 | 18 | 0.013296699 | 0.01480815 | 0.013797388 | 0.010735109 | 0.009994209 | 0.010650093 |
| NITMOv2_2687 | A0A0K2GE18 | Putative RsbT co-antagonist protein rsbRA | 31.76 | 284 | -0.616 | -1.53 | 2.31E-04 | 9.74E-04 | no | no | 0 | 11 | 13 | 17 | 21 | 23 | 21 | 0.013287003 | 0.012650323 | 0.01356521 | 0.023574949 | 0.022138403 | 0.021496541 |
| rpoD | A0A0K2G796 | RNA polymerase sigma factor | 48.621 | 431 | 2.15 | 4.44 | 8.02E-05 | 4.86E-04 | yes | no | 0 | 25 | 29 | 24 | 2 | 9 | 3 | 0.013263184 | 0.022532526 | 0.01200784 | 0.002101178 | 0.002912997 | 0.002429622 |
| trpC | A0A0K2GCM6 | Indole-3-glycerol phosphate synthase | 30.626 | 277 | 0.107 | 1.08 | 1.41E-01 | 1.75E-01 | no | no | 0 | 11 | 9 | 10 | 10 | 9 | 11 | 0.013229669 | 0.013583263 | 0.012662768 | 0.013370095 | 0.013805389 | 0.011730764 |
| NITMOv2_3618 | A0A0K2GGC8 | Uncharacterized protein | 44.801 | 417 | 0.0408 | 1.03 | 5.38E-01 | 5.79E-01 | no | no | 0 | 18 | 18 | 17 | 16 | 17 | 15 | 0.013212806 | 0.013744391 | 0.013043531 | 0.014186578 | 0.01438833 | 0.011579182 |
| gltA | A0A0K2GD35 | Citrate synthase | 42.067 | 377 | 0.1 | 1.07 | 3.46E-01 | 3.89E-01 | no | no | 0 | 20 | 25 | 25 | 17 | 18 | 17 | 0.01318435 | 0.01728354 | 0.018962613 | 0.015384946 | 0.014707923 | 0.011615504 |
| valS | A0A0K2GDE5 | Valine--tRNA ligase | 104.04 | 913 | -0.0247 | -1.02 | 7.82E-01 | 8.09E-01 | no | no | 0 | 33 | 36 | 40 | 47 | 45 | 41 | 0.013161795 | 0.013131947 | 0.014488743 | 0.016371122 | 0.016591761 | 0.014753046 |
| NITMOv2_4412 | A0A0K2GJI1 | Uncharacterized protein | 39.729 | 362 | 1.17 | 2.25 | 1.69E-04 | 7.81E-04 | yes | no | 0 | 9 | 8 | 8 | 7 | 8 | 5 | 0.013128491 | 0.019155482 | 0.015464999 | 0.007467103 | 0.007440424 | 0.007192665 |
| nuoH | A0A0K2G7P7 | NADH-quinone oxidoreductase subunit H | 39.264 | 358 | -0.218 | -1.16 | 1.56E-01 | 1.92E-01 | no | no | 0 | 16 | 10 | 11 | 13 | 17 | 15 | 0.013123854 | 0.012733624 | 0.011639539 | 0.01435642 | 0.013660431 | 0.01409853 |
| modA.1 | A0A0K2GEJ2 | Molybdenum ABC transporter, periplasmic binding protein | 29.824 | 274 | 1.02 | 2.03 | 1.50E-05 | 1.53E-04 | yes | no | 0 | 15 | 16 | 15 | 5 | 8 | 7 | 0.013049656 | 0.013647597 | 0.013585533 | 0.006478706 | 0.007341765 | 0.007070213 |
| glmS | A0A0K2GGT3 | Glutamine--fructose-6-phosphate aminotransferase [isomerizing] | 66.858 | 609 | -0.109 | -1.08 | 1.27E-01 | 1.59E-01 | no | no | 0 | 25 | 26 | 28 | 26 | 29 | 27 | 0.013009396 | 0.012572301 | 0.013640174 | 0.019870124 | 0.018353635 | 0.017283183 |
| ecfE | A0A0K2GA62 | Zinc metalloprotease | 49.993 | 463 | -0.406 | -1.33 | 1.81E-02 | 2.92E-02 | no | no | 0 | 6 | 7 | 6 | 13 | 13 | 12 | 0.012951429 | 0.014722306 | 0.011646249 | 0.018832121 | 0.02069814 | 0.019639081 |
| NITMOv2_3951 | A0A0K2GHC4 | PPK2 domain-containing protein | 39.783 | 337 | 0.821 | 1.77 | 2.33E-04 | 9.80E-04 | no | no | 0 | 16 | 19 | 18 | 15 | 11 | 13 | 0.012939836 | 0.014868768 | 0.01282765 | 0.010133038 | 0.007785962 | 0.006985522 |
| NITMOv2_2483 | A0A0K2GD63 | Phosphopentomutase | 40.331 | 384 | 0.583 | 1.50 | 7.40E-04 | 2.28E-03 | no | no | 0 | 23 | 26 | 22 | 15 | 13 | 14 | 0.012939625 | 0.01221074 | 0.01452958 | 0.00963299 | 0.010141188 | 0.01001122 |
| NITMOv2_1067 | A0A0K2G969 | WD_REPEATS_REGION domain-containing protein | 47.432 | 444 | 0.996 | 1.99 | 2.50E-05 | 2.20E-04 | no | no | 0 | 20 | 17 | 19 | 6 | 9 | 9 | 0.012925924 | 0.010354637 | 0.011684594 | 0.005019875 | 0.006036362 | 0.006596768 |
| NITMOv2_4093 | A0A0K2GHQ3 | Histidine kinase | 90.787 | 830 | 1.95 | 3.86 | 9.51E-08 | 8.21E-06 | yes | no | 0 | 49 | 47 | 42 | 10 | 11 | 11 | 0.012918968 | 0.013941499 | 0.012966074 | 0.003558083 | 0.003042729 | 0.002635686 |
| NITMOv2_0023 | A0A0K2G7B0 | Uncharacterized protein | 10.097 | 91 | -0.202 | -1.15 | 3.55E-02 | 5.22E-02 | no | no | 0 | 3 | 3 | 4 | 6 | 6 | 6 | 0.012894727 | 0.013151893 | 0.014339582 | 0.016839038 | 0.018588241 | 0.015929557 |
| NITMOv2_0867 | A0A0K2G9M6 | RDD domain-containing protein | 16.093 | 152 | 0.838 | 1.79 | 1.87E-05 | 1.77E-04 | no | no | 0 | 2 | 2 | 2 | 0 | 1 | 1 | 0.012893884 | 0.012511095 | 0.013117152 | 0.008336153 | 0.008272109 | 0.00875757 |
| NITMOv2_3044 | A0A0K2GF29 | Putative transcriptional regulator, XRE family | 13.603 | 120 | 0.473 | 1.39 | 8.44E-03 | 1.54E-02 | no | no | 0 | 4 | 6 | 4 | 4 | 5 | 6 | 0.012867114 | 0.006756629 | 0.010102874 | 0.006543414 | 0.004266564 | 0.006107139 |
| NITMOv2_1877 | A0A0K2GCH2 | Uncharacterized protein | 42.661 | 392 | -0.268 | -1.20 | 2.01E-02 | 3.19E-02 | no | no | 0 | 11 | 12 | 11 | 15 | 15 | 17 | 0.012851937 | 0.012419581 | 0.012429248 | 0.016341507 | 0.014407907 | 0.01396457 |
| dat | A0A0K2GIU1 | D-alanine aminotransferase | 31.882 | 288 | 0.129 | 1.09 | 1.14E-01 | 1.45E-01 | no | no | 0 | 12 | 11 | 13 | 12 | 10 | 14 | 0.01276172 | 0.013110633 | 0.012499227 | 0.010380322 | 0.013258493 | 0.013094459 |
| NITMOv2_2242 | A0A0K2GCI4 | Uncharacterized protein | 33.062 | 305 | 0.341 | 1.27 | 3.05E-03 | 6.70E-03 | no | no | 0 | 10 | 10 | 12 | 15 | 10 | 12 | 0.012715136 | 0.01287207 | 0.012919485 | 0.011713585 | 0.009397285 | 0.010236345 |
| NITMOv2_0971 | A0A0K2G8Y3 | TPR_REGION domain-containing protein | 53.369 | 465 | -0.133 | -1.10 | 1.09E-01 | 1.39E-01 | no | no | 0 | 24 | 32 | 31 | 31 | 31 | 29 | 0.012711553 | 0.013869539 | 0.013980867 | 0.011899271 | 0.012363884 | 0.014465886 |
| rpiA | A0A0K2G6W0 | Ribose-5-phosphate isomerase A | 25.023 | 233 | -0.578 | -1.49 | 4.88E-03 | 9.74E-03 | no | no | 0 | 10 | 11 | 13 | 15 | 14 | 13 | 0.012696165 | 0.012557048 | 0.01305235 | 0.008876329 | 0.007190902 | 0.00819296 |
| NITMOv2_4339 | A0A0K2GIE2 | Uncharacterized protein | 22.311 | 196 | 0.0584 | 1.04 | 3.94E-01 | 4.38E-01 | no | no | 0 | 6 | 8 | 10 | 11 | 9 | 11 | 0.012691317 | 0.013502308 | 0.013165467 | 0.015638153 | 0.016486111 | 0.017459219 |
| NITMOv2_0471 | A0A0K2G8K2 | Transcriptional regulator, MarR family | 17.412 | 153 | -2.64 | -6.23 | 1.45E-03 | 3.74E-03 | yes | yes | 3 | 1 | 2 | 2 | 3 | 2 | 1 | 0.012686891 | 0.010767235 | 0.010337927 | 0.007272533 | 0.007849508 | 0.006865767 |
| ureB | A0A0K2G9Q4 | Urease subunit beta | 17.731 | 164 | 0.264 | 1.20 | 1.69E-02 | 2.76E-02 | no | no | 0 | 5 | 6 | 6 | 8 | 7 | 8 | 0.012680567 | 0.014108103 | 0.013707853 | 0.011311119 | 0.011385842 | 0.012735554 |
| NITMOv2_3123 | A0A0K2GFZ0 | AbiEi_1 domain-containing protein | 29.809 | 268 | 0.37 | 1.29 | 7.62E-03 | 1.42E-02 | no | no | 0 | 17 | 18 | 20 | 16 | 16 | 18 | 0.012675508 | 0.012020476 | 0.011592566 | 0.010402385 | 0.009170758 | 0.009079793 |
| udg | A0A0K2GIT0 | UDP-glucose 6-dehydrogenase | 47.518 | 438 | -0.631 | -1.55 | 1.67E-04 | 7.81E-04 | no | no | 0 | 28 | 26 | 26 | 40 | 40 | 41 | 0.012658013 | 0.012792483 | 0.013274174 | 0.025399228 | 0.024481354 | 0.021701526 |
| NITMOv2_3055 | A0A0K2GF36 | PINc domain-containing protein | 16.747 | 152 | 0.094 | 1.07 | 6.53E-01 | 6.91E-01 | no | no | 0 | 7 | 7 | 6 | 12 | 7 | 9 | 0.012633561 | 0.013909821 | 0.01300787 | 0.012130415 | 0.009104572 | 0.007010515 |
| NITMOv2_0817 | A0A0K2G9H3 | YciI-like protein | 10.304 | 92 | 0.00684 | 1.00 | 9.40E-01 | 9.48E-01 | no | no | 0 | 5 | 7 | 6 | 6 | 6 | 5 | 0.012628502 | 0.012411564 | 0.009723645 | 0.014077151 | 0.014393613 | 0.013431247 |
| NITMOv2_3323 | A0A0K2GFI8 | Putative Signal peptide peptidase SppA | 33.784 | 317 | -0.665 | -1.59 | 2.47E-05 | 2.20E-04 | no | no | 0 | 16 | 19 | 17 | 21 | 20 | 21 | 0.01262787 | 0.012949505 | 0.013500983 | 0.022073473 | 0.02300691 | 0.021002057 |
| asd | A0A0K2G9M1 | Aspartate-semialdehyde dehydrogenase | 36.535 | 341 | -0.422 | -1.34 | 1.10E-03 | 3.02E-03 | no | no | 0 | 16 | 21 | 22 | 28 | 28 | 25 | 0.012624076 | 0.012941292 | 0.012577259 | 0.016821269 | 0.01347228 | 0.015956888 |
| NITMOv2_3548 | A0A0K2GG63 | Peptidase_M23 domain-containing protein | 34.618 | 323 | -0.444 | -1.36 | 1.56E-02 | 2.59E-02 | no | no | 0 | 9 | 13 | 11 | 12 | 10 | 11 | 0.012564001 | 0.010725779 | 0.009429541 | 0.021763997 | 0.011709939 | 0.00937954 |
| argS | A0A0K2GGE3 | Arginine--tRNA ligase | 64.191 | 576 | -0.15 | -1.11 | 6.45E-02 | 8.77E-02 | no | no | 0 | 29 | 22 | 25 | 32 | 26 | 28 | 0.012514677 | 0.012297366 | 0.012262257 | 0.013204844 | 0.011359895 | 0.011502402 |
| galT | A0A0K2GGP8 | UDP-glucose--hexose-1-phosphate uridylyltransferase | 38.24 | 334 | 0.366 | 1.29 | 1.89E-03 | 4.60E-03 | no | no | 0 | 11 | 10 | 10 | 13 | 12 | 11 | 0.012403803 | 0.012496039 | 0.013247524 | 0.010270303 | 0.009889181 | 0.009098674 |
| NITMOv2_0506 | A0A0K2G8N2 | Uncharacterized protein | 90.799 | 832 | 0.0454 | 1.03 | 6.20E-01 | 6.61E-01 | no | no | 0 | 64 | 65 | 68 | 65 | 69 | 61 | 0.012337826 | 0.013121192 | 0.015411509 | 0.013773747 | 0.014169727 | 0.013634255 |
| NITMOv2_2039 | A0A0K2GBX7 | Uncharacterized protein | 24.628 | 229 | 0.665 | 1.59 | 3.95E-04 | 1.41E-03 | no | no | 0 | 9 | 10 | 8 | 7 | 8 | 8 | 0.012287026 | 0.011908038 | 0.012142621 | 0.007254024 | 0.008575698 | 0.00877681 |
| NITMOv2_1321 | A0A0K2GA47 | Uncharacterized protein | 16.113 | 147 | 0.533 | 1.45 | 3.35E-02 | 4.97E-02 | no | no | 0 | 3 | 3 | 2 | 1 | 2 | 2 | 0.012262997 | 0.011505804 | 0.010543838 | 0.010138813 | 0.009311988 | 0.008944934 |
| lnt | A0A0K2GCV3 | Apolipoprotein N-acyltransferase | 59.888 | 537 | 0.469 | 1.38 | 4.73E-04 | 1.63E-03 | no | no | 0 | 17 | 19 | 23 | 18 | 16 | 18 | 0.012262575 | 0.012691974 | 0.011766652 | 0.01020056 | 0.00932333 | 0.009318764 |
| pyrE | A0A0K2G6V4 | Orotate phosphoribosyltransferase | 20.621 | 190 | 0.0863 | 1.06 | 2.99E-01 | 3.41E-01 | no | no | 0 | 4 | 2 | 2 | 6 | 5 | 3 | 0.012262153 | 0.012085592 | 0.012008031 | 0.013142801 | 0.01427196 | 0.014228175 |
| glnS | A0A0K2GIC4 | Glutamine--tRNA ligase | 64.601 | 566 | 0.341 | 1.27 | 3.11E-03 | 6.80E-03 | no | no | 0 | 18 | 14 | 18 | 15 | 17 | 15 | 0.012227795 | 0.012580122 | 0.012188827 | 0.00960604 | 0.009594447 | 0.008561755 |
| nrdZ | A0A0K2GB93 | Vitamin B12-dependent ribonucleotide reductase | 63.362 | 577 | 0.925 | 1.90 | 2.02E-05 | 1.90E-04 | no | no | 0 | 30 | 36 | 32 | 20 | 21 | 22 | 0.012203765 | 0.012301081 | 0.012560004 | 0.005651858 | 0.006561506 | 0.005013522 |
| exbD.1 | A0A0K2GGM3 | Biopolymer transport protein ExbD | 16.538 | 147 | 0.223 | 1.17 | 1.10E-01 | 1.40E-01 | no | no | 0 | 7 | 5 | 7 | 5 | 8 | 5 | 0.012163716 | 0.0128883 | 0.013608157 | 0.008947849 | 0.013095046 | 0.008914546 |

| | | | | | | | | | | | | | | | | | | | | | | |
|---|---|---|---|---|---|---|---|---|---|---|---|---|---|---|---|---|---|---|---|---|---|---|
| purA | A0A0K2GHK8 | Adenylosuccinate synthetase | 47.796 | 438 | -0.386 | -1.31 | 3.35E-03 | 7.21E-03 | no | no | 0 | 14 | 16 | 15 | 13 | 16 | 19 | 0.012141583 | 0.012474724 | 0.011596784 | 0.013963134 | 0.013529455 | 0.013462714 |
| StomI | A0A0K2GK56 | Stomatin-like protein 2 | 34.129 | 312 | 0.165 | 1.12 | 1.09E-01 | 1.40E-01 | no | no | 0 | 11 | 11 | 19 | 21 | 25 | 21 | 0.012097317 | 0.01278427 | 0.015390227 | 0.01264838 | 0.014057241 | 0.013571141 |
| NITMOv2_3446 | A0A0K2GFV3 | TonB_C domain-containing protein | 16.94 | 153 | 0.896 | 1.86 | 3.04E-04 | 1.17E-03 | no | no | 0 | 5 | 5 | 4 | 2 | 3 | 2 | 0.012081087 | 0.011547455 | 0.010578732 | 0.00655763 | 0.00750226 | 0.005195851 |
| NITMOv2_3166 | A0A0K2GF28 | Uncharacterized protein | 13.182 | 113 | -0.333 | -1.26 | 6.43E-03 | 1.23E-02 | no | no | 0 | 3 | 4 | 2 | 3 | 3 | 4 | 0.012062116 | 0.010284241 | 0.011666188 | 0.015010317 | 0.014372483 | 0.015574248 |
| hslU | A0A0K2G716 | ATP-dependent protease ATPase subunit HslU | 52.284 | 465 | 0.0773 | 1.06 | 2.24E-01 | 2.65E-01 | no | no | 0 | 26 | 30 | 32 | 32 | 28 | 25 | 0.0120364 | 0.013064876 | 0.012987356 | 0.012812002 | 0.01158549 | 0.009972741 |
| NITMOv2_4293 | A0A0K2GI97 | Putative Outer membrane efflux protein | 54.39 | 492 | 0.553 | 1.47 | 3.48E-04 | 1.30E-03 | no | no | 0 | 19 | 20 | 15 | 14 | 11 | 9 | 0.012015743 | 0.012247698 | 0.012189785 | 0.007977516 | 0.00756767 | 0.008388775 |
| GGPS | A0A0K2G8U0 | Geranylgeranyl pyrophosphate synthase, chloroplastic | 32.492 | 297 | 0.192 | 1.14 | 5.37E-02 | 7.47E-02 | no | no | 0 | 13 | 10 | 12 | 8 | 8 | 7 | 0.011970845 | 0.009653221 | 0.012020494 | 0.0089449 | 0.014639561 | 0.010754923 |
| NITMOv2_0492 | A0A0K2G7K3 | Putative 2-oxoglutarate-Fe(II) oxygenase superfamily | 28.94 | 252 | -0.23 | -1.17 | 1.01E-02 | 1.80E-02 | no | no | 0 | 12 | 12 | 12 | 14 | 16 | 15 | 0.011909506 | 0.011305958 | 0.012082804 | 0.014101732 | 0.015988933 | 0.014381195 |
| cysNC | A0A0K2G723 | ATP-sulfurylase large subunit | 68.502 | 616 | -1.03 | -2.04 | 8.94E-06 | 1.10E-04 | yes | no | 0 | 33 | 35 | 30 | 62 | 58 | 65 | 0.011900652 | 0.015859394 | 0.011336807 | 0.027729921 | 0.025461727 | 0.028947955 |
| NITMOv2_1455 | A0A0K2GBB8 | Uncharacterized protein | 10.268 | 91 | 0.583 | 1.50 | 6.20E-04 | 2.00E-03 | no | no | 0 | 2 | 3 | 4 | 3 | 2 | 2 | 0.011848166 | 0.01160514 | 0.010260663 | 0.009609742 | 0.00962723 | 0.009936598 |
| NITMOv2_0568 | A0A0K2G824 | Uncharacterized protein | 12.837 | 122 | -3.35 | -10.20 | 5.29E-07 | 1.95E-05 | yes | no | 0 | 2 | 2 | 2 | 11 | 12 | 13 | 0.011844794 | 0.010144036 | 0.007458623 | 0.085082156 | 0.088980873 | 0.071921257 |
| argK | A0A0K2G6C8 | LAO/AO transport system ATPase | 27.639 | 263 | 3.17 | 9.00 | 1.76E-04 | 7.99E-04 | yes | no | 0 | 10 | 9 | 10 | 0 | 0 | 0 | 0.011836151 | 0.012712115 | 0.011366716 | 0.000611977 | 0.000635052 | 0.00066784 |
| macA | A0A0K2G941 | Putative Efflux transporter, RND family, MFP subunit, Macrolide-specific efflux protein | 44.014 | 415 | 0.533 | 1.45 | 3.13E-04 | 1.19E-03 | no | no | 0 | 10 | 13 | 11 | 9 | 8 | 7 | 0.011773337 | 0.011724422 | 0.011203559 | 0.0080172 | 0.009642456 | 0.008670541 |
| forD | A0A0K2G932 | 2-oxoglutarate:ferredoxin oxidoreductase delta subunit | 27.954 | 256 | -2.05 | -4.14 | 2.57E-05 | 2.23E-04 | yes | no | 0 | 9 | 12 | 14 | 60 | 47 | 41 | 0.01176891 | 0.008980941 | 0.013282226 | 0.065987943 | 0.061693571 | 0.054465095 |
| NITMOv2_2415 | A0A0K2GCZ6 | Putative Helicase (C-terminal) | 68.696 | 619 | -0.0393 | -1.03 | 6.56E-01 | 6.93E-01 | no | no | 0 | 34 | 28 | 29 | 40 | 34 | 27 | 0.011760268 | 0.010818467 | 0.012145497 | 0.014783763 | 0.013234101 | 0.011741912 |
| nuoC | A0A0K2G6P8 | NADH-quinone oxidoreductase subunit C | 19.767 | 165 | -0.432 | -1.35 | 1.17E-03 | 3.16E-03 | no | no | 0 | 10 | 9 | 10 | 12 | 11 | 14 | 0.011750572 | 0.012370108 | 0.012369622 | 0.013945069 | 0.014417074 | 0.015546018 |
| NITMOv2_3097 | A0A0K2GEW9 | Uncharacterized protein | 29.949 | 262 | 0.71 | 1.64 | 4.26E-04 | 1.51E-03 | no | no | 0 | 8 | 8 | 8 | 7 | 5 | 5 | 0.011715792 | 0.010662814 | 0.010402155 | 0.008505254 | 0.008009226 | 0.007742171 |
| NITMOv2_1038 | A0A0K2G9E0 | Uncharacterized protein | 22.19 | 197 | 0.203 | 1.15 | 8.63E-02 | 1.14E-01 | no | no | 0 | 10 | 10 | 10 | 9 | 8 | 7 | 0.011701458 | 0.011125471 | 0.011339875 | 0.009336248 | 0.008895135 | 0.00892965 |
| NITMOv2_1961 | A0A0K2GBQ2 | N-acetyltransferase domain-containing protein | 30.674 | 285 | 0.42 | 1.34 | 8.85E-03 | 1.60E-02 | no | no | 0 | 11 | 12 | 13 | 8 | 11 | 14 | 0.011682909 | 0.01120936 | 0.0101347 | 0.009851992 | 0.010569693 | 0.009108024 |
| ddlB | A0A0K2G7W2 | D-alanine--D-alanine ligase | 35.139 | 329 | 0.177 | 1.13 | 9.47E-02 | 1.23E-01 | no | no | 0 | 6 | 5 | 7 | 4 | 4 | 7 | 0.011678061 | 0.012395138 | 0.012265325 | 0.006547857 | 0.006666224 | 0.01272962 |
| NITMOv2_1436 | A0A0K2GA86 | 7,8-dihydroneopterin aldolase | 35.106 | 310 | 0.882 | 1.84 | 1.87E-05 | 1.77E-04 | no | no | 0 | 6 | 9 | 7 | 4 | 3 | 3 | 0.011626207 | 0.012625489 | 0.011211803 | 0.007054716 | 0.008060653 | 0.006427205 |
| NITMOv2_4080 | A0A0K2GHY9 | Putative 2-deoxycytidine 5-triphosphate deaminase | 43.485 | 384 | -0.395 | -1.31 | 1.13E-03 | 3.06E-03 | no | no | 0 | 15 | 15 | 19 | 24 | 21 | 29 | 0.011580255 | 0.011824736 | 0.011606179 | 0.015925418 | 0.017051651 | 0.015451796 |
| NITMOv2_3355 | A0A0K2GGJ7 | Uncharacterized protein | 26.724 | 230 | 0.335 | 1.26 | 1.19E-02 | 2.06E-02 | no | no | 0 | 14 | 12 | 15 | 8 | 11 | 9 | 0.011574353 | 0.011305763 | 0.011545786 | 0.00568932 | 0.007838477 | 0.008400823 |
| NITMOv2_0292 | A0A0K2G6Y0 | Uncharacterized protein | 17.557 | 165 | -0.35 | -1.27 | 7.18E-02 | 9.66E-02 | no | no | 0 | 1 | 2 | 0 | 1 | 2 | 1 | 0.011569294 | 0.015378356 | 0.001765908 | 0.002841847 | 0.002903209 | 0.003080541 |
| cysS | A0A0K2GIP6 | Cysteine--tRNA ligase | 55.902 | 491 | 0.615 | 1.53 | 3.75E-05 | 2.87E-04 | no | no | 0 | 25 | 29 | 21 | 18 | 14 | 17 | 0.011537676 | 0.011732048 | 0.012169079 | 0.008383981 | 0.008251911 | 0.008531187 |
| NITMOv2_0163 | A0A0K2G6N0 | Putative NADH-quinone oxidoreductase, subunit G | 98.157 | 901 | -1.14 | -2.20 | 2.01E-06 | 4.22E-05 | yes | no | 0 | 36 | 46 | 45 | 82 | 80 | 82 | 0.011532406 | 0.011525359 | 0.012122874 | 0.034431775 | 0.033994544 | 0.032200757 |
| NITMOv2_1575 | A0A0K2GAV9 | RHH_1 domain-containing protein | 14.067 | 123 | 0.585 | 1.50 | 9.41E-04 | 2.70E-03 | no | no | 0 | 8 | 6 | 7 | 4 | 2 | 5 | 0.011482239 | 0.013304417 | 0.010453153 | 0.008992864 | 0.009017099 | 0.007756376 |
| NITMOv2_1744 | A0A0K2GBB5 | Uncharacterized protein | 15.472 | 150 | 0.599 | 1.51 | 9.61E-05 | 5.43E-04 | no | no | 0 | 7 | 8 | 10 | 5 | 6 | 6 | 0.011466641 | 0.01269608 | 0.012530478 | 0.009867244 | 0.008398268 | 0.008242768 |
| NITMOv2_4862 | A0A0K2GJU8 | Uncharacterized protein | 56.766 | 495 | 0.142 | 1.10 | 9.59E-02 | 1.25E-01 | no | no | 0 | 18 | 21 | 25 | 21 | 20 | 23 | 0.011456101 | 0.010971774 | 0.011648166 | 0.010416748 | 0.009730239 | 0.010328769 |
| tpm | A0A0K2GB89 | Thiopurine S-methyltransferase | 23.794 | 212 | 0.396 | 1.32 | 1.72E-02 | 2.81E-02 | no | no | 0 | 11 | 13 | 14 | 13 | 10 | 9 | 0.011445773 | 0.011009318 | 0.014326353 | 0.012575083 | 0.010754736 | 0.00880522 |
| malQ | A0A0K2G9G5 | 4-alpha-glucanotransferase | 83.891 | 749 | 0.191 | 1.14 | 3.96E-02 | 5.72E-02 | no | no | 0 | 28 | 28 | 39 | 42 | 38 | 40 | 0.011384433 | 0.011529269 | 0.012797741 | 0.012271086 | 0.012021919 | 0.010479271 |
| pyrD | A0A0K2G972 | Dihydroorotate dehydrogenase | 32.904 | 309 | 0.311 | 1.24 | 3.86E-02 | 5.60E-02 | no | no | 0 | 12 | 13 | 11 | 14 | 14 | 10 | 0.011367992 | 0.011506 | 0.010601931 | 0.011917928 | 0.010388844 | 0.010015716 |
| ybeZ | A0A0K2GJL4 | Putative enzyme with nucleoside triphosphate hydrolase domain | 39.607 | 359 | -0.123 | -1.09 | 3.01E-01 | 3.43E-01 | no | no | 0 | 16 | 17 | 18 | 25 | 21 | 21 | 0.011320143 | 0.013376964 | 0.012708589 | 0.01469788 | 0.013885093 | 0.011446301 |
| aroE | A0A0K2G7A2 | Shikimate dehydrogenase (NADP(+)) | 30.491 | 287 | 0.268 | 1.20 | 1.10E-02 | 1.93E-02 | no | no | 0 | 10 | 10 | 14 | 10 | 10 | 12 | 0.011303912 | 0.011832362 | 0.010935913 | 0.010636639 | 0.0116351 | 0.010571155 |
| yddE | A0A0K2GC12 | Putative isomerase | 33.497 | 309 | -0.664 | -1.58 | 3.86E-02 | 5.60E-02 | no | no | 0 | 21 | 22 | 24 | 15 | 17 | 19 | 0.011300118 | 0.011283862 | 0.013265355 | 0.010011468 | 0.010072515 | 0.009820799 |
| NITMOv2_2494 | A0A0K2GE99 | Alpha,alpha-trehalose-phosphate synthase (UDP-forming) | 85.311 | 749 | 0.0351 | 1.02 | 6.68E-01 | 7.05E-01 | no | no | 0 | 22 | 25 | 30 | 25 | 23 | 24 | 0.01126302 | 0.011312216 | 0.011804996 | 0.005992133 | 0.006094314 | 0.00531237 |
| NITMOv2_0358 | A0A0K2G7E9 | Putative Acyltransferase | 25.187 | 230 | 0.239 | 1.18 | 3.59E-02 | 5.27E-02 | no | no | 0 | 8 | 7 | 9 | 9 | 8 | 7 | 0.011251426 | 0.009402143 | 0.010808417 | 0.008731216 | 0.007816104 | 0.00906379 |
| zraR | A0A0K2GBJ1 | Transcriptional regulatory protein ZraR | 48.529 | 445 | 0.405 | 1.32 | 2.25E-03 | 5.27E-03 | no | no | 0 | 20 | 21 | 17 | 13 | 14 | 13 | 0.011242995 | 0.011546868 | 0.011171158 | 0.007516856 | 0.007925483 | 0.007349641 |
| NITMOv2_2407 | A0A0K2GDZ0 | Multidrug resistance ABC transporter ATP-binding and permease | 63.749 | 574 | -0.948 | -1.93 | 5.27E-05 | 3.64E-04 | no | no | 0 | 7 | 6 | 7 | 12 | 12 | 14 | 0.011242784 | 0.009895109 | 0.009586946 | 0.008300023 | 0.008275993 | 0.009177971 |
| NITMOv2_1240 | A0A0K2G9N9 | Nucleotide-binding protein NITMOv2_1240 | 32.656 | 288 | 0.179 | 1.13 | 1.02E-01 | 1.31E-01 | no | no | 0 | 14 | 15 | 15 | 15 | 12 | 12 | 0.011229504 | 0.011662435 | 0.012449571 | 0.010485307 | 0.009748106 | 0.008742106 |
| NITMOv2_1451 | A0A0K2GAA2 | PPM-type phosphatase domain-containing protein | 26.587 | 249 | 0.642 | 1.56 | 3.26E-04 | 1.22E-03 | no | no | 0 | 6 | 6 | 5 | 5 | 4 | 4 | 0.01119641 | 0.01273734 | 0.012086446 | 0.007685069 | 0.008139736 | 0.008795151 |
| NITMOv2_3871 | A0A0K2GH16 | Uncharacterized protein | 26.142 | 237 | 0.477 | 1.39 | 2.23E-03 | 5.22E-03 | no | no | 0 | 5 | 6 | 5 | 6 | 4 | 3 | 0.011173435 | 0.010622532 | 0.009651749 | 0.008685017 | 0.0075888 | 0.007735338 |
| trmI | A0A0K2G779 | tRNA (adenine(58)-N(1))-methyltransferase TrmI | 32.089 | 292 | 0.229 | 1.17 | 7.37E-02 | 9.87E-02 | no | no | 0 | 17 | 16 | 18 | 15 | 13 | 11 | 0.011165847 | 0.012518526 | 0.013560417 | 0.009281017 | 0.00886344 | 0.007956327 |
| NITMOv2_4432 | A0A0K2GJK4 | Glycosyl transferase, family 2 | 27.85 | 244 | 0.0639 | 1.05 | 6.09E-01 | 6.50E-01 | no | no | 0 | 12 | 11 | 11 | 13 | 9 | 8 | 0.01115067 | 0.011353671 | 0.010910222 | 0.012769357 | 0.010787208 | 0.009840579 |
| NITMOv2_0714 | A0A0K2G890 | Histidine kinase | 67.605 | 618 | 0.213 | 1.16 | 1.01E-02 | 1.80E-02 | no | no | 0 | 37 | 37 | 32 | 24 | 28 | 23 | 0.011137812 | 0.011801271 | 0.012514757 | 0.009679929 | 0.010364296 | 0.009411367 |
| NITMOv2_2911 | A0A0K2GFD8 | Putative heavy metal efflux pump protein CzcC | 47.623 | 424 | 0.721 | 1.65 | 9.13E-04 | 2.65E-03 | no | no | 0 | 14 | 17 | 19 | 14 | 14 | 10 | 0.011131277 | 0.011051751 | 0.011440338 | 0.00662293 | 0.007651103 | 0.006753024 |
| NITMOv2_0130 | A0A0K2G6L3 | Putative Regulatory protein, MerR family | 34.353 | 306 | 0.484 | 1.40 | 3.49E-03 | 7.48E-03 | no | no | 0 | 14 | 13 | 15 | 12 | 11 | 11 | 0.011095443 | 0.01303222 | 0.013769396 | 0.008743358 | 0.009102707 | 0.007413295 |
| ftsE | A0A0K2GG77 | Cell division ATP-binding protein FtsE | 24.767 | 225 | 0.157 | 1.11 | 6.53E-02 | 8.85E-02 | no | no | 0 | 12 | 15 | 14 | 9 | 10 | 8 | 0.011072046 | 0.011557623 | 0.013404546 | 0.013073354 | 0.012323333 | 0.011484601 |
| NITMOv2_4098 | A0A0K2GHQ8 | Uncharacterized protein | 30.607 | 287 | 0.169 | 1.12 | 1.13E-01 | 1.43E-01 | no | no | 0 | 7 | 12 | 12 | 11 | 7 | 6 | 0.011053497 | 0.012331586 | 0.011298463 | 0.009056091 | 0.008440994 | 0.009629839 |
| hemY | A0A0K2G9A8 | Coproporphyrinogen III oxidase | 51.722 | 483 | 0.376 | 1.30 | 1.16E-03 | 3.15E-03 | no | no | 0 | 29 | 27 | 29 | 26 | 22 | 24 | 0.01104633 | 0.01055155 | 0.010250885 | 0.008289806 | 0.008551461 | 0.009634694 |
| tolQ | A0A0K2GHI7 | Protein TolQ | 24.557 | 231 | 0.394 | 1.31 | 3.09E-02 | 4.64E-02 | no | no | 0 | 8 | 7 | 9 | 6 | 10 | 8 | 0.01102504 | 0.01200933 | 0.011472164 | 0.008405896 | 0.011663329 | 0.011111491 |
| NITMOv2_1733 | A0A0K2GB28 | Putative NAD-dependent alcohol dehydrogenase | 36.784 | 342 | 0.375 | 1.30 | 1.61E-03 | 4.08E-03 | no | no | 0 | 12 | 8 | 14 | 9 | 12 | 9 | 0.011020614 | 0.011719534 | 0.012820556 | 0.013872068 | 0.011767736 | 0.010969439 |
| pycA | A0A0K2GAB2 | Biotin carboxylase | 54.851 | 497 | 0.0526 | 1.04 | 3.89E-01 | 4.33E-01 | no | no | 0 | 14 | 20 | 18 | 16 | 14 | 15 | 0.011018506 | 0.010968449 | 0.011257817 | 0.011288463 | 0.009567569 | 0.010044845 |
| kdsA | A0A0K2GGE4 | 2-dehydro-3-deoxyphosphooctonate aldolase | 29.337 | 275 | -1.04 | -2.06 | 9.76E-05 | 5.47E-04 | yes | no | 0 | 5 | 3 | 6 | 14 | 17 | 13 | 0.010973819 | 0.010380058 | 0.010435323 | 0.029166244 | 0.023286573 | 0.024132586 |
| argH | A0A0K2G7A4 | Argininosuccinate lyase | 53.068 | 469 | 0.044 | 1.03 | 5.43E-01 | 5.84E-01 | no | no | 0 | 14 | 18 | 18 | 19 | 19 | 20 | 0.010929975 | 0.012392009 | 0.012051361 | 0.01143639 | 0.011761522 | 0.011349742 |
| thrS | A0A0K2G831 | Threonine--tRNA ligase | 72.734 | 643 | -0.671 | -1.59 | 1.72E-04 | 7.89E-04 | no | no | 0 | 21 | 17 | 24 | 54 | 53 | 45 | 0.010920279 | 0.009921312 | 0.011512809 | 0.020922934 | 0.021611705 | 0.019739776 |
| NITMOv2_1505 | A0A0K2GAE4 | Putative CRISPR-associated protein Cas8a1/Csx13 | 56.813 | 503 | 0.372 | 1.29 | 1.71E-03 | 4.26E-03 | no | no | 0 | 33 | 29 | 33 | 28 | 30 | 31 | 0.010886764 | 0.011336854 | 0.011264144 | 0.008716408 | 0.009117622 | 0.009289994 |
| NITMOv2_4525 | A0A0K2GJ72 | Putative Glutathione transferase FosA | 16.134 | 142 | -1.16 | -2.23 | 3.75E-02 | 5.47E-02 | no | yes | 1 | 0 | 1 | 2 | 2 | 3 | 2 | 0.010880651 | 0.011128795 | 0.010286545 | 0.007898889 | 0.007862093 | 0.007907778 |
| NITMOv2_0554 | A0A0K2G7T3 | Putative TonB-dependent receptor | 89.088 | 796 | 0.25 | 1.19 | 5.07E-03 | 1.00E-02 | no | no | 0 | 31 | 26 | 31 | 21 | 27 | 24 | 0.010841444 | 0.011282884 | 0.011550579 | 0.008931709 | 0.009667781 | 0.009958895 |
| NITMOv2_0612 | A0A0K2G7W5 | Sortilin-Vps10 domain-containing protein | 36.893 | 338 | 0.275 | 1.21 | 4.61E-03 | 9.30E-03 | no | no | 0 | 17 | 16 | 29 | 18 | 17 | 16 | 0.010829429 | 0.011017531 | 0.011495363 | 0.008896171 | 0.009271903 | 0.008046773 |
| NITMOv2_3688 | A0A0K2GGJ3 | Putative Type IV pilus biogenesis/stability protein PilW | 25.628 | 234 | -0.943 | -1.92 | 1.68E-04 | 7.81E-04 | no | no | 0 | 5 | 6 | 8 | 15 | 13 | 12 | 0.01076261 | 0.009778174 | 0.012115205 | 0.024125786 | 0.025676135 | 0.023729807 |
| drrA | A0A0K2G7I3 | Daunorubicin/doxorubicin resistance ATP-binding protein DrrA | 35.251 | 322 | -0.798 | -1.74 | 1.04E-04 | 5.73E-04 | no | no | 0 | 8 | 9 | 12 | 20 | 17 | 20 | 0.010743428 | 0.012972775 | 0.013295072 | 0.024396763 | 0.021400404 | 0.023762173 |
| gltX | A0A0K2GIC9 | Glutamate--tRNA ligase | 53.508 | 471 | 0.00998 | 1.01 | 8.97E-01 | 9.12E-01 | no | no | 0 | 36 | 36 | 38 | 31 | 35 | 32 | 0.010739001 | 0.011100637 | 0.01227702 | 0.011546113 | 0.010610244 | 0.010235266 |
| NITMOv2_2400 | A0A0K2GCY2 | Uncharacterized protein | 9.8762 | 90 | -0.78 | -1.72 | 1.01E-03 | 2.83E-03 | no | no | 0 | 4 | 5 | 3 | 6 | 5 | 5 | 0.010730148 | 0.011320624 | 0.013358532 | 0.018983157 | 0.016190911 | 0.016430334 |
| NITMOv2_1002 | A0A0K2G9A6 | TPR_REGION domain-containing protein | 59.038 | 526 | 0.335 | 1.26 | 1.38E-03 | 3.63E-03 | no | no | 0 | 19 | 19 | 21 | 15 | 18 | 23 | 0.01069052 | 0.009255485 | 0.011049222 | 0.007483984 | 0.008223013 | 0.0070249 |
| NITMOv2_4173 | A0A0K2GHV9 | Putative Cell division protein ZapB | 9.8993 | 84 | 0.748 | 1.68 | 5.48E-02 | 7.60E-02 | no | no | 0 | 2 | 2 | 3 | 3 | 3 | 2 | 0.010677451 | 0.009450833 | 0.009887952 | 0.012508746 | 0.013410754 | 0.014779838 |
| guaA | A0A0K2GB03 | GMP synthase [glutamine-hydrolyzing] | 57.996 | 518 | -0.398 | -1.32 | 1.11E-03 | 3.02E-03 | no | no | 0 | 19 | 19 | 26 | 31 | 27 | 27 | 0.010670706 | 0.00960981 | 0.010135276 | 0.013765454 | 0.012958478 | 0.013922314 |
| NITMOv2_1168 | A0A0K2G9R1 | Response regulatory domain-containing protein | 14.244 | 127 | 0.645 | 1.56 | 4.83E-03 | 9.68E-03 | no | no | 0 | 6 | 4 | 4 | 1 | 1 | 1 | 0.010644358 | 0.008804952 | 0.009386403 | 0.007375889 | 0.006813047 | 0.006400952 |
| NITMOv2_0210 | A0A0K2G711 | TPR_REGION domain-containing protein | 41.83 | 381 | 0.238 | 1.18 | 4.83E-02 | 6.83E-02 | no | no | 0 | 25 | 21 | 24 | 21 | 16 | 18 | 0.010607681 | 0.010911742 | 0.0107509 | 0.007439562 | 0.007350932 | 0.007424982 |
| NITMOv2_3903 | A0A0K2GH53 | MCP methyltransferase/methylesterase, CheR/CheB with PAS/PAC sensor | 124.4 | 1116 | 0.346 | 1.27 | 5.43E-03 | 1.06E-02 | no | no | 0 | 30 | 38 | 41 | 34 | 33 | 33 | 0.010583862 | 0.010039616 | 0.011297887 | 0.008948737 | 0.008009071 | 0.007790541 |
| NITMOv2_0793 | A0A0K2G8E1 | TPR_REGION domain-containing protein | 45.934 | 414 | 0.666 | 1.59 | 2.78E-04 | 1.10E-03 | no | no | 0 | 18 | 22 | 27 | 16 | 18 | 15 | 0.010565734 | 0.010624488 | 0.011572244 | 0.00710595 | 0.007481596 | 0.006366428 |
| nnr | A0A0K2GIS6 | Bifunctional NAD(P)H-hydrate repair enzyme | 53.39 | 518 | -0.142 | -1.10 | 1.45E-01 | 1.79E-01 | no | no | 0 | 17 | 14 | 24 | 20 | 20 | 17 | 0.010555827 | 0.010074227 | 0.012729871 | 0.013386976 | 0.013718849 | 0.012053167 |
| NITMOv2_3441 | A0A0K2GFU8 | Uncharacterized protein | 31.389 | 286 | 0.53 | 1.44 | 3.23E-04 | 1.22E-03 | no | no | 0 | 9 | 7 | 9 | 9 | 9 | 4 | 0.010555405 | 0.008992283 | 0.010067022 | 0.007824555 | 0.007920511 | 0.007004042 |
| NITMOv2_3028 | A0A0K2GEQ5 | Uncharacterized protein | 17.422 | 159 | 0.734 | 1.66 | 2.17E-03 | 5.13E-03 | no | no | 0 | 6 | 8 | 8 | 5 | 4 | 3 | 0.010545498 | 0.01076176 | 0.005746339 | 0.004174962 | 0.006359838 | 0.005971381 |
| pdhA | A0A0K2GIK6 | Pyruvate dehydrogenase E1 component | 100.25 | 893 | 0.433 | 1.35 | 6.48E-04 | 2.06E-03 | no | no | 0 | 33 | 37 | 35 | 31 | 34 | 33 | 0.010530322 | 0.011781912 | 0.011099262 | 0.008012462 | 0.008534215 | 0.007722931 |
| NITMOv2_2468 | A0A0K2GD49 | Haloacid dehalogenase, subfamily IA | 26.881 | 240 | -0.155 | -1.11 | 4.35E-02 | 6.22E-02 | no | no | 0 | 17 | 13 | 17 | 13 | 14 | 16 | 0.010519782 | 0.010236529 | 0.009614746 | 0.011002235 | 0.011690363 | 0.010105262 |
| leuS | A0A0K2GJ43 | Leucine--tRNA ligase | 91.838 | 811 | 0.256 | 1.19 | 4.20E-03 | 8.65E-03 | no | no | 0 | 15 | 17 | 20 | 18 | 23 | 22 | 0.01048184 | 0.010590268 | 0.010474435 | 0.010393205 | 0.009754166 | 0.01042353 |
| yhbC | A0A0K2GJ05 | Ribosome maturation factor RimP | 17.966 | 163 | -0.16 | -1.12 | 1.67E-01 | 2.04E-01 | no | no | 0 | 5 | 5 | 6 | 8 | 7 | 8 | 0.010460129 | 0.010587921 | 0.013481619 | 0.017231435 | 0.014954336 | 0.014914517 |
| NITMOv2_1725 | A0A0K2GB22 | Uncharacterized protein | 94.596 | 851 | 0.0859 | 1.06 | 2.19E-01 | 2.61E-01 | no | no | 0 | 20 | 23 | 24 | 21 | 22 | 25 | 0.010459286 | 0.011422502 | 0.01121813 | 0.009719021 | 0.010370045 | 0.009980113 |
| NITMOv2_1449 | A0A0K2GA80 | Uncharacterized protein | 36.678 | 331 | -0.698 | -1.62 | 1.44E-04 | 7.13E-04 | no | no | 0 | 10 | 12 | 12 | 23 | 19 | 18 | 0.010429776 | 0.014041422 | 0.018212398 | 0.034357738 | 0.034083103 | 0.027320656 |

| | | | | | | | | | | | | | | | | | | | | | | |
|---|---|---|---|---|---|---|---|---|---|---|---|---|---|---|---|---|---|---|---|---|---|---|
| nuoB | A0A0K2G6K9 | NADH-quinone oxidoreductase subunit B | 19.365 | 175 | -0.622 | -1.54 | 8.78E-04 | 2.57E-03 | no | no | 0 | 5 | 8 | 13 | 14 | 15 | 14 | 0.010429354 | 0.011426022 | 0.011621325 | 0.022791635 | 0.021024413 | 0.020842024 |
| NITMOv2_1447 | A0A0K2GAG8 | Putative Response regulator with HD domain | 41.719 | 380 | -0.913 | -1.88 | 2.15E-05 | 1.99E-04 | no | no | 0 | 15 | 16 | 21 | 33 | 34 | 32 | 0.010427036 | 0.0097068 | 0.009702555 | 0.021309409 | 0.023210443 | 0.021875944 |
| map | A0A0K2G9K7 | Methionine aminopeptidase | 27.156 | 254 | -0.223 | -1.17 | 5.49E-03 | 1.07E-02 | no | no | 0 | 8 | 6 | 9 | 11 | 11 | 11 | 0.010353049 | 0.01183862 | 0.01195895 | 0.015503405 | 0.014656496 | 0.013972482 |
| NITMOv2_1438 | A0A0K2GA93 | Molybdopterin synthase catalytic subunit (Modular protein) | 29.232 | 258 | -0.656 | -1.58 | 1.70E-03 | 4.25E-03 | no | no | 0 | 12 | 14 | 12 | 29 | 21 | 19 | 0.010347358 | 0.006871413 | 0.010822221 | 0.012859238 | 0.016829474 | 0.015897191 |
| NITMOv2_3858 | A0A0K2GH08 | Uncharacterized protein | 16.358 | 144 | 0.399 | 1.32 | 1.79E-01 | 2.17E-01 | no | no | 0 | 6 | 6 | 10 | 6 | 3 | 2 | 0.010310259 | 0.009898042 | 0.006620024 | 0.008831166 | 0.008600713 | 0.007232044 |
| NITMOv2_4831 | A0A0K2GK14 | Uncharacterized protein | 53.571 | 484 | 0.228 | 1.17 | 1.53E-02 | 2.56E-02 | no | no | 0 | 27 | 24 | 25 | 27 | 27 | 27 | 0.010297612 | 0.010606498 | 0.011120351 | 0.011459193 | 0.012412514 | 0.010932218 |
| NITMOv2_2482 | A0A0K2GD66 | YjgF_endoribonc domain-containing protein | 16.092 | 153 | 0.226 | 1.17 | 3.98E-02 | 5.74E-02 | no | no | 0 | 9 | 10 | 12 | 6 | 7 | 7 | 0.010246601 | 0.012215433 | 0.010359209 | 0.01000673 | 0.009555761 | 0.008084533 |
| NITMOv2_4316 | A0A0K2GJ76 | Putative Ferredoxin reductase | 25.633 | 233 | 0.518 | 1.43 | 2.80E-04 | 1.10E-03 | no | no | 0 | 13 | 12 | 11 | 6 | 8 | 6 | 0.01024365 | 0.009546454 | 0.010214074 | 0.006665428 | 0.006960958 | 0.007191406 |
| fliC | A0A0K2GCI5 | Flagellin | 28.624 | 274 | 0.873 | 1.83 | 1.60E-04 | 7.61E-04 | no | no | 0 | 7 | 6 | 10 | 8 | 8 | 9 | 0.010201071 | 0.006999299 | 0.012586078 | 0.006537047 | 0.006053608 | 0.006145979 |
| NITMOv2_4861 | A0A0K2GK44 | TPR_REGION domain-containing protein | 39.676 | 351 | 0.00411 | 1.00 | 9.44E-01 | 9.50E-01 | no | no | 0 | 8 | 9 | 12 | 13 | 14 | 13 | 0.010190321 | 0.009226935 | 0.00929591 | 0.007887339 | 0.008332237 | 0.008473467 |
| NITMOv2_0363 | A0A0K2G7F4 | Uncharacterized protein | 12.324 | 106 | -1.69 | -3.23 | 1.12E-04 | 6.04E-04 | yes | no | 0 | 4 | 8 | 4 | 11 | 11 | 10 | 0.010189689 | 0.028015577 | 0.016089251 | 0.101813099 | 0.08872607 | 0.082298215 |
| purD | A0A0K2GC88 | Phosphoribosylamine--glycine ligase | 44.587 | 424 | -0.09 | -1.06 | 3.02E-01 | 3.44E-01 | no | no | 0 | 21 | 19 | 17 | 21 | 18 | 16 | 0.010121183 | 0.010007547 | 0.009565856 | 0.012685251 | 0.01009784 | 0.008508171 |
| NITMOv2_4047 | A0A0K2GHN4 | Thiamine pyrophosphate enzyme, central domain family | 59.257 | 545 | 1.71 | 3.27 | 2.93E-05 | 2.43E-04 | yes | no | 0 | 12 | 16 | 8 | 2 | 2 | 0 | 0.010087246 | 0.012374606 | 0.009693353 | 0.00336455 | 0.00313797 | 0.002065142 |
| NITMOv2_1264 | A0A0K2GAR3 | Uncharacterized protein | 29.206 | 273 | -0.582 | -1.50 | 2.30E-04 | 9.72E-04 | no | no | 0 | 4 | 6 | 11 | 11 | 8 | 12 | 0.010060054 | 0.00995827 | 0.009159402 | 0.012719456 | 0.012038388 | 0.01106402 |
| NITMOv2_3523 | A0A0K2GG39 | Uma2 domain-containing protein | 22.568 | 197 | -0.256 | -1.19 | 8.49E-02 | 1.12E-01 | no | no | 0 | 7 | 8 | 9 | 8 | 9 | 9 | 0.010051623 | 0.009170423 | 0.009665169 | 0.008568334 | 0.008449384 | 0.007852755 |
| NITMOv2_1311 | A0A0K2GA38 | Uncharacterized protein | 8.065 | 70 | 0.592 | 1.51 | 5.43E-03 | 1.06E-02 | no | no | 0 | 3 | 3 | 2 | 2 | 2 | 1 | 0.010017475 | 0.008513004 | 0.010154256 | 0.011495175 | 0.012931133 | 0.01030845 |
| NITMOv2_3828 | A0A0K2GHW2 | Putative Universal stress protein | 34.4 | 321 | 0.841 | 1.79 | 3.57E-05 | 2.79E-04 | no | no | 0 | 10 | 8 | 10 | 5 | 4 | 4 | 0.009998293 | 0.009661434 | 0.010620911 | 0.002234297 | 0.003492986 | 0.003800329 |
| NITMOv2_1138 | A0A0K2G9N3 | Uncharacterized protein | 13.902 | 127 | 0.433 | 1.35 | 1.37E-01 | 1.71E-01 | no | no | 0 | 2 | 4 | 3 | 6 | 5 | 4 | 0.009990073 | 0.009576959 | 0.008607047 | 0.009722871 | 0.009295364 | 0.00749403 |
| devR | A0A0K2G6X7 | Transcriptional regulatory protein DevR (DosR) | 23.422 | 218 | -0.527 | -1.44 | 1.01E-02 | 1.80E-02 | no | no | 0 | 4 | 2 | 3 | 10 | 9 | 9 | 0.009968994 | 0.008742964 | 0.009636027 | 0.020335078 | 0.019189516 | 0.016216357 |
| kbl | A0A0K2GJ14 | 2-amino-3-ketobutyrate coenzyme A ligase | 42.446 | 396 | 0.51 | 1.42 | 8.62E-04 | 2.54E-03 | no | no | 0 | 18 | 19 | 20 | 12 | 13 | 10 | 0.009964567 | 0.010039029 | 0.010083127 | 0.006322191 | 0.006833245 | 0.005851986 |
| NITMOv2_3296 | A0A0K2GFQ7 | Uncharacterized protein | 47.189 | 414 | -0.371 | -1.29 | 1.04E-02 | 1.85E-02 | no | no | 0 | 21 | 16 | 14 | 27 | 27 | 24 | 0.00995993 | 0.008601195 | 0.009959848 | 0.015284255 | 0.015939215 | 0.017552182 |
| NITMOv2_1180 | A0A0K2G9G2 | Uncharacterized protein | 16.66 | 151 | 0.567 | 1.48 | 6.23E-05 | 4.08E-04 | no | no | 0 | 9 | 7 | 4 | 4 | 5 | 6 | 0.009908709 | 0.011000714 | 0.009472487 | 0.012976217 | 0.013630911 | 0.014180884 |
| NITMOv2_4364 | A0A0K2GIG5 | HTH_37 domain-containing protein | 12.782 | 117 | 1.16 | 2.23 | 3.92E-06 | 6.39E-05 | yes | no | 0 | 4 | 4 | 5 | 2 | 3 | 3 | 0.009822075 | 0.007795163 | 0.011137415 | 0.003731034 | 0.003681758 | 0.004946991 |
| cbiC | A0A0K2GAX8 | Cobalt-precorrin-8X methylmutase | 23.096 | 216 | -1.92 | -3.78 | 1.11E-07 | 8.87E-06 | yes | no | 0 | 13 | 11 | 12 | 37 | 37 | 38 | 0.009791932 | 0.011301461 | 0.012443052 | 0.054183439 | 0.054521779 | 0.053404204 |
| NITMOv2_1466 | A0A0K2GAB4 | Uncharacterized protein | 32.682 | 294 | 0.0846 | 1.06 | 2.98E-01 | 3.41E-01 | no | no | 0 | 12 | 12 | 11 | 12 | 8 | 10 | 0.009782868 | 0.010728322 | 0.009566815 | 0.010505741 | 0.011620447 | 0.010283995 |
| murC | A0A0K2G878 | UDP-N-acetylmuramate--L-alanine ligase | 53.573 | 499 | -0.387 | -1.31 | 6.16E-04 | 1.99E-03 | no | no | 0 | 15 | 16 | 17 | 20 | 19 | 21 | 0.009766216 | 0.009780325 | 0.009479773 | 0.011006677 | 0.011287027 | 0.011262173 |
| ilvD.1 | A0A0K2GGT9 | Dihydroxy-acid dehydratase | 58.654 | 557 | -1.45 | -2.73 | 1.51E-06 | 3.61E-05 | yes | no | 0 | 18 | 18 | 20 | 44 | 51 | 47 | 0.00974851 | 0.01000872 | 0.01032489 | 0.035222493 | 0.036276901 | 0.032288865 |
| leuD | A0A0K2GJQ3 | 3-isopropylmalate dehydratase small subunit | 23.203 | 205 | -0.961 | -1.95 | 7.60E-06 | 9.86E-05 | no | no | 0 | 12 | 13 | 12 | 24 | 21 | 25 | 0.009731015 | 0.009339178 | 0.009765441 | 0.021167257 | 0.0223062 | 0.020405081 |
| NITMOv2_0926 | A0A0K2G8U6 | Phosphodiesterase/alkaline phosphatase D | 53.769 | 491 | 0.417 | 1.34 | 9.31E-04 | 2.69E-03 | no | no | 0 | 23 | 22 | 18 | 12 | 13 | 13 | 0.009667146 | 0.009932653 | 0.009576785 | 0.00607387 | 0.006089187 | 0.005797143 |
| NITMOv2_2362 | A0A0K2GDU1 | dITP/XTP pyrophosphatase | 20.92 | 196 | -0.2 | -1.15 | 5.97E-02 | 8.19E-02 | no | no | 0 | 8 | 5 | 5 | 8 | 6 | 4 | 0.009650915 | 0.010683346 | 0.010235739 | 0.011393448 | 0.010633238 | 0.009759124 |
| NITMOv2_2300 | A0A0K2GCM9 | Uncharacterized protein | 46.389 | 424 | 0.0792 | 1.06 | 4.44E-01 | 4.87E-01 | no | no | 0 | 13 | 13 | 16 | 14 | 16 | 19 | 0.009624988 | 0.007760551 | 0.008294729 | 0.007589413 | 0.008831124 | 0.007348562 |
| NITMOv2_4497 | A0A0K2GIV6 | Uncharacterized protein | 39.563 | 365 | -0.00729 | -1.01 | 9.33E-01 | 9.44E-01 | no | no | 0 | 8 | 7 | 11 | 12 | 14 | 14 | 0.009609812 | 0.008802801 | 0.009644655 | 0.010299621 | 0.009261493 | 0.009134097 |
| NITMOv2_2100 | A0A0K2GD45 | Uncharacterized protein | 135.2 | 1224 | 0.805 | 1.75 | 1.20E-04 | 6.31E-04 | no | no | 0 | 29 | 40 | 45 | 26 | 29 | 26 | 0.00959611 | 0.00932197 | 0.009433951 | 0.006676978 | 0.006049258 | 0.004701188 |
| proB | A0A0K2G8X7 | Glutamate 5-kinase | 39.999 | 373 | 0.167 | 1.12 | 3.59E-02 | 5.27E-02 | no | no | 0 | 13 | 11 | 10 | 11 | 13 | 10 | 0.009544046 | 0.009974109 | 0.008220724 | 0.008462905 | 0.009316804 | 0.00912169 |
| NITMOv2_0578 | A0A0K2G834 | NTP_transf_2 domain-containing protein | 12.356 | 109 | 0.795 | 1.74 | 4.16E-05 | 3.06E-04 | no | no | 0 | 3 | 3 | 4 | 3 | 3 | 3 | 0.009536668 | 0.008671004 | 0.00931489 | 0.006804914 | 0.005679481 | 0.007386862 |
| rfbD | A0A0K2GG00 | dTDP-4-dehydrorhamnose reductase | 30.624 | 283 | -0.344 | -1.27 | 8.42E-03 | 1.54E-02 | no | no | 0 | 5 | 7 | 6 | 8 | 9 | 9 | 0.009457623 | 0.009816109 | 0.009642929 | 0.013339296 | 0.016959983 | 0.014387128 |
| glgX | A0A0K2GH55 | Glycogen operon protein GlgX-like protein | 81.013 | 715 | 0.576 | 1.49 | 1.15E-04 | 6.13E-04 | no | no | 0 | 24 | 29 | 27 | 17 | 15 | 23 | 0.009445397 | 0.009842703 | 0.010090987 | 0.005803486 | 0.007707657 | 0.00780061 |
| pepP | A0A0K2GFL6 | Xaa-Pro aminopeptidase | 42.727 | 383 | -0.626 | -1.54 | 1.05E-04 | 5.76E-04 | no | no | 0 | 10 | 12 | 14 | 17 | 21 | 21 | 0.009417362 | 0.008636979 | 0.011047113 | 0.015752171 | 0.017245861 | 0.014897795 |
| NITMOv2_1524 | A0A0K2GAR2 | Uncharacterized protein | 38.681 | 350 | 0.552 | 1.47 | 3.51E-04 | 1.30E-03 | no | no | 0 | 9 | 10 | 11 | 7 | 6 | 10 | 0.009407455 | 0.007152605 | 0.008896358 | 0.006527718 | 0.00596567 | 0.007732641 |
| fabF.2 | A0A0K2GCE9 | 3-oxoacyl-[acyl-carrier-protein] synthase 2 | 43.705 | 418 | -0.825 | -1.77 | 6.05E-04 | 1.96E-03 | no | no | 0 | 12 | 13 | 16 | 23 | 24 | 20 | 0.009393543 | 0.00973007 | 0.011326838 | 0.020215138 | 0.018628637 | 0.016566451 |
| phoB | A0A0K2G852 | DNA-binding response regulator in two-component regulatory system with PhoR (Or CreC) | 26.576 | 236 | -0.172 | -1.13 | 2.03E-01 | 2.43E-01 | no | no | 0 | 12 | 15 | 16 | 13 | 13 | 9 | 0.00938469 | 0.011546282 | 0.011930575 | 0.011567584 | 0.011800519 | 0.011038127 |
| NITMOv2_1338 | A0A0K2G9W8 | Uncharacterized protein | 38.937 | 359 | 0.356 | 1.28 | 1.59E-02 | 2.62E-02 | no | no | 0 | 7 | 7 | 7 | 6 | 5 | 5 | 0.009383004 | 0.009445162 | 0.009443729 | 0.008309351 | 0.00824088 | 0.007124157 |
| fdxB | A0A0K2GI98 | 2Fe-2S ferredoxin | 14.629 | 138 | 0.869 | 1.83 | 2.88E-03 | 6.40E-03 | no | no | 0 | 3 | 2 | 1 | 1 | 1 | 1 | 0.009349067 | 0.008223991 | 0.008031684 | 0.004710251 | 0.004825423 | 0.00436494 |
| NITMOv2_3319 | A0A0K2GGF8 | Uncharacterized protein | 29.151 | 266 | 0.715 | 1.64 | 4.14E-05 | 3.06E-04 | no | no | 0 | 6 | 8 | 6 | 2 | 3 | 4 | 0.009329675 | 0.008791263 | 0.006955923 | 0.004931327 | 0.00467021 | 0.005802538 |
| carA | A0A0K2GGG2 | Carbamoyl-phosphate synthase small chain | 42.918 | 392 | -0.512 | -1.43 | 7.64E-04 | 2.32E-03 | no | no | 0 | 9 | 12 | 14 | 14 | 15 | 15 | 0.009319768 | 0.009278363 | 0.012053087 | 0.020420961 | 0.022226963 | 0.019178762 |
| NITMOv2_1431 | A0A0K2GA81 | Uncharacterized protein | 15.698 | 137 | 1.01 | 2.01 | 7.02E-03 | 1.32E-02 | yes | no | 0 | 1 | 0 | 3 | 2 | 2 | 2 | 0.009277821 | 0.004457619 | 0.004006256 | 0.00180651 | 0.001600136 | 0.001532449 |
| NITMOv2_0432 | A0A0K2G7D6 | YCII domain-containing protein | 16.114 | 143 | 0.194 | 1.14 | 1.41E-01 | 1.75E-01 | no | no | 0 | 7 | 6 | 8 | 6 | 10 | 6 | 0.009262644 | 0.008345619 | 0.010824714 | 0.006827718 | 0.010946927 | 0.009525728 |
| thrC.1 | A0A0K2GIS5 | Threonine synthase | 37.09 | 352 | -0.79 | -1.73 | 4.28E-04 | 1.51E-03 | no | no | 0 | 9 | 9 | 9 | 15 | 18 | 14 | 0.009231237 | 0.007960202 | 0.00911569 | 0.020906646 | 0.021447015 | 0.016331976 |
| adhC | A0A0K2GIJ3 | Alcohol dehydrogenase, NADP-dependent | 37.829 | 347 | 0.15 | 1.11 | 3.74E-02 | 5.47E-02 | no | no | 0 | 15 | 12 | 13 | 12 | 12 | 16 | 0.009229551 | 0.009564053 | 0.009779628 | 0.007542029 | 0.008349483 | 0.008546471 |
| NITMOv2_0946 | A0A0K2G8W4 | Uncharacterized protein | 19.935 | 171 | -0.242 | -1.18 | 4.93E-02 | 6.94E-02 | no | no | 0 | 5 | 5 | 8 | 9 | 10 | 10 | 0.009210791 | 0.009869102 | 0.011312458 | 0.012994726 | 0.012374449 | 0.011090812 |
| NITMOv2_0069 | A0A0K2G6C0 | Oxidoreductase, zinc-binding dehydrogenase family | 34.681 | 332 | 1.01 | 2.01 | 8.96E-04 | 2.61E-03 | yes | no | 0 | 6 | 11 | 10 | 6 | 5 | 4 | 0.009205099 | 0.008901941 | 0.008844784 | 0.002256656 | 0.002207626 | 0.002722176 |
| NITMOv2_0271 | A0A0K2G701 | Aminopeptidase | 101.24 | 901 | 0.189 | 1.14 | 7.41E-02 | 9.92E-02 | no | no | 0 | 29 | 32 | 30 | 33 | 31 | 30 | 0.00920004 | 0.009468823 | 0.008660538 | 0.008243458 | 0.008359892 | 0.007582138 |
| NITMOv2_2020 | A0A0K2GCX6 | Uncharacterized protein | 39.362 | 366 | 0.0974 | 1.07 | 4.01E-01 | 4.45E-01 | no | no | 0 | 22 | 21 | 16 | 15 | 16 | 16 | 0.009158304 | 0.010569344 | 0.010214074 | 0.010412454 | 0.01022462 | 0.00872916 |
| NITMOv2_4629 | A0A0K2GJ80 | Putative Peptidylprolyl isomerase | 37.019 | 325 | 0.302 | 1.23 | 1.04E-02 | 1.85E-02 | no | no | 0 | 16 | 13 | 13 | 13 | 14 | 14 | 0.009144814 | 0.009273279 | 0.009393689 | 0.008567742 | 0.009045066 | 0.007727426 |
| cpsB | A0A0K2GG84 | Mannose-1-phosphate guanylyltransferase | 58.278 | 525 | -0.355 | -1.28 | 6.53E-03 | 1.24E-02 | no | no | 0 | 17 | 20 | 22 | 25 | 26 | 25 | 0.009132588 | 0.00924297 | 0.009777136 | 0.01320899 | 0.012532924 | 0.011837213 |
| ilvE.1 | A0A0K2GI80 | Putative branched-chain-amino-acid aminotransferase | 32.174 | 290 | 0.661 | 1.58 | 1.85E-03 | 4.51E-03 | no | no | 0 | 9 | 7 | 6 | 2 | 2 | 2 | 0.009127108 | 0.009134639 | 0.0071858 | 0.004409068 | 0.005327417 | 0.005615173 |
| carB | A0A0K2GFT3 | Carbamoyl-phosphate synthase large chain | 117.95 | 1084 | -0.427 | -1.34 | 1.35E-03 | 3.54E-03 | no | no | 0 | 37 | 41 | 31 | 58 | 60 | 55 | 0.009116779 | 0.008376319 | 0.007951544 | 0.012003219 | 0.01275681 | 0.011894213 |
| lolC | A0A0K2GDK8 | Lipoprotein-releasing system, transmembrane protein LolC | 46.632 | 425 | 0.00674 | 1.00 | 9.35E-01 | 9.44E-01 | no | no | 0 | 10 | 11 | 10 | 9 | 10 | 11 | 0.00909296 | 0.00943167 | 0.008353205 | 0.009489654 | 0.008935376 | 0.00849954 |
| NITMOv2_0398 | A0A0K2G7I2 | DZANK-type domain-containing protein | 23.908 | 216 | 0.544 | 1.46 | 9.62E-04 | 2.73E-03 | no | no | 0 | 5 | 4 | 3 | 2 | 2 | 0 | 0.009077573 | 0.005316448 | 0.004937266 | 0.00570087 | 0.007628419 | 0.006769387 |
| NITMOv2_2522 | A0A0K2GDB0 | Uncharacterized protein | 17.65 | 163 | 0.695 | 1.62 | 3.59E-05 | 2.79E-04 | no | no | 0 | 7 | 7 | 6 | 3 | 3 | 4 | 0.009065347 | 0.009184698 | 0.00934154 | 0.005647711 | 0.005417686 | 0.005476538 |
| cmk | A0A0K2G758 | Cytidylate kinase | 23.875 | 223 | -0.542 | -1.46 | 8.25E-04 | 2.45E-03 | no | no | 0 | 6 | 6 | 11 | 14 | 13 | 10 | 0.009062396 | 0.009297918 | 0.009462134 | 0.013829126 | 0.012101933 | 0.012137139 |
| NITMOv2_3577 | A0A0K2GGL0 | Putative Outer membrane efflux protein | 57.121 | 520 | 0.473 | 1.39 | 9.45E-04 | 2.70E-03 | no | no | 0 | 19 | 22 | 24 | 30 | 26 | 30 | 0.009052067 | 0.011086167 | 0.010615351 | 0.009620848 | 0.008018393 | 0.007499065 |
| NITMOv2_1662 | A0A0K2GB38 | HTH_37 domain-containing protein | 11.913 | 107 | 0.779 | 1.72 | 8.21E-05 | 4.96E-04 | no | no | 0 | 4 | 5 | 4 | 2 | 1 | 0 | 0.009028459 | 0.009338004 | 0.018063621 | 0.005961481 | 0.005184168 | 0.006471079 |
| glmU | A0A0K2GFU2 | Bifunctional protein GlmU | 55.889 | 521 | -0.066 | -1.05 | 4.39E-01 | 4.82E-01 | no | no | 0 | 13 | 18 | 21 | 26 | 24 | 21 | 0.008981875 | 0.010184318 | 0.010432255 | 0.011261366 | 0.010345186 | 0.009645303 |
| NITMOv2_3653 | A0A0K2GGG1 | Uncharacterized protein | 26.505 | 237 | 0.873 | 1.83 | 7.43E-06 | 9.75E-05 | no | no | 0 | 3 | 3 | 2 | 1 | 1 | 1 | 0.00897239 | 0.009567182 | 0.009218262 | 0.005535175 | 0.006039935 | 0.005453702 |
| NITMOv2_2380 | A0A0K2GCV9 | Uncharacterized protein | 10.502 | 89 | 0.153 | 1.11 | 2.87E-01 | 3.30E-01 | no | no | 0 | 5 | 5 | 6 | 5 | 4 | 4 | 0.008970914 | 0.009959247 | 0.009658459 | 0.009121985 | 0.007854946 | 0.006145439 |
| top6A | A0A0K2GCU4 | Type 2 DNA topoisomerase 6 subunit A | 42.054 | 367 | -0.665 | -1.59 | 2.11E-04 | 9.17E-04 | no | no | 0 | 14 | 14 | 15 | 24 | 22 | 20 | 0.008938031 | 0.009384153 | 0.008895207 | 0.014703359 | 0.015355031 | 0.013451386 |
| hisG | A0A0K2G8S0 | ATP phosphoribosyltransferase | 25.933 | 236 | 0.303 | 1.23 | 2.38E-03 | 5.51E-03 | no | no | 0 | 8 | 9 | 8 | 8 | 9 | 7 | 0.008895031 | 0.00933292 | 0.010286929 | 0.008804069 | 0.008789019 | 0.006432779 |
| NITMOv2_1081 | A0A0K2GAA0 | Methyltransf_25 domain-containing protein | 24.415 | 219 | 0.0398 | 1.03 | 7.29E-01 | 7.61E-01 | no | no | 0 | 4 | 4 | 8 | 8 | 9 | 9 | 0.008885124 | 0.009259591 | 0.01115467 | 0.009683779 | 0.005782491 | 0.00988733 |
| purQ | A0A0K2G9V6 | Phosphoribosylformylglycinamidine synthase subunit PurQ | 25.486 | 239 | 0.0512 | 1.04 | 5.22E-01 | 5.64E-01 | no | no | 0 | 11 | 13 | 16 | 9 | 8 | 10 | 0.008867417 | 0.010129371 | 0.010411357 | 0.009230671 | 0.010758465 | 0.008552225 |
| NITMOv2_3029 | A0A0K2GF16 | Uncharacterized protein | 55.855 | 530 | 0.132 | 1.10 | 4.19E-01 | 4.63E-01 | no | no | 0 | 10 | 8 | 15 | 9 | 5 | 7 | 0.008859197 | 0.007264457 | 0.009748569 | 0.005244652 | 0.005322601 | 0.005412525 |
| NITMOv2_1720 | A0A0K2GB17 | Uncharacterized protein | 11.572 | 100 | 0.367 | 1.29 | 1.81E-03 | 4.45E-03 | no | no | 0 | 3 | 3 | 4 | 4 | 4 | 3 | 0.008836642 | 0.008190553 | 0.009281339 | 0.007358565 | 0.007299971 | 0.006675705 |
| moaC | A0A0K2GAF4 | Cyclic pyranopterin monophosphate synthase | 18.076 | 170 | -0.335 | -1.26 | 9.99E-03 | 1.78E-02 | no | no | 0 | 3 | 6 | 5 | 7 | 8 | 7 | 0.008823573 | 0.008809645 | 0.008258877 | 0.014066194 | 0.013956096 | 0.013736388 |
| atoC.5 | A0A0K2GF92 | Acetoacetate metabolism regulatory protein AtoC | 51.768 | 463 | 0.393 | 1.31 | 2.13E-03 | 5.07E-03 | no | no | 0 | 20 | 22 | 20 | 19 | 22 | 15 | 0.008796171 | 0.008688016 | 0.008196758 | 0.009438272 | 0.009653332 | 0.008050369 |
| NITMOv2_3952 | A0A0K2GI95 | DrsE domain-containing protein | 12.407 | 119 | 1.29 | 2.45 | 9.46E-06 | 1.13E-04 | yes | no | 0 | 3 | 2 | 2 | 2 | 1 | 1 | 0.008770877 | 0.009066198 | 0.008409763 | 0.004832413 | 0.00527568 | 0.005096955 |
| NITMOv2_1050 | A0A0K2G939 | PINc domain-containing protein | 16.262 | 145 | -0.212 | -1.16 | 2.73E-01 | 3.16E-01 | no | no | 0 | 3 | 3 | 6 | 6 | 7 | 3 | 0.008761602 | 0.008316287 | 0.008652102 | 0.010036345 | 0.009172001 | 0.008546651 |
| NITMOv2_0247 | A0A0K2G6T5 | N-acetyltransferase, GNAT family | 18.409 | 158 | 0.00222 | 1.00 | 9.74E-01 | 9.77E-01 | no | no | 0 | 7 | 7 | 7 | 4 | 6 | 4 | 0.008756121 | 0.004319174 | 0.004084863 | 0.004551071 | 0.004267496 | 0.004129206 |
| phrA | A0A0K2G6U9 | Deoxyribodipyrimidine photo-lyase | 54.559 | 480 | 0.539 | 1.45 | 2.34E-04 | 9.81E-04 | no | no | 0 | 17 | 17 | 17 | 15 | 12 | 9 | 0.00875317 | 0.009007731 | 0.008962502 | 0.005869675 | 0.005614538 | 0.005237388 |
| aroB | A0A0K2G9M9 | 3-dehydroquinate synthase | 41.413 | 376 | -0.167 | -1.12 | 3.93E-02 | 5.69E-02 | no | no | 0 | 19 | 17 | 20 | 20 | 19 | 20 | 0.008749376 | 0.00894633 | 0.009821424 | 0.009593898 | 0.009795649 | 0.009955299 |
| pncB | A0A0K2GH92 | Nicotinate phosphoribosyltransferase | 49.902 | 457 | 0.911 | 1.88 | 1.60E-05 | 1.60E-04 | no | no | 0 | 16 | 19 | 19 | 5 | 6 | 6 | 0.0087342 | 0.00939745 | 0.008900767 | 0.00276855 | 0.0027626 | 0.002432499 |

| | | | | | | | | | | | | | | | | | | | | | | | |
|---|---|---|---|---|---|---|---|---|---|---|---|---|---|---|---|---|---|---|---|---|---|---|---|
| gltD | A0A0K2GGQ9 | Glutamate synthase [NADPH] small chain | 52.38 | 478 | -0.795 | -1.74 | 3.63E-05 | 2.81E-04 | no | no | 0 | 14 | 14 | 13 | 30 | 32 | 30 | 0.008728719 | 0.00854742 | 0.00879647 | 0.017212186 | 0.017158855 | 0.01792655 |
| NITMOv2_0509 | A0A0K2G7P5 | UPF0114 protein NITMOv2_0509 | 21.428 | 189 | 0.0661 | 1.05 | 3.38E-01 | 3.81E-01 | no | no | 0 | 4 | 4 | 6 | 7 | 7 | 5 | 0.008724925 | 0.008805929 | 0.00922919 | 0.009693108 | 0.011026631 | 0.010046464 |
| lon | A0A0K2GCP9 | Lon protease | 87.807 | 794 | 0.437 | 1.35 | 3.36E-02 | 4.97E-02 | no | no | 0 | 35 | 31 | 41 | 24 | 23 | 22 | 0.008715018 | 0.007048185 | 0.009831394 | 0.004446235 | 0.005055368 | 0.004149884 |
| NITMOv2_3142 | A0A0K2GF10 | SpoVT-AbrB domain-containing protein | 10.807 | 98 | 0.338 | 1.26 | 4.59E-03 | 9.25E-03 | no | no | 0 | 4 | 5 | 3 | 2 | 2 | 2 | 0.008706797 | 0.008494037 | 0.006977013 | 0.004938138 | 0.005569792 | 0.004837845 |
| NITMOv2_3837 | A0A0K2GH09 | DLH domain-containing protein | 23.003 | 212 | 0.656 | 1.58 | 3.15E-04 | 1.19E-03 | no | no | 0 | 10 | 9 | 10 | 7 | 5 | 7 | 0.008705743 | 0.00902005 | 0.009413436 | 0.006078312 | 0.005439127 | 0.005139031 |
| hypB | A0A088ND36 | Hydrogenase nickel incorporation protein | 28.59 | 258 | 1.82 | 3.53 | 6.85E-06 | 9.26E-05 | yes | no | 0 | 7 | 7 | 6 | 2 | 2 | 2 | 0.008693307 | 0.010399612 | 0.00816129 | 0.002517267 | 0.002382725 | 0.001978113 |
| NITMOv2_3646 | A0A0K2GGF1 | FGE-sulfatase domain-containing protein | 34.454 | 313 | 0.018 | 1.01 | 7.96E-01 | 8.21E-01 | no | no | 0 | 6 | 10 | 9 | 10 | 10 | 9 | 0.008677076 | 0.008850709 | 0.009290925 | 0.008913644 | 0.008497548 | 0.008781126 |
| NITMOv2_1890 | A0A0K2GBI1 | Uncharacterized protein | 23.467 | 203 | 0.775 | 1.71 | 3.97E-03 | 8.25E-03 | no | no | 0 | 7 | 6 | 6 | 4 | 5 | 5 | 0.008669698 | 0.009159473 | 0.009446796 | 0.004806056 | 0.007083698 | 0.006061287 |
| NITMOv2_3926 | A0A0K2GHA0 | Uncharacterized protein | 9.7723 | 86 | 0.403 | 1.32 | 7.01E-02 | 9.46E-02 | no | no | 0 | 2 | 3 | 3 | 3 | 2 | 1 | 0.008662321 | 0.011310847 | 0.008564101 | 0.006009606 | 0.005800824 | 0.004936922 |
| cinA | A0A0K2GDC1 | CinA-like protein | 45.179 | 430 | 0.468 | 1.38 | 8.92E-03 | 1.62E-02 | no | no | 0 | 13 | 14 | 11 | 7 | 10 | 8 | 0.008639345 | 0.009129554 | 0.008920515 | 0.005230289 | 0.005877576 | 0.006377037 |
| nuoF.1 | A0A0K2GHL7 | NADH-quinone oxidoreductase, subunit F | 47.063 | 430 | -0.556 | -1.47 | 1.68E-04 | 7.81E-04 | no | no | 0 | 14 | 14 | 14 | 25 | 25 | 25 | 0.008612364 | 0.008252931 | 0.009322368 | 0.015509328 | 0.016094583 | 0.014778939 |
| NITMOv2_1011 | A0A0K2G917 | Putative Mercuric reductase | 55.901 | 517 | 0.0962 | 1.07 | 1.98E-01 | 2.37E-01 | no | no | 0 | 10 | 11 | 10 | 5 | 7 | 6 | 0.008586859 | 0.009657132 | 0.009262742 | 0.008252935 | 0.009201987 | 0.008801624 |
| NITMOv2_2403 | A0A0K2GCZ1 | MPN domain-containing protein | 18.889 | 169 | 0.15 | 1.11 | 1.32E-01 | 1.65E-01 | no | no | 0 | 5 | 5 | 7 | 6 | 8 | 6 | 0.008542804 | 0.009134639 | 0.008151512 | 0.008312313 | 0.008244764 | 0.008185947 |
| bipA | A0A0K2GJ89 | GTP-binding protein | 68.412 | 618 | -1.16 | -2.23 | 1.98E-06 | 4.19E-05 | yes | no | 0 | 11 | 17 | 15 | 30 | 34 | 28 | 0.008498117 | 0.008659467 | 0.009473062 | 0.020194407 | 0.0219473 | 0.019488039 |
| NITMOv2_1027 | A0A0K2G931 | Uncharacterized protein | 12.086 | 107 | -2.03 | -4.08 | 2.28E-06 | 4.57E-05 | yes | no | 0 | 2 | 2 | 2 | 4 | 4 | 5 | 0.008492215 | 0.009328032 | 0.007913966 | 0.03791745 | 0.036865746 | 0.035912079 |
| NITMOv2_0570 | A0A0K2G7R1 | Uncharacterized protein | 16.093 | 149 | 0.193 | 1.14 | 2.81E-01 | 3.23E-01 | no | no | 0 | 3 | 3 | 3 | 4 | 4 | 4 | 0.008486946 | 0.008387856 | 0.008828104 | 0.00786113 | 0.008081939 | 0.007115706 |
| focA | A0A0K2GGZ4 | Formate transporter | 30.609 | 298 | 1.01 | 2.01 | 1.02E-04 | 5.64E-04 | yes | no | 0 | 5 | 4 | 5 | 3 | 2 | 3 | 0.008485048 | 0.010182167 | 0.008827337 | 0.005356597 | 0.005162727 | 0.004731037 |
| NITMOv2_3158 | A0A0K2GG18 | ANK_REP_REGION domain-containing protein | 58.376 | 527 | -0.000555 | -1.00 | 9.94E-01 | 9.94E-01 | no | no | 0 | 24 | 23 | 24 | 21 | 19 | 20 | 0.00847746 | 0.008988372 | 0.008954258 | 0.007590005 | 0.008576941 | 0.008161133 |
| NITMOv2_2323 | A0A0K2GCR3 | Cupin_7 domain-containing protein | 13.751 | 125 | 0.177 | 1.13 | 2.93E-01 | 3.35E-01 | no | no | 0 | 5 | 2 | 4 | 3 | 3 | 2 | 0.008470715 | 0.006008674 | 0.008605897 | 0.006034038 | 0.005459169 | 0.003834494 |
| NITMOv2_3585 | A0A0K2GH91 | Uncharacterized protein | 21.082 | 191 | 0.464 | 1.38 | 3.95E-04 | 1.41E-03 | no | no | 0 | 4 | 4 | 6 | 4 | 4 | 5 | 0.008470293 | 0.008470963 | 0.008281884 | 0.006068391 | 0.006767369 | 0.006381173 |
| NITMOv2_2600 | A0A0K2GDI5 | Uncharacterized protein | 21.521 | 189 | 0.043 | 1.03 | 5.81E-01 | 6.22E-01 | no | no | 0 | 10 | 8 | 10 | 9 | 9 | 7 | 0.008449636 | 0.008365955 | 0.009610528 | 0.00826715 | 0.008129947 | 0.008116 |
| trpE | A0A0K2GCN8 | Anthranilate synthase component 1 | 55.765 | 498 | -0.385 | -1.31 | 2.35E-03 | 5.44E-03 | no | no | 0 | 10 | 13 | 14 | 27 | 24 | 23 | 0.008442469 | 0.008125632 | 0.01026699 | 0.015062143 | 0.014302878 | 0.011783449 |
| NITMOv2_0929 | A0A0K2G8S9 | Bifunctional deaminase-reductase domain protein | 20.507 | 186 | 1.07 | 2.10 | 1.37E-01 | 1.71E-01 | no | yes | 1 | 3 | 3 | 5 | 1 | 2 | 2 | 0.008429611 | 0.009125252 | 0.00931489 | 0.004685227 | 0.005735725 | 0.005348872 |
| NITMOv2_0117 | A0A0K2G7H6 | Uncharacterized protein | 42.94 | 378 | 0.195 | 1.14 | 1.02E-01 | 1.31E-01 | no | no | 0 | 10 | 14 | 12 | 14 | 13 | 13 | 0.008423288 | 0.007870447 | 0.008177203 | 0.008129737 | 0.008080851 | 0.008548089 |
| NITMOv2_3257 | A0A0K2GG98 | Sigma-54 dependent transcriptional regulator | 51.594 | 467 | 0.374 | 1.30 | 2.17E-03 | 5.13E-03 | no | no | 0 | 16 | 16 | 19 | 16 | 16 | 15 | 0.008409586 | 0.008568539 | 0.008450409 | 0.007410391 | 0.007364604 | 0.007214782 |
| atoC.7 | A0A0K2GID8 | Acetoacetate metabolism regulatory protein AtoC | 49.677 | 451 | 0.162 | 1.12 | 5.08E-02 | 7.13E-02 | no | no | 0 | 21 | 24 | 20 | 25 | 27 | 19 | 0.008405371 | 0.011158518 | 0.010712172 | 0.009293307 | 0.00831266 | 0.008185228 |
| ispU | A0A0K2GA60 | Isoprenyl transferase | 29.897 | 261 | -0.582 | -1.50 | 1.36E-02 | 2.30E-02 | no | no | 0 | 4 | 3 | 2 | 4 | 6 | 4 | 0.00835689 | 0.004532708 | 0.009287474 | 0.008659696 | 0.010682179 | 0.009414064 |
| NITMOv2_1885 | A0A0K2GBH6 | Plasmid stabilization system protein | 11.543 | 97 | 0.979 | 1.97 | 2.74E-05 | 2.33E-04 | no | no | 0 | 5 | 4 | 4 | 1 | 3 | 3 | 0.008325061 | 0.008924038 | 0.00824814 | 0.004574319 | 0.004026365 | 0.004578017 |
| prsA | A0A0K2G8Y7 | Ribose-phosphate pyrophosphokinase | 33.926 | 313 | -1.63 | -3.10 | 1.23E-06 | 3.15E-05 | yes | no | 0 | 6 | 6 | 9 | 24 | 25 | 24 | 0.008295339 | 0.006703246 | 0.008407079 | 0.038416461 | 0.038389908 | 0.044059367 |
| NITMOv2_3630 | A0A0K2GHD4 | Sigma-54 dependent transcriptional regulator | 53.035 | 482 | 0.158 | 1.12 | 1.18E-01 | 1.50E-01 | no | no | 0 | 19 | 19 | 23 | 17 | 19 | 18 | 0.008293653 | 0.009098072 | 0.009075236 | 0.007688919 | 0.006855463 | 0.006370924 |
| ispD | A0A0K2GGX3 | 2-C-methyl-D-erythritol 4-phosphate cytidylyltransferase | 23.929 | 218 | -0.329 | -1.26 | 1.79E-02 | 2.90E-02 | no | no | 0 | 9 | 9 | 10 | 11 | 12 | 11 | 0.008285222 | 0.009178636 | 0.010724826 | 0.011584316 | 0.011243369 | 0.010142124 |
| radA | A0A0K2GBC9 | DNA repair protein RadA | 48.583 | 453 | -0.0229 | -1.02 | 8.16E-01 | 8.38E-01 | no | no | 0 | 10 | 13 | 14 | 22 | 17 | 16 | 0.008270045 | 0.007998333 | 0.009130452 | 0.008422925 | 0.007955314 | 0.007241574 |
| tpiA | A0A0K2G987 | Triosephosphate isomerase | 27.608 | 258 | -0.204 | -1.15 | 7.04E-02 | 9.49E-02 | no | no | 0 | 11 | 12 | 9 | 10 | 8 | 9 | 0.008266672 | 0.007534307 | 0.009125084 | 0.010757764 | 0.011108976 | 0.009534898 |
| topA | A0A0K2G8P4 | DNA topoisomerase 1 | 87.265 | 780 | -0.416 | -1.33 | 5.30E-04 | 1.78E-03 | no | no | 0 | 21 | 30 | 29 | 37 | 40 | 37 | 0.008262878 | 0.008732014 | 0.008261753 | 0.012445814 | 0.013030724 | 0.016574723 |
| polA | A0A0K2GIU7 | DNA polymerase I | 97.192 | 878 | -0.0282 | -1.02 | 6.92E-01 | 7.27E-01 | no | no | 0 | 29 | 29 | 27 | 29 | 25 | 29 | 0.008260138 | 0.008435178 | 0.007538763 | 0.010391724 | 0.009751058 | 0.009373067 |
| panB | A0A0K2G807 | 3-methyl-2-oxobutanoate hydroxymethyltransferase | 27.891 | 261 | -0.746 | -1.68 | 6.61E-04 | 2.09E-03 | no | no | 0 | 4 | 6 | 6 | 11 | 11 | 11 | 0.008259295 | 0.003786708 | 0.005829931 | 0.009170849 | 0.010465596 | 0.010504445 |
| thrH | A0A0K2GIK2 | Homoserine kinase | 23.282 | 205 | 0.0276 | 1.02 | 8.07E-01 | 8.30E-01 | no | no | 0 | 7 | 7 | 8 | 7 | 8 | 7 | 0.008254868 | 0.007743735 | 0.007533395 | 0.008766458 | 0.008016063 | 0.007267647 |
| NITMOv2_0828 | A0A0K2G8H6 | Uncharacterized protein | 11.932 | 102 | -0.68 | -1.60 | 2.37E-02 | 3.68E-02 | no | no | 0 | 1 | 1 | 1 | 0 | 2 | 0 | 0.008250863 | 0.007845026 | 0.005966246 | 0.005690505 | 0.006603456 | 0.00591474 |
| NITMOv2_0999 | A0A0K2G8Z2 | Uncharacterized protein | 16.423 | 151 | 0.704 | 1.63 | 2.25E-04 | 9.63E-04 | no | no | 0 | 3 | 3 | 3 | 4 | 3 | 3 | 0.008247701 | 0.008650276 | 0.008357039 | 0.005624168 | 0.005256259 | 0.004543673 |
| NITMOv2_2263 | A0A0K2GCT6 | TPR_REGION domain-containing protein | 15.009 | 142 | -0.358 | -1.28 | 1.90E-03 | 4.62E-03 | no | no | 0 | 3 | 4 | 5 | 7 | 4 | 5 | 0.008246648 | 0.007890588 | 0.008702909 | 0.013218023 | 0.01319992 | 0.011118683 |
| NITMOv2_2773 | A0A0K2GEZ0 | Prevent-host-death family protein | 9.0274 | 82 | 1.02 | 2.03 | 2.32E-03 | 5.39E-03 | yes | no | 0 | 2 | 4 | 4 | 3 | 1 | 1 | 0.008223461 | 0.008415624 | 0.006513617 | 0.004496876 | 0.004526805 | 0.004693456 |
| NITMOv2_1187 | A0A0K2G9G7 | Uncharacterized protein | 34.097 | 299 | 0.416 | 1.33 | 2.53E-03 | 5.76E-03 | no | no | 0 | 10 | 10 | 8 | 1 | 4 | 4 | 0.0082125 | 0.007985231 | 0.007900162 | 0.003668843 | 0.005608012 | 0.004979358 |
| aceF | A0A0K2GHK3 | Dihydrolipoamide acetyltransferase component of pyruvate dehydrogenase complex | 47.483 | 443 | 0.525 | 1.44 | 1.98E-02 | 3.15E-02 | no | no | 0 | 10 | 11 | 14 | 13 | 14 | 10 | 0.008198588 | 0.008054258 | 0.010329875 | 0.006015084 | 0.005572744 | 0.004617576 |
| NITMOv2_3919 | A0A0K2GHG9 | Putative Fimbriae Z protein | 21.479 | 195 | -0.125 | -1.09 | 2.47E-01 | 2.88E-01 | no | no | 0 | 10 | 10 | 8 | 13 | 9 | 8 | 0.008163176 | 0.007951598 | 0.007936014 | 0.009801795 | 0.009484136 | 0.007574047 |
| NITMOv2_0334 | A0A0K2G838 | Putative Sulfite reductase, contains SirA-like domain | 90.617 | 832 | -1.42 | -2.68 | 4.44E-07 | 1.75E-05 | yes | no | 0 | 29 | 28 | 31 | 80 | 77 | 79 | 0.008122704 | 0.007386867 | 0.008160139 | 0.024484127 | 0.025227121 | 0.025477581 |
| NITMOv2_2416 | A0A0K2GCZ4 | Putative Helicase (N-terminal) | 39.083 | 353 | -0.401 | -1.32 | 1.49E-02 | 2.50E-02 | no | no | 0 | 11 | 12 | 11 | 11 | 13 | 17 | 0.008111533 | 0.006628548 | 0.009057981 | 0.007064489 | 0.009429602 | 0.009395184 |
| NITMOv2_4828 | A0A0K2GJT0 | TPR_REGION domain-containing protein | 63.389 | 575 | 0.452 | 1.37 | 2.51E-03 | 5.74E-03 | no | no | 0 | 16 | 17 | 15 | 10 | 11 | 11 | 0.008099939 | 0.007934781 | 0.007804684 | 0.005196676 | 0.005869807 | 0.005697887 |
| NITMOv2_3593 | A0A0K2GGA6 | PDZ domain-containing protein | 44.999 | 412 | 0.409 | 1.33 | 6.62E-03 | 1.26E-02 | no | no | 0 | 7 | 4 | 6 | 5 | 5 | 8 | 0.00806811 | 0.006982873 | 0.007396312 | 0.006668685 | 0.007160916 | 0.006163241 |
| NITMOv2_4496 | A0A0K2GIV2 | Uncharacterized protein | 27.301 | 254 | -0.0348 | -1.02 | 7.60E-01 | 7.88E-01 | no | no | 0 | 7 | 6 | 8 | 9 | 11 | 9 | 0.008063895 | 0.0076882 | 0.007670669 | 0.005339568 | 0.00843074 | 0.007518844 |
| pgm | A0A0K2GD12 | Phosphoglucomutase | 59.165 | 547 | 0.178 | 1.13 | 6.15E-02 | 8.41E-02 | no | no | 0 | 19 | 18 | 19 | 19 | 17 | 21 | 0.007993281 | 0.007647136 | 0.008067728 | 0.006319674 | 0.006476054 | 0.006723715 |
| NITMOv2_2424 | A0A0K2GE06 | Putative Steroid Delta-isomerase | 14.171 | 130 | 1.39 | 2.62 | 3.09E-03 | 6.76E-03 | yes | yes | 3 | 2 | 2 | 2 | 1 | 1 | 1 | 0.007982741 | 0.009167099 | 0.008325597 | 0.003866819 | 0.005431825 | 0.005316685 |
| gpsA | A0A0K2G7I1 | Glycerol-3-phosphate dehydrogenase [NAD(P)+] | 35.827 | 341 | 0.0527 | 1.04 | 5.29E-01 | 5.71E-01 | no | no | 0 | 9 | 9 | 13 | 13 | 14 | 12 | 0.007944589 | 0.009651461 | 0.007959596 | 0.00721745 | 0.007913364 | 0.009387272 |
| NITMOv2_0151 | A0A0K2G6J1 | Rhodanese-like sulfurtransferase | 15.687 | 142 | 1.79 | 3.46 | 4.99E-03 | 9.93E-03 | yes | yes | 1 | 3 | 3 | 3 | 1 | 1 | 1 | 0.007933417 | 0.007442988 | 0.009082713 | 0.003847865 | 0.001041355 | 0.003756096 |
| NITMOv2_0702 | A0A0K2G855 | Uncharacterized protein | 45.728 | 402 | 1.71 | 3.27 | 1.44E-05 | 1.48E-04 | yes | no | 0 | 8 | 10 | 7 | 0 | 1 | 0 | 0.007928358 | 0.010198202 | 0.008140775 | 0.002415984 | 0.002029884 | 0.002243336 |
| leuC | A0A0K2GK01 | 3-isopropylmalate dehydratase large subunit | 50.342 | 467 | -1.33 | -2.51 | 2.56E-05 | 2.23E-04 | yes | no | 0 | 9 | 8 | 10 | 33 | 33 | 28 | 0.007924985 | 0.008373191 | 0.007599539 | 0.022178606 | 0.022727248 | 0.020198297 |
| NITMOv2_2059 | A0A0K2GBZ5 | RCK C-terminal domain-containing protein | 23.687 | 215 | 0.469 | 1.38 | 2.86E-03 | 6.37E-03 | no | no | 0 | 4 | 4 | 4 | 7 | 7 | 4 | 0.007888098 | 0.008821964 | 0.007392478 | 0.005697761 | 0.005784976 | 0.005668038 |
| ftsY | A0A0K2GIP2 | Signal recognition particle receptor FtsY | 32.783 | 311 | -0.12 | -1.09 | 3.79E-01 | 4.22E-01 | no | no | 0 | 4 | 3 | 9 | 17 | 14 | 12 | 0.007886411 | 0.010027297 | 0.010996881 | 0.012186091 | 0.012349745 | 0.009624265 |
| NITMOv2_1337 | A0A0K2G9Z4 | Uncharacterized protein | 17.521 | 154 | 0.393 | 1.31 | 6.54E-02 | 8.86E-02 | no | no | 0 | 6 | 3 | 7 | 2 | 5 | 5 | 0.007881142 | 0.008848753 | 0.009310289 | 0.007581713 | 0.008140978 | 0.006664737 |
| NITMOv2_4094 | A0A0K2GHS2 | Uncharacterized protein | 21.196 | 195 | 0.764 | 1.70 | 3.09E-02 | 4.64E-02 | no | no | 0 | 2 | 4 | 3 | 1 | 2 | 1 | 0.007865122 | 0.008001071 | 0.007234306 | 0.003096535 | 0.003055003 | 0.002847145 |
| NITMOv2_3048 | A0A0K2GFT0 | Uncharacterized protein | 42.063 | 364 | 0.515 | 1.43 | 7.37E-04 | 2.27E-03 | no | no | 0 | 16 | 18 | 20 | 10 | 8 | 11 | 0.007862171 | 0.006337383 | 0.007954995 | 0.00544174 | 0.005768197 | 0.005182905 |
| NITMOv2_1322 | A0A0K2G9Y2 | O-methyltransferase, family 2:Generic methyltransferase | 39.385 | 355 | 0.439 | 1.36 | 9.90E-04 | 2.79E-03 | no | no | 0 | 8 | 8 | 7 | 8 | 7 | 6 | 0.00785121 | 0.007682921 | 0.007914541 | 0.005734631 | 0.005654778 | 0.005377821 |
| NITMOv2_3340 | A0A0K2GFJ8 | Uncharacterized protein | 83.684 | 731 | -0.631 | -1.55 | 3.49E-04 | 1.30E-03 | no | no | 0 | 24 | 26 | 24 | 39 | 38 | 36 | 0.007825283 | 0.008353832 | 0.007556402 | 0.0160498 | 0.014363316 | 0.013869809 |
| trpD.1 | A0A0K2GCM4 | Anthranilate phosphoribosyltransferase | 36 | 341 | -0.186 | -1.14 | 4.71E-02 | 6.67E-02 | no | no | 0 | 5 | 6 | 7 | 4 | 4 | 5 | 0.007801253 | 0.008005177 | 0.007968607 | 0.008076874 | 0.008458862 | 0.00866209 |
| aroK | A0A0K2G759 | Shikimate kinase | 18.337 | 166 | 0.287 | 1.22 | 2.42E-02 | 3.76E-02 | no | no | 0 | 6 | 6 | 7 | 6 | 6 | 7 | 0.007774694 | 0.009180983 | 0.008023824 | 0.003990165 | 0.005150453 | 0.005349411 |
| corA | A0A0K2GKM9 | Magnesium transport protein CorA | 39.841 | 351 | 0.202 | 1.15 | 9.09E-03 | 1.64E-02 | no | no | 0 | 8 | 7 | 13 | 10 | 9 | 8 | 0.007766895 | 0.005919701 | 0.007476453 | 0.006273178 | 0.00570434 | 0.004568847 |
| actP | A0A0K2GE73 | P-type copper-exporting ATPase | 95.814 | 897 | 0.734 | 1.66 | 9.21E-04 | 2.67E-03 | no | no | 0 | 15 | 17 | 16 | 10 | 6 | 6 | 0.007753615 | 0.008173736 | 0.007450762 | 0.005086213 | 0.004715112 | 0.004638254 |
| NITMOv2_3659 | A0A0K2GGI4 | Uncharacterized protein | 33.339 | 303 | -0.42 | -1.34 | 1.90E-02 | 3.05E-02 | no | no | 0 | 13 | 13 | 12 | 14 | 17 | 15 | 0.007715252 | 0.008244132 | 0.006730456 | 0.012604994 | 0.011416294 | 0.011945999 |
| NITMOv2_2228 | A0A0K2GCQ4 | Putative cyclase | 29.983 | 276 | 0.441 | 1.36 | 3.07E-03 | 6.74E-03 | no | no | 0 | 5 | 6 | 5 | 4 | 5 | 4 | 0.007714619 | 0.007177048 | 0.007408966 | 0.005507929 | 0.005630385 | 0.006203698 |
| NITMOv2_0938 | A0A0K2G9V2 | Uncharacterized protein | 9.2015 | 77 | 0.286 | 1.22 | 7.30E-02 | 9.81E-02 | no | no | 0 | 2 | 3 | 3 | 2 | 1 | 1 | 0.007707242 | 0.006521585 | 0.007215901 | 0.005674661 | 0.005350723 | 0.005265439 |
| speB | A0A0K2GIH9 | Agmatinase 1 | 33.613 | 310 | -0.111 | -1.08 | 3.01E-01 | 3.43E-01 | no | no | 0 | 6 | 6 | 5 | 2 | 5 | 3 | 0.007691011 | 0.008293213 | 0.007055811 | 0.008256785 | 0.008465698 | 0.00715904 |
| glmM | A0A0K2GC66 | Phosphoglucosamine mutase | 49.774 | 457 | -0.445 | -1.36 | 9.35E-04 | 2.69E-03 | no | no | 0 | 9 | 10 | 12 | 20 | 18 | 18 | 0.007685531 | 0.007365162 | 0.008854945 | 0.012557462 | 0.013085724 | 0.01206917 |
| lpd.1 | A0A0K2GHL2 | Dihydrolipoyl dehydrogenase | 49.863 | 471 | 0.645 | 1.56 | 7.15E-04 | 2.22E-03 | no | no | 0 | 13 | 10 | 11 | 8 | 11 | 9 | 0.007684266 | 0.006712241 | 0.007212066 | 0.004304675 | 0.004653741 | 0.004831912 |
| NITMOv2_0496 | A0A0K2G8M3 | DUF4149 domain-containing protein | 18.999 | 169 | 1.54 | 2.91 | 6.24E-02 | 8.51E-02 | no | yes | 2 | 3 | 2 | 1 | 2 | 2 | 1 | 0.007672251 | 0.007525703 | 0.008162248 | 0.004932215 | 0.005415045 | 0.00619219 |
| NITMOv2_3607 | A0A0K2GGN8 | Uncharacterized protein | 20.83 | 189 | 0.221 | 1.17 | 8.07E-02 | 1.07E-01 | no | no | 0 | 11 | 13 | 11 | 8 | 8 | 8 | 0.007659393 | 0.007379436 | 0.008066962 | 0.007436896 | 0.006941381 | 0.006772624 |
| ligA | A0A0K2GCN1 | DNA ligase | 78.759 | 707 | 0.317 | 1.25 | 1.90E-03 | 4.62E-03 | no | no | 0 | 18 | 23 | 20 | 15 | 14 | 14 | 0.007658128 | 0.008347183 | 0.007345314 | 0.006870511 | 0.005834694 | 0.005718745 |
| metH | A0A0K2GCU2 | Methionine synthase | 135.29 | 1232 | -0.0582 | -1.04 | 4.62E-01 | 5.05E-01 | no | no | 0 | 20 | 26 | 35 | 30 | 29 | 27 | 0.007657285 | 0.008094932 | 0.008132148 | 0.008831018 | 0.008581758 | 0.006850842 |
| nagZ | A0A0K2G6R4 | Beta-N-acetylglucosaminidase | 40.05 | 367 | -0.291 | -1.22 | 5.00E-03 | 9.93E-03 | no | no | 0 | 18 | 16 | 13 | 20 | 21 | 19 | 0.007598686 | 0.0088906 | 0.008345536 | 0.011799469 | 0.011170657 | 0.01054778 |
| rmlB | A0A0K2GGY4 | dTDP-glucose 4,6-dehydratase | 37.954 | 341 | -0.61 | -1.53 | 1.90E-04 | 8.40E-04 | no | no | 0 | 8 | 9 | 10 | 26 | 23 | 18 | 0.00757297 | 0.007643421 | 0.009141764 | 0.015004394 | 0.012917305 | 0.013053282 |
| kdsD | A0A0K2GFH1 | D-arabinose 5-phosphate isomerase | 36.112 | 342 | -0.416 | -1.33 | 1.76E-03 | 4.35E-03 | no | no | 0 | 12 | 11 | 14 | 19 | 17 | 18 | 0.007560744 | 0.006412081 | 0.006190179 | 0.015263525 | 0.015948537 | 0.014019772 |
| NITMOv2_4623 | A0A0K2GJG6 | Uncharacterized protein | 34.945 | 332 | 0.0691 | 1.05 | 4.94E-01 | 5.37E-01 | no | no | 0 | 5 | 5 | 4 | 5 | 5 | 5 | 0.007511631 | 0.007207749 | 0.00797666 | 0.008561522 | 0.008264185 | 0.007370859 |

| | | | | | | | | | | | | | | | | | | | | | | | |
|---|---|---|---|---|---|---|---|---|---|---|---|---|---|---|---|---|---|---|---|---|---|---|---|
| ispH | A0A0K2G964 | 4-hydroxy-3-methylbut-2-enyl diphosphate reductase | 34.435 | 313 | -1.23 | -2.35 | 3.02E-05 | 2.47E-04 | yes | no | 0 | 14 | 9 | 10 | 27 | 27 | 27 | 0.007511209 | 0.00751182 | 0.008034177 | 0.023115918 | 0.021826113 | 0.019673246 |
| NITMOv2_2457 | A0A0K2GD37 | Putative RNA polymerase sigma factor, sigma-24-like protein | 22.146 | 199 | 0.143 | 1.10 | 7.08E-02 | 9.54E-02 | no | no | 0 | 5 | 5 | 11 | 8 | 8 | 9 | 0.007501302 | 0.008108815 | 0.009460409 | 0.007600222 | 0.008015752 | 0.007122179 |
| NITMOv2_1843 | A0A0K2GBD9 | Uncharacterized protein | 26.411 | 244 | -0.407 | -1.33 | 2.79E-03 | 6.24E-03 | no | no | 0 | 7 | 7 | 7 | 9 | 7 | 7 | 0.007491395 | 0.007558555 | 0.00607572 | 0.009330622 | 0.008973441 | 0.007816793 |
| NITMOv2_0517 | A0A0K2G7M7 | Transcriptional regulator (Modular protein) | 36.128 | 324 | 1.36 | 2.57 | 5.25E-04 | 1.77E-03 | yes | no | 0 | 7 | 9 | 10 | 3 | 2 | 1 | 0.00748739 | 0.00961509 | 0.009006407 | 0.002838737 | 0.001829926 | 0.001781327 |
| degT | A0A0K2GGR5 | Pleiotropic regulatory protein | 40.728 | 372 | -0.504 | -1.42 | 2.36E-04 | 9.85E-04 | no | no | 0 | 9 | 9 | 12 | 11 | 11 | 13 | 0.007482753 | 0.008029815 | 0.009133712 | 0.011028296 | 0.01166799 | 0.010935994 |
| lpxB | A0A0K2GG48 | Lipid-A-disaccharide synthase | 41.119 | 376 | -0.164 | -1.12 | 1.60E-01 | 1.96E-01 | no | no | 0 | 10 | 9 | 8 | 12 | 13 | 12 | 0.007482753 | 0.008258211 | 0.008367968 | 0.009895378 | 0.009205561 | 0.008381403 |
| NITMOv2_0339 | A0A0K2G844 | Iron-sulfur cluster carrier protein | 33.49 | 308 | -1.18 | -2.27 | 7.87E-07 | 2.72E-05 | yes | no | 0 | 7 | 7 | 8 | 20 | 17 | 16 | 0.007478748 | 0.007324097 | 0.007664725 | 0.017465393 | 0.01803047 | 0.016814952 |
| hisS | A0A0K2GJW2 | Histidine--tRNA ligase | 46.829 | 427 | -0.14 | -1.10 | 1.62E-01 | 1.98E-01 | no | no | 0 | 15 | 14 | 24 | 24 | 25 | 19 | 0.007439963 | 0.007364771 | 0.008633121 | 0.009778991 | 0.009736764 | 0.009596214 |
| NITMOv2_4524 | A0A0K2GIW5 | Uncharacterized protein | 60.613 | 542 | -0.143 | -1.10 | 1.48E-01 | 1.82E-01 | no | no | 0 | 17 | 23 | 19 | 19 | 24 | 22 | 0.007437223 | 0.007652807 | 0.006826126 | 0.0102013 | 0.010202092 | 0.008657774 |
| NITMOv2_2524 | A0A0K2GEC6 | RNA 2,3-cyclic phosphodiesterase | 22.852 | 204 | 0.419 | 1.34 | 4.38E-03 | 8.91E-03 | no | no | 0 | 7 | 7 | 8 | 7 | 6 | 7 | 0.007426894 | 0.007667081 | 0.007634817 | 0.006398597 | 0.005188829 | 0.005493261 |
| purN | A0A0K2G999 | Phosphoribosylglycinamide formyltransferase | 50.964 | 459 | -0.175 | -1.13 | 1.22E-01 | 1.54E-01 | no | no | 0 | 15 | 13 | 15 | 21 | 19 | 21 | 0.007426473 | 0.007869665 | 0.007761162 | 0.009648982 | 0.009921808 | 0.008898183 |
| hemB | A0A0K2GB49 | Delta-aminolevulinic acid dehydratase | 35.707 | 323 | -0.645 | -1.56 | 2.70E-04 | 1.08E-03 | no | no | 0 | 35 | 41 | 37 | 31 | 35 | 32 | 0.007423522 | 0.00744729 | 0.008362216 | 0.013113926 | 0.012801867 | 0.010727952 |
| NITMOv2_1134 | A0A0K2G9E9 | Farnesyl-diphosphate farnesyltransferase | 39.669 | 352 | 0.142 | 1.10 | 9.59E-02 | 1.25E-01 | no | no | 0 | 6 | 6 | 7 | 7 | 7 | 6 | 0.007405183 | 0.007702475 | 0.007461307 | 0.00746044 | 0.00757171 | 0.007731023 |
| NITMOv2_3294 | A0A0K2GGD1 | mRNA interferase | 12.316 | 112 | 0.224 | 1.17 | 1.12E-01 | 1.43E-01 | no | no | 0 | 3 | 5 | 6 | 4 | 5 | 4 | 0.007385369 | 0.010181972 | 0.009815097 | 0.009039063 | 0.008888455 | 0.009447869 |
| NITMOv2_2240 | A0A0K2GCG9 | Putative Oxidoreductase, GFO/IDH/MOCA family | 36.018 | 336 | 0.153 | 1.11 | 3.06E-01 | 3.47E-01 | no | no | 0 | 11 | 9 | 13 | 13 | 9 | 13 | 0.007378835 | 0.007261328 | 0.008348795 | 0.005230733 | 0.00483257 | 0.004625847 |
| NITMOv2_2395 | A0A0K2GCX7 | Uncharacterized protein | 14.575 | 125 | -2.03 | -4.08 | 1.41E-03 | 3.68E-03 | yes | yes | 3 | 3 | 2 | 3 | 3 | 2 | 2 | 0.007367452 | 0.006979158 | 0.006352569 | 0.006121402 | 0.006457099 | 0.006938231 |
| NITMOv2_4145 | A0A0K2GIT5 | Uncharacterized protein | 52.298 | 458 | 0.258 | 1.20 | 4.28E-02 | 6.13E-02 | no | no | 0 | 16 | 19 | 17 | 12 | 13 | 11 | 0.007367241 | 0.008001071 | 0.007574424 | 0.005818738 | 0.005811389 | 0.006455615 |
| NITMOv2_3826 | A0A0K2GGX9 | Uncharacterized protein | 50.332 | 458 | 0.182 | 1.13 | 1.59E-01 | 1.95E-01 | no | no | 0 | 21 | 20 | 24 | 15 | 13 | 13 | 0.007355016 | 0.006934183 | 0.00757845 | 0.002426942 | 0.002497076 | 0.00251719 |
| lolD | A0A0K2GCM8 | Lipoprotein-releasing system ATP-binding protein LolD | 24.14 | 220 | -0.15 | -1.11 | 5.26E-02 | 7.36E-02 | no | no | 0 | 10 | 9 | 10 | 9 | 12 | 10 | 0.007326981 | 0.007169031 | 0.007123681 | 0.007154074 | 0.009083131 | 0.006400773 |
| murF | A0A0K2G873 | UDP-N-acetylmuramoyl-tripeptide--D-alanyl-D-alanine ligase | 51.42 | 481 | -0.279 | -1.21 | 9.68E-03 | 1.73E-02 | no | no | 0 | 13 | 9 | 9 | 13 | 13 | 15 | 0.007319814 | 0.007460783 | 0.007273801 | 0.009370009 | 0.011364867 | 0.010388646 |
| NITMOv2_3740 | A0A0K2GGR4 | Putative Nucleotidyl transferase | 26.882 | 246 | 0.0991 | 1.07 | 2.31E-01 | 2.73E-01 | no | no | 0 | 2 | 4 | 4 | 3 | 2 | 1 | 0.007304216 | 0.006874738 | 0.007585544 | 0.007212119 | 0.007629662 | 0.007746487 |
| NITMOv2_0421 | A0A0K2G8D1 | Glutamine amidotransferase | 24.262 | 227 | 0.557 | 1.47 | 5.88E-04 | 1.93E-03 | no | no | 0 | 3 | 4 | 4 | 3 | 3 | 5 | 0.007286088 | 0.007014747 | 0.0072692 | 0.005093616 | 0.004383401 | 0.004355769 |
| NITMOv2_2001 | A0A0K2GBU9 | Putative Phosphoglycolate phosphatase | 25.464 | 231 | 0.318 | 1.25 | 3.61E-03 | 7.67E-03 | no | no | 0 | 6 | 7 | 7 | 4 | 2 | 4 | 0.007261004 | 0.007862821 | 0.007134226 | 0.006125696 | 0.006001249 | 0.006226894 |
| lepA | A0A0K2GGF0 | Elongation factor 4 | 66.486 | 599 | -1.14 | -2.20 | 2.60E-06 | 4.99E-05 | yes | no | 0 | 18 | 17 | 18 | 38 | 43 | 39 | 0.00722243 | 0.007401337 | 0.00829147 | 0.01785631 | 0.019389941 | 0.018263518 |
| NITMOv2_3417 | A0A0K2GG24 | PilZ domain-containing protein | 12.679 | 114 | -2.41 | -5.31 | 2.76E-07 | 1.39E-05 | yes | no | 0 | 2 | 2 | 3 | 9 | 7 | 9 | 0.007206621 | 0.006790067 | 0.006950747 | 0.043064521 | 0.039360958 | 0.037147389 |
| thi4 | A0A0K2G8T3 | Thiamine thiazole synthase | 28.855 | 266 | -0.452 | -1.37 | 1.08E-02 | 1.90E-02 | no | no | 0 | 3 | 3 | 3 | 9 | 8 | 10 | 0.007177743 | 0.006131084 | 0.00812352 | 0.013275476 | 0.014374192 | 0.011788843 |
| NITMOv2_3322 | A0A0K2GFH7 | Putative Peptidase M48, Ste24p | 34.233 | 316 | -0.204 | -1.15 | 1.58E-02 | 2.61E-02 | no | no | 0 | 11 | 10 | 12 | 11 | 9 | 9 | 0.007152238 | 0.007000081 | 0.007609892 | 0.011524642 | 0.011691917 | 0.010064625 |
| parB | A0A0K2G720 | Chromosomal partitioning protein ParB | 30.959 | 282 | -0.895 | -1.86 | 3.78E-04 | 1.37E-03 | no | no | 0 | 7 | 10 | 7 | 20 | 22 | 20 | 0.007106708 | 0.008625247 | 0.008105881 | 0.016868653 | 0.015628479 | 0.014632392 |
| NITMOv2_0476 | A0A0K2G8K6 | Saccharop_dh_N domain-containing protein | 44.063 | 414 | 1.04 | 2.06 | 2.44E-04 | 1.01E-03 | yes | no | 0 | 7 | 10 | 9 | 7 | 8 | 6 | 0.007105654 | 0.008032944 | 0.006096043 | 0.003357887 | 0.003573156 | 0.003184113 |
| merP | A0A0K2GAJ4 | Mercuric transport protein periplasmic component | 12.617 | 117 | 0.767 | 1.70 | 1.85E-05 | 1.77E-04 | no | no | 0 | 5 | 4 | 4 | 1 | 0 | 0 | 0.007086894 | 0.008136387 | 0.007974934 | 0.004861732 | 0.005460879 | 0.005075917 |
| lpd.2 | A0A0K2GJQ9 | Dihydrolipoyl dehydrogenase | 48.774 | 455 | 0.526 | 1.44 | 2.27E-04 | 9.65E-04 | no | no | 0 | 9 | 11 | 13 | 9 | 11 | 3 | 0.007031878 | 0.008144404 | 0.007879456 | 0.004532858 | 0.005763536 | 0.005556734 |
| trpB | A0A0K2GDL4 | Tryptophan synthase beta chain | 41.258 | 379 | -0.241 | -1.18 | 1.02E-01 | 1.31E-01 | no | no | 0 | 10 | 10 | 11 | 14 | 14 | 13 | 0.007009745 | 0.005293569 | 0.00604121 | 0.008447801 | 0.008600091 | 0.007468317 |
| NITMOv2_1028 | A0A0K2G9D0 | Uncharacterized protein | 98.148 | 885 | 0.481 | 1.40 | 1.86E-04 | 8.26E-04 | no | no | 0 | 24 | 25 | 27 | 17 | 16 | 15 | 0.006993304 | 0.006815683 | 0.006334739 | 0.005246577 | 0.004882754 | 0.004843959 |
| NITMOv2_2092 | A0A0K2GCD0 | Uncharacterized protein | 13.54 | 128 | -0.544 | -1.46 | 8.65E-03 | 1.58E-02 | no | no | 0 | 6 | 5 | 4 | 4 | 5 | 1 | 0.006969906 | 0.009666909 | 0.00657401 | 0.004087598 | 0.014956201 | 0.0026605 |
| hslV | A0A0K2G6S8 | ATP-dependent protease subunit HslV | 19.24 | 178 | -0.868 | -1.83 | 6.61E-03 | 1.26E-02 | no | no | 0 | 4 | 7 | 10 | 6 | 5 | 3 | 0.006949038 | 0.009115084 | 0.01463752 | 0.008473566 | 0.00824927 | 0.006163421 |
| uvrA | A0A0K2G7P1 | Excinuclease ABC, subunit A | 91.869 | 836 | 0.136 | 1.10 | 1.77E-01 | 2.14E-01 | no | no | 0 | 20 | 21 | 24 | 20 | 19 | 15 | 0.006941239 | 0.007466845 | 0.00828591 | 0.008238868 | 0.008653072 | 0.007504459 |
| NITMOv2_1136 | A0A0K2GAD6 | TFIIS N-terminal domain-containing protein | 15.301 | 137 | 0.805 | 1.75 | 2.20E-01 | 2.61E-01 | no | no | 0 | 2 | 1 | 1 | 2 | 2 | 2 | 0.006929014 | 0.007457654 | 0.006722404 | 0.007161626 | 0.001195356 | 0.00682387 |
| speE | A0A0K2GJR7 | Polyamine aminopropyltransferase | 56.31 | 516 | -0.24 | -1.18 | 7.31E-02 | 9.82E-02 | no | no | 0 | 14 | 14 | 14 | 11 | 13 | 13 | 0.006908356 | 0.006121893 | 0.005599862 | 0.005939566 | 0.007402514 | 0.00568548 |
| NITMOv2_4539 | A0A0K2GIZ2 | Sigma54 dependent transcriptional regulator | 53.286 | 472 | -1.1 | -2.14 | 1.28E-04 | 6.56E-04 | yes | no | 0 | 10 | 10 | 15 | 19 | 26 | 21 | 0.0069014 | 0.007202078 | 0.00795442 | 0.015513771 | 0.01636337 | 0.015197901 |
| yabD | A0A0K2GG65 | Putative deoxyribonuclease YabD | 29.117 | 258 | 0.674 | 1.60 | 2.83E-04 | 1.11E-03 | no | no | 0 | 6 | 6 | 4 | 4 | 4 | 2 | 0.006900979 | 0.006804342 | 0.005794462 | 0.002885381 | 0.002981359 | 0.003047456 |
| NITMOv2_4472 | A0A0K2GIT3 | t(6)A37 threonylcarbamoyladenosine biosynthesis protein TsaE | 20.965 | 196 | -0.115 | -1.08 | 3.56E-01 | 3.98E-01 | no | no | 0 | 4 | 6 | 5 | 3 | 4 | 4 | 0.006869571 | 0.006897812 | 0.004066457 | 0.006773966 | 0.005327728 | 0.005586943 |
| vapC | A0A0K2G8M8 | Ribonuclease VapC | 14.047 | 124 | 1.04 | 2.06 | 1.52E-04 | 7.40E-04 | yes | no | 0 | 4 | 4 | 3 | 1 | 1 | 1 | 0.006851444 | 0.006533513 | 0.0059233 | 0.003943669 | 0.003650529 | 0.003776414 |
| argD | A0A0K2G7C3 | Acetylornithine aminotransferase | 44.172 | 405 | -0.728 | -1.66 | 9.42E-05 | 5.41E-04 | no | no | 0 | 6 | 6 | 10 | 22 | 22 | 24 | 0.006829522 | 0.006776183 | 0.007533778 | 0.015957994 | 0.015808706 | 0.013588403 |
| NITMOv2_3122 | A0A0K2GEZ1 | Uncharacterized protein | 30.204 | 262 | 0.587 | 1.50 | 3.57E-03 | 7.62E-03 | no | no | 0 | 7 | 9 | 5 | 6 | 5 | 3 | 0.006822144 | 0.006713609 | 0.007379249 | 0.003262527 | 0.00370382 | 0.003248126 |
| NITMOv2_0173 | A0A0K2G6N9 | Putative Histidine kinase | 105.15 | 956 | -1.67 | -3.18 | 1.36E-07 | 1.05E-05 | yes | no | 0 | 25 | 23 | 24 | 75 | 75 | 77 | 0.006786521 | 0.008542727 | 0.007707288 | 0.0297541 | 0.028213296 | 0.029043256 |
| dfrA | A0A0K2G7R6 | Putative dihydroflavonol-4-reductase | 35.618 | 327 | -0.256 | -1.19 | 6.36E-02 | 8.65E-02 | no | no | 0 | 7 | 7 | 8 | 11 | 10 | 9 | 0.006765021 | 0.007899192 | 0.005887448 | 0.009185657 | 0.008929938 | 0.007751701 |
| NITMOv2_4820 | A0A0K2GJQ6 | Putative Inositol-phosphate phosphatase | 28.679 | 262 | 0.188 | 1.14 | 8.76E-02 | 1.15E-01 | no | no | 0 | 7 | 8 | 8 | 6 | 6 | 7 | 0.006761227 | 0.007078299 | 0.006753847 | 0.00550941 | 0.004504588 | 0.004898622 |
| ispE | A0A0K2G7X7 | 4-diphosphocytidyl-2-C-methyl-D-erythritol kinase | 32.074 | 303 | 0.0376 | 1.03 | 9.27E-01 | 9.40E-01 | no | no | 0 | 5 | 6 | 11 | 15 | 13 | 11 | 0.006758276 | 0.006539575 | 0.005964521 | 0.007004815 | 0.007361031 | 0.006190752 |
| NITMOv2_1933 | A0A0K2GCN0 | Uncharacterized protein | 9.9453 | 89 | 0.395 | 1.31 | 3.87E-02 | 5.62E-02 | no | no | 0 | 2 | 1 | 1 | 1 | 2 | 2 | 0.006754271 | 0.006957257 | 0.006255365 | 0.004262474 | 0.004958263 | 0.004952026 |
| NITMOv2_2466 | A0A0K2GDD7 | 2-dehydropantoate 2-reductase | 49.028 | 450 | -0.179 | -1.13 | 2.30E-02 | 3.58E-02 | no | no | 0 | 21 | 23 | 18 | 23 | 20 | 18 | 0.006738883 | 0.006674696 | 0.007307544 | 0.009933581 | 0.010751784 | 0.007256319 |
| NITMOv2_2233 | A0A0K2GCQ9 | Uncharacterized protein | 30.594 | 276 | 2.78 | 6.87 | 3.12E-04 | 1.19E-03 | yes | yes | 3 | 2 | 2 | 2 | 0 | 2 | 1 | 0.006731295 | 0.006201089 | 0.00600785 | 0.002933801 | 0.002837954 | 0.003183034 |
| NITMOv2_3590 | A0A0K2GH96 | Uncharacterized protein | 50.828 | 455 | -0.117 | -1.08 | 1.75E-01 | 2.12E-01 | no | no | 0 | 8 | 9 | 8 | 13 | 11 | 10 | 0.006697569 | 0.004189528 | 0.004232682 | 0.004732463 | 0.004932316 | 0.003820289 |
| NITMOv2_3876 | A0A0K2GH20 | Malto-oligosyltrehalose synthase | 108.94 | 964 | -0.255 | -1.19 | 8.55E-03 | 1.56E-02 | no | no | 0 | 24 | 26 | 32 | 28 | 34 | 27 | 0.006664897 | 0.006477001 | 0.006221814 | 0.004682561 | 0.003949147 | 0.005766395 |
| glgA | A0A0K2GFS6 | Glycogen synthase | 54.097 | 493 | 0.0512 | 1.04 | 5.04E-01 | 5.46E-01 | no | no | 0 | 19 | 20 | 19 | 16 | 15 | 16 | 0.006661102 | 0.007052291 | 0.006743877 | 0.006991192 | 0.006119329 | 0.006351684 |
| nuoB.1 | A0A0K2G7Q1 | NADH-quinone oxidoreductase subunit B | 17.79 | 159 | -0.257 | -1.19 | 1.31E-02 | 2.23E-02 | no | no | 0 | 3 | 2 | 3 | 3 | 3 | 3 | 0.006615572 | 0.006948067 | 0.006363689 | 0.007259947 | 0.007918025 | 0.009445172 |
| NITMOv2_3500 | A0A0K2GH06 | Putative SAM-dependent methyltransferase | 33.751 | 300 | -0.383 | -1.30 | 1.79E-03 | 4.40E-03 | no | no | 0 | 6 | 5 | 5 | 7 | 7 | 7 | 0.006610092 | 0.005337762 | 0.006168515 | 0.009992515 | 0.010112911 | 0.009881216 |
| NITMOv2_1688 | A0A0K2GAY3 | Rhodanese-like protein | 13.796 | 121 | 0.0903 | 1.06 | 4.55E-01 | 4.99E-01 | no | no | 0 | 1 | 1 | 2 | 0 | 0 | 0 | 0.006586273 | 0.005633034 | 0.005948799 | 0.006756938 | 0.005782801 | 0.005598271 |
| NITMOv2_4437 | A0A0K2GIP5 | acidPPc domain-containing protein | 40.415 | 376 | 0.274 | 1.21 | 1.44E-01 | 1.78E-01 | no | no | 0 | 4 | 4 | 3 | 4 | 5 | 3 | 0.006576155 | 0.007440251 | 0.006815198 | 0.008019273 | 0.005726713 | 0.005304098 |
| NITMOv2_1001 | A0A0K2G907 | Dipeptide epimerase | 39.707 | 368 | 0.489 | 1.40 | 4.80E-04 | 1.65E-03 | no | no | 0 | 11 | 11 | 9 | 10 | 8 | 8 | 0.006527674 | 0.007059331 | 0.006361772 | 0.00540176 | 0.005339847 | 0.005180747 |
| NITMOv2_4429 | A0A0K2GIV9 | Uncharacterized protein | 20.587 | 183 | 0.188 | 1.14 | 3.41E-01 | 3.84E-01 | no | no | 0 | 1 | 2 | 1 | 4 | 2 | 1 | 0.006493737 | 0.006639107 | 0.005905853 | 0.006003238 | 0.005951997 | 0.004848814 |
| thiF | A0A0K2GCY6 | Adenylyltransferase ThiF | 29.368 | 273 | -1.15 | -2.22 | 1.76E-05 | 1.71E-04 | yes | no | 0 | 9 | 7 | 10 | 16 | 15 | 16 | 0.006481511 | 0.005908946 | 0.006787398 | 0.018028076 | 0.018690784 | 0.017205324 |
| Uxs | A0A0K2GFL2 | UDP-glucuronic acid decarboxylase 1 | 37.449 | 339 | -0.691 | -1.61 | 3.77E-05 | 2.87E-04 | no | no | 0 | 12 | 16 | 15 | 12 | 13 | 11 | 0.006452212 | 0.005742929 | 0.007440217 | 0.012532586 | 0.012422924 | 0.010791245 |
| plsX | A0A0K2G901 | Phosphate acyltransferase | 36.054 | 339 | -1.03 | -2.04 | 6.16E-06 | 8.54E-05 | yes | no | 0 | 7 | 6 | 6 | 15 | 15 | 14 | 0.006437878 | 0.007533525 | 0.008107607 | 0.014057309 | 0.013910573 | 0.013252154 |
| NITMOv2_2443 | A0A0K2GD21 | Uncharacterized protein | 15.503 | 140 | 0.52 | 1.43 | 5.77E-03 | 1.11E-02 | no | no | 0 | 4 | 3 | 2 | 3 | 2 | 3 | 0.006423966 | 0.006575946 | 0.005434405 | 0.00424589 | 0.004132636 | 0.003749622 |
| NITMOv2_2679 | A0A0K2GDQ9 | Response regulator receiver protein | 17.596 | 157 | 0.108 | 1.08 | 2.00E-01 | 2.39E-01 | no | no | 0 | 4 | 5 | 4 | 5 | 5 | 5 | 0.00641954 | 0.006880995 | 0.006600276 | 0.00610926 | 0.007537529 | 0.007352159 |
| NITMOv2_0525 | A0A0K2G7M0 | Putative N-acetyltransferase | 17.608 | 159 | -0.106 | -1.08 | 6.27E-01 | 6.68E-01 | no | no | 0 | 3 | 5 | 7 | 6 | 3 | 5 | 0.00640078 | 0.008168652 | 0.007691183 | 0.004983745 | 0.005992859 | 0.005558532 |
| hypE | A0A088NNR9 | Hydrogenase maturation protein | 38.832 | 366 | -0.217 | -1.16 | 6.03E-02 | 8.27E-02 | no | no | 0 | 4 | 3 | 5 | 0 | 0 | 1 | 0.00639298 | 0.011419178 | 0.012584353 | 0.005609064 | 0.004907923 | 0.005076097 |
| deoC | A0A0K2GE71 | Deoxyribose-phosphate aldolase | 24.015 | 224 | 0.328 | 1.26 | 9.99E-03 | 1.78E-02 | no | no | 0 | 9 | 8 | 6 | 5 | 5 | 5 | 0.006388765 | 0.006505355 | 0.006458209 | 0.006981715 | 0.00563862 | 0.005447409 |
| maeA | A0A0K2GC35 | Malate dehydrogenase (Oxaloacetate-decarboxylating) | 51.028 | 478 | 0.56 | 1.47 | 2.50E-03 | 5.72E-03 | no | no | 0 | 18 | 17 | 17 | 9 | 9 | 7 | 0.006369794 | 0.007756054 | 0.007952694 | 0.003006061 | 0.004483302 | 0.003788821 |
| NITMOv2_2441 | A0A0K2GDA6 | Uncharacterized protein | 31.301 | 284 | -0.612 | -1.53 | 4.66E-03 | 9.37E-03 | no | no | 0 | 5 | 7 | 7 | 13 | 12 | 8 | 0.006357779 | 0.007399382 | 0.007856449 | 0.010753026 | 0.011650278 | 0.009750852 |
| smpB | A0A0K2G668 | SsrA-binding protein | 18.13 | 158 | 0.962 | 1.95 | 3.65E-02 | 5.36E-02 | no | yes | 1 | 4 | 4 | 2 | 1 | 2 | 1 | 0.006352088 | 0.00625545 | 0.004696077 | 0.002202313 | 0.003108295 | 0.004908691 |
| NITMOv2_4309 | A0A0K2GIB2 | Uncharacterized protein | 15.904 | 141 | -1.07 | -2.10 | 1.81E-02 | 2.92E-02 | yes | no | 0 | 4 | 4 | 4 | 4 | 3 | 3 | 0.006307401 | 0.004096645 | 0.004529853 | 0.002365491 | 0.002290747 | 0.00219209 |
| NITMOv2_1312 | A0A0K2G9X1 | Plasmid stabilization system | 12.396 | 105 | 2.21 | 4.63 | 6.22E-02 | 8.49E-02 | no | yes | 2 | 2 | 2 | 3 | 2 | 2 | 1 | 0.006290959 | 0.006708916 | 0.005489621 | 0.001828573 | 0.005342488 | 0.002046442 |
| NITMOv2_4013 | A0A0K2GIF6 | DUF218 domain-containing protein | 27.461 | 253 | -1.09 | -2.13 | 5.67E-03 | 1.10E-02 | yes | no | 0 | 6 | 11 | 14 | 9 | 12 | 11 | 0.006290538 | 0.008777184 | 0.008754099 | 0.007526777 | 0.00769647 | 0.007520822 |
| ydaO | A0A0K2GGT7 | Putative amino acid permease YdaO | 65.709 | 604 | 0.455 | 1.37 | 6.02E-03 | 1.16E-02 | no | no | 0 | 6 | 6 | 6 | 8 | 6 | 7 | 0.006276204 | 0.006302185 | 0.006165639 | 0.005592332 | 0.005753903 | 0.004953644 |
| engB | A0A0K2GJ84 | Probable GTP-binding protein EngB | 22.169 | 200 | 0.185 | 1.14 | 5.56E-02 | 7.69E-02 | no | no | 0 | 11 | 10 | 8 | 7 | 7 | 6 | 0.006256179 | 0.006178601 | 0.006595483 | 0.004403885 | 0.006278581 | 0.006075492 |
| NITMOv2_1174 | A0A0K2G9I9 | ABC-type transport system, permease component | 28.145 | 255 | -1.29 | -2.45 | 3.04E-03 | 6.70E-03 | yes | no | 0 | 2 | 2 | 2 | 5 | 4 | 3 | 0.006245851 | 0.007210878 | 0.007358735 | 0.01682275 | 0.015699948 | 0.013544529 |
| NITMOv2_0869 | A0A0K2G8W5 | Cell shape-determining protein MreC | 31.644 | 292 | 0.0704 | 1.05 | 3.56E-01 | 3.98E-01 | no | no | 0 | 10 | 11 | 10 | 9 | 10 | 8 | 0.006231517 | 0.006783419 | 0.007265749 | 0.00811108 | 0.007905751 | 0.006257462 |
| zraR.1 | A0A0K2GGP2 | Transcriptional regulatory protein ZraR | 50.879 | 461 | -0.249 | -1.19 | 5.54E-03 | 1.08E-02 | no | no | 0 | 24 | 21 | 23 | 26 | 26 | 23 | 0.006230885 | 0.006006914 | 0.00666297 | 0.008645481 | 0.008906943 | 0.008008113 |
| moeB | A0A0K2GDY5 | Molybdopterin synthase sulfurylase | 28.67 | 267 | -0.755 | -1.69 | 5.62E-04 | 1.85E-03 | no | no | 0 | 3 | 4 | 5 | 13 | 9 | 5 | 0.00621613 | 0.006047978 | 0.006617723 | 0.012876267 | 0.01224394 | 0.011244732 |

Table S1

| | | | | | | | | | | | | | | | | | | | | | | | |
|---|---|---|---|---|---|---|---|---|---|---|---|---|---|---|---|---|---|---|---|---|---|---|---|
| NITMOv2_0503 | A0A0K2G7W3 | Uncharacterized protein | 62.899 | 580 | 0.371 | 1.29 | 1.07E-02 | 1.89E-02 | no | no | 0 | 12 | 11 | 16 | 9 | 9 | 6 | 0.006202218 | 0.006096082 | 0.006110422 | 0.006016121 | 0.005535766 | 0.004598695 |
| NITMOv2_1708 | A0A0K2GB04 | Uncharacterized protein | 13.265 | 119 | 1.41 | 2.66 | 3.04E-02 | 4.58E-02 | yes | yes | 1 | 2 | 2 | 2 | 1 | 2 | 2 | 0.006199899 | 0.005443552 | 0.006842231 | 0.002540219 | 0.003631885 | 0.003881784 |
| NITMOv2_2030 | A0A0K2GCY5 | Uncharacterized protein | 12.715 | 109 | 0.845 | 1.80 | 6.45E-05 | 4.18E-04 | no | no | 0 | 4 | 4 | 4 | 2 | 2 | 1 | 0.006180928 | 0.006252126 | 0.006603919 | 0.002801126 | 0.002890158 | 0.00335925 |
| NITMOv2_1146 | A0A0K2GAE6 | Putative Cytochrome c | 13.644 | 130 | 0.435 | 1.35 | 5.58E-02 | 7.71E-02 | no | no | 0 | 2 | 3 | 2 | 1 | 1 | 1 | 0.006169756 | 0.006867698 | 0.005511861 | 0.004777477 | 0.004897358 | 0.004444237 |
| NITMOv2_0011 | A0A0K2G6A6 | Putative Quercetin 2,3-dioxygenase | 32.309 | 294 | 0.918 | 1.89 | 1.66E-04 | 7.81E-04 | no | no | 0 | 7 | 8 | 8 | 2 | 3 | 2 | 0.006167438 | 0.008170998 | 0.009140805 | 0.003852307 | 0.00378166 | 0.003419308 |
| murD | A0A0K2G7V7 | UDP-N-acetylmuramoylalanine--D-glutamate ligase | 50.555 | 468 | -0.482 | -1.40 | 6.15E-04 | 1.99E-03 | no | no | 0 | 11 | 10 | 10 | 17 | 14 | 13 | 0.006133079 | 0.006103317 | 0.005143177 | 0.009596564 | 0.009683162 | 0.007571709 |
| NITMOv2_1120 | A0A0K2G9A4 | Uncharacterized protein | 22.241 | 204 | 0.754 | 1.69 | 6.69E-05 | 4.28E-04 | no | no | 0 | 3 | 3 | 5 | 3 | 2 | 2 | 0.006107785 | 0.006698944 | 0.005920616 | 0.003416376 | 0.003555133 | 0.003402945 |
| NITMOv2_0123 | A0A0K2G6J5 | Putative Histidine kinase with N-terminal NAD-binding region | 41.097 | 369 | 0.359 | 1.28 | 7.45E-03 | 1.39E-02 | no | no | 0 | 4 | 7 | 7 | 7 | 6 | 6 | 0.006101461 | 0.00741483 | 0.006005166 | 0.004722245 | 0.002025068 | 0.002480149 |
| NITMOv2_1124 | A0A0K2G9E1 | Response regulator | 23.629 | 212 | 0.0688 | 1.05 | 4.79E-01 | 5.22E-01 | no | no | 0 | 5 | 6 | 5 | 9 | 7 | 7 | 0.00608249 | 0.006208128 | 0.005575322 | 0.00761888 | 0.007112752 | 0.005741581 |
| galE | A0A0K2GF41 | UDP-glucose 4-epimerase | 35.977 | 328 | 0.0226 | 1.02 | 7.85E-01 | 8.11E-01 | no | no | 0 | 7 | 7 | 10 | 9 | 10 | 6 | 0.006065627 | 0.006053062 | 0.006161421 | 0.006362911 | 0.006760377 | 0.006093114 |
| NITMOv2_1472 | A0A0K2GAL8 | Uncharacterized protein | 37.041 | 354 | -0.524 | -1.44 | 3.19E-03 | 6.95E-03 | no | no | 0 | 11 | 18 | 15 | 18 | 17 | 21 | 0.006020729 | 0.006747047 | 0.006284891 | 0.011177407 | 0.01029609 | 0.00971489 |
| mgtE | A0A0K2GGS2 | Magnesium transporter MgtE | 53.259 | 486 | -0.375 | -1.30 | 2.47E-02 | 3.81E-02 | no | no | 0 | 18 | 13 | 19 | 23 | 23 | 22 | 0.005998597 | 0.00505481 | 0.006688469 | 0.005440555 | 0.005560469 | 0.005311291 |
| cztS | A0A0K2G7R8 | Histidine kinase | 51.838 | 467 | 0.00278 | 1.00 | 9.68E-01 | 9.72E-01 | no | no | 0 | 11 | 10 | 12 | 14 | 17 | 13 | 0.005987214 | 0.005838941 | 0.005366535 | 0.006078016 | 0.006360615 | 0.006256743 |
| NITMOv2_1751 | A0A0K2GB24 | Putative ABC transporter, periplasmic substrate-binding protein, MCE domain-containing | 34.711 | 325 | 0.127 | 1.09 | 3.51E-01 | 3.93E-01 | no | no | 0 | 10 | 9 | 9 | 7 | 8 | 8 | 0.005982366 | 0.005281641 | 0.004961039 | 0.003960994 | 0.00396111 | 0.004121294 |
| NITMOv2_4005 | A0A0K2GHP8 | Transcriptional regulator, NifA subfamily, Fis Family | 88.428 | 797 | 0.442 | 1.36 | 2.27E-04 | 9.65E-04 | no | no | 0 | 19 | 21 | 20 | 14 | 9 | 13 | 0.005972037 | 0.006576924 | 0.006803695 | 0.005702203 | 0.004660111 | 0.004670261 |
| gltB | A0A0K2GHL8 | Glutamate synthase, large subunit | 166.52 | 1506 | -0.981 | -1.97 | 9.92E-06 | 1.18E-04 | no | no | 0 | 20 | 27 | 24 | 63 | 66 | 61 | 0.005967611 | 0.006558543 | 0.005682304 | 0.01390272 | 0.014837344 | 0.014019952 |
| NITMOv2_1463 | A0A0K2GAB7 | HIRAN domain family | 28.83 | 256 | 0.0855 | 1.06 | 2.35E-01 | 2.77E-01 | no | no | 0 | 17 | 21 | 18 | 11 | 14 | 9 | 0.005953699 | 0.006270702 | 0.007117163 | 0.006179447 | 0.005986955 | 0.005633155 |
| nuoI.1 | A0A0K2G8Q5 | NADH-quinone oxidoreductase subunit I | 21.27 | 182 | -0.353 | -1.28 | 9.62E-04 | 2.73E-03 | no | no | 0 | 4 | 6 | 6 | 7 | 6 | 7 | 0.005941262 | 0.006624637 | 0.006198999 | 0.009333583 | 0.009718431 | 0.010019132 |
| NITMOv2_4174 | A0A0K2GI71 | Putative Soluble lytic murein transglycosylase | 84.894 | 758 | 0.129 | 1.09 | 1.22E-01 | 1.54E-01 | no | no | 0 | 20 | 23 | 19 | 17 | 19 | 16 | 0.005939154 | 0.006979158 | 0.006742918 | 0.004744901 | 0.005356005 | 0.005269754 |
| trpD | A0A0K2G7C8 | Anthranilate phosphoribosyltransferase | 37.059 | 346 | -0.82 | -1.77 | 4.02E-05 | 3.00E-04 | no | no | 0 | 5 | 6 | 5 | 10 | 12 | 13 | 0.005937047 | 0.006267574 | 0.006667763 | 0.010701348 | 0.010255383 | 0.013554059 |
| NITMOv2_3189 | A0A0K2GF54 | Uncharacterized protein | 48.484 | 429 | 0.685 | 1.61 | 1.10E-03 | 3.01E-03 | no | no | 0 | 6 | 9 | 7 | 2 | 6 | 5 | 0.00591934 | 0.007267194 | 0.005675593 | 0.00546558 | 0.005257346 | 0.004291037 |
| accD | A0A0K2GA92 | Acetyl-coenzyme A carboxylase carboxyl transferase subunit beta | 30.865 | 278 | -0.196 | -1.15 | 1.54E-02 | 2.56E-02 | no | no | 0 | 7 | 4 | 9 | 8 | 9 | 12 | 0.005918076 | 0.007351083 | 0.007514031 | 0.010333679 | 0.00995661 | 0.01019337 |
| NITMOv2_3165 | A0A0K2GFD9 | Putative 23S rRNA methyltransferase RlmB | 28.894 | 267 | 0.415 | 1.33 | 5.31E-03 | 1.04E-02 | no | no | 0 | 7 | 7 | 8 | 4 | 5 | 3 | 0.005905218 | 0.007120536 | 0.006216254 | 0.00484189 | 0.005423746 | 0.004160853 |
| NITMOv2_2101 | A0A0K2GC44 | Uncharacterized protein | 18.546 | 173 | 0.696 | 1.62 | 1.24E-04 | 6.44E-04 | no | no | 0 | 4 | 5 | 6 | 3 | 4 | 4 | 0.005890041 | 0.006037027 | 0.005928668 | 0.003940856 | 0.004485633 | 0.002911878 |
| NITMOv2_3575 | A0A0K2GH85 | Putative Peptidase, M28 family | 51.513 | 485 | 0.477 | 1.39 | 9.92E-04 | 2.80E-03 | no | no | 0 | 11 | 15 | 12 | 10 | 11 | 8 | 0.005877604 | 0.006616228 | 0.006220088 | 0.004701811 | 0.005442856 | 0.004724923 |
| NITMOv2_3356 | A0A0K2GFL3 | Uncharacterized protein | 66.477 | 593 | 0.361 | 1.28 | 3.73E-03 | 7.87E-03 | no | no | 0 | 16 | 16 | 20 | 16 | 15 | 16 | 0.005866644 | 0.006639694 | 0.005838367 | 0.004592384 | 0.004139473 | 0.004863019 |
| relA | A0A0K2GGB9 | GTP pyrophosphokinase | 81.44 | 726 | -0.132 | -1.10 | 1.89E-01 | 2.26E-01 | no | no | 0 | 14 | 15 | 17 | 15 | 16 | 21 | 0.005828491 | 0.005933585 | 0.005867509 | 0.007375741 | 0.006850646 | 0.006732706 |
| NITMOv2_0472 | A0A0K2G7I7 | Uncharacterized protein | 23.735 | 210 | 0.312 | 1.24 | 1.28E-02 | 2.19E-02 | no | no | 0 | 5 | 5 | 4 | 5 | 2 | 2 | 0.005808888 | 0.006759367 | 0.007012098 | 0.005061928 | 0.005846347 | 0.005273171 |
| NITMOv2_0744 | A0A0K2G8B8 | Uncharacterized protein | 70.606 | 666 | 0.0468 | 1.03 | 5.34E-01 | 5.76E-01 | no | no | 0 | 15 | 13 | 17 | 12 | 11 | 14 | 0.005806991 | 0.00569639 | 0.005949183 | 0.006351658 | 0.006363412 | 0.006829445 |
| NITMOv2_4295 | A0A0K2GJ58 | 4HB_MCP_1 domain-containing protein | 28.941 | 260 | -0.116 | -1.08 | 6.34E-01 | 6.74E-01 | no | no | 0 | 0 | 0 | 2 | 1 | 2 | 2 | 0.005797927 | 0.005843439 | 0.005741163 | 0.004219977 | 0.004207369 | 0.004529108 |
| NITMOv2_1692 | A0A0K2GBX5 | SGNH_hydro domain-containing protein | 22.855 | 211 | -0.0834 | -1.06 | 5.02E-01 | 5.45E-01 | no | no | 0 | 5 | 7 | 9 | 8 | 9 | 8 | 0.005785279 | 0.004575532 | 0.005968547 | 0.005162619 | 0.005829878 | 0.00523505 |
| secG | A0A0K2G8Z0 | Protein-export membrane protein SecG | 12.102 | 117 | 0.0205 | 1.01 | 8.62E-01 | 8.80E-01 | no | no | 0 | 3 | 2 | 2 | 3 | 1 | 2 | 0.005753872 | 0.006246455 | 0.006477956 | 0.006575695 | 0.009204007 | 0.007343348 |
| NITMOv2_0666 | A0A0K2G911 | Uncharacterized protein | 26.09 | 232 | -0.339 | -1.26 | 1.05E-02 | 1.86E-02 | no | no | 0 | 10 | 9 | 9 | 9 | 9 | 9 | 0.005742068 | 0.006001829 | 0.006053864 | 0.00941073 | 0.009304686 | 0.008422041 |
| NITMOv2_0429 | A0A0K2G7F3 | BNR repeat protein | 38.195 | 352 | 0.477 | 1.39 | 2.63E-03 | 5.96E-03 | no | no | 0 | 5 | 5 | 5 | 4 | 6 | 5 | 0.005737009 | 0.005553056 | 0.00498002 | 0.004181773 | 0.004070023 | 0.004054044 |
| NITMOv2_2517 | A0A0K2GDA5 | Uncharacterized protein | 47.672 | 426 | 0.503 | 1.42 | 1.52E-03 | 3.92E-03 | no | no | 0 | 7 | 7 | 7 | 9 | 11 | 8 | 0.005714666 | 0.003194992 | 0.004716592 | 0.003658182 | 0.003292872 | 0.003440166 |
| NITMOv2_1058 | A0A0K2G9F8 | Uncharacterized protein | 50.704 | 479 | 0.641 | 1.56 | 1.71E-04 | 7.86E-04 | no | no | 0 | 18 | 12 | 14 | 8 | 11 | 8 | 0.005711504 | 0.005809023 | 0.006335698 | 0.003890511 | 0.004128131 | 0.003853014 |
| hom | A0A0K2GJN8 | Homoserine dehydrogenase | 46.272 | 437 | -1.12 | -2.17 | 2.13E-06 | 4.38E-05 | yes | no | 0 | 9 | 11 | 11 | 27 | 26 | 25 | 0.00570771 | 0.006249193 | 0.005791011 | 0.018599644 | 0.018535416 | 0.017642627 |
| sdhC | A0A0K2GJ95 | Succinate dehydrogenase/fumarate reductase, cytochrome b558 subunit | 24.523 | 225 | 0.814 | 1.76 | 2.44E-04 | 1.01E-03 | no | no | 0 | 6 | 6 | 7 | 4 | 4 | 2 | 0.005705602 | 0.00575388 | 0.004486715 | 0.003843275 | 0.002268064 | 0.003439806 |
| NITMOv2_3106 | A0A0K2GEX3 | HTH cro/C1-type domain-containing protein | 11.914 | 105 | 1.28 | 2.43 | 1.40E-03 | 3.67E-03 | yes | yes | 1 | 2 | 2 | 2 | 1 | 1 | 1 | 0.005703915 | 0.006662963 | 0.006283932 | 0.003974321 | 0.003527944 | 0.003882324 |
| treS | A0A0K2GH45 | Maltose alpha-D-glucosyltransferase | 65.226 | 559 | 0.225 | 1.17 | 5.71E-02 | 7.86E-02 | no | no | 0 | 12 | 13 | 18 | 14 | 13 | 12 | 0.005696538 | 0.005169203 | 0.005377271 | 0.004766816 | 0.005080537 | 0.003872974 |
| hisZ | A0A0K2GGU9 | ATP phosphoribosyltransferase regulatory subunit | 37.889 | 344 | 0.209 | 1.16 | 5.60E-02 | 7.73E-02 | no | no | 0 | 5 | 7 | 9 | 7 | 7 | 8 | 0.005670611 | 0.006943569 | 0.00716241 | 0.006399042 | 0.007067074 | 0.007501762 |
| gspF | A0A0K2GC48 | General secretion pathway protein F | 44.847 | 410 | 0.565 | 1.48 | 9.41E-04 | 2.70E-03 | no | no | 0 | 9 | 10 | 8 | 7 | 7 | 5 | 0.005632458 | 0.005334829 | 0.005207788 | 0.003620571 | 0.003640896 | 0.00311039 |
| vapC.4 | A0A0K2GIA7 | Ribonuclease VapC | 14.027 | 127 | 0.374 | 1.30 | 7.58E-03 | 1.41E-02 | no | no | 0 | 3 | 2 | 2 | 4 | 1 | 2 | 0.005629297 | 0.00432504 | 0.00589435 | 0.005215482 | 0.005335652 | 0.004481458 |
| NITMOv2_0798 | A0A0K2G8E6 | Uncharacterized protein | 15.263 | 140 | 0.0371 | 1.03 | 7.33E-01 | 7.65E-01 | no | no | 0 | 4 | 5 | 5 | 5 | 6 | 5 | 0.005619179 | 0.006171953 | 0.004713333 | 0.004651022 | 0.006364189 | 0.006215566 |
| NITMOv2_1383 | A0A0K2GA37 | Uncharacterized protein | 26.437 | 244 | 0.291 | 1.22 | 2.07E-01 | 2.47E-01 | no | no | 0 | 5 | 6 | 11 | 9 | 6 | 5 | 0.005579761 | 0.005591187 | 0.006685977 | 0.002648609 | 0.003201205 | 0.00328265 |
| NITMOv2_1259 | A0A0K2GAQ7 | Uncharacterized protein | 15.715 | 141 | -0.62 | -1.54 | 2.94E-05 | 2.43E-04 | no | no | 0 | 4 | 4 | 4 | 6 | 4 | 5 | 0.005578918 | 0.006130497 | 0.005259553 | 0.011764079 | 0.010902336 | 0.009361559 |
| NITMOv2_1981 | A0A0K2GBS0 | Uncharacterized protein | 47.09 | 419 | 0.516 | 1.43 | 2.12E-04 | 9.18E-04 | no | no | 0 | 16 | 12 | 14 | 13 | 13 | 16 | 0.005556364 | 0.004960363 | 0.00584316 | 0.003470571 | 0.003884203 | 0.003650546 |
| hemD | A0A0K2GBE6 | SUMT | 56.198 | 521 | -0.309 | -1.24 | 4.51E-03 | 9.11E-03 | no | no | 0 | 18 | 19 | 16 | 25 | 22 | 20 | 0.005550251 | 0.006624441 | 0.006902049 | 0.011766448 | 0.011830505 | 0.010613411 |
| NITMOv2_3414 | A0A0K2GFS7 | Uncharacterized protein | 53.326 | 491 | 0.163 | 1.12 | 8.10E-02 | 1.07E-01 | no | no | 0 | 7 | 5 | 7 | 5 | 3 | 3 | 0.005539923 | 0.00563714 | 0.006328795 | 0.004963755 | 0.004801652 | 0.005751831 |
| hisC | A0A0K2G736 | Histidinol-phosphate aminotransferase | 39.261 | 358 | -0.931 | -1.91 | 4.06E-05 | 3.03E-04 | no | no | 0 | 7 | 7 | 8 | 20 | 24 | 16 | 0.005523903 | 0.005879419 | 0.006002865 | 0.017520181 | 0.016594868 | 0.013542011 |
| NITMOv2_4545 | A0A0K2GJ92 | Uncharacterized protein | 35.693 | 337 | -0.549 | -1.46 | 7.54E-04 | 2.30E-03 | no | no | 0 | 0 | 0 | 0 | 4 | 5 | 5 | 0.005514206 | 0.005540346 | 0.005269714 | 0.004940804 | 0.005058941 | 0.006255844 |
| NITMOv2_1289 | A0A0K2GAU2 | Putative (Glutamate-ammonia-ligase) adenylyltransferase, subunit A (Modular protein) | 84.919 | 751 | -0.297 | -1.23 | 3.16E-02 | 4.72E-02 | no | no | 0 | 40 | 42 | 45 | 25 | 33 | 32 | 0.005497765 | 0.005992052 | 0.005845077 | 0.004491249 | 0.004460774 | 0.004521916 |
| NITMOv2_4392 | A0A0K2GJF7 | Uncharacterized protein | 46.568 | 430 | 0.407 | 1.33 | 3.81E-03 | 8.00E-03 | no | no | 0 | 7 | 7 | 6 | 5 | 5 | 5 | 0.005474789 | 0.005127748 | 0.004686108 | 0.005172392 | 0.003929415 | 0.003804105 |
| NITMOv2_4436 | A0A0K2GIN5 | Putative Glycosyl transferase, family 39 | 64.058 | 576 | 0.284 | 1.22 | 2.30E-02 | 3.58E-02 | no | no | 0 | 5 | 4 | 7 | 6 | 11 | 7 | 0.005468255 | 0.005757008 | 0.005593919 | 0.004644506 | 0.006853288 | 0.006637225 |
| NITMOv2_1022 | A0A0K2G9C5 | Putative Glycosyl hydrolase, family 13 | 13.013 | 124 | 0.296 | 1.23 | 2.21E-01 | 2.62E-01 | no | no | 0 | 3 | 4 | 3 | 3 | 2 | 2 | 0.005467412 | 0.006918735 | 0.006322085 | 0.003805516 | 0.004319078 | 0.003443402 |
| purK | A0A0K2GJ53 | N5-carboxyaminoimidazole ribonucleotide synthase | 40.507 | 377 | -0.0838 | -1.06 | 4.76E-01 | 5.19E-01 | no | no | 0 | 9 | 12 | 11 | 14 | 13 | 13 | 0.005463617 | 0.005663147 | 0.005904319 | 0.006906937 | 0.006405051 | 0.0054751 |
| NITMOv2_1082 | A0A0K2G983 | FGE-sulfatase domain-containing protein | 34.062 | 299 | 0.962 | 1.95 | 9.09E-06 | 1.10E-04 | no | no | 0 | 7 | 8 | 11 | 3 | 5 | 4 | 0.005460034 | 0.004655705 | 0.004593505 | 0.001521762 | 0.002607388 | 0.00263227 |
| NITMOv2_1012 | A0A0K2G9B6 | Uncharacterized protein | 28.523 | 254 | 0.937 | 1.91 | 1.22E-03 | 3.24E-03 | no | no | 0 | 1 | 2 | 3 | 2 | 0 | 0 | 0.005457505 | 0.00550417 | 0.005408522 | 0.002541552 | 0.003347872 | 0.004786779 |
| ribF | A0A0K2GC60 | Riboflavin biosynthesis protein | 34.489 | 315 | -0.205 | -1.15 | 4.76E-02 | 6.73E-02 | no | no | 0 | 9 | 9 | 7 | 8 | 7 | 9 | 0.005452024 | 0.005633229 | 0.005582032 | 0.005309509 | 0.005156823 | 0.006064883 |
| thiL | A0A0K2GG58 | Thiamine-monophosphate kinase | 37.031 | 347 | 0.315 | 1.24 | 1.23E-02 | 2.13E-02 | no | no | 0 | 8 | 6 | 9 | 5 | 7 | 7 | 0.005450338 | 0.006626788 | 0.00746284 | 0.004764595 | 0.004490139 | 0.005324057 |
| NITMOv2_3991 | A0A0K2GHE1 | Putative Histidine kinase | 70.11 | 637 | -0.205 | -1.15 | 3.35E-02 | 4.97E-02 | no | no | 0 | 10 | 8 | 8 | 14 | 11 | 12 | 0.005430313 | 0.005613479 | 0.005973723 | 0.006452052 | 0.006954588 | 0.006988578 |
| thiC | A0A0K2G9W3 | Phosphomethylpyrimidine synthase | 70.166 | 635 | -1.65 | -3.14 | 7.50E-07 | 2.63E-05 | yes | no | 0 | 7 | 7 | 6 | 31 | 30 | 30 | 0.005418298 | 0.005515707 | 0.006222772 | 0.021119873 | 0.020287968 | 0.02063524 |
| mobB | A0A0K2GHH0 | Molybdopterin-guanine dinucleotide biosynthesis protein B | 19.301 | 177 | -0.508 | -1.42 | 5.31E-04 | 1.78E-03 | no | no | 0 | 2 | 2 | 3 | 6 | 4 | 6 | 0.005394901 | 0.005359663 | 0.005183822 | 0.013584359 | 0.013137928 | 0.013408231 |
| NITMOv2_1061 | A0A0K2GA82 | SpoVT-AbrB domain-containing protein | 9.1406 | 78 | 0.406 | 1.33 | 2.94E-02 | 4.45E-02 | no | no | 0 | 1 | 3 | 1 | 1 | 1 | 0 | 0.005373611 | 0.005517663 | 0.006727389 | 0.002871314 | 0.0025976 | 0.002025044 |
| NITMOv2_4807 | A0A0K2GJP3 | TPR_REGION domain-containing protein | 25.096 | 226 | 0.36 | 1.28 | 9.72E-02 | 1.26E-01 | no | no | 0 | 6 | 5 | 5 | 4 | 3 | 3 | 0.005350424 | 0.005040144 | 0.004246869 | 0.005132116 | 0.004418048 | 0.003581858 |
| rffH | A0A0K2GG89 | Glucose-1-phosphate thymidylyltransferase | 26.494 | 243 | -1.26 | -2.39 | 7.59E-06 | 9.86E-05 | yes | no | 0 | 3 | 2 | 2 | 8 | 8 | 7 | 0.005343468 | 0.006047391 | 0.006309431 | 0.014881492 | 0.015418887 | 0.013072702 |
| NITMOv2_4084 | A0A0K2GIN6 | Uncharacterized protein | 27.318 | 250 | -0.169 | -1.12 | 2.35E-01 | 2.77E-01 | no | no | 0 | 19 | 20 | 19 | 10 | 14 | 13 | 0.005335248 | 0.006119938 | 0.006107163 | 0.005359114 | 0.005392983 | 0.00512069 |
| NITMOv2_1474 | A0A0K2GAC5 | Uncharacterized protein | 25.959 | 234 | 0.116 | 1.08 | 1.73E-01 | 2.09E-01 | no | no | 0 | 4 | 4 | 4 | 3 | 3 | 2 | 0.005325762 | 0.005382542 | 0.005231753 | 0.005308028 | 0.005456373 | 0.005265798 |
| NITMOv2_3525 | A0A0K2GH39 | Putative Phosphatidylinositol alpha-mannosyltransferase | 40.932 | 371 | -0.16 | -1.12 | 7.81E-02 | 1.04E-01 | no | no | 0 | 12 | 11 | 7 | 10 | 9 | 10 | 0.005306791 | 0.00550417 | 0.004901605 | 0.006624411 | 0.006813047 | 0.00620208 |
| NITMOv2_1373 | A0A0K2GA26 | PSII_BNR domain-containing protein | 115.34 | 1035 | 0.115 | 1.08 | 2.47E-01 | 2.88E-01 | no | no | 0 | 16 | 16 | 14 | 13 | 9 | 13 | 0.005259996 | 0.006496556 | 0.006112915 | 0.005707385 | 0.006147761 | 0.006058051 |
| hisF | A0A0K2G8I8 | Imidazole glycerol phosphate synthase subunit HisF | 27.608 | 260 | 0.00678 | 1.00 | 9.40E-01 | 9.49E-01 | no | no | 0 | 4 | 3 | 5 | 4 | 6 | 5 | 0.005255148 | 0.005444138 | 0.005091603 | 0.007002742 | 0.006950548 | 0.006743674 |
| rpoE | A0A0K2GA48 | RNA polymerase sigma factor | 22.418 | 200 | 0.646 | 1.56 | 7.22E-05 | 4.50E-04 | no | no | 0 | 5 | 4 | 5 | 2 | 1 | 1 | 0.00525304 | 0.004325236 | 0.005702051 | 0.002835331 | 0.002065308 | 0.002172131 |
| gspE | A0A0K2GD48 | General secretion pathway protein E | 66.52 | 609 | -0.197 | -1.15 | 3.31E-02 | 4.92E-02 | no | no | 0 | 11 | 12 | 7 | 7 | 11 | 10 | 0.005248403 | 0.005562247 | 0.005055559 | 0.006520167 | 0.007087427 | 0.006148676 |
| NITMOv2_0222 | A0A0K2G6R1 | Putative periplasmic serine endoprotease DegP-like | 51.751 | 492 | -2.88 | -7.36 | 1.37E-05 | 1.44E-04 | yes | no | 0 | 9 | 10 | 10 | 68 | 62 | 50 | 0.005236177 | 0.006399175 | 0.006639388 | 0.097122097 | 0.088319005 | 0.057144296 |
| NITMOv2_1123 | A0A0K2G9L9 | Response regulator, contains GGDEF domain | 36.786 | 334 | 0.101 | 1.07 | 4.56E-01 | 4.99E-01 | no | no | 0 | 12 | 13 | 14 | 9 | 13 | 12 | 0.005236177 | 0.005070258 | 0.007409925 | 0.004775552 | 0.004026365 | 0.003781269 |
| NITMOv2_2130 | A0A0K2GD68 | Putative Response regulator, CheY-like | 22.845 | 210 | -0.559 | -1.47 | 8.54E-04 | 2.52E-03 | no | no | 0 | 5 | 4 | 5 | 8 | 8 | 7 | 0.005208353 | 0.004652381 | 0.00532378 | 0.009678893 | 0.01029205 | 0.008653279 |
| NITMOv2_3721 | A0A0K2GGM1 | Putative Phosphate-binding protein PstS | 31.05 | 292 | 0.465 | 1.38 | 8.78E-02 | 1.15E-01 | no | no | 0 | 6 | 9 | 8 | 4 | 4 | 3 | 0.005206246 | 0.006732773 | 0.004102501 | 0.003429999 | 0.002961006 | 0.003117403 |
| NITMOv2_4440 | A0A0K2GIW9 | Putative Response regulator, CheY-like | 28.685 | 261 | -0.0564 | -1.04 | 7.32E-01 | 7.64E-01 | no | no | 0 | 6 | 5 | 7 | 6 | 6 | 5 | 0.0051995 | 0.005029781 | 0.004904289 | 0.00455196 | 0.004634631 | 0.004148626 |
| NITMOv2_3885 | A0A0K2GI23 | Putative Small-conductance mechanosensitive ion channel | 33.582 | 308 | 0.317 | 1.25 | 1.79E-02 | 2.89E-02 | no | no | 0 | 5 | 5 | 5 | 6 | 3 | 4 | 0.005187275 | 0.006010825 | 0.006669872 | 0.005494158 | 0.005022274 | 0.004160853 |
| top6B | A0A0K2GCK6 | Type 2 DNA topoisomerase 6 subunit B | 74.807 | 681 | -0.806 | -1.75 | 4.01E-05 | 3.00E-04 | no | no | 0 | 10 | 8 | 11 | 25 | 21 | 20 | 0.005170833 | 0.005847154 | 0.006121159 | 0.012771282 | 0.011720038 | 0.011431197 |

| | | | | | | | | | | | | | | | | | | | | | | | |
|---|---|---|---|---|---|---|---|---|---|---|---|---|---|---|---|---|---|---|---|---|---|---|---|
| NITMOv2_4058 | A0A0K2GIL3 | AAA domain-containing protein | 47.19 | 431 | 0.624 | 1.54 | 9.38E-05 | 5.41E-04 | no | no | 0 | 14 | 12 | 10 | 6 | 6 | 8 | 0.00514849 | 0.005093137 | 0.005084509 | 0.003698902 | 0.003261954 | 0.002758678 |
| NITMOv2_0580 | A0A0K2G7S0 | Uncharacterized protein | 10.6 | 96 | -0.6 | -1.52 | 5.94E-04 | 1.94E-03 | no | no | 0 | 2 | 2 | 4 | 4 | 3 | 4 | 0.005146171 | 0.005616217 | 0.005532759 | 0.007106098 | 0.00721716 | 0.008495044 |
| NITMOv2_2790 | A0A0K2GEA6 | Uncharacterized protein | 9.0762 | 78 | 0.0447 | 1.03 | 4.38E-01 | 4.81E-01 | no | no | 0 | 3 | 2 | 2 | 0 | 0 | 0 | 0.005135421 | 0.005139285 | 0.005092562 | 0.004111586 | 0.003888709 | 0.003824784 |
| folC | A0A0K2GAG2 | Bifunctional protein FolC: Folylpolyglutamate synthase and Dihydrofolate synthase | 46.864 | 433 | -3.06 | -8.34 | 2.60E-02 | 3.98E-02 | yes | yes | 2 | 0 | 0 | 1 | 8 | 6 | 4 | 0.005118558 | 0.005558531 | 0.005454728 | 0.002364306 | 0.008048845 | 0.009936419 |
| NITMOv2_4865 | A0A0K2GJV2 | Uncharacterized protein | 12.817 | 113 | -0.693 | -1.62 | 4.15E-03 | 8.56E-03 | no | no | 0 | 4 | 8 | 7 | 12 | 13 | 14 | 0.005114764 | 0.004985197 | 0.006983532 | 0.007755405 | 0.010359635 | 0.009110721 |
| trpA | A0A0K2GCL9 | Tryptophan synthase alpha chain | 31.694 | 302 | -0.522 | -1.44 | 2.84E-03 | 6.34E-03 | no | no | 0 | 9 | 8 | 5 | 9 | 7 | 6 | 0.005112656 | 0.005013159 | 0.005326656 | 0.00940303 | 0.009160038 | 0.00806835 |
| NITMOv2_0397 | A0A0K2G799 | Uncharacterized protein | 30.992 | 276 | 1.12 | 2.17 | 8.30E-02 | 1.10E-01 | no | yes | 1 | 3 | 2 | 2 | 1 | 1 | 0 | 0.005105278 | 0.00433736 | 0.005074156 | 0.002502756 | 0.002544774 | 0.002325331 |
| NITMOv2_1462 | A0A0K2GAI4 | HipA_C domain-containing protein | 34.535 | 307 | 0.449 | 1.37 | 9.37E-03 | 1.69E-02 | no | no | 0 | 5 | 4 | 7 | 8 | 3 | 4 | 0.005098322 | 0.004347919 | 0.005205487 | 0.003557491 | 0.00312756 | 0.003516586 |
| NITMOv2_0044 | A0A0K2G6C4 | Putative 2-succinylbenzoate--CoA ligase | 63.858 | 572 | 4.57 | 23.75 | 7.19E-05 | 4.49E-04 | yes | no | 0 | 11 | 12 | 12 | 0 | 0 | 1 | 0.005085886 | 0.005817432 | 0.005327231 | 0.000224126 | 0.00037467 | 0.000423601 |
| NITMOv2_4296 | A0A0K2GI77 | Pseudouridine synthase | 29.118 | 257 | 0.46 | 1.38 | 3.60E-03 | 7.67E-03 | no | no | 0 | 5 | 7 | 7 | 4 | 3 | 3 | 0.005081249 | 0.005479727 | 0.004656391 | 0.003766868 | 0.003822056 | 0.003112368 |
| trxB | A0A0K2G8Z8 | Thioredoxin reductase | 32.573 | 306 | 0.219 | 1.16 | 2.37E-02 | 3.68E-02 | no | no | 0 | 5 | 6 | 3 | 3 | 4 | 2 | 0.005063332 | 0.007708537 | 0.007407624 | 0.003856009 | 0.004105136 | 0.002083303 |
| murE | A0A0K2G7X0 | UDP-N-acetylmuramoyl-L-alanyl-D-glutamate--2,6-diaminopimelate ligase | 53.494 | 500 | -0.638 | -1.56 | 2.96E-04 | 1.15E-03 | no | no | 0 | 8 | 9 | 12 | 19 | 18 | 13 | 0.005059537 | 0.005984426 | 0.005967588 | 0.01030051 | 0.010088673 | 0.008808277 |
| NITMOv2_0595 | A0A0K2G7T2 | Putative Acriflavine resistance protein acrA | 43.555 | 395 | 0.536 | 1.45 | 2.55E-03 | 5.79E-03 | no | no | 0 | 10 | 10 | 10 | 6 | 7 | 6 | 0.005028762 | 0.006024904 | 0.004833543 | 0.003335972 | 0.003500599 | 0.003759512 |
| NITMOv2_4368 | A0A0K2GIP9 | Fido domain-containing protein | 38.009 | 334 | 0.294 | 1.23 | 1.43E-02 | 2.41E-02 | no | no | 0 | 2 | 1 | 4 | 4 | 2 | 1 | 0.004997987 | 0.004944328 | 0.0038176 | 0.003172201 | 0.003265527 | 0.002546679 |
| NITMOv2_1669 | A0A0K2GAU4 | Transcriptional regulator, Crp/Fnr family | 24.755 | 220 | 0.179 | 1.13 | 4.26E-01 | 4.69E-01 | no | no | 0 | 4 | 3 | 5 | 4 | 6 | 2 | 0.004994193 | 0.005173897 | 0.004967175 | 0.004840557 | 0.004444616 | 0.004131543 |
| NITMOv2_1254 | A0A0K2GAQ2 | Uncharacterized protein | 17.804 | 158 | -0.101 | -1.07 | 1.37E-01 | 1.70E-01 | no | no | 0 | 2 | 3 | 4 | 2 | 1 | 2 | 0.004980914 | 0.00520929 | 0.005448401 | 0.00595082 | 0.006117464 | 0.005874822 |
| NITMOv2_3444 | A0A0K2GFW2 | Putative P-loop guanosine triphosphatase, CobW-like | 35.032 | 323 | 0.585 | 1.50 | 2.41E-04 | 1.00E-03 | no | no | 0 | 3 | 4 | 5 | 2 | 3 | 3 | 0.004933065 | 0.005482465 | 0.005808458 | 0.002865983 | 0.003698849 | 0.003228886 |
| gcvP | A0A0K2G6R6 | Glycine dehydrogenase (decarboxylating) | 105.61 | 962 | -1.08 | -2.11 | 3.63E-06 | 6.27E-05 | yes | no | 0 | 13 | 16 | 16 | 41 | 43 | 35 | 0.004928427 | 0.005138894 | 0.006173499 | 0.017946635 | 0.016242183 | 0.016043917 |
| NITMOv2_3519 | A0A0K2GG51 | Putative Mannosyltransferase | 41.132 | 372 | -0.24 | -1.18 | 2.84E-02 | 4.30E-02 | no | no | 0 | 7 | 8 | 7 | 8 | 8 | 7 | 0.004917256 | 0.004979135 | 0.004650639 | 0.006620117 | 0.006642142 | 0.005999612 |
| pyrG | A0A0K2GFH0 | CTP synthase | 59.51 | 533 | -1.83 | -3.56 | 4.98E-07 | 1.87E-05 | yes | no | 0 | 10 | 11 | 9 | 41 | 40 | 31 | 0.004905873 | 0.004718475 | 0.00456839 | 0.02651719 | 0.025572038 | 0.023485263 |
| NITMOv2_2372 | A0A0K2GDV4 | Uncharacterized protein | 28.278 | 248 | -0.228 | -1.17 | 3.84E-02 | 5.58E-02 | no | no | 0 | 7 | 6 | 9 | 12 | 9 | 9 | 0.004903133 | 0.005645353 | 0.006030665 | 0.007862166 | 0.008014198 | 0.007232044 |
| NITMOv2_0015 | A0A0K2G6I8 | Sigma-54 dependent transcriptional regulator | 78.825 | 718 | -0.204 | -1.15 | 5.26E-02 | 7.36E-02 | no | no | 0 | 11 | 15 | 16 | 28 | 26 | 25 | 0.004899339 | 0.003953702 | 0.005846802 | 0.008360437 | 0.007884155 | 0.007782629 |
| gcp | A0A0K2GC22 | tRNA N6-adenosine threonylcarbamoyltransferase | 37.834 | 362 | -0.262 | -1.20 | 1.38E-01 | 1.72E-01 | no | no | 0 | 3 | 4 | 4 | 8 | 6 | 5 | 0.004879946 | 0.005390755 | 0.003703524 | 0.007424162 | 0.007625778 | 0.004969288 |
| NITMOv2_0393 | A0A0K2G7H8 | Uncharacterized protein | 19.783 | 181 | -0.823 | -1.77 | 1.32E-02 | 2.26E-02 | no | no | 0 | 3 | 3 | 4 | 8 | 8 | 6 | 0.004878471 | 0.006277742 | 0.00862181 | 0.011076569 | 0.010499156 | 0.008853949 |
| NITMOv2_3422 | A0A0K2GG25 | Uncharacterized protein | 19.345 | 174 | 0.313 | 1.24 | 8.10E-02 | 1.07E-01 | no | no | 0 | 1 | 1 | 1 | 2 | 1 | 1 | 0.004869828 | 0.003970715 | 0.003489944 | 0.005782903 | 0.004685902 | 0.003878368 |
| fabH | A0A0K2G811 | 3-oxoacyl-[acyl-carrier-protein] synthase 3 | 34.264 | 324 | -0.591 | -1.51 | 2.03E-03 | 4.87E-03 | no | no | 0 | 4 | 3 | 3 | 5 | 7 | 6 | 0.004862029 | 0.004552654 | 0.004780819 | 0.008203478 | 0.008759499 | 0.008233777 |
| NITMOv2_3909 | A0A0K2GH59 | Putative Cation efflux system protein CzcB | 43.609 | 400 | 0.505 | 1.42 | 3.43E-03 | 7.37E-03 | no | no | 0 | 13 | 13 | 15 | 4 | 5 | 4 | 0.004851279 | 0.0051123 | 0.004882625 | 0.002975262 | 0.002446582 | 0.002291166 |
| NITMOv2_3961 | A0A0K2GHB1 | Putative Quinol-cytochrome c reductase, iron-sulfur subunit (Rieske iron-sulfur protein) | 19.317 | 174 | 0.306 | 1.24 | 9.00E-02 | 1.18E-01 | no | no | 0 | 4 | 4 | 3 | 0 | 2 | 0 | 0.004850225 | 0.004366691 | 0.002736662 | 0.002657494 | 0.002887827 | 0.002289008 |
| NITMOv2_3409 | A0A0K2GFS2 | NAD-dependent epimerase/dehydratase | 38.34 | 342 | -1.66 | -3.16 | 8.41E-06 | 1.05E-04 | yes | no | 0 | 6 | 4 | 3 | 18 | 20 | 19 | 0.004841583 | 0.003631642 | 0.00376756 | 0.018048807 | 0.017638942 | 0.016144792 |
| NITMOv2_1137 | A0A0K2G9D9 | Ferredoxin, 2Fe-2S | 12.794 | 116 | -1.52 | -2.87 | 1.45E-05 | 1.49E-04 | yes | no | 0 | 1 | 1 | 2 | 5 | 5 | 5 | 0.004823666 | 0.003980883 | 0.003120493 | 0.010617241 | 0.009809011 | 0.008406397 |
| devR.1 | A0A0K2G7M3 | Transcriptional regulatory protein DevR (DosR) | 24.513 | 227 | -0.226 | -1.17 | 6.51E-02 | 8.84E-02 | no | no | 0 | 4 | 5 | 6 | 6 | 6 | 5 | 0.004820083 | 0.004028205 | 0.003692596 | 0.004676787 | 0.004687767 | 0.004598516 |
| NITMOv2_4632 | A0A0K2GJ78 | Putative Glyoxalase | 16.228 | 144 | 0.708 | 1.63 | 1.64E-03 | 4.13E-03 | no | no | 0 | 2 | 3 | 3 | 0 | 1 | 1 | 0.004814813 | 0.004429265 | 0.004219261 | 0.00363908 | 0.003506814 | 0.002772343 |
| NITMOv2_0966 | A0A0K2G8W3 | ABC transporter domain-containing protein | 25.241 | 232 | -0.902 | -1.87 | 2.11E-02 | 3.34E-02 | no | no | 0 | 2 | 2 | 3 | 2 | 2 | 2 | 0.00480596 | 0.003552055 | 0.004397947 | 0.004664348 | 0.004675182 | 0.003204252 |
| NITMOv2_4719 | A0A0K2GJG1 | Uncharacterized protein | 13.149 | 120 | 1.02 | 2.03 | 1.73E-02 | 2.81E-02 | yes | yes | 3 | 3 | 4 | 4 | 0 | 0 | 0 | 0.004793523 | 0.00480471 | 0.004799608 | 0.000556597 | 0 | 0.000232929 |
| folP | A0A0K2GB58 | Dihydropteroate synthase | 31.835 | 297 | 0.615 | 1.53 | 1.25E-02 | 2.15E-02 | no | no | 0 | 7 | 7 | 14 | 4 | 4 | 5 | 0.004789308 | 0.004683668 | 0.006624817 | 0.001916825 | 0.002032681 | 0.001973978 |
| NITMOv2_4172 | A0A0K2GIV8 | Putative Cell division protein ZapA | 11.254 | 98 | -3.31 | -9.92 | 1.83E-03 | 4.47E-03 | yes | no | 0 | 2 | 1 | 6 | 14 | 15 | 14 | 0.004785724 | 0.002713954 | 0.003829103 | 0.011711068 | 0.011081475 | 0.014617648 |
| NITMOv2_0940 | A0A0K2G928 | YCII domain-containing protein | 17.735 | 158 | 0.417 | 1.34 | 2.59E-02 | 3.97E-02 | no | no | 0 | 5 | 5 | 5 | 2 | 3 | 1 | 0.004782562 | 0.004707133 | 0.004865561 | 0.001281526 | 0.001973486 | 0.001511735 |
| NITMOv2_1328 | A0A0K2G9V9 | Uncharacterized protein | 12.651 | 108 | 2.77 | 6.82 | 1.47E-08 | 3.40E-06 | yes | yes | 3 | 3 | 3 | 2 | 0 | 0 | 1 | 0.004766964 | 0.011841358 | 0.005555383 | 0.000144296 | 0 | 0.000375322 |
| NITMOv2_1648 | A0A0K2GBT5 | Uncharacterized protein | 34.23 | 312 | 1.65 | 3.14 | 1.19E-01 | 1.50E-01 | no | yes | 1 | 3 | 4 | 3 | 0 | 0 | 1 | 0.004727336 | 0.005312537 | 0.005559984 | 0.00081438 | 0.001789064 | 0.002524562 |
| NITMOv2_3660 | A0A0K2GHG0 | Plastocyanin-like domain-containing protein | 178.27 | 1614 | -3.74 | -13.36 | 8.60E-10 | 1.79E-06 | yes | no | 0 | 12 | 14 | 13 | 199 | 205 | 193 | 0.004716375 | 0.004662745 | 0.004612103 | 0.078910408 | 0.080062743 | 0.075253176 |
| NITMOv2_2318 | A0A0K2GCQ8 | Uncharacterized protein | 10.017 | 86 | -1.23 | -2.35 | 6.89E-02 | 9.30E-02 | no | yes | 3 | 1 | 2 | 1 | 2 | 2 | 1 | 0.004715321 | 0.00470068 | 0.004517583 | 0.00289234 | 0.004588797 | 0.003920444 |
| ribBA | A0A0K2G9V5 | Riboflavin biosynthesis protein RibBA | 44.345 | 402 | -0.971 | -1.96 | 3.53E-04 | 1.31E-03 | no | no | 0 | 3 | 2 | 5 | 14 | 12 | 10 | 0.004696983 | 0.00456204 | 0.004707964 | 0.010465169 | 0.010585696 | 0.009607362 |
| NITMOv2_4639 | A0A0K2GJ90 | TPM_phosphatase domain-containing protein | 28.971 | 283 | 1.33 | 2.51 | 1.59E-04 | 7.59E-04 | yes | no | 0 | 1 | 3 | 2 | 0 | 1 | 0 | 0.004685178 | 0.004112875 | 0.004113046 | 0.003368844 | 0.003069763 | 0.003625013 |
| cobP | A0A0K2GAN4 | Adenosylcobinamide kinase | 20.352 | 187 | -1.11 | -2.16 | 4.48E-06 | 7.02E-05 | yes | no | 0 | 5 | 5 | 4 | 12 | 10 | 12 | 0.004676325 | 0.005867491 | 0.006017628 | 0.012642457 | 0.014095617 | 0.012954925 |
| accA | A0A0K2GBY5 | Acetyl-coenzyme A carboxylase carboxyl transferase subunit alpha | 35.501 | 321 | -0.522 | -1.44 | 1.42E-04 | 7.06E-04 | no | no | 0 | 5 | 5 | 6 | 10 | 14 | 12 | 0.004667472 | 0.004733923 | 0.005126305 | 0.008648738 | 0.0092009 | 0.008330157 |
| NITMOv2_3506 | A0A0K2GG16 | Glyco_trans_2-like domain-containing protein | 39.75 | 341 | -0.47 | -1.39 | 1.58E-03 | 4.04E-03 | no | no | 0 | 4 | 3 | 4 | 7 | 5 | 4 | 0.004631638 | 0.005466235 | 0.004950303 | 0.006373277 | 0.006540842 | 0.006009501 |
| NITMOv2_3560 | A0A0K2GH69 | Cell division protein FtsX | 32.948 | 299 | 0.0754 | 1.05 | 5.04E-01 | 5.46E-01 | no | no | 0 | 3 | 3 | 3 | 6 | 5 | 5 | 0.004621731 | 0.004632044 | 0.004681315 | 0.004272395 | 0.005015127 | 0.004670261 |
| trxC | A0A0K2GHB9 | Thioredoxin | 15.924 | 144 | 1.1 | 2.14 | 6.95E-04 | 2.17E-03 | yes | yes | 2 | 2 | 1 | 0 | 0 | 0 | 0 | 0.004617516 | 0.004582376 | 0.005249967 | 0.001197124 | 0.000959709 | 0.001999871 |
| merA | A0A0K2GIB4 | Mercuric reductase | 59.027 | 554 | 0.27 | 1.21 | 7.37E-02 | 9.87E-02 | no | no | 0 | 6 | 9 | 12 | 7 | 8 | 9 | 0.004612246 | 0.005375698 | 0.005414657 | 0.00452812 | 0.004868305 | 0.00433545 |
| lig | A0A0K2GHU0 | Probable DNA ligase | 56.287 | 511 | 0.916 | 1.89 | 6.61E-05 | 4.24E-04 | no | no | 0 | 13 | 18 | 16 | 8 | 9 | 7 | 0.00460255 | 0.005396817 | 0.005795996 | 0.002697918 | 0.002705736 | 0.002466123 |
| panC | A0A0K2G7K0 | Pantothenate synthetase | 31.717 | 292 | 0.136 | 1.10 | 2.50E-01 | 2.92E-01 | no | no | 0 | 5 | 4 | 4 | 2 | 3 | 2 | 0.004599177 | 0.004269897 | 0.004656774 | 0.003522101 | 0.002225027 | 0.00289012 |
| NITMOv2_4292 | A0A0K2GIH5 | Putative Efflux transporter, membrane fusion protein | 53.045 | 478 | -0.625 | -1.54 | 4.41E-05 | 3.19E-04 | no | no | 0 | 12 | 15 | 12 | 19 | 14 | 12 | 0.004578309 | 0.005880592 | 0.00517692 | 0.008375837 | 0.008465853 | 0.007882784 |
| coaE | A0A0K2G924 | Dephospho-CoA kinase | 22.505 | 199 | 0.388 | 1.31 | 1.66E-03 | 4.16E-03 | no | no | 0 | 4 | 3 | 3 | 1 | 3 | 2 | 0.004572196 | 0.004927316 | 0.005127072 | 0.003902801 | 0.004006011 | 0.004483616 |
| NITMOv2_3203 | A0A0K2GG47 | Uncharacterized protein | 28.284 | 252 | 0.484 | 1.40 | 1.69E-02 | 2.76E-02 | no | no | 0 | 4 | 3 | 3 | 3 | 3 | 2 | 0.004564819 | 0.004539748 | 0.00318798 | 0.003252013 | 0.003314779 | 0.001879217 |
| NITMOv2_4212 | A0A0K2GI32 | TPR_REGION domain-containing protein | 50.011 | 454 | 0.503 | 1.42 | 3.23E-04 | 1.22E-03 | no | no | 0 | 9 | 8 | 9 | 7 | 8 | 6 | 0.004553436 | 0.004420661 | 0.00423805 | 0.003822989 | 0.003571291 | 0.003000165 |
| NITMOv2_0530 | A0A0K2G7M5 | Putative tRNA-processing ribonuclease BN | 31.782 | 284 | 1.21 | 2.31 | 4.73E-02 | 6.70E-02 | no | yes | 3 | 1 | 0 | 1 | 1 | 1 | 0 | 0.004545848 | 0.005924981 | 0.005026609 | 0.003002804 | 0.003217829 | 0.002611412 |
| NITMOv2_3971 | A0A0K2GHC1 | Uncharacterized protein | 73.038 | 652 | 0.384 | 1.30 | 7.65E-04 | 2.32E-03 | no | no | 0 | 14 | 15 | 18 | 20 | 17 | 17 | 0.004505587 | 0.004120501 | 0.004631275 | 0.003821064 | 0.003375062 | 0.003369859 |
| NITMOv2_0415 | A0A0K2G795 | Putative DNA recombination protein RmuC | 51.45 | 465 | -1.73 | -3.32 | 2.41E-07 | 1.33E-05 | yes | no | 0 | 3 | 3 | 5 | 13 | 13 | 13 | 0.004495259 | 0.004152962 | 0.003744745 | 0.009894341 | 0.01026657 | 0.010536811 |
| treZ | A0A0K2GH32 | Malto-oligosyltrehalose trehalohydrolase | 69.023 | 627 | 0.578 | 1.49 | 1.02E-02 | 1.82E-02 | no | no | 0 | 9 | 7 | 8 | 3 | 3 | 4 | 0.004480504 | 0.004477369 | 0.004028112 | 0.00160113 | 0.001901861 | 0.001591607 |
| NITMOv2_4823 | A0A0K2GJS5 | Aminotransferase | 42.053 | 386 | -0.696 | -1.62 | 1.01E-03 | 2.84E-03 | no | no | 0 | 6 | 5 | 6 | 11 | 8 | 9 | 0.004479239 | 0.004496337 | 0.004236899 | 0.00879015 | 0.00759082 | 0.00706356 |
| NITMOv2_0464 | A0A0K2G7K2 | Putative phytochrome sensor protein | 51.283 | 460 | 0.393 | 1.31 | 3.32E-03 | 7.17E-03 | no | no | 0 | 13 | 14 | 13 | 9 | 9 | 9 | 0.004477763 | 0.006599216 | 0.004840637 | 0.002818895 | 0.004969605 | 0.003922062 |
| recX | A0A0K2GD81 | Regulatory protein RecX | 16.935 | 144 | 0.153 | 1.11 | 2.41E-01 | 2.82E-01 | no | no | 0 | 3 | 4 | 4 | 1 | 3 | 3 | 0.004457528 | 0.00359703 | 0.003843674 | 0.00315843 | 0.002903675 | 0.003395033 |
| NITMOv2_3621 | A0A0K2GGD0 | Uncharacterized protein | 42.063 | 382 | 0.0708 | 1.05 | 6.34E-01 | 6.74E-01 | no | no | 0 | 3 | 3 | 1 | 2 | 2 | 1 | 0.004444037 | 0.005139285 | 0.004195487 | 0.004710844 | 0.004278993 | 0.00479541 |
| NITMOv2_3939 | A0A0K2GHI2 | Uncharacterized protein | 11.212 | 96 | 1.78 | 3.43 | 2.13E-03 | 5.07E-03 | yes | yes | 3 | 1 | 2 | 2 | 0 | 0 | 0 | 0.004436238 | 0.004099774 | 0.004267384 | 0.002531186 | 0.002676837 | 0.002690709 |
| NITMOv2_3980 | A0A0K2GHM1 | Uncharacterized protein | 16.961 | 155 | 1.67 | 3.18 | 3.48E-05 | 2.74E-04 | yes | no | 0 | 4 | 4 | 3 | 2 | 1 | 1 | 0.004417689 | 0.004683277 | 0.00338488 | 0.001585434 | 0.001174754 | 0.002045543 |
| NITMOv2_1091 | A0A0K2GAA8 | Aminopyrimidine aminohydrolase | 27.293 | 236 | 0.331 | 1.26 | 1.56E-01 | 1.92E-01 | no | no | 0 | 3 | 2 | 4 | 3 | 3 | 3 | 0.004416846 | 0.004690708 | 0.003740527 | 0.003661587 | 0.004210787 | 0.00579067 |
| cobU | A0A0K2GBB1 | Adenosylcobinamide kinase | 22.641 | 214 | -1.54 | -2.91 | 2.32E-05 | 2.09E-04 | yes | no | 0 | 5 | 6 | 6 | 10 | 11 | 13 | 0.004415581 | 0.003710446 | 0.004357685 | 0.011314524 | 0.010180651 | 0.009630918 |
| truD | A0A0K2G7H7 | tRNA pseudouridine synthase D | 39.81 | 362 | 0.422 | 1.34 | 6.05E-04 | 1.96E-03 | no | no | 0 | 7 | 6 | 7 | 2 | 3 | 5 | 0.004415159 | 0.004447842 | 0.00439603 | 0.003380542 | 0.003133775 | 0.003609369 |
| mraW | A0A0K2G7Z3 | Ribosomal RNA small subunit methyltransferase H | 32.477 | 296 | 0.464 | 1.38 | 1.75E-03 | 4.35E-03 | no | no | 0 | 4 | 6 | 6 | 7 | 4 | 3 | 0.004404831 | 0.005302564 | 0.004582194 | 0.004874022 | 0.004507229 | 0.004244645 |
| NITMOv2_1251 | A0A0K2G9Y4 | Putative urea ABC transporter, urea binding protein UrtA | 48.472 | 438 | 0.277 | 1.21 | 1.56E-02 | 2.60E-02 | no | no | 0 | 9 | 9 | 10 | 9 | 6 | 6 | 0.004393659 | 0.005107021 | 0.005052108 | 0.004348061 | 0.003562125 | 0.003641016 |
| nuoI | A0A0K2G7L4 | NADH-quinone oxidoreductase subunit I | 22.918 | 195 | -0.0751 | -1.05 | 7.59E-01 | 7.87E-01 | no | no | 0 | 3 | 1 | 3 | 5 | 9 | 7 | 0.004390708 | 0.004730207 | 0.003064893 | 0.006042922 | 0.00793698 | 0.006468742 |
| NITMOv2_3956 | A0A0K2GHA5 | Uncharacterized protein | 15.037 | 140 | 2.2 | 4.59 | 4.87E-03 | 9.73E-03 | yes | yes | 2 | 2 | 3 | 0 | 0 | 0 | 0 | 0.004386281 | 0.005144369 | 0.003834088 | 0.000258775 | 0.000375618 | 0.000318106 |
| NITMOv2_4423 | A0A0K2GIK8 | PINc domain-containing protein | 14.685 | 131 | 0.207 | 1.15 | 1.17E-01 | 1.48E-01 | no | no | 0 | 2 | 1 | 1 | 3 | 3 | 2 | 0.004385017 | 0.004090779 | 0.004717742 | 0.006314639 | 0.005947802 | 0.004439741 |
| thiE.1 | A0A0K2GG85 | Thiamine-phosphate synthase | 22.065 | 208 | 0.593 | 1.51 | 4.82E-03 | 9.65E-03 | no | no | 0 | 4 | 3 | 6 | 4 | 4 | 4 | 0.004381223 | 0.005635185 | 0.003850001 | 0.003027236 | 0.002811541 | 0.002897313 |
| NITMOv2_3852 | A0A0K2GH24 | HlyD_D23 domain-containing protein | 37.036 | 332 | 0.874 | 1.83 | 2.76E-05 | 2.33E-04 | no | no | 0 | 9 | 13 | 9 | 6 | 7 | 7 | 0.004381012 | 0.004306464 | 0.003902917 | 0.002632914 | 0.002714436 | 0.002590913 |
| rlp | A0A0K2G8H0 | Ribulose bisphosphate carboxylase-like protein (RuBisCO type IV-like protein) | 41.2 | 388 | 0.618 | 1.53 | 1.46E-04 | 7.22E-04 | no | no | 0 | 9 | 9 | 9 | 6 | 7 | 5 | 0.004380801 | 0.004397196 | 0.00429595 | 0.002724868 | 0.003200583 | 0.003336594 |
| tnaA | A0A0K2G6N4 | Tryptophanase | 52.717 | 467 | 3.66 | 12.64 | 1.10E-04 | 5.96E-04 | yes | yes | 3 | 11 | 9 | 10 | 0 | 0 | 0 | 0.0043378 | 0.005283792 | 0.004817055 | 5.644602E-06 | 0 | 0 |
| NITMOv2_0120 | A0A0K2G6K3 | TnsA endonuclease | 31.069 | 266 | 0.848 | 1.80 | 2.17E-03 | 5.13E-03 | no | no | 0 | 4 | 4 | 5 | 3 | 1 | 1 | 0.004318829 | 0.004585114 | 0.004136628 | 0.001805769 | 0.002133204 | 0.00187616 |
| PTS | A0A0K2GHM0 | 6-carboxy-5,6,7,8-tetrahydropterin synthase | 15.483 | 136 | 0.501 | 1.42 | 3.13E-02 | 4.68E-02 | no | no | 0 | 2 | 2 | 1 | 2 | 1 | 0 | 0.004309555 | 0.004679366 | 0.003564525 | 0.005333201 | 0.002641569 | 0.005026109 |
| NITMOv2_1064 | A0A0K2G981 | Uncharacterized protein | 27.09 | 242 | 0.908 | 1.88 | 7.09E-02 | 9.55E-02 | no | yes | 1 | 5 | 5 | 5 | 1 | 2 | 1 | 0.004301123 | 0.004474045 | 0.004288665 | 0.001802511 | 0.002614845 | 0.00194341 |

| | | | | | | | | | | | | | | | | | | | | | | | |
|---|---|---|---|---|---|---|---|---|---|---|---|---|---|---|---|---|---|---|---|---|---|---|---|
| NITMOv2_4581 | A0A0K2GJC6 | Uncharacterized protein | 22.654 | 209 | -0.00783 | -1.01 | 9.75E-01 | 9.77E-01 | no | no | 0 | 1 | 3 | 3 | 4 | 2 | 2 | 0.004299858 | 0.003598399 | 0.004576634 | 0.005424267 | 0.004448345 | 0.004391732 |
| NITMOv2_3622 | A0A0K2GGQ1 | B12-binding domain-containing protein | 67.759 | 593 | 0.228 | 1.17 | 7.55E-02 | 1.01E-01 | no | no | 0 | 11 | 13 | 11 | 14 | 16 | 15 | 0.004298172 | 0.004698529 | 0.004840062 | 0.004345248 | 0.004321719 | 0.004920379 |
| murG | A0A0K2G7X5 | UDP-N-acetylglucosamine--N-acetylmuramyl-(pentapeptide) pyrophosphoryl-undecaprenol N | 39.249 | 375 | 0.218 | 1.16 | 5.60E-02 | 7.73E-02 | no | no | 0 | 8 | 10 | 7 | 12 | 8 | 8 | 0.004293113 | 0.004454882 | 0.004135286 | 0.00571553 | 0.004940085 | 0.003249565 |
| mutL | A0A0K2GGW3 | DNA mismatch repair protein MutL | 64.263 | 593 | -0.526 | -1.44 | 6.28E-04 | 2.02E-03 | no | no | 0 | 16 | 17 | 18 | 24 | 24 | 17 | 0.004285314 | 0.004665091 | 0.004900263 | 0.009558212 | 0.009111718 | 0.008021239 |
| recN | A0A0K2GA97 | DNA repair protein RecN | 61.107 | 559 | -0.582 | -1.50 | 2.47E-04 | 1.02E-03 | no | no | 0 | 5 | 6 | 7 | 6 | 8 | 8 | 0.004274142 | 0.004195004 | 0.004048819 | 0.005196676 | 0.005510907 | 0.006758958 |
| NITMOv2_0700 | A0A0K2G837 | Uncharacterized protein | 28.189 | 240 | 0.849 | 1.80 | 6.42E-06 | 8.83E-05 | no | no | 0 | 10 | 9 | 8 | 6 | 4 | 4 | 0.004261074 | 0.00440756 | 0.004250895 | 0.002783653 | 0.002397019 | 0.002745551 |
| pyk | A0A0K2GJ54 | Pyruvate kinase | 52.285 | 496 | 0.318 | 1.25 | 6.48E-02 | 8.80E-02 | no | no | 0 | 14 | 20 | 20 | 8 | 8 | 9 | 0.004252431 | 0.005778714 | 0.005617884 | 0.004306156 | 0.004618939 | 0.003762389 |
| NITMOv2_3245 | A0A0K2GFA4 | PilZ domain-containing protein | 13.315 | 118 | 0.278 | 1.21 | 3.72E-02 | 5.44E-02 | no | no | 0 | 6 | 4 | 5 | 4 | 3 | 3 | 0.004228191 | 0.002777897 | 0.00423364 | 0.006280434 | 0.00251743 | 0.002682797 |
| NITMOv2_3872 | A0A0K2GHC9 | Glycosyl transferase, family 2 | 46.799 | 426 | 0.0452 | 1.03 | 6.85E-01 | 7.21E-01 | no | no | 0 | 11 | 9 | 11 | 12 | 14 | 10 | 0.004226504 | 0.004090974 | 0.003410763 | 0.004739274 | 0.004875451 | 0.004752794 |
| rtcB | A0A0K2GFB1 | RNA-splicing ligase RtcB | 52.575 | 483 | 0.465 | 1.38 | 2.77E-03 | 6.20E-03 | no | no | 0 | 8 | 10 | 11 | 7 | 5 | 4 | 0.004219127 | 0.003831683 | 0.004460066 | 0.002747819 | 0.003213168 | 0.00232587 |
| NITMOv2_1007 | A0A0K2G9B1 | Uncharacterized protein | 11.955 | 110 | 0.693 | 1.62 | 4.50E-03 | 9.10E-03 | no | no | 0 | 0 | 3 | 2 | 0 | 1 | 1 | 0.004212803 | 0.003589209 | 0.002899244 | 0.001169522 | 0.002012172 | 0.001209273 |
| cysD.1 | A0A0K2GET3 | ATP-sulfurylase small subunit | 30.667 | 265 | 0.593 | 1.51 | 1.44E-02 | 2.42E-02 | no | no | 0 | 4 | 6 | 8 | 0 | 1 | 4 | 0.004202475 | 0.003840483 | 0.004702596 | 0.002569093 | 0.002275211 | 0.002935253 |
| rmlC.1 | A0A0K2GIC6 | dTDP-4-dehydrorhamnose 3,5-epimerase | 21.409 | 190 | -0.316 | -1.24 | 6.34E-02 | 8.65E-02 | no | no | 0 | 9 | 9 | 13 | 8 | 9 | 8 | 0.004199524 | 0.011550584 | 0.012355051 | 0.007016364 | 0.0064007 | 0.005189019 |
| NITMOv2_4827 | A0A0K2GJR4 | Uncharacterized protein | 18.86 | 180 | 0.155 | 1.11 | 1.34E-01 | 1.68E-01 | no | no | 0 | 6 | 5 | 5 | 4 | 4 | 4 | 0.004196994 | 0.004142207 | 0.006510166 | 0.003999494 | 0.004478486 | 0.00386722 |
| NITMOv2_0412 | A0A0K2G7B5 | SLT domain-containing protein | 40.403 | 352 | -0.346 | -1.27 | 1.79E-02 | 2.89E-02 | no | no | 0 | 8 | 9 | 8 | 14 | 16 | 13 | 0.004195097 | 0.004104662 | 0.003453325 | 0.006209062 | 0.007104673 | 0.007377152 |
| hemC | A0A0K2GB73 | Porphobilinogen deaminase | 33.917 | 315 | -0.13 | -1.09 | 4.91E-02 | 6.92E-02 | no | no | 0 | 8 | 5 | 8 | 12 | 10 | 7 | 0.004194254 | 0.004552262 | 0.004306495 | 0.005456695 | 0.00550174 | 0.005383036 |
| NITMOv2_2984 | A0A0K2GEV7 | Transcriptional activator protein, LysR family | 33.651 | 299 | 0.967 | 1.95 | 1.59E-04 | 7.59E-04 | no | no | 0 | 7 | 7 | 9 | 3 | 3 | 5 | 0.004182661 | 0.004459184 | 0.004199705 | 0.001976499 | 0.001962765 | 0.002003287 |
| ureG | A0A0K2G9S2 | Urease accessory protein UreG | 24.392 | 223 | -0.112 | -1.08 | 2.85E-01 | 3.28E-01 | no | no | 0 | 2 | 2 | 5 | 3 | 4 | 4 | 0.004164322 | 0.004559693 | 0.004170947 | 0.00539406 | 0.006064484 | 0.005769272 |
| NITMOv2_2506 | A0A0K2GDK2 | Uncharacterized protein | 25.607 | 230 | 2.71 | 6.54 | 3.11E-04 | 1.19E-03 | yes | yes | 3 | 4 | 3 | 4 | 1 | 1 | 2 | 0.004162636 | 0.004376077 | 0.004222328 | 0.000800832 | 0.002193642 | 0.000651046 |
| NITMOv2_0344 | A0A0K2G849 | Uncharacterized protein | 68.433 | 615 | -0.0618 | -1.04 | 5.43E-01 | 5.84E-01 | no | no | 0 | 12 | 12 | 12 | 15 | 16 | 12 | 0.004155047 | 0.004271657 | 0.004038082 | 0.004569433 | 0.004611947 | 0.003674641 |
| NITMOv2_2062 | A0A0K2GCA3 | Uncharacterized protein | 13.916 | 120 | 1.43 | 2.69 | 5.58E-03 | 1.08E-02 | yes | yes | 2 | 1 | 1 | 1 | 1 | 1 | 1 | 0.004150199 | 0.004594891 | 0.004081412 | 0.000878156 | 0.002486045 | 0.003941842 |
| NITMOv2_0305 | A0A0K2G712 | N-acetyltransferase, GCN5-related | 16.585 | 143 | -0.49 | -1.40 | 7.36E-02 | 9.87E-02 | no | no | 0 | 1 | 1 | 1 | 6 | 5 | 6 | 0.004149356 | 0.004734314 | 0.003794018 | 0.005504227 | 0.004374079 | 0.005179668 |
| putA | A0A0K2GCA8 | Bifunctional protein PutA | 109.41 | 999 | 0.695 | 1.62 | 1.97E-03 | 4.75E-03 | no | no | 0 | 21 | 27 | 23 | 21 | 15 | 13 | 0.004148091 | 0.004944719 | 0.004067608 | 0.002844808 | 0.00262308 | 0.002109556 |
| tatA.1 | A0A0K2GIN4 | Sec-independent protein translocase protein TatA | 10.536 | 96 | -0.338 | -1.26 | 1.11E-02 | 1.94E-02 | no | no | 0 | 2 | 1 | 4 | 5 | 4 | 4 | 0.004133547 | 0.004039155 | 0.004365738 | 0.006632999 | 0.006200586 | 0.007305228 |
| NITMOv2_0128 | A0A0K2G6J9 | AAA_11 domain-containing protein | 53.897 | 485 | -0.335 | -1.26 | 1.41E-02 | 2.38E-02 | no | no | 0 | 7 | 9 | 11 | 15 | 16 | 15 | 0.004133125 | 0.004050301 | 0.0040701 | 0.006198105 | 0.005977633 | 0.005305357 |
| NITMOv2_4560 | A0A0K2GJA6 | Putative Peptidase, family M48 | 41.61 | 377 | -0.102 | -1.07 | 3.19E-01 | 3.61E-01 | no | no | 0 | 4 | 4 | 3 | 5 | 4 | 3 | 0.004125748 | 0.00383403 | 0.004401398 | 0.005817997 | 0.00609043 | 0.005294388 |
| NITMOv2_3709 | A0A0K2GGN3 | Putative Hydrolase, beta-lactamase-like | 23.167 | 213 | -0.189 | -1.14 | 4.45E-02 | 6.35E-02 | no | no | 0 | 10 | 11 | 10 | 6 | 9 | 6 | 0.004113733 | 0.004384486 | 0.004734422 | 0.005195788 | 0.004615521 | 0.004451969 |
| rlpA | A0A0K2GDM7 | Probable endolytic peptidoglycan transglycosylase RlpA | 24.548 | 224 | -0.0296 | -1.02 | 7.39E-01 | 7.70E-01 | no | no | 0 | 4 | 4 | 4 | 4 | 6 | 3 | 0.004106355 | 0.003770673 | 0.003652143 | 0.003367511 | 0.002853957 | 0.00262256 |
| tatC | A0A0K2GGI3 | Sec-independent protein translocase protein TatC | 34.474 | 311 | -0.522 | -1.44 | 1.63E-03 | 4.11E-03 | no | no | 0 | 1 | 1 | 1 | 3 | 2 | 2 | 0.004100664 | 0.004151593 | 0.003539409 | 0.006351066 | 0.006577975 | 0.005836702 |
| yrbE | A0A0K2G968 | Toluene transporter subunit: membrane component of ABC superfamily | 27.687 | 256 | -0.511 | -1.43 | 5.27E-02 | 7.37E-02 | no | no | 0 | 11 | 12 | 14 | 11 | 12 | 13 | 0.004100664 | 0.004553827 | 0.005440732 | 0.005790011 | 0.005169253 | 0.005003273 |
| NITMOv2_0454 | A0A0K2G7H5 | Uncharacterized protein | 46.959 | 419 | 0.585 | 1.50 | 1.13E-04 | 6.07E-04 | no | no | 0 | 5 | 4 | 5 | 5 | 4 | 5 | 0.004097713 | 0.004550111 | 0.004596381 | 0.002895598 | 0.00287229 | 0.002549736 |
| nuoF | A0A0K2G7R9 | NADH-quinone oxidoreductase subunit F | 47.363 | 435 | -0.588 | -1.50 | 3.01E-03 | 6.63E-03 | no | no | 0 | 4 | 4 | 4 | 9 | 7 | 7 | 0.004091811 | 0.00459978 | 0.003822777 | 0.007912067 | 0.007159207 | 0.006555411 |
| NITMOv2_1868 | A0A0K2GBG1 | Uncharacterized protein | 9.9482 | 92 | 0.145 | 1.11 | 3.21E-01 | 3.63E-01 | no | no | 0 | 2 | 3 | 4 | 3 | 1 | 2 | 0.00408612 | 0.004702636 | 0.004068374 | 0.003313464 | 0.003132532 | 0.002934714 |
| nuoM.1 | A0A0K2G7K9 | NADH-quinone oxidoreductase, membrane subunit M | 62.827 | 568 | -0.264 | -1.20 | 5.81E-02 | 7.99E-02 | no | no | 0 | 3 | 5 | 2 | 5 | 3 | 2 | 0.004085066 | 0.006865743 | 0.003853452 | 0.005261533 | 0.005622151 | 0.004964793 |
| NITMOv2_1083 | A0A0K2G9I1 | Peptidase_S9 domain-containing protein | 27.96 | 254 | 0.69 | 1.61 | 1.59E-02 | 2.62E-02 | no | no | 0 | 3 | 3 | 2 | 2 | 1 | 0 | 0.00408359 | 0.003986163 | 0.004160977 | 0.002250733 | 0.002297739 | 0.001672487 |
| NITMOv2_0588 | A0A0K2G843 | Putative FMN reductase | 26.718 | 236 | 1.15 | 2.22 | 8.82E-04 | 2.58E-03 | yes | no | 0 | 9 | 8 | 9 | 4 | 4 | 1 | 0.004067992 | 0.004890358 | 0.005129756 | 0.000918033 | 0.001147953 | 0.001393112 |
| NITMOv2_4074 | A0A0K2GHM6 | Histidine kinase | 114.26 | 1034 | 0.282 | 1.22 | 1.30E-02 | 2.22E-02 | no | no | 0 | 21 | 29 | 27 | 26 | 24 | 17 | 0.004057031 | 0.004077677 | 0.00416577 | 0.00449273 | 0.004116944 | 0.004024555 |
| NITMOv2_2495 | A0A0K2GD95 | Putative Trehalose-phosphatase | 29.675 | 272 | -0.419 | -1.34 | 5.64E-04 | 1.86E-03 | no | no | 0 | 2 | 4 | 4 | 9 | 9 | 6 | 0.00404607 | 0.004041502 | 0.004172097 | 0.004929254 | 0.005703097 | 0.00548427 |
| NITMOv2_2259 | A0A0K2GCJ2 | Uncharacterized protein | 8.6527 | 74 | 0.953 | 1.94 | 1.64E-01 | 2.00E-01 | no | yes | 3 | 0 | 1 | 1 | 0 | 0 | 2 | 0.004035741 | 0.006661595 | 0.003764301 | 0.003171461 | 0.003680826 | 0.00239276 |
| NITMOv2_4304 | A0A0K2GIC0 | Transcriptional regulator, AbrB family | 7.6691 | 69 | 0.457 | 1.37 | 5.18E-01 | 5.61E-01 | no | no | 0 | 2 | 3 | 1 | 2 | 1 | 1 | 0.004025624 | 0.007667277 | 0.003118959 | 0.00732436 | 0.002178882 | 0.00231616 |
| NITMOv2_3375 | A0A0K2GGM0 | PMT_2 domain-containing protein | 61.02 | 550 | 0.157 | 1.11 | 1.29E-01 | 1.61E-01 | no | no | 0 | 6 | 9 | 8 | 5 | 4 | 5 | 0.004023516 | 0.002873713 | 0.003624534 | 0.003349002 | 0.004172877 | 0.00296672 |
| xthA | A0A0K2GJN9 | Exodeoxyribonuclease III | 29.751 | 257 | -0.286 | -1.22 | 3.19E-03 | 6.96E-03 | no | no | 0 | 13 | 11 | 9 | 9 | 8 | 9 | 0.004001594 | 0.004109355 | 0.003979798 | 0.004871949 | 0.004633854 | 0.004721507 |
| purF | A0A0K2GAX0 | Amidophosphoribosyltransferase | 52.217 | 476 | -1.05 | -2.07 | 2.97E-05 | 2.44E-04 | yes | no | 0 | 6 | 7 | 4 | 15 | 15 | 15 | 0.004001172 | 0.00430529 | 0.004290966 | 0.008047111 | 0.007636032 | 0.010191212 |
| vapC.2 | A0A0K2GE12 | Ribonuclease VapC | 15.901 | 139 | 0.0561 | 1.04 | 6.68E-01 | 7.05E-01 | no | no | 0 | 2 | 1 | 3 | 2 | 1 | 2 | 0.003999908 | 0.004329538 | 0.003962926 | 0.004159414 | 0.004025899 | 0.003845462 |
| ureA | A0A0K2G9N5 | Urease subunit gamma | 11.083 | 100 | 0.324 | 1.25 | 6.69E-03 | 1.27E-02 | no | no | 0 | 2 | 4 | 2 | 1 | 0 | 0 | 0.003993162 | 0.004701854 | 0.004456615 | 0.0040528 | 0.003456941 | 0.003074428 |
| adh | A0A0K2GJZ1 | Alcohol dehydrogenase | 35.03 | 329 | -0.0485 | -1.03 | 6.64E-01 | 7.02E-01 | no | no | 0 | 6 | 7 | 8 | 6 | 5 | 7 | 0.003992952 | 0.004465832 | 0.004557078 | 0.004738978 | 0.004939929 | 0.005065308 |
| obg | A0A0K2GA07 | GTPase Obg | 37.081 | 341 | 0.297 | 1.23 | 4.21E-03 | 8.66E-03 | no | no | 0 | 10 | 10 | 9 | 10 | 9 | 9 | 0.003991476 | 0.004627547 | 0.004451438 | 0.003594658 | 0.003324101 | 0.003870816 |
| NITMOv2_3912 | A0A0K2GH57 | Response regulatory domain-containing protein | 16.663 | 152 | -2.7 | -6.50 | 3.74E-06 | 6.29E-05 | yes | no | 0 | 7 | 7 | 11 | 33 | 31 | 33 | 0.003990001 | 0.003351232 | 0.004772383 | 0.033512232 | 0.026499586 | 0.024494009 |
| NITMOv2_4337 | A0A0K2GIC2 | DUF4440 domain-containing protein | 16.976 | 155 | 0.524 | 1.44 | 1.29E-03 | 3.42E-03 | no | no | 0 | 1 | 2 | 3 | 1 | 0 | 1 | 0.003989579 | 0.004837952 | 0.006529914 | 0.003029901 | 0.003904711 | 0.002489679 |
| NITMOv2_0365 | A0A0K2G745 | Uncharacterized protein | 45.66 | 423 | -0.543 | -1.46 | 3.90E-03 | 8.14E-03 | no | no | 0 | 9 | 8 | 4 | 11 | 9 | 9 | 0.003982201 | 0.004638497 | 0.003727106 | 0.009146713 | 0.005110834 | 0.005019815 |
| NITMOv2_1853 | A0A0K2GBE7 | HD-GYP domain-containing protein | 75.424 | 693 | 0.889 | 1.85 | 1.19E-05 | 1.34E-04 | no | no | 0 | 17 | 19 | 13 | 8 | 9 | 8 | 0.003979883 | 0.004226877 | 0.004078344 | 0.002208828 | 0.002245691 | 0.002190651 |
| rnd | A0A0K2GC38 | Ribonuclease D | 44.235 | 396 | -0.298 | -1.23 | 5.87E-02 | 8.06E-02 | no | no | 0 | 12 | 12 | 16 | 22 | 19 | 19 | 0.003978829 | 0.004919689 | 0.005682879 | 0.008597949 | 0.00875142 | 0.006263756 |
| NITMOv2_3176 | A0A0K2GF38 | Uncharacterized protein | 53.275 | 481 | 0.125 | 1.09 | 2.37E-01 | 2.78E-01 | no | no | 0 | 6 | 6 | 7 | 4 | 6 | 6 | 0.003964284 | 0.003186584 | 0.004451246 | 0.001992343 | 0.002309702 | 0.002452817 |
| hemH | A0A0K2GAC0 | Ferrochelatase | 34.4 | 314 | -1.09 | -2.13 | 1.75E-03 | 4.34E-03 | yes | no | 0 | 3 | 2 | 5 | 9 | 9 | 8 | 0.003955853 | 0.003035428 | 0.003219614 | 0.005604029 | 0.005142529 | 0.005193873 |
| prfB | A0A0K2GCL6 | Peptide chain release factor 2 | 42.295 | 381 | -0.236 | -1.18 | 1.50E-02 | 2.52E-02 | no | no | 0 | 3 | 4 | 4 | 4 | 4 | 5 | 0.003953745 | 0.004829153 | 0.004296142 | 0.004230786 | 0.004412299 | 0.003695859 |
| glnEb | A0A0K2G9U0 | (Glutamate-ammonia-ligase) adenylyltransferase, subunit B | 64.663 | 574 | 0.347 | 1.27 | 2.90E-03 | 6.44E-03 | no | no | 0 | 10 | 11 | 14 | 10 | 13 | 8 | 0.003951848 | 0.004384486 | 0.004622839 | 0.003192487 | 0.003598947 | 0.00348386 |
| NITMOv2_3722 | A0A0K2GGY9 | Uncharacterized protein | 11.409 | 101 | 1.34 | 2.53 | 4.09E-02 | 5.89E-02 | no | no | 0 | 2 | 4 | 6 | 4 | 3 | 3 | 0.003939201 | 0.010757653 | 0.003578712 | 0.002230003 | 0.002003783 | 0.001398902 |
| NITMOv2_4127 | A0A0K2GI30 | Uncharacterized protein | 36.529 | 348 | 0.556 | 1.47 | 2.18E-04 | 9.37E-04 | no | no | 0 | 4 | 5 | 4 | 2 | 3 | 3 | 0.003925921 | 0.003605634 | 0.003631628 | 0.002706655 | 0.002757629 | 0.002617166 |
| nth.1 | A0A0K2GJT8 | Endonuclease III | 24.884 | 219 | -0.492 | -1.41 | 8.15E-04 | 2.43E-03 | no | no | 0 | 7 | 8 | 5 | 6 | 8 | 7 | 0.003908847 | 0.004220033 | 0.00358619 | 0.004852403 | 0.005188518 | 0.004066991 |
| NITMOv2_1672 | A0A0K2GJD0 | Putative Cytochrome P460 | 22.655 | 205 | -0.573 | -1.49 | 1.37E-02 | 2.32E-02 | no | no | 0 | 6 | 6 | 5 | 5 | 5 | 5 | 0.003904842 | 0.004930835 | 0.005790436 | 0.007078556 | 0.006218298 | 0.007089093 |
| NITMOv2_4528 | A0A0K2GJX4 | Putative Uroporphyrinogen III synthase HemD | 30.64 | 279 | 0.137 | 1.10 | 3.74E-01 | 4.17E-01 | no | no | 0 | 4 | 3 | 6 | 5 | 5 | 5 | 0.003899151 | 0.0036291 | 0.00420469 | 0.0032415 | 0.003010568 | 0.00272811 |
| NITMOv2_0520 | A0A0K2G7L5 | Putative Cytochrome P450 | 57.742 | 515 | 1.39 | 2.62 | 7.32E-06 | 9.68E-05 | yes | no | 0 | 10 | 11 | 16 | 8 | 4 | 4 | 0.003894092 | 0.003934735 | 0.005219675 | 0.001490518 | 0.001582114 | 0.001156588 |
| NITMOv2_2701 | A0A0K2GER9 | Putative Histidine kinase family | 73.83 | 685 | -0.761 | -1.69 | 1.40E-03 | 3.67E-03 | no | no | 0 | 14 | 14 | 18 | 21 | 22 | 19 | 0.003858469 | 0.003200467 | 0.004290199 | 0.00545092 | 0.005374028 | 0.004934584 |
| NITMOv2_1004 | A0A0K2G8Z6 | Uncharacterized protein | 115.96 | 1050 | 0.338 | 1.26 | 3.24E-03 | 7.03E-03 | no | no | 0 | 18 | 16 | 18 | 20 | 13 | 14 | 0.003842238 | 0.003724329 | 0.00368416 | 0.003482565 | 0.003022997 | 0.00332095 |
| pncA | A0A0K2GGY0 | Nicotine deamidase | 20.956 | 193 | 1.33 | 2.51 | 2.09E-04 | 9.09E-04 | yes | yes | 1 | 5 | 6 | 4 | 0 | 0 | 0 | 0.003830223 | 0.003284551 | 0.00341824 | 0.000302427 | 0.000622078 | 0.000717972 |
| NITMOv2_1927 | A0A0K2GBK0 | Sigma subunit DNA-directed RNA polymerase | 22.67 | 202 | 0.12 | 1.09 | 4.68E-01 | 5.12E-01 | no | no | 0 | 3 | 2 | 1 | 1 | 2 | 0 | 0.003812096 | 0.002763818 | 0.001984339 | 0.004067312 | 0.003062772 | 0.003922422 |
| NITMOv2_1929 | A0A0K2GBM1 | Putative chemiosmotic efflux system B protein C | 49.272 | 446 | 1.02 | 2.03 | 1.07E-02 | 1.89E-02 | yes | no | 0 | 7 | 9 | 7 | 0 | 0 | 0 | 0.003811674 | 0.003225497 | 0.002516947 | 0.001032391 | 0.000585551 | 0.000826201 |
| NITMOv2_0261 | A0A0K2G6Z1 | Uncharacterized protein | 47.284 | 425 | 0.295 | 1.23 | 1.33E-02 | 2.27E-02 | no | no | 0 | 11 | 12 | 12 | 9 | 8 | 5 | 0.00381062 | 0.003597421 | 0.003848851 | 0.002937503 | 0.002628829 | 0.002660321 |
| NITMOv2_1266 | A0A0K2G9Z6 | Uncharacterized protein | 35.268 | 322 | -0.0896 | -1.06 | 4.82E-01 | 5.25E-01 | no | no | 0 | 3 | 3 | 3 | 5 | 2 | 5 | 0.00380788 | 0.004245258 | 0.004674029 | 0.005216666 | 0.004112905 | 0.005089942 |
| NITMOv2_1099 | A0A0K2G9B5 | CxxC_CXXC_SSSS domain-containing protein | 11.247 | 101 | -0.72 | -1.65 | 7.86E-04 | 2.37E-03 | no | no | 0 | 2 | 3 | 4 | 7 | 8 | 6 | 0.003803664 | 0.005231386 | 0.005578389 | 0.011682342 | 0.010397856 | 0.008538199 |
| NITMOv2_0779 | A0A0K2G8F2 | Uncharacterized protein | 56.172 | 498 | 0.0222 | 1.02 | 8.04E-01 | 8.28E-01 | no | no | 0 | 9 | 8 | 7 | 12 | 13 | 9 | 0.003797551 | 0.003622647 | 0.003640064 | 0.004190362 | 0.004791553 | 0.004502136 |
| prfA | A0A0K2G8F6 | Peptide chain release factor 1 | 39.975 | 359 | -0.616 | -1.53 | 1.00E-04 | 5.60E-04 | no | no | 0 | 5 | 4 | 3 | 10 | 12 | 9 | 0.003776894 | 0.003872943 | 0.003220764 | 0.006928704 | 0.007740906 | 0.007533769 |
| NITMOv2_3673 | A0A0K2GGH9 | ABC-type transport system, ATP-binding component | 27.109 | 243 | -0.474 | -1.39 | 1.46E-03 | 3.77E-03 | no | no | 0 | 3 | 3 | 2 | 3 | 4 | 4 | 0.003772257 | 0.003826599 | 0.003654443 | 0.004363609 | 0.005556585 | 0.005376023 |
| NITMOv2_3466 | A0A0K2GFX5 | Uncharacterized protein | 34.571 | 315 | 0.451 | 1.37 | 1.95E-03 | 4.70E-03 | no | no | 0 | 4 | 4 | 4 | 2 | 2 | 4 | 0.003749913 | 0.003878418 | 0.004282721 | 0.004332957 | 0.003319751 | 0.005190996 |
| NITMOv2_0142 | A0A0K2G7J6 | Uncharacterized protein | 19.86 | 177 | -0.0316 | -1.02 | 7.56E-01 | 7.85E-01 | no | no | 0 | 4 | 3 | 3 | 3 | 4 | 2 | 0.003748438 | 0.003497107 | 0.005360016 | 0.003785674 | 0.005285157 | 0.003203713 |
| NITMOv2_0792 | A0A0K2G9E2 | Putative transmembrane signal transduction receptor and sigma-54 dependent response regu | 92.237 | 846 | 0.194 | 1.14 | 6.12E-02 | 8.38E-02 | no | no | 0 | 12 | 11 | 12 | 9 | 8 | 7 | 0.003743379 | 0.003943925 | 0.003719246 | 0.003763315 | 0.003379878 | 0.002875196 |
| NITMOv2_3338 | A0A0K2GFK4 | MazG nucleotide pyrophosphohydrolase | 13.433 | 115 | 0.838 | 1.79 | 2.04E-01 | 2.43E-01 | no | yes | 1 | 1 | 0 | 1 | 3 | 3 | 1 | 0.003729678 | 0.003757376 | 0.003602103 | 0.003788635 | 0.004025588 | 0.001663676 |
| NITMOv2_0524 | A0A0K2G7Q7 | Putative lactoylglutathione lyase | 15.64 | 142 | 0.147 | 1.11 | 2.89E-01 | 3.32E-01 | no | no | 0 | 4 | 2 | 4 | 5 | 5 | 5 | 0.00372757 | 0.00417936 | 0.005110584 | 0.007387439 | 0.007187174 | 0.007648489 |
| NITMOv2_0797 | A0A0K2G9F1 | Uncharacterized protein | 33.542 | 291 | -0.456 | -1.37 | 1.58E-03 | 4.04E-03 | no | no | 0 | 3 | 2 | 4 | 5 | 4 | 3 | 0.003726727 | 0.003575521 | 0.004293458 | 0.004763262 | 0.005131187 | 0.005387891 |
| NITMOv2_1093 | A0A0K2G9J1 | Putative thiol peroxidase (Modular protein) | 23.754 | 220 | -1.3 | -2.46 | 7.88E-03 | 1.45E-02 | yes | no | 0 | 1 | 1 | 2 | 5 | 5 | 5 | 0.003725884 | 0.003054396 | 0.004138737 | 0.006137246 | 0.00579927 | 0.006850483 |

| | | | | | | | | | | | | | | | | | | | | | | | |
|---|---|---|---|---|---|---|---|---|---|---|---|---|---|---|---|---|---|---|---|---|---|---|---|
| NITMOv2_1165 | A0A0K2G9E7 | Uncharacterized protein | 16.39 | 145 | 0.443 | 1.36 | 1.11E-01 | 1.41E-01 | no | no | 0 | 3 | 3 | 6 | 6 | 7 | 7 | 0.003718084 | 0.007224175 | 0.007790304 | 0.007697063 | 0.008564667 | 0.009329193 |
| NITMOv2_0145 | A0A0K2G6M9 | Uncharacterized protein | 37.546 | 326 | -0.573 | -1.49 | 4.14E-03 | 8.55E-03 | no | no | 0 | 7 | 5 | 6 | 10 | 7 | 8 | 0.00371682 | 0.005055397 | 0.003872625 | 0.005145887 | 0.003879697 | 0.004416546 |
| shc | A0A0K2G895 | Squalene--hopene cyclase | 81.818 | 725 | -0.663 | -1.58 | 1.51E-04 | 7.40E-04 | no | no | 0 | 13 | 14 | 14 | 21 | 21 | 21 | 0.003709653 | 0.004190506 | 0.003813766 | 0.009140346 | 0.00813119 | 0.008807198 |
| NITMOv2_3972 | A0A0K2GHE4 | Response regulator, LuxR family | 23.531 | 210 | -0.142 | -1.10 | 2.72E-01 | 3.15E-01 | no | no | 0 | 6 | 6 | 5 | 6 | 6 | 5 | 0.003680564 | 0.004180142 | 0.004732121 | 0.004784881 | 0.004455181 | 0.004466174 |
| rsmA | A0A0K2G7J8 | Ribosomal RNA small subunit methyltransferase A | 29.13 | 265 | 0.847 | 1.80 | 2.35E-03 | 5.44E-03 | no | no | 0 | 6 | 3 | 4 | 1 | 1 | 0 | 0.003680353 | 0.003720223 | 0.003714069 | 0.002948905 | 0.002808745 | 0.002436994 |
| ogt | A0A0K2GK77 | Methylated-DNA--protein-cysteine methyltransferase | 19.288 | 175 | -1.14 | -2.20 | 5.39E-04 | 1.80E-03 | yes | no | 0 | 1 | 1 | 1 | 4 | 4 | 2 | 0.003679089 | 0.002844577 | 0.002692758 | 0.006930037 | 0.006551873 | 0.006698362 |
| NITMOv2_2011 | A0A0K2GBV8 | Phosphohydrolase | 17.899 | 159 | 0.801 | 1.74 | 2.84E-02 | 4.30E-02 | no | yes | 1 | 2 | 2 | 1 | 0 | 0 | 0 | 0.003678878 | 0.003318185 | 0.003100746 | 0.001410543 | 0.00177881 | 0.001171116 |
| NITMOv2_3415 | A0A0K2GGR2 | SLBB domain-containing protein | 23.436 | 215 | -1.46 | -2.75 | 2.51E-06 | 4.86E-05 | yes | no | 0 | 6 | 7 | 5 | 9 | 12 | 10 | 0.003667074 | 0.003781037 | 0.00293548 | 0.007295041 | 0.007627642 | 0.007600839 |
| NITMOv2_3977 | A0A0K2GHE9 | RND-type efflux transporter, membrane-fusion protein | 37.878 | 354 | 1.03 | 2.04 | 3.96E-04 | 1.41E-03 | yes | no | 0 | 4 | 4 | 4 | 0 | 2 | 0 | 0.003665177 | 0.00410564 | 0.003009102 | 0.001479502 | 0.002588122 | 0.002082764 |
| NITMOv2_3464 | A0A0K2GFY4 | Putative N-acetylmuramoyl-L-alanine amidase AmiB | 45.905 | 423 | -0.769 | -1.70 | 5.60E-04 | 1.85E-03 | no | no | 0 | 9 | 13 | 18 | 19 | 21 | 22 | 0.003664755 | 0.003387212 | 0.004043067 | 0.005093765 | 0.006219541 | 0.005805594 |
| NITMOv2_4488 | A0A0K2GJR2 | Putative 2Fe-2S ferredoxin | 11.22 | 105 | -0.548 | -1.46 | 3.17E-02 | 4.74E-02 | no | no | 0 | 1 | 2 | 2 | 3 | 4 | 4 | 0.003645573 | 0.003900906 | 0.003638722 | 0.009846513 | 0.015450893 | 0.012966613 |
| hemA | A0A0K2GC53 | Glutamyl-tRNA reductase | 50.956 | 460 | -0.799 | -1.74 | 1.12E-05 | 1.28E-04 | no | no | 0 | 11 | 7 | 7 | 15 | 11 | 13 | 0.003630607 | 0.003522528 | 0.003479783 | 0.005646379 | 0.00549941 | 0.005423314 |
| NITMOv2_0585 | A0A0K2G7S2 | Uncharacterized protein | 39.662 | 353 | -0.0146 | -1.01 | 9.11E-01 | 9.24E-01 | no | no | 0 | 7 | 6 | 9 | 8 | 11 | 8 | 0.003615642 | 0.005455675 | 0.005393568 | 0.00524791 | 0.005060184 | 0.003440166 |
| NITMOv2_4594 | A0A0K2GJ56 | Uncharacterized protein | 17.317 | 158 | 0.322 | 1.25 | 3.69E-02 | 5.40E-02 | no | no | 0 | 2 | 2 | 3 | 4 | 3 | 2 | 0.00360194 | 0.003167811 | 0.002954269 | 0.002980148 | 0.00305143 | 0.002569875 |
| potA | A0A0K2G9S3 | Spermidine/putrescine import ATP-binding protein PotA | 41.072 | 374 | -0.344 | -1.27 | 6.96E-03 | 1.31E-02 | no | no | 0 | 8 | 10 | 13 | 14 | 14 | 15 | 0.003581283 | 0.003631446 | 0.004282721 | 0.004793469 | 0.004634786 | 0.004482896 |
| NITMOv2_0667 | A0A0K2G820 | Uncharacterized protein | 37.606 | 348 | -0.552 | -1.47 | 1.81E-04 | 8.15E-04 | no | no | 0 | 6 | 7 | 7 | 11 | 10 | 11 | 0.003578965 | 0.004104467 | 0.004112279 | 0.008258858 | 0.009882344 | 0.008742106 |
| NITMOv2_3243 | A0A0K2GFA2 | TPR_REGION domain-containing protein | 63.718 | 565 | 0.118 | 1.09 | 1.52E-01 | 1.87E-01 | no | no | 0 | 9 | 8 | 12 | 16 | 14 | 11 | 0.003578543 | 0.003632619 | 0.004670003 | 0.003406455 | 0.00395676 | 0.004208683 |
| NITMOv2_2237 | A0A0K2GCH9 | AAA domain-containing protein | 55.161 | 509 | -0.801 | -1.74 | 3.06E-05 | 2.49E-04 | no | no | 0 | 4 | 4 | 6 | 16 | 12 | 9 | 0.003569901 | 0.003677008 | 0.004703555 | 0.010017539 | 0.008985404 | 0.007807443 |
| NITMOv2_3594 | A0A0K2GGC1 | UPF0301 protein NITMOv2_3594 | 20.321 | 186 | 0.101 | 1.07 | 1.40E-01 | 1.74E-01 | no | no | 0 | 3 | 2 | 2 | 1 | 1 | 1 | 0.003563999 | 0.003711424 | 0.003482467 | 0.003351223 | 0.003641829 | 0.003217558 |
| ylyB | A0A0K2G730 | Pseudouridine synthase | 35.196 | 320 | 0.0738 | 1.05 | 4.88E-01 | 5.31E-01 | no | no | 0 | 7 | 6 | 3 | 3 | 0 | 1 | 0.003560837 | 0.004021556 | 0.003050514 | 0.003334047 | 0.002905694 | 0.003223492 |
| NITMOv2_0013 | A0A0K2G7A1 | Putative Glyoxalase | 16.639 | 143 | -0.326 | -1.25 | 1.70E-01 | 2.07E-01 | no | no | 0 | 4 | 7 | 6 | 3 | 4 | 3 | 0.003539336 | 0.004959776 | 0.004588329 | 0.00251786 | 0.002090167 | 0.002069638 |
| NITMOv2_3182 | A0A0K2GF51 | Uncharacterized protein | 30.661 | 298 | 2 | 4.00 | 3.54E-03 | 7.57E-03 | yes | yes | 3 | 2 | 2 | 1 | 1 | 1 | 1 | 0.003529429 | 0.003905403 | 0.003675341 | 0.001732028 | 0.00158693 | 0.001460938 |
| pyrB | A0A0K2GFI1 | Aspartate carbamoyltransferase | 33.097 | 306 | -1.13 | -2.19 | 2.14E-05 | 1.98E-04 | yes | no | 0 | 10 | 7 | 10 | 17 | 19 | 14 | 0.003527954 | 0.003963871 | 0.003287676 | 0.010251053 | 0.010324211 | 0.010226815 |
| iaaA | A0A0K2GJH6 | Isoaspartyl peptidase | 30.798 | 293 | 1.29 | 2.45 | 8.26E-02 | 1.09E-01 | no | yes | 1 | 0 | 1 | 1 | 1 | 0 | 2 | 0.003526268 | 0.003854757 | 0.003632778 | 0.001935038 | 0.001368001 | 0.001876879 |
| NITMOv2_4818 | A0A0K2GJS0 | Uncharacterized protein | 13.794 | 122 | -0.827 | -1.77 | 8.78E-02 | 1.15E-01 | no | no | 0 | 1 | 2 | 3 | 3 | 5 | 5 | 0.003520998 | 0.004081784 | 0.00464968 | 0.002522302 | 0.003660473 | 0.003308363 |
| pepM | A0A0K2G853 | Phosphoenolpyruvate mutase | 61.752 | 559 | -2.44 | -5.43 | 2.84E-07 | 1.39E-05 | yes | no | 0 | 12 | 12 | 12 | 44 | 41 | 43 | 0.00350561 | 0.003353187 | 0.003256042 | 0.023025593 | 0.022839113 | 0.021683545 |
| NITMOv2_0153 | A0A0K2G6M1 | Putative Hybrid histidine kinase | 58.733 | 544 | -0.9 | -1.87 | 2.19E-03 | 5.15E-03 | no | no | 0 | 17 | 17 | 23 | 25 | 25 | 23 | 0.003504135 | 0.003570436 | 0.004330269 | 0.006091343 | 0.005711798 | 0.005587482 |
| NITMOv2_0800 | A0A0K2G8G9 | Putative ATPase, AAA family | 63.883 | 582 | -1.55 | -2.93 | 5.58E-06 | 7.92E-05 | yes | no | 0 | 7 | 9 | 8 | 31 | 37 | 32 | 0.003496968 | 0.003977559 | 0.003504707 | 0.016428871 | 0.017390353 | 0.014774983 |
| ychF | A0A0K2G823 | Ribosome-binding ATPase YchF | 39.66 | 363 | -0.679 | -1.60 | 3.00E-05 | 2.46E-04 | no | no | 0 | 7 | 8 | 8 | 11 | 12 | 12 | 0.003488958 | 0.00299847 | 0.00367074 | 0.0064522 | 0.006767214 | 0.00651765 |
| NITMOv2_1030 | A0A0K2G918 | J domain-containing protein | 24.524 | 212 | -0.136 | -1.10 | 3.42E-01 | 3.84E-01 | no | no | 0 | 2 | 1 | 4 | 8 | 2 | 2 | 0.003481791 | 0.003339695 | 0.002994339 | 0.003796187 | 0.003276247 | 0.003288224 |
| NITMOv2_4660 | A0A0K2GJB3 | Putative Exopolyphosphatase | 57.491 | 517 | 1.07 | 2.10 | 4.77E-02 | 6.74E-02 | no | no | 0 | 3 | 2 | 3 | 1 | 1 | 2 | 0.003476943 | 0.003413024 | 0.003497038 | 0.000734583 | 0.002344349 | 0.002350864 |
| NITMOv2_1432 | A0A0K2GAE9 | Uncharacterized protein | 37.639 | 337 | -0.411 | -1.33 | 4.34E-03 | 8.86E-03 | no | no | 0 | 6 | 6 | 5 | 9 | 9 | 7 | 0.003475257 | 0.003862383 | 0.003753948 | 0.008242422 | 0.007466215 | 0.006797978 |
| NITMOv2_1428 | A0A0K2GA83 | Putative beta-phosphoglucomutase | 24.672 | 228 | -0.565 | -1.48 | 6.99E-03 | 1.31E-02 | no | no | 0 | 2 | 4 | 6 | 9 | 9 | 8 | 0.003457972 | 0.003753857 | 0.003805905 | 0.006640551 | 0.007427683 | 0.005296546 |
| NITMOv2_3488 | A0A0K2GFZ4 | Epimerase domain-containing protein | 41.594 | 368 | -1.19 | -2.28 | 2.49E-04 | 1.02E-03 | yes | no | 0 | 2 | 3 | 2 | 7 | 6 | 6 | 0.003456708 | 0.002092515 | 0.002109535 | 0.007353826 | 0.007730496 | 0.007270164 |
| ywlC.1 | A0A0K2GFE1 | Threonylcarbamoyl-AMP synthase | 36.553 | 343 | 2.16 | 4.47 | 4.51E-05 | 3.25E-04 | yes | yes | 1 | 6 | 6 | 5 | 1 | 0 | 0 | 0.003436683 | 0.003573956 | 0.003544586 | 0.000258035 | 0.000337537 | 0.000138743 |
| dnaX | A0A0K2G706 | DNA polymerase III subunit gamma/tau | 63.411 | 592 | -0.92 | -1.89 | 1.18E-05 | 1.34E-04 | no | no | 0 | 9 | 6 | 9 | 26 | 26 | 21 | 0.003432045 | 0.003155492 | 0.00293222 | 0.006986305 | 0.007123783 | 0.007568832 |
| trpH | A0A0K2GB99 | Phosphoesterase | 30.85 | 283 | 0.609 | 1.53 | 9.15E-04 | 2.66E-03 | no | no | 0 | 4 | 3 | 2 | 4 | 3 | 1 | 0.003425722 | 0.00300199 | 0.002996064 | 0.0023763 | 0.001935887 | 0.001968044 |
| surE | A0A0K2GIR4 | 5-nucleotidase SurE | 29.92 | 272 | 0.0263 | 1.02 | 7.99E-01 | 8.24E-01 | no | no | 0 | 3 | 5 | 4 | 5 | 1 | 1 | 0.00341708 | 0.004585114 | 0.004185518 | 0.004622887 | 0.004517328 | 0.004007833 |
| moaA | A0A0K2GA69 | GTP 3,8-cyclase | 39.903 | 354 | -0.759 | -1.69 | 1.75E-04 | 7.96E-04 | no | no | 0 | 1 | 2 | 2 | 7 | 11 | 8 | 0.003402957 | 0.003051854 | 0.003204851 | 0.005893219 | 0.006380036 | 0.006219702 |
| NITMOv2_2252 | A0A0K2GCJ5 | Uncharacterized protein | 11.115 | 98 | 0.413 | 1.33 | 4.13E-03 | 8.55E-03 | no | no | 0 | 2 | 2 | 3 | 2 | 2 | 1 | 0.003385672 | 0.003256784 | 0.005452235 | 0.002878125 | 0.002669846 | 0.002778457 |
| NITMOv2_4294 | A0A0K2GIB1 | Putative ABC-type transport system, ATPase and permease component | 61.789 | 554 | -0.471 | -1.39 | 6.39E-04 | 2.05E-03 | no | no | 0 | 6 | 5 | 4 | 10 | 11 | 10 | 0.00338251 | 0.00336492 | 0.003145034 | 0.00503646 | 0.004875762 | 0.005118352 |
| rluB | A0A0K2G9Z3 | Pseudouridine synthase | 31.285 | 281 | -0.408 | -1.33 | 1.22E-01 | 1.54E-01 | no | no | 0 | 4 | 2 | 7 | 5 | 4 | 5 | 0.003381246 | 0.004672326 | 0.005207788 | 0.003908872 | 0.003524836 | 0.004559496 |
| NITMOv2_1714 | A0A0K2GB84 | Transcriptional regulator, AraC family | 36.513 | 339 | 0.771 | 1.71 | 1.10E-04 | 5.97E-04 | no | no | 0 | 9 | 8 | 8 | 4 | 4 | 5 | 0.003379138 | 0.003249744 | 0.004028688 | 0.002618254 | 0.002455282 | 0.002582102 |
| NITMOv2_4300 | A0A0K2GJ63 | Uncharacterized protein | 9.7998 | 90 | -0.428 | -1.35 | 3.24E-02 | 4.83E-02 | no | no | 0 | 1 | 0 | 1 | 2 | 1 | 1 | 0.003372603 | 0.00262811 | 0.002546281 | 0.005659557 | 0.003971054 | 0.004693636 |
| ruvB | A0A0K2G7E4 | Holliday junction ATP-dependent DNA helicase RuvB | 36.981 | 334 | -0.713 | -1.64 | 4.74E-04 | 1.63E-03 | no | no | 0 | 3 | 3 | 4 | 8 | 8 | 6 | 0.003370285 | 0.003020958 | 0.003138899 | 0.006369131 | 0.005750951 | 0.005209158 |
| psd | A0A0K2G8J0 | Phosphatidylserine decarboxylase proenzyme | 23.489 | 216 | -2.61 | -6.11 | 4.44E-03 | 9.01E-03 | yes | yes | 3 | 0 | 1 | 0 | 3 | 4 | 4 | 0.003356583 | 0.003907359 | 0.003672657 | 0.006394007 | 0.005071681 | 0.005337903 |
| NITMOv2_1331 | A0A0K2GA57 | Uncharacterized protein | 56.367 | 496 | 0.337 | 1.26 | 1.70E-02 | 2.78E-02 | no | no | 0 | 6 | 9 | 7 | 6 | 8 | 10 | 0.003350892 | 0.002846728 | 0.003281349 | 0.003054038 | 0.003155371 | 0.002966001 |
| NITMOv2_2114 | A0A0K2GC43 | Uncharacterized protein | 20.582 | 182 | 1.34 | 2.53 | 1.16E-02 | 2.02E-02 | yes | yes | 3 | 1 | 2 | 4 | 0 | 1 | 1 | 0.00334499 | 0.003078643 | 0.002325415 | 0.001640074 | 0.001798231 | 0.002257541 |
| NITMOv2_4247 | A0A0K2GI54 | Putative Leu/Ile/Val-binding protein, periplasmic-binding component of ABC transport system | 44.199 | 414 | 0.385 | 1.31 | 4.19E-03 | 8.64E-03 | no | no | 0 | 7 | 7 | 6 | 7 | 6 | 5 | 0.003342671 | 0.003390145 | 0.002975358 | 0.003017907 | 0.003009791 | 0.002568616 |
| NITMOv2_0169 | A0A0K2G6X2 | OmpA-like domain-containing protein | 55.135 | 519 | -0.296 | -1.23 | 2.18E-02 | 3.43E-02 | no | no | 0 | 7 | 8 | 8 | 12 | 11 | 11 | 0.003332132 | 0.003448026 | 0.003573152 | 0.004171556 | 0.00478394 | 0.00356046 |
| NITMOv2_4256 | A0A0K2GIE5 | Putative Transcriptional regulator | 26.249 | 233 | 0.645 | 1.56 | 9.59E-03 | 1.72E-02 | no | no | 0 | 2 | 3 | 2 | 2 | 2 | 1 | 0.003326862 | 0.003023109 | 0.002567371 | 0.002769734 | 0.002666272 | 0.002368126 |
| pgl | A0A0K2G6Y4 | 6-phosphogluconolactonase | 27.575 | 251 | 0.204 | 1.15 | 3.49E-01 | 3.91E-01 | no | no | 0 | 4 | 3 | 3 | 1 | 2 | 2 | 0.003324544 | 0.002863154 | 0.003165932 | 0.00228331 | 0.002199857 | 0.001810349 |
| NITMOv2_0005 | A0A0K2G6I1 | Uncharacterized protein | 21.313 | 189 | 1.06 | 2.08 | 1.67E-01 | 2.03E-01 | no | yes | 2 | 2 | 2 | 2 | 0 | 1 | 1 | 0.00331295 | 0.002741917 | 0.003191239 | 0 | 0.003284793 | 0.004021498 |
| NITMOv2_4028 | A0A0K2GIJ0 | Putative Nitrite oxidoreductase, alpha subunit | 131.62 | 1145 | -0.803 | -1.74 | 1.54E-02 | 2.56E-02 | no | no | 0 | 7 | 6 | 3 | 5 | 7 | 9 | 0.003309156 | 0.003708295 | 0.003677067 | 0.002658679 | 0.00251541 | 0.009945049 |
| NITMOv2_4718 | A0A0K2GKD1 | Uncharacterized protein | 41.738 | 369 | 0.897 | 1.86 | 5.24E-02 | 7.34E-02 | no | no | 0 | 1 | 2 | 1 | 0 | 0 | 0 | 0.003276063 | 0.004628133 | 0.003425525 | 0.001610903 | 0.001679374 | 0.001339475 |
| NITMOv2_4308 | A0A0K2GII8 | Uncharacterized protein | 19.175 | 177 | -0.541 | -1.45 | 7.29E-05 | 4.52E-04 | no | no | 0 | 1 | 2 | 4 | 6 | 5 | 4 | 0.003270371 | 0.003147279 | 0.003419198 | 0.006815131 | 0.007072357 | 0.003626271 |
| NITMOv2_2118 | A0A0K2GC55 | Uncharacterized protein | 24.938 | 228 | 0.515 | 1.43 | 3.06E-03 | 6.73E-03 | no | no | 0 | 4 | 7 | 5 | 4 | 5 | 3 | 0.003265734 | 0.003317403 | 0.003192006 | 0.002324474 | 0.002250973 | 0.002913496 |
| NITMOv2_4532 | A0A0K2GJ01 | CHASE3 domain-containing protein | 24.331 | 213 | -1.09 | -2.13 | 2.75E-03 | 6.17E-03 | yes | no | 0 | 3 | 3 | 5 | 11 | 13 | 12 | 0.003265102 | 0.002628501 | 0.005086618 | 0.010924792 | 0.007594549 | 0.006184998 |
| NITMOv2_3741 | A0A0K2GHM3 | Putative Phosphotransferase | 41.306 | 367 | -0.951 | -1.93 | 7.80E-04 | 2.36E-03 | no | no | 0 | 6 | 6 | 6 | 22 | 20 | 18 | 0.003252244 | 0.003005901 | 0.003096528 | 0.007548544 | 0.00817749 | 0.00712128 |
| NITMOv2_2936 | A0A0K2GEG0 | Peptidase_M78 domain-containing protein | 21.854 | 193 | 0.136 | 1.10 | 1.15E-01 | 1.46E-01 | no | no | 0 | 4 | 3 | 4 | 3 | 2 | 3 | 0.003238332 | 0.00318893 | 0.005732727 | 0.003919089 | 0.002758872 | 0.00345509 |
| pth | A0A0K2G8A0 | Peptidyl-tRNA hydrolase | 21.01 | 193 | -0.395 | -1.31 | 2.32E-01 | 2.74E-01 | no | yes | 1 | 0 | 0 | 1 | 1 | 3 | 0 | 0.003227792 | 0.002673672 | 0.002966156 | 0.003906207 | 0.002632247 | 0.00282305 |
| NITMOv2_0386 | A0A0K2G894 | Uncharacterized protein | 5.9363 | 53 | -0.183 | -1.14 | 3.49E-02 | 5.13E-02 | no | no | 0 | 3 | 3 | 0 | 2 | 3 | 2 | 0.003224841 | 0.004606233 | 0.005127264 | 0.006297463 | 0.005736657 | 0.005453163 |
| NITMOv2_4584 | A0A0K2GK28 | Putative HAD-like hydrolase, purine nucleotidase | 26.413 | 232 | 0.141 | 1.10 | 2.80E-01 | 3.23E-01 | no | no | 0 | 2 | 1 | 2 | 3 | 3 | 4 | 0.003217885 | 0.002982631 | 0.00316689 | 0.003507442 | 0.003424313 | 0.003035948 |
| NITMOv2_2444 | A0A0K2GE28 | TPR_REGION domain-containing protein | 28.278 | 254 | 0.166 | 1.12 | 1.61E-01 | 1.97E-01 | no | no | 0 | 4 | 4 | 3 | 3 | 2 | 2 | 0.00321388 | 0.003258544 | 0.002926852 | 0.003046486 | 0.002797248 | 0.002554771 |
| NITMOv2_1112 | A0A0K2G9B4 | Histidine kinase | 85.587 | 769 | -0.559 | -1.47 | 4.86E-04 | 1.66E-03 | no | no | 0 | 14 | 12 | 12 | 22 | 24 | 23 | 0.003213248 | 0.003171527 | 0.003486302 | 0.005853239 | 0.005601331 | 0.007362408 |
| sdhB | A0A0K2GJ86 | Succinate dehydrogenase/fumarate reductase, Fe-S protein subunit | 28.73 | 262 | 0.855 | 1.81 | 8.95E-04 | 2.61E-03 | no | no | 0 | 4 | 4 | 2 | 0 | 0 | 1 | 0.003212194 | 0.003807045 | 0.003282116 | 0.002480841 | 0.002118444 | 0.002505862 |
| pafA | A0A0K2GG80 | Pup--protein ligase | 53.081 | 458 | -0.582 | -1.50 | 1.02E-03 | 2.84E-03 | no | no | 0 | 5 | 5 | 9 | 13 | 16 | 10 | 0.003209454 | 0.003575521 | 0.004096941 | 0.006667501 | 0.005116427 | 0.00515881 |
| NITMOv2_3963 | A0A0K2GIA5 | Putative Cytochrome c55x | 12.598 | 119 | 0.734 | 1.66 | 2.66E-02 | 4.05E-02 | no | no | 0 | 2 | 1 | 1 | 1 | 1 | 1 | 0.003206713 | 0.003131245 | 0.00238159 | 0.001580696 | 0.002413644 | 0.00148131 |
| NITMOv2_2053 | A0A0K2GBZ6 | Putative Ribosomal RNA small subunit methyltransferase D | 20.261 | 189 | 0.649 | 1.57 | 3.30E-04 | 1.23E-03 | no | no | 0 | 4 | 3 | 3 | 2 | 3 | 2 | 0.003204606 | 0.003197339 | 0.002900203 | 0.001222503 | 0.0014881 | 0.001972359 |
| NITMOv2_1643 | A0A0K2GBT1 | hydrogenase maturation protein | 31.829 | 290 | 1.46 | 2.75 | 1.60E-03 | 4.08E-03 | yes | no | 0 | 4 | 4 | 4 | 3 | 2 | 1 | 0.003202076 | 0.003124792 | 0.002525766 | 0.001776895 | 0.001478514 | 0.000820429 |
| NITMOv2_4849 | A0A0K2GKR5 | Hydrolase_4 domain-containing protein | 28.441 | 250 | 0.402 | 1.32 | 3.65E-03 | 7.75E-03 | no | no | 0 | 6 | 11 | 7 | 7 | 7 | 6 | 0.003198071 | 0.006140275 | 0.003519662 | 0.003505665 | 0.003449328 | 0.002867644 |
| NITMOv2_1464 | A0A0K2GA96 | Type I restriction enzyme R Protein | 119.1 | 1060 | 0.144 | 1.10 | 9.65E-02 | 1.25E-01 | no | no | 0 | 25 | 28 | 33 | 24 | 24 | 24 | 0.003196174 | 0.003764416 | 0.003642173 | 0.003549347 | 0.003554822 | 0.002655825 |
| mazG | A0A0K2G994 | Nucleoside triphosphate pyrophosphohydrolase | 33.512 | 298 | -0.296 | -1.23 | 1.54E-02 | 2.56E-02 | no | no | 0 | 4 | 5 | 5 | 7 | 9 | 9 | 0.003192169 | 0.002984978 | 0.002963088 | 0.006070464 | 0.005675287 | 0.005055958 |
| NITMOv2_2897 | A0A0K2GEC4 | Uncharacterized protein | 11.49 | 102 | 0.915 | 1.89 | 1.16E-01 | 1.47E-01 | no | yes | 3 | 0 | 2 | 0 | 0 | 0 | 1 | 0.00318964 | 0.005105065 | 0.003444314 | 0.00266934 | 0.004259572 | 0.002433937 |
| NITMOv2_3731 | A0A0K2GHL4 | DUF4365 domain-containing protein | 52.059 | 458 | 0.226 | 1.17 | 2.14E-01 | 2.55E-01 | no | no | 0 | 6 | 5 | 3 | 4 | 2 | 4 | 0.003172566 | 0.002628306 | 0.00230145 | 0.002690663 | 0.002009842 | 0.001898636 |
| NITMOv2_2036 | A0A0K2GBY1 | Putative Oxidoreductase with FAD/NAD(P)-binding domain | 43.03 | 401 | 0.747 | 1.68 | 8.68E-05 | 5.13E-04 | no | no | 0 | 6 | 6 | 6 | 4 | 2 | 4 | 0.003166453 | 0.003290418 | 0.003303973 | 0.00117703 | 0.001342054 | 0.001453961 |
| NITMOv2_1181 | A0A0K2GAI1 | Uncharacterized protein | 47.398 | 453 | 0.0957 | 1.07 | 2.53E-01 | 2.94E-01 | no | no | 0 | 6 | 7 | 7 | 6 | 8 | 6 | 0.003166242 | 0.002980285 | 0.002708671 | 0.002500979 | 0.004382313 | 0.004132802 |
| NITMOv2_1387 | A0A0K2GAB1 | Putative Histidine kinase, contains GAF domain | 83.034 | 747 | -0.595 | -1.51 | 9.84E-04 | 2.78E-03 | no | no | 0 | 10 | 11 | 10 | 20 | 24 | 19 | 0.003162026 | 0.002867847 | 0.003556856 | 0.007004519 | 0.005760739 | 0.00529295 |
| NITMOv2_1379 | A0A0K2GA03 | Exodeoxyribonuclease III | 32.363 | 291 | 0.672 | 1.59 | 1.68E-04 | 7.81E-04 | no | no | 0 | 5 | 6 | 6 | 4 | 3 | 5 | 0.00315233 | 0.00374799 | 0.004272177 | 0.001870774 | 0.002334251 | 0.002003826 |
| mraY | A0A0K2G7Z7 | Phospho-N-acetylmuramoyl-pentapeptide-transferase | 39.291 | 358 | 0.694 | 1.62 | 1.13E-02 | 1.98E-02 | no | yes | 1 | 3 | 3 | 2 | 3 | 3 | 2 | 0.003147271 | 0.003313296 | 0.002644252 | 0.002299894 | 0.001959037 | 0.001891803 |

| | | | | | | | | | | | | | | | | | | | | | | | |
|---|---|---|---|---|---|---|---|---|---|---|---|---|---|---|---|---|---|---|---|---|---|---|---|
| cls | A0A0K2GBV4 | Phospholipase D Active site motif family protein | 48.287 | 436 | 0.408 | 1.33 | 3.21E-03 | 7.00E-03 | no | no | 0 | 6 | 7 | 8 | 2 | 3 | 4 | 0.003137153 | 0.003053809 | 0.002894259 | 0.00179185 | 0.002049772 | 0.001830308 |
| NITMOv2_3293 | A0A0K2GFF7 | DUF2294 domain-containing protein | 13.006 | 119 | 1.31 | 2.48 | 2.21E-03 | 5.18E-03 | yes | no | 0 | 2 | 1 | 2 | 0 | 1 | 1 | 0.003125982 | 0.002367841 | 0.003010252 | 0.00090133 | 0.001644571 | 0.001063517 |
| NITMOv2_4604 | A0A0K2GJ61 | Putative Phosphoglycolate phosphatase | 24.072 | 216 | -0.882 | -1.84 | 1.45E-03 | 3.75E-03 | no | no | 0 | 2 | 3 | 2 | 7 | 7 | 6 | 0.003107011 | 0.003275947 | 0.003303973 | 0.004194804 | 0.006254033 | 0.00497738 |
| NITMOv2_0925 | A0A0K2G915 | Uncharacterized protein | 9.0023 | 78 | -0.319 | -1.25 | 6.29E-01 | 6.69E-01 | no | yes | 4 | 1 | 1 | 1 | 1 | 1 | 0 | 0.003104271 | 0.003914203 | 0.004044026 | 0.002396587 | 0.003274228 | 0.002191011 |
| NITMOv2_2043 | A0A0K2GBY8 | DUF3883 domain-containing protein | 120.01 | 1088 | 0.323 | 1.25 | 4.39E-03 | 8.93E-03 | no | no | 0 | 12 | 9 | 14 | 12 | 11 | 12 | 0.003091834 | 0.003144737 | 0.003545928 | 0.003168203 | 0.002767261 | 0.002582642 |
| NITMOv2_2684 | A0A0K2GDR4 | Putative anti-sigma regulatory factor, serine/threonine protein kinase | 34.656 | 334 | -0.049 | -1.03 | 6.39E-01 | 6.79E-01 | no | no | 0 | 2 | 1 | 3 | 4 | 4 | 4 | 0.003089094 | 0.002616964 | 0.002211723 | 0.00285547 | 0.003312137 | 0.002959708 |
| uppP | A0A0K2G9W9 | Undecaprenyl-diphosphatase | 31.34 | 291 | 0.0865 | 1.06 | 2.97E-01 | 3.40E-01 | no | no | 0 | 3 | 2 | 3 | 4 | 4 | 2 | 0.00308804 | 0.003206529 | 0.003014662 | 0.02008039 | 0.018914514 | 0.020541738 |
| NITMOv2_1626 | A0A0K2GAS9 | HlyD_D23 domain-containing protein | 43.932 | 406 | 0.496 | 1.41 | 2.29E-04 | 9.71E-04 | no | no | 0 | 3 | 4 | 8 | 3 | 3 | 2 | 0.003076868 | 0.003008248 | 0.003144267 | 0.002031287 | 0.002346991 | 0.001609103 |
| yaaI | A0A0K2G7Y3 | Putative isochorismatase family protein YaaI | 21.525 | 190 | 1.58 | 2.99 | 5.84E-03 | 1.13E-02 | yes | yes | 3 | 2 | 2 | 2 | 0 | 0 | 0 | 0.003076868 | 0.00433208 | 0.003144842 | 0.000483597 | 0.000509328 | 0.000413694 |
| mfd | A0A0K2GJ73 | Transcription-repair-coupling factor | 128.18 | 1155 | -0.265 | -1.20 | 1.04E-01 | 1.33E-01 | no | no | 0 | 12 | 11 | 13 | 12 | 14 | 16 | 0.003066329 | 0.003172505 | 0.003196607 | 0.002479064 | 0.003222024 | 0.002597566 |
| lptB | A0A0K2G9P8 | Putative lipopolysaccharide transport protein B: ATP-binding component of ABC superfamily | 28.465 | 260 | 0.198 | 1.15 | 2.89E-01 | 3.32E-01 | no | no | 0 | 1 | 2 | 2 | 2 | 1 | 2 | 0.003064853 | 0.003016265 | 0.00291324 | 0.002649498 | 0.003534314 | 0.002830243 |
| NITMOv2_1625 | A0A0K2GB07 | Putative Cation efflux system protein CzcC | 46.06 | 417 | 0.418 | 1.34 | 1.03E-03 | 2.87E-03 | no | no | 0 | 6 | 10 | 7 | 3 | 3 | 4 | 0.003033235 | 0.002947825 | 0.003088283 | 0.001260248 | 0.002305352 | 0.002155947 |
| recR | A0A0K2G6P5 | Recombination protein RecR | 21.981 | 200 | -1.24 | -2.36 | 5.09E-03 | 1.01E-02 | yes | no | 0 | 3 | 4 | 4 | 5 | 9 | 5 | 0.003030916 | 0.003308994 | 0.003736501 | 0.006932999 | 0.0072299 | 0.005710654 |
| nuoG | A0A0K2G7Z6 | NADH-quinone oxidoreductase | 96.191 | 887 | -1.29 | -2.45 | 1.24E-05 | 1.38E-04 | yes | no | 0 | 11 | 12 | 17 | 44 | 37 | 42 | 0.003029019 | 0.003086465 | 0.003158071 | 0.008780525 | 0.009554052 | 0.007292101 |
| NITMOv2_3438 | A0A0K2GFU0 | Shikimate kinase | 21.063 | 195 | 0.322 | 1.25 | 3.47E-03 | 7.45E-03 | no | no | 0 | 4 | 5 | 5 | 4 | 4 | 4 | 0.003025225 | 0.002976765 | 0.003563183 | 0.00333227 | 0.002802996 | 0.003183933 |
| NITMOv2_2449 | A0A0K2GE32 | Uncharacterized protein | 39.544 | 362 | -0.43 | -1.35 | 3.95E-02 | 5.72E-02 | no | no | 0 | 9 | 9 | 8 | 7 | 5 | 8 | 0.003016161 | 0.003270081 | 0.003535958 | 0.004185031 | 0.003500288 | 0.003335155 |
| NITMOv2_1396 | A0A0K2GA44 | DUF2235 domain-containing protein | 38.902 | 339 | 0.268 | 1.20 | 4.18E-02 | 5.99E-02 | no | no | 0 | 3 | 3 | 5 | 4 | 3 | 4 | 0.003014053 | 0.002409883 | 0.003112824 | 0.002719389 | 0.003307632 | 0.002610692 |
| NITMOv2_0333 | A0A0K2G719 | Putative HTH-type transcriptional regulator, rrf2 family | 16.37 | 149 | -0.666 | -1.59 | 2.55E-04 | 1.04E-03 | no | no | 0 | 2 | 2 | 1 | 5 | 4 | 4 | 0.003007097 | 0.001853874 | 0.002326374 | 0.006268292 | 0.00688405 | 0.006845268 |
| NITMOv2_3970 | A0A0K2GHL1 | Uncharacterized protein | 35.749 | 318 | -0.931 | -1.91 | 9.80E-03 | 1.75E-02 | no | no | 0 | 3 | 4 | 6 | 5 | 5 | 5 | 0.003001195 | 0.00273957 | 0.003210028 | 0.001336787 | 0.001498013 | 0.001927586 |
| NITMOv2_4852 | A0A0K2GJU0 | DUF2294 domain-containing protein | 13.676 | 123 | 1.31 | 2.48 | 4.69E-04 | 1.62E-03 | yes | no | 0 | 3 | 3 | 3 | 2 | 2 | 2 | 0.002987705 | 0.003312318 | 0.0030601 | 0.001507251 | 0.001447503 | 0.001537592 |
| NITMOv2_4605 | A0A0K2GK46 | Uncharacterized protein | 22.19 | 195 | -1.47 | -2.77 | 7.58E-03 | 1.41E-02 | yes | no | 0 | 3 | 3 | 5 | 3 | 4 | 4 | 0.002984965 | 0.001925775 | 0.003448916 | 0.001591357 | 0.002149984 | 0.001255089 |
| dnaA | A0A0K2G696 | Chromosomal replication initiator protein DnaA | 50.568 | 447 | -1.73 | -3.32 | 3.07E-06 | 5.51E-05 | yes | no | 0 | 2 | 4 | 5 | 28 | 27 | 21 | 0.002974425 | 0.00233851 | 0.003124328 | 0.013544823 | 0.0135293 | 0.010057252 |
| NITMOv2_2112 | A0A0K2GCE7 | Uncharacterized protein | 30.839 | 296 | 0.407 | 1.33 | 1.45E-03 | 3.74E-03 | no | no | 0 | 5 | 3 | 4 | 3 | 5 | 4 | 0.002959881 | 0.002548524 | 0.002288029 | 0.002754483 | 0.002521935 | 0.013636413 |
| NITMOv2_0039 | A0A0K2G6C1 | 4-carboxymuconolactone decarboxylase | 12.065 | 110 | 0.557 | 1.47 | 7.35E-03 | 1.38E-02 | no | no | 0 | 2 | 2 | 2 | 3 | 1 | 1 | 0.002958616 | 0.003584516 | 0.001989516 | 0.00285621 | 0.001982497 | 0.001502816 |
| NITMOv2_2104 | A0A0K2GC34 | Uncharacterized protein | 31.357 | 297 | -0.248 | -1.19 | 6.79E-02 | 9.17E-02 | no | no | 0 | 4 | 4 | 4 | 2 | 3 | 3 | 0.002955665 | 0.002986738 | 0.002763887 | 0.002132274 | 0.001950802 | 0.002461268 |
| pilC | A0A0K2G7Z0 | Type IV pilus biogenesis protein PilC | 43.25 | 404 | 0.0388 | 1.03 | 7.52E-01 | 7.82E-01 | no | no | 0 | 2 | 2 | 3 | 3 | 3 | 3 | 0.002948077 | 0.003182868 | 0.001932382 | 0.001899352 | 0.001752552 | 0.001807292 |
| NITMOv2_3463 | A0A0K2GFW7 | Uncharacterized protein | 18.949 | 169 | 0.549 | 1.46 | 1.45E-03 | 3.74E-03 | no | no | 0 | 4 | 5 | 4 | 3 | 3 | 4 | 0.002945758 | 0.00328416 | 0.002997598 | 0.001673094 | 0.00199648 | 0.002033855 |
| NITMOv2_2538 | A0A0K2GDC7 | Response regulator, LuxR family | 23.437 | 217 | -0.861 | -1.82 | 2.66E-03 | 6.01E-03 | no | no | 0 | 1 | 1 | 3 | 6 | 6 | 4 | 0.002943861 | 0.003213764 | 0.003367241 | 0.007724457 | 0.006089187 | 0.005233612 |
| NITMOv2_3513 | A0A0K2GG31 | Glycosyl transferase family 2 | 30.682 | 267 | 0.377 | 1.30 | 1.76E-02 | 2.86E-02 | no | no | 0 | 3 | 3 | 5 | 3 | 4 | 3 | 0.002924679 | 0.003349081 | 0.007501569 | 0.008768086 | 0.008719103 | 0.002763712 |
| NITMOv2_3559 | A0A0K2GG90 | Putative Murein hydrolase EnvC | 44.515 | 393 | 0.0305 | 1.02 | 6.81E-01 | 7.17E-01 | no | no | 0 | 6 | 7 | 6 | 6 | 9 | 4 | 0.00291941 | 0.003413219 | 0.003660003 | 0.003865634 | 0.00426066 | 0.003447898 |
| kdsC | A0A0K2GFY2 | 3-deoxy-D-manno-octulosonate 8-phosphate phosphatase KdsC | 22.034 | 200 | -3.58 | -11.96 | 1.46E-03 | 3.75E-03 | yes | yes | 3 | 2 | 5 | 4 | 8 | 8 | 5 | 0.002907816 | 0.003319358 | 0.003403285 | 0.011059836 | 0.011415983 | 0.009765237 |
| NITMOv2_2029 | A0A0K2GBW7 | HTH cro/C1-type domain-containing protein | 10.68 | 95 | 0.191 | 1.14 | 5.73E-01 | 6.15E-01 | no | yes | 3 | 1 | 1 | 2 | 1 | 1 | 1 | 0.002900228 | 0.002562799 | 0.002702919 | 0.001540568 | 0.001444022 | 0.001114152 |
| devS | A0A0K2GAG9 | CRISPR-associated protein Cas5/DevS | 23.523 | 209 | 0.32 | 1.25 | 2.06E-02 | 3.26E-02 | no | no | 0 | 5 | 5 | 5 | 3 | 3 | 4 | 0.002899596 | 0.003073168 | 0.002885632 | 0.002754483 | 0.003092758 | 0.003119561 |
| NITMOv2_3728 | A0A0K2GGN2 | NTP_transf_2 domain-containing protein | 11.782 | 105 | -0.553 | -1.47 | 1.84E-01 | 2.22E-01 | no | yes | 3 | 1 | 0 | 1 | 1 | 0 | 0 | 0.002897277 | 0.003471882 | 0.003259684 | 0.002519488 | 0.001961833 | 0.002139944 |
| NITMOv2_2393 | A0A0K2GCY0 | Putative 2-octaprenyl-3-methyl-6-methoxy-1, 4-benzoquinol hydroxylase | 45.636 | 423 | 0.299 | 1.23 | 8.79E-02 | 1.15E-01 | no | no | 0 | 6 | 5 | 5 | 7 | 7 | 4 | 0.002885895 | 0.00252369 | 0.00282984 | 0.003000287 | 0.002620749 | 0.003619618 |
| NITMOv2_2417 | A0A0K2GE02 | Uncharacterized protein | 37.679 | 338 | 0.528 | 1.44 | 1.33E-02 | 2.26E-02 | no | no | 0 | 3 | 2 | 6 | 3 | 1 | 3 | 0.002874723 | 0.004654141 | 0.003798044 | 0.002842587 | 0.002374802 | 0.002792482 |
| NITMOv2_3505 | A0A0K2GH13 | Methyltransf_11 domain-containing protein | 33.865 | 291 | -0.912 | -1.88 | 5.12E-06 | 7.60E-05 | no | no | 0 | 5 | 6 | 5 | 11 | 12 | 14 | 0.002866291 | 0.002961708 | 0.002940848 | 0.005857385 | 0.00817112 | 0.008359106 |
| NITMOv2_4283 | A0A0K2GI87 | dTTP/UTP pyrophosphatase | 23.063 | 210 | 0.393 | 1.31 | 8.14E-03 | 1.50E-02 | no | no | 0 | 3 | 4 | 3 | 2 | 1 | 1 | 0.002850061 | 0.002819352 | 0.003076205 | 0.00090743 | 0.001141909 | 0.000901362 |
| lpxK | A0A0K2GG44 | Tetraacyldisaccharide 4-kinase | 38.6 | 354 | -0.482 | -1.40 | 1.65E-02 | 2.71E-02 | no | no | 0 | 8 | 5 | 4 | 8 | 4 | 5 | 0.002845845 | 0.003421236 | 0.00337376 | 0.003748359 | 0.00394433 | 0.004515442 |
| NITMOv2_2539 | A0A0K2GED7 | Response regulator, LuxR family | 23.391 | 213 | -1.14 | -2.20 | 8.84E-06 | 1.09E-04 | yes | no | 0 | 6 | 7 | 5 | 11 | 10 | 11 | 0.00284184 | 0.001863358 | 0.003257192 | 0.009697847 | 0.009409248 | 0.008591963 |
| nuoE | A0A0K2G7N5 | NADH-quinone oxidoreductase, subunit E | 19.698 | 176 | -0.557 | -1.47 | 1.16E-03 | 3.15E-03 | no | no | 0 | 3 | 3 | 4 | 4 | 5 | 5 | 0.002833198 | 0.001202067 | 0.002864734 | 0.00498182 | 0.004366466 | 0.005458018 |
| NITMOv2_3654 | A0A0K2GGH7 | Putative O-methyltransferase, family 2 | 35.89 | 327 | -0.589 | -1.50 | 2.67E-03 | 6.04E-03 | no | no | 0 | 3 | 3 | 2 | 6 | 4 | 3 | 0.002826242 | 0.002704568 | 0.002423386 | 0.005303734 | 0.005168942 | 0.004580894 |
| NITMOv2_3766 | A0A0K2GHP7 | Trypsin-like serine protease | 34.638 | 332 | -0.324 | -1.25 | 3.46E-02 | 5.10E-02 | no | no | 0 | 2 | 3 | 3 | 6 | 7 | 5 | 0.002817389 | 0.002892486 | 0.002622395 | 0.003978467 | 0.003690614 | 0.003706288 |
| hisH | A0A0K2G8K5 | Imidazole glycerol phosphate synthase subunit HisH | 22.101 | 202 | -0.33 | -1.26 | 1.27E-02 | 2.18E-02 | no | no | 0 | 2 | 2 | 4 | 4 | 3 | 6 | 0.002809589 | 0.002833627 | 0.002842494 | 0.006160642 | 0.004274954 | 0.004816628 |
| NITMOv2_4344 | A0A0K2GIE7 | Uncharacterized protein | 9.8893 | 91 | 0.0952 | 1.07 | 4.67E-01 | 5.10E-01 | no | no | 0 | 2 | 2 | 2 | 2 | 2 | 2 | 0.002801158 | 0.003454674 | 0.00330704 | 0.002520969 | 0.002718476 | 0.0029099 |
| NITMOv2_4215 | A0A0K2GIA9 | Glycoside hydrolase 15-related protein | 78.308 | 690 | 0.662 | 1.58 | 2.53E-04 | 1.03E-03 | no | no | 0 | 12 | 13 | 12 | 6 | 6 | 7 | 0.002799893 | 0.003099762 | 0.003198525 | 0.001800438 | 0.001858824 | 0.001963369 |
| pilH | A0A0K2GBL7 | Protein PilH | 13.267 | 120 | -0.603 | -1.52 | 1.32E-01 | 1.65E-01 | no | no | 0 | 1 | 3 | 2 | 6 | 4 | 6 | 0.002794413 | 0.006307269 | 0.005582224 | 0.005939714 | 0.006333115 | 0.005504769 |
| ybgC | A0A0K2GJG0 | Acyl-CoA thioester hydrolase YbgC | 14.651 | 133 | -0.637 | -1.56 | 3.92E-01 | 4.35E-01 | no | yes | 4 | 1 | 1 | 1 | 1 | 2 | 3 | 0.002793359 | 0.002705741 | 0.00301332 | 0.002019589 | 0.002850539 | 0.002748968 |
| NITMOv2_2678 | A0A0K2GDQ6 | Nitrogen regulation protein B | 40.539 | 365 | -0.266 | -1.20 | 2.69E-02 | 4.09E-02 | no | no | 0 | 5 | 5 | 5 | 7 | 7 | 6 | 0.002786403 | 0.002846924 | 0.003140624 | 0.003239279 | 0.003166402 | 0.002810104 |
| NITMOv2_1163 | A0A0K2G9Q6 | Putative Histidine kinase | 40.811 | 384 | 0.0465 | 1.03 | 7.54E-01 | 7.84E-01 | no | no | 0 | 4 | 2 | 5 | 3 | 2 | 3 | 0.002786192 | 0.002305267 | 0.002279402 | 0.002284346 | 0.002266355 | 0.001883352 |
| zitB | A0A0K2GFU5 | Zinc transporter ZitB | 34.185 | 313 | -0.109 | -1.08 | 8.45E-01 | 8.64E-01 | no | yes | 1 | 2 | 1 | 2 | 2 | 2 | 1 | 0.002782187 | 0.002873518 | 0.004200664 | 0.003739179 | 0.003801858 | 0.003476847 |
| NITMOv2_0795 | A0A0K2G8G5 | HIT domain-containing protein | 17.011 | 149 | 0.804 | 1.75 | 1.94E-03 | 4.69E-03 | no | no | 0 | 3 | 2 | 2 | 1 | 1 | 0 | 0.002778814 | 0.003417912 | 0.003664605 | 0.00135854 | 0.001653894 | 0.001819159 |
| ribD | A0A0K2GFG4 | Riboflavin biosynthesis protein RibD | 36.849 | 346 | -0.223 | -1.17 | 1.82E-01 | 2.20E-01 | no | no | 0 | 9 | 8 | 9 | 6 | 8 | 7 | 0.002767432 | 0.004413426 | 0.005083167 | 0.005069925 | 0.005353519 | 0.003939144 |
| NITMOv2_0417 | A0A0K2G7C0 | PilZ domain-containing protein | 19.113 | 170 | -0.103 | -1.07 | 2.36E-01 | 2.77E-01 | no | no | 0 | 2 | 2 | 1 | 2 | 2 | 3 | 0.002762795 | 0.003136916 | 0.002677995 | 0.003657589 | 0.003468593 | 0.003687048 |
| NITMOv2_3277 | A0A0K2GFD1 | Putative Transcriptional regulator, Crp/Fnr family | 26.141 | 238 | -1.13 | -2.19 | 2.95E-02 | 4.46E-02 | yes | yes | 4 | 1 | 1 | 1 | 2 | 1 | 1 | 0.002750358 | 0.002712976 | 0.002451761 | 0.002329509 | 0.00227692 | 0.002193528 |
| NITMOv2_0730 | A0A0K2G867 | Uncharacterized protein | 18.87 | 174 | -2.41 | -5.31 | 1.27E-04 | 6.52E-04 | yes | yes | 3 | 0 | 0 | 0 | 2 | 2 | 3 | 0.002740662 | 0.002786892 | 0.002309502 | 0.007804417 | 0.008468805 | 0.008471309 |
| cas6 | A0A0K2GAP6 | CRISPR-associated protein, Cas6-related | 21.83 | 197 | 0.353 | 1.28 | 1.72E-03 | 4.28E-03 | no | no | 0 | 3 | 2 | 2 | 2 | 3 | 3 | 0.002738975 | 0.0025452 | 0.002318322 | 0.003397126 | 0.002176086 | 0.003514069 |
| NITMOv2_0705 | A0A0K2G842 | Putative Cytochrome c biogenesis protein, ResB-like | 54.363 | 478 | -0.395 | -1.31 | 8.87E-03 | 1.61E-02 | no | no | 0 | 6 | 8 | 7 | 11 | 11 | 11 | 0.002728015 | 0.002611489 | 0.002650579 | 0.004367607 | 0.003722309 | 0.003669786 |
| NITMOv2_0249 | A0A0K2G6W1 | Uncharacterized protein | 9.4049 | 86 | -0.984 | -1.98 | 1.25E-05 | 1.39E-04 | no | no | 0 | 4 | 4 | 3 | 2 | 4 | 4 | 0.002719583 | 0.002718843 | 0.002199645 | 0.004297272 | 0.004656693 | 0.004551045 |
| NITMOv2_3729 | A0A0K2GGQ5 | Piwi domain-containing protein | 53.874 | 480 | 0.0394 | 1.03 | 7.18E-01 | 7.52E-01 | no | no | 0 | 8 | 8 | 8 | 7 | 5 | 7 | 0.002713892 | 0.003286116 | 0.003567401 | 0.00264239 | 0.003235696 | 0.003360869 |
| gppA | A0A0K2GAY6 | Exopolyphosphatase | 34.956 | 319 | 0.232 | 1.17 | 1.85E-01 | 2.22E-01 | no | no | 0 | 4 | 4 | 4 | 4 | 3 | 3 | 0.002712627 | 0.002860808 | 0.003175326 | 0.002624029 | 0.001835364 | 0.002139405 |
| smc | A0A0K2G869 | Chromosome partition protein Smc | 135.85 | 1228 | -2.12 | -4.35 | 8.34E-08 | 7.62E-06 | yes | no | 0 | 7 | 7 | 7 | 60 | 76 | 76 | 0.002705039 | 0.002425331 | 0.002322348 | 0.014061307 | 0.014405576 | 0.013644145 |
| NITMOv2_4834 | A0A0K2GKQ2 | Uncharacterized protein | 78.34 | 725 | 0.079 | 1.06 | 3.51E-01 | 3.93E-01 | no | no | 0 | 8 | 6 | 6 | 4 | 3 | 1 | 0.002702931 | 0.002909107 | 0.002979193 | 0.002296932 | 0.002113162 | 0.002139225 |
| NITMOv2_4638 | A0A0K2GJI0 | TPM_phosphatase domain-containing protein | 24.142 | 215 | -0.0361 | -1.03 | 9.72E-01 | 9.75E-01 | no | yes | 3 | 2 | 0 | 1 | 2 | 0 | 0 | 0.002691759 | 0.00205458 | 0.001491993 | 0.004173777 | 0.001251568 | 0.001403416 |
| NITMOv2_2113 | A0A0K2GC50 | Uncharacterized protein | 57.362 | 526 | 0.279 | 1.21 | 1.42E-02 | 2.39E-02 | no | no | 0 | 7 | 9 | 9 | 4 | 6 | 4 | 0.002688386 | 0.002462093 | 0.002458855 | 0.001355045 | 0.002519915 | 0.001927946 |
| atpB | A0A0K2GI02 | ATP synthase subunit a | 26.762 | 249 | -0.758 | -1.69 | 8.36E-02 | 1.10E-01 | no | yes | 2 | 1 | 1 | 1 | 4 | 2 | 2 | 0.002674264 | 0.002417314 | 0.002926085 | 0.002987552 | 0.00242685 | 0.002384848 |
| NITMOv2_2190 | A0A0K2GCB5 | Rhs family protein | 135.28 | 1247 | 4.1 | 17.15 | 1.11E-06 | 3.12E-05 | yes | yes | 3 | 2 | 1 | 0 | 0 | 0 | 0 | 0.002672788 | 0.002502767 | 0.00275756 | 0.003254382 | 0.003269256 | 0.000138687 |
| NITMOv2_1014 | A0A0K2G905 | DUF547 domain-containing protein | 32.378 | 283 | -0.021 | -1.01 | 7.91E-01 | 8.18E-01 | no | no | 0 | 4 | 5 | 6 | 4 | 7 | 6 | 0.002666254 | 0.002524276 | 0.002714422 | 0.002889971 | 0.002826767 | 0.002581743 |
| NITMOv2_0155 | A0A0K2G6N8 | Putative Sensor histidine kinase | 55.398 | 495 | -0.0779 | -1.06 | 2.32E-01 | 2.74E-01 | no | no | 0 | 6 | 6 | 5 | 6 | 8 | 5 | 0.002663935 | 0.001599412 | 0.001678693 | 0.001929559 | 0.002399194 | 0.002375858 |
| NITMOv2_4608 | A0A0K2GJ64 | Putative Guanyl-specific ribonuclease Sa3 | 15.797 | 139 | 0.238 | 1.18 | 1.16E-01 | 1.47E-01 | no | no | 0 | 3 | 3 | 2 | 3 | 3 | 2 | 0.00266267 | 0.00225423 | 0.002390793 | 0.002311148 | 0.002103063 | 0.001844153 |
| NITMOv2_3317 | A0A0K2GFH2 | L-2-hydroxyglutarate oxidase | 44.437 | 399 | -0.4 | -1.32 | 7.01E-03 | 1.32E-02 | no | no | 0 | 4 | 3 | 2 | 11 | 10 | 9 | 0.002652974 | 0.002599756 | 0.002586351 | 0.006164936 | 0.005643281 | 0.004582512 |
| NITMOv2_0229 | A0A0K2G6U2 | Selenocysteine lyase | 48.558 | 436 | 0.634 | 1.55 | 4.53E-04 | 1.58E-03 | no | no | 0 | 4 | 5 | 4 | 5 | 3 | 5 | 0.002650023 | 0.002588415 | 0.002828306 | 0.002359272 | 0.004561919 | 0.004598156 |
| NITMOv2_4340 | A0A0K2GIF4 | Putative Fe-S oxidoreductase | 48.572 | 432 | -0.394 | -1.31 | 6.66E-03 | 1.26E-02 | no | no | 0 | 5 | 5 | 5 | 8 | 9 | 7 | 0.002648969 | 0.002360411 | 0.002577148 | 0.003696977 | 0.004723191 | 0.003711502 |
| queC | A0A0K2G9F9 | 7-cyano-7-deazaguanine synthase | 24.447 | 232 | 0.127 | 1.09 | 1.85E-01 | 2.22E-01 | no | no | 0 | 3 | 3 | 4 | 2 | 3 | 4 | 0.002638851 | 0.002716301 | 0.002535928 | 0.003645892 | 0.002486201 | 0.00395281 |
| NITMOv2_1078 | A0A0K2G9H6 | Putative DNA mismatch repair protein, MutS family | 67.656 | 612 | -0.255 | -1.19 | 2.48E-01 | 2.90E-01 | no | no | 0 | 4 | 4 | 2 | 1 | 3 | 2 | 0.002618194 | 0.002965619 | 0.002948325 | 0.002103251 | 0.00169895 | 0.001474406 |
| NITMOv2_1893 | A0A0K2GBI6 | J domain-containing protein | 16.551 | 144 | 0.895 | 1.86 | 2.12E-02 | 3.35E-02 | no | yes | 3 | 3 | 3 | 2 | 0 | 0 | 0 | 0.002606179 | 0.001762124 | 0.001994309 | 0.000604544 | 0.000541738 | 0.000874157 |
| NITMOv2_1984 | A0A0K2GBS7 | Ribonuclease BN | 49.048 | 447 | 0.371 | 1.29 | 2.44E-03 | 5.61E-03 | no | no | 0 | 3 | 5 | 5 | 5 | 5 | 5 | 0.002602174 | 0.002367255 | 0.003648883 | 0.002197722 | 0.003010723 | 0.002811363 |
| pucG | A0A0K2G960 | Purine catabolism protein PucG | 42.286 | 398 | -0.193 | -1.14 | 1.51E-02 | 2.53E-02 | no | no | 0 | 3 | 1 | 5 | 3 | 3 | 2 | 0.002561492 | 0.002619506 | 0.002341712 | 0.002928767 | 0.003122122 | 0.006683257 |
| NITMOv2_2392 | A0A0K2GDX5 | Uncharacterized protein | 28.475 | 253 | 0.175 | 1.13 | 2.61E-01 | 3.03E-01 | no | no | 0 | 3 | 3 | 5 | 4 | 5 | 3 | 0.002560017 | 0.002990258 | 0.003090968 | 0.003553197 | 0.005372319 | 0.002828445 |
| mntB | A0A0K2G809 | Manganese transport system, ATPase component | 29.424 | 265 | -0.0113 | -1.01 | 8.95E-01 | 9.10E-01 | no | no | 0 | 3 | 3 | 4 | 4 | 5 | 4 | 0.002559174 | 0.001740536 | 0.001835504 | 0.001909569 | 0.002876175 | 0.002934354 |

| | | | | | | | | | | | | | | | | | | | | | | |
|---|---|---|---|---|---|---|---|---|---|---|---|---|---|---|---|---|---|---|---|---|---|---|
| NITMOv2_2478 | A0A0K2GD58 | Ribosomal RNA small subunit methyltransferase E | 26.54 | 244 | -0.551 | -1.47 | 5.01E-02 | 7.05E-02 | no | no | 0 | 2 | 3 | 4 | 10 | 6 | 6 | 0.002548634 | 0.003621278 | 0.003631053 | 0.007037243 | 0.004952048 | 0.00582915 |
| hoxA | A0A088N9I8 | Hydrogenase transcriptional regulatory protein | 55.316 | 490 | 0.811 | 1.75 | 7.61E-05 | 4.65E-04 | no | no | 0 | 6 | 6 | 4 | 1 | 2 | 2 | 0.002547159 | 0.002202998 | 0.002367211 | 0.00066598 | 0.001048424 | 0.000851051 |
| NITMOv2_4805 | A0A0K2GJP1 | Uncharacterized protein | 16.312 | 145 | 0.882 | 1.84 | 5.45E-02 | 7.56E-02 | no | yes | 2 | 2 | 1 | 1 | 1 | 1 | 1 | 0.002542311 | 0.002493185 | 0.002145579 | 0.000514204 | 0.000530722 | 0.002150733 |
| NITMOv2_1994 | A0A0K2GBT3 | TPR_REGION domain-containing protein | 24.703 | 227 | 0.479 | 1.39 | 1.12E-02 | 1.97E-02 | no | no | 0 | 2 | 2 | 3 | 2 | 3 | 3 | 0.002537252 | 0.002017036 | 0.002293398 | 0.001005648 | 0.001855406 | 0.001685739 |
| NITMOv2_2050 | A0A0K2GD03 | CxxC_CXXC_SSSS domain-containing protein | 12.514 | 125 | -3.4 | -10.56 | 3.79E-04 | 1.37E-03 | yes | yes | 3 | 1 | 1 | 1 | 5 | 6 | 5 | 0.002534301 | 0.001988291 | 0.001748768 | 0.027599615 | 0.027243799 | 0.024335774 |
| thiE | A0A0K2GC15 | Thiamine-phosphate synthase | 22.793 | 213 | 0.789 | 1.73 | 1.24E-03 | 3.29E-03 | no | no | 0 | 2 | 4 | 1 | 0 | 0 | 0 | 0.002530717 | 0.003591555 | 0.003664605 | 0.001842047 | 0.002878505 | 0.003055907 |
| NITMOv2_2531 | A0A0K2GDM3 | Uncharacterized protein | 15.404 | 141 | -1.89 | -3.71 | 5.08E-06 | 7.60E-05 | yes | no | 0 | 1 | 0 | 0 | 2 | 4 | 2 | 0.002523129 | 0.002672694 | 0.002285153 | 0.008615866 | 0.008684301 | 0.00681434 |
| NITMOv2_0183 | A0A0K2G6Q0 | Uncharacterized protein | 14.771 | 147 | -0.603 | -1.52 | 1.67E-02 | 2.74E-02 | no | no | 0 | 0 | 1 | 2 | 2 | 0 | 2 | 0.002518492 | 0.002429829 | 0.002620286 | 0.003613463 | 0.002968619 | 0.003501302 |
| nubp | A0A0K2GGU8 | Iron-sulfur cluster carrier protein | 34.76 | 319 | -2.32 | -4.99 | 1.82E-04 | 8.17E-04 | yes | no | 0 | 3 | 4 | 5 | 6 | 7 | 6 | 0.002484344 | 0.003567308 | 0.003144842 | 0.009961715 | 0.009707089 | 0.009384395 |
| NITMOv2_4135 | A0A0K2GIS9 | Uncharacterized protein | 18.6 | 166 | 0.97 | 1.96 | 6.08E-04 | 1.97E-03 | no | no | 0 | 2 | 6 | 3 | 1 | 2 | 2 | 0.002466216 | 0.004182684 | 0.003003158 | 0.001009854 | 0.00172816 | 0.001667668 |
| NITMOv2_1647 | A0A088N9K0 | Rieske domain-containing protein | 31.74 | 294 | 1.4 | 2.64 | 1.17E-03 | 3.16E-03 | yes | no | 0 | 2 | 1 | 2 | 0 | 0 | 0 | 0.002466005 | 0.002019969 | 0.002068697 | 0.000901552 | 0.000613129 | 0.00045176 |
| ppsA | A0A0K2G7D9 | Phosphoenolpyruvate synthase | 97.233 | 876 | 0.325 | 1.25 | 1.49E-02 | 2.49E-02 | no | no | 0 | 8 | 6 | 8 | 2 | 5 | 4 | 0.002453569 | 0.002320715 | 0.002452528 | 0.001218639 | 0.001352759 | 0.001405717 |
| NITMOv2_2097 | A0A0K2GCD5 | J domain-containing protein | 23.839 | 209 | -0.82 | -1.77 | 1.10E-03 | 3.01E-03 | no | no | 0 | 3 | 3 | 4 | 6 | 5 | 6 | 0.002453358 | 0.002207691 | 0.002068314 | 0.003660255 | 0.003856703 | 0.003228527 |
| NITMOv2_4489 | A0A0K2GIS7 | Uncharacterized protein | 19 | 170 | 1.34 | 2.53 | 1.96E-02 | 3.13E-02 | yes | yes | 2 | 1 | 2 | 2 | 2 | 1 | 1 | 0.002446402 | 0.002005694 | 0.002347847 | 0.000586982 | 0.001308029 | 0.000532334 |
| NITMOv2_3974 | A0A0K2GHC0 | Uncharacterized protein | 16.967 | 151 | -2.41 | -5.31 | 1.41E-03 | 3.68E-03 | yes | yes | 2 | 0 | 0 | 0 | 22 | 17 | 11 | 0.002424269 | 0.002119305 | 0.002228978 | 0.01503549 | 0.01498541 | 0.014001791 |
| NITMOv2_2535 | A0A0K2GDD0 | Putative 6-phosphogluconolactonase | 41.237 | 404 | 0.301 | 1.23 | 2.05E-03 | 4.90E-03 | no | no | 0 | 2 | 3 | 2 | 0 | 1 | 1 | 0.002417103 | 0.002369406 | 0.002395011 | 0.001069306 | 0.002098246 | 0.001389408 |
| kdtX | A0A0K2GG56 | Lipopolysaccharide core biosynthesis glycosyltransferase KdtX | 29.126 | 247 | 0.832 | 1.78 | 9.36E-02 | 1.22E-01 | no | yes | 2 | 2 | 2 | 2 | 0 | 1 | 2 | 0.002415838 | 0.002778092 | 0.002248726 | 0.000615368 | 0.00306417 | 0.001936037 |
| NITMOv2_1731 | A0A0K2GB05 | Uncharacterized protein | 47.487 | 412 | -0.739 | -1.67 | 6.89E-05 | 4.37E-04 | no | no | 0 | 5 | 6 | 3 | 11 | 11 | 11 | 0.002413519 | 0.002686578 | 0.002186224 | 0.00487802 | 0.005702165 | 0.005142627 |
| NITMOv2_2944 | A0A0K2GEQ7 | Uncharacterized protein | 23.195 | 209 | -0.64 | -1.56 | 2.16E-03 | 5.13E-03 | no | no | 0 | 1 | 3 | 6 | 3 | 2 | 2 | 0.002402558 | 0.002100337 | 0.002115861 | 0.001340193 | 0.001321142 | 0.002580304 |
| NITMOv2_0314 | A0A0K2G816 | Beta_helix domain-containing protein | 28.92 | 265 | 0.658 | 1.58 | 3.98E-02 | 5.74E-02 | no | no | 0 | 2 | 1 | 1 | 2 | 1 | 2 | 0.002390543 | 0.002483603 | 0.002210381 | 0.001837161 | 0.001976438 | 0.001725082 |
| NITMOv2_1644 | A0A088NA15 | Uncharacterized protein | 37.378 | 338 | 1.91 | 3.76 | 4.21E-05 | 3.07E-04 | yes | yes | 3 | 4 | 2 | 4 | 0 | 0 | 0 | 0.0023897 | 0.001707626 | 0.002208847 | 0.000109973 | 0.000152607 | 0 |
| NITMOv2_4169 | A0A0K2GI66 | Uncharacterized protein | 27.931 | 259 | -0.744 | -1.67 | 1.27E-02 | 2.18E-02 | no | no | 0 | 0 | 0 | 1 | 5 | 4 | 2 | 0.002378318 | 0.002305854 | 0.001666711 | 0.005285521 | 0.005907251 | 0.004123092 |
| NITMOv2_3631 | A0A0K2GGD8 | Histidine kinase | 118.47 | 1059 | 0.12 | 1.09 | 2.89E-01 | 3.32E-01 | no | no | 0 | 4 | 8 | 6 | 13 | 12 | 9 | 0.002370097 | 0.002577464 | 0.002567946 | 0.002731975 | 0.002711018 | 0.002257541 |
| crtB | A0A0K2GA75 | Phytoene synthase | 32.591 | 286 | -0.126 | -1.09 | 2.13E-01 | 2.54E-01 | no | no | 0 | 3 | 3 | 3 | 6 | 4 | 4 | 0.002366303 | 0.002180315 | 0.00268854 | 0.003153692 | 0.003732097 | 0.003181236 |
| NITMOv2_0596 | A0A0K2G8V6 | Putative Transcriptional regulator acrR | 24.49 | 219 | -0.369 | -1.29 | 5.51E-02 | 7.63E-02 | no | no | 0 | 5 | 3 | 3 | 3 | 4 | 3 | 0.002358082 | 0.002536205 | 0.002014248 | 0.000999562 | 0.00184453 | 0.00164609 |
| fdsB | A0A0K2GGX0 | Formate dehydrogenase, beta subunit | 56.395 | 528 | 0.338 | 1.26 | 7.32E-02 | 9.83E-02 | no | no | 0 | 6 | 6 | 5 | 4 | 3 | 5 | 0.00235218 | 0.002158218 | 0.002377564 | 0.001405124 | 0.001520463 | 0.00160067 |
| NITMOv2_2469 | A0A0K2GE55 | Putative Response regulator, CheY like (Modular protein) | 42.005 | 382 | -0.618 | -1.53 | 8.79E-03 | 1.60E-02 | no | no | 0 | 2 | 0 | 2 | 3 | 3 | 3 | 0.002347332 | 0.002233698 | 0.002318513 | 0.003827875 | 0.003664046 | 0.004305781 |
| sdhA | A0A0K2GJI6 | Succinate dehydrogenase or fumarate reductase, flavoprotein subunit | 71.022 | 636 | -0.00307 | -1.00 | 9.78E-01 | 9.80E-01 | no | no | 0 | 3 | 4 | 5 | 7 | 3 | 5 | 0.00234691 | 0.002396 | 0.002512346 | 0.003556455 | 0.002819931 | 0.002601522 |
| nadE | A0A0K2G9R3 | Glutamine-dependent NAD(+) synthetase | 64.659 | 590 | 0.397 | 1.32 | 3.79E-03 | 7.98E-03 | no | no | 0 | 11 | 11 | 12 | 9 | 7 | 7 | 0.002337003 | 0.003249744 | 0.003571235 | 0.001633262 | 0.002010463 | 0.001907987 |
| NITMOv2_1386 | A0A0K2GA34 | Uncharacterized protein | 27.132 | 249 | -2.82 | -7.06 | 3.94E-07 | 1.61E-05 | yes | no | 0 | 1 | 1 | 2 | 17 | 16 | 16 | 0.002328783 | 0.002183639 | 0.001926055 | 0.032169196 | 0.025853254 | 0.021757268 |
| NITMOv2_3869 | A0A0K2GH40 | Maltokinase | 61.028 | 558 | 0.175 | 1.13 | 1.00E-01 | 1.30E-01 | no | no | 0 | 13 | 11 | 13 | 6 | 8 | 8 | 0.002326886 | 0.002270069 | 0.002554333 | 0.002060013 | 0.001955463 | 0.00178771 |
| NITMOv2_3256 | A0A0K2GFC2 | Putative Sensor histidine kinase | 78.303 | 711 | 0.267 | 1.20 | 2.23E-02 | 3.49E-02 | no | no | 0 | 9 | 7 | 8 | 3 | 3 | 4 | 0.002323934 | 0.004400911 | 0.004421529 | 0.001428016 | 0.00163354 | 0.00156548 |
| fdsA | A0A0K2GHV6 | Formate dehydrogenase, alpha subunit | 99.835 | 916 | 0.738 | 1.67 | 4.08E-05 | 3.03E-04 | no | no | 0 | 12 | 11 | 9 | 6 | 6 | 5 | 0.002318032 | 0.002445668 | 0.002813927 | 0.001597872 | 0.001380508 | 0.001259386 |
| apaH | A0A0K2G916 | Ap4A hydrolase | 29.934 | 269 | -0.944 | -1.92 | 3.14E-03 | 6.87E-03 | no | yes | 2 | 3 | 4 | 3 | 5 | 2 | 3 | 0.002316557 | 0.001259811 | 0.001093515 | 0.00243094 | 0.001400581 | 0.001274724 |
| NITMOv2_1424 | A0A0K2GA54 | Uncharacterized protein | 22.697 | 210 | 0.795 | 1.74 | 1.01E-01 | 1.30E-01 | no | yes | 2 | 2 | 1 | 2 | 2 | 0 | 1 | 0.002312552 | 0.003638877 | 0.003162481 | 0.001778671 | 0.001239496 | 0.000814675 |
| NITMOv2_3496 | A0A0K2GG07 | Uncharacterized protein | 27.747 | 241 | -1.92 | -3.78 | 2.46E-06 | 4.80E-05 | yes | no | 0 | 5 | 7 | 7 | 14 | 13 | 16 | 0.002311709 | 0.00245916 | 0.002512154 | 0.008399381 | 0.008790728 | 0.008366838 |
| NITMOv2_0613 | A0A0K2G868 | Uncharacterized protein | 39.204 | 358 | 0.0779 | 1.06 | 2.61E-01 | 3.03E-01 | no | no | 0 | 2 | 2 | 2 | 1 | 2 | 4 | 0.002310023 | 0.0008592 | 0.002368553 | 0.000232196 | 0.001870321 | 0.002557108 |
| nadD | A0A0K2G904 | Probable nicotinate-nucleotide adenylyltransferase | 25.524 | 228 | -0.86 | -1.82 | 6.03E-03 | 1.16E-02 | no | no | 0 | 6 | 4 | 6 | 5 | 6 | 5 | 0.002308969 | 0.002801949 | 0.003416323 | 0.002692736 | 0.003356417 | 0.003649287 |
| NITMOv2_1443 | A0A0K2GA99 | Uncharacterized protein | 90.986 | 813 | 0.0286 | 1.02 | 7.43E-01 | 7.74E-01 | no | no | 0 | 8 | 9 | 9 | 8 | 8 | 7 | 0.002307704 | 0.00262635 | 0.002630256 | 0.002292934 | 0.002036565 | 0.002102723 |
| NITMOv2_4039 | A0A0K2GHJ5 | Response regulatory domain-containing protein | 16.289 | 152 | -1.85 | -3.61 | 3.76E-06 | 6.29E-05 | yes | no | 0 | 0 | 0 | 1 | 2 | 2 | 2 | 0.002300537 | 0.002074134 | 0.002611659 | 0.008242126 | 0.008385217 | 0.007119482 |
| NITMOv2_1029 | A0A0K2G952 | Uncharacterized protein | 24.582 | 228 | 0.601 | 1.52 | 1.68E-01 | 2.04E-01 | no | yes | 1 | 3 | 1 | 1 | 0 | 2 | 1 | 0.002299483 | 0.001171914 | 0.000812716 | 0.000527071 | 0.000616858 | 0.00069073 |
| NITMOv2_1094 | A0A0K2G9B0 | Putative 2-hydroxy-6-ketonona-2,4-dienedioate hydrolase | 32.742 | 294 | 0.561 | 1.48 | 5.50E-04 | 1.83E-03 | no | no | 0 | 7 | 6 | 5 | 3 | 3 | 3 | 0.002298218 | 0.00263163 | 0.002169736 | 0.002890859 | 0.002773787 | 0.002553872 |
| NITMOv2_0218 | A0A0K2G7R4 | Uncharacterized protein | 26.136 | 232 | 0.269 | 1.20 | 2.16E-02 | 3.40E-02 | no | no | 0 | 3 | 2 | 3 | 1 | 2 | 2 | 0.002272292 | 0.002257359 | 0.003701799 | 0.001967615 | 0.001579006 | 0.001909425 |
| NITMOv2_0725 | A0A0K2G862 | Putative Surface antigen D15 | 44.186 | 400 | 0.191 | 1.14 | 2.03E-02 | 3.22E-02 | no | no | 0 | 8 | 9 | 8 | 7 | 6 | 6 | 0.002268076 | 0.001981055 | 0.002494324 | 0.001670873 | 0.001894403 | 0.001773973 |
| yloB | A0A0K2GI47 | Calcium-transporting ATPase | 99.174 | 924 | 0.421 | 1.34 | 7.84E-04 | 2.37E-03 | no | no | 0 | 7 | 7 | 8 | 3 | 3 | 3 | 0.002266811 | 0.00221825 | 0.002813544 | 0.001575217 | 0.001925011 | 0.001977754 |
| NITMOv2_3515 | A0A0K2GH28 | Methyltransf_11 domain-containing protein | 23.847 | 206 | 0.368 | 1.29 | 7.89E-03 | 1.45E-02 | no | no | 0 | 2 | 3 | 3 | 4 | 4 | 2 | 0.002265968 | 0.002452707 | 0.003798044 | 0.002903446 | 0.002921697 | 0.002933815 |
| NITMOv2_0239 | A0A0K2G6V2 | Uncharacterized protein | 43.849 | 408 | -0.859 | -1.81 | 2.99E-04 | 1.16E-03 | no | no | 0 | 3 | 4 | 9 | 10 | 11 | 9 | 0.002257958 | 0.002047931 | 0.002464415 | 0.006080237 | 0.005422036 | 0.004552304 |
| NITMOv2_0150 | A0A0K2G6N3 | Uncharacterized protein | 11.264 | 103 | 0.4 | 1.32 | 1.18E-01 | 1.50E-01 | no | no | 0 | 1 | 2 | 2 | 2 | 3 | 0 | 0.00225585 | 0.003328353 | 0.001856459 | 0.004343619 | 0.002041226 | 0.003688486 |
| NITMOv2_2214 | A0A0K2GCE6 | Uncharacterized protein | 10.291 | 93 | 1.46 | 2.75 | 1.63E-02 | 2.68E-02 | yes | yes | 3 | 3 | 1 | 1 | 2 | 3 | 1 | 0.002247419 | 0.00187597 | 0.00139941 | 0.001752758 | 0.001102989 | 0.000261788 |
| scpB | A0A0K2G9Y9 | Segregation and condensation protein B | 30.862 | 283 | -2.72 | -6.59 | 5.60E-08 | 6.98E-06 | yes | no | 0 | 0 | 0 | 0 | 9 | 12 | 12 | 0.002243835 | 0.001356781 | 0.001118669 | 0.01313747 | 0.012349124 | 0.010899492 |
| NITMOv2_1479 | A0A0K2GAC8 | Uncharacterized protein | 22.351 | 196 | 1.23 | 2.35 | 1.57E-02 | 2.60E-02 | yes | yes | 1 | 1 | 2 | 1 | 1 | 0 | 1 | 0.002238776 | 0.002395804 | 0.002249493 | 0.000525842 | 0.000701844 | 0.001078837 |
| NITMOv2_1641 | A0A088ND26 | Uncharacterized protein | 12.252 | 104 | 0.646 | 1.56 | 1.33E-02 | 2.26E-02 | no | no | 0 | 2 | 2 | 0 | 0 | 0 | 0 | 0.002232242 | 0.002356109 | 0.00162758 | 0.001283022 | 0.001219609 | 0.001060136 |
| bioA | A0A0K2GFS8 | Adenosylmethionine-8-amino-7-oxononanoate aminotransferase | 50.585 | 455 | -0.418 | -1.34 | 4.66E-03 | 9.37E-03 | no | no | 0 | 3 | 5 | 7 | 8 | 7 | 8 | 0.002227394 | 0.001964434 | 0.002402297 | 0.002954087 | 0.002810143 | 0.002906843 |
| ftsK | A0A0K2G7I9 | DNA translocase FtsK 2 | 85.76 | 809 | -1.18 | -2.27 | 2.70E-06 | 5.04E-05 | yes | no | 0 | 8 | 7 | 7 | 18 | 15 | 13 | 0.002221492 | 0.002436868 | 0.002680296 | 0.006010346 | 0.00677001 | 0.006528799 |
| NITMOv2_1356 | A0A0K2GA84 | Uncharacterized protein | 30.614 | 275 | 0.106 | 1.08 | 3.30E-01 | 3.72E-01 | no | no | 0 | 2 | 2 | 4 | 1 | 4 | 4 | 0.002208212 | 0.002444495 | 0.002386575 | 0.001661841 | 0.00208706 | 0.002401751 |
| NITMOv2_0634 | A0A0K2G814 | Putative Esterase, beta-lactamase family | 42.06 | 383 | -0.167 | -1.12 | 3.03E-01 | 3.45E-01 | no | no | 0 | 6 | 4 | 5 | 6 | 5 | 4 | 0.002201678 | 0.002427873 | 0.002119504 | 0.002357643 | 0.002344194 | 0.001364756 |
| ndh | A0A0K2GB79 | NADH dehydrogenase | 45.843 | 418 | 0.486 | 1.40 | 1.29E-02 | 2.21E-02 | no | no | 0 | 2 | 1 | 1 | 1 | 1 | 1 | 0.002200202 | 0.002101902 | 0.002302792 | 0.001460178 | 0.001504647 | 0.001294036 |
| NITMOv2_2034 | A0A0K2GBX2 | Putative Acetyltransferase, GNAT family | 17.884 | 159 | 0.3 | 1.23 | 9.43E-02 | 1.23E-01 | no | no | 0 | 2 | 2 | 0 | 0 | 2 | 2 | 0.00219704 | 0.001203905 | 0.000995793 | 0.001606757 | 0.001302 | 0.001566757 |
| NITMOv2_1071 | A0A0K2GA91 | Putative Amidohydrolase | 36.834 | 323 | -0.0405 | -1.03 | 7.10E-01 | 7.44E-01 | no | no | 0 | 3 | 2 | 4 | 3 | 3 | 3 | 0.00219683 | 0.002400302 | 0.002131199 | 0.002256952 | 0.002057385 | 0.001902772 |
| NITMOv2_1320 | A0A0K2G9X2 | TPR_REGION domain-containing protein | 31.067 | 274 | -0.453 | -1.37 | 2.16E-02 | 3.40E-02 | no | no | 0 | 2 | 2 | 2 | 3 | 3 | 3 | 0.002194722 | 0.002463462 | 0.001995459 | 0.003777678 | 0.003834174 | 0.002868723 |
| NITMOv2_3652 | A0A0K2GGS8 | PCP_red domain-containing protein | 18.621 | 164 | -0.985 | -1.98 | 3.60E-04 | 1.32E-03 | no | no | 0 | 2 | 3 | 4 | 8 | 8 | 6 | 0.002190717 | 0.002211993 | 0.002308544 | 0.007508268 | 0.005791036 | 0.004958499 |
| ybgJ | A0A0K2GCN7 | Putative hydrolase subunit | 26.351 | 242 | 1.13 | 2.19 | 8.90E-02 | 1.17E-01 | no | yes | 1 | 1 | 2 | 2 | 1 | 1 | 1 | 0.00218629 | 0.002536791 | 0.002580791 | 0.000564919 | 0.001084858 | 0.001406293 |
| NITMOv2_4280 | A0A0K2GJ44 | Tn7-like transposition protein A | 31.533 | 277 | 0.837 | 1.79 | 9.26E-05 | 5.38E-04 | no | no | 0 | 6 | 6 | 4 | 5 | 2 | 3 | 0.002182074 | 0.002244258 | 0.002397503 | 0.001415593 | 0.001355587 | 0.001301336 |
| NITMOv2_4550 | A0A0K2GJ96 | Hydrolase_4 domain-containing protein | 29.036 | 254 | 0.53 | 1.44 | 2.43E-02 | 3.76E-02 | no | no | 0 | 2 | 2 | 3 | 3 | 2 | 3 | 0.002181442 | 0.002780243 | 0.002926085 | 0.001509176 | 0.001515398 | 0.00240265 |
| ackA | A0A0K2GDC4 | Acetate kinase | 42.976 | 401 | 0.0915 | 1.07 | 3.78E-01 | 4.20E-01 | no | no | 0 | 6 | 4 | 3 | 3 | 2 | 3 | 0.002180599 | 0.002175622 | 0.002260613 | 0.00209022 | 0.001690094 | 0.001664593 |
| NITMOv2_4357 | A0A0K2GIE3 | Putative transcription regulator with HTH domain | 46.75 | 413 | -0.389 | -1.31 | 1.75E-02 | 2.84E-02 | no | no | 0 | 3 | 3 | 4 | 8 | 3 | 5 | 0.002179756 | 0.002754432 | 0.002884673 | 0.003318943 | 0.003202914 | 0.003192025 |
| nuoL | A0A0K2G6W3 | NADH-quinone oxidoreductase, membrane subunit L | 71.378 | 657 | -1.28 | -2.43 | 2.24E-01 | 2.66E-01 | no | yes | 1 | 3 | 5 | 3 | 3 | 6 | 5 | 0.002174486 | 0.007258981 | 0.003954682 | 0.008005206 | 0.006129738 | 0.008765122 |
| NITMOv2_2313 | A0A0K2GCQ3 | DNA_pol_B_exo2 domain-containing protein | 27.182 | 237 | -0.814 | -1.76 | 8.66E-03 | 1.58E-02 | no | no | 0 | 1 | 2 | 2 | 6 | 3 | 7 | 0.002166265 | 0.00161267 | 0.002414183 | 0.008199924 | 0.005141131 | 0.007286707 |
| NITMOv2_0243 | A0A0K2G7T9 | Cytokinin riboside 5-monophosphate phosphoribohydrolase | 31.525 | 284 | -0.0822 | -1.06 | 4.79E-01 | 5.22E-01 | no | no | 0 | 7 | 10 | 11 | 7 | 11 | 9 | 0.002143711 | 0.001974602 | 0.00185784 | 0.002338689 | 0.00233394 | 0.002582102 |
| NITMOv2_1850 | A0A0K2GBE2 | Putative Response regulator | 46.178 | 428 | 0.104 | 1.07 | 4.50E-01 | 4.93E-01 | no | no | 0 | 7 | 5 | 4 | 4 | 4 | 4 | 0.002134015 | 0.001986922 | 0.001983189 | 0.002230595 | 0.003537887 | 0.001924349 |
| cysH | A0A0K2G739 | Phosphoadenosine phosphosulfate reductase | 27.064 | 238 | -2.12 | -4.35 | 6.99E-06 | 9.34E-05 | yes | no | 0 | 1 | 2 | 2 | 9 | 7 | 12 | 0.002129799 | 0.002311134 | 0.002259462 | 0.010002436 | 0.009329389 | 0.010363652 |
| ybaB | A0A0K2G6T8 | Nucleoid-associated protein NITMOv2_0205 | 11.061 | 107 | -2.05 | -4.14 | 1.90E-03 | 4.62E-03 | yes | yes | 1 | 2 | 2 | 4 | 8 | 8 | 6 | 0.002129799 | 0.001321271 | 0.002502376 | 0.003623384 | 0.003200272 | 0.002590194 |
| NITMOv2_1391 | A0A0K2GA39 | Uncharacterized protein | 11.561 | 106 | 0.961 | 1.95 | 1.01E-02 | 1.80E-02 | no | yes | 3 | 3 | 5 | 2 | 0 | 0 | 0 | 0.002123897 | 0.003047943 | 0.003143692 | 0.000150133 | 8.143464E-05 | 0 |
| ykoU | A0A0K2GDN5 | 3-phosphoesterase | 99.667 | 895 | 1.12 | 2.17 | 3.06E-04 | 1.18E-03 | yes | no | 0 | 7 | 11 | 10 | 4 | 5 | 3 | 0.002123054 | 0.002288646 | 0.001912654 | 0.000320537 | 0.000508038 | 0.000527623 |
| NITMOv2_0263 | A0A0K2G7W1 | SWIM-type domain-containing protein | 33.341 | 302 | -1.2 | -2.30 | 3.13E-05 | 2.54E-04 | yes | no | 0 | 2 | 2 | 2 | 6 | 8 | 7 | 0.002122 | 0.001827162 | 0.002325991 | 0.006427916 | 0.00588783 | 0.005452803 |
| NITMOv2_1883 | A0A0K2GBH7 | Type III restriction protein res subunit | 106.05 | 944 | -0.286 | -1.22 | 1.33E-02 | 2.26E-02 | no | no | 0 | 9 | 9 | 12 | 14 | 14 | 13 | 0.002113358 | 0.00214981 | 0.002365869 | 0.002623289 | 0.002911132 | 0.002767309 |
| NITMOv2_3982 | A0A0K2GHF4 | Uncharacterized protein | 25.499 | 237 | 0.92 | 1.89 | 7.58E-02 | 1.01E-01 | no | yes | 2 | 3 | 2 | 3 | 1 | 1 | 1 | 0.002109985 | 0.002289624 | 0.002327141 | 0.001498514 | 0.000612368 | 0.00059543 |
| cobD.1 | A0A0K2GBD4 | L-threonine-O-3-phosphate decarboxylase | 40.999 | 372 | -1.02 | -2.03 | 1.20E-04 | 6.31E-04 | yes | no | 0 | 7 | 6 | 7 | 9 | 10 | 10 | 0.002108088 | 0.001977536 | 0.002249876 | 0.004095298 | 0.00392724 | 0.004098458 |
| czcA.3 | A0A0K2GHG2 | Cation efflux system protein CzcA | 114.36 | 1044 | 0.424 | 1.34 | 8.81E-04 | 2.58E-03 | no | no | 0 | 8 | 10 | 12 | 5 | 4 | 4 | 0.002084185 | 0.002121456 | 0.00243048 | 0.001465375 | 0.00161878 | 0.001428194 |
| NITMOv2_2683 | A0A0K2GDR0 | Histidine kinase | 79.796 | 723 | -0.56 | -1.47 | 7.71E-03 | 1.42E-02 | no | no | 0 | 2 | 2 | 2 | 7 | 7 | 3 | 0.002079105 | 0.00190364 | 0.001974178 | 0.00222097 | 0.002004404 | 0.001757861 |

| | | | | | | | | | | | | | | | | | | | | | | | |
|---|---|---|---|---|---|---|---|---|---|---|---|---|---|---|---|---|---|---|---|---|---|---|---|
| NITMOv2_1523 | A0A0K2GAI2 | Histidine kinase | 56.736 | 512 | -0.272 | -1.21 | 1.91E-02 | 3.06E-02 | no | no | 0 | 6 | 5 | 7 | 13 | 11 | 10 | 0.002075205 | 0.001975385 | 0.001940818 | 0.003132665 | 0.00308297 | 0.002729189 |
| dnaJ | A0A0K2GD60 | Chaperone protein DnaJ | 41.214 | 375 | -0.514 | -1.43 | 3.78E-03 | 7.96E-03 | no | no | 0 | 5 | 5 | 5 | 7 | 8 | 7 | 0.002068397 | 0.00152448 | 0.001963058 | 0.004632512 | 0.005135848 | 0.004994282 |
| NITMOv2_1400 | A0A0K2GB61 | Uncharacterized protein | 12.361 | 109 | 0.0447 | 1.03 | 6.48E-01 | 6.86E-01 | no | no | 0 | 2 | 5 | 1 | 5 | 2 | 3 | 0.002068186 | 0.005787122 | 0.002024984 | 0.003803887 | 0.005179973 | 0.005333588 |
| NITMOv2_2991 | A0A0K2GEL7 | Putative Universal stress protein | 32.385 | 296 | -0.868 | -1.83 | 1.99E-02 | 3.17E-02 | no | no | 0 | 5 | 7 | 9 | 12 | 10 | 9 | 0.002065424 | 0.002827565 | 0.003487069 | 0.003736957 | 0.004542031 | 0.003422005 |
| NITMOv2_3278 | A0A0K2GFE2 | Putative Transcriptional regulator, Crp/Fnr family | 26.378 | 239 | 0.612 | 1.53 | 1.69E-04 | 7.81E-04 | no | no | 0 | 2 | 1 | 2 | 0 | 0 | 1 | 0.002061061 | 0.002132015 | 0.001530529 | 0.001250105 | 0.001314834 | 0.001947365 |
| NITMOv2_1005 | A0A0K2GA36 | DctA-YdbH domain-containing protein | 100.14 | 933 | 0.198 | 1.15 | 2.37E-02 | 3.68E-02 | no | no | 0 | 13 | 14 | 16 | 16 | 16 | 13 | 0.002058742 | 0.002245431 | 0.002445243 | 0.002346241 | 0.002348389 | 0.002166377 |
| nuoM | A0A0K2G6M5 | NADH-quinone oxidoreductase, membrane subunit M | 59.172 | 534 | -0.138 | -1.10 | 4.29E-01 | 4.72E-01 | no | no | 0 | 5 | 4 | 4 | 5 | 5 | 5 | 0.002056571 | 0.000986343 | 0.001207667 | 0.003147325 | 0.003112489 | 0.003222593 |
| rbfA | A0A0K2GIT8 | Ribosome-binding factor A | 16.076 | 145 | -1.05 | -2.07 | 2.56E-05 | 2.23E-04 | yes | no | 0 | 2 | 2 | 2 | 2 | 3 | 5 | 0.002053009 | 0.001945877 | 0.002547431 | 0.005462322 | 0.004900776 | 0.004893587 |
| NITMOv2_3599 | A0A0K2GGC5 | Hist_deacetyl domain-containing protein | 33.749 | 314 | 0.355 | 1.28 | 4.41E-03 | 8.97E-03 | no | no | 0 | 3 | 5 | 8 | 5 | 7 | 4 | 0.002033216 | 0.005002991 | 0.004396989 | 0.003314501 | 0.003154439 | 0.002973194 |
| NITMOv2_4232 | A0A0K2GI50 | DNA-binding transcriptional regulator, tetR-type | 21.303 | 195 | -0.725 | -1.65 | 3.94E-03 | 8.20E-03 | no | no | 0 | 1 | 0 | 4 | 6 | 4 | 5 | 0.002008744 | 0.00153457 | 0.003056457 | 0.009375044 | 0.01117221 | 0.010092855 |
| NITMOv2_1246 | A0A0K2G9X9 | Putative Type IV pili biogenesis protein PilQ | 70.352 | 656 | -0.735 | -1.66 | 8.74E-04 | 2.57E-03 | no | no | 0 | 11 | 15 | 14 | 24 | 25 | 29 | 0.00200164 | 0.003068671 | 0.001845244 | 0.004938138 | 0.004949251 | 0.005843355 |
| dnaC | A0A0K2GGT0 | Replicative DNA helicase | 51.352 | 466 | -1.47 | -2.77 | 2.65E-06 | 5.00E-05 | yes | no | 0 | 1 | 2 | 3 | 18 | 18 | 21 | 0.001990405 | 0.001913827 | 0.001874175 | 0.005576932 | 0.005060806 | 0.005081131 |
| NITMOv2_3916 | A0A0K2GI58 | PilZ domain-containing protein | 15.343 | 139 | -0.233 | -1.18 | 7.58E-02 | 1.01E-01 | no | no | 0 | 1 | 3 | 2 | 2 | 1 | 3 | 0.001987918 | 0.001716035 | 0.001364593 | 0.002523338 | 0.00237822 | 0.002759217 |
| NITMOv2_4131 | A0A0K2GHS1 | Hemoglobin-like protein HbN (Modular protein) (Modular protein) | 33.067 | 306 | -1.13 | -2.19 | 2.30E-02 | 3.58E-02 | yes | no | 0 | 1 | 2 | 3 | 6 | 3 | 2 | 0.001987328 | 0.00203131 | 0.002107042 | 0.001285805 | 0.000985407 | 0.001236119 |
| NITMOv2_3217 | A0A0K2GFI6 | Putative Proline iminopeptidase | 31.259 | 278 | -0.128 | -1.09 | 6.99E-01 | 7.34E-01 | no | no | 0 | 2 | 1 | 0 | 3 | 1 | 1 | 0.001981404 | 0.002037959 | 0.001521825 | 0.00263321 | 0.002360042 | 0.002292065 |
| uvrB | A0A0K2GG86 | UvrABC system protein B | 74.919 | 665 | 0.0224 | 1.02 | 8.27E-01 | 8.47E-01 | no | no | 0 | 6 | 5 | 5 | 6 | 5 | 3 | 0.001969874 | 0.00224504 | 0.001932765 | 0.001834496 | 0.001749445 | 0.001708449 |
| NITMOv2_3504 | A0A0K2GG22 | Uncharacterized protein | 40.937 | 350 | -0.00407 | -1.00 | 9.64E-01 | 9.68E-01 | no | no | 0 | 6 | 5 | 3 | 4 | 8 | 5 | 0.001967366 | 0.001763122 | 0.00170912 | 0.002122353 | 0.002823194 | 0.001977754 |
| NITMOv2_0312 | A0A0K2G738 | DUF2726 domain-containing protein | 18.301 | 169 | 0.0793 | 1.06 | 3.32E-01 | 3.74E-01 | no | no | 0 | 2 | 2 | 3 | 1 | 1 | 1 | 0.001950545 | 0.002346723 | 0.002184115 | 0.000975796 | 0.001659642 | 0.001346703 |
| acrB | A0A0K2G7X4 | Acriflavin resistance protein acrB | 116.84 | 1062 | 0.33 | 1.26 | 2.64E-02 | 4.03E-02 | no | no | 0 | 12 | 14 | 10 | 8 | 5 | 6 | 0.00194907 | 0.002196936 | 0.001780345 | 0.002101178 | 0.001557721 | 0.000912187 |
| uvrC | A0A0K2G7N2 | UvrABC system protein C | 68.317 | 607 | 0.132 | 1.10 | 4.16E-01 | 4.59E-01 | no | no | 0 | 5 | 8 | 8 | 3 | 6 | 3 | 0.001947109 | 0.001875071 | 0.001604515 | 0.001097973 | 0.001045472 | 0.001173742 |
| NITMOv2_3665 | A0A0K2GHG4 | Uncharacterized protein | 59.451 | 520 | 0.741 | 1.67 | 3.23E-04 | 1.22E-03 | no | no | 0 | 4 | 7 | 7 | 3 | 5 | 4 | 0.001944959 | 0.002396586 | 0.002708287 | 0.001489778 | 0.00156145 | 0.001610146 |
| NITMOv2_3137 | A0A0K2GF06 | Uncharacterized protein | 41.898 | 359 | -0.6 | -1.52 | 3.53E-03 | 7.56E-03 | no | no | 0 | 1 | 1 | 3 | 2 | 3 | 3 | 0.001939521 | 0.00188827 | 0.001872468 | 0.003104975 | 0.003259468 | 0.003205151 |
| NITMOv2_0209 | A0A0K2G6S3 | Uncharacterized protein | 27.722 | 253 | 0.659 | 1.58 | 6.00E-05 | 4.00E-04 | no | yes | 1 | 2 | 2 | 1 | 2 | 0 | 1 | 0.001939226 | 0.00196463 | 0.001562279 | 0.001500143 | 0.000871258 | 0.00140088 |
| NITMOv2_2998 | A0A0K2GEN2 | Uncharacterized protein | 33.873 | 306 | -0.0902 | -1.06 | 5.77E-01 | 6.18E-01 | no | no | 0 | 7 | 6 | 6 | 5 | 6 | 5 | 0.001935853 | 0.002008236 | 0.002469016 | 0.001992195 | 0.001904502 | 0.001918416 |
| NITMOv2_0701 | A0A0K2G951 | Uncharacterized protein | 34.966 | 321 | 0.727 | 1.66 | 2.45E-04 | 1.01E-03 | no | no | 0 | 3 | 2 | 2 | 2 | 2 | 1 | 0.001926663 | 0.001709191 | 0.001948295 | 0.001606609 | 0.002155266 | 0.000967479 |
| ispG | A0A0K2G879 | 4-hydroxy-3-methylbut-2-en-1-yl diphosphate synthase (flavodoxin) | 42.452 | 394 | -1.85 | -3.61 | 9.53E-07 | 2.93E-05 | yes | no | 0 | 4 | 2 | 4 | 21 | 22 | 24 | 0.001918 | 0.002381725 | 0.00242607 | 0.012374738 | 0.012544732 | 0.011496648 |
| thiI | A0A0K2GA22 | Probable tRNA sulfurtransferase | 43.46 | 395 | -0.362 | -1.29 | 1.53E-02 | 2.56E-02 | no | no | 0 | 3 | 4 | 4 | 5 | 6 | 3 | 0.001914901 | 0.001878454 | 0.002889083 | 0.003129556 | 0.003156459 | 0.002298718 |
| NITMOv2_2985 | A0A0K2GEK3 | Uncharacterized protein | 20.487 | 180 | 0.691 | 1.61 | 4.36E-03 | 8.89E-03 | no | no | 0 | 2 | 3 | 2 | 1 | 1 | 1 | 0.001913952 | 0.001878512 | 0.002085377 | 0.001147134 | 0.000901353 | 0.000708945 |
| era | A0A0K2GFT4 | GTPase Era | 32.802 | 294 | -1.35 | -2.55 | 1.26E-05 | 1.39E-04 | yes | no | 0 | 3 | 1 | 1 | 7 | 7 | 8 | 0.001909104 | 0.002179141 | 0.002525383 | 0.007441635 | 0.005976701 | 0.006524303 |
| NITMOv2_4503 | A0A0K2GJS8 | Uncharacterized protein | 44.367 | 414 | 1.34 | 2.53 | 5.03E-02 | 7.06E-02 | no | yes | 2 | 3 | 2 | 2 | 1 | 1 | 0 | 0.001908177 | 0.001759485 | 0.001601544 | 0.000263913 | 0.000915336 | 0.00066989 |
| tatA | A0A0K2G6Z2 | Sec-independent protein translocase protein TatA | 27.951 | 272 | 0.702 | 1.63 | 2.47E-02 | 3.81E-02 | no | no | 0 | 4 | 3 | 2 | 0 | 0 | 0 | 0.001904678 | 0.001848144 | 0.00122822 | 0.001052973 | 0.001495309 | 0.001144235 |
| NITMOv2_3535 | A0A0K2GH48 | Uncharacterized protein | 30.221 | 280 | -0.588 | -1.50 | 8.90E-05 | 5.23E-04 | no | no | 0 | 0 | 2 | 1 | 5 | 4 | 7 | 0.001897384 | 0.001901058 | 0.001531814 | 0.005185423 | 0.004993842 | 0.005938835 |
| NITMOv2_4044 | A0A0K2GHJ8 | Uncharacterized protein | 15.146 | 136 | -1.87 | -3.66 | 6.57E-06 | 8.94E-05 | yes | no | 0 | 2 | 1 | 1 | 6 | 4 | 5 | 0.00189319 | 0.001955341 | 0.00219466 | 0.006361283 | 0.006258383 | 0.005161148 |
| NITMOv2_4844 | A0A0K2GKR1 | XRE family transcriptional regulator | 13.514 | 122 | -1.06 | -2.08 | 1.29E-03 | 3.42E-03 | yes | no | 0 | 6 | 5 | 3 | 6 | 6 | 2 | 0.001891546 | 0.001885141 | 0.00127617 | 0.003778122 | 0.00425025 | 0.002244775 |
| NITMOv2_3141 | A0A0K2GF04 | Putative PIN domain protein | 15.294 | 136 | -0.42 | -1.34 | 2.37E-01 | 2.78E-01 | no | no | 0 | 0 | 2 | 3 | 2 | 2 | 3 | 0.001891398 | 0.002440388 | 0.00251944 | 0.002114949 | 0.002106481 | 0.002088878 |
| NITMOv2_4110 | A0A0K2GIQ7 | DNA-3-methyladenine glycosylase I | 25.261 | 222 | 0.702 | 1.63 | 5.51E-04 | 1.84E-03 | no | no | 0 | 3 | 3 | 4 | 2 | 2 | 1 | 0.001882145 | 0.001827632 | 0.00230816 | 0.001326689 | 0.001231075 | 0.001338719 |
| NITMOv2_1021 | A0A0K2G926 | Putative Glycosyl hydrolase, family 13 | 24.776 | 226 | 0.161 | 1.12 | 4.26E-01 | 4.69E-01 | no | no | 0 | 2 | 1 | 2 | 3 | 3 | 1 | 0.001879594 | 0.001756063 | 0.002095155 | 0.001568998 | 0.003883115 | 0.002243696 |
| NITMOv2_3949 | A0A0K2GHJ0 | Uncharacterized protein | 20.902 | 193 | -0.202 | -1.15 | 5.19E-01 | 5.61E-01 | no | no | 0 | 10 | 10 | 11 | 5 | 5 | 5 | 0.001865408 | 0.001510577 | 0.002385617 | 0.001432458 | 0.001585687 | 0.001263756 |
| NITMOv2_2534 | A0A0K2GED2 | Oxidoreductase, Glucose/ribitol dehydrogenase family | 29.944 | 287 | -0.456 | -1.37 | 5.67E-02 | 7.81E-02 | no | no | 0 | 3 | 4 | 2 | 5 | 3 | 4 | 0.001864902 | 0.001706864 | 0.001534939 | 0.002411986 | 0.001856183 | 0.001670527 |
| NITMOv2_2377 | A0A0K2GDW0 | Uncharacterized protein | 27.514 | 245 | -0.356 | -1.28 | 5.36E-02 | 7.47E-02 | no | no | 0 | 7 | 7 | 6 | 4 | 3 | 3 | 0.001863089 | 0.00227398 | 0.002586543 | 0.002411246 | 0.002810609 | 0.003268445 |
| NITMOv2_0119 | A0A0K2GJK8 | Putative Transposon Tn7 transposition protein TnsB | 83.928 | 729 | -0.44 | -1.36 | 9.00E-03 | 1.63E-02 | no | no | 0 | 3 | 5 | 4 | 4 | 7 | 5 | 0.001854995 | 0.001770943 | 0.001714699 | 0.00151732 | 0.001595786 | 0.001730278 |
| NITMOv2_3927 | A0A0K2GI69 | Putative NAD(P)H dehydrogenase (Quinone) | 21.729 | 193 | 0.533 | 1.45 | 3.27E-02 | 4.87E-02 | no | no | 0 | 2 | 3 | 2 | 2 | 3 | 1 | 0.001852908 | 0.000728518 | 0.000821267 | 0.000530981 | 0.000391466 | 0.000332796 |
| NITMOv2_0029 | A0A0K2G6B3 | Metallo-beta-lactamase superfamily hydrolase | 32.857 | 292 | -0.188 | -1.14 | 1.94E-01 | 2.33E-01 | no | no | 0 | 4 | 5 | 7 | 5 | 5 | 2 | 0.001851686 | 0.002903436 | 0.001974753 | 0.003540611 | 0.003172617 | 0.003254779 |
| truB | A0A0K2GIZ8 | tRNA pseudouridine synthase B | 34.701 | 325 | -1.25 | -2.38 | 2.08E-04 | 9.04E-04 | yes | no | 0 | 1 | 1 | 1 | 7 | 5 | 6 | 0.00184884 | 0.00166226 | 0.00156456 | 0.005901807 | 0.005643281 | 0.006265014 |
| NITMOv2_4758 | A0A0K2GKH1 | Uncharacterized protein | 24.661 | 226 | 0.234 | 1.18 | 5.31E-02 | 7.42E-02 | no | no | 0 | 4 | 4 | 4 | 1 | 2 | 4 | 0.001848566 | 0.001918149 | 0.002246617 | 0.001436886 | 0.00151978 | 0.001457755 |
| NITMOv2_3416 | A0A0K2GFS0 | Putative Polysaccharide export protein | 23.616 | 223 | -3.74 | -13.36 | 1.95E-08 | 3.98E-06 | yes | no | 0 | 4 | 2 | 3 | 24 | 28 | 22 | 0.001846901 | 0.001883851 | 0.002245658 | 0.033636615 | 0.033268975 | 0.031842931 |
| murA | A0A0K2G8H1 | UDP-N-acetylglucosamine 1-carboxyvinyltransferase | 45.153 | 427 | -0.604 | -1.52 | 9.47E-04 | 2.71E-03 | no | no | 0 | 4 | 3 | 4 | 8 | 10 | 10 | 0.001845805 | 0.00225071 | 0.003418623 | 0.005031573 | 0.005644368 | 0.005379799 |
| NITMOv2_3901 | A0A0K2GH46 | ATPase (AAA+ superfamily) | 42.988 | 390 | -0.187 | -1.14 | 4.96E-01 | 5.39E-01 | no | no | 0 | 8 | 13 | 13 | 5 | 9 | 8 | 0.0018422 | 0.002211211 | 0.003017729 | 0.002986368 | 0.002042159 | 0.001939454 |
| NITMOv2_2396 | A0A0K2GCX3 | Putative Thiamine biosynthesis protein ThiS | 7.3584 | 66 | -0.685 | -1.61 | 8.95E-02 | 1.17E-01 | no | yes | 1 | 1 | 1 | 0 | 2 | 3 | 1 | 0.001834422 | 0.005026261 | 0.004195871 | 0.007493313 | 0.008433226 | 0.007847361 |
| NITMOv2_1427 | A0A0K2GAE3 | ABC-type transport system periplasmic binding protein | 33.82 | 312 | -1.35 | -2.55 | 6.05E-03 | 1.16E-02 | yes | no | 0 | 7 | 7 | 9 | 9 | 11 | 11 | 0.001827614 | 0.001944919 | 0.00281201 | 0.005320171 | 0.004769801 | 0.004447473 |
| sthA | A0A0K2G993 | Putative soluble pyridine nucleotide transhydrogenase | 50.659 | 467 | -1.07 | -2.10 | 1.09E-03 | 3.01E-03 | yes | no | 0 | 3 | 4 | 2 | 6 | 6 | 6 | 0.001813554 | 0.001853756 | 0.001760157 | 0.00187877 | 0.00183179 | 0.001877958 |
| NITMOv2_2731 | A0A0K2GDW1 | Uncharacterized protein | 7.6635 | 73 | -0.277 | -1.21 | 3.71E-01 | 4.14E-01 | no | yes | 3 | 0 | 2 | 1 | 1 | 1 | 0 | 0.001812268 | 0.001272913 | 0.001922412 | 0.000661079 | 0.000801964 | 0.000501676 |
| NITMOv2_2894 | A0A0K2GEB0 | Histidine kinase | 85.301 | 780 | 0.64 | 1.56 | 1.44E-03 | 3.74E-03 | no | no | 0 | 8 | 7 | 9 | 5 | 4 | 8 | 0.001804216 | 0.002122434 | 0.001931999 | 0.002150487 | 0.001203202 | 0.001296445 |
| prfC | A0A0K2GBV2 | Peptide chain release factor 3 | 59.427 | 542 | -0.975 | -1.97 | 5.67E-06 | 8.00E-05 | no | no | 0 | 5 | 2 | 4 | 11 | 10 | 12 | 0.001803036 | 0.001833302 | 0.001987982 | 0.004844703 | 0.004652498 | 0.004411511 |
| mutS | A0A0K2GA17 | Endonuclease MutS2 | 88.12 | 809 | 0.337 | 1.26 | 7.75E-03 | 1.43E-02 | no | no | 0 | 6 | 9 | 9 | 6 | 5 | 6 | 0.00180234 | 0.003091158 | 0.002089212 | 0.001557152 | 0.001457897 | 0.001416704 |
| NITMOv2_4208 | A0A0K2GHZ1 | Putative Nitrite oxidoreductase, membrane subunit | 28.733 | 262 | 0.309 | 1.24 | 3.82E-03 | 8.01E-03 | no | no | 0 | 2 | 2 | 2 | 2 | 3 | 2 | 0.001798377 | 0.001755046 | 0.001591497 | 0.001252948 | 0.001467048 | 0.001438587 |
| NITMOv2_1126 | A0A0K2GAD0 | Uncharacterized protein | 23.24 | 208 | 1.39 | 2.62 | 7.64E-03 | 1.42E-02 | yes | yes | 3 | 2 | 2 | 1 | 0 | 0 | 0 | 0.001795911 | 0.002624004 | 0.001721774 | 0 | 6.868514E-05 | 0 |
| ffh | A0A0K2GG99 | Signal recognition particle protein | 48.865 | 448 | -1.45 | -2.73 | 1.17E-06 | 3.13E-05 | yes | no | 0 | 4 | 5 | 5 | 13 | 11 | 10 | 0.001792201 | 0.00184042 | 0.001872775 | 0.005942232 | 0.00896272 | 0.008605629 |
| rlmI | A0A0K2GG23 | Ribosomal RNA large subunit methyltransferase I | 43.779 | 396 | 0.0579 | 1.04 | 6.46E-01 | 6.84E-01 | no | no | 0 | 3 | 4 | 4 | 4 | 2 | 3 | 0.001790325 | 0.001713395 | 0.002202904 | 0.002344612 | 0.001466333 | 0.001948624 |
| bioD | A0A0K2GD19 | ATP-dependent dethiobiotin synthetase BioD | 27.995 | 257 | 0.126 | 1.09 | 9.07E-01 | 9.21E-01 | no | yes | 3 | 1 | 1 | 2 | 2 | 1 | 2 | 0.001784634 | 0.001789579 | 0.00214577 | 0.001952955 | 0.001602467 | 0.001692356 |
| NITMOv2_2264 | A0A0K2GCJ6 | DNA helicase | 124.89 | 1123 | -0.834 | -1.78 | 1.26E-04 | 6.50E-04 | no | no | 0 | 2 | 3 | 3 | 10 | 12 | 13 | 0.001770427 | 0.001728061 | 0.001676297 | 0.002874275 | 0.002086283 | 0.002902527 |
| selO | A0A0K2GK34 | Protein adenylyltransferase SelO | 56.992 | 497 | 0.03 | 1.02 | 8.16E-01 | 8.38E-01 | no | no | 0 | 4 | 4 | 3 | 2 | 1 | 2 | 0.001770111 | 0.001988486 | 0.001602617 | 0.001151753 | 0.000876136 | 0.00132063 |
| murB | A0A0K2G802 | UDP-N-acetylenolpyruvoylglucosamine reductase | 33.576 | 314 | -0.183 | -1.14 | 3.10E-01 | 3.52E-01 | no | no | 0 | 9 | 10 | 9 | 2 | 2 | 3 | 0.001766401 | 0.001892376 | 0.002478027 | 0.002209717 | 0.00234233 | 0.002114051 |
| NITMOv2_3195 | A0A0K2GFG9 | Putative HTH-type transcriptional regulator rrf2-like | 15.683 | 144 | -1.42 | -2.68 | 4.45E-02 | 6.35E-02 | no | yes | 2 | 0 | 0 | 1 | 3 | 2 | 4 | 0.001766232 | 0.002251102 | 0.00282313 | 0.004954426 | 0.00423347 | 0.004834069 |
| NITMOv2_1542 | A0A0K2GBK9 | Toxin-antitoxin system, toxin component, PIN family | 16.192 | 144 | 1.28 | 2.43 | 8.02E-02 | 1.07E-01 | no | yes | 2 | 2 | 2 | 2 | 0 | 0 | 1 | 0.00175114 | 0.002110701 | 0.002364719 | 0.000551148 | 0.00133121 | 0.001549243 |
| NITMOv2_0708 | A0A0K2G8G3 | Thiol-disulfide oxidoreductase ResA (Modular protein) | 20.853 | 185 | -2.39 | -5.24 | 1.56E-02 | 2.59E-02 | yes | yes | 1 | 1 | 1 | 1 | 4 | 3 | 1 | 0.001747198 | 0.001952428 | 0.001819917 | 0.006740057 | 0.009219233 | 0.003890595 |
| waaE | A0A0K2GH34 | Lipopolysaccharide core biosynthesis glycosyltransferase WaaE | 28.81 | 253 | -0.483 | -1.40 | 8.04E-04 | 2.41E-03 | no | no | 0 | 1 | 1 | 2 | 3 | 3 | 3 | 0.001741043 | 0.001634219 | 0.002325991 | 0.003833798 | 0.004054331 | 0.00278547 |
| NITMOv2_2743 | A0A0K2GDX8 | AMP-dependent synthetase and ligase | 98.636 | 900 | 0.409 | 1.33 | 2.30E-02 | 3.59E-02 | no | no | 0 | 3 | 4 | 6 | 4 | 3 | 0 | 0.001736216 | 0.001806532 | 0.001725282 | 0.000931093 | 0.001056115 | 0.000861426 |
| atoC.6 | A0A0K2GGS6 | Acetoacetate metabolism regulatory protein AtoC | 50.673 | 452 | -0.0568 | -1.04 | 7.81E-01 | 8.09E-01 | no | no | 0 | 5 | 5 | 7 | 7 | 7 | 8 | 0.001734277 | 0.002151179 | 0.00188192 | 0.001661248 | 0.001880886 | 0.001553379 |
| nadB | A0A0K2GI08 | L-aspartate oxidase | 61.581 | 562 | -0.275 | -1.21 | 9.84E-02 | 1.27E-01 | no | no | 0 | 6 | 5 | 6 | 4 | 4 | 4 | 0.001733855 | 0.001317829 | 0.001426941 | 0.001114927 | 0.001005589 | 0.001381227 |
| NITMOv2_0861 | A0A0K2G8J4 | SGL domain-containing protein | 42.398 | 407 | -0.3 | -1.23 | 1.65E-02 | 2.71E-02 | no | no | 0 | 2 | 3 | 2 | 4 | 4 | 3 | 0.001727173 | 0.00173643 | 0.001584576 | 0.002357643 | 0.002924339 | 0.00277576 |
| NITMOv2_4097 | A0A0K2GI04 | Penicillin-insensitive transglycosylase | 82.843 | 761 | -4 | -16.00 | 8.02E-06 | 1.01E-04 | yes | yes | 3 | 1 | 0 | 0 | 5 | 7 | 6 | 0.001724918 | 0 | 0 | 0.003778122 | 0.005095297 | 0.000969349 |
| NITMOv2_2497 | A0A0K2GD88 | Uncharacterized protein | 33.742 | 293 | -1.9 | -3.73 | 1.41E-05 | 1.47E-04 | yes | yes | 1 | 1 | 0 | 1 | 2 | 4 | 3 | 0.001703249 | 0.000310739 | 0.001893194 | 0.001559669 | 0.002066551 | 0.00210596 |
| NITMOv2_1065 | A0A0K2G954 | Uncharacterized protein | 26.527 | 239 | -0.0895 | -1.06 | 4.89E-01 | 5.32E-01 | no | no | 0 | 2 | 1 | 2 | 1 | 5 | 2 | 0.001701984 | 0.000877132 | 0.002226103 | 0.002287752 | 0.002481695 | 0.001764299 |
| NITMOv2_0452 | A0A0K2G7F5 | YCII domain-containing protein | 13.237 | 121 | 0.583 | 1.50 | 1.89E-02 | 3.04E-02 | no | yes | 2 | 1 | 1 | 2 | 1 | 1 | 1 | 0.00170131 | 0.001823017 | 0.001947336 | 0.001969392 | 0.001373796 | 0.001244911 |
| NITMOv2_1614 | A0A0K2GAR6 | Uncharacterized protein | 79.122 | 715 | -0.547 | -1.46 | 1.71E-03 | 4.26E-03 | no | no | 0 | 4 | 5 | 7 | 17 | 12 | 11 | 0.001701289 | 0.0021678 | 0.002546089 | 0.003441401 | 0.003239891 | 0.003068314 |
| NITMOv2_4784 | A0A0K2GKJ5 | PBP_domain domain-containing protein | 52.377 | 489 | 0.478 | 1.39 | 8.03E-03 | 1.48E-02 | no | no | 0 | 3 | 2 | 2 | 1 | 3 | 2 | 0.001697368 | 0.001304063 | 0.001682355 | 0.001529758 | 0.001450455 | 0.001446768 |
| NITMOv2_2658 | A0A0K2GDN9 | Uncharacterized protein | 27.24 | 246 | 1.16 | 2.23 | 1.09E-02 | 1.92E-02 | yes | yes | 3 | 2 | 2 | 0 | 0 | 1 | 0 | 0.001692225 | 0.002225681 | 0.001573974 | 0.000652683 | 0.000824616 | 0.00055686 |
| NITMOv2_4619 | A0A0K2GJ74 | NfeD domain-containing protein | 15.358 | 144 | -0.236 | -1.18 | 2.11E-01 | 2.52E-01 | no | yes | 1 | 2 | 2 | 2 | 1 | 1 | 1 | 0.001692077 | 0.001925521 | 0.002420894 | 0.002040615 | 0.001829926 | 0.002102543 |
| NITMOv2_0859 | A0A0K2G8V7 | Cupin_2 domain-containing protein | 13.164 | 120 | -0.23 | -1.17 | 5.32E-01 | 5.74E-01 | no | yes | 2 | 1 | 1 | 0 | 1 | 2 | 1 | 0.001688536 | 0.001948849 | 0.001533213 | 0.000190439 | 0.001151231 | 0.000958992 |

| | | | | | | | | | | | | | | | | | | | | | | | |
|---|---|---|---|---|---|---|---|---|---|---|---|---|---|---|---|---|---|---|---|---|---|---|---|
| mltG | A0A0K2GD72 | Endolytic murein transglycosylase | 39.698 | 360 | -0.211 | -1.16 | 1.51E-01 | 1.86E-01 | no | no | 0 | 3 | 3 | 3 | 2 | 2 | 2 | 0.001684236 | 0.001677552 | 0.001654459 | 0.001155352 | 0.001625306 | 0.001394965 |
| recG | A0A0K2G9Y6 | ATP-dependent DNA helicase RecG | 91.998 | 832 | -0.489 | -1.40 | 2.03E-03 | 4.87E-03 | no | no | 0 | 10 | 10 | 13 | 21 | 20 | 19 | 0.001682149 | 0.001598259 | 0.001931232 | 0.004319038 | 0.004078724 | 0.004014845 |
| NITMOv2_1041 | A0A0K2GA66 | Putative NAD-dependent epimerase/dehydratase | 28.984 | 260 | 2.15 | 4.44 | 3.81E-05 | 2.90E-04 | yes | yes | 3 | 1 | 1 | 2 | 0 | 0 | 0 | 0.001663073 | 0.001882736 | 0.001731072 | 0.000698453 | 0.000315522 | 0.000727088 |
| NITMOv2_3950 | A0A0K2GH99 | Putative Cytochrome c (Modular protein) | 19.192 | 178 | 0.786 | 1.72 | 1.61E-03 | 4.08E-03 | no | no | 0 | 1 | 2 | 2 | 1 | 0 | 1 | 0.00164777 | 0.001916976 | 0.001721678 | 0.001340119 | 0.001167809 | 0.001090129 |
| NITMOv2_3892 | A0A0K2GHE8 | D-xylulose 5-phosphate/D-fructose 6-phosphate phosphoketolase | 89.671 | 822 | 0.168 | 1.12 | 1.25E-01 | 1.57E-01 | no | no | 0 | 6 | 5 | 6 | 5 | 5 | 4 | 0.001634848 | 0.001699648 | 0.001543298 | 0.000953674 | 0.000977623 | 0.00097442 |
| NITMOv2_3849 | A0A0K2GGZ7 | Uncharacterized protein | 30.471 | 266 | 1.08 | 2.11 | 9.55E-02 | 1.24E-01 | no | no | 0 | 2 | 2 | 3 | 2 | 3 | 2 | 0.001620346 | 0.001776184 | 0.002171461 | 0.00043605 | 0.001577142 | 0.001444161 |
| NITMOv2_4701 | A0A0K2G7F2 | PqiB family protein | 60.277 | 547 | 0.000201 | 1.00 | 9.98E-01 | 9.98E-01 | no | no | 0 | 3 | 3 | 1 | 5 | 6 | 4 | 0.001617901 | 0.001951509 | 0.001420979 | 0.003185972 | 0.003361389 | 0.002800754 |
| NITMOv2_4850 | A0A0K2GJT7 | Putative Carboxylesterase | 25.65 | 237 | -0.049 | -1.03 | 7.93E-01 | 8.19E-01 | no | no | 0 | 2 | 2 | 3 | 4 | 4 | 1 | 0.001616426 | 0.001771354 | 0.002274992 | 0.002991254 | 0.002911132 | 0.00189468 |
| NITMOv2_0427 | A0A0K2G7D0 | Uncharacterized protein | 22.653 | 207 | 0.672 | 1.59 | 4.32E-02 | 6.18E-02 | no | yes | 4 | 2 | 2 | 1 | 0 | 1 | 0 | 0.001612484 | 0.001923018 | 0.00174641 | 0.000307018 | 0.001523369 | 0.000222374 |
| NITMOv2_3271 | A0A0K2GFN3 | Uncharacterized protein | 178.95 | 1622 | -0.273 | -1.21 | 6.83E-03 | 1.29E-02 | no | no | 0 | 13 | 12 | 13 | 14 | 13 | 11 | 0.001602556 | 0.001687251 | 0.001647097 | 0.001818355 | 0.0018697 | 0.001832645 |
| nadA | A0A0K2G935 | Quinolinate synthase A | 41.044 | 367 | -0.213 | -1.16 | 1.00E-01 | 1.30E-01 | no | no | 0 | 1 | 3 | 2 | 5 | 2 | 4 | 0.001586009 | 0.001500115 | 0.001191159 | 0.002012037 | 0.00298835 | 0.002591632 |
| argF | A0A0K2G776 | Ornithine carbamoyltransferase | 35.475 | 325 | -0.96 | -1.95 | 2.84E-04 | 1.11E-03 | no | no | 0 | 2 | 2 | 2 | 8 | 8 | 10 | 0.001584913 | 0.002443126 | 0.002483204 | 0.007170362 | 0.007191835 | 0.008063136 |
| NITMOv2_1752 | A0A0K2GC30 | ABC transporter, ATP-binding protein | 28.167 | 262 | 0.456 | 1.37 | 5.28E-03 | 1.04E-02 | no | no | 0 | 3 | 3 | 4 | 3 | 3 | 2 | 0.001583732 | 0.001365503 | 0.001515632 | 0.001558633 | 0.001410416 | 0.001182319 |
| NITMOv2_0498 | A0A0K2G7V8 | Putative Universal stress protein UspA (Modular protein) | 76.557 | 701 | -2.57 | -5.94 | 3.72E-06 | 6.29E-05 | yes | no | 0 | 1 | 1 | 3 | 37 | 35 | 36 | 0.001582636 | 0.00173819 | 0.00240939 | 0.014931838 | 0.014418627 | 0.015357035 |
| NITMOv2_3508 | A0A0K2GG29 | ABC transporter related protein | 65.629 | 605 | -1.81 | -3.51 | 2.21E-07 | 1.33E-05 | yes | no | 0 | 1 | 1 | 1 | 6 | 7 | 6 | 0.001576123 | 0.001561379 | 0.001506219 | 0.004293422 | 0.004597498 | 0.004416366 |
| NITMOv2_1425 | A0A0K2GB88 | Putative Pytoene desaturase | 48.566 | 441 | -0.665 | -1.59 | 5.43E-02 | 7.54E-02 | no | no | 0 | 2 | 1 | 1 | 2 | 3 | 3 | 0.001573868 | 0.001569123 | 0.000834707 | 0.001359946 | 0.001216051 | 0.001064901 |
| NITMOv2_0469 | A0A0K2G7K7 | Transglut_core2 domain-containing protein | 32.903 | 292 | -0.962 | -1.95 | 2.57E-05 | 2.23E-04 | no | no | 0 | 2 | 2 | 1 | 6 | 5 | 5 | 0.00157216 | 0.001551543 | 0.001409993 | 0.003546978 | 0.003573933 | 0.002937411 |
| nusB | A0A0K2GAV3 | Transcription antitermination protein NusB | 17.358 | 152 | -1.13 | -2.19 | 1.02E-02 | 1.80E-02 | yes | yes | 2 | 0 | 0 | 1 | 3 | 3 | 1 | 0.001570959 | 0.001776008 | 0.002142894 | 0.002242293 | 0.00245839 | 0.002457852 |
| cdsA | A0A0K2GB72 | Phosphatidate cytidylyltransferase | 32.258 | 308 | -0.0967 | -1.07 | 6.91E-01 | 7.27E-01 | no | yes | 4 | 1 | 0 | 1 | 0 | 1 | 0 | 0.001569462 | 0.001670277 | 0.001458825 | 0.001207637 | 0.001259709 | 0.001262875 |
| NITMOv2_3843 | A0A0K2GHY0 | APH domain-containing protein | 41.835 | 361 | 0.511 | 1.43 | 2.03E-03 | 4.87E-03 | no | no | 0 | 2 | 5 | 4 | 1 | 3 | 3 | 0.001561558 | 0.002444886 | 0.002628339 | 0.001179162 | 0.002200013 | 0.001701706 |
| NITMOv2_4498 | A0A0K2GJS3 | Uncharacterized protein | 38.101 | 360 | -1.48 | -2.79 | 6.18E-05 | 4.07E-04 | yes | no | 0 | 14 | 13 | 13 | 23 | 23 | 21 | 0.001560082 | 0.001547867 | 0.001339151 | 0.003517511 | 0.003371488 | 0.003044219 |
| mrdA | A0A0K2G8K4 | Penicillin-binding protein 2 | 67.438 | 622 | -0.172 | -1.13 | 2.61E-02 | 3.99E-02 | no | no | 0 | 4 | 7 | 6 | 5 | 7 | 7 | 0.001548194 | 0.001416989 | 0.001518604 | 0.001670873 | 0.001933867 | 0.002160802 |
| NITMOv2_2703 | A0A0K2GE31 | Histidine kinase | 227.25 | 2099 | -0.74 | -1.67 | 1.42E-03 | 3.69E-03 | no | no | 0 | 13 | 18 | 22 | 55 | 46 | 35 | 0.001541259 | 0.001788503 | 0.001811519 | 0.004064202 | 0.004322341 | 0.003004481 |
| NITMOv2_3517 | A0A0K2GGE9 | Uncharacterized protein | 42.194 | 362 | -0.787 | -1.73 | 7.08E-04 | 2.20E-03 | no | no | 0 | 9 | 10 | 7 | 7 | 7 | 9 | 0.001540711 | 0.000728538 | 0.001263017 | 0.001014251 | 0.00136219 | 0.001893062 |
| NITMOv2_1037 | A0A0K2G946 | Putative Imidazolonepropionase | 41.768 | 402 | -0.563 | -1.48 | 1.77E-04 | 8.03E-04 | no | no | 0 | 5 | 5 | 5 | 5 | 9 | 9 | 0.001539235 | 0.001554222 | 0.00160534 | 0.003139921 | 0.00352437 | 0.003170627 |
| fabH.1 | A0A0K2G958 | Beta-ketoacyl-[acyl-carrier-protein] synthase III | 34.334 | 330 | -2.2 | -4.59 | 9.64E-07 | 2.93E-05 | yes | no | 0 | 1 | 0 | 2 | 10 | 9 | 9 | 0.001537507 | 0.001479466 | 0.001475064 | 0.007165327 | 0.007630905 | 0.007129731 |
| tesA | A0A0K2G755 | Esterase TesA | 25.287 | 231 | 0.376 | 1.30 | 5.22E-02 | 7.31E-02 | no | no | 0 | 2 | 3 | 3 | 2 | 2 | 1 | 0.001534176 | 0.001889893 | 0.002081735 | 0.001683015 | 0.001232504 | 0.001110951 |
| NITMOv2_0830 | A0A0K2G8J5 | Uncharacterized protein | 45.134 | 395 | 0.0705 | 1.05 | 6.89E-01 | 7.25E-01 | no | no | 0 | 1 | 2 | 1 | 2 | 2 | 2 | 0.001533207 | 0.001748221 | 0.001490843 | 0.001439981 | 0.001308386 | 0.001441985 |
| NITMOv2_3437 | A0A0K2GG36 | Uncharacterized protein | 25.981 | 237 | -0.574 | -1.49 | 2.43E-02 | 3.76E-02 | no | no | 0 | 1 | 2 | 1 | 2 | 2 | 3 | 0.001533101 | 0.00209232 | 0.001501349 | 0.002997621 | 0.003107673 | 0.00293831 |
| NITMOv2_2514 | A0A0K2GEB6 | Putative Amylo-alpha-1,6-glucosidase | 80.005 | 722 | 0.238 | 1.18 | 1.71E-01 | 2.08E-01 | no | no | 0 | 5 | 4 | 9 | 3 | 4 | 2 | 0.001525323 | 0.001765879 | 0.002365486 | 0.001227553 | 0.001147751 | 0.000927327 |
| ubiA | A0A0K2G893 | 4-hydroxybenzoate octaprenyltransferase | 31.918 | 298 | -0.599 | -1.51 | 4.50E-03 | 9.10E-03 | no | no | 0 | 3 | 4 | 4 | 0 | 0 | 1 | 0.001524754 | 0.00162454 | 0.001504129 | 0.001307839 | 0.000998178 | 0.001043863 |
| NITMOv2_3475 | A0A0K2GGX8 | Uncharacterized protein | 27.909 | 250 | 0.0392 | 1.03 | 9.38E-01 | 9.47E-01 | no | yes | 1 | 2 | 2 | 3 | 1 | 2 | 3 | 0.001521676 | 0.001418652 | 0.001577866 | 0.000754203 | 0.00016121 | 0.000663183 |
| NITMOv2_3851 | A0A0K2GH03 | Transcriptional regulator | 24.41 | 216 | 0.648 | 1.57 | 2.61E-05 | 2.24E-04 | no | yes | 2 | 2 | 2 | 3 | 1 | 1 | 1 | 0.001518768 | 0.001614997 | 0.001492319 | 0.00080323 | 0.00079881 | 0.0009518 |
| trmFO | A0A0K2G6Q1 | Methylenetetrahydrofolate--tRNA-(uracil-5-)-methyltransferase TrmFO | 48.44 | 438 | -1.46 | -2.75 | 1.06E-06 | 3.10E-05 | yes | no | 0 | 2 | 1 | 1 | 8 | 11 | 5 | 0.001516196 | 0.001673817 | 0.001572018 | 0.00440892 | 0.004421 | 0.003920264 |
| NITMOv2_0515 | A0A0K2G7L0 | Putative two-component sensor histidine kinase | 30.051 | 275 | -1.4 | -2.64 | 6.48E-04 | 2.06E-03 | yes | no | 0 | 2 | 3 | 4 | 11 | 12 | 10 | 0.001515163 | 0.001586839 | 0.00204895 | 0.005485422 | 0.005277544 | 0.004793072 |
| NITMOv2_0605 | A0A0K2G7U2 | 1L-myo-inositol-1-phosphate cytidylyltransferase | 51.301 | 478 | -0.344 | -1.27 | 1.32E-03 | 3.49E-03 | no | no | 0 | 3 | 3 | 1 | 4 | 2 | 3 | 0.001513498 | 0.001602287 | 0.001481985 | 0.002803495 | 0.0029102 | 0.002662478 |
| NITMOv2_0969 | A0A0K2G8X3 | Putative ABC transporter, permease component | 92.539 | 870 | 0.12 | 1.09 | 2.50E-01 | 2.91E-01 | no | no | 0 | 12 | 15 | 13 | 7 | 7 | 8 | 0.001512086 | 0.00192204 | 0.002066397 | 0.001908977 | 0.001661973 | 0.00142805 |
| NITMOv2_3667 | A0A0K2GGU2 | NFACT-R_1 domain-containing protein | 53.861 | 478 | 0.543 | 1.46 | 2.26E-04 | 9.65E-04 | no | no | 0 | 9 | 7 | 5 | 4 | 3 | 4 | 0.001502874 | 0.001807862 | 0.002040131 | 0.001642443 | 0.00142269 | 0.001416938 |
| NITMOv2_2394 | A0A0K2GD67 | PLD phosphodiesterase domain-containing protein | 21.505 | 190 | -0.152 | -1.11 | 7.23E-01 | 7.56E-01 | no | yes | 3 | 0 | 1 | 1 | 2 | 0 | 0 | 0.001499375 | 0.002345158 | 0.001991049 | 0.002432273 | 0.001666323 | 0.002412539 |
| NITMOv2_0190 | A0A0K2G6S4 | Uncharacterized protein | 28.87 | 275 | -1.11 | -2.16 | 5.03E-05 | 3.53E-04 | yes | no | 0 | 1 | 2 | 0 | 3 | 3 | 4 | 0.00149088 | 0.001786137 | 0.001338825 | 0.002158335 | 0.002908336 | 0.002873758 |
| NITMOv2_2367 | A0A0K2GDU8 | Uncharacterized protein | 29.523 | 261 | -0.217 | -1.16 | 7.20E-02 | 9.68E-02 | no | no | 0 | 0 | 1 | 2 | 2 | 0 | 2 | 0.001488098 | 0.002771248 | 0.001380218 | 0.001757201 | 0.002319646 | 0.00242117 |
| NITMOv2_3682 | A0A0K2GGV3 | Uncharacterized protein | 58.623 | 520 | 0.242 | 1.18 | 2.93E-02 | 4.43E-02 | no | no | 0 | 2 | 1 | 0 | 2 | 3 | 2 | 0.001486243 | 0.001658701 | 0.001517454 | 0.001534052 | 0.001580094 | 0.001500424 |
| NITMOv2_4210 | A0A0K2GIA4 | MULTIHEME_CYTC domain-containing protein | 49.009 | 445 | 0.182 | 1.13 | 1.55E-01 | 1.91E-01 | no | no | 0 | 3 | 3 | 3 | 2 | 1 | 2 | 0.001466513 | 0.001730564 | 0.001559997 | 0.001298081 | 0.001421618 | 0.001306173 |
| NITMOv2_0041 | A0A0K2G6C9 | Inositol-1-monophosphatase | 35.781 | 326 | 0.0411 | 1.03 | 7.82E-01 | 8.09E-01 | no | no | 0 | 0 | 2 | 2 | 2 | 1 | 0 | 0.001465354 | 0.001224692 | 0.001550104 | 0.001866628 | 0.001341277 | 0.001302937 |
| NITMOv2_3289 | A0A0K2GGC7 | UPF0753 protein NITMOv2_3289 | 121.54 | 1084 | 1.11 | 2.16 | 4.97E-03 | 9.89E-03 | yes | no | 0 | 4 | 3 | 6 | 1 | 0 | 2 | 0.001461391 | 0.001414819 | 0.001521461 | 0.000443913 | 0.000355156 | 0.000305123 |
| NITMOv2_2937 | A0A0K2GFG3 | Helix-turn-helix domain protein | 13.135 | 121 | -1.21 | -2.31 | 1.21E-03 | 3.22E-03 | yes | no | 0 | 0 | 1 | 1 | 3 | 2 | 2 | 0.001458672 | 0.004748589 | 0.004334295 | 0.011777554 | 0.005062515 | 0.00504373 |
| cebB | A0A0K2GBU7 | Chemiosmotic efflux system B protein B | 46.652 | 418 | 0.42 | 1.34 | 3.27E-02 | 4.86E-02 | no | no | 0 | 2 | 1 | 4 | 0 | 1 | 1 | 0.001456691 | 0.001382945 | 0.001250076 | 0.000710225 | 0.000423658 | 0.000257293 |
| atoC | A0A0K2G818 | Acetoacetate metabolism regulatory protein AtoC | 51.273 | 461 | -2.32 | -4.99 | 5.63E-03 | 1.09E-02 | yes | yes | 2 | 2 | 1 | 1 | 4 | 4 | 4 | 0.0014561 | 0.001242662 | 0.001373163 | 0.002385333 | 0.002090944 | 0.002228052 |
| NITMOv2_2276 | A0A0K2GDJ8 | Putative Aldo-keto reductase | 34.674 | 312 | -0.901 | -1.87 | 2.97E-03 | 6.56E-03 | no | no | 0 | 3 | 3 | 3 | 3 | 4 | 5 | 0.0014561 | 0.001501347 | 0.001569679 | 0.002169588 | 0.002118444 | 0.00274699 |
| NITMOv2_1090 | A0A0K2G975 | Uncharacterized protein | 35.903 | 340 | -0.705 | -1.63 | 1.42E-02 | 2.40E-02 | no | no | 0 | 1 | 3 | 4 | 13 | 9 | 9 | 0.001452412 | 0.002200456 | 0.002057961 | 0.003870225 | 0.003646179 | 0.00275616 |
| priA | A0A0K2G6Q3 | Probable primosomal protein N | 72.985 | 679 | 0.447 | 1.36 | 5.25E-03 | 1.04E-02 | no | no | 0 | 11 | 12 | 12 | 5 | 2 | 5 | 0.001448386 | 0.001998654 | 0.00192433 | 0.001385637 | 0.001259134 | 0.001397158 |
| NITMOv2_3392 | A0A0K2GG03 | TPR_REGION domain-containing protein | 24.369 | 213 | -2.26 | -4.79 | 3.06E-06 | 5.51E-05 | yes | no | 0 | 1 | 1 | 1 | 6 | 6 | 5 | 0.001434727 | 0.001637015 | 0.001474143 | 0.007158664 | 0.007510029 | 0.006465145 |
| NITMOv2_3518 | A0A0K2GG35 | Putative Lipopolysaccharide heptosyltransferase III | 38.03 | 345 | -0.853 | -1.81 | 1.67E-01 | 2.03E-01 | no | yes | 3 | 1 | 1 | 1 | 0 | 1 | 2 | 0.001432872 | 0.001080321 | 0.001012416 | 0.000817623 | 0.000985593 | 0.000918714 |
| NITMOv2_1525 | A0A0K2GAI8 | Putative Transcriptional regulator ModE | 41.799 | 389 | -0.0616 | -1.04 | 6.16E-01 | 6.56E-01 | no | no | 0 | 3 | 5 | 2 | 4 | 4 | 0 | 0.001426632 | 0.00153543 | 0.001563199 | 0.001647033 | 0.001714021 | 0.001187605 |
| gmd | A0A0K2GG40 | GDP-mannose 4,6-dehydratase | 41.327 | 363 | -1.91 | -3.76 | 8.04E-06 | 1.01E-04 | yes | no | 0 | 2 | 0 | 3 | 9 | 7 | 5 | 0.001425831 | 0.000992913 | 0.001695028 | 0.006884282 | 0.006438455 | 0.005471503 |
| NITMOv2_4512 | A0A0K2GIX0 | Uncharacterized protein | 15.078 | 138 | -1.99 | -3.97 | 1.05E-03 | 2.92E-03 | yes | no | 0 | 3 | 3 | 3 | 6 | 8 | 6 | 0.001421974 | 0.002512544 | 0.00279648 | 0.003763167 | 0.004372836 | 0.00339773 |
| lspA | A0A0K2G8T8 | Lipoprotein signal peptidase | 19.627 | 178 | -0.457 | -1.37 | 7.69E-02 | 1.03E-01 | no | no | 0 | 0 | 1 | 1 | 5 | 3 | 1 | 0.001418032 | 0.001526983 | 0.001811098 | 0.002089776 | 0.001834587 | 0.001536099 |
| NITMOv2_1459 | A0A0K2GA90 | Site-specific DNA-methyltransferase (adenine-specific) | 61.035 | 548 | -0.665 | -1.59 | 4.58E-04 | 1.59E-03 | no | no | 0 | 6 | 11 | 7 | 12 | 14 | 14 | 0.00141272 | 0.001662241 | 0.002164368 | 0.003290661 | 0.003515825 | 0.002990635 |
| NITMOv2_0996 | A0A0K2G902 | Glyco_hydro_63 domain-containing protein | 103.13 | 892 | 0.24 | 1.18 | 1.64E-01 | 2.00E-01 | no | no | 0 | 3 | 4 | 6 | 5 | 4 | 4 | 0.001410128 | 0.001505297 | 0.001457713 | 0.000678922 | 0.000855317 | 0.000658904 |
| NITMOv2_2711 | A0A0K2GET0 | Putative Uncharacterized lipoprotein yeaY | 18.244 | 160 | -0.917 | -1.89 | 2.77E-03 | 6.20E-03 | no | no | 0 | 3 | 4 | 3 | 4 | 3 | 4 | 0.001409811 | 0.001544601 | 0.00177281 | 0.002401029 | 0.002266199 | 0.001950422 |
| fdx | A0A0K2GIL7 | 2Fe-2S ferredoxin | 12.597 | 115 | -1.75 | -3.36 | 4.46E-02 | 6.36E-02 | no | yes | 1 | 0 | 0 | 0 | 3 | 2 | 2 | 0.001400937 | 0.00244997 | 0.001564273 | 0.006067947 | 0.004260815 | 0.006351324 |
| NITMOv2_2702 | A0A0K2GDT8 | Putative Histidine kinase family | 42.728 | 378 | -2.54 | -5.82 | 9.36E-05 | 5.41E-04 | yes | yes | 3 | 0 | 0 | 0 | 2 | 3 | 2 | 0.001400684 | 0.001350739 | 0.00137995 | 0.001996785 | 0.001910406 | 0.001900434 |
| nuoM.2 | A0A0K2G7P2 | NADH-quinone oxidoreductase, membrane subunit M | 56.506 | 516 | -1.74 | -3.34 | 2.86E-03 | 6.37E-03 | yes | yes | 3 | 2 | 2 | 1 | 2 | 2 | 3 | 0.001398155 | 0.001548395 | 0.001275901 | 0.002670376 | 0.002692529 | 0.002499209 |
| NITMOv2_1416 | A0A0K2GA65 | DUF5615 domain-containing protein | 12.009 | 107 | 0.338 | 1.26 | 1.07E-01 | 1.37E-01 | no | yes | 3 | 2 | 0 | 0 | 0 | 0 | 0 | 0.001375959 | 0.001438362 | 0.001443851 | 0.000482693 | 0.000337304 | 0.000389617 |
| xseA | A0A0K2GHV3 | Exodeoxyribonuclease 7 large subunit | 45.812 | 415 | 0.0408 | 1.03 | 7.68E-01 | 7.96E-01 | no | no | 0 | 6 | 6 | 5 | 3 | 5 | 2 | 0.001372165 | 0.001397846 | 0.001447015 | 0.00149422 | 0.002094207 | 0.00140926 |
| NITMOv2_3499 | A0A0K2GG19 | Glyco_trans_2-like domain-containing protein | 38.635 | 338 | -3.21 | -9.25 | 3.56E-06 | 6.20E-05 | yes | no | 0 | 9 | 13 | 12 | 27 | 29 | 29 | 0.001370942 | 0.001349625 | 0.001466034 | 0.012736781 | 0.012677261 | 0.011896551 |
| NITMOv2_1818 | A0A0K2GBB6 | Uncharacterized protein | 52.39 | 479 | 0.00547 | 1.00 | 9.45E-01 | 9.51E-01 | no | no | 0 | 7 | 10 | 6 | 3 | 3 | 3 | 0.001362237 | 0.001378115 | 0.001280349 | 0.00102857 | 0.001013994 | 0.001034639 |
| NITMOv2_3698 | A0A0K2GGK2 | Uncharacterized protein | 37.003 | 350 | -0.45 | -1.37 | 6.35E-02 | 8.65E-02 | no | no | 0 | 1 | 2 | 2 | 4 | 2 | 1 | 0.001359096 | 0.001471781 | 0.002309311 | 0.002791798 | 0.002556427 | 0.002661759 |
| NITMOv2_4224 | A0A0K2GI05 | Efflux transporter, RND family, MFP subunit | 60.974 | 569 | 1.14 | 2.20 | 1.17E-02 | 2.04E-02 | yes | yes | 2 | 2 | 2 | 2 | 2 | 1 | 1 | 0.00135585 | 0.001249799 | 0.001211846 | 0.001054009 | 0.000129187 | 0.000149026 |
| atoC.2 | A0A0K2GCK2 | Acetoacetate metabolism regulatory protein AtoC | 53.068 | 470 | -2.18 | -4.53 | 4.91E-06 | 7.39E-05 | yes | no | 0 | 4 | 4 | 4 | 24 | 24 | 23 | 0.001342802 | 0.001283237 | 0.001648612 | 0.008239904 | 0.008830347 | 0.007604974 |
| NITMOv2_1859 | A0A0K2GBN2 | Lipopolysaccharide export system permease protein LptF | 40.447 | 363 | 0.151 | 1.11 | 1.46E-01 | 1.80E-01 | no | no | 0 | 3 | 4 | 4 | 4 | 2 | 2 | 0.001341074 | 0.001346535 | 0.001333073 | 0.00136581 | 0.001477846 | 0.001593729 |
| epsC | A0A0K2GFY8 | Putative polysaccharide biosynthesis protein EpsC | 69.363 | 633 | -1.66 | -3.16 | 2.36E-06 | 4.65E-05 | yes | no | 0 | 5 | 3 | 6 | 24 | 20 | 14 | 0.001329586 | 0.001253124 | 0.002128899 | 0.006935368 | 0.007347669 | 0.007001165 |
| NITMOv2_4246 | A0A0K2GID7 | Pyridoxamine 5-phosphate oxidase-related FMN-binding | 22.645 | 199 | 0.995 | 1.99 | 2.68E-04 | 1.08E-03 | no | no | 0 | 2 | 2 | 4 | 0 | 0 | 0 | 0.00132948 | 0.001420235 | 0.001841639 | 0.000471973 | 0.000329008 | 0.000315049 |
| NITMOv2_3743 | A0A0K2GH10 | Glycoside hydrolase, family 57 | 83.802 | 733 | -0.131 | -1.10 | 3.03E-01 | 3.44E-01 | no | no | 0 | 4 | 2 | 6 | 8 | 9 | 8 | 0.001324569 | 0.001295146 | 0.001155211 | 0.001923785 | 0.001833499 | 0.001658138 |
| NITMOv2_4163 | A0A0K2GHU8 | Divalent-cation tolerance protein CutA (Modular protein) | 15.395 | 140 | -1.43 | -2.69 | 3.73E-03 | 7.87E-03 | yes | no | 0 | 2 | 2 | 2 | 4 | 3 | 3 | 0.001322229 | 0.001525086 | 0.001562586 | 0.002326547 | 0.001851211 | 0.001917157 |
| NITMOv2_3413 | A0A0K2GFR4 | Putative Lipopolysaccharide biosynthesis protein | 88.477 | 802 | -1.17 | -2.25 | 3.33E-04 | 1.24E-03 | yes | no | 0 | 3 | 3 | 2 | 11 | 11 | 11 | 0.001318119 | 0.001772449 | 0.00118652 | 0.005111385 | 0.005545088 | 0.006303494 |
| czcA.2 | A0A0K2GED4 | Cation efflux system protein CzcA | 114.04 | 1048 | -0.652 | -1.57 | 3.41E-02 | 5.03E-02 | no | no | 0 | 1 | 2 | 7 | 1 | 2 | 1 | 0.001317318 | 0.001501582 | 0.001386449 | 0.000783211 | 0.000994542 | 0.000930654 |
| soj | A0A0K2G784 | Sporulation initiation inhibitor protein soj | 27.681 | 258 | -0.492 | -1.41 | 5.30E-03 | 1.04E-02 | no | no | 0 | 1 | 1 | 3 | 0 | 0 | 0 | 0.001315969 | 0.001514996 | 0.001626276 | 0.001945848 | 0.002447825 | 0.001890904 |
| hrpB | A0A0K2GCM0 | ATP-dependent RNA helicase HrpB | 92.051 | 840 | 0.0163 | 1.01 | 8.10E-01 | 8.33E-01 | no | no | 0 | 6 | 6 | 9 | 6 | 5 | 8 | 0.0013158 | 0.001153279 | 0.001154962 | 0.000981423 | 0.000910908 | 0.000939159 |
| NITMOv2_1891 | A0A0K2GBG3 | DNA polymerase, beta-like region | 12.549 | 114 | 0.273 | 1.21 | 2.31E-01 | 2.72E-01 | no | yes | 1 | 1 | 2 | 1 | 0 | 0 | 0 | 0.00130817 | 0.001162567 | 0.00099911 | 0.000824375 | 0.00073573 | 0.000588885 |

| | | | | | | | | | | | | | | | | | | | | | | |
|---|---|---|---|---|---|---|---|---|---|---|---|---|---|---|---|---|---|---|---|---|---|---|
| NITMOv2_1572 | A0A0K2GBN4 | HTH cro/C1-type domain-containing protein | 24.855 | 225 | 0.269 | 1.20 | 3.22E-02 | 4.81E-02 | no | no | 0 | 1 | 2 | 2 | 2 | 1 | 1 | 0.001303448 | 0.001673132 | 0.00154719 | 0.000856641 | 0.000762982 | 0.000897802 |
| NITMOv2_0528 | A0A0K2G7Y6 | Response regulatory domain-containing protein | 28.773 | 259 | -0.0195 | -1.01 | 8.30E-01 | 8.50E-01 | no | no | 0 | 2 | 2 | 1 | 1 | 3 | 1 | 0.001299843 | 0.001224379 | 0.001026392 | 0.001307868 | 0.001525016 | 0.001456119 |
| NITMOv2_4137 | A0A0K2GI39 | Putative DeoR-family transcriptional regulator | 25.901 | 235 | 0.0702 | 1.05 | 8.03E-01 | 8.28E-01 | no | yes | 2 | 1 | 2 | 2 | 1 | 2 | 1 | 0.001286142 | 0.001341509 | 0.001269958 | 0.000717451 | 0.001022446 | 0.000836918 |
| NITMOv2_3683 | A0A0K2GGI8 | Uncharacterized protein | 63.591 | 575 | -0.952 | -1.93 | 1.63E-04 | 7.72E-04 | no | no | 0 | 2 | 2 | 4 | 6 | 6 | 4 | 0.001285426 | 0.000963445 | 0.001335393 | 0.001953548 | 0.001950647 | 0.001903671 |
| NITMOv2_0583 | A0A0K2G839 | Uncharacterized protein | 23.557 | 208 | 0.271 | 1.21 | 4.20E-01 | 4.64E-01 | no | yes | 3 | 0 | 3 | 1 | 0 | 0 | 0 | 0.001279629 | 0.001835903 | 0.001342084 | 0.001050648 | 0.001094864 | 0.000792198 |
| NITMOv2_3712 | A0A0K2GGY2 | FGE-sulfatase domain-containing protein | 33.284 | 300 | 0.753 | 1.69 | 1.34E-01 | 1.67E-01 | no | no | 0 | 3 | 5 | 0 | 1 | 0 | 0 | 0.001274507 | 0.001243522 | 0.001079691 | 0.000965061 | 0.00095194 | 0.000809316 |
| NITMOv2_3669 | A0A0K2GGJ4 | Fe2OG dioxygenase domain-containing protein | 25.916 | 227 | -1.04 | -2.06 | 5.32E-03 | 1.04E-02 | yes | no | 0 | 2 | 3 | 3 | 8 | 7 | 8 | 0.001271724 | 0.001618497 | 0.001879639 | 0.003853048 | 0.00337848 | 0.003404563 |
| NITMOv2_3188 | A0A0K2GG33 | Uncharacterized protein | 29.123 | 265 | -0.169 | -1.12 | 1.96E-01 | 2.35E-01 | no | no | 0 | 3 | 2 | 1 | 2 | 3 | 3 | 0.001271493 | 0.001405003 | 0.001349025 | 0.001273678 | 0.001586153 | 0.003714739 |
| NITMOv2_1019 | A0A0K2G909 | Uncharacterized protein | 31.62 | 298 | 0.578 | 1.49 | 1.60E-02 | 2.63E-02 | no | yes | 1 | 1 | 0 | 1 | 1 | 2 | 1 | 0.001270186 | 0.000955271 | 0.001140372 | 0.000303271 | 0.00093572 | 0.000726639 |
| NITMOv2_2706 | A0A0K2GES5 | Nitrogen regulation protein B | 43.715 | 392 | 0.595 | 1.51 | 4.69E-04 | 1.62E-03 | no | no | 0 | 2 | 3 | 4 | 0 | 1 | 1 | 0.001269532 | 0.001178524 | 0.001361161 | 0.000587323 | 0.000910333 | 0.000793942 |
| NITMOv2_1042 | A0A0K2G950 | Putative ATP-dependent DNA helicase Lhr | 159.48 | 1451 | 0.0898 | 1.06 | 2.79E-01 | 3.22E-01 | no | no | 0 | 13 | 14 | 14 | 8 | 8 | 3 | 0.00124917 | 0.001216987 | 0.001129482 | 0.001307543 | 0.001382341 | 0.001085256 |
| NITMOv2_1033 | A0A0K2G9D5 | Putative ATP dependent transcriptional activator | 120.9 | 1067 | 0.535 | 1.45 | 2.52E-03 | 5.75E-03 | no | no | 0 | 7 | 9 | 8 | 2 | 5 | 3 | 0.001243479 | 0.001331478 | 0.001501138 | 0.000809953 | 0.000984599 | 0.000952213 |
| NITMOv2_1478 | A0A0K2GAE5 | DNA-binding response regulator, CheY like | 16.194 | 146 | -2.04 | -4.11 | 3.69E-03 | 7.81E-03 | yes | no | 0 | 2 | 1 | 3 | 5 | 3 | 3 | 0.001240359 | 0.001621059 | 0.003442972 | 0.010520993 | 0.00954069 | 0.008093704 |
| ruvA | A0A0K2G854 | Holliday junction ATP-dependent DNA helicase RuvA | 21.773 | 202 | -1.37 | -2.58 | 5.11E-05 | 3.55E-04 | yes | no | 0 | 1 | 1 | 2 | 6 | 5 | 6 | 0.001232434 | 0.001107013 | 0.001489021 | 0.003895545 | 0.003844584 | 0.003453472 |
| NITMOv2_0445 | A0A0K2G7C7 | Putative tetR-family transcriptional regulator | 21.537 | 193 | -0.613 | -1.53 | 4.19E-05 | 3.07E-04 | no | no | 0 | 2 | 3 | 1 | 6 | 3 | 3 | 0.001231274 | 0.001071952 | 0.001242963 | 0.002048463 | 0.002475014 | 0.001651395 |
| tilS | A0A0K2GB77 | tRNA(Ile)-lysidine synthase | 54.243 | 487 | -0.287 | -1.22 | 1.29E-02 | 2.21E-02 | no | no | 0 | 6 | 5 | 3 | 8 | 6 | 6 | 0.001222379 | 0.001115226 | 0.001120413 | 0.001158802 | 0.001058663 | 0.001146087 |
| NITMOv2_1186 | A0A0K2G9J8 | DNA helicase | 75.28 | 672 | -0.326 | -1.25 | 6.49E-02 | 8.81E-02 | no | no | 0 | 4 | 6 | 6 | 4 | 10 | 6 | 0.00121985 | 0.001672252 | 0.001191696 | 0.001946292 | 0.002010153 | 0.001370186 |
| NITMOv2_1519 | A0A0K2GAI3 | Putative acetyltransferase | 17.925 | 159 | 0.389 | 1.31 | 2.13E-01 | 2.54E-01 | no | yes | 3 | 1 | 0 | 1 | 1 | 0 | 1 | 0.001219828 | 0.001273675 | 0.001540365 | 0.000503853 | 0.001072024 | 0.001012037 |
| NITMOv2_1615 | A0A0K2GAZ8 | PepSY domain-containing protein | 53.192 | 481 | -0.461 | -1.38 | 1.82E-02 | 2.94E-02 | no | no | 0 | 5 | 5 | 5 | 8 | 10 | 8 | 0.001218458 | 0.001424674 | 0.001585209 | 0.002241849 | 0.00236144 | 0.003752499 |
| msbA | A0A0K2GGG4 | Lipid A export ATP-binding/permease protein MsbA | 63.849 | 581 | -1.86 | -3.63 | 4.69E-06 | 7.17E-05 | yes | no | 0 | 0 | 1 | 2 | 12 | 12 | 12 | 0.001217636 | 0.001270449 | 0.00144412 | 0.004949836 | 0.005437263 | 0.005138851 |
| NITMOv2_0591 | A0A0K2G8V3 | SCP2 domain-containing protein | 13.127 | 121 | -0.487 | -1.40 | 3.02E-01 | 3.44E-01 | no | yes | 2 | 1 | 1 | 1 | 2 | 1 | 1 | 0.001213905 | 0.000956855 | 0.001160024 | 0.002143971 | 0.001532178 | 0.001966066 |
| NITMOv2_0513 | A0A0K2G7X2 | Uncharacterized protein | 100.65 | 889 | -0.887 | -1.85 | 9.54E-05 | 5.41E-04 | no | no | 0 | 3 | 2 | 5 | 20 | 17 | 18 | 0.001212303 | 0.001254668 | 0.001512277 | 0.004517458 | 0.004368796 | 0.004369794 |
| NITMOv2_2498 | A0A0K2GD89 | Uncharacterized protein | 47.825 | 430 | -1.63 | -3.10 | 1.18E-06 | 3.13E-05 | yes | no | 0 | 2 | 2 | 1 | 9 | 12 | 9 | 0.001211292 | 0.000975236 | 0.00108088 | 0.004360203 | 0.004395053 | 0.004880641 |
| dop | A0A0K2G8D5 | Pup deamidase/depupylase | 57.393 | 506 | -1.25 | -2.38 | 5.50E-06 | 7.92E-05 | yes | no | 0 | 5 | 5 | 5 | 14 | 15 | 10 | 0.001202059 | 0.0012445 | 0.001355486 | 0.004361832 | 0.004079501 | 0.003411036 |
| atoC.3 | A0A0K2GEA7 | Acetoacetate metabolism regulatory protein AtoC | 50.159 | 458 | -0.29 | -1.22 | 6.09E-02 | 8.34E-02 | no | no | 0 | 3 | 4 | 6 | 6 | 7 | 5 | 0.001198729 | 0.001399566 | 0.001685327 | 0.001829017 | 0.001493414 | 0.00169793 |
| yggS | A0A0K2G7W8 | Pyridoxal phosphate homeostasis protein | 26.282 | 239 | -0.13 | -1.09 | 3.25E-01 | 3.67E-01 | no | no | 0 | 3 | 3 | 3 | 3 | 4 | 2 | 0.001198349 | 0.001135739 | 0.001347798 | 0.001553006 | 0.001354903 | 0.001173454 |
| mobA | A0A0K2GGG3 | Probable molybdenum cofactor guanylyltransferase | 23.387 | 213 | -0.0341 | -1.02 | 5.54E-01 | 5.95E-01 | no | no | 0 | 1 | 2 | 1 | 1 | 1 | 1 | 0.001173034 | 0.001130342 | 0.001207303 | 0.001146823 | 0.001224938 | 0.001209452 |
| hmuV | A0A0K2GBK2 | Hemin import ATP-binding protein HmuV | 32.638 | 296 | -0.752 | -1.68 | 2.04E-03 | 4.88E-03 | no | no | 0 | 3 | 3 | 4 | 7 | 6 | 5 | 0.001169977 | 0.001242662 | 0.001378684 | 0.002133902 | 0.001689628 | 0.001845052 |
| NITMOv2_1057 | A0A0K2G961 | Putative 6-phosphofructokinase (Modular protein) | 84.261 | 776 | -0.464 | -1.38 | 1.24E-02 | 2.14E-02 | no | no | 0 | 6 | 5 | 5 | 13 | 12 | 8 | 0.001165487 | 0.001260183 | 0.001256537 | 0.001854634 | 0.002193953 | 0.001892703 |
| yfhA | A0A0K2GCF9 | Putative DNA-binding response regulator in two-component system | 52.769 | 474 | -1.84 | -3.58 | 1.36E-06 | 3.39E-05 | yes | no | 0 | 4 | 5 | 6 | 24 | 24 | 19 | 0.001160513 | 0.001003062 | 0.001689411 | 0.008888619 | 0.009130673 | 0.007221075 |
| rlmL | A0A0K2GCQ2 | Ribosomal RNA large subunit methyltransferase L | 43.554 | 390 | 0.131 | 1.10 | 1.72E-01 | 2.09E-01 | no | no | 0 | 3 | 4 | 3 | 2 | 2 | 2 | 0.001155391 | 0.001211258 | 0.001082107 | 0.000844277 | 0.001101855 | 0.001236136 |
| NITMOv2_1813 | A0A0K2GBB2 | HTH luxR-type domain-containing protein | 41.469 | 365 | -0.329 | -1.26 | 2.64E-01 | 3.06E-01 | no | no | 0 | 0 | 1 | 1 | 4 | 3 | 0 | 0.001147676 | 0.001226197 | 0.003347302 | 0.003217956 | 0.001987624 | 0.001465163 |
| NITMOv2_4839 | A0A0K2GKQ6 | S5A_REDUCTASE domain-containing protein | 29.924 | 263 | -0.219 | -1.16 | 4.42E-01 | 4.85E-01 | no | yes | 1 | 3 | 2 | 4 | 2 | 2 | 2 | 0.001146327 | 0.001796579 | 0.00232484 | 0.001778375 | 0.002432132 | 0.002222658 |
| ureF | A0A0K2G9Z2 | Urease accessory protein UreF | 24.524 | 228 | -2.08 | -4.23 | 2.97E-03 | 6.57E-03 | yes | yes | 3 | 0 | 0 | 0 | 2 | 2 | 0 | 0.001137579 | 0.00148191 | 0.000798452 | 0.002592785 | 0.002921076 | 0.002881669 |
| NITMOv2_3447 | A0A0K2GG49 | Putative TonB-dependent siderophore receptor | 89.781 | 807 | 0.409 | 1.33 | 7.71E-04 | 2.34E-03 | no | no | 0 | 5 | 5 | 6 | 4 | 4 | 4 | 0.001126133 | 0.001298392 | 0.001220225 | 0.000979765 | 0.000807308 | 0.000715401 |
| NITMOv2_1995 | A0A0K2GCV6 | Uncharacterized protein | 21.27 | 190 | 0.413 | 1.33 | 3.04E-01 | 3.46E-01 | no | yes | 1 | 1 | 0 | 0 | 1 | 2 | 0 | 0.001123098 | 0.001352616 | 0.001008504 | 0.000815609 | 0.00105467 | 0.00089203 |
| NITMOv2_3407 | A0A0K2GG17 | Oxidoreductase domain protein | 46.094 | 426 | -1.62 | -3.07 | 9.67E-07 | 2.93E-05 | yes | no | 0 | 2 | 2 | 4 | 18 | 18 | 19 | 0.001110851 | 0.001148684 | 0.00148187 | 0.006818833 | 0.007527119 | 0.007147532 |
| NITMOv2_1358 | A0A0K2G9Y7 | Conserved exported protein | 38.692 | 345 | -0.431 | -1.35 | 3.94E-03 | 8.20E-03 | no | no | 0 | 0 | 0 | 0 | 1 | 2 | 3 | 0.001107963 | 0.000980887 | 0.000895809 | 0.001409433 | 0.001287427 | 0.001468184 |
| NITMOv2_4803 | A0A0K2GJQ5 | Uncharacterized protein | 31.517 | 285 | 0.132 | 1.10 | 1.71E-01 | 2.08E-01 | no | no | 0 | 2 | 3 | 1 | 2 | 1 | 1 | 0.00110771 | 0.000859298 | 0.001164491 | 0.0012301 | 0.001006972 | 0.001196524 |
| holB | A0A0K2GFV6 | DNA polymerase III subunit delta | 39.64 | 364 | -0.667 | -1.59 | 1.18E-04 | 6.25E-04 | no | no | 0 | 4 | 5 | 5 | 7 | 7 | 5 | 0.001104317 | 0.001309734 | 0.000959557 | 0.00184338 | 0.002051636 | 0.001878857 |
| NITMOv2_0362 | A0A0K2G763 | Uncharacterized protein | 37.552 | 333 | -3.68 | -12.82 | 1.69E-04 | 7.81E-04 | yes | no | 0 | 2 | 2 | 2 | 24 | 23 | 21 | 0.001103874 | 0.000621849 | 0.001101203 | 0.017502412 | 0.017614083 | 0.017750155 |
| tdh | A0A0K2GJ09 | Threonine 3-dehydrogenase, NAD(P)-binding | 36.428 | 344 | -0.058 | -1.04 | 7.29E-01 | 7.61E-01 | no | no | 0 | 5 | 5 | 6 | 2 | 3 | 3 | 0.00110048 | 0.001342702 | 0.001350904 | 0.001505326 | 0.001060403 | 0.001134363 |
| NITMOv2_0242 | A0A0K2G6T0 | Putative RNA-metabolising metallo-beta-lactamase | 51.35 | 465 | -1.16 | -2.23 | 1.82E-06 | 4.01E-05 | yes | no | 0 | 3 | 2 | 2 | 7 | 9 | 6 | 0.001086042 | 0.00097465 | 0.001029134 | 0.003091797 | 0.003088718 | 0.002867464 |
| NITMOv2_3261 | A0A0K2GFC7 | Uncharacterized protein | 59.889 | 551 | -0.35 | -1.27 | 1.74E-02 | 2.83E-02 | no | no | 0 | 5 | 4 | 5 | 7 | 4 | 3 | 0.001085683 | 0.000966046 | 0.001131936 | 0.001614013 | 0.001205346 | 0.000935059 |
| NITMOv2_0805 | A0A0K2G8H4 | Uncharacterized protein | 56.569 | 498 | -1.51 | -2.85 | 6.16E-05 | 4.06E-04 | yes | no | 0 | 1 | 3 | 1 | 13 | 13 | 7 | 0.001076851 | 0.001070231 | 0.000754241 | 0.002842883 | 0.002759027 | 0.002140843 |
| NITMOv2_2691 | A0A0K2GER0 | Uncharacterized protein | 11.459 | 105 | -0.521 | -1.43 | 3.00E-01 | 3.42E-01 | no | no | 0 | 0 | 2 | 3 | 2 | 2 | 2 | 0.001075186 | 0.001799923 | 0.002481287 | 0.001271264 | 0.000888131 | 0.001030162 |
| NITMOv2_1729 | A0A0K2GBA0 | Elp3 domain-containing protein | 74.381 | 650 | -0.835 | -1.78 | 5.81E-04 | 1.91E-03 | no | no | 0 | 3 | 3 | 4 | 6 | 11 | 10 | 0.001071666 | 0.001316363 | 0.001377361 | 0.002936911 | 0.002804239 | 0.002782593 |
| dnaE | A0A0K2GC83 | DNA polymerase III subunit alpha | 127.2 | 1153 | -2.05 | -4.14 | 1.59E-07 | 1.13E-05 | yes | no | 0 | 5 | 5 | 7 | 37 | 43 | 33 | 0.001068736 | 0.001202634 | 0.001239071 | 0.007359157 | 0.007159673 | 0.006900111 |
| NITMOv2_3202 | A0A0K2GF71 | PBPb domain-containing protein | 40.451 | 362 | -0.69 | -1.61 | 4.85E-02 | 6.84E-02 | no | no | 0 | 4 | 3 | 3 | 4 | 2 | 3 | 0.001056068 | 0.000904743 | 0.00177028 | 0.001102978 | 0.000880223 | 0.001010436 |
| hisI | A0A0K2G8U1 | Histidine biosynthesis bifunctional protein HisIE | 25.566 | 230 | -0.815 | -1.76 | 6.36E-03 | 1.22E-02 | no | no | 0 | 3 | 2 | 3 | 2 | 2 | 2 | 0.001052337 | 0.000908653 | 0.001212613 | 0.002013666 | 0.001840957 | 0.001773721 |
| NITMOv2_1243 | A0A0K2G9M4 | Putative Type IV pilus biogenesis protein PilN | 20.958 | 183 | -1.8 | -3.48 | 2.17E-03 | 5.13E-03 | yes | yes | 2 | 0 | 0 | 0 | 0 | 2 | 0 | 0.001042303 | 0.000495332 | 0.000604735 | 0.004120322 | 0.004877161 | 0.004024375 |
| NITMOv2_0107 | A0A0K2G7G9 | Cytochrome_P460 domain-containing protein | 22.365 | 204 | 0.0218 | 1.02 | 8.27E-01 | 8.48E-01 | no | yes | 1 | 2 | 0 | 1 | 1 | 1 | 0 | 0.001041059 | 0.001106994 | 0.001069971 | 0.00074021 | 0.001503187 | 0.00185656 |
| NITMOv2_1330 | A0A0K2G9Y1 | Putative site-specific recombinase transmembrane protein | 78.408 | 714 | -0.341 | -1.27 | 1.23E-02 | 2.13E-02 | no | no | 0 | 3 | 2 | 2 | 5 | 5 | 3 | 0.001032944 | 0.000976468 | 0.001047041 | 0.0018034 | 0.001634628 | 0.001341956 |
| nuoL.1 | A0A0K2G7Z1 | NADH-quinone oxidoreductase, membrane subunit L | 67.807 | 630 | -0.68 | -1.60 | 6.78E-03 | 1.28E-02 | no | yes | 1 | 1 | 1 | 1 | 2 | 2 | 2 | 0.001031911 | 0.000806853 | 0.0006514 | 0.000835363 | 0.00145465 | 0.001706903 |
| rsmB | A0A0K2G746 | 16S rRNA m5C967 methyltransferase | 51.246 | 466 | 0.193 | 1.14 | 1.20E-01 | 1.51E-01 | no | no | 0 | 3 | 5 | 4 | 2 | 2 | 3 | 0.001028855 | 0.001286816 | 0.001191505 | 0.000947188 | 0.000873184 | 0.000803419 |
| NITMOv2_0781 | A0A0K2G9D2 | HEPN_Apea domain-containing protein | 53.678 | 470 | -0.776 | -1.71 | 3.72E-04 | 1.35E-03 | no | no | 0 | 7 | 8 | 6 | 9 | 10 | 11 | 0.001024154 | 0.001024631 | 0.001198579 | 0.00242324 | 0.002211044 | 0.00199106 |
| NITMOv2_3388 | A0A0K2GFN9 | N-acetyltransferase domain-containing protein | 38.42 | 346 | -2.87 | -7.31 | 2.64E-05 | 2.26E-04 | yes | yes | 3 | 1 | 0 | 0 | 1 | 1 | 1 | 0.001020044 | 0.001059633 | 0.000893719 | 0.001752462 | 0.001605729 | 0.001468652 |
| NITMOv2_2614 | A0A0K2GEK0 | Putative Heavy metal efflux system, outer membrane lipoprotein | 46.609 | 425 | -2.14 | -4.41 | 1.16E-06 | 3.12E-05 | yes | no | 0 | 4 | 4 | 5 | 15 | 15 | 14 | 0.001019496 | 0.001021776 | 0.000900181 | 0.006620413 | 0.006992342 | 0.006749968 |
| NITMOv2_1486 | A0A0K2GAE7 | Putative Histidine kinase | 51.537 | 471 | 1.83 | 3.56 | 1.07E-03 | 2.95E-03 | yes | yes | 2 | 3 | 4 | 3 | 0 | 0 | 0 | 0.001006279 | 0.001785296 | 0.001488235 | 0 | 0.000198794 | 0 |
| NITMOv2_4301 | A0A0K2GI83 | Thioredoxin domain-containing protein | 23.858 | 214 | -1.37 | -2.58 | 9.32E-04 | 2.69E-03 | yes | no | 0 | 1 | 2 | 1 | 4 | 4 | 5 | 0.001000251 | 0.001153807 | 0.001038778 | 0.004131872 | 0.005266513 | 0.005176971 |
| NITMOv2_4207 | A0A0K2GIY2 | TPR_REGION domain-containing protein | 66.263 | 588 | -1.28 | -2.43 | 1.64E-02 | 2.70E-02 | yes | yes | 3 | 0 | 1 | 0 | 3 | 1 | 1 | 0.000999281 | 0.000998936 | 0.000271289 | 0.000673606 | 0.000498965 | 0.000492577 |
| NITMOv2_4542 | A0A0K2GJ11 | Uncharacterized protein | 22.933 | 196 | -1.91 | -3.76 | 1.33E-05 | 1.42E-04 | yes | no | 0 | 1 | 0 | 1 | 6 | 8 | 5 | 0.000994939 | 0.001034818 | 0.001214569 | 0.003814548 | 0.004705168 | 0.004093423 |
| NITMOv2_4556 | A0A0K2GJ21 | DUF4105 domain-containing protein | 76.262 | 671 | -0.586 | -1.50 | 2.55E-02 | 3.92E-02 | no | no | 0 | 1 | 2 | 2 | 3 | 2 | 1 | 0.000992178 | 0.000839411 | 0.000889291 | 0.000824035 | 0.000800301 | 0.000717217 |
| NITMOv2_1252 | A0A0K2G9R2 | Uncharacterized protein | 15.607 | 145 | -0.0916 | -1.07 | 6.55E-01 | 6.93E-01 | no | yes | 2 | 2 | 2 | 1 | 2 | 0 | 1 | 0.000990365 | 0.000982608 | 0.000993454 | 0.000935224 | 0.000227909 | 0.000171762 |
| NITMOv2_1889 | A0A0K2GBR2 | Site-specific DNA-methyltransferase (adenine-specific) | 56.558 | 501 | -1.29 | -2.45 | 3.54E-06 | 6.20E-05 | yes | no | 0 | 4 | 6 | 5 | 12 | 10 | 9 | 0.00095822 | 0.001045593 | 0.001043992 | 0.002325215 | 0.003167645 | 0.002873758 |
| NITMOv2_1865 | A0A0K2GBF4 | Histidine kinase domain-containing protein | 86.443 | 779 | 0.126 | 1.09 | 3.48E-01 | 3.90E-01 | no | no | 0 | 2 | 4 | 8 | 7 | 7 | 7 | 0.000946374 | 0.001154374 | 0.000984788 | 0.000817312 | 0.000895324 | 0.000702274 |
| NITMOv2_1728 | A0A0K2GB23 | B12-binding domain-containing protein | 73.245 | 632 | -0.346 | -1.27 | 9.11E-03 | 1.64E-02 | no | no | 0 | 0 | 0 | 0 | 0 | 3 | 3 | 0.000945678 | 0.001174711 | 0.001090428 | 0.001965838 | 0.002062978 | 0.001623919 |
| NITMOv2_4178 | A0A0K2GHW5 | Asparagine synthetase domain-containing protein | 29.732 | 269 | -1.27 | -2.41 | 4.81E-06 | 7.30E-05 | yes | no | 0 | 1 | 3 | 3 | 6 | 7 | 7 | 0.000944308 | 0.000941368 | 0.000876311 | 0.003625902 | 0.004359785 | 0.003952091 |
| NITMOv2_4068 | A0A0K2GIM2 | VWFA domain-containing protein | 109.79 | 1006 | -0.693 | -1.62 | 9.68E-05 | 5.45E-04 | no | no | 0 | 5 | 3 | 3 | 4 | 5 | 6 | 0.000941568 | 0.001039335 | 0.00091617 | 0.00139849 | 0.001422799 | 0.0013168 |
| phoU | A0A0K2GJG8 | Phosphate-specific transport system accessory protein PhoU | 26.379 | 230 | 0.0318 | 1.02 | 9.54E-01 | 9.59E-01 | no | yes | 4 | 0 | 2 | 2 | 0 | 0 | 0 | 0.000934317 | 0.000944399 | 0.000854608 | 0.000876424 | 0.000726983 | 0.000515575 |
| NITMOv2_3749 | A0A0K2GGQ3 | Putative Sensor histidine kinase | 51.162 | 464 | 0.322 | 1.25 | 2.62E-01 | 3.04E-01 | no | no | 0 | 1 | 3 | 3 | 0 | 0 | 0 | 0.000931429 | 0.001214601 | 0.00157777 | 0.000450917 | 0.000427589 | 0.000423637 |
| NITMOv2_3658 | A0A0K2GGG6 | fn3_3 domain-containing protein | 34.49 | 312 | -3.01 | -8.06 | 1.61E-07 | 1.13E-05 | yes | no | 0 | 0 | 1 | 0 | 15 | 16 | 15 | 0.000930923 | 0.001333179 | 0.00137161 | 0.015583366 | 0.016593315 | 0.015104219 |
| cobD.2 | A0A0K2GBL2 | Cobalamin biosynthesis protein CobD | 33.87 | 318 | 0.233 | 1.18 | 1.83E-02 | 2.95E-02 | no | no | 0 | 2 | 3 | 3 | 1 | 1 | 3 | 0.000922028 | 0.000860843 | 0.000942264 | 0.000725965 | 0.000754188 | 0.007770402 |
| NITMOv2_1467 | A0A0K2GAI9 | Putative Peptidase M22 | 28.069 | 261 | -2.93 | -7.62 | 9.53E-04 | 2.72E-03 | yes | no | 0 | 2 | 3 | 3 | 8 | 8 | 5 | 0.000910898 | 0.00095621 | 0.002004278 | 0.00357526 | 0.002785129 | 0.003246328 |
| NITMOv2_1334 | A0A0K2GAZ2 | Uncharacterized protein | 35.775 | 333 | -0.104 | -1.07 | 4.21E-01 | 4.64E-01 | no | no | 0 | 1 | 2 | 1 | 2 | 1 | 2 | 0.000908664 | 0.000988123 | 0.000762331 | 0.001043348 | 0.000925015 | 0.000964548 |
| NITMOv2_2460 | A0A0K2GD50 | Putative 4-phosphopantetheinyl transferase HetI | 28.956 | 262 | -1.02 | -2.03 | 6.53E-02 | 8.85E-02 | no | no | 0 | 2 | 4 | 3 | 1 | 3 | 2 | 0.000908369 | 0.000928814 | 0.001167558 | 0.001181887 | 0.001239076 | 0.000980785 |
| NITMOv2_2552 | A0A0K2GDD9 | Uncharacterized protein | 73.104 | 665 | 2.27 | 4.82 | 2.77E-04 | 1.10E-03 | yes | yes | 3 | 2 | 1 | 2 | 1 | 1 | 2 | 0.000898714 | 0.000950265 | 0.00089949 | 0.000710832 | 0.000896629 | 0.000755139 |
| NITMOv2_2557 | A0A0K2GDE2 | Uncharacterized protein | 107.06 | 972 | -0.483 | -1.40 | 1.17E-03 | 3.16E-03 | no | no | 0 | 7 | 5 | 8 | 8 | 10 | 6 | 0.000897091 | 0.000813619 | 0.000900698 | 0.00102857 | 0.001150625 | 0.00079254 |
| cysD | A0A0K2G762 | ATP-sulfurylase small subunit | 30.742 | 265 | -3.6 | -12.13 | 2.35E-07 | 1.33E-05 | yes | no | 0 | 0 | 0 | 1 | 16 | 18 | 20 | 0.000895215 | 0.001078738 | 0.001008255 | 0.01350736 | 0.01133022 | 0.013306278 |
| NITMOv2_3492 | A0A0K2GGA1 | Asparagine synthetase | 67.657 | 606 | -1.98 | -3.94 | 2.90E-05 | 2.41E-04 | yes | no | 0 | 1 | 1 | 1 | 10 | 11 | 9 | 0.000892897 | 0.000775645 | 0.000957391 | 0.003644707 | 0.003183803 | 0.00344592 |

| | | | | | | | | | | | | | | | | | | | | | | | |
|---|---|---|---|---|---|---|---|---|---|---|---|---|---|---|---|---|---|---|---|---|---|---|---|
| NITMOv2_0499 | A0A0K2G7N6 | Radical SAM superfamily protein | 61.779 | 542 | -1.19 | -2.28 | 1.11E-05 | 1.28E-04 | yes | no | 0 | 1 | 2 | 2 | 10 | 6 | 3 | 0.000889208 | 0.001185113 | 0.000975605 | 0.004263955 | 0.004888658 | 0.002312923 |
| NITMOv2_4365 | A0A0K2GIH4 | Uncharacterized protein | 12.994 | 112 | -0.117 | -1.08 | 7.92E-01 | 8.19E-01 | no | yes | 2 | 1 | 1 | 2 | 0 | 2 | 0 | 0.00088807 | 0.001100326 | 0.001021216 | 0.000965757 | 0.001300012 | 0.001260645 |
| NITMOv2_2902 | A0A0K2GEC9 | Uncharacterized protein | 12.2 | 109 | -0.654 | -1.57 | 1.55E-02 | 2.59E-02 | no | yes | 2 | 1 | 1 | 1 | 2 | 1 | 0 | 0.000884065 | 0.000868371 | 0.000537248 | 0.001470484 | 0.000603559 | 0.00119415 |
| aglA | A0A0K2GHC2 | Putative alpha-glucosidase | 63.218 | 555 | 0.503 | 1.42 | 2.00E-03 | 4.80E-03 | no | no | 0 | 2 | 2 | 3 | 2 | 3 | 2 | 0.000876582 | 0.000835285 | 0.000897305 | 0.000548202 | 0.000740158 | 0.000806242 |
| NITMOv2_2503 | A0A0K2GD94 | Histidine kinase | 57.46 | 528 | -0.226 | -1.17 | 1.37E-01 | 1.71E-01 | no | no | 0 | 6 | 7 | 6 | 3 | 4 | 6 | 0.000875127 | 0.000947586 | 0.001075512 | 0.001125737 | 0.000967819 | 0.001088511 |
| NITMOv2_3391 | A0A0K2GFP3 | Putative methyltransferase | 30.738 | 272 | -2.05 | -4.14 | 2.75E-02 | 4.18E-02 | yes | yes | 1 | 1 | 2 | 0 | 5 | 3 | 3 | 0.000869541 | 0.001338224 | 0.000679008 | 0.005591591 | 0.003187377 | 0.003227808 |
| hypD | A0A088N9K3 | Hydrogenase expression/formation protein | 42.094 | 381 | 1.2 | 2.30 | 3.20E-02 | 4.79E-02 | yes | yes | 2 | 8 | 9 | 5 | 0 | 0 | 2 | 0.000868656 | 0.000535438 | 0.000797934 | 6.262665E-05 | 6.493455E-05 | 0.001348951 |
| NITMOv2_0303 | A0A0K2G6Z0 | Uncharacterized protein | 15.274 | 140 | -1.08 | -2.11 | 3.41E-02 | 5.03E-02 | no | no | 0 | 3 | 3 | 5 | 3 | 5 | 3 | 0.000868466 | 0.000553311 | 0.001357921 | 0.001436901 | 0.001233794 | 0.000937037 |
| NITMOv2_1263 | A0A0K2G9P5 | Putative urease accessory protein UreD | 31.466 | 293 | -0.423 | -1.34 | 9.74E-02 | 1.26E-01 | no | no | 0 | 1 | 1 | 2 | 2 | 2 | 6 | 0.000860709 | 0.001326726 | 0.000637385 | 0.001142765 | 0.001203792 | 0.001760595 |
| NITMOv2_4033 | A0A0K2GIJ5 | Putative Nitrite oxidoreductase, alpha subunit | 131.92 | 1145 | 0.376 | 1.30 | 1.58E-02 | 2.62E-02 | no | no | 0 | 3 | 4 | 4 | 3 | 2 | 5 | 0.000857168 | 0.000811781 | 0.000993205 | 0.000718946 | 0.000850547 | 0.000910407 |
| NITMOv2_4069 | A0A0K2GHM2 | Ferredoxin-thioredoxin reductase subunit B | 15.358 | 137 | -1.61 | -3.05 | 6.41E-04 | 2.05E-03 | yes | yes | 1 | 0 | 1 | 0 | 4 | 3 | 1 | 0.000854786 | 0.001137499 | 0.000476491 | 0.003648705 | 0.003143252 | 0.00332131 |
| tgt | A0A0K2GHD8 | Queuine tRNA-ribosyltransferase | 42.328 | 381 | -1.18 | -2.27 | 1.71E-03 | 4.26E-03 | yes | no | 0 | 0 | 1 | 0 | 3 | 3 | 3 | 0.000841549 | 0.000586026 | 0.000564492 | 0.002596191 | 0.002269928 | 0.00236381 |
| NITMOv2_0807 | A0A0K2G9G1 | Putative Hydrolase, alpha/beta fold family | 36.187 | 325 | 0.298 | 1.23 | 5.57E-02 | 7.70E-02 | no | no | 0 | 1 | 2 | 2 | 2 | 1 | 0 | 0.000824285 | 0.000810862 | 0.000732039 | 0.000704109 | 0.000635673 | 0.000527443 |
| gmhB | A0A0K2GGF9 | LPS heptosyltransferase II and D,D-heptose 1,7-bisphosphate phosphatase (Modular protein) | 64.512 | 594 | -0.936 | -1.91 | 1.41E-04 | 7.03E-04 | no | no | 0 | 3 | 5 | 6 | 9 | 14 | 11 | 0.000815875 | 0.001511926 | 0.001110789 | 0.003367363 | 0.003435034 | 0.00300538 |
| NITMOv2_3035 | A0A0K2GER1 | RNA polymerase sigma factor | 22.653 | 197 | -0.134 | -1.10 | 8.56E-01 | 8.74E-01 | no | yes | 2 | 1 | 1 | 1 | 1 | 0 | 0 | 0.000815791 | 0.000566745 | 0.000571106 | 0.000650802 | 0.000543851 | 8.753075E-05 |
| NITMOv2_3347 | A0A0K2GFV9 | Putative membrane protein | 70.494 | 656 | -1.79 | -3.46 | 3.70E-05 | 2.86E-04 | yes | no | 0 | 4 | 4 | 8 | 20 | 16 | 14 | 0.000802047 | 0.000734659 | 0.00102645 | 0.004119434 | 0.003624272 | 0.002879332 |
| kdpE | A0A0K2GAN2 | DNA-binding response regulator in two-component regulatory system with KdpD | 26.095 | 234 | 0.514 | 1.43 | 2.15E-02 | 3.39E-02 | no | yes | 3 | 3 | 2 | 2 | 2 | 1 | 1 | 0.000800993 | 0.000915263 | 0.000599577 | 0.0010213 | 0.000439754 | 0.000451886 |
| mutS.1 | A0A0K2GIR2 | DNA mismatch repair protein MutS | 98.113 | 884 | -1.5 | -2.83 | 5.90E-04 | 1.93E-03 | yes | no | 0 | 3 | 4 | 6 | 12 | 13 | 15 | 0.000789885 | 0.00101851 | 0.001018819 | 0.001969243 | 0.001893627 | 0.001838939 |
| coaBC | A0A0K2GCB7 | Coenzyme A biosynthesis bifunctional protein CoaBC | 43.977 | 417 | -1.44 | -2.71 | 7.85E-04 | 2.37E-03 | yes | no | 0 | 9 | 12 | 9 | 12 | 10 | 13 | 0.000789569 | 0.001555708 | 0.001231556 | 0.002796536 | 0.003535401 | 0.003227987 |
| NITMOv2_2056 | A0A0K2GC03 | MPN domain-containing protein | 25.752 | 237 | -1.86 | -3.63 | 1.35E-05 | 1.43E-04 | yes | no | 0 | 2 | 2 | 2 | 3 | 5 | 4 | 0.000789021 | 0.00047347 | 0.000706041 | 0.000930545 | 0.00090982 | 0.000883561 |
| glnD | A0A0K2GAR8 | Bifunctional uridylyltransferase/uridylyl-removing enzyme | 99.658 | 891 | -0.443 | -1.36 | 8.26E-04 | 2.45E-03 | no | no | 0 | 3 | 3 | 4 | 6 | 7 | 3 | 0.000788915 | 0.000917512 | 0.00091548 | 0.001456595 | 0.001714176 | 0.001307953 |
| fabG.3 | A0A0K2GC46 | 3-oxoacyl-(Acyl-carrier-protein) reductase | 25.827 | 242 | -1.05 | -2.07 | 9.67E-03 | 1.73E-02 | yes | no | 0 | 4 | 6 | 7 | 6 | 4 | 4 | 0.000785163 | 0.000704232 | 0.000227327 | 0.001115831 | 0.000331913 | 0.000803419 |
| NITMOv2_1352 | A0A0K2GA06 | Peptidoglycan-binding lipoprotein, OmpA family (Modular protein) | 49.31 | 442 | 0.61 | 1.53 | 2.19E-03 | 5.16E-03 | no | yes | 1 | 0 | 1 | 0 | 0 | 2 | 0 | 0.00077865 | 0.000723884 | 0.000899663 | 0.000305448 | 0.000404641 | 0.000431333 |
| cbiA | A0A0K2GBA6 | Cobyrinate a,c-diamide synthase | 49.672 | 468 | -1.29 | -2.45 | 9.48E-05 | 5.41E-04 | yes | no | 0 | 1 | 1 | 1 | 9 | 7 | 6 | 0.000777048 | 0.000925236 | 0.000869562 | 0.00232714 | 0.001902638 | 0.001963369 |
| hypF | A0A088NA18 | Carbamoyltransferase | 87.86 | 800 | 2.39 | 5.24 | 2.19E-04 | 9.41E-04 | yes | yes | 3 | 4 | 4 | 3 | 0 | 0 | 0 | 0.00075481 | 0.000641189 | 0.000961532 | 0 | 6.767058E-05 | 0 |
| NITMOv2_1938 | A0A0K2GCN5 | Putative Micrococcal nuclease-like nuclease | 27.824 | 248 | -0.22 | -1.16 | 9.22E-02 | 1.20E-01 | no | no | 0 | 1 | 2 | 2 | 2 | 3 | 4 | 0.000751795 | 0.000712542 | 0.000610026 | 0.001622305 | 0.001762341 | 0.001327535 |
| NITMOv2_3412 | A0A0K2GG21 | Putative Undecaprenyl-phosphate galactose phosphotransferase RfbP | 40.214 | 356 | -0.74 | -1.67 | 3.29E-03 | 7.13E-03 | no | no | 0 | 2 | 2 | 1 | 1 | 0 | 3 | 0.000750678 | 0.001236053 | 0.000735912 | 0.001699007 | 0.00140821 | 0.001693543 |
| NITMOv2_4520 | A0A0K2GJ47 | Putative Glutathione-regulated potassium-efflux system | 72.243 | 666 | 0.0332 | 1.02 | 7.12E-01 | 7.47E-01 | no | no | 0 | 3 | 3 | 2 | 4 | 3 | 2 | 0.000741572 | 0.000714361 | 0.000669863 | 0.000794316 | 0.000785526 | 0.000634845 |
| fhs | A0A0K2GHY3 | Formate--tetrahydrofolate ligase | 58.732 | 560 | -0.366 | -1.29 | 2.29E-02 | 3.58E-02 | no | no | 0 | 3 | 4 | 3 | 2 | 5 | 3 | 0.000739338 | 0.000907441 | 0.000693023 | 0.001778375 | 0.001672227 | 0.001476869 |
| NITMOv2_4198 | A0A0K2GHY2 | Transcriptional regulator, MerR family | 16.849 | 147 | 0.371 | 1.29 | 2.75E-01 | 3.18E-01 | no | yes | 4 | 0 | 1 | 2 | 1 | 0 | 0 | 0.000732888 | 0.001421291 | 0.001449775 | 9.884124E-05 | 0.000100281 | 0.000468716 |
| NITMOv2_2715 | A0A0K2GDU6 | OMP_b-brl domain-containing protein | 37.743 | 349 | 0.594 | 1.51 | 2.16E-01 | 2.57E-01 | no | yes | 3 | 1 | 1 | 2 | 0 | 1 | 0 | 0.000724267 | 0.000215313 | 0.000562536 | 0.000505142 | 0.00019898 | 0.000385913 |
| mviN | A0A0K2GJ81 | Probable lipid II flippase MurJ | 57.093 | 538 | -0.237 | -1.18 | 2.63E-01 | 3.05E-01 | no | no | 0 | 3 | 4 | 3 | 5 | 4 | 2 | 0.000723571 | 0.000771147 | 0.000678874 | 0.000988486 | 0.00100374 | 0.000854809 |
| NITMOv2_0038 | A0A0K2G7C2 | Putative Radical SAM protein | 70.67 | 618 | -1.35 | -2.55 | 1.43E-05 | 1.48E-04 | yes | no | 0 | 2 | 4 | 3 | 12 | 12 | 9 | 0.000721906 | 0.000552216 | 0.001071601 | 0.002182915 | 0.002602882 | 0.002237043 |
| NITMOv2_1991 | A0A0K2GBU1 | Uncharacterized protein | 9.3475 | 84 | -0.813 | -1.76 | 1.71E-01 | 2.07E-01 | no | yes | 4 | 0 | 0 | 0 | 1 | 2 | 2 | 0.000721611 | 0.000681666 | 0.000555749 | 0.003148953 | 0.00209545 | 0.003192385 |
| silA | A0A0K2GBM3 | Putative cation efflux system protein SilA | 116.48 | 1052 | 0.17 | 1.13 | 8.30E-02 | 1.10E-01 | no | no | 0 | 6 | 7 | 9 | 2 | 4 | 2 | 0.000721063 | 0.000601786 | 0.000496257 | 0.000443454 | 0.000398084 | 0.000456363 |
| mtgA | A0A0K2GK69 | Biosynthetic peptidoglycan transglycosylase | 27.495 | 246 | -0.285 | -1.22 | 4.14E-01 | 4.58E-01 | no | yes | 3 | 2 | 2 | 1 | 0 | 0 | 0 | 0.000719566 | 0.000724432 | 0.000777842 | 0 | 0 | 0.000100389 |
| NITMOv2_4595 | A0A0K2GK38 | Uncharacterized protein | 26.816 | 246 | -0.41 | -1.33 | 2.05E-02 | 3.24E-02 | no | no | 0 | 3 | 2 | 3 | 1 | 3 | 4 | 0.00071261 | 0.000467819 | 0.000487821 | 0.0009244 | 0.000960299 | 0.001198142 |
| NITMOv2_4734 | A0A0K2GJH4 | Uncharacterized protein | 54.406 | 479 | 1.18 | 2.27 | 1.34E-04 | 6.79E-04 | yes | yes | 3 | 1 | 2 | 1 | 1 | 0 | 1 | 0.000710502 | 0.000740349 | 0.000985287 | 0.000761074 | 0.000901384 | 0.000875595 |
| NITMOv2_3379 | A0A0K2GFP0 | Glycosyl transferase, group 1 | 48.086 | 431 | -0.788 | -1.73 | 1.49E-02 | 2.49E-02 | no | no | 0 | 0 | 0 | 0 | 4 | 3 | 1 | 0.000705001 | 0.000745198 | 0.000730352 | 0.001431733 | 0.001540692 | 0.001373009 |
| NITMOv2_3987 | A0A0K2GHG1 | Uncharacterized protein | 181.26 | 1646 | 0.954 | 1.94 | 4.96E-04 | 1.68E-03 | no | no | 0 | 6 | 3 | 8 | 1 | 1 | 1 | 0.000703588 | 0.00047482 | 0.000841206 | 0.000241747 | 0.000302719 | 0.000131248 |
| NITMOv2_0720 | A0A0K2G857 | Putative Adenosylhomocysteine nucleosidase | 29.815 | 279 | -2.52 | -5.74 | 8.66E-05 | 5.13E-04 | yes | yes | 3 | 1 | 1 | 1 | 1 | 1 | 2 | 0.000702429 | 0.000604954 | 0.000666086 | 0.003873482 | 0.003990164 | 0.003614943 |
| NITMOv2_2212 | A0A0K2GCF3 | Uncharacterized protein | 55.643 | 508 | 1.56 | 2.95 | 9.12E-02 | 1.19E-01 | no | yes | 2 | 1 | 4 | 2 | 0 | 0 | 0 | 0.000698867 | 0.000761761 | 0.000754969 | 5.775648E-05 | 5.56078E-05 | 0.000120533 |
| NITMOv2_4056 | A0A0K2GHL6 | Putative ABC-type transport system, ATPase component | 69.415 | 613 | -2.45 | -5.46 | 1.44E-05 | 1.48E-04 | yes | no | 0 | 1 | 1 | 0 | 3 | 4 | 3 | 0.000698803 | 0.000944066 | 0.000556305 | 0.001941257 | 0.002164433 | 0.00180873 |
| NITMOv2_2555 | A0A0K2GDE7 | Uncharacterized protein | 72.692 | 674 | 1.64 | 3.12 | 2.43E-02 | 3.76E-02 | yes | yes | 2 | 2 | 2 | 0 | 0 | 0 | 0 | 0.000696864 | 0.00065386 | 0.000538302 | 0.00015576 | 0.000149438 | 0.000147498 |
| NITMOv2_3369 | A0A0K2GFN2 | Putative General secretion pathway protein D | 84.165 | 770 | -0.208 | -1.16 | 1.01E-01 | 1.30E-01 | no | no | 0 | 3 | 2 | 4 | 6 | 5 | 6 | 0.000696843 | 0.00068063 | 0.001018436 | 0.001514951 | 0.001404901 | 0.001201738 |
| NITMOv2_4516 | A0A0K2GIX2 | Formimidoyltetrahydrofolate cyclodeaminase | 52.329 | 491 | -1.13 | -2.19 | 1.55E-02 | 2.59E-02 | yes | no | 0 | 0 | 1 | 3 | 4 | 1 | 2 | 0.000694419 | 0.001031787 | 0.001026661 | 0.001279956 | 0.001225295 | 0.00130513 |
| NITMOv2_2945 | A0A0K2GEG1 | Exonuclease domain-containing protein | 20.882 | 186 | -0.19 | -1.14 | 1.45E-01 | 1.79E-01 | no | yes | 1 | 1 | 1 | 1 | 1 | 2 | 2 | 0.000690878 | 0.001069469 | 0.000635851 | 0.000903595 | 0.000889793 | 0.000740484 |
| scpA | A0A0K2GAZ7 | Segregation and condensation protein A | 30.84 | 268 | -1.77 | -3.41 | 4.58E-04 | 1.59E-03 | yes | no | 0 | 0 | 0 | 0 | 3 | 4 | 4 | 0.00069014 | 0.000521222 | 0.000467422 | 0.004875355 | 0.004458599 | 0.004560395 |
| NITMOv2_3481 | A0A0K2GFZ1 | Ribosomal protein S23 | 14.462 | 121 | -0.42 | -1.34 | 2.23E-02 | 3.49E-02 | no | no | 0 | 2 | 0 | 2 | 2 | 2 | 1 | 0.00067958 | 0.001565212 | 0.001218844 | 0.001186255 | 0.000966794 | 0.001312683 |
| napG | A0A0K2GGD4 | Ferredoxin-type protein NapG | 21.258 | 200 | 0.221 | 1.17 | 7.46E-01 | 7.76E-01 | no | yes | 4 | 1 | 2 | 3 | 1 | 1 | 1 | 0.000678758 | 0.001709347 | 0.002032078 | 0.00079522 | 0.000730727 | 0.000741509 |
| NITMOv2_0006 | A0A0K2G6A1 | Uncharacterized protein | 17.415 | 158 | 0.0996 | 1.07 | 6.68E-01 | 7.05E-01 | no | yes | 3 | 1 | 0 | 5 | 4 | 3 | 4 | 0.000676291 | 0.000436591 | 0.000627166 | 0.001217839 | 0.000589544 | 0.000291619 |
| NITMOv2_1821 | A0A0K2GB97 | Uncharacterized protein | 16.817 | 157 | -2.06 | -4.17 | 1.93E-06 | 4.12E-05 | yes | no | 0 | 0 | 0 | 0 | 2 | 3 | 4 | 0.000667038 | 0.000600906 | 0.000564511 | 0.002352608 | 0.004269671 | 0.003977984 |
| cobT | A0A0K2GBR3 | Nicotinate-nucleotide--dimethylbenzimidazole phosphoribosyltransferase | 36.415 | 352 | -3.76 | -13.55 | 3.93E-06 | 6.39E-05 | yes | no | 0 | 3 | 1 | 3 | 26 | 24 | 19 | 0.000666616 | 0.001145027 | 0.001500448 | 0.024433782 | 0.02309547 | 0.020351137 |
| vapC.1 | A0A0K2G949 | Ribonuclease VapC | 14.724 | 132 | 0.0463 | 1.03 | 9.42E-01 | 9.50E-01 | no | yes | 2 | 2 | 1 | 2 | 0 | 1 | 0 | 0.000663475 | 0.000789861 | 0.000563725 | 0.000784588 | 0.000801482 | 0.000145957 |
| NITMOv2_0267 | A0A0K2G6V5 | Putative Uncharacterized ABC-type transport system, permease component YbbP | 95.437 | 893 | 0.496 | 1.41 | 4.69E-02 | 6.64E-02 | no | no | 0 | 3 | 2 | 2 | 3 | 2 | 2 | 0.000659766 | 0.000680121 | 0.00192989 | 0.000906971 | 0.000900545 | 0.000640527 |
| NITMOv2_0468 | A0A0K2G7T0 | Transport permease protein | 30.814 | 280 | -0.879 | -1.84 | 1.06E-01 | 1.36E-01 | no | no | 0 | 1 | 1 | 2 | 0 | 3 | 2 | 0.000654749 | 0.000622143 | 0.000939273 | 0.001680498 | 0.00172847 | 0.001636506 |
| cbiH | A0A0K2GAP9 | Cobalt-precorrin-3B C(17)-methyltransferase | 29.767 | 278 | -3.38 | -10.41 | 6.43E-04 | 2.05E-03 | yes | yes | 2 | 0 | 0 | 0 | 4 | 7 | 5 | 0.000654728 | 0.000296034 | 0.000327579 | 0.015330158 | 0.01474288 | 0.014073176 |
| NITMOv2_0757 | A0A0K2G8A7 | Uncharacterized protein | 49.078 | 457 | -0.21 | -1.16 | 2.53E-01 | 2.95E-01 | no | no | 0 | 4 | 3 | 2 | 0 | 0 | 0 | 0.000651819 | 0.000775879 | 0.000775349 | 0.000592269 | 0.000748129 | 0.000358977 |
| NITMOv2_3507 | A0A0K2GGE1 | Glycos_transf_1 domain-containing protein | 40.053 | 359 | -1.47 | -2.77 | 2.26E-05 | 2.06E-04 | yes | no | 0 | 3 | 2 | 2 | 8 | 7 | 5 | 0.000642165 | 0.000701142 | 0.000980379 | 0.002497573 | 0.002557359 | 0.002515572 |
| NITMOv2_2759 | A0A0K2GE79 | Nitrogen regulation protein B | 102.11 | 922 | -0.127 | -1.09 | 3.33E-01 | 3.75E-01 | no | no | 0 | 3 | 3 | 4 | 4 | 3 | 3 | 0.000621739 | 0.000673199 | 0.000633589 | 0.00048527 | 0.000638439 | 0.000808867 |
| lgt | A0A0K2GDI3 | Phosphatidylglycerol--prolipoprotein diacylglyceryl transferase | 30.818 | 275 | -1.79 | -3.46 | 8.63E-03 | 1.58E-02 | yes | no | 0 | 6 | 7 | 8 | 9 | 9 | 9 | 0.000618093 | 0.000532564 | 0.001124267 | 0.001744466 | 0.001424306 | 0.001082631 |
| NITMOv2_4682 | A0A0K2G6I3 | Uncharacterized protein | 18.321 | 165 | -0.995 | -1.99 | 3.81E-04 | 1.37E-03 | no | no | 0 | 2 | 2 | 1 | 3 | 2 | 2 | 0.000617081 | 0.000635205 | 0.000836394 | 0.002458482 | 0.001368933 | 0.002521146 |
| NITMOv2_1324 | A0A0K2GAY1 | Uncharacterized protein | 14.923 | 133 | -0.105 | -1.08 | 8.53E-01 | 8.71E-01 | no | yes | 2 | 2 | 2 | 2 | 0 | 1 | 1 | 0.000616132 | 0.00081137 | 0.000738212 | 0.000680995 | 0.000374126 | 0.000330603 |
| NITMOv2_3167 | A0A0K2GF37 | HTH cro/C1-type domain-containing protein | 19.799 | 176 | -0.787 | -1.73 | 1.41E-01 | 1.75E-01 | no | yes | 1 | 1 | 0 | 3 | 1 | 1 | 1 | 0.000615606 | 0.000604954 | 0.000672796 | 0.000593379 | 0.000140585 | 0.000163688 |
| NITMOv2_2319 | A0A0K2GCZ2 | Putative Nudix hydrolase, MutT family | 19.723 | 178 | -1.5 | -2.83 | 6.95E-04 | 2.17E-03 | yes | no | 0 | 8 | 10 | 15 | 17 | 15 | 18 | 0.000608333 | 0.000797819 | 0.001246338 | 0.00307773 | 0.002649959 | 0.00218202 |
| NITMOv2_3486 | A0A0K2GFZ6 | Putative NAD-dependent epimerase/dehydratase | 36.563 | 335 | -0.448 | -1.36 | 1.77E-02 | 2.87E-02 | no | no | 0 | 1 | 2 | 0 | 3 | 4 | 3 | 0.000606499 | 0.000726113 | 0.000692467 | 0.001354008 | 0.001775547 | 0.002074672 |
| alr | A0A0K2GBC5 | Alanine racemase | 40.875 | 386 | -3.21 | -9.25 | 1.69E-05 | 1.66E-04 | yes | yes | 1 | 1 | 0 | 0 | 7 | 6 | 5 | 0.000605551 | 0.00066921 | 0.000555768 | 0.005487051 | 0.006384231 | 0.005299243 |
| NITMOv2_4487 | A0A0K2GIU6 | Inhibitor_I42 domain-containing protein | 12.794 | 114 | -3.37 | -10.34 | 1.68E-05 | 1.66E-04 | yes | yes | 3 | 0 | 0 | 0 | 5 | 5 | 4 | 0.00060456 | 0.000167161 | 5.830122E-05 | 0.018161344 | 0.006616196 | 0.004711258 |
| pabB | A0A0K2GHW7 | Para-aminobenzoate synthase component 1 | 54.189 | 490 | -0.407 | -1.33 | 5.12E-02 | 7.18E-02 | no | no | 0 | 3 | 3 | 3 | 4 | 3 | 3 | 0.000596002 | 0.000863326 | 0.00074197 | 0.00037297 | 0.000387535 | 0.000386147 |
| NITMOv2_1888 | A0A0K2GBI2 | AAA_15 domain-containing protein | 69.567 | 622 | -1.29 | -2.45 | 2.32E-03 | 5.39E-03 | yes | no | 0 | 1 | 0 | 1 | 4 | 3 | 6 | 0.000578865 | 0.000466529 | 0.000524843 | 0.002598412 | 0.002611272 | 0.002447783 |
| NITMOv2_4156 | A0A0K2GHX8 | Uncharacterized protein | 12.775 | 118 | 0.331 | 1.26 | 1.61E-01 | 1.97E-01 | no | yes | 1 | 2 | 2 | 2 | 0 | 0 | 0 | 0.00057701 | 0.000650418 | 0.001195722 | 0.000454781 | 0.000564701 | 0.000659083 |
| dbh | A0A0K2GIQ3 | DNA polymerase IV | 40.916 | 369 | 0.558 | 1.47 | 5.79E-04 | 1.90E-03 | no | no | 0 | 3 | 2 | 2 | 0 | 1 | 1 | 0.000576315 | 0.000573178 | 0.00066808 | 0.000288183 | 0.000505257 | 0.000362501 |
| NITMOv2_1783 | A0A0K2GB82 | Putative Competence protein ComEC/Rec2 related protein | 92.658 | 855 | 0.0845 | 1.06 | 4.79E-01 | 5.22E-01 | no | no | 0 | 1 | 0 | 1 | 5 | 3 | 1 | 0.000569928 | 0.000490268 | 0.000403233 | 0.000548039 | 0.000548496 | 0.000478822 |
| NITMOv2_3043 | A0A0K2GER8 | Putative Patatin-like phospholipase | 35.515 | 316 | -0.13 | -1.09 | 3.12E-01 | 3.53E-01 | no | no | 0 | 1 | 2 | 0 | 5 | 3 | 1 | 0.000567061 | 0.000628165 | 0.000821708 | 0.001050604 | 0.001072226 | 0.000657177 |
| NITMOv2_0331 | A0A0K2G7C4 | HTH_8 domain-containing protein | 21.609 | 196 | -2.07 | -4.20 | 4.66E-04 | 1.62E-03 | yes | yes | 4 | 0 | 0 | 0 | 2 | 1 | 1 | 0.000562571 | 0.000321983 | 0 | 0.003514994 | 0.003325965 | 0.003102658 |
| NITMOv2_3976 | A0A0K2GHC6 | RND-type efflux transporter, transmembrane pump | 119.54 | 1097 | 2.01 | 4.03 | 1.43E-03 | 3.71E-03 | yes | yes | 3 | 2 | 2 | 1 | 0 | 0 | 0 | 0.000562023 | 0.000648795 | 0.000602894 | 0 | 7.260352E-05 | 1.636992E-05 |
| NITMOv2_2888 | A0A0K2GEK9 | Uncharacterized protein | 80.884 | 732 | -0.359 | -1.28 | 2.11E-02 | 3.34E-02 | no | no | 0 | 3 | 3 | 0 | 5 | 2 | 4 | 0.000559915 | 0.000554426 | 0.000544418 | 0.000813359 | 0.000798499 | 0.000673594 |
| NITMOv2_0864 | A0A0K2G8W1 | Putative Amidase enhancer protein LytB | 44.047 | 406 | -1.58 | -2.99 | 4.31E-03 | 8.81E-03 | yes | yes | 3 | 1 | 0 | 0 | 2 | 1 | 1 | 0.000559578 | 0.000720951 | 0.000339811 | 0.000970184 | 0.001090544 | 0.001003262 |
| mltC | A0A0K2GB66 | Membrane-bound lytic murein transglycosylase C | 45.924 | 410 | -1 | -2.00 | 4.16E-05 | 3.06E-04 | yes | no | 0 | 6 | 5 | 5 | 7 | 6 | 9 | 0.000557365 | 0.000447307 | 0.00076051 | 0.001822798 | 0.00195826 | 0.001871125 |
| NITMOv2_3491 | A0A0K2GG01 | Putative Glycosyl transferase, group 1 | 44.975 | 394 | -1.26 | -2.39 | 8.59E-05 | 5.11E-04 | yes | no | 0 | 2 | 3 | 2 | 7 | 8 | 7 | 0.000546488 | 0.000814792 | 0.00105308 | 0.003180345 | 0.003949613 | 0.003642634 |

Table S1

| | | | | | | | | | | | | | | | | | | | | | |
|---|---|---|---|---|---|---|---|---|---|---|---|---|---|---|---|---|---|---|---|---|---|
| dnaG | A0A0K2G8I1 | DNA primase | 65.86 | 606 | -2.94 | -7.67 | 8.04E-07 | 2.73E-05 | yes | no | 0 | 1 | 1 | 1 | 15 | 16 | 14 | 0.000540165 | 0.00052965 | 0.000436765 | 0.00353069 | 0.003393861 | 0.003643893 |
| NITMOv2_0564 | A0A0K2G7U3 | Uncharacterized protein | 55.321 | 508 | 0.107 | 1.08 | 4.43E-01 | 4.86E-01 | no | yes | 1 | 4 | 4 | 5 | 0 | 0 | 3 | 0.000536075 | 0.00057093 | 0.000752975 | 0.000235009 | 0.000154927 | 0.000538771 |
| NITMOv2_0466 | A0A0K2G8J8 | Putative Peptidase S16, lon-like | 26.504 | 230 | -1.27 | -2.41 | 8.12E-04 | 2.43E-03 | yes | no | 0 | 3 | 4 | 3 | 6 | 4 | 4 | 0.000531733 | 0.000203698 | 0.00030277 | 0.001669392 | 0.001482227 | 0.001137797 |
| ftsZ | A0A0K2G883 | Cell division protein FtsZ | 42.335 | 399 | -7.87 | -233.94 | 3.52E-09 | 1.79E-06 | yes | no | 0 | 0 | 0 | 0 | 37 | 31 | 42 | 0.0005307 | 0.000501922 | 0.000515698 | 0.045417426 | 0.04283965 | 0.039215228 |
| NITMOv2_1307 | A0A0K2G9W7 | Uncharacterized protein | 24.934 | 218 | -0.487 | -1.40 | 3.75E-02 | 5.47E-02 | no | no | 0 | 0 | 1 | 0 | 1 | 0 | 2 | 0.000529541 | 0.000557437 | 0.000667466 | 0.000678048 | 0.000571087 | 0.000602389 |
| NITMOv2_0566 | A0A0K2G8T2 | Putative TonB-dependent receptor | 75.482 | 685 | 0.674 | 1.60 | 2.54E-02 | 3.91E-02 | no | yes | 1 | 1 | 1 | 2 | 0 | 0 | 0 | 0.000525915 | 0.000499497 | 0.000567578 | 0.000316065 | 0.000311047 | 0.000326107 |
| rfbF | A0A0K2GFR5 | Glucose-1-phosphate cytidylyltransferase | 30.439 | 267 | -2.73 | -6.63 | 2.84E-08 | 4.25E-06 | yes | no | 0 | 1 | 1 | 1 | 6 | 4 | 5 | 0.000517336 | 0.000601024 | 0.000688173 | 0.005144998 | 0.004916779 | 0.005002733 |
| NITMOv2_3955 | A0A0K2GHJ4 | Putative Sulfite:cytochrome c oxidoreductase, subunit A | 43.815 | 392 | 0.567 | 1.48 | 3.45E-01 | 3.88E-01 | no | yes | 2 | 1 | 1 | 0 | 1 | 1 | 1 | 0.000510127 | 0.000657321 | 0.000524326 | 0.000256095 | 0.000429577 | 0.000110119 |
| NITMOv2_3900 | A0A0K2GI41 | Putative Response regulator/sensory box histidine kinase | 72.861 | 655 | -1.19 | -2.28 | 1.16E-04 | 6.18E-04 | yes | no | 0 | 1 | 0 | 1 | 2 | 3 | 3 | 0.00050901 | 0.000375268 | 0.000366882 | 0.000492585 | 0.000429422 | 0.00087279 |
| trmD | A0A0K2GH73 | tRNA (guanine-N(1)-)-methyltransferase | 29.197 | 261 | -3.48 | -11.16 | 2.31E-05 | 2.09E-04 | yes | no | 0 | 1 | 1 | 3 | 25 | 24 | 25 | 0.000497101 | 0.000605951 | 0.001204599 | 0.013144282 | 0.012656442 | 0.012154581 |
| NITMOv2_3588 | A0A0K2GGA2 | Helix-turn-helix, type 11 domain protein | 35.683 | 314 | -0.0648 | -1.05 | 7.34E-01 | 7.66E-01 | no | no | 0 | 1 | 1 | 0 | 4 | 1 | 1 | 0.000496026 | 0.000689859 | 0.000837813 | 0.000927776 | 0.001021499 | 0.000870722 |
| cobM | A0A0K2GAQ3 | Precorrin-4 C(11)-methyltransferase | 28.253 | 263 | -4.09 | -17.03 | 2.70E-08 | 4.25E-06 | yes | no | 0 | 0 | 0 | 0 | 13 | 15 | 13 | 0.000495225 | 0.000642362 | 0.000664226 | 0.011845816 | 0.010597193 | 0.011890257 |
| NITMOv2_1374 | A0A0K2G9Z9 | CMP/dCMP-type deaminase domain-containing protein | 42.934 | 381 | -2.39 | -5.24 | 3.91E-06 | 6.39E-05 | yes | no | 0 | 3 | 3 | 2 | 8 | 6 | 7 | 0.000489997 | 0.000573198 | 0.000631557 | 0.007533737 | 0.006806832 | 0.007499964 |
| msbA.1 | A0A0K2GHZ7 | Lipid A export ATP-binding/permease protein MsbA | 61.554 | 562 | -1.12 | -2.17 | 1.06E-04 | 5.77E-04 | yes | no | 0 | 0 | 0 | 0 | 5 | 3 | 3 | 0.000489807 | 0.000673023 | 0.000485597 | 0.001243737 | 0.001457586 | 0.003282111 |
| NITMOv2_4157 | A0A0K2GIU5 | Putative Methyltransferase, UbiE-family | 42.781 | 395 | -2.32 | -4.99 | 7.54E-04 | 2.30E-03 | yes | yes | 1 | 0 | 0 | 0 | 6 | 6 | 4 | 0.00048595 | 0.000347012 | 0.000153992 | 0.00297615 | 0.004025432 | 0.003337493 |
| NITMOv2_3131 | A0A0K2GEZ4 | HTH cro/C1-type domain-containing protein | 24.652 | 225 | -1.96 | -3.89 | 7.37E-04 | 2.27E-03 | yes | yes | 3 | 0 | 0 | 0 | 2 | 1 | 0 | 0.000481523 | 0.000489701 | 0.00045542 | 0.003423928 | 0.003470613 | 0.003447178 |
| NITMOv2_3380 | A0A0K2GGM5 | Putative Methyltransferase type 11 | 29.626 | 259 | -0.523 | -1.44 | 3.93E-01 | 4.36E-01 | no | yes | 4 | 1 | 1 | 1 | 0 | 2 | 2 | 0.000478762 | 0.000283695 | 0.000291535 | 0.000204535 | 0.001008432 | 0.000880108 |
| rnc | A0A0K2G815 | Ribonuclease 3 | 26.681 | 241 | -0.912 | -1.88 | 8.51E-02 | 1.12E-01 | no | yes | 4 | 0 | 0 | 0 | 2 | 1 | 1 | 0.000478551 | 0.000254266 | 0.000249873 | 0.001170174 | 0.001229521 | 0.001266471 |
| NITMOv2_4840 | A0A0K2GJS6 | dTDP-4-dehydrorhamnose reductase | 32.078 | 291 | -1.23 | -2.35 | 1.28E-02 | 2.19E-02 | yes | no | 0 | 2 | 1 | 2 | 4 | 5 | 2 | 0.000472923 | 0.000485438 | 0.00069749 | 0.000962055 | 0.001077991 | 0.000946783 |
| udgA | A0A0K2GAD7 | Type-4 uracil-DNA glycosylase | 23.849 | 216 | -1.96 | -3.89 | 3.25E-04 | 1.22E-03 | yes | yes | 2 | 0 | 0 | 0 | 4 | 4 | 4 | 0.00047168 | 0.000433834 | 0.000739401 | 0.003810699 | 0.004005856 | 0.003221694 |
| NITMOv2_4096 | A0A0K2GHP5 | Uncharacterized protein | 15.525 | 147 | -1.92 | -3.78 | 4.30E-03 | 8.81E-03 | yes | yes | 2 | 0 | 0 | 0 | 1 | 2 | 2 | 0.000470541 | 0.000445684 | 0.000805028 | 0.00313074 | 0.00335968 | 0.003305486 |
| NITMOv2_1236 | A0A0K2G9X0 | OstA-like_N domain-containing protein | 20.93 | 196 | -0.202 | -1.15 | 5.25E-01 | 5.67E-01 | no | no | 0 | 1 | 2 | 3 | 2 | 2 | 2 | 0.000460107 | 0.001157522 | 0.001080324 | 0.000566829 | 0.001459854 | 0.001688022 |
| NITMOv2_1498 | A0A0K2GAP2 | Putative Transcriptional regulator | 37.421 | 328 | -1.15 | -2.22 | 2.54E-02 | 3.91E-02 | yes | yes | 1 | 2 | 2 | 5 | 4 | 4 | 4 | 0.000459053 | 0.000646449 | 0.000531534 | 0.000898428 | 0.000678026 | 0.000272379 |
| NITMOv2_3760 | A0A0K2GGT5 | Putative PP-loop ATPase, YdaO type | 34.751 | 312 | -0.722 | -1.65 | 2.43E-03 | 5.59E-03 | no | no | 0 | 2 | 3 | 5 | 5 | 4 | 2 | 0.000458758 | 0.000369715 | 0.001023056 | 0.001779116 | 0.001716352 | 0.001316818 |
| NITMOv2_0551 | A0A0K2G8R9 | Uncharacterized protein | 46.31 | 425 | -2.34 | -5.06 | 1.51E-04 | 7.40E-04 | yes | yes | 1 | 0 | 0 | 0 | 6 | 5 | 6 | 0.000457978 | 0.000190174 | 0.000606767 | 0.00376598 | 0.004581184 | 0.006674626 |
| nrdJ | A0A0K2GCN6 | Vitamin B12-dependent ribonucleotide reductase | 130.54 | 1176 | -5.03 | -32.67 | 1.71E-05 | 1.67E-04 | yes | no | 0 | 2 | 0 | 3 | 53 | 52 | 50 | 0.00045722 | 0.000186351 | 0.000515621 | 0.015445656 | 0.015093702 | 0.014906246 |
| NITMOv2_0404 | A0A0K2G7C9 | Radical SAM (Modular protein) | 73.298 | 637 | -5.37 | -41.36 | 1.52E-06 | 3.61E-05 | yes | no | 0 | 0 | 0 | 0 | 19 | 22 | 19 | 0.00045684 | 0.000519326 | 0.000357833 | 0.006422289 | 0.007271694 | 0.007020225 |
| rbpA | A0A0K2G8D2 | Putative RNA-binding protein RbpA | 10.476 | 100 | -5.59 | -48.17 | 3.82E-08 | 5.36E-06 | yes | yes | 2 | 0 | 0 | 1 | 14 | 13 | 12 | 0.000454774 | 0.00034318 | 0.000226445 | 0.082841788 | 0.085544131 | 0.080816563 |
| NITMOv2_0791 | A0A0K2G8C5 | GCN5-related N-acetyltransferase | 20.562 | 183 | -0.198 | -1.15 | 7.46E-01 | 7.76E-01 | no | yes | 4 | 0 | 1 | 0 | 2 | 1 | 1 | 0.000449589 | 0.000375229 | 0.000299875 | 0.001227656 | 0.001347119 | 0.001490553 |
| cobI | A0A0K2GAM2 | Precorrin-2 C(20)-methyltransferase | 28.446 | 260 | -4.62 | -24.59 | 2.49E-07 | 1.33E-05 | yes | no | 0 | 2 | 1 | 2 | 23 | 26 | 21 | 0.000449146 | 0.000507045 | 0.000660986 | 0.029090726 | 0.02949819 | 0.026964628 |
| NITMOv2_3484 | A0A0K2GG05 | Uncharacterized protein | 46.572 | 425 | -1.95 | -3.86 | 2.97E-06 | 5.41E-05 | yes | no | 0 | 0 | 0 | 0 | 3 | 3 | 3 | 0.000443434 | 0.00055071 | 0.000513071 | 0.002336913 | 0.002325239 | 0.002193708 |
| NITMOv2_2265 | A0A0K2GCJ3 | PDDEXK_1 domain-containing protein | 117.31 | 1059 | -1.2 | -2.30 | 2.13E-05 | 1.98E-04 | yes | no | 0 | 10 | 10 | 9 | 13 | 16 | 15 | 0.000436815 | 0.000436826 | 0.000611176 | 0.001426373 | 0.001546565 | 0.001317519 |
| NITMOv2_3989 | A0A0K2GHD5 | Sensory sigma-54 dependent transcriptional regulator | 50.875 | 465 | 0.915 | 1.89 | 2.20E-02 | 3.46E-02 | no | no | 0 | 3 | 0 | 4 | 1 | 0 | 1 | 0.000435783 | 0.000284712 | 0.000523099 | 0.000170922 | 0.000185525 | 0.00022232 |
| cobT.1 | A0A0K2GCC8 | Nicotinate-nucleotide--dimethylbenzimidazole phosphoribosyltransferase | 36.798 | 352 | -3.74 | -13.36 | 6.07E-05 | 4.02E-04 | yes | no | 0 | 0 | 0 | 0 | 7 | 5 | 5 | 0.000434729 | 0.000121034 | 0.000524575 | 0.003853936 | 0.003279044 | 0.002678302 |
| NITMOv2_1960 | A0A0K2GCQ6 | Uncharacterized protein | 28.097 | 249 | -0.949 | -1.93 | 1.68E-04 | 7.81E-04 | no | no | 0 | 1 | 1 | 1 | 2 | 2 | 3 | 0.000427456 | 0.000542146 | 0.000334577 | 0.001019005 | 0.001096107 | 0.00100916 |
| NITMOv2_2335 | A0A0K2GCR6 | B12-binding domain-containing protein | 80.488 | 710 | -0.473 | -1.39 | 5.97E-03 | 1.15E-02 | no | no | 0 | 0 | 0 | 1 | 1 | 1 | 2 | 0.000427309 | 0.000427107 | 0.000448269 | 0.000850955 | 0.000753706 | 0.000807087 |
| miaB | A0A0K2GCN9 | tRNA-2-methylthio-N(6)-dimethylallyladenosine synthase | 49.509 | 443 | -1.81 | -3.51 | 2.26E-05 | 2.06E-04 | yes | yes | 3 | 0 | 0 | 0 | 2 | 2 | 2 | 0.000426529 | 0.000411151 | 0.000420776 | 0.001575661 | 0.001267773 | 0.001460866 |
| NITMOv2_3382 | A0A0K2GFZ2 | Acetyltransf_6 domain-containing protein | 46.068 | 408 | -0.946 | -1.93 | 5.60E-04 | 1.85E-03 | no | no | 0 | 0 | 0 | 0 | 3 | 3 | 3 | 0.000424674 | 0.000174083 | 0.000348055 | 0.000821769 | 0.000839469 | 0.000787487 |
| uvrA.1 | A0A0K2GG96 | UvrABC system protein A | 105.8 | 963 | -0.0675 | -1.05 | 5.60E-01 | 6.01E-01 | no | no | 0 | 4 | 5 | 4 | 4 | 6 | 6 | 0.000418751 | 0.000485262 | 0.000495931 | 0.001395247 | 0.00059666 | 0.000622096 |
| fadD | A0A0K2GFN8 | Long-chain fatty acid-CoA ligase | 57.482 | 525 | -1.44 | -2.71 | 7.22E-04 | 2.23E-03 | yes | no | 0 | 2 | 2 | 2 | 7 | 7 | 3 | 0.000415315 | 0.000423118 | 0.000342188 | 0.001982126 | 0.001581803 | 0.001452145 |
| NITMOv2_1603 | A0A0K2GBQ9 | Cob(I)yrinic acid a,c-diamide adenosyltransferase (Modular protein) | 48.337 | 433 | -2.17 | -4.50 | 2.99E-03 | 6.61E-03 | yes | no | 0 | 0 | 0 | 0 | 9 | 10 | 10 | 0.000414662 | 0.0003414 | 0.000339159 | 0.004793469 | 0.004866595 | 0.006692608 |
| NITMOv2_2688 | A0A0K2GDR7 | Uncharacterized protein | 105.05 | 939 | 0.18 | 1.13 | 1.69E-01 | 2.06E-01 | no | no | 0 | 2 | 3 | 1 | 9 | 13 | 11 | 0.000407685 | 0.00158197 | 0.001396419 | 0.002166331 | 0.002143769 | 0.001659846 |
| NITMOv2_3399 | A0A0K2GFR0 | UDP-N-acetylglucosamine 2-epimerase-like protein | 40.803 | 372 | -2.5 | -5.66 | 1.05E-06 | 3.10E-05 | yes | no | 0 | 1 | 1 | 1 | 16 | 13 | 12 | 0.000404986 | 0.000841445 | 0.000671876 | 0.007084479 | 0.006787101 | 0.006243437 |
| NITMOv2_3343 | A0A0K2GFK9 | Putative Helicase | 140.01 | 1244 | -3.28 | -9.71 | 2.07E-06 | 4.31E-05 | yes | no | 0 | 0 | 0 | 0 | 18 | 14 | 20 | 0.000403406 | 0.000394881 | 0.000383198 | 0.002970672 | 0.002884564 | 0.003521621 |
| NITMOv2_3390 | A0A0K2GGN6 | Asparagine synthase (glutamine-hydrolyzing) | 70.766 | 628 | -1.4 | -2.64 | 3.94E-05 | 2.98E-04 | yes | no | 0 | 1 | 0 | 1 | 3 | 9 | 8 | 0.000402373 | 0.00038223 | 0.000496046 | 0.002566872 | 0.002117512 | 0.002033675 |
| NITMOv2_3389 | A0A0K2GFQ0 | NodB homology domain-containing protein | 40.94 | 363 | -0.229 | -1.17 | 4.97E-01 | 5.39E-01 | no | yes | 3 | 0 | 1 | 0 | 3 | 3 | 2 | 0.000397714 | 0.000380744 | 0.000395487 | 0.000649855 | 0.000664587 | 0.000616234 |
| ispB | A0A0K2GGN4 | Octaprenyl-diphosphate synthase | 37.172 | 334 | -1.5 | -2.83 | 1.79E-05 | 1.73E-04 | yes | no | 0 | 0 | 0 | 0 | 3 | 4 | 3 | 0.000395923 | 0.000702003 | 0.000703606 | 0.002970375 | 0.002209335 | 0.002503165 |
| NITMOv2_3526 | A0A0K2GG45 | Putative Lipopolysaccharide heptosyltransferase II | 38.24 | 348 | -0.947 | -1.93 | 8.93E-03 | 1.62E-02 | no | yes | 1 | 1 | 0 | 0 | 3 | 2 | 3 | 0.000384624 | 0.000385476 | 0.000231966 | 0.001470899 | 0.00125048 | 0.001256401 |
| phoP | A0A0K2GJF5 | Alkaline phosphatase synthesis transcriptional regulatory protein PhoP | 27.113 | 240 | -1.69 | -3.23 | 1.58E-04 | 7.58E-04 | yes | no | 0 | 0 | 0 | 1 | 2 | 3 | 2 | 0.000383844 | 0.000389387 | 0.000390464 | 0.000891276 | 0.001102011 | 0.000801764 |
| NITMOv2_2920 | A0A0K2GEE3 | Putative Secretion protein HlyD | 48.641 | 435 | -0.0417 | -1.03 | 9.21E-01 | 9.34E-01 | no | yes | 3 | 3 | 0 | 0 | 0 | 0 | 0 | 0.000381989 | 0.000290461 | 0.000293452 | 0.000565289 | 0.000118041 | 0.000130139 |
| NITMOv2_4036 | A0A0K2GHJ7 | Histidine kinase | 76.059 | 686 | 0.0214 | 1.01 | 9.27E-01 | 9.40E-01 | no | no | 0 | 1 | 0 | 2 | 1 | 2 | 0 | 0.000374717 | 0.000286765 | 0.000527048 | 0.000485122 | 0.00057654 | 0.000548139 |
| NITMOv2_3234 | A0A0K2GFK1 | Histidine kinase | 71.375 | 643 | -1.21 | -2.31 | 1.79E-04 | 8.10E-04 | yes | no | 0 | 2 | 3 | 3 | 9 | 10 | 6 | 0.000372736 | 0.000502626 | 0.000400606 | 0.001764456 | 0.001346871 | 0.001220727 |
| NITMOv2_3346 | A0A0K2GFK3 | OmpA/MotB domain protein | 27.014 | 240 | -1.5 | -2.83 | 1.07E-04 | 5.80E-04 | yes | no | 0 | 1 | 2 | 1 | 3 | 3 | 3 | 0.000367024 | 0.000502802 | 0.000432183 | 0.001330924 | 0.001489436 | 0.001473237 |
| aroF.1 | A0A0K2GFW6 | Phospho-2-dehydro-3-deoxyheptonate aldolase | 38.286 | 352 | -3.43 | -10.78 | 8.73E-06 | 1.08E-04 | yes | yes | 3 | 0 | 0 | 0 | 4 | 5 | 5 | 0.000367002 | 8.192312E-05 | 0.000129484 | 0.004113955 | 0.005974526 | 0.003927816 |
| NITMOv2_3514 | A0A0K2GG46 | Putative Glycosyl transferase group 1 | 42.298 | 385 | -2.11 | -4.32 | 6.44E-03 | 1.23E-02 | yes | no | 0 | 3 | 3 | 4 | 7 | 6 | 7 | 0.000355536 | 0.000536064 | 0.00065533 | 0.001449813 | 0.000829262 | 0.001132061 |
| yuiH | A0A0K2G938 | Putative oxidoreductase YuiH | 24.959 | 214 | -2.36 | -5.13 | 9.42E-04 | 2.70E-03 | yes | no | 0 | 0 | 0 | 2 | 2 | 3 | 2 | 0.000348622 | 0.000301666 | 0.000497427 | 0.001123975 | 0.001045332 | 0.001127116 |
| NITMOv2_3905 | A0A0K2GH76 | Uncharacterized protein | 14.785 | 130 | 0.319 | 1.25 | 1.59E-01 | 1.95E-01 | no | yes | 2 | 1 | 2 | 3 | 1 | 0 | 0 | 0.000343036 | 0.000284223 | 0.000691317 | 0.000755032 | 0.000662226 | 0.00062233 |
| NITMOv2_1460 | A0A0K2GBC3 | Type I restriction-modification system DNA specificity subunit | 46.576 | 418 | -0.626 | -1.54 | 3.86E-04 | 1.39E-03 | no | no | 0 | 2 | 3 | 2 | 4 | 4 | 4 | 0.000340359 | 0.000309077 | 0.000439009 | 0.002119539 | 0.000944918 | 0.000789106 |
| NITMOv2_4211 | A0A0K2GI19 | MULTIHEME_CYTC domain-containing protein | 45.046 | 410 | -0.355 | -1.28 | 1.28E-02 | 2.19E-02 | no | no | 0 | 2 | 4 | 2 | 2 | 2 | 1 | 0.000336776 | 0.000860491 | 0.000454232 | 0.000537644 | 0.000950791 | 0.000822748 |
| ymxG | A0A0K2GIT6 | Putative zinc protease YmxG | 46.803 | 417 | -2.7 | -6.50 | 1.03E-02 | 1.83E-02 | yes | yes | 1 | 0 | 0 | 0 | 8 | 5 | 5 | 0.000335827 | 0.000502607 | 0.000899395 | 0.007362711 | 0.005855203 | 0.007066617 |
| NITMOv2_0544 | A0A0K2G7S4 | Uncharacterized protein | 43.389 | 389 | -0.576 | -1.49 | 5.92E-02 | 8.12E-02 | no | yes | 1 | 1 | 0 | 1 | 5 | 4 | 3 | 0.000333213 | 0.000304227 | 0.000349263 | 0.00197487 | 0.001687764 | 0.001639437 |
| NITMOv2_0728 | A0A0K2G8I2 | Radical SAM protein | 38.691 | 338 | -3.44 | -10.85 | 4.15E-03 | 8.57E-03 | yes | yes | 2 | 0 | 0 | 0 | 12 | 13 | 17 | 0.000324318 | 0.000255087 | 0.000366786 | 0.013073798 | 0.010525879 | 0.010973935 |
| rlmN | A0A0K2GHY1 | Probable dual-specificity RNA methyltransferase RlmN | 39.878 | 361 | -1.12 | -2.17 | 7.50E-02 | 1.00E-01 | no | yes | 3 | 1 | 1 | 1 | 2 | 1 | 0 | 0.000320081 | 0.000757615 | 0.005446867 | 0.001425218 | 0.000977887 | 0.001393148 |
| NITMOv2_3366 | A0A0K2GFM2 | LysM domain-containing protein | 28.82 | 264 | -0.321 | -1.25 | 2.71E-01 | 3.14E-01 | no | yes | 3 | 1 | 0 | 0 | 1 | 0 | 1 | 0.000318816 | 0 | 0.000608281 | 0.002119095 | 0.001099727 | 0.001666175 |
| NITMOv2_3516 | A0A0K2GG38 | B12-binding domain-containing protein | 56.301 | 501 | -0.0757 | -1.05 | 8.76E-01 | 8.93E-01 | no | yes | 2 | 1 | 0 | 0 | 0 | 4 | 1 | 0.000313631 | 0.000274544 | 0.000163609 | 0.000831513 | 0.001411923 | 0.001042425 |
| recQ | A0A0K2GIU9 | DNA helicase | 68.889 | 619 | -2.81 | -7.01 | 1.56E-05 | 1.58E-04 | yes | yes | 3 | 0 | 0 | 0 | 2 | 5 | 4 | 0.000300668 | 0 | 0.000231123 | 0.001125766 | 0.001875449 | 0.002066761 |
| NITMOv2_3254 | A0A0K2GFM1 | UPF0753 protein NITMOv2_3254 | 122.81 | 1100 | -0.892 | -1.86 | 5.03E-02 | 7.06E-02 | no | yes | 3 | 0 | 0 | 0 | 1 | 2 | 1 | 0.000300415 | 0.000325581 | 0.000439335 | 0.000299747 | 0.000263364 | 0.000281082 |
| NITMOv2_4540 | A0A0K2GJ87 | Histidine kinase | 75.583 | 685 | -2.51 | -5.70 | 1.34E-02 | 2.27E-02 | yes | yes | 2 | 0 | 0 | 0 | 9 | 6 | 5 | 0.000293206 | 0.000452058 | 0.000407163 | 0.003257788 | 0.002810143 | 0.002203598 |
| NITMOv2_3959 | A0A0K2GHA2 | Putative Quinol-cytochrome c reductase, cytochrome c subunit | 30.231 | 278 | -0.3 | -1.23 | 5.70E-01 | 6.11E-01 | no | no | 0 | 2 | 2 | 3 | 3 | 2 | 2 | 0.00029055 | 0.000147788 | 0.000407815 | 0.000369994 | 0.000194474 | 0.000181754 |
| NITMOv2_3383 | A0A0K2GFN5 | Glyco_trans_4-like_N domain-containing protein | 47.449 | 424 | -1.09 | -2.13 | 7.00E-05 | 4.42E-04 | yes | no | 0 | 1 | 2 | 1 | 4 | 5 | 4 | 0.000290339 | 0.000494198 | 0.000317207 | 0.001920527 | 0.001633385 | 0.001409979 |
| NITMOv2_2232 | A0A0K2GCH3 | Histidine kinase | 56.349 | 509 | -2.11 | -4.32 | 7.43E-05 | 4.59E-04 | yes | yes | 1 | 1 | 0 | 1 | 7 | 6 | 5 | 0.00028939 | 0.000102569 | 0.000325738 | 0.001742097 | 0.001550745 | 0.001371068 |
| NITMOv2_0079 | A0A0K2G6C7 | ABC_trans_aux domain-containing protein | 21.824 | 203 | -2.37 | -5.17 | 4.42E-06 | 7.00E-05 | yes | no | 0 | 1 | 5 | 3 | 11 | 12 | 15 | 0.000283594 | 0.000351353 | 0.000561616 | 0.003562822 | 0.003356573 | 0.003358171 |
| ubiE | A0A0K2G7J7 | Demethylmenaquinone methyltransferase | 28.289 | 258 | 0.0633 | 1.04 | 8.04E-01 | 8.28E-01 | no | yes | 3 | 0 | 0 | 0 | 2 | 1 | 1 | 0.000283088 | 7.8857E-05 | 0.000315347 | 0.000900545 | 0.000910395 | 0.000719141 |
| NITMOv2_1155 | A0A0K2G9D8 | Uncharacterized protein | 363.78 | 3578 | -0.877 | -1.84 | 1.06E-03 | 2.94E-03 | no | no | 0 | 2 | 0 | 2 | 7 | 6 | 11 | 0.000276237 | 0.000329765 | 0.000339638 | 0.001135243 | 0.000638345 | 0.000593758 |
| bioB | A0A0K2GA04 | Biotin synthase | 36.364 | 330 | -2.36 | -5.13 | 4.09E-03 | 8.47E-03 | yes | yes | 3 | 0 | 0 | 0 | 4 | 2 | 1 | 0.000276174 | 0.000219283 | 0.000267512 | 0.004463263 | 0.003766434 | 0.00301527 |
| pilT | A0A0K2GHR5 | Twitching motility protein | 38.552 | 352 | -0.611 | -1.53 | 8.03E-02 | 1.07E-01 | no | yes | 2 | 0 | 0 | 1 | 3 | 3 | 3 | 0.000265614 | 0.000299652 | 0.000317801 | 0.000284007 | 0.00063182 | 0.000549722 |
| NITMOv2_3494 | A0A0K2GG15 | Glyco_trans_2-like domain-containing protein | 35.885 | 310 | -0.65 | -1.57 | 8.15E-03 | 1.50E-02 | no | yes | 3 | 0 | 0 | 1 | 1 | 1 | 1 | 0.000260702 | 0 | 5.479652E-05 | 0.000806251 | 0.00137984 | 0.000774721 |
| NITMOv2_0099 | A0A0K2GJN2 | Uncharacterized protein | 48.746 | 439 | -0.593 | -1.51 | 1.36E-01 | 1.69E-01 | no | yes | 2 | 1 | 1 | 0 | 1 | 1 | 1 | 0.000250606 | 0.000504621 | 0.000248205 | 0.000770846 | 0.000651894 | 0.000835497 |
| desC | A0A0K2G9C0 | Fatty acid desaturase, type 1 | 38.591 | 339 | -1.75 | -3.36 | 2.75E-05 | 2.33E-04 | yes | yes | 3 | 0 | 0 | 0 | 5 | 5 | 7 | 0.000245315 | 0.000247383 | 0.000244908 | 0.001696786 | 0.001758456 | 0.00215397 |
| NITMOv2_4035 | A0A0K2GHV0 | Sigma54 dependent transcriptional regulator | 54.183 | 488 | -0.0272 | -1.02 | 9.43E-01 | 9.50E-01 | no | yes | 1 | 2 | 0 | 1 | 1 | 0 | 1 | 0.00024464 | 0.000319245 | 0.000420872 | 0.000372852 | 0.000311358 | 0.000464185 |

| | | | | | | | | | | | | | | | | | | | | | |
|---|---|---|---|---|---|---|---|---|---|---|---|---|---|---|---|---|---|---|---|---|---|
| queA | A0A0K2G8N3 | S-adenosylmethionine:tRNA ribosyltransferase-isomerase | 37.345 | 339 | -0.283 | -1.22 | 2.86E-01 | 3.29E-01 | no | yes | 4 | 2 | 2 | 2 | 1 | 2 | 2 | 0.000237895 | 6.343445E-05 | 7.002512E-05 | 0.000806384 | 0.000839066 | 0.000310625 |
| ftsA | A0A0K2G7Y0 | Cell division protein FtsA | 43.877 | 413 | -3.99 | -15.89 | 5.15E-06 | 7.60E-05 | yes | yes | 2 | 0 | 0 | 0 | 7 | 8 | 12 | 0.000236188 | 3.79805E-05 | 0.000278747 | 0.004176739 | 0.004474602 | 0.003631666 |
| NITMOv2_1448 | A0A0K2GAA4 | Uncharacterized protein | 185.11 | 1679 | -0.257 | -1.19 | 4.32E-02 | 6.18E-02 | no | no | 0 | 10 | 9 | 10 | 2 | 5 | 6 | 0.000233785 | 0.000212224 | 0.000241092 | 0.000224511 | 0.000203253 | 0.000187508 |
| ybbA | A0A0K2G6Z6 | Putative transporter subunit: ATP-binding component of ABC superfamily | 24.523 | 234 | -0.837 | -1.79 | 3.13E-02 | 4.68E-02 | no | yes | 4 | 0 | 0 | 0 | 1 | 1 | 0 | 0.000206743 | 0.000263143 | 0.000423939 | 0.001293683 | 0.000934679 | 0.000749511 |
| NITMOv2_3459 | A0A0K2GFX8 | Cobalamin synthesis protein P47K | 36.174 | 323 | -0.705 | -1.63 | 4.25E-03 | 8.73E-03 | no | no | 0 | 1 | 3 | 2 | 1 | 2 | 1 | 0.000202803 | 0.000189263 | 0.000223473 | 0.000303538 | 0.000366063 | 0.000278313 |
| ybgK | A0A0K2GCD8 | Putative hydrolase subunit | 34.98 | 329 | 0.483 | 1.40 | 1.09E-01 | 1.39E-01 | no | yes | 2 | 1 | 0 | 1 | 2 | 1 | 2 | 0.000199321 | 8.415233E-05 | 0.00035281 | 0.000219639 | 0.000144214 | 0.000154351 |
| cusA | A0A0K2GEE0 | Copper/silver efflux system, membrane component | 115.43 | 1043 | -0.249 | -1.19 | 2.18E-01 | 2.59E-01 | no | no | 0 | 2 | 3 | 4 | 2 | 2 | 1 | 0.000198252 | 0.00026856 | 0.000359673 | 0.000531736 | 0.000273805 | 0.000189972 |
| yiaD | A0A0K2GAD1 | Inner membrane lipoprotein YiaD | 24.093 | 230 | -3.01 | -8.06 | 8.49E-08 | 7.62E-06 | yes | yes | 3 | 0 | 0 | 0 | 3 | 2 | 2 | 0.000197373 | 0.000249182 | 0.000363105 | 0.007692325 | 0.008436489 | 0.006762734 |
| NITMOv2_0191 | A0A0K2G6N1 | PilZ domain-containing protein | 18.585 | 172 | -1.59 | -3.01 | 2.17E-03 | 5.13E-03 | yes | no | 0 | 2 | 2 | 3 | 4 | 4 | 3 | 0.000184962 | 0.000135936 | 0.000425607 | 0.000942139 | 0.000963981 | 0.000686703 |
| NITMOv2_1687 | A0A0K2GBX0 | Elp3 domain-containing protein | 38.394 | 338 | 0.192 | 1.14 | 6.77E-01 | 7.14E-01 | no | yes | 2 | 0 | 0 | 0 | 1 | 1 | 2 | 0.000180919 | 1.920984E-05 | 3.040736E-05 | 0.000464154 | 0.000382408 | 0.000404811 |
| NITMOv2_1809 | A0A0K2GBI0 | Uncharacterized protein | 23.956 | 213 | -0.837 | -1.79 | 1.04E-01 | 1.34E-01 | no | yes | 1 | 1 | 1 | 1 | 5 | 4 | 2 | 0.00017817 | 0.000108521 | 0.000243163 | 0.000861246 | 0.000863878 | 0.000638908 |
| dxs | A0A0K2G973 | 1-deoxy-D-xylulose-5-phosphate synthase | 70.402 | 648 | -7.59 | -192.67 | 1.65E-04 | 7.80E-04 | yes | yes | 1 | 0 | 0 | 1 | 60 | 56 | 50 | 0.000174498 | 5.703821E-05 | 5.430379E-05 | 0.030808391 | 0.027424026 | 0.027327848 |
| NITMOv2_2681 | A0A0K2GDR8 | MetOD1 domain-containing protein | 19.952 | 180 | -2.39 | -5.24 | 1.22E-03 | 3.25E-03 | yes | yes | 3 | 0 | 0 | 0 | 3 | 2 | 4 | 0.000168044 | 0 | 0.000162271 | 0.006559258 | 0.005191936 | 0.004769517 |
| NITMOv2_4417 | A0A0K2GJI4 | Uncharacterized protein | 29.096 | 260 | -0.992 | -1.99 | 4.68E-02 | 6.64E-02 | no | yes | 4 | 0 | 0 | 0 | 1 | 2 | 1 | 0.00016503 | 0.000236471 | 0.000161897 | 0.001810508 | 0.001627481 | 0.001348249 |
| pilM | A0A0K2G9Q3 | Type IV pilus biogenesis protein PilM | 39.84 | 368 | -2.35 | -5.10 | 5.51E-06 | 7.92E-05 | yes | no | 0 | 0 | 2 | 2 | 5 | 5 | 8 | 0.000165021 | 0.000651767 | 0.000726977 | 0.002403102 | 0.002771767 | 0.002934894 |
| NITMOv2_2442 | A0A0K2GD22 | Uncharacterized protein | 66.744 | 595 | -0.394 | -1.31 | 7.49E-02 | 1.00E-01 | no | no | 0 | 2 | 1 | 2 | 2 | 0 | 1 | 0.000162895 | 0.000222529 | 0.000242377 | 0.000282734 | 0.000329443 | 0.00026373 |
| rmlC | A0A0K2GFQ8 | dTDP-4-dehydrorhamnose 3,5-epimerase | 21.239 | 184 | -1.66 | -3.16 | 1.18E-03 | 3.17E-03 | yes | yes | 2 | 0 | 0 | 0 | 3 | 2 | 2 | 0.000162113 | 0.000275326 | 0 | 0.002797572 | 0.002423587 | 0.001926507 |
| rhlE | A0A0K2GAC2 | ATP-dependent RNA helicase RhlE | 46.369 | 430 | -7.65 | -200.85 | 5.49E-05 | 3.74E-04 | yes | yes | 1 | 0 | 0 | 0 | 33 | 35 | 35 | 0.000155184 | 6.466442E-05 | 0.000106721 | 0.019876047 | 0.019324686 | 0.020029274 |
| mutT | A0A0K2GCY3 | Mutator MutT protein | 15.92 | 138 | -0.608 | -1.52 | 4.26E-01 | 4.69E-01 | no | yes | 4 | 0 | 0 | 0 | 0 | 1 | 1 | 0.000144562 | 0.000138748 | 0 | 0.001407774 | 0.001596097 | 0.001690252 |
| pal | A0A0K2GBE8 | Peptidoglycan-associated protein | 21.898 | 199 | -2.27 | -4.82 | 1.17E-02 | 2.04E-02 | yes | yes | 4 | 0 | 0 | 0 | 2 | 2 | 1 | 0.000140408 | 0 | 0 | 0.003294955 | 0.004282101 | 0.003112188 |
| NITMOv2_2921 | A0A0K2GFE8 | Putative Cation efflux system protein CzcC | 46.141 | 419 | -0.483 | -1.40 | 1.58E-01 | 1.93E-01 | no | no | 0 | 1 | 2 | 5 | 2 | 0 | 1 | 0.000140161 | 0.000167274 | 0.000232752 | 0.000183213 | 0.000254322 | 0.000334361 |
| NITMOv2_0278 | A0A0K2G7X8 | YkuD domain-containing protein | 41.952 | 378 | -0.0167 | -1.01 | 9.81E-01 | 9.83E-01 | no | yes | 4 | 0 | 1 | 0 | 1 | 1 | 1 | 0.00013491 | 0.00010409 | 0 | 0.001313095 | 0.001170077 | 0.000316487 |
| nemA | A0A0K2G676 | N-ethylmaleimide reductase | 39.478 | 363 | -5.7 | -51.98 | 1.36E-04 | 6.84E-04 | yes | yes | 2 | 0 | 0 | 0 | 21 | 25 | 27 | 0.000129967 | 9.460415E-05 | 0.000183805 | 0.018424916 | 0.01990887 | 0.018815542 |
| trmE | A0A0K2G715 | tRNA modification GTPase MnmE | 50.315 | 475 | -3.2 | -9.19 | 2.17E-04 | 9.36E-04 | yes | no | 0 | 0 | 0 | 0 | 14 | 9 | 11 | 0.000127057 | 0.000123193 | 0.000221556 | 0.02075561 | 0.001226103 | 0.000947574 |
| NITMOv2_2463 | A0A0K2GD42 | Putative Multi-domain non-ribosomal peptide synthetase | 339.61 | 3095 | 0.0981 | 1.07 | 5.07E-01 | 5.49E-01 | no | no | 0 | 0 | 0 | 2 | 1 | 3 | 4 | 0.000119464 | 9.716186E-05 | 8.806823E-05 | 0.000190927 | 0.000145844 | 0.000139185 |
| NITMOv2_3395 | A0A0K2GGP1 | Uncharacterized protein | 47.132 | 426 | -0.824 | -1.77 | 1.33E-03 | 3.52E-03 | no | no | 0 | 2 | 2 | 1 | 4 | 6 | 2 | 0.000112892 | 7.716359E-05 | 0.000137176 | 0.001110337 | 0.001071558 | 0.000694093 |
| NITMOv2_3004 | A0A0K2GEZ8 | UPF0753 protein NITMOv2_3004 | 124.49 | 1097 | -0.905 | -1.87 | 1.89E-03 | 4.60E-03 | no | no | 0 | 0 | 1 | 1 | 1 | 1 | 2 | 0.000106987 | 0.000139106 | 9.462709E-05 | 0.00032566 | 0.000347962 | 0.000305357 |
| NITMOv2_1573 | A0A0K2GAM6 | PINc domain-containing protein | 15.132 | 135 | -0.35 | -1.27 | 7.60E-02 | 1.01E-01 | no | yes | 4 | 3 | 5 | 6 | 6 | 6 | 7 | 0.000100152 | 0.000121626 | 0.000193085 | 0.000902692 | 0.000746295 | 0.0001931 |
| NITMOv2_2993 | A0A0K2GEL9 | Putative Mannose-1-phosphate guanylyltransferase (GDP) | 37.21 | 332 | -3.15 | -8.88 | 2.97E-03 | 6.56E-03 | yes | yes | 3 | 3 | 0 | 1 | 0 | 0 | 0 | 9.801839E-05 | 6.025295E-05 | 7.460348E-05 | 3.185232E-05 | 1.632142E-05 | 2.319936E-05 |
| NITMOv2_4138 | A0A0K2GHU2 | Uncharacterized protein | 23.362 | 212 | -2.33 | -5.03 | 3.23E-03 | 7.03E-03 | yes | no | 0 | 0 | 1 | 2 | 3 | 3 | 3 | 9.685484E-05 | 0.000107162 | 0.000158022 | 0.000472743 | 0.000512047 | 0.01371517 |
| smf | A0A0K2G8W8 | DprA_WH domain-containing protein | 39.076 | 372 | -2.15 | -4.44 | 1.10E-05 | 1.28E-04 | yes | no | 0 | 2 | 2 | 2 | 5 | 7 | 4 | 9.526761E-05 | 0.000230703 | 0.000277731 | 0.000972628 | 0.000991917 | 0.000978538 |
| pmtA | A0A0K2G7A6 | Phosphatidylethanolamine N-methyltransferase | 26.576 | 242 | -4.56 | -23.59 | 4.40E-04 | 1.54E-03 | yes | yes | 3 | 0 | 0 | 0 | 14 | 15 | 15 | 9.328832E-05 | 0 | 0 | 0.016793135 | 0.016377353 | 0.011699117 |
| prmA | A0A0K2G996 | Ribosomal protein L11 methyltransferase | 31.295 | 289 | -1.05 | -2.07 | 2.88E-01 | 3.31E-01 | no | no | 0 | 1 | 1 | 2 | 2 | 2 | 1 | 9.25084E-05 | 0.000885814 | 0.000102039 | 0.000721464 | 0.001402384 | 0.000407113 |
| asnB | A0A0K2GGP6 | Asparagine synthetase | 71.221 | 631 | -1.44 | -2.71 | 8.48E-05 | 5.06E-04 | yes | yes | 3 | 1 | 0 | 0 | 2 | 2 | 2 | 8.389772E-05 | 0 | 5.778357E-05 | 0.000434169 | 0.000421281 | 0.000429985 |
| NITMOv2_3405 | A0A0K2GGQ2 | Putative Polysaccharide pyruvyl transferase | 50.502 | 449 | -0.0623 | -1.04 | 8.04E-01 | 8.28E-01 | no | yes | 3 | 0 | 0 | 0 | 3 | 1 | 3 | 8.365532E-05 | 9.58439E-05 | 5.179029E-05 | 0.000465917 | 0.000484873 | 0.000496084 |
| NITMOv2_4856 | A0A0K2GK39 | UPF0753 protein NITMOv2_4856 | 121.81 | 1079 | 0.367 | 1.29 | 5.19E-01 | 5.61E-01 | no | yes | 4 | 1 | 1 | 1 | 1 | 2 | 1 | 8.097831E-05 | 0.00012771 | 7.246001E-05 | 0.000335359 | 0.000500922 | 0.000348296 |
| NITMOv2_0175 | A0A0K2G6Q8 | Uncharacterized protein | 145.33 | 1296 | -3.04 | -8.22 | 6.14E-03 | 1.18E-02 | yes | yes | 2 | 0 | 0 | 0 | 5 | 3 | 2 | 7.553788E-05 | 0 | 0.000245981 | 0.000675783 | 0.000816832 | 0.000638459 |
| NITMOv2_0260 | A0A0K2G752 | AAA domain-containing protein | 34.573 | 313 | -2.63 | -6.19 | 2.37E-05 | 2.13E-04 | yes | yes | 3 | 0 | 0 | 0 | 3 | 4 | 3 | 5.890041E-05 | 0.000101732 | 0.00013829 | 0.002540515 | 0.002459943 | 0.001961031 |
| rimO | A0A0K2G721 | Ribosomal protein S12 methylthiotransferase RimO | 54.811 | 495 | -2.86 | -7.26 | 8.55E-07 | 2.82E-05 | yes | yes | 3 | 0 | 0 | 0 | 5 | 5 | 2 | 5.483431E-05 | 0.0002826 | 0 | 0.001233609 | 0.001824643 | 0.001580261 |
| pykA | A0A0K2GA77 | Pyruvate kinase | 51.321 | 482 | -1.42 | -2.68 | 4.52E-05 | 3.25E-04 | yes | no | 0 | 2 | 2 | 3 | 6 | 3 | 2 | 5.162191E-05 | 0.000127835 | 0.000131325 | 0.000448784 | 0.000253297 | 0.000450034 |
| gspE.1 | A0A0K2GFM8 | General secretion pathway protein E | 63.584 | 574 | -2.11 | -4.32 | 1.28E-03 | 3.39E-03 | yes | yes | 1 | 0 | 0 | 0 | 2 | 3 | 2 | 5.146382E-05 | 0.000190554 | 0.000120945 | 0.000842663 | 0.000676022 | 0.000729282 |
| NITMOv2_4162 | A0A0K2GIV0 | Peptidase_M23 domain-containing protein | 36.217 | 333 | -2.63 | -6.19 | 8.57E-04 | 2.53E-03 | yes | yes | 3 | 0 | 0 | 0 | 0 | 3 | 1 | 5.02539E-05 | 0 | 0 | 0.000801261 | 0.009248132 | 0.000442859 |
| kamA | A0A0K2GHN2 | L-lysine 2,3-aminomutase | 43.085 | 389 | -1.85 | -3.61 | 1.20E-04 | 6.32E-04 | yes | yes | 3 | 0 | 0 | 0 | 3 | 3 | 4 | 4.261074E-05 | 4.704982E-05 | 5.992512E-05 | 0.001142943 | 0.001321204 | 0.0011677 |
| NITMOv2_3397 | A0A0K2GG08 | Asparagine synthetase | 71.031 | 631 | -2.38 | -5.21 | 1.07E-03 | 2.96E-03 | yes | yes | 1 | 1 | 1 | 1 | 5 | 7 | 7 | 3.271425E-05 | 0.000169337 | 0.000156109 | 0.001030377 | 0.001304269 | 0.000967191 |
| NITMOv2_0022 | A0A0K2G680 | Multidrug efflux system protein | 111.48 | 1031 | -6.69 | -103.25 | 1.16E-06 | 3.12E-05 | yes | yes | 3 | 0 | 0 | 0 | 45 | 49 | 44 | 3.10954E-05 | 2.809575E-05 | 0.00010481 | 0.015296101 | 0.016643032 | 0.016151445 |
| NITMOv2_2696 | A0A0K2GER4 | Putative Cyanobacterial phytochrome B | 86.394 | 768 | 0.134 | 1.10 | 8.18E-01 | 8.39E-01 | no | yes | 3 | 0 | 0 | 0 | 1 | 2 | 1 | 2.830668E-05 | 0 | 0 | 0.000129306 | 0.000218945 | 0.000187113 |
| waaA | A0A0K2GG61 | 3-deoxy-D-manno-octulosonic acid transferase | 51.217 | 467 | -0.396 | -1.32 | 1.83E-01 | 2.21E-01 | no | yes | 3 | 1 | 0 | 0 | 2 | 1 | 3 | 2.154461E-05 | 0.00014832 | 4.384527E-05 | 0.000539065 | 0.000621395 | 0.000381525 |
| cobL | A0A0K2GAP8 | Precorrin-6Y C(5,15)-methyltransferase (Decarboxylating) | 45.379 | 418 | -3.74 | -13.36 | 1.53E-06 | 3.61E-05 | yes | yes | 3 | 0 | 0 | 0 | 17 | 17 | 14 | 2.067849E-05 | 0 | 2.83099E-05 | 0.008529983 | 0.007363983 | 0.006832501 |
| NITMOv2_3384 | A0A0K2GFP5 | Acetyltransf_6 domain-containing protein | 43.623 | 379 | -2.09 | -4.26 | 4.38E-04 | 1.54E-03 | yes | yes | 3 | 0 | 0 | 0 | 3 | 2 | 1 | 1.634279E-05 | 0 | 0 | 0.002266577 | 0.002243826 | 0.001414762 |
| NITMOv2_2680 | A0A0K2GEP9 | Putative sensor signal transduction histidine kinase | 71.672 | 650 | -2.26 | -4.79 | 6.43E-03 | 1.23E-02 | yes | yes | 3 | 0 | 1 | 0 | 7 | 6 | 2 | 1.294532E-05 | 0.001031005 | 0 | 0.00113046 | 0.000743654 | 0.000543824 |
| cobQ.1 | A0A0K2GCD3 | Cobyric acid synthase | 55.452 | 505 | -0.346 | -1.27 | 5.10E-01 | 5.52E-01 | no | yes | 3 | 0 | 0 | 0 | 2 | 1 | 2 | 1.258276E-05 | 5.533893E-05 | 0 | 0.00039182 | 0.000390502 | 0.000403355 |
| NITMOv2_1887 | A0A0K2GCI1 | Methylase_S domain-containing protein | 50.151 | 450 | -2.02 | -4.06 | 5.41E-03 | 1.06E-02 | yes | yes | 2 | 0 | 0 | 0 | 5 | 2 | 4 | 1.203029E-05 | 1.456939E-05 | 6.07016E-05 | 0.002040763 | 0.00161008 | 0.001513137 |
| gidA | A0A0K2G6X5 | tRNA uridine 5-carboxymethylaminomethyl modification enzyme MnmG | 69.019 | 624 | -2.69 | -6.45 | 1.48E-04 | 7.30E-04 | yes | yes | 1 | 0 | 0 | 0 | 7 | 6 | 6 | 8.821466E-06 | 4.764036E-06 | 6.406444E-05 | 0.001216817 | 0.000828252 | 0.000768733 |
| NITMOv2_3914 | A0A0K2GH64 | Putative Histidine kinase | 92.176 | 841 | 0.438 | 1.35 | 5.95E-01 | 6.36E-01 | no | yes | 2 | 1 | 1 | 0 | 1 | 1 | 2 | 7.347638E-06 | 2.303312E-05 | 8.172793E-06 | 7.947753E-05 | 8.870898E-05 | 7.964239E-05 |
| NITMOv2_1003 | A0A0K2G933 | Cellulose synthase (UDP-forming) | 130.51 | 1148 | 1.99 | 3.97 | 1.58E-02 | 2.61E-02 | yes | yes | 3 | 0 | 0 | 1 | 3 | 2 | 2 | 6.802963E-06 | 7.275798E-06 | 4.000888E-06 | 2.653496E-05 | 6.198877E-05 | 4.23062E-05 |
| NITMOv2_2523 | A0A0K2GDB4 | Nitrogen regulation protein B | 174.97 | 1590 | 0.519 | 1.43 | 4.34E-01 | 4.77E-01 | no | yes | 4 | 0 | 0 | 0 | 1 | 2 | 1 | 4.748204E-06 | 0 | 0 | 9.084522E-05 | 5.536698E-05 | 4.610203E-05 |
| NITMOv2_1715 | A0A0K2GB12 | Methyltransferase, ubiE family | 30.229 | 273 | 1.5 | 2.83 | 1.08E-01 | 1.38E-01 | no | yes | 4 | 0 | 0 | 0 | 0 | 2 | 1 | 3.826008E-06 | 8.352072E-06 | 4.121865E-06 | 1.649847E-05 | 7.309759E-05 | 2.137427E-05 |
| NITMOv2_1866 | A0A0K2GBE0 | Putative FAD/FMN-containing dehydrogenase, glycolate oxidase, fused subunits GlcD and Gl | 105.56 | 965 | -1.62 | -3.07 | 2.83E-02 | 4.30E-02 | yes | yes | 3 | 0 | 0 | 0 | 7 | 3 | 4 | 2.451883E-06 | 0 | 6.994268E-06 | 0.001362153 | 0.000349221 | 0.000435253 |
| NITMOv2_3973 | A0A0K2GIB5 | Response regulatory domain-containing protein | 17.669 | 162 | -4.97 | -31.34 | 4.20E-07 | 1.68E-05 | yes | yes | 3 | 0 | 0 | 0 | 15 | 13 | 12 | 0 | 0 | 0 | 0.023022631 | 0.023524286 | 0.018738222 |
| NITMOv2_3975 | A0A0K2GHL5 | RRM domain-containing protein | 11.163 | 106 | -2.98 | -7.89 | 1.76E-03 | 4.36E-03 | yes | yes | 3 | 0 | 0 | 0 | 17 | 18 | 16 | 0 | 2.979503E-05 | 0 | 0.018201324 | 0.014061436 | 0.012427716 |
| NITMOv2_0021 | A0A0K2G6B4 | Putative RND efflux system, membrane fusion protein | 39.671 | 375 | -4.74 | -26.72 | 3.98E-09 | 1.79E-06 | yes | yes | 3 | 0 | 0 | 0 | 14 | 14 | 14 | 0 | 0 | 0 | 0.017289184 | 0.020291076 | 0.019196743 |
| NITMOv2_0590 | A0A0K2G7S7 | Uncharacterized protein | 15.796 | 146 | -2.08 | -4.23 | 1.47E-02 | 2.47E-02 | yes | yes | 3 | 0 | 0 | 0 | 5 | 3 | 4 | 0 | 0 | 0.000224988 | 0.008335561 | 0.00738387 | 0.006684876 |
| NITMOv2_1497 | A0A0K2GAF8 | Uncharacterized protein | 31.92 | 295 | -2.61 | -6.11 | 7.14E-05 | 4.48E-04 | yes | yes | 3 | 0 | 0 | 0 | 7 | 5 | 5 | 0 | 0 | 0 | 0.006853779 | 0.005857067 | 0.003626991 |
| cobQ | A0A0K2GAR5 | Cobyric acid synthase | 55.704 | 512 | -4.23 | -18.77 | 9.05E-07 | 2.93E-05 | yes | yes | 3 | 0 | 0 | 0 | 18 | 9 | 9 | 0 | 0 | 9.601901E-05 | 0.006551855 | 0.005715216 | 0.005013342 |
| cobA | A0A0K2GAM7 | SUMT | 51.509 | 491 | -4.29 | -19.56 | 1.61E-06 | 3.75E-05 | yes | yes | 3 | 0 | 0 | 0 | 21 | 14 | 14 | 0 | 0 | 0 | 0.006342625 | 0.006263355 | 0.005628479 |
| NITMOv2_3778 | A0A0K2GH38 | Putative Lytic murein transglycosylase | 47.76 | 428 | -2.81 | -7.01 | 3.20E-06 | 5.70E-05 | yes | yes | 3 | 0 | 0 | 0 | 7 | 8 | 11 | 0 | 0 | 0 | 0.004914446 | 0.00433679 | 0.004061057 |
| NITMOv2_1592 | A0A0K2GAL9 | FAD-binding FR-type domain-containing protein | 25.657 | 241 | -1.71 | -3.27 | 3.67E-02 | 5.38E-02 | no | yes | 3 | 0 | 0 | 0 | 9 | 3 | 2 | 0 | 0 | 0 | 0.004461338 | 0.004080277 | 0.003443762 |
| cbiX | A0A0K2GAX3 | Sirohydrochlorin cobaltochelatase | 43.134 | 393 | -3.72 | -13.18 | 6.28E-04 | 2.02E-03 | yes | yes | 3 | 0 | 0 | 0 | 7 | 8 | 9 | 0 | 0 | 0 | 0.003168943 | 0.002625566 | 0.002530676 |
| NITMOv2_3512 | A0A0K2GGE6 | Putative transcriptional regulator, AsnC family | 20.958 | 186 | -1.58 | -2.99 | 1.75E-03 | 4.34E-03 | yes | yes | 3 | 0 | 0 | 0 | 2 | 2 | 2 | 0 | 7.511429E-05 | 0 | 0.002958678 | 0.003130823 | 0.002859912 |
| NITMOv2_3385 | A0A0K2GGN1 | Diaminopimelate decarboxylase | 46.076 | 427 | -1.83 | -3.56 | 4.68E-03 | 9.40E-03 | yes | yes | 3 | 0 | 0 | 0 | 3 | 2 | 3 | 0 | 0 | 0 | 0.002592637 | 0.002375112 | 0.001906908 |
| NITMOv2_4144 | A0A0K2GHW8 | Transcriptional regulator, NifA subfamily, Fis Family | 59.273 | 531 | -2.21 | -4.63 | 1.10E-03 | 3.02E-03 | yes | yes | 3 | 0 | 0 | 0 | 4 | 7 | 9 | 0 | 0 | 0 | 0.00190335 | 0.001778188 | 0.001608438 |
| NITMOv2_1610 | A0A0K2GAZ4 | Aminotransferase | 38.937 | 363 | -2.02 | -4.06 | 1.01E-04 | 5.64E-04 | yes | yes | 3 | 0 | 0 | 0 | 3 | 3 | 3 | 0 | 0 | 0 | 0.001802067 | 0.001512058 | 0.000814279 |
| NITMOv2_0497 | A0A0K2G7K8 | Putative Radical SAM superfamily protein, Coenzyme PQQ synthesis protein E | 51.013 | 460 | -2.44 | -5.43 | 6.82E-05 | 4.34E-04 | yes | yes | 3 | 0 | 0 | 0 | 4 | 2 | 2 | 0 | 5.69991E-05 | 0 | 0.001575661 | 0.001515321 | 0.001573176 |
| cobB | A0A0K2GAQ6 | Cobyrinate a,c-diamide synthase | 49.873 | 458 | -2.02 | -4.06 | 8.74E-03 | 1.59E-02 | yes | yes | 3 | 0 | 0 | 0 | 0 | 2 | 2 | 0 | 4.956452E-05 | 0 | 0.001570034 | 0.0020723 | 0.002145339 |
| NITMOv2_4158 | A0A0K2GHU4 | DUF3641 domain-containing protein | 37.605 | 344 | -1.42 | -2.68 | 6.44E-03 | 1.23E-02 | yes | yes | 3 | 0 | 0 | 0 | 2 | 1 | 0 | 0 | 0 | 0 | 0.001442409 | 0.001669586 | 0.001105935 |
| NITMOv2_3394 | A0A0K2GFQ5 | Asparagine synthetase | 71.672 | 628 | -1.44 | -2.71 | 2.54E-02 | 3.91E-02 | yes | yes | 3 | 0 | 0 | 0 | 7 | 8 | 6 | 0 | 2.589979E-06 | 8.530166E-05 | 0.001335188 | 0.001344198 | 0.000949732 |
| NITMOv2_3398 | A0A0K2GFP8 | FemAB-related protein, PEP-CTERM system-associated | 38.806 | 351 | -0.84 | -1.79 | 2.47E-01 | 2.89E-01 | no | yes | 3 | 0 | 0 | 0 | 5 | 4 | 3 | 0 | 0 | 0 | 0.001160594 | 0.000877566 | 0.000768068 |
| cbiG | A0A0K2GBQ5 | Uroporphyrin-III C-methyltransferase | 40.842 | 379 | -1.56 | -2.95 | 5.36E-06 | 7.80E-05 | yes | yes | 3 | 0 | 0 | 0 | 3 | 2 | 3 | 0 | 0 | 0 | 0.001132756 | 0.000955188 | 0.000915513 |
| cbiD | A0A0K2GAP4 | Cobalt-precorrin-5B C(1)-methyltransferase | 37.37 | 356 | -0.635 | -1.55 | 2.75E-04 | 1.09E-03 | no | yes | 3 | 0 | 0 | 0 | 1 | 2 | 2 | 0 | 0 | 0 | 0.000967208 | 0.001126528 | 0.000903502 |
| hlyB | A0A0K2GAF6 | Type I secretion system ATPase HlyB | 79.971 | 724 | -1.63 | -3.10 | 5.47E-03 | 1.07E-02 | yes | yes | 3 | 0 | 0 | 0 | 3 | 3 | 3 | 0 | 0 | 0 | 0.000859869 | 0.00069058 | 0.000734587 |

| | | | | | | | | | | | | | | | | | | | | | | | |
|---|---|---|---|---|---|---|---|---|---|---|---|---|---|---|---|---|---|---|---|---|---|---|---|
| NITMOv2_3490 | A0A0K2GGZ6 | Putative UDP-alpha-N-acetylglucosamine 3-alpha-N-acetyl-L-fucosaminyltransferase | 47.289 | 421 | -0.854 | -1.81 | 2.28E-01 | 2.70E-01 | no | yes | 4 | 0 | 0 | 0 | 3 | 2 | 0 | 0 | 0 | 0 | 0.000812189 | 0.000977685 | 0.000434516 |
| mutY | A0A0K2GCP6 | A/G-specific adenine glycosylase | 32.31 | 285 | -0.85 | -1.80 | 6.44E-02 | 8.76E-02 | no | yes | 3 | 0 | 0 | 0 | 2 | 2 | 3 | 0 | 0 | 0 | 0.000657421 | 0.000641002 | 0.001026709 |
| NITMOv2_3522 | A0A0K2GGF4 | Glyco_trans_2-like domain-containing protein | 32.352 | 281 | -1.14 | -2.20 | 1.09E-01 | 1.39E-01 | no | yes | 4 | 0 | 0 | 0 | 1 | 2 | 1 | 0 | 0 | 6.731607E-05 | 0.000505601 | 0.000901399 | 0.001543148 |
| NITMOv2_1605 | A0A0K2GAY9 | Putative Xylose isomerase domain protein TIM barrel | 29.908 | 261 | 0.512 | 1.43 | 2.45E-02 | 3.78E-02 | no | yes | 4 | 0 | 0 | 0 | 2 | 1 | 1 | 0 | 0 | 0 | 0.000499352 | 0.000418251 | 0.000320138 |
| NITMOv2_0106 | A0A0K2GKB0 | Putative Regulatory protein Crp | 25.047 | 222 | 1.13 | 2.19 | 1.31E-04 | 6.66E-04 | yes | yes | 3 | 0 | 0 | 0 | 2 | 1 | 1 | 0 | 2.475782E-05 | 7.683898E-05 | 0.000454248 | 0.000405169 | 0.000309708 |
| NITMOv2_4242 | A0A0K2GI49 | Sigma-54 factor interaction domain-containing protein | 45.167 | 405 | 0.222 | 1.17 | 4.21E-01 | 4.64E-01 | no | yes | 3 | 0 | 0 | 0 | 1 | 2 | 2 | 0 | 0 | 7.679105E-05 | 0.000408493 | 0.000296178 | 0.000206011 |
| NITMOv2_4083 | A0A0K2GHR3 | Glycosyl transferase, family 2 | 25.951 | 230 | 0.678 | 1.60 | 6.75E-02 | 9.13E-02 | no | yes | 3 | 0 | 0 | 0 | 1 | 2 | 1 | 0 | 0 | 0 | 0.000312497 | 0.000584184 | 0.000589083 |
| NITMOv2_2946 | A0A0K2GEH0 | Uncharacterized protein | 41.821 | 386 | -0.098 | -1.07 | 8.79E-01 | 8.96E-01 | no | yes | 3 | 0 | 0 | 0 | 1 | 3 | 1 | 0 | 0 | 0 | 0.000302827 | 0.000343224 | 0.000338532 |
| NITMOv2_3396 | A0A0K2GFP9 | Glycosyl transferase family 2 | 41.978 | 385 | 0.914 | 1.88 | 9.56E-02 | 1.24E-01 | no | yes | 4 | 0 | 0 | 0 | 1 | 1 | 0 | 0 | 0 | 0 | 0.000258568 | 0.000269626 | 0.000164197 |
